# ZNF423 depletion induces the integrated stress response and represents a potential vulnerability in NF1-associated MPNST

**DOI:** 10.64898/2026.03.03.709360

**Authors:** Sarah K. Morrow, Christine A. Berryhill, Emily E. White, Colin J. Beach, Shelley A. H. Dixon, Emily Y. Zhang, Christopher Davis, Erin M. Mannion, Brooke E. Hickey, Henry Mang, Marisa D. Ciesielski, Silpa Gampala, Mohammad Reza Saadatzadeh, Emmanuel A. Gyabaah, Christopher D. Collier, Aaron Cohen-Gadol, Christine A. Pratilas, Karen E. Pollok, Melissa L. Fishel, D. Wade Clapp, Dana K. Mitchell, Steven D. Rhodes, Steven P. Angus

## Abstract

Malignant peripheral nerve sheath tumors (MPNST) are aggressive sarcomas with limited systemic therapies and represent the leading cause of mortality for individuals with neurofibromatosis type 1 (NF1). Malignant progression can reactivate developmental precursor programs that are largely absent from normal nerve and benign tumors, creating tumor-selective vulnerabilities. Zinc finger protein 423 (ZNF423; also known as OAZ/ROAZ) is a developmentally regulated transcription factor that delays olfactory precursor differentiation and has been implicated in B-cell malignancy. Here, we asked whether ZNF423 is reactivated and functionally required in NF1-associated MPNST. In genetically defined models, *Nf1* loss reduced *Zfp423* in a benign tumor cell-of-origin context, whereas combined *Nf1* and *Cdkn2a* loss induced marked *Zfp423* upregulation during transformation. ZNF423 depletion impaired DNA synthesis and proliferation, induced DNA damage signaling, and activated the integrated stress response (ISR), increasing sensitivity to cytotoxic agents. In an orthotopic MPNST model, shRNA-mediated suppression of ZNF423 reduced tumor initiation in vivo; however, tumors that eventually emerged showed restoration of ZNF423 expression. ZNF423 is developmentally restricted in the peripheral nerve lineage yet elevated in MPNST, with single-cell analyses of patient nerve sheath tumors revealing localized expression restricted to malignant cells rather than SOX10-positive benign tumor cells. These data identify ZNF423 as a putative malignant biomarker, a potential dependency in NF1-MPNST, and nominate downstream stress and genome maintenance pathways as cooperative therapeutic vulnerabilities.

**STATEMENT OF SIGNIFICANCE:** ZNF423 is a developmentally restricted transcription factor selectively reactivated in NF1-associated malignant peripheral nerve sheath tumors. Targeted ablation triggers the integrated stress response, impairs DNA synthesis, sensitizes cells to chemotherapy and PARP inhibition, and restricts in vivo growth. ZNF423 represents a candidate biomarker and therapeutic vulnerability in this aggressive sarcoma.

## INTRODUCTION

Neurofibromatosis type 1 (NF1) is a common autosomal dominant cancer predisposition syndrome, affecting ∼1 in 2500 individuals worldwide^1^. NF1-related tumorigenesis arises from biallelic inactivation of the *NF1* tumor suppressor gene, leading to hyperactive Ras signaling, and aberrant cell proliferation^2^. The hallmark manifestation of NF1 is the development of plexiform neurofibromas (PNF), benign Schwann cell (SC) precursor-derived tumors that occur in ∼50% of NF1 patients^3,4^. PNFs carry an 8-13% lifetime risk of malignant transformation into malignant peripheral nerve sheath tumors (MPNST), highly aggressive sarcomas and the leading cause of NF1-associated mortality^5,6^. MPNST are frequently refractory to therapy, with surgical resection often infeasible, and consequently, 5-year survival rates are extremely poor^7^. While MEK1/2 kinase inhibitors are effective in PNF, they have shown limited efficacy in MPNST, despite extensive combination strategies^8–11^. Therefore, elucidating molecular drivers of MPNST tumorigenesis remains essential for identifying actionable therapeutic vulnerabilities in NF1-associated MPNST.

Deletion of chromosome 9p21.3, which harbors the *CDKN2A/B* (*INK4A/ARF and INK4B*) tumor suppressor loci, is a frequent event driving the progression of PNF to atypical neurofibromas (ANF) or atypical neurofibromatous neoplasms of uncertain biological potential (ANNUBP), and eventually to MPNST^12,13^. Our group and others have confirmed that combined inactivation of *Nf1* and *Cdkn2a* in genetically engineered mouse models of NF1 is sufficient to drive the development of MPNST that faithfully recapitulate human MPNST^12,14^. In addition to *CDKN2A/B* deletion, 60-90% of MPNST patients harbor somatic mutations in polycomb repressive complex 2 (PRC2) core components *EED* or *SUZ12*, resulting in the loss of H3K27me3-mediated chromatin repression^15–18^. Functional PRC2 loss promotes H3K27 acetylation and recruitment of BET bromodomain protein BRD4, driving oncogenic transcriptional programs, and creating a vulnerability that can be exploited through BRD4 inhibition^18–21^.

Although PRC2 inactivation is implicated as an early driver of MPNST, it is not universal, suggesting alternative pathways contribute to malignant progression^22^. Accumulating evidence demonstrates that MPNST evolution is accompanied by aberrant reactivation of developmental and stem-like programs, resulting in dedifferentiated transcriptional states that shape tumor heterogeneity and influence therapeutic response^23–25^. Notably, recent work by our group identified the embryonic factor delta-like non-canonical Notch ligand 1 (DLK1), as a marker of early malignant transition that defines a subset of MPNST with embryonic transcriptional signatures and worse clinical outcome^26^. These developmental programs are normally transient and largely extinguished in mature peripheral nerve and benign tumors, however their reactivation in malignancy may create tumor-specific dependencies and vulnerabilities that are not shared by normal tissue. Defining the regulators that orchestrate the acquisition of these precursor-like states is a priority for identifying new therapeutic opportunities in NF1-associated MPNST.

Another treatment-refractory pediatric solid tumor, neuroblastoma (NB), shares a SC precursor lineage origin and enrichment of dedifferentiation programs similar to those observed in MPNST^27^. In select cases, all-trans retinoic acid (ATRA) has been used to successfully drive NB to a more differentiated, less aggressive state. Seminal studies by the Bernards group used siRNA screening to identify NF1 and the zinc finger protein 423 (ZNF423, also known as OAZ/ROAZ) as critical mediators of RA-driven differentiation^28,29^. Importantly, they demonstrated that NF1 loss impairs RA responsiveness in part through repression of ZNF423-dependent programs. Given the shared neural crest-derived lineage of neuroblastoma and peripheral nerve sheath tumors, these findings suggested that NF1 status could similarly intersect with ZNF423-regulated differentiation during PNF transformation to MPNST. Recent studies have explored the use of RA in MPNST models to induce differentiation and growth arrest, which were successful in vitro but not in xenograft studies^30,31^.

In the present study, we demonstrate that ZNF423 is selectively reactivated during malignant progression in NF1-associated tumors. Loss of *Nf1* in a PNF murine tumor cell-of-origin model was associated with reduced *Zfp423* (murine ortholog of ZNF423) expression, whereas combined loss of *Nf1* and *Cdkn2a* markedly upregulated *Zfp423*. Peak expression of *Zfp423* was found to occur early in murine Schwann cell development, and ZNF423 expression was elevated in human MPNST relative to PNF and normal nerve tissue. Functionally, ZNF423 was required for MPNST cell viability, proliferation, and DNA synthesis. Accordingly, depletion of ZNF423 triggered cellular stress and impaired tumor cell survival. Suppression of ZNF423 in an NF1 MPNST xenograft model significantly delayed tumor growth in vivo, with tumors resuming growth when the cells reacquired expression of ZNF423. Moreover, ZNF423 loss enhanced DNA damage and apoptosis when combined with standard-of-care chemotherapy. Lastly, integrative single-cell RNAseq analysis of 71 patient PNF and MPNST tumors demonstrated that ZNF423 expression was absent in PNF, yet exclusively annotated malignant cell populations in MPNST. Taken together, our data suggest that ZNF423 may serve as a candidate biomarker and therapeutic vulnerability for NF1 MPNST, meriting further mechanistic evaluation and screening for direct inhibitor molecules.

## RESULTS

### ZNF423 is a developmentally restricted neuronal precursor that is aberrantly upregulated during NF1-tumorigenesis and progression to MPNST

During neurogenesis, ZNF423 controls lineage specification by delaying olfactory precursor differentiation via repression of early B-cell factor (EBF)-dependent programs, thereby prolonging a progenitor-like state^32^. In cancer, however, ZNF423 function is context dependent. In neuroblastoma, ZNF423 functions like a tumor suppressor, as it is required for retinoic acid (RA)-induced differentiation and its expression is associated with improved clinical outcome^28^. In contrast, in B-cell malignancy, aberrant ZNF423 activity interferes with B-lineage transcriptional circuitry, impedes differentiation, and is linked to worse prognosis^33–35^. To examine the developmental regulation of *Zfp423* within the murine peripheral nervous system, we analyzed transcriptomic data from the Sciatic Nerve Atlas (SNAT), which revealed that *Zfp423* expression peaks at embryonic day 13.5 (E13.5) and declines to low levels by postnatal day 1 (P1) (**Figure 1A**)^36^. At birth, *Zfp423* expression remains uniformly low across all postnatal endpoints. Consistent with this, merged t-SNE plot of Schwann cells specifically at P1, P5, P14 and P60 reveals expression of *Zfp423* is restricted to a few immature Schwann cells (iSC) (**Figure 1B**), suggesting that *Zfp423* expression is restricted to the prenatal period, being largely extinguished upon peripheral nerve maturation.

**Figure 1.**
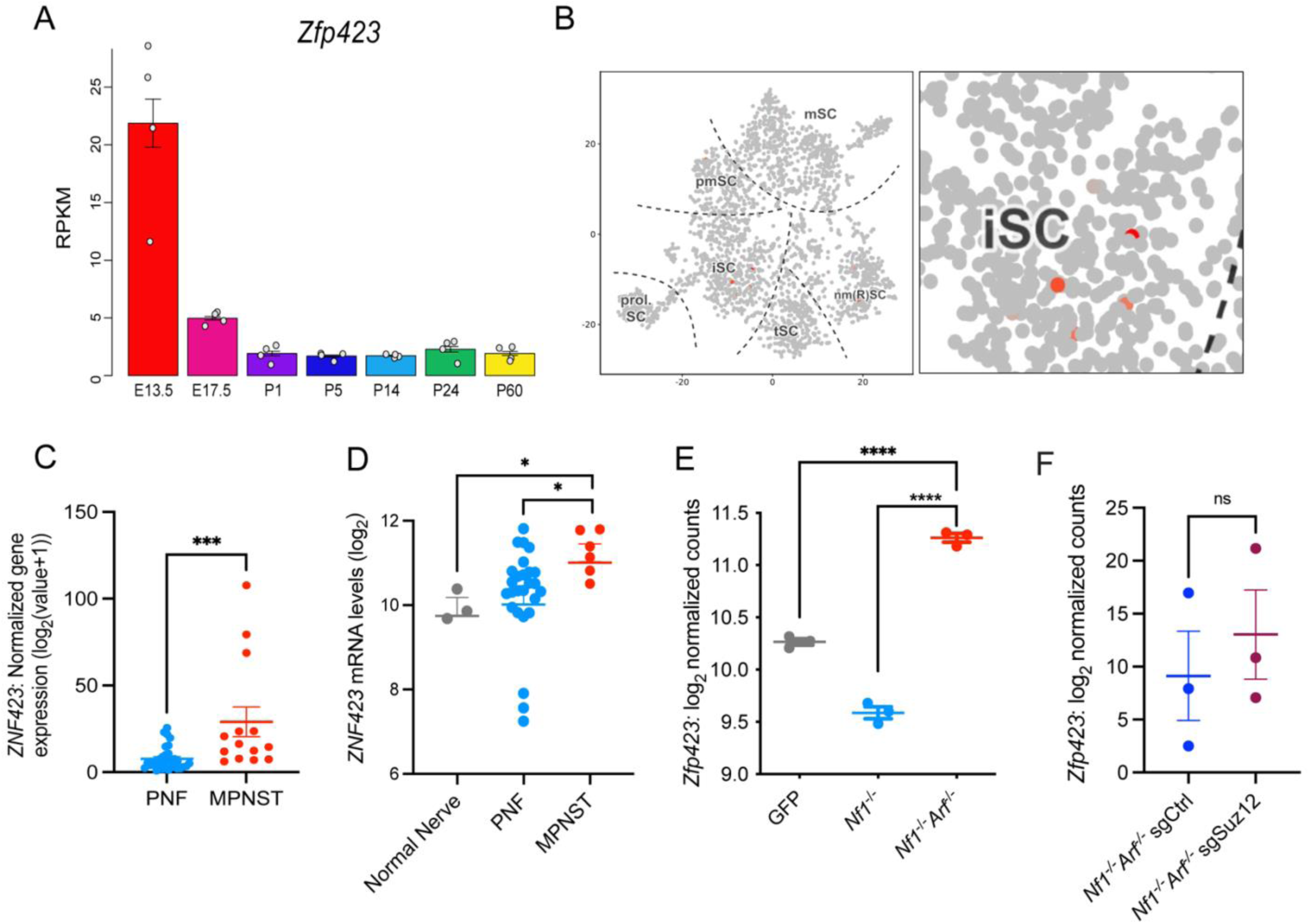
*ZNF423/Zfp423* is aberrantly upregulated in both human and genetically engineered mouse models (GEMM) of MPNST compared to PNF and normal nerves. (A) Bar plot showing Zfp423 transcript expression (RPKM) across embryonic (E13.5, E17.5) and postnatal (P1, P5, P14, P24, and P60) murine developmental timepoints. **(B)** Merged tSNE plot generated from scRNAseq data of murine Schwann cells at P1, P5, P14, and P60 showing Zfp423 expression restricted to the immature Schwann cells (iSC) postnatally. The color gradient of grey to red indicates relative transcript abundance of Zfp423 per cell, with grey representing no detectable expression and dark red indicating high expression. Data were obtained from the Sciatic Nerve ATlas (SNAT). Cell populations include: proliferating Schwann cells (prol. SC), immature Schwann cells (iSC), pro-myelinating Schwann cells (pmSCs), transition SCs (tSC), mature non-myelinating SCs (nm(R)SC) and myelinating SCs (mSC). **(C)** Dot plot comparing log_2_-normalized ZNF423 gene expression between PNF (*n*=32), and MPNST (*n*=14). P-values represent two-tailed unpaired t-test with Welch’s correction. **(D)** Dot plot comparing ZNF423 mRNA levels in normal nerve (*n*=3), PNF (*n*=26), and MPNST (*n*=6) (GEO accession: GSE41747). Error bars reflect standard error of the mean (SEM). P-values represent two-tailed unpaired t-test with Welch’s correction. **(E)** Dot plot depicting log_2_-normalized counts of Zfp423 for each sample (*n*=3 per group). Error bars reflect standard error of the mean (SEM). P-values represent unpaired, two-tailed t-test with Welch’s correction. **(F)** Dot plot depicting log_2_-normalized counts of Zfp423 following CRISPR-mediated knockout of Suz12 compared to control (*n*=3 per group). Error bars reflect standard error of the mean (SEM). P-values represent two-tailed unpaired t-test with Welch’s correction. ****=p-value≤0.0001, ***=p-value≤0.001, *=p-value≤0.05, ns=not significant.

Given this tightly regulated developmental expression pattern, we next examined *ZNF423* expression across stages of the PNST continuum. In an independent bulk RNAseq cohort comparing PNF and MPNST, *ZNF423* expression was significantly elevated in MPNST relative to benign tumors (**Figure 1C**)^37,38^. Analysis of a second independent dataset comprising normal peripheral nerves, PNF, and MPNST similarly revealed consistent *ZNF423* upregulation in human MPNST compared to PNF and normal nerve tissue suggesting that ZNF423 reactivation in MPNST may play a functionally relevant role in maintaining tumors (**Figure 1D**)^11^.

Building upon this observation, to further interrogate *ZNF423* regulation during MPNST progression we examined gene expression changes by RNA-seq in Schwann cell precursors (SCPs) derived from genetically engineered mouse models of NF1-tumorigenesis ^14^. Embryonic day 13.5 (E13.5) dorsal root ganglia (DRG) were harvested from *Nf1^flox/flox^* and *Nf1^flox/flox^;Arf^flox/flox^*mouse embryos, and either infected *ex vivo* with adenovirus expressing Cre recombinase (Ad-Cre-GFP) to excise floxed alleles or with control adenovirus (Ad-GFP) to serve as wild type controls ^26^. E13.5 DRG cultures are a tractable platform to study NF1 tumor progression because *Nf1^flox/flox^* and *Nf1^flox/flox^;Arf^flox/flox^*SCP represent the tumor cells-of-origin for PNF and ANNUBP/MPNST, respectively^3^. These models faithfully recapitulate the progression from PNF to MPNST in human patients, where the loss of CDKN2A similarly accompanies the development of ANNUBP/MPNST^4,14,39^. Among transcription factors (TFs) upregulated upon combined ablation of *Nf1* and *Arf*, *Zfp423* showed a significant increase in expression between PNF (*Nf1*^-/-^) and MPNST (*Nf1*^-/-^;*Arf*^-/-^) models (**Figure 1E**). Interestingly, we observed significantly lower *Zfp423* expression in our PNF model relative to wild-type controls but was significantly elevated in our MPNST model compared to both wild-type and PNF. Reduced Zfp423 expression in PNF is consistent with prior observations in neuroblastoma where NF1 loss repressed ZNF423^29^. The subsequent upregulation in MPNST suggests that an acquisition of transcriptional programs typically associated with development may accompany malignant transformation. Importantly, PRC2 loss has been implicated in the de-repression of developmental TFs in MPNST^21^. However, *Nf1^-/-^Arf^-/-^* cells do not exhibit inherent loss of PRC2 and CRISPR-mediated knockout of *Suz12* did not significantly alter *Zfp423* expression (**Figure 1F**). This suggested that functional PRC2 loss is not essential for the increase in *Zfp423* expression in this model. Consistent with these findings, restoration of *SUZ12* in two PRC2-deficient human NF1 MPNST cell lines (ST88-14 and NF.90.8) by ectopic expression did not result in a significant change in ZNF423 protein abundance (**Supplemental Figure S1A**). In contrast, transcriptomic profiling by RNAseq following *SUZ12* restoration identified 14 transcription factors commonly downregulated across both cell lines, among which was ZNF423 (**Supplemental Figure S1B-D**). This discrepancy between ZNF423 mRNA and protein levels may imply that ZNF423 is subject to post-transcriptional or post-translational regulation affecting protein abundance irrespective of changes in transcript levels. Taken together, our gene expression data suggests that ZNF423 upregulation is associated with the evolution of PNF to MPNST.

### ZNF423 depletion impedes DNA synthesis and proliferative capacity in MPNST

Given the observed upregulation of ZNF423 in human and murine MPNST, we first assessed ZNF423 protein abundance across a panel of 7 human MPNST cell lines. We observed variable protein levels, consistent with the known genetic and transcriptional heterogeneity of MPNST (**Figure 2A**)^24^. ZNF423 was detected in the majority of human MPNST cell lines, though notably absent in the JH-2-079 cell line, suggesting that while frequently expressed, it is not universal across every tumor. In contrast to our above findings, ZNF423 was also detected in the immortalized Schwann cell line (ipn02.3) and in one of two immortalized PNF lines (ipNF95.6, but not ipNF05.5). Given that these lines were immortalized by ectopic hTERT and mCdk4 expression, ZNF423 expression in these cells could reflect the reactivation of transcriptional programs associated with de-differentiation^40^. The use of overexpressed mCdk4 is akin to the loss of CDKN2A (and p16ink4a expression) observed in ANNUBP/MPNST, which both drive cell cycle progression. It is also possible that PNF cell lines such as ipNF05.5, which exhibited no observable ZNF423 expression despite similar immortalization techniques, acquire ZNF423-independent programs of de-differentiation to allow for survival and proliferation *in vitro*. Taken together, the prevalence of detectable ZNF423 across the majority of human MPNST cell models motivated further investigation into its functional role in MPNST growth.

**Figure 2.**
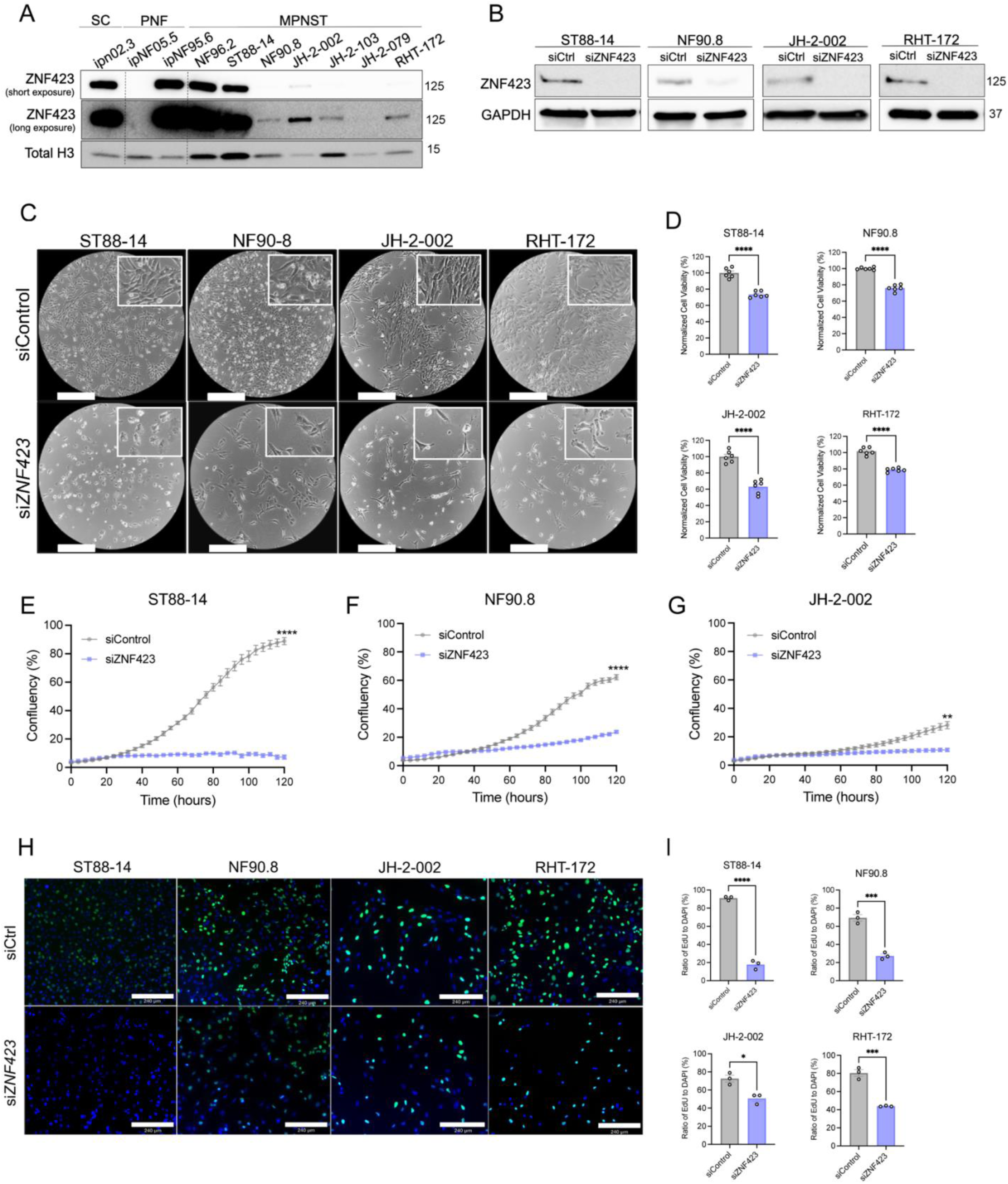
ZNF423 is required for MPNST cell viability and proliferation. **(A)** Representative immunoblot of ZNF423 protein expression in human Schwann, PNF, and MPNST cell lines. Total histone H3 serves as loading control. **(B)** Immunoblots showing ZNF423 in four human MPNST cell lines following siRNA-mediated depletion of ZNF423 compared to non-targeting control. GAPDH serves as loading control. **(C)** Representative phase-contrast images (4x) of ST88-14, NF90.8, JH-2-002, RHT-172 cells following siRNA-mediated depletion of ZNF423 compared to control showing reduced cell number and altered morphology. Scale bars represent 360μm. **(D)** Cell viability measured by CellTiter-Glo assay following ZNF423 knockdown (*n*=6). Error bars reflect SEM. P-values represent two-tailed unpaired t-test with Welch’s correction. **(E-G)** Cell confluency over time following transfection with siZNF423 or a non-targeting siControl, measured by live-cell imaging (incucyte) every 4 hours over 5 days (*n*=6). Error bars reflect SEM. P-values represent two-way repeated measures ANOVA followed by Bonferroni’s post hoc correction for multiple comparisons. **(H)** Representative immunofluorescent images (10x) showing DAPI stained nuclei in blue, and EdU positive cells in green. Scale bars represent 240 μm. **(I)** Quantification of EdU positive cells expressed as a percentage of total DAPI positive nuclei (*n*=3). Error bars reflect SEM. P-values represent two-tailed unpaired t-test with Welch’s correction. ****=p-value ≤0.0001, ***=p-value≤0.001, **=p-value≤0.01, *=p-value≤0.05, ns=not significant.

To evaluate ZNF423 in MPNST, we selected widely propagated human MPNST cell lines ST88-14 and NF90.8, and more recently derived patient MPNST cell lines, JH-2-002 and RHT-172, representing ZNF423 low-expressing and high-expressing cells, respectively. Efficient siRNA-mediated knockdown of ZNF423 was confirmed by immunoblotting (**Figure 2B**) and resulted in reduced cell density (**Figure 2C**). Depletion of ZNF423 led to a modest but significant reduction in MPNST cell viability compared to siRNA control populations in all cell lines tested (**Figure 2C-D**). Notably, this effect was observed even in cell lines with low baseline ZNF423 expression, suggesting MPNST are at least partially dependent on ZNF423 for proliferation regardless of relative expression level. Additionally, live-cell imaging revealed that ZNF423 depletion induced striking morphological changes in cells and reduced MPNST cell growth over 4-5 days (**Figure 2E-G**), demonstrating that ZNF423 is required for MPNST to maintain their full proliferative capacity.

ZNF423 has been implicated in the regulation of cell cycle progression by controlling DNA replication and S-phase dynamics during neural progenitor development^41^. We therefore hypothesized that the growth impairment following ZNF423 loss may reflect impaired or slowed DNA synthesis. To directly assess S-phase entry and progression we performed 5-ethynyl-2′-deoxyuridine (EdU) incorporation assays. EdU/DAPI staining revealed a significant reduction in EdU incorporation in ZNF423-depleted cells compared to controls, indicating diminished DNA replication activity and impaired proliferation (**Figure 2H-I**).

To determine whether this reduction in DNA synthesis reflected a broader defect in cell cycle progression, we performed cell cycle distribution analysis using propidium iodide (PI) staining and flow cytometry to evaluate DNA content and reveal the distribution of cells in G_0_/G_1_, S, or G_2_/M phase of the cell cycle. Despite the marked decrease in EdU incorporation, cell cycle distribution profiles were not significantly altered in ZNF423-depleted cells compared to controls (**Supplemental Figure S2**). Although not statistically significant, we observed a modest increase in the proportion of cells in S phase, which may reflect delayed S-phase progression, potentially due to replication stress or an accumulation of DNA damage. Collectively, these findings indicate that ZNF423 loss impedes DNA synthesis without inducing overt cell cycle arrest, and its loss attenuates the overall proliferative capacity of MPNST cells, potentially through disruption of replication rate rather than phase-specific blockade.

### ZNF423 coordinates proliferative and lineage-associated transcriptional programs in MPNST, and its loss activates stress-responsive and pro-apoptotic signaling pathways

To delineate the transcriptional programs underlying the phenotypic consequences of ZNF423 depletion, we performed RNA sequencing of two human NF1 MPNST cell lines (ST88-14 and NF90.8). Differential gene expression (DEG) analysis by DESeq2 revealed widespread transcriptomic reprogramming following ZNF423 knockdown in both models (**Figure 3A-B**) underscoring its regulatory influence. ZNF423 was not present in the DEG list for the NF90.8 cell line due to raw count cut-off filtering, but examination of raw count data confirmed reduced ZNF423 transcript abundance. The absence of ZNF423 from the DEG list likely reflects its comparatively low baseline expression in this cell line. Nine hundred seventy-seven and 2775 genes were significantly downregulated in the ST88-14 and NF90.8 cell lines, respectively, with 445 genes overlapping between the two lines (**Figure 3C**). Pathway enrichment analysis of the overlapping downregulated genes using the Molecular Signatures (MSigDB) Hallmark database revealed a significant suppression of developmental and proliferative signaling networks including Notch, IL6-JAK-STAT3, and Wnt/β-catenin, as well as a reduction in inflammatory response signatures (**Figure 3D**).

**Figure 3.**
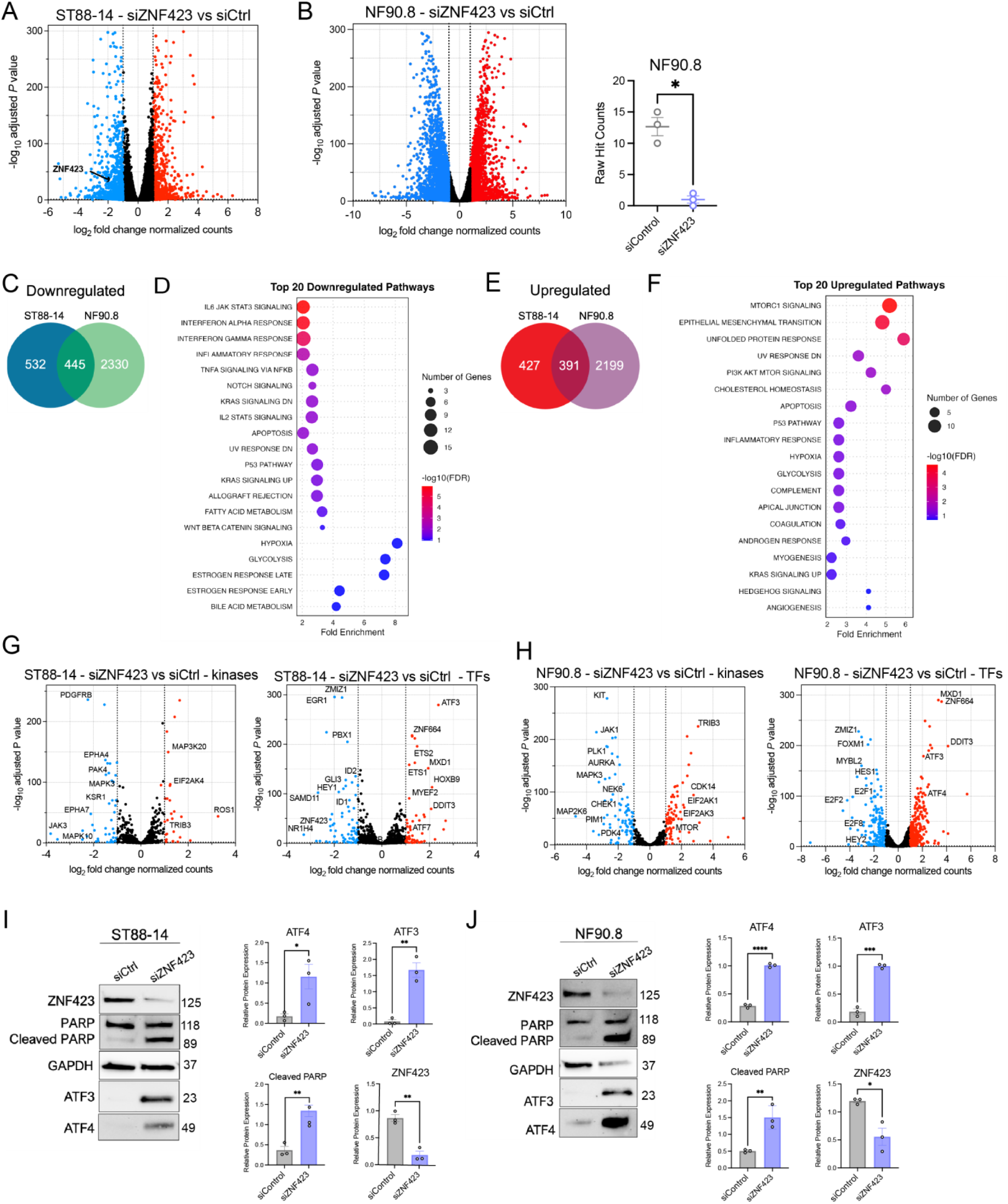
RNA sequencing of ZNF423-depleted cells reveals a stress-induced transcriptional phenotype characterized by activation of the integrated stress response. **(A-B)** Volcano plots of whole-transcriptome differentially expressed genes (DEGs) identified by RNAseq in ST88-14 and NF90.8 cells following siRNA-mediated depletion of ZNF423 (*n*=3) compared to control (*n*=3). Dashed vertical lines reflect log_2_FC of −1 and 1. Dashed horizontal line reflects false discovery rate (FDR) of 0.05. Downregulated genes with log_2_FC≤-1 and -log_10_ adjusted p-value above FDR threshold are denoted in blue. Upregulated genes log_2_FC≥1 and -log_10_ adjusted p-value above FDR threshold are denoted in red. Raw RNAseq hit counts for ZNF423 in NF90.8 cells transfected with control siRNA (*n*=3) or siZNF423 (*n*=3). P-values represent unpaired, two-tailed t-tests between groups. **(C)** Venn diagram showing overlap of significantly downregulated DEGs common among ST88-14 and NF90.8 cell lines. Downregulated DEGs were defined as log_2_FC≥-1 and adjusted p-value of ≤0.05. **(D)** Dot plot of top enriched Hallmark gene signatures of shared downregulated genes (FDR≤0.05). **(E)** Venn diagram showing overlap of significantly upregulated DEGs common among ST88-14 and NF90.8 cell lines. Upregulated DEGs were defined as log_2_FC≥1 and adjusted p-value of ≤0.05. **(F)** Dot plot of top enriched Hallmark gene signatures of shared upregulated genes (FDR≤0.05). **(G-H)** Volcano plots of differentially expressed transcription factors and kinases in ST88-14 and NF90.8 cells following siRNA-mediated depletion of ZNF423 compared to control. **(I-J)** Immunoblots showing cleavage of PARP, ATF3, ATF4 and ZNF423 protein expression following siRNA-mediated depletion of ZNF423 (*n*=3) compared to controls (*n*=3). GAPDH serves as loading control. Protein expression was normalized to total protein and quantified using ImageJ. Error bars reflect SEM. P-values represent unpaired, two-tailed t test with Welch’s correction. ****=p-value ≤0.0001, ***=p-value≤0.001, **=p-value≤0.01, *=p-value≤0.05, ns=not significant.

Conversely, a total of 818 genes were significantly upregulated in the ST88-14 cell line, while 2590 were upregulated in the NF90.8 cell line, with 391 genes overlapping (**Figure 3E**). Enrichment analysis of shared upregulated genes revealed significant activation of stress-associated pathways, including the unfolded protein response (UPR), epithelial-to-mesenchymal transition (EMT), mTORC1 signaling and the p53 pathway (**Figure 3F**). Notably, both cell lines showed a robust induction of genes associated with the integrated stress response (ISR), indicative of a conserved stress-adaptive program triggered by *ZNF423* loss.

To mechanistically connect the observed growth inhibition with these transcriptional shifts, we focused on differentially expressed TFs and kinases which function as key regulatory nodes controlling cell fate decisions and proliferative signaling. ZNF423 depletion downregulated critical regulators of cell cycle (kinases *AURKA, CHEK1, E2F2, CDK6*), dedifferentiation and developmental identity (TFs *ID1/2, HEY1/2, HES1, GLI3*) and survival signaling (kinases *PAK4, MAPK3, MAPK10*) (**Figure 3G-H**). This concerted attenuation of mitogenic and developmental regulators aligns with prior studies implicating ZNF423 in neurogenesis, embryogenesis, and cell cycle control, reinforcing its role as a central coordinator of proliferative and developmental transcriptional programs in MPNST^41–44^.

In contrast, genes upregulated following ZNF423-depletion were enriched for ISR-related factors, including bona fide stress-sensing kinases *EIF2AK4* (GCN2), *EIF2AK3* (PERK), and *EIF2AK1* (HRI) and the pseudokinase *TRIB3*. Downstream ISR transcriptional effectors *ATF3, ATF4,* and *DDIT3* (CHOP), which coordinate adaptation to proteotoxic or genotoxic stress, were also upregulated (**Figure 3G-H**)^45^. The ISR is initiated by phosphorylation of eIF2α by one of four key stress-sensing kinases (PERK, GCN2, PKR, HRI), which attenuate global protein synthesis while triggering selective translation of ATF4^46,47^. As the master regulator of the ISR, ATF4 modulates either pro-survival or pro-apoptotic signals. Under prolonged stress, ATF4 induces transcriptional activation of *ATF3, DDIT3,* and *TRIB3,* triggering pro-apoptotic cascades that eliminate damaged cells^48,49^. Consistent with the transcriptional data, immunoblotting confirmed increased ATF4 and ATF3 protein levels following ZNF423 depletion, accompanied by accumulation of cleaved PARP (**Figure 3I-J**) demonstrating that ISR activation induces apoptosis and impedes the proliferative capacity of MPNST cells.

Importantly, RNAseq analysis of additional patient-derived xenoline models (JH-2-002 and RHT-172) by RNAseq revealed a similar transcriptional reprogramming following ZNF423 depletion. While changes were not statistically significant across most ISR genes, JH-2-002 cells demonstrated modest, but significant induction of *ATF3* and *DDIT3* supporting partial engagement of the ISR (*ATF3* log₂FC=0.599, padj<0.05, *DDIT3* log₂FC=0.783, padj<0.05). This transcriptional signature was confirmed at the protein level, with immunoblot analysis confirming increased ATF3, accompanied by accumulation of cleaved PARP, further supporting activation of pro-apoptotic stress signaling (**Supplemental Figure S3**). Similarly, in RHT-172 cells, RNAseq demonstrated detectable induction of *ATF3*, *ATF4* and *DDIT3* (*ATF3* log₂FC=0.6, padj<0.05, *ATF4* log₂FC=0.2, padj<0.05, and *DDIT3* log₂FC=0.40, padj<0.05) as well as downregulation of proliferative and upregulation of stress-related gene programs following ZNF423 depletion (**Supplemental Figure S4**). Collectively, these data extend our findings across multiple in vitro models and reinforce ISR engagement as a conserved transcriptional program and mechanistically relevant consequence of ZNF423 depletion.

### Loss of ZNF423 impairs the DNA damage response and enhances sensitivity to cytotoxic and PARP-directed therapies in MPNST

Within the DNA damage response (DDR) pathway, ZNF423 functions as a scaffold protein, where it interacts with the DNA double-strand damage sensor poly (ADP) ribose polymerase 1 (PARP1), to facilitate the repair preservation of DNA integrity^50^. Since unresolved DNA damage can trigger ISR, we hypothesized that the stress phenotype observed upon depletion of ZNF423 in our MPNST cell models may reflect impaired DNA repair capacity and consequently DNA damage accumulation^51^.

Consistent with this hypothesis, depletion of ZNF423 led to increased levels of γH2A.x by immunoblotting (**Figure 4A-B**) and immunofluorescence staining (**Figure 4C**). These findings parallel prior observations in neural progenitor models where *Zfp423* mutations delayed cell cycle progression at the G_2_-M checkpoint, and elevated γH2A.x and 53BP1 foci^41^. Moreover, *Zfp423*-deficient P19 cells exhibit enhanced sensitivity to ionizing radiation, characterized by γH2A.x accumulation, suggesting that cells lacking ZNF423 not only have a reduced capacity to resolve DNA damage, but also have increased sensitivity to DNA damage-inducing agents^52^. These findings support a functional role for ZNF423 in maintaining genomic stability in MPNST, and that loss or inhibition of ZNF423 may sensitize MPNST cells to DNA-damaging therapies.

**Figure 4.**
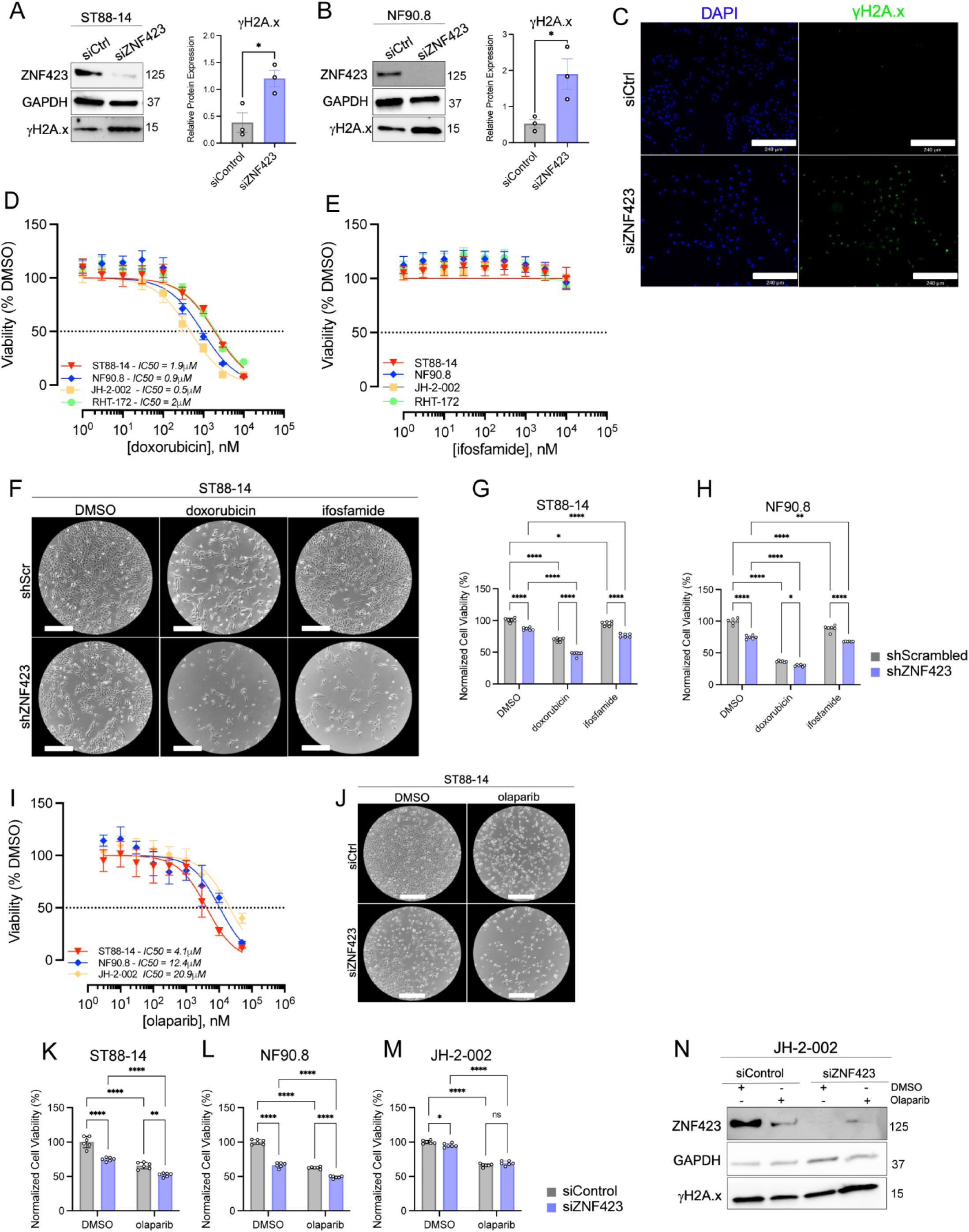
ZNF423 depletion increases DNA damage and sensitizes MPNST cells to cytotoxic and PARP-directed therapies. **(A-B)** Immunoblots showing increased protein levels of γH2A.x following siRNA-mediated depletion of ZNF423 (*n*=3) compared to control (*n*=3). GAPDH serves as loading control. Protein expression was normalized to total protein and quantified using ImageJ. Error bars reflect SEM. P-values represent unpaired, two-tailed t test with Welch’s correction. **(C)** Immunofluorescence staining for phosphorylated γH2A.x in ST88-14 cells (10x) following ZNF423 knockdown. DAPI stained nuclei are in blue, and phosphorylated γH2A.x positive cells are shown in green. Scale bars represent 240μm. **(D-E)** CellTiter-Glo viability assays were used to generate dose response curves for four MPNST cell lines treated with doxorubicin or ifosfamide. IC₅₀ values were calculated using nonlinear regression (log[inhibitor] vs. normalized response, variable slope model). Error bars reflect SEM. **(F)** Representative phase-contrast images of ST88-14 cells showing enhanced growth inhibition following combined ZNF423 knockdown and treatment with doxorubicin or ifosfamide compared to scrambled controls. Scale bars represent 360μm. **(G-H)** Quantification of cell viability by CellTiter-Glo assay following ZNF423 depletion in combination with doxorubicin or ifosfamide treatment (*n*=6). P-values represent two-way ANOVA with Tukey’s multiple comparisons tests across groups. Error bars reflect SEM. **(I)** Dose-response curves for three MPNST cell lines treated with the PARP inhibitor olaparib. IC₅₀ values were calculated using nonlinear regression (log[inhibitor] vs. normalized response, variable slope model). Error bars reflect SEM. **(J)** Representative images showing enhanced growth inhibition following ZNF423 siRNA-mediated knockdown combined with olaparib treatment. Scale bars, 360μm. **(K-M)** Quantification of cell viability by CellTiter-Glo assay following combined ZNF423 depletion and PARP inhibitor treatment (*n*=6). P-values represent two-way ANOVA with Tukey’s multiple comparisons tests across groups. Error bars reflect SEM. **(N)** Immunoblots of JH-2-002 cells showing increased γH2A.x following ZNF423 knockdown and treatment with olaparib. ****=p-value ≤0.0001, ***=p-value≤0.001, **=p-value≤0.01, *=p-value≤0.05, ns=not significant.

To determine whether this DDR impairment translated to improved therapeutic responsiveness, we evaluated the impact of ZNF423 depletion on sensitivity to DNA damaging agents. Despite their limited clinical efficacy due to intrinsic or acquired resistance, doxorubicin and ifosfamide are the current standard-of-care chemotherapies in the management of MPNST. As such, we selected these agents to assess whether ZNF423 modulation alters treatment response^53^. To establish baseline response to these agents, dose response curves were generated with increasing concentrations of doxorubicin (**Figure 4D**) or ifosfamide (**Figure 4E**), and IC_50_ values were calculated for each condition in a panel of human MPNST cell lines. Consistent with our hypothesis, treatment of ZNF423-depleted cells with doxorubicin or ifosfamide resulted in increased cellular rounding, and reduced confluence compared to control cells, reflecting enhanced cytotoxic stress and diminished proliferative capacity (**Figure 4F**). ATP-based cell viability assays were also performed to directly compare treatment effects between control and ZNF423-depleted cells. Across both agents, ZNF423 knockdown significantly potentiated drug-induced cytotoxicity in both cell models, resulting in further reductions in cell viability compared to treated controls (**Figure 4G-H**). These findings identify ZNF423 as a modulator of chemotherapeutic response in MPNST.

Given the established interaction between ZNF423 and PARP1^52^, we next evaluated whether loss of ZNF423 augments sensitivity to PARP inhibition. To determine the sensitivity of human MPNST cells to the FDA-approved PARP inhibitor olaparib (AZD2281), we generated dose response curves and calculated baseline IC50s for each line tested (**Figure 4I**). Treatment of ZNF423 knockdown cells with olaparib also resulted in morphologic changes including cellular rounding and decreased confluence (**Figure 4J**). Notably, ZNF423 knockdown significantly increased olaparib sensitivity in two of the three cell lines tested, with combined knockdown and treatment resulting in greater reductions in cell viability (**Figure 4K-M**). Although JH-2-002 cells did not demonstrate further reductions in short-term viability, ZNF423 depletion in this line enhanced γH2AX accumulation under PARP inhibition (**Figure 4N**) indicating exacerbated DNA damage despite preserved metabolic activity.

Collectively, these findings demonstrate that ZNF423 stabilizes DNA repair capacity in MPNST and suggest that its suppression lowers the threshold for therapeutic response to both conventional cytotoxic agents and PARP-directed strategies.

### Sustained ZNF423 suppression delays tumor initiation and impairs MPNST tumorigenicity in vivo

While *in vitro* assays demonstrated that ZNF423 is required for MPNST cell proliferation and induces activation of stress response pathways, we next sought to test whether sustained ZNF423 suppression similarly impaired tumor growth *in vivo*. To assess long-term effects of *ZNF423* knockdown, we generated stable shRNA-mediated knockdown models in ST88-14 MPNST cells which we confirmed by immunoblotting (**Figure 5A**). Stable *ZNF423* knockdown recapitulated the phenotypes observed following transient siRNA-mediated knockdown, with cells exhibiting altered morphology, reduced cell size, and diminished proliferative capacity *in vitro* (**Figure 5B**). Prior to *in vivo* implantation, we characterized the transcriptional impact of sustained *ZNF423* suppression which recapitulated key features of the transient knockdown, including induction of stress-associated gene programs such as the unfolded protein response (**Supplemental Figure S5**). Notably ISR-related transcripts including *ATF3* and *ATF4* were modestly induced but not statistically significant. Pathways such as E2F Targets and G2/M checkpoint were also upregulated, suggesting that stable knockdown of *ZNF423* (as opposed to transient knockdown) could select for surviving cells that are persisting, despite their impaired proliferative capacity.

**Figure 5.**
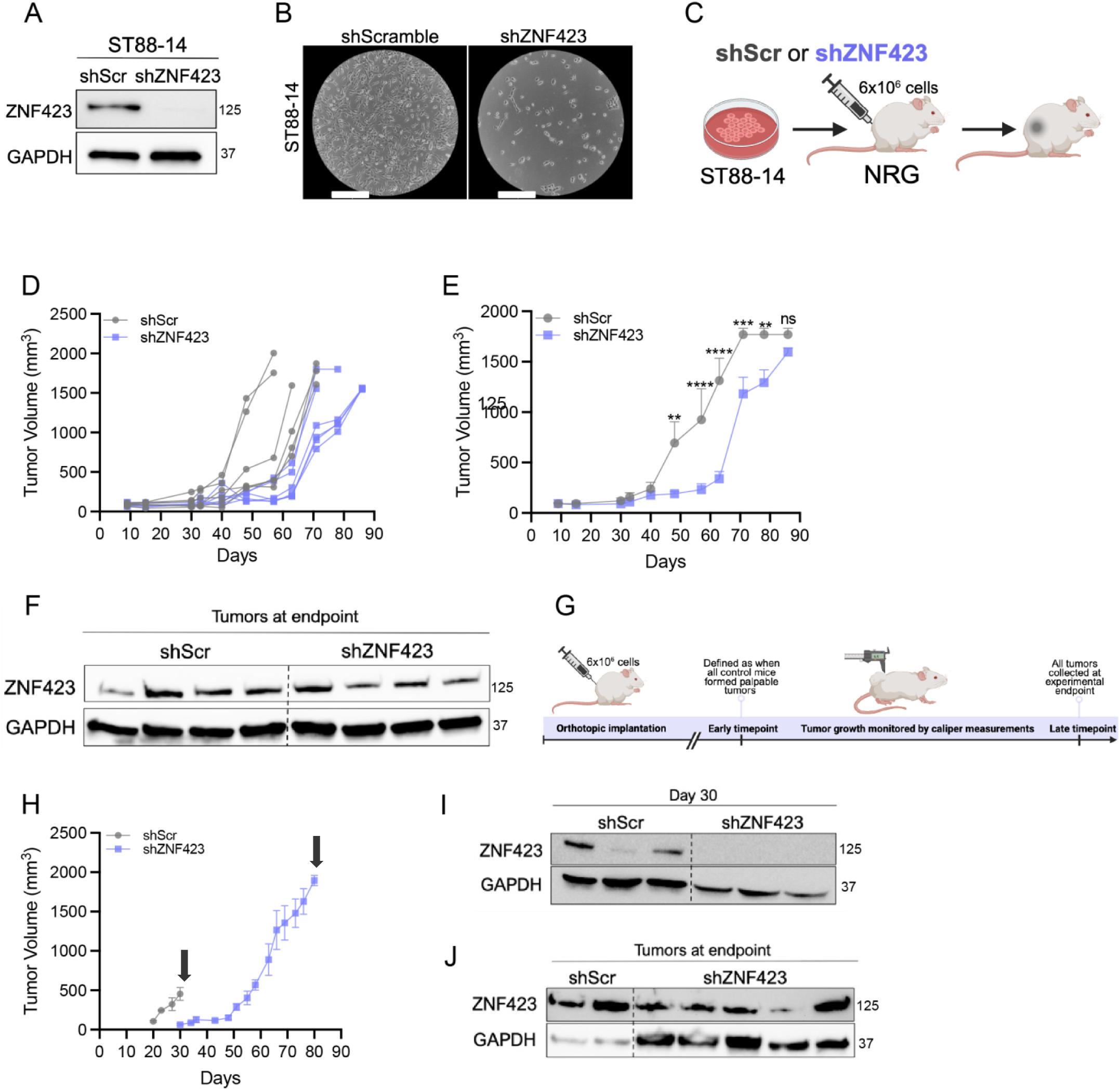
ZNF423 knockdown delays MPNST tumor growth in mice. **(A)** Immunoblot confirming shRNA mediated knockdown of ZNF423 (shZNF423) in ST88-14 cells compared to scrambled control (shScr). GAPDH serves as loading control. **(B)** Representative phase-contrast images showing reduced cell density and altered cellular morphology following ZNF423 knockdown. **(C)** Experimental schematic of orthotopic implantation of 6 × 10^6^ shScr or shZNF423 ST88-14 cells into the sciatic nerve of NRG mice. **(D)** Individual tumor growth trajectories showing variability in tumor growth among shScr mice (*n*=6) and shZNF423 mice (*n*=6). **(E)** Tumor volume over time showing delayed tumor growth in shZNF423 tumors relative to shScr controls. Error bars reflect SEM **. (F)** Immunoblots of tumors collected at experimental endpoints demonstrating restoration of ZNF423 expression in shZNF423 tumors. **(G)** Schematic of the second in vivo experiment defining early and late tissue collection time points. Early time point was defined as when all control tumors were palpable. Late time point was humane endpoint. **(H)** Tumor volume over time of shScr (*n*=11) and shZNF423 (*n*=11). Arrows indicate time points at which tumors were harvested. Error bars reflect SEM. **(I)** Immunoblots of tumors collected at the early timepoint (day 30) showing sustained ZNF423 knockdown in shZNF423 tumors. **(J)** Immunoblots of tumors collected at endpoint confirming re-expression of ZNF423 in shZNF423 tumors. P-values represent two-way ANOVA with repeated measures followed by Šídák’s multiple comparisons test between groups at each time point. ****=p - value ≤0.0001, ***=p-value≤0.001, **=p-value≤0.01, *=p-value≤0.05, ns=not significant.

After confirming both the phenotypic and transcriptional effects of stable ZNF423 suppression, we selected the ST88-14 cell line for orthotopic implantation based on its robust tumorigenic capacity and established use in NOD.Cg-*Rag1^tm1Mom^ Il2rg^tm1Wjl^*/SzJ (NRG) mouse models (**Figure 5C**)^7^. To preserve knockdown efficiency and minimize loss of shRNA expression prior to implant, cells expressing either scrambled control or ZNF423 shRNA were harvested immediately following puromycin selection and implanted into the sciatic nerve of NRG mice. Control cells efficiently formed tumors 30 days post-implantation, whereas tumor formation was markedly delayed in ZNF423-silenced cells (**Figure 5D**). Individual tumor growth trajectories revealed that control tumors grew rapidly and reached humane endpoints earlier, whereas most ZNF423-depleted tumors remained stable or regressed until week 8, after which they grew rapidly to humane endpoint (**Figure 5E**). Immunoblotting of tumors harvested at experimental endpoints (1,500-2,000mm^3^ tumor volume) revealed shZNF423 tissue had restored ZNF423 expression, suggesting that the eventual tumor outgrowth resulted from loss of shRNA-mediated suppression and escape from selective pressure (**Figure 5F**). We have observed that the antibody used to detect ZNF423 does not react with murine ZNF423 (ZFP423) (**data not shown**), ensuring that the source of the observed ZNF423 protein was the implanted ST88-14 cells.

To confirm our hypothesis that the tumor regrowth coincided with the reappearance of ZNF423 expression, we performed a second orthotopic experiment to monitor ZNF423 expression at early and late timepoints (**Figure 5G**). In this experiment, mice were monitored until all control animals developed palpable tumors (week 4). ZNF423-depleted mice showed no detectable tumor growth at this timepoint, consistent with our previously observed delay in tumor formation (**Figure 5H**). To determine whether ZNF423 expression remained suppressed at this early timepoint prior to potential outgrowth, we harvested tumors from control mice and collected tissue from the implantation sites of five ZNF423-depleted mice. Western blot analysis of these tissues showed robust expression of ZNF423 in the controls, while expression remained undetectable in the ZNF423-depleted tumors (**Figure 5I**). The remaining shZNF423 mice were monitored until they reached experimental endpoints, where immunoblotting showed restored ZNF423 expression, confirming eventual tumor growth is only observed when re-expression of ZNF423 can be detected (**Figure 5J**). Together, these data demonstrate that sustained suppression of ZNF423 significantly impairs tumor initiation and early growth in an orthotopic MPNST model, establishing ZNF423 as a critical dependency for MPNST tumor maintenance *in vivo* and a potentially exploitable vulnerability.

### ZNF423 defines a developmentally reprogrammed late malignant cell state in human MPNST

While the preceding findings established ZNF423 as a central regulator of proliferative and stress-associated programs in MPNST models, we next examined its clinical relevance in human tumors. We leveraged publicly available human single-cell RNA sequencing datasets to interrogate *ZNF423* expression directly in patient-derived peripheral nerve tumors. We compiled, integrated, and batch-corrected datasets derived from multiple independent cohorts encompassing 46 PNF and 25 MPNST samples (363,793 cells, 10,000 highly variable genes) (**Figure 6A-B**). Pseudobulked differential expression analysis revealed that *ZNF423* expression was significantly elevated in MPNST relative to PNF (*P* = 8.9 × 10^-22^) (**Supplemental Figure S6A, Supplemental Table 1**) supporting its enrichment in malignant tumor states. Overlay of tumor type demonstrated that PNF and MPNST cells occupy overlapping transcriptional space, consistent with their shared SC lineage origin, while unsupervised Leiden clustering revealed a heterogeneous tumor environment characterized by multiple transcriptionally distinct cellular populations (**Figure 6C**). Clusters were annotated based on the expression of established lineage and cell-type specific marker genes (**Supplemental Table 2 and 3).** These clusters broadly included malignant cells, Schwann cells (SC), fibroblasts, endothelial cells, B cells, T cells, smooth muscle cells and mast cells. Annotation was validated by marker gene expression mapped onto UMAP projections (**Supplemental Figure S7**).

**Figure 6.**
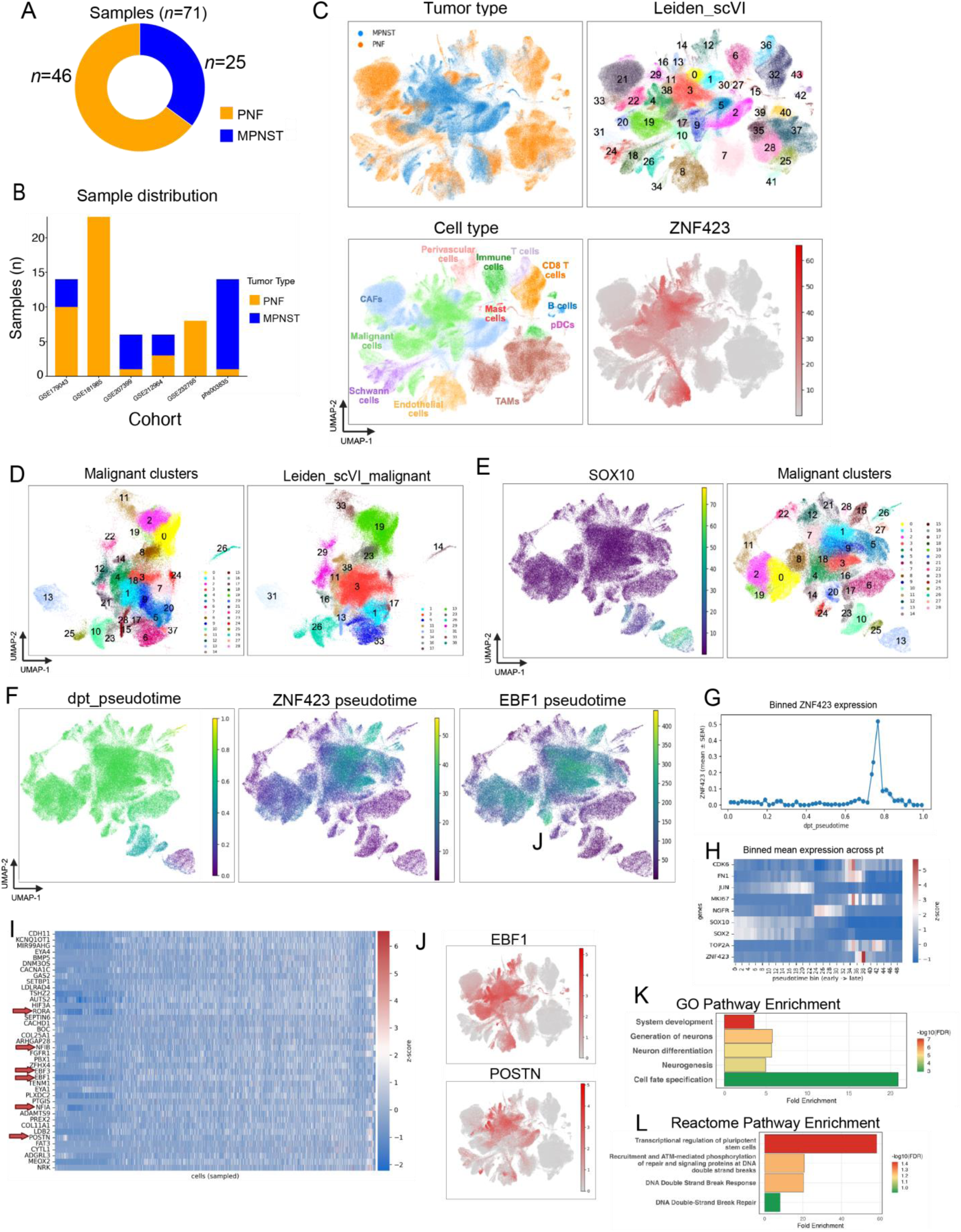
Single-cell RNA sequencing of human PNF and MPNST tumors identifies ZNF423-associated malignant cell states and pseudotime progression. **(A)** Circle plot showing the total number of samples divided into PNF (*n*=46) and MPNST (*n*=25). **(B)** Bar plot of the number of samples (y-axis) obtained from each cohort (x-axis). Orange bars represent PNF samples and blue bars represent MPNST samples. **(C)** UMAPs of batch-normalized scRNAseq data comprising 363,793 cells and 10,000 highly variable genes (HVGs). Top left: UMAP colored by tumor type of origin (orange, PNF; blue, MPNST). Top right: UMAP showing 43 clusters identified using Leiden resolution 1.00. Bottom left: UMAP of cell type annotations. Cell type was manually assigned using inferCNV score, percent expression of marker genes (Supplemental Table 2), and z-score of the scanpy-calculated marker panel scores. Bottom right: UMAP of scVI-normalized ZNF423 RNA expression. **(D)** Left: UMAP of clusters identified by PAGA analysis within the malignant annotation (*n* = 105,062 cells). Right: Corresponding Leiden (resolution 1.00) cluster numbers. **(E)** Malignant PAGA clusters arranged by pseudotime. Left: SOX10 scVI-normalized RNA expression. Right: Overlay of cluster numbers. **(F)** Left: Malignant PAGA clusters ordered by pseudotime. The root cluster was defined as the cluster with the lowest cycling score. Pseudotime 0 represents early-stage malignant clusters and pseudotime 1 represents late-stage clusters. Middle: ZNF423 and Right: EBF1 scVI-normalized RNA expression along pseudotime in malignant clusters. **(G)** Binned expression of ZNF423 across pseudotime. Pseudotime was divided into 50 bins and the z-score of the mean ZNF423 expression across bins was calculated. Plot shows pseudotime (x-axis) versus z-scored mean expression. **(H)** Binned mean expression of selected marker genes across pseudotime in malignant cells. Pseudotime was divided into 50 bins and the z-score of the mean expression for each gene across bins was calculated. Heatmap displays genes (y-axis) across pseudotime (x-axis; early to late). **(I)** Heatmap of the z-score of the top 40 genes most positively correlated with ZNF423 expression across malignant cells (Spearman correlation, *p* ≤ 0.05). Rows represent genes and columns represent individual malignant cells ( *n* = 105,062 cells). **(J)** UMAP of log1p-normalized RNA expression of EBF1 (top) and POSTN (bottom) across all cells. **(K)** GO pathways analysis of top 40 genes correlated with ZNF423 in malignant cells (FDR≤0.05). **(L)** Reactome pathway analysis of top 40 genes correlated with ZNF423 in malignant cells (FDR≤0.05).

To ensure accurate classification of malignant populations within this heterogeneous environment, inferCNV analysis was performed to assess copy number variations (CNV) across clusters (**Supplemental Table 4)**. Clusters exhibiting inferCNV scores associated with genomic instability were classified as malignant using predefined thresholds (non-malignant ≤0.02; malignant ≥0.04) which was resolved based on CNV heatmap patterns (**Supplemental Figure S6B**). Integration of inferCNV profiles with marker-based annotation enabled robust discrimination of malignant from non-malignant populations. Importantly, *ZNF423* expression was heterogeneous within the malignant cells and was minimal or absent in non-malignant populations, including PNF-derived SC populations, further supporting its activation is specifically associated with malignant transformation rather than benign nerve sheath biology (**Figure 6C**). Notably, *ZNF423* was detected in a subset of endothelial and perivascular cells, which is consistent with prior reports demonstrating *Zfp423* localization to defined endothelial cell populations during development^54^.

To resolve intratumoral heterogeneity within the malignant populations, malignant cells were subset and reclustered prior to PAGA analysis, revealing multiple transcriptional states (**Figure 6D**). We next sought to determine whether these malignant transcriptional states reflected ordered transitions along a malignant continuum. To establish a biologically relevant starting point for trajectory inference, we examined canonical cell cycle and SC lineage markers across the malignant clusters. Cycling genes (*MKI67, TOP2A, UBE2C, BIRC5, CENPF*) were used to define end-stage malignant clusters. Pseudotime (PT) was subsequently computed using PAGA for the malignant cells relative to this reference state, ordering malignant cells along a continuous transcriptional trajectory with the lowest cycling score constituting the root cluster. Cluster 13, which corresponds to cluster 31 of the initial clustering, was identified as the root cluster, with a PT of 0 and low expression of cycling genes. Notably, this cluster also retained strong *SOX10* expression, consistent with a preserved SC lineage identity, and therefore represented the earliest malignant state (**Figure 6E**). Diffusion pseudotime (DPT) analysis ordered this SC population along a continuous transcriptional trajectory, revealing a gradient from a distinct region of low PT (early) to high PT (late), with most malignant cells distributed across mid-to-late PT values (**Figure 6F**). Overlay of *ZNF423* and *EBF1* onto the PT UMAP demonstrated their activation is spatially restricted within the late malignant state.

Specifically, mapping *ZNF423* onto this malignant trajectory revealed minimal expression across early PT states (0.0-0.6) followed by a sharp, restricted activation within a late PT interval (0.7-0.8), resulting in a moderately positive correlation with PT (rho=0.26) (**Supplemental Figure S6C**). Binned expression analysis confirmed *ZNF423* expression within this narrow PT window, demonstrating that *ZNF423* does not increase gradually with progression but instead marks a discrete transcriptional state along the malignant continuum (**Figure 6G**).

Because late PT states were enriched for proliferating cells and cycling scores peaked within the same late PT interval, we next evaluated the relationship between PT and cell cycle activity. The overall correlation between cell cycle score and PT was weak (rho=–0.048), validating that PT does not simply reflect cell cycle, and is a true reflection of time (**Supplemental Figure S6D**). Heatmap visualization of marker genes across PT further supported this transition (**Figure 6H**), with early states retaining high *SOX10* and *SOX2* expression consistent with SC lineage identity, whereas mid-trajectory states were enriched for *NGFR* suggestive of a dedifferentiated phenotype. Late PT states were enriched for canonical proliferation markers (*MKI67, TOP2A CDK6*), and concurrent ZNF423 activation (**Supplemental Figure S6E**). *ZNF423* peaked at bin 38, aligning with proliferative markers. However, direct comparison of *ZNF423* expression and cell cycle score revealed a weak correlation (rho=0.021), suggesting *ZNF423* activation is not strongly associated with cell cycle even though they peak at the same PT window (**Supplemental Figure S6F**). These findings suggest that *ZNF423* activation is not a direct marker of cell cycle activity but rather delineates a discrete malignant state that is permissive for proliferation.

To define the malignant transcriptional state associated with ZNF423 activation, we analyzed gene networks that correlated with *ZNF423* in malignant cells (**Supplemental Table 5**). The top 40 co-expressed genes formed a transcriptional program enriched for developmental transcription factors and neuronal regulatory genes, including established ZNF423 interactors *EBF1* and *EBF3*, as well as *NFIA, NFIB, PBX1*, and *RORA* (**Figure 6I**). Importantly, all genes remained significantly correlated with *ZNF423* after adjusting for cell cycle score using partial Spearman correlation. ZNF423 expression did not positively correlate with *SOX10* (Spearman rho=-0.219), demonstrating that *ZNF423* activation represents a transcriptional state distinct from benign Schwann lineage identity (**Supplemental Figure S6G**).

Consistent with this, interrogation of the SNAT demonstrated that several *ZNF423*-associated genes including *Ebf1*, *Ebf3*, and *Postn* peaked during embryonic peripheral nerve development and declined to minimal levels postnatally (**Supplemental Figure S8**). Overlay of UMAPs further reinforced this concept, revealing overlapping expression of *ZNF423* with *EBF1* and *POSTN* within malignant cells of MPNST (**Figure 6J**). This coordinated expression suggests that *ZNF423* reactivation in MPNST reflects engagement of a developmentally restricted transcription program normally confined to immature peripheral nerve. In the context of malignant progression, this developmental rewiring event appears to re-establish a transcriptional state that allows proliferative expansion.

Gene ontology (GO) enrichment analysis of *ZNF423* co-expressed genes revealed significant enrichment for system development, generation of neurons, neuron differentiation, neurogenesis, and cell fate specification, consistent with the reactivation of developmental transcriptional programs (**Figure 6K**). Reactome pathway analysis further identified enrichment of transcriptional regulation of pluripotent stem cells, DNA double strand break response, and DNA double strand break repair, aligning with established roles of ZNF423 in recruitment of DNA repair genes like *PARP1* (**Figure 6L**)^50^. Notably, enrichment of DNA damage response programs directly parallels our functional data demonstrating increased γH2AX accumulation and enhanced sensitivity to DNA-damaging agents and PARP inhibition upon ZNF423 depletion.

Taken together, our findings demonstrate that ZNF423 defines a dominant late malignant transcriptional state characterized by coordinated activation of developmental transcription factors, proliferative expansion, and DNA damage response programs. ZNF423 activation is independent of cell cycle dynamics and SC lineage loss and instead marks a malignant reprogramming event that enables proliferative capacity within the evolving PNF-MPNST continuum that confer a targetable dependency, either alone or in combination with drugs such as PARPi.

## DISCUSSION

Accumulating evidence suggests that transcriptional dysregulation is a defining feature of NF1-associated tumor progression. Prior work from our group has demonstrated that overexpression of developmentally restricted proteins such as DLK1 marks more aggressive MPNST subsets with worse clinical outcome and is associated with reactivation of embryonic transcriptional programs^26^. In addition, complementary single-cell analyses have further revealed that progression along the PNF to MPNST continuum is marked by increased phenotypic heterogeneity accompanied by loss of SC identity and emergence of malignant stem-like states^24^. Despite these advances, the transcriptional dependencies that sustain tumor growth and confer therapeutic resistance in MPNST remain incompletely defined. Standard chemotherapy has minimal to no long-term impact on overall survival, underscoring the urgent need to identify biomarkers and potential vulnerabilities to improve medical therapies. Given that dedifferentiated transcriptional programs are co-opted by numerous cancers, we reasoned that identifying key lineage-restricted TFs in NF1 MPNST might reveal actionable vulnerabilities.

Here, we examined the function of ZNF423, a multifunctional Krüppel-like C2H2 zinc finger TF that is preferentially expressed in immature neuronal and glial cell populations^43,55^. ZNF423 integrates multiple developmental signaling pathways through interactions with EBF proteins, bone morphogenetic protein (BMP)-dependent SMADs, Notch, retinoic acid receptors and PARP1, and exhibits context-dependent roles in cancer^28,32–35^. By combining human cell and mouse model systems with single-cell profiling of clinical specimens, we present evidence that ZNF423 is a developmentally restricted, tumor-enriched transcriptional dependency that becomes selectively reactivated during a late malignant state within the PNF to MPNST continuum. Specifically, our studies revealed that ZNF423 is functionally required for proliferative competence and DNA damage response programs. Sustained suppression of ZNF423 slows tumor growth in MPNST xenograft models and sensitizes MPNST cells to both cytotoxic chemotherapy and PARP inhibition. Importantly, the functional consequence of ZNF423 depletion was validated and comparable in a panel of four human MPNST cell lines that exhibit heterogeneous growth rates and ZNF423 endogenous expression, further supporting that ZNF423 may represent a biologically meaningful axis for molecular stratification and therapeutic exploitation for these refractory tumors.

These findings extend an emerging paradigm in oncology focused on targeting dysregulated gene expression programs, with particular emphasis on developmental transcription factors that stabilize malignant states^56^. ZNF423 has long been recognized as a regulator of neuronal and glial lineage specification^43,55^, and its reactivation in MPNST reflects re-engagement of embryonic transcriptional circuitry, consistent with mounting evidence that developmental reprogramming promotes heterogeneity in NF1-driven tumors^26^. This resonates with emerging efforts to exploit developmental reprograming therapeutically, to shift tumor cells toward a more mature, less proliferative state^57^. Neuroblastoma, another dedifferentiated pediatric tumor, can sometimes be driven to a more differentiated state through treatment with ATRA^27^. Attempts to target similar differentiation programs in MPNST using ATRA have yielded mixed results^30^.

Despite advances in our molecular understanding, systemic therapy in MPNST largely remains dependent on DNA-damaging cytotoxic agents, with limited durable responses and poor overall survival^58^. Doxorubicin- and ifosfamide-based regimens remain the backbone of systemic treatment, yet intrinsic and acquired resistance are common. Our data nominate ZNF423 as a candidate therapeutic vulnerability that may modulate response to these agents. We demonstrate that ZNF423 suppression amplifies γH2A.x accumulation and enhances MPNST chemosensitivity, and that this vulnerability extends beyond conventional cytotoxic agents.

PARP inhibitors, including olaparib (AZD2281), exploit defects in DNA repair through synthetic lethality and have transformed the treatment landscapes of cancers with DNA repair deficiencies, most notably BRCA1/2-mutant breast, ovarian, pancreatic and prostate cancers^59^. Although PARP inhibition has shown limited efficacy as a monotherapy in MPNST, prior preclinical work demonstrated that olaparib reduces MPNST cell proliferation and promotes apoptosis, supporting further exploration of DNA repair-directed strategies in this disease^60^. Our data extend these observations by demonstrating that attenuation of ZNF423 enhances PARP inhibitor sensitivity and exacerbates DNA damage accumulation, suggesting that ZNF423-dependent tumors may be particularly susceptible to PARP-directed therapy. ZNF423 status may stratify tumors based on intrinsic DNA repair capacity and identify patients most likely to benefit from DNA damage-inducing or PARP-targeted combination strategies.

Sustained ZNF423 suppression in orthotopic xenograft models significantly delayed tumor initiation, with tumor regrowth occurring only upon restoration of ZNF423 expression, reinforcing that ZNF423 is required for maintenance of tumorigenic competence. While the siRNA- and shRNA-based suppression strategies employed here represent proof-of-principle rather than deployable therapeutic regimens, the perception of transcription factors as “undruggable” is rapidly dissolving. Proteolysis-targeting chimeras (PROTACs)^61^, molecular glues^62^, and small molecules designed to disrupt transcription factor-cofactor interactions^63^ have demonstrated clinical feasibility. Several cancer subtypes are now recognized as transcription factor-driven malignancies with emerging strategies to target these dependencies. Specifically, selective degradation of zinc finger TFs IKZF1 and IKZF3 in multiple myeloma by lenalidomide, an FDA-approved molecular glue^64^, illustrates that zinc finger TFs can be selectively targeted. In the context of ZNF423, both targeted degradation and disruption of critical cofactor interactions, including PARP1, represent rational therapeutic strategies.

The re-emergence of a developmentally restricted transcription factor raises important questions regarding the epigenetic mechanisms regarding its activation. The mechanistic consequences of PRC2 loss in MPNST have been well documented, with loss of H3K27me3 repressive mark and gain of H3K27ac associated with activation of developmentally regulated genes that transcriptionally resemble mesenchymal states^17,65^. Others have reported ZNF423 among transcription factors upregulated in PRC2-deficient cell lines and patient tumors supporting its activation in PRC2-loss transcriptional signatures^17,20,21^. However, in our models, ZNF423 was not strictly dependent on PRC2 status, as restoration of *SUZ12* did not consistently suppress ZNF423 protein levels. It remains possible that PRC2 deficiency contributes to a broader epigenetic context that permits developmental transcription factor activation, but additional regulatory mechanisms are likely involved. Clarifying whether ZNF423 is directly subject to PRC2-mediated repression will require focused analyses of PRC2 occupancy and H3K27me3 enrichment at the ZNF423 locus and associated regulatory elements.

While this study presents several novel findings, we acknowledge several limitations. Our inability to identify a reliable antibody to detect murine ZFP423 in mouse tumor tissues from our genetically engineered mouse models of PNF and MPNST limited *in situ* validation of ZFP423. Despite testing multiple commercially available antibodies in mouse tissues with reported high endogenous ZFP423 expression, we were unable to obtain specific and reproducible immunostaining. Furthermore, the antibody used to detect human ZNF423 protein was tested but not suitable for immunohistochemical staining of patient tissues. This reagent will be critical to pursue broader analyses of patient NF1 tumors and explore the use of ZNF423 as a biomarker. Beyond these technical constraints, the present work establishes ZNF423 as a tumor dependency and modulator of therapeutic response. Future studies should prioritize combinatorial in vivo testing integrating ZNF423 suppression with anthracycline-based chemotherapy or PARP inhibition in orthotopic and genetically engineered NF1 models. Given the developmental role of ZNF423 and its integration within retinoic acid signaling networks, stratified evaluation of differentiation-based approaches such as ATRA in ZNF423-expressing tumors may also be warranted. Acknowledging these limitations, our data nonetheless establish ZNF423 as a state-defining regulator of proliferative and DNA repair programs in NF1-MPNST and support its exploitation as a therapeutic vulnerability.

## MATERIALS AND METHODS

### Culture of human MPNST cell lines

Human MPNST cell lines JH-2-002, JH-2-103 and JH-2-079 were obtained from the Johns Hopkins NF1 Biospecimen Repository^37,38^. The NF90.8 line was obtained from Dr. Verena Staedtke (Johns Hopkins University). RHT-172 cells were derived from patient-derived xenografts of NF1-MPNST from Dr. Melissa Fishel (Indiana University)^66^. ST88-14 MPNST cells were obtained from Dr. Andrew Tee (Cardiff University). Immortalized human Schwann cell lines (iPNF02.3, iPNF05.5, iP95.6) were obtained from Dr. Margaret Wallace (University of Florida). Cells were cultured in either DMEM (ST-8814, NF90.8, iPNF02.3, iPNF05.5, iP95.6), DMEM/F12 (JH-2-002, JH-2-103, JH-2-079) or RPMI (RHT-172) media supplemented with 10% FBS (Harvest MidSci), 1% glutamine (Gibco), 1% penicillin/streptomycin (Lonza). Cells were authenticated by STR profiling, regularly tested for mycoplasma and confirmed negative (InvivoGen MycoStrip, Cat. rep-mys-10) and maintained in 5% CO_2_ at 37°C. Trypsin-EDTA 0.05% (Gibco) was used to dissociate cells for passaging upon reaching confluence.

### Surgical implantation of MPNST cells

All animal studies were performed in accordance with the guidelines of Institutional Animal Care and Use Committee (IACUC) of Indiana University School of Medicine (IACUC protocol #22027). NOD.Cg-*Rag1^tm1Mom^ Il2rg^tm1Wjl^*/SzJ (NRG) male and female mice were used for all studies and were obtained from the on-site breeding colony maintained by the Preclinical Modeling and Therapeutics Core (PMTC) at the IUSCCC. Tumor cells were orthotopically implanted into the sciatic nerve as described previously. Recipient mice were anesthetized, and scalpel and forceps were used to dissect through the dorsal skin of the thigh, exposing the sciatic nerve. 6×10^6^ cells in 40μL of Leibovitz’s L-15 media (11415064, ThermoFisher) were implanted into the muscular pocket surrounding the sciatic nerve, and the skin was closed with surgical wound clips, that were removed after seven days. Animals were monitored biweekly for tumor development using digital calipers. Tumor volume was calculated using the ellipsoid formula of V= (length x width^2^) / 2. Mice were euthanized once tumor volumes exceeded 1,500mm^3^ or other signs of distress meeting humane endpoints.

### siRNA transfection

Human MPNST cell lines were reverse transfected with 2μM siRNA against ZNF423 or negative control using Lipofectamine RNAiMAX transfection reagent (13778150, Invitrogen). For ST88-14 cells, a pooled ZNF423 siRNA was used (sc-38144, Santa Cruz Biotechnology). For NF90.8, JH-2-002 and RHT-172 cells, a single ZNF423 siRNA sequence was used (SASI_Hs01_00160938, Sigma-Aldrich). Corresponding controls were CtrlA (sc-37007, Santa Cruz Biotechnology), or negative control (SIC001-10NMOL, Sigma-Aldrich). Cells were plated in the transfection complex at a density of 1×10^6^ cells per 10cm plate or 500,000 cells per 6cm plate, and cell lysates were collected 72 hours post-transfection to confirm protein knockdown by western blotting.

### Viral production and transduction of MPNST cells

Lentiviral particles were generated using the lentiviral vector pLKO.1-puro to either express shZNF423 (TRCN0000018174, Sigma-Aldrich) or scramble shRNA (a gift from David Sabatini, Addgene plasmid #1864; http://n2t.net/addgene:1864; RRID:Addgene_1864), which were co-transfected with the lentiviral packaging plasmids psPAX2 (Addgene plasmid #12260; http://n2t.net/addgene:12260; RRID:Addgene_12260) and pMD2.G (Addgene plasmid #12259; http://n2t.net/addgene:12259; RRID:Addgene_12259), gifts from Didier Trono, into HEK293T cells using Lipofectamine 3000 (Invitrogen). 24 hours post-transfection, culture media was replaced with fresh media. 48 hours post-transfection, supernatants containing lentivirus particles were collected and filtered through a 0.45-μm filter and stored at −80°C until use. MPNST cell lines were transduced with viral supernatant containing 2μg/ml of polybrene (sc-255611, Santa Cruz) for 24 hours. After culturing for another 24 hours, cells were selected with 2μg/ml of puromycin (MIR5940, Fisher) in culture media to select for lentivirus-infected cells.

### Cell viability assay

Human MPNST cell lines were plated at a density of 1,000cells/well in a 96-well plate and transfected with siRNA against ZNF423 or control. After 72 hours, cell viability was determined using the CellTiter-Glo 2.0 assay (Promega) according to the manufacturer’s instructions. Briefly, CellTiter-Glo reagent was equilibrated to room temperature, and 20μL was added directly to each well. Plates were incubated at room temperature for 10 minutes before luminescence was measured using a plate reader (BioTek Synergy H1). Cell viability was normalized to control wells.

### Drug treatment

Doxorubicin (hydrochloride) (15007, Cayman Chemical) was dissolved in 100% DMSO and stored as a 10 μM stock at −20°C until use. PARP inhibitor olaparib (HY-10162, MedChemExpress) was dissolved in 100% DMSO and stored as a 10μM stock at −20°C until use. Ifosfamide (HY-17419, MedChemExpress) was dissolved in 100% DMSO and stored as 10μM stock at −20°C until use.

### Proliferation assay

Cell proliferation was measured using the Incucyte S3 Live-Cell Analysis System (Sartorius). Human MPNST cell lines were transfected with siRNA against ZNF423 or control and plated onto a 96-well plate at a density of 1,000cells/well in 100 μL of media. Plates were placed into the Incucyte system and maintained at 5% CO_2_ at 37°C. Brightfield phase-contrast images were captured every 4 hours over a period of 5 days. Confluency was quantified using the Incucyte integrated image analysis software. An analysis definition was created within the software, and threshold parameters were optimized to distinguish cells from the background, ensuring accurate segmentation. Data were extracted as confluency values over time and analyzed using GraphPad Prism to generate growth curves.

### EdU labeling

Human MPNST cell lines were seeded at a density of 2×10^5^ cells/well onto four 12mm glass coverslips (05-8697, Fisher) per well of a 6-well plate and transfected with siRNA. After 48 hours, EdU labeling was performed for 24 hours before cells were fixed in 3.7% paraformaldehyde for 15 minutes and then permeabilized with 0.5% Triton X-100 in PBS. EdU-labeled cells were detected using the Click-iT EdU Alexa Fluor 488 Imaging Kit (C10337, Thermo Fisher) according to the manufacturer’s instructions. Afterwards, cells were mounted onto microscope slides (12-544-2, Fisher) using ProLong Diamond Antifade Mountant with DAPI (P36966, Fisher). Images were acquired by ECHO microscope, and EdU positive cells were counted manually.

### PI staining and flow cytometry

Human MPNST cells were transfected with siRNA against ZNF423 or control and plated onto 6cm plates. Cells were harvested by trypsinization and resuspended in 200μL of PBS to achieve single cells. Cells were fixed in 300μL of ice-cold 100% ethanol, with gentle agitation, before being pelleted. Cell pellets were resuspended in 500μL FxCycle PI/RNase Staining solution (F10797, Thermo Fisher Scientific and incubated in the dark for 30 minutes. Stained cells were analyzed on an Attune NxT Flow Cytometer (Thermo Fisher Scientific), and data were analyzed with FlowJo 10 software.

### Western blotting

Cell lysates were collected, and protein concentrations were determined using Pierce Coomassie Protein Assay Kit (PI23200, Thermo Fisher Scientific). Isolated proteins were fractioned using 4-20% Mini-PROTEAN TGX Stain-free gels (4568094, Bio-Rad) and electro-transferred to nitrocellulose membranes (1704158, Bio-Rad). Immunoblots were performed using primary antibodies against ZNF423 (ABN410, Sigma-Aldrich), GAPDH (sc-365062, Santa Cruz Biotechnology), SUZ12 (3737S, Cell Signaling Technology), ATF3 (18665S, Cell Signaling Technology), ATF4 (11815S, Cell Signaling Technology), PARP (9542S, Cell Signaling Technology), and γH2A.x (9718S, Cell Signaling Technology). Following incubation with primary antibodies at 4°C overnight, blots were washed 3x with TBST, before being incubated with appropriate horseradish peroxidase (HRP) – conjugated secondary antibodies, either anti-rabbit (7074V, Cell Signaling Technology) or anti-mouse (7076V, Cell Signaling Technology). Signals were detected using enhanced chemiluminescence substrate (SuperSignal West Pico PLUS Chemiluminescent Substrate 34580 and/or SuperSignal West Femto Maximum Sensitivity Substrate 34095), and images were collected on a Bio-Rad ChemiDoc. Band intensities were quantified using ImageJ, and signals were normalized to the corresponding loading control.

### Immunofluorescence staining

Human MPNST cell lines were seeded at a density of 2×10^5^ cells/well onto three 12mm glass coverslips (05-8697, Fisher) per well of a 6-well plate and transfected with siRNA. After 72 hours, coverslips were placed into individual wells of a 24-well plate, and cells were fixed in 4% formaldehyde in PBS for 15 minutes at room temperature. After fixing, cells were washed in 0.05% Triton X-100 in PBS 3x for 5 minutes each, before being permeabilized and blocked in 5% BSA, 0.5% Triton X-100 and 0.05% SDS in water for 1 hour. Incubation with primary antibodies against γH2A.x (9718S, Cell Signaling Technology) was performed at 4°C overnight in a humidified incubation chamber. Following incubation with primary antibodies, coverslips were washed in 0.05% Triton X-100 in PBS 3x for 5 minutes each before being incubated with anti-rabbit Alexa Fluor 488 secondary antibodies for 1 hour in the dark at room temperature. Coverslips were washed in 0.05% Triton X-100 in PBS 3x for 5 minutes, and mounted onto microscope slides (12-544-2, Fisher) using ProLong Diamond Antifade Mountant with DAPI (P36966, Fisher). Images were acquired by ECHO microscope.

### Bulk RNA sequencing and corresponding analysis

Frozen cell pellets (*n=*3 per condition) of human MPNST cell lines treated with siZNF423 or siControl (ST88-14, NF90.8, RHT-172) were submitted to GENEWIZ (Azenta Life Sciences, South Plainfield, NJ, USA) for total RNA extraction, library preparation and bulk RNA sequencing. Sequencing and primary data processing were performed by the vendor using standard protocols. Briefly, total RNA was extracted using RNeasy Kit (Qiagen), and RNA quality and integrity were assessed using an Agilent Bioanalyzer. Sequencing libraries were generated using poly(A), and indexed, quantified and pooled prior to sequencing. Pooled libraries were sequenced on an Illumina platform using paired end (2 x 150bp) configuration. Reads were aligned to the human reference genome (GRCh38/hg38) using STAR aligner, and differential gene expression analysis was performed using DESeq2.

Frozen cell pellets (*n*=3 per condition) of the JH-2-002 cell line treated with siZNF423 or siControl were submitted to Novogene (Sacramento, CA, USA) for total RNA sequencing. Total RNA extraction, library preparation, sequencing, and primary data processing were performed by the vendor using standard protocols. Briefly, RNA quality was assessed prior to library construction, and libraries were prepared using poly(A) mRNA enrichment, followed by indexing, quantification, and pooling. Libraries were sequenced on an Illumina platform using paired-end (2 × 150 bp) sequencing. Raw reads were aligned to the human reference genome (GRCh38/hg38) using a standard RNAseq alignment pipeline, and differential expression analysis was conducted using DESeq2.

Total RNA from ST88-14 and NF90.8 NLS-YFP (*n*=3 per cell line) and HA-SUZ12 (*n*=3 per cell line) was extracted using RNeasy Kit with on-column DNase I treatment according to the manufacturer’s instructions. Total RNA was submitted to the Indiana University Center for Medical Genomics for bulk RNAseq. Briefly, RNA quality and integrity were assessed using Agilent Bioanalyzer. Libraries were generated and indexed libraries were quantified, and quality assessed by Qubit and Agilent TapeStation. Pooled libraries were sequenced on an Illumina NovaSeq 6000 using paired end (2 x 100bp) configuration. QC-passed reads were aligned to the human reference genome (GRCh38/hg38) and translated to transcriptome coordinates using Salmon (v1.4)^67^, and differential gene expression analysis was performed using DESeq2.

Subsets of transcription factors and kinases were interrogated by differential gene expression and pathway analysis. Differentially expressed genes with p-values≤0.05 (false discovery rate) and log_2_ fold changes of ≤-1 or ≥1 were queried using ShinyGO 0.80. Dot plots reflecting Hallmark Molecular Signatures Database (HMSigDB) enrichment of dysregulated genes were plotted using R.

### Single cell RNA sequencing and corresponding analysis

Tumor samples from NF1 patients were obtained for single nucleus sequencing from the IU Simon Comprehensive Cancer Center (IUSCCC) Biospecimen Repository and the Indiana Pediatric Biobank with informed consent under IRB approval (1501467439). Tumor samples were sent to Novogene for single nucleus capture, library preparation and sequencing. Briefly, single nuclei suspensions were processed using 10x Genomics Chromium Chips for 3′ library preparation per manufacturer’s instructions. Library quality was assessed using Qubit 2.0 (concentration), Agilent 2100 (insert size), and qPCR (effective concentration). Libraries were sequenced on an Illumina sequencing platform.

We assembled transcriptomes from *n*=71 human peripheral nerve sheath tumors (*n*=46 PNF; *n=*25 MPNST) by integrating profiled tumors from the IUSCCC Biobank, phs003835^26^, with five publicly available scRNAseq datasets: GSE212964^68^, GSE232766^69^, GSE207399^23^, GSE179043^21^ and GSE181985^70^. Reads were aligned to the GRCh38 human reference genome for tumors from the IUSCCC Biobank and public cohorts that had available raw FASTQ files. When raw FASTQs were not available, we used the authors filtered feature-barcode matrices as provided.

### scRNAseq Data Processing

366,397 total cells from 71 samples spanning 65,490 genes were analyzed using scanpy Python (v.3.11.5) package (v.1.11.4). Data processing and quality control were conducted following established single-cell RNAseq best-practice workflows implemented in Scanpy and described by Luecken and Theis^71^. Figures were generated using scanpy and MATLAB (via the MATLAB Engine API for Python).

### Quality Control

Quality control was performed using scanpy. Mitochondrial genes were identified by the “MT-” prefix. Cells with mitochondrial gene expression greater than 3 median absolute deviations (MADs) from the median or exceeding 8% of total counts were removed. Cells were additionally filtered if they exceeded 5 MADs for total counts, number of genes detected, or percent of counts in the top 20 expressed genes. Doublets were identified using scanpy’s implementation of Scrublet, with each sample specified as the batch key. A total of 2,604 predicted doublets were removed from downstream analysis.

### Normalization and data integration

Filtered samples were concatenated and normalized using total-count normalization followed by log transformation. The top 10,000 highly variable genes were identified, and fifty principal components were computed using scanpy’s experimental Pearson residuals recipe. To mitigate batch effects, scVI (v1.4.0) was applied with sample specified as the batch key and percent mitochondrial counts included as a covariate. The learned latent space was used for downstream analyses, and scVI-normalized expression values were used for visualization and clustering.

### Clustering and Annotation

Principal component analysis, nearest neighbor graph construction, and UMAP embedding were performed using default scanpy parameters. Leiden clustering was performed on the scVI latent space at multiple resolutions (0.1, 0.5, 1.0, 1.5, and 2.0), with a resolution of 1.0 selected for downstream analyses. The top 100 differentially expressed genes per cluster were identified using scanpy’s rank_genes_groups function. Initial cell type annotations were assigned based on canonical marker gene expression, followed by refinement using established markers for tumor microenvironment populations and cancer-associated fibroblasts (Table X). To distinguish malignant from non-malignant cells, inferCNVpy (v.0.6.1) was performed using annotated non-tumor immune cells as the reference population and the Gencode v38 gene annotation. Clusters were classified as malignant based on mean and median CNV scores, the proportion of cells within each cluster with CNV score ≥ 0.04, and visual inspection of the CNV dendrogram.

### Trajectory and Correlation Analysis

Clusters identified as malignant were subjected to trajectory analysis using Scanpy’s PAGA implementation with default parameters. A cycling score was calculated using the genes MKI67, TOP2A, UBE2C, BIRC5, and CENPF. The root cluster was defined as the cluster with the lowest average cycling score. Spearman partial correlation analysis was performed within malignant clusters to identify genes correlated with ZNF423 expression while controlling for cell cycle score. The cell cycle score was computed from canonical cell cycle genes and regressed out prior to correlation testing. P-values were adjusted using the Benjamini–Hochberg method to control the false discovery rate.

### Differential Analysis

Cells were pseudobulked by summing raw counts per sample within each cluster (minimum number of 10 cells per sample). Differential expression analysis between MPNST and PNF tumor types was performed using *edgeR* (v.4.8.0). Lowly expressed genes were removed, and libraries normalized using the trimmed mean of M-values (TMM) method. A quasi-likelihood negative binomal generalized linear model was fit: dispersion was estimated, the GLM was fit with glmQLFit() and differential expression between MPNST and PNF was tested with glmQLFTest(). logCPM values were computed and exported for plotting in R (v.4.5.0). Barplots were generated using *ggpubr*, with each point representing an individual biological sample. All code used can be found at: https://github.com/AngusLab/Cancer_scRNAseq.

### Statistical analysis

All statistical analyses were performed using GraphPad Prism v10.0 (GraphPad Software), R (v.4.5.0) or Python (v.3.11.5) depending on the dataset. Details of the statistical tests used for individual experiments are specified in the corresponding figure legends. Data are presented as mean ± standard error of the mean (SEM) unless otherwise stated. The number of independent biological replicates (*n*) is indicated in the figure legends. All statistical tests were two-tailed, and p values of ≤0.05 was considered statistically significant. Two group comparisons were performed using unpaired two-tailed Student’s t-tests. Multiple group comparisons were analyzed using one-way or two-way analysis of variance (ANOVA) followed by Tukey’s or Bonferroni’s post hoc correction for multiple comparisons where appropriate. Growth curves were analyzed using two-way repeated measures ANOVA. Dose response curves and IC_50_ values were generated using nonlinear regression (log[inhibitor] vs. normalized response-variable slope model).

## Data availability

Mouse scRNAseq data from embryonic and postnatal Schwann cells are publicly available through the Sciatic Nerve ATlas (SNAT) web portal (https://www.snat.ethz.ch) based on data generated by Gerber and colleagues^36^. Bulk RNAseq of human PNF and MPNST were obtained from Johns Hopkins 2024 cohort on cBioPortal^37,38^. Normal nerve, PNF and MPNST expression data are publicly available in Gene Expression Omnibus (GEO): GSE41747. Bulk RNAseq of *Nf1*^−/−^ GFP, *Nf1*^−/−^ Cre+, *Nf1*^−/−^;*Arf*^−/−^ Cre+ DNSCs are available in GEO: GSE232451. Bulk RNAseq data of *Nf1*^−/−^;*Arf*^−/−^ sgControl and sgSUZ12 have been deposited in GEO: GSE320227. Bulk RNAseq of ST88-14 and NF90.8 NLS-YFP and HA-SUZ12 have been deposited in GEO: GSE320094. Bulk RNAseq data of human MPNST cell lines with siRNA depletion have been deposited in GEO: GSE320122 (ST88-14), GSE320235 (NF90.8), GSE320095 (JH-2-002), and GSE320236 (RHT-172). Bulk RNAseq data of ST88-14 shRNA-depleted ZNF423 have been deposited in GEO: GSE320231. Human scRNAseq data are available under the accession code phs003835 in dbGaP or are publicly available in GEO: GSE212964, GSE232766, GSE207399, GSE179043 and GSE181985. All other data presented in this manuscript is available from the authors upon request.

## Supporting information

Supplemental Table 1. ZNF423 count by tumor type.

Supplemental Table 2. Marker genes.

Supplemental Table 3. Detailed cell type annotation.

Supplemental Table 4. CNV score summary.

Supplemental Table 5. Spearman correlation of ZNF423.

## ACKNOWLEDGEMENTS

The authors thank Katie Jackson and Heather Daniel for administrative support.

## AUTHOR CONTRIBUTIONS

Conceptualization: SKM, SDR, SPA

Data curation: SKM, CAB, EEW, CJB, EYZ

Formal analysis: SKM, CAB, CJB, EYZ, SDR, SPA

Funding acquisition: SPA

Investigation: SKM, CAB, CJB, SAHD, EYZ, CD, EMM, HM, MDC, BJH

Methodology: MRS, SG, KEP, MLF, CAP

Project administration: SPA

Resources: MRS, SG, CDC, AC, CAP, KEP, MLF

Software: SKM, CAB, SDR

Supervision: DKM, DWC, SDR, SPA

Validation: SKM, CAB, CJB, SAHD, EZ, CD, EMM, HM, MDC, EAG

Visualization: SKM, CAB

Writing – original draft: SKM, CAB, SPA

Writing – review and editing: SKM, CAB, DKM, SDR, DWC, SPA

