## Supplemental Figures for "ZNF423 depletion induces the integrated stress response and represents a potential vulnerability in NF1-associated MPNST"

**Supplemental Figure S1. SUZ12 restoration in MPNST cells does not alter ZNF423 protein levels despite transcriptional changes. (A)** Representative immunoblot showing ZNF423 and SUZ12 protein expression in human MPNST cell lines (ST88-14, NF90.8 and JH-2-002) following restoration of SUZ12 expression (HA-SUZ12) or control (NLS-YFP). GAPDH serves as loading control. **(B-C)** Volcano plots of whole transcriptome differentially expressed transcription factors in ST88-14 and NF90.8 MPNST cells comparing NLS-YFP ($n$=3) and HA-SUZ12 ($n$=3). Dashed vertical lines reflect $\log_2$FC of -1 and 1. Dashed horizontal line reflects false discovery rate (FDR) of 0.05. Downregulated genes with $\log_2 FC \leq -1$ and adjusted p-value $\leq 0.05$ are denoted in blue**.** Upregulated genes $\log_2 FC \geq 1$ and adjusted p-value $\leq 0.05$ are denoted in green. **(D)** Venn diagram showing overlap of significantly downregulated genes among ST88-14 and NF90.8 cell lines following SUZ12 restoration. Thresholds for DEGs were defined as $\log_2 FC \leq -1$ and adjusted p-value $\leq 0.05$.

**Supplemental Figure S2. ZNF423 depletion does not significantly alter cell-cycle distribution in ST88-14 cells. (A)** Representative flow cytometry histograms showing PI-stained ST88-14 cells showing DNA content distribution. G0/G1 (purple) and G2/M (green) populations were manually gated in FlowJo, and S phase (yellow) was calculated as the fraction of cells between these peaks. **(B)** Quantification of the mean percentage of cells in G0/G1, S, and G2/M phases following transfection with siControl (*n*=3) and siZNF423 (*n*=3). **(C)** Bar graph showing percentage of cells in each cell cycle phase. Error bars reflect SEM. P-values represent unpaired, two-tailed t test with Welch's correction. ns=not significant.

**Supplemental Figure S3. Transcriptomic changes following ZNF423 depletion in JH-2-002 cells. (A)** Volcano plot of whole-transcriptome differentially expressed genes (DEGs) identified by RNAseq in JH-2-002 cells following siRNA-mediated depletion of ZNF423 (*n*=3) compared to control (*n*=3). Dashed vertical lines reflect $log_2FC$ of -1 and 1. Dashed horizontal line reflects false discovery rate (FDR) of 0.05. Downregulated genes with $log_2FC \leq -1$ and $-log_{10}$ adjusted p-value above FDR threshold are denoted in blue. Upregulated genes $log_2FC \geq 1$ and $-log_{10}$adjusted p-value above FDR threshold are denoted in red**. (B-C)** Dot plots of top enriched Hallmark gene signatures of significantly downregulated and upregulated genes (FDR≤0.05). **(D-E)** Volcano plots of differentially expressed transcription factors and kinases in JH-2-002 cells following siRNA-mediated depletion of ZNF423 compared to control. **(F)** Immunoblots showing cleavage of PARP and ATF3 induction following siRNA-mediated depletion of ZNF423. GAPDH serves as loading control.

**Supplemental Figure S4. Transcriptomic changes following ZNF423 depletion in RHT-172 cells. (A)** Volcano plot of whole-transcriptome differentially expressed genes (DEGs) identified by RNAseq in RHT-172 cells following siRNA-mediated depletion of ZNF423 ($n$=3) compared to control ($n$=3). Dashed vertical lines reflect $log_2FC$ of -1 and 1. Dashed horizontal line reflects false discovery rate (FDR) of 0.05. Downregulated genes with $log_2FC \leq -1$ and $-log_{10}$ adjusted p-value above FDR threshold are denoted in blue. Upregulated genes $log_2FC \geq 1$ and $-log_{10}$adjusted p-value above FDR threshold are denoted in red. **(B-C)** Dot plots of top enriched Hallmark gene signatures of significantly downregulated and upregulated genes (FDR≤0.05). **(D-E)** Volcano plots of differentially expressed transcription factors and kinases in RHT-172 cells following siRNA-mediated depletion of ZNF423 compared to control.

**Supplemental Figure S5. Transcriptomic changes following stable shRNA-mediated ZNF423 knockdown in ST88-14 cells.** **(A)** Volcano plot of whole-transcriptome differentially expressed genes (DEGs) identified by RNAseq in ST88-14 cells following shRNA-mediated knockdown of ZNF423 (*n*=3) compared to scrambled control (*n*=3). Dashed vertical lines reflect $\log_2$FC of -1 and 1. Dashed horizontal line reflects false discovery rate (FDR) of 0.05. Downregulated genes with $\log_2 FC \leq -1$ and $-\log_{10}$ adjusted p-value above FDR threshold are denoted in blue. Upregulated genes $\log_2 FC \geq 1$ and $-\log_{10}$adjusted p-value above FDR threshold are denoted in red. **(B-C)** Dot plots of top enriched Hallmark gene signatures of significantly downregulated and upregulated genes (FDR$\leq$0.05). **(D-E)** Volcano plots of differentially expressed transcription factors and kinases in cells following shRNA knockdown of ZNF423.

**Supplemental Figure S6. Single-cell analyses of ZNF423 expression and malignant cell-state correlations. (A)** Boxplot of logCPM ZNF423 expression (y-axis) in PNF (orange) and MPNST (blue) tumor samples (x-axis). RNA counts were pseudobulked per sample across all cells. Each dot represents one pseudobulked biological sample. Differential expression between MPNST and PNF samples was performed using edgeR, and p-values were FDR-adjusted. **(B)** Dendrogram of inferred copy number variation (CNV) profiles across annotated cell types. Euclidean distances between annotation-level averaged chromosome CNV scores were computed, and hierarchical clustering was performed to generate the dendrogram. **(C)** Spearman correlation between log-normalized ZNF423 expression (y-axis) and PAGA-derived pseudotime (x-axis) across malignant cells. **(D)** Spearman correlation between cell cycle score (Scanpy score of S and G2M genes; y-axis) and pseudotime (x-axis) across malignant cells. **(E)** Scatter plot of log-normalized expression of the indicated gene (y-axis) versus PAGA-derived pseudotime (x-axis) across malignant cells. **(F)** Scatter plot and Spearman correlation between cell cycle score (x-axis) and log-normalized ZNF423 expression (y-axis) across malignant cells. **(G)** Scatter plot and partial Spearman correlation between log-normalized ZNF423 and SOX10 expression in malignant cells. Colors indicate Leiden resolution 1.00 clusters. Partial Spearman correlation was performed controlling for cell cycle score.

**Supplemental Figure S7. UMAP visualization of integrated single-cell RNAseq data highlighting canonical marker genes defining major cell populations.** Representative canonical markers used for annotation are displayed for each major lineage, including Schwann cells (SOX10, MPZ), malignant cells (PDGFRA, EGFR), CAFs (COL1A1, DCN), perivascular cells (ACTA2, MYH11, PDGFRB), endothelial cells (PECAM1, VWF, CDH5), T cells (TRAC, CD3D, CD3E), mixed immune cells (IL2RA, FOXP3, CTLA4), B cells (MS4A1, CD79A), TAMs (CD68, CSF1R, C1QA, MRC1), pDCs (CLEC4C) and mast cells (TPSAB1, KIT, CPA3). Marker-based annotation criteria are provided in Supplemental Table 2.

**Supplemental Figure S8. ZNF423 is co-expressed with developmental genes. (A-C)** Bar plots depicting Ebf1, Postn, and Ebf3 transcript expression (RPKM) at days of embryonic (E13.5, E17.5) and postnatal (P1, P5, P14, P24, and P60) development. These data were generated using the Sciatic Nerve ATlas (SNAT).

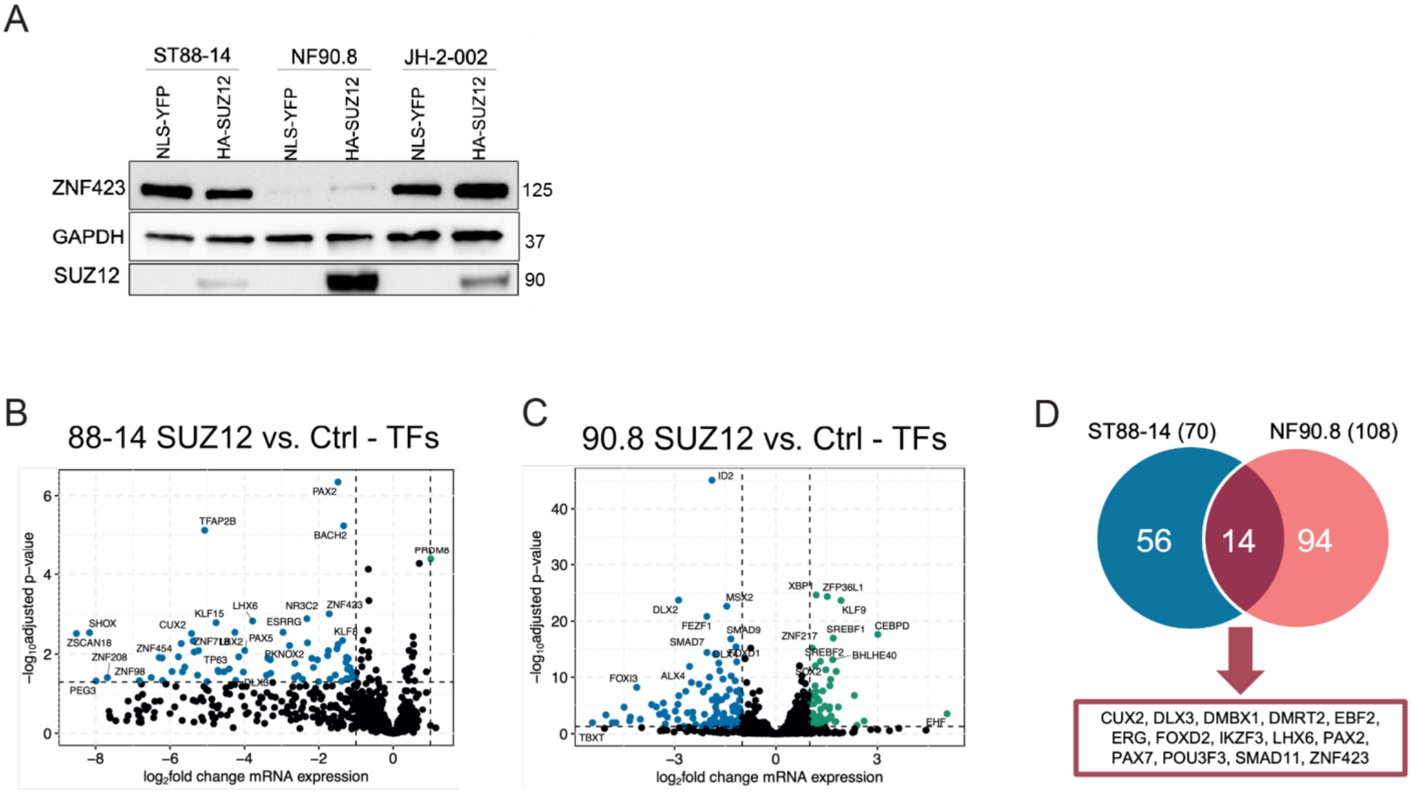


**Supplemental Figure S1**

A

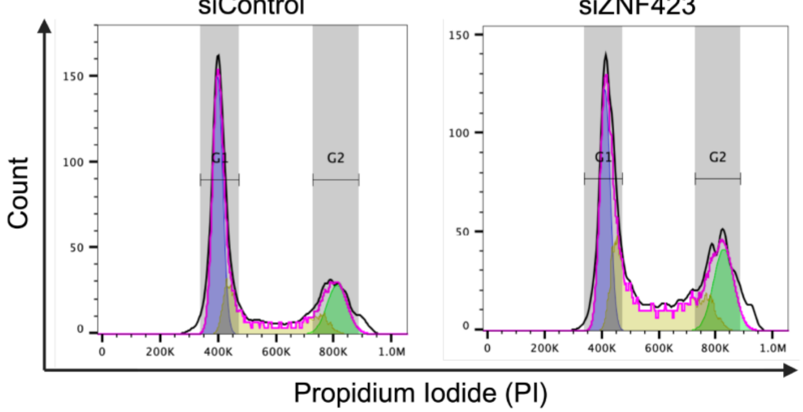


B

| | GO/G1 | S | G2/M |
|---|---|---|---|
| siControl | 39.7% | 33.6% | 14.96% |
| siZNF423 | 29.6% | 44.2% | 26.05% |

C

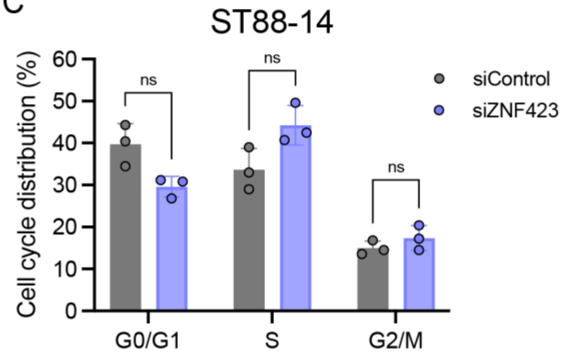


**Supplemental Figure S2**

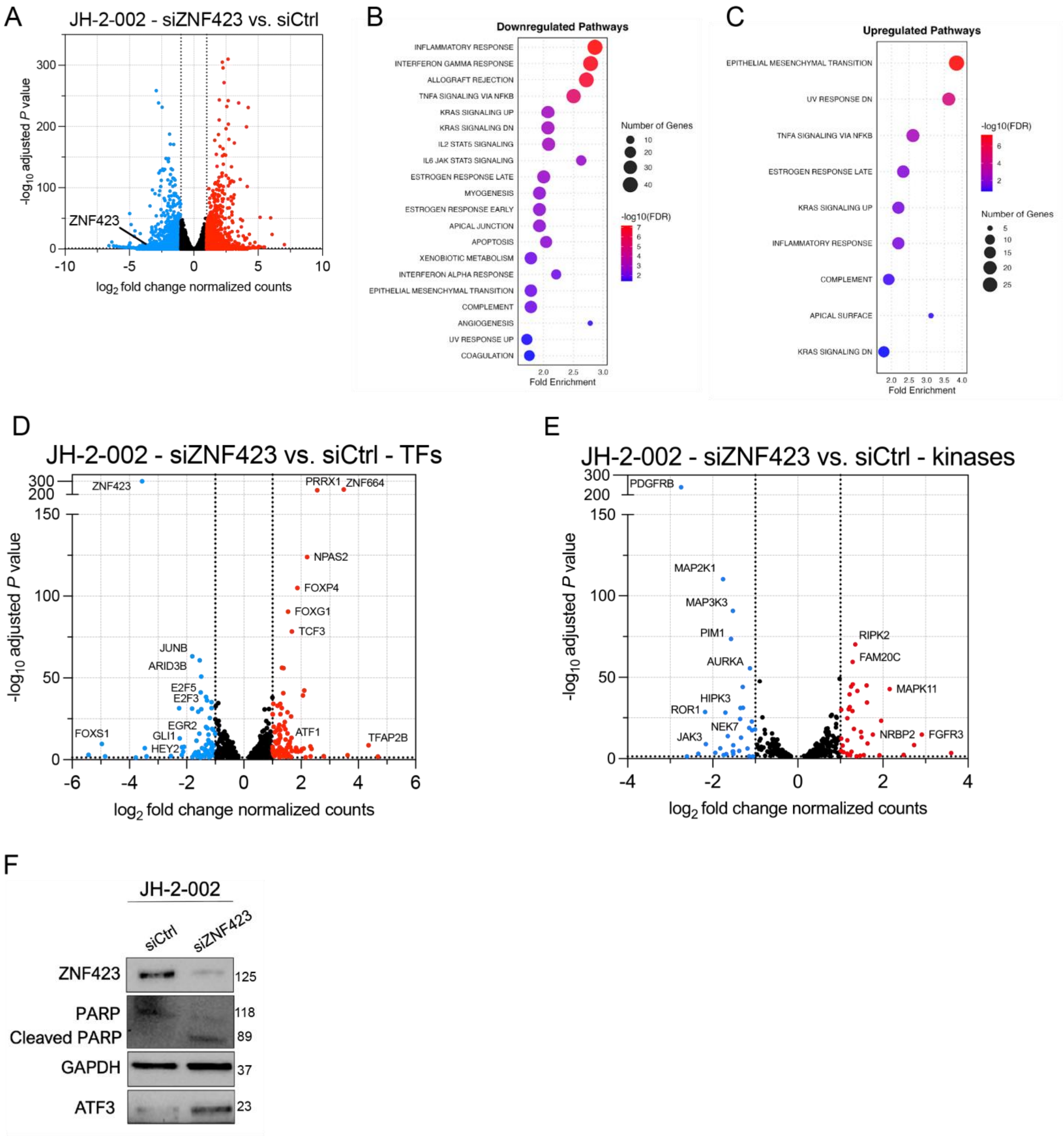


**Supplemental Figure S3**

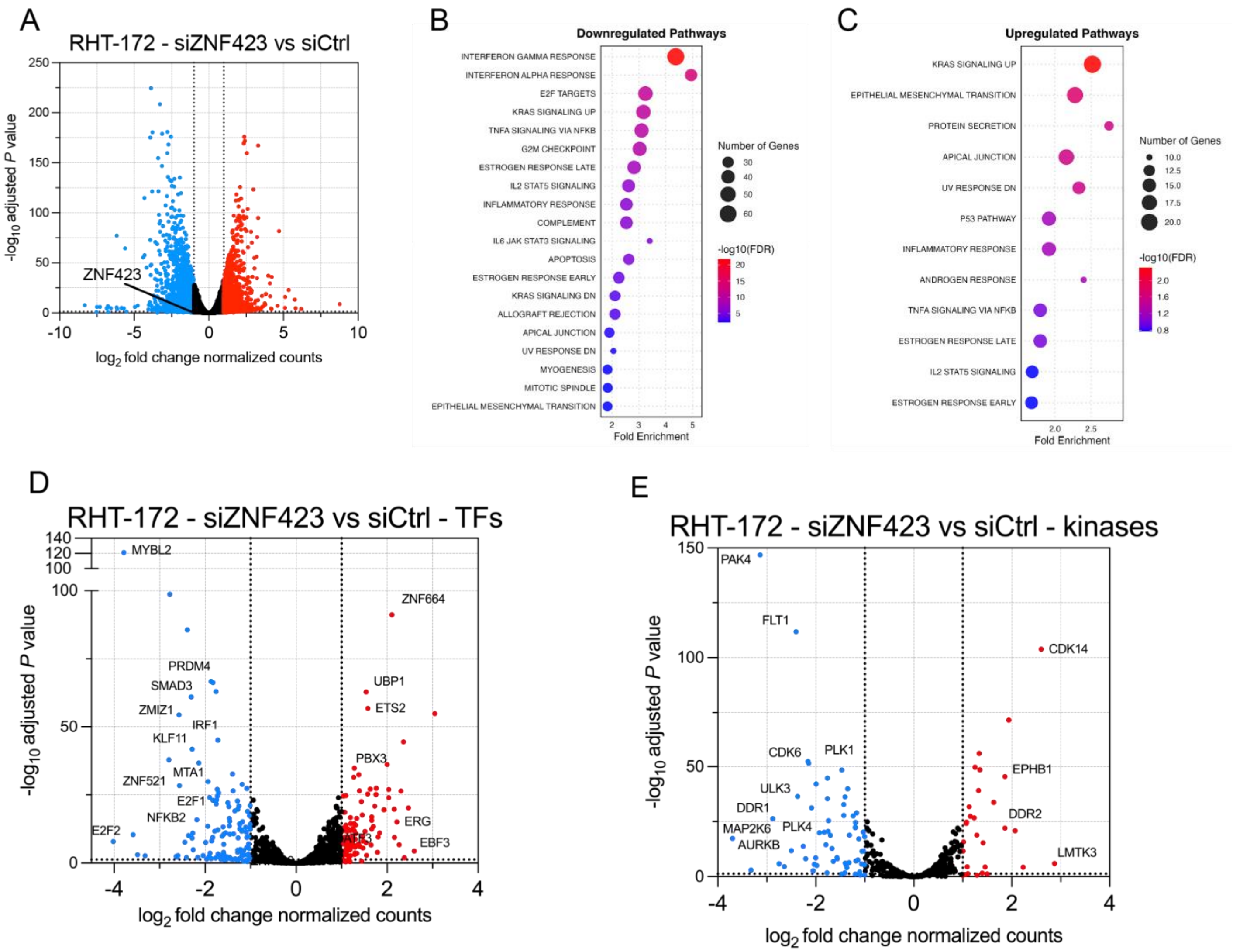


**Supplemental Figure S4**

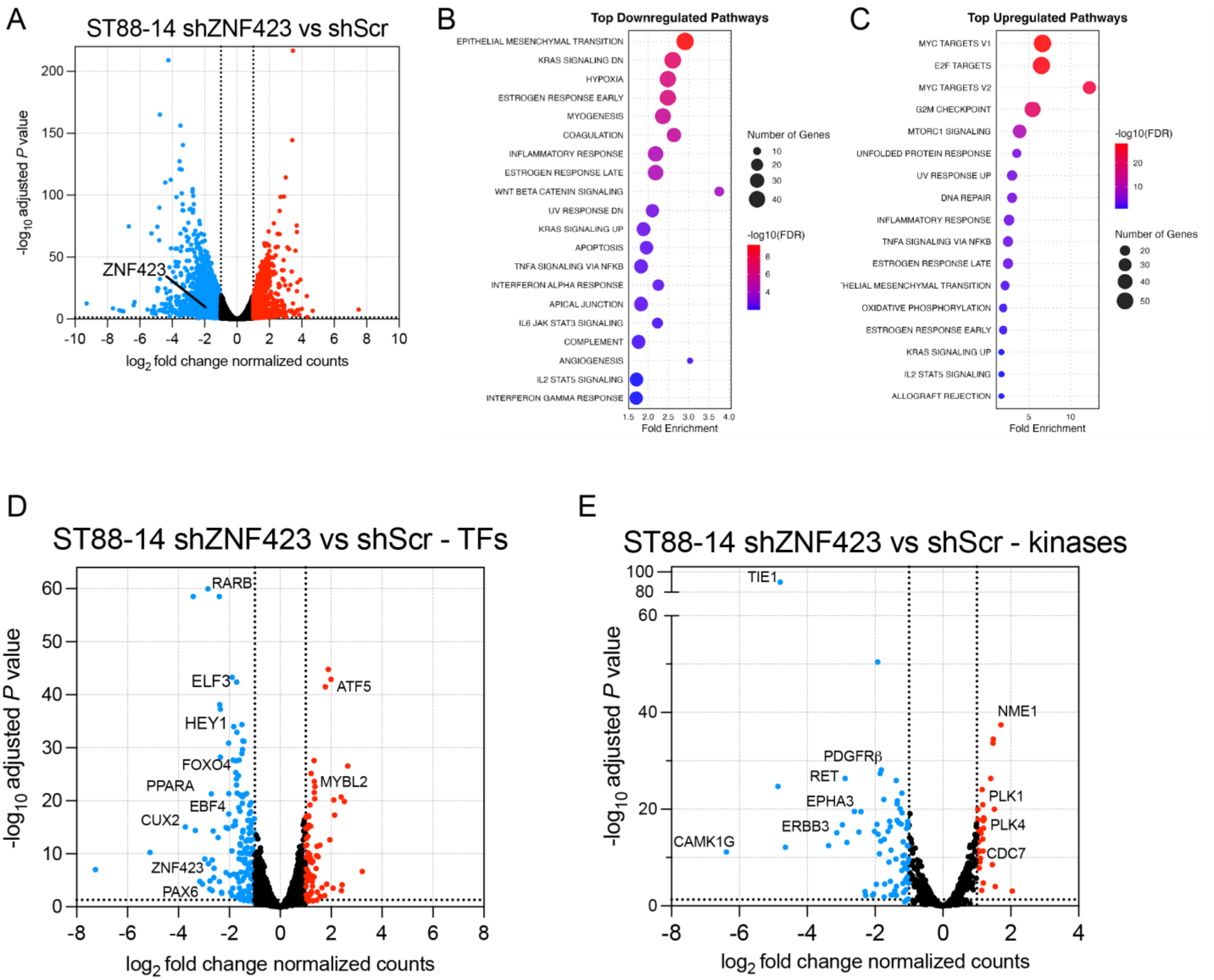


**Supplemental Figure S5**

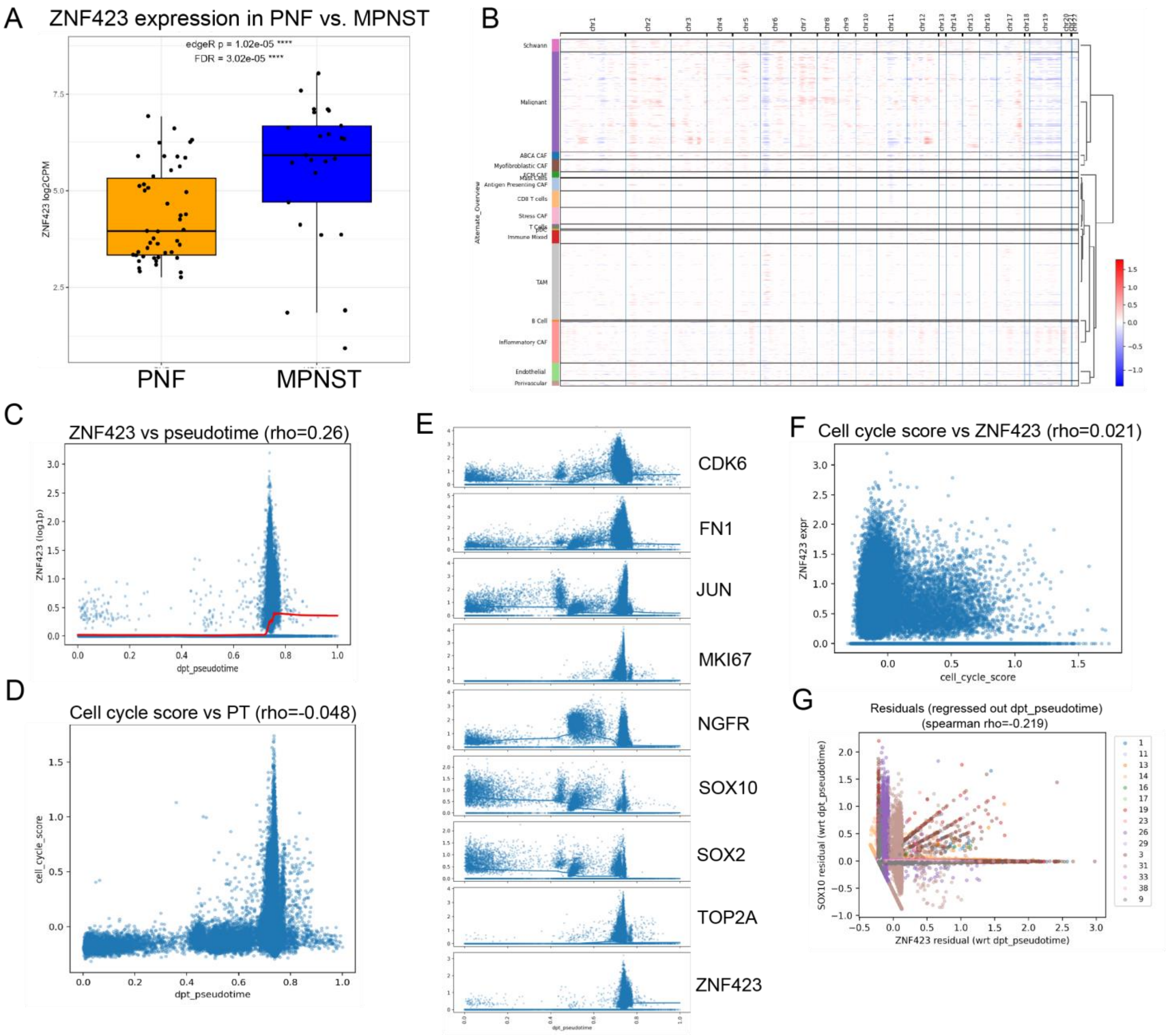


**Supplemental Figure S6**

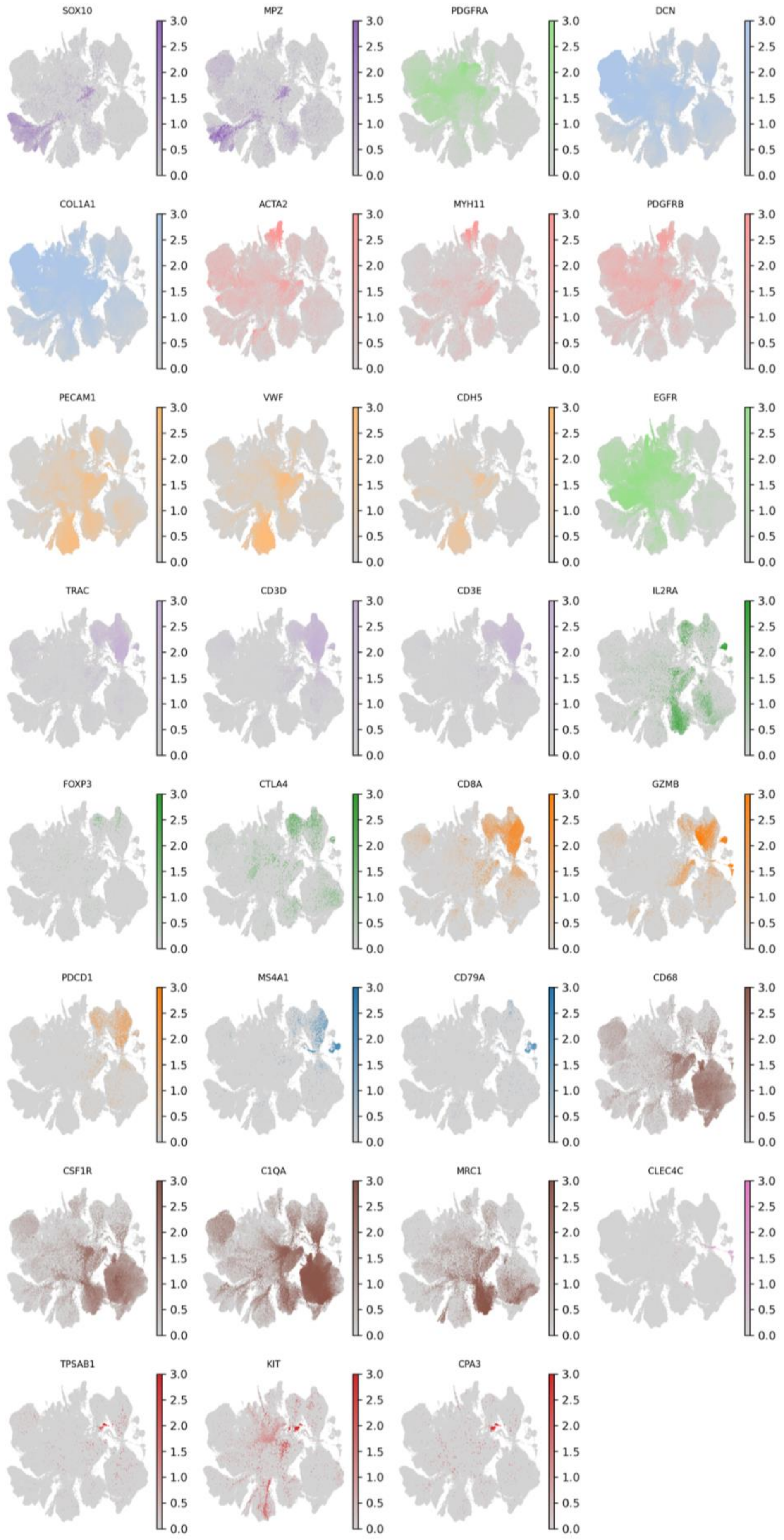


**Supplemental Figure S7**

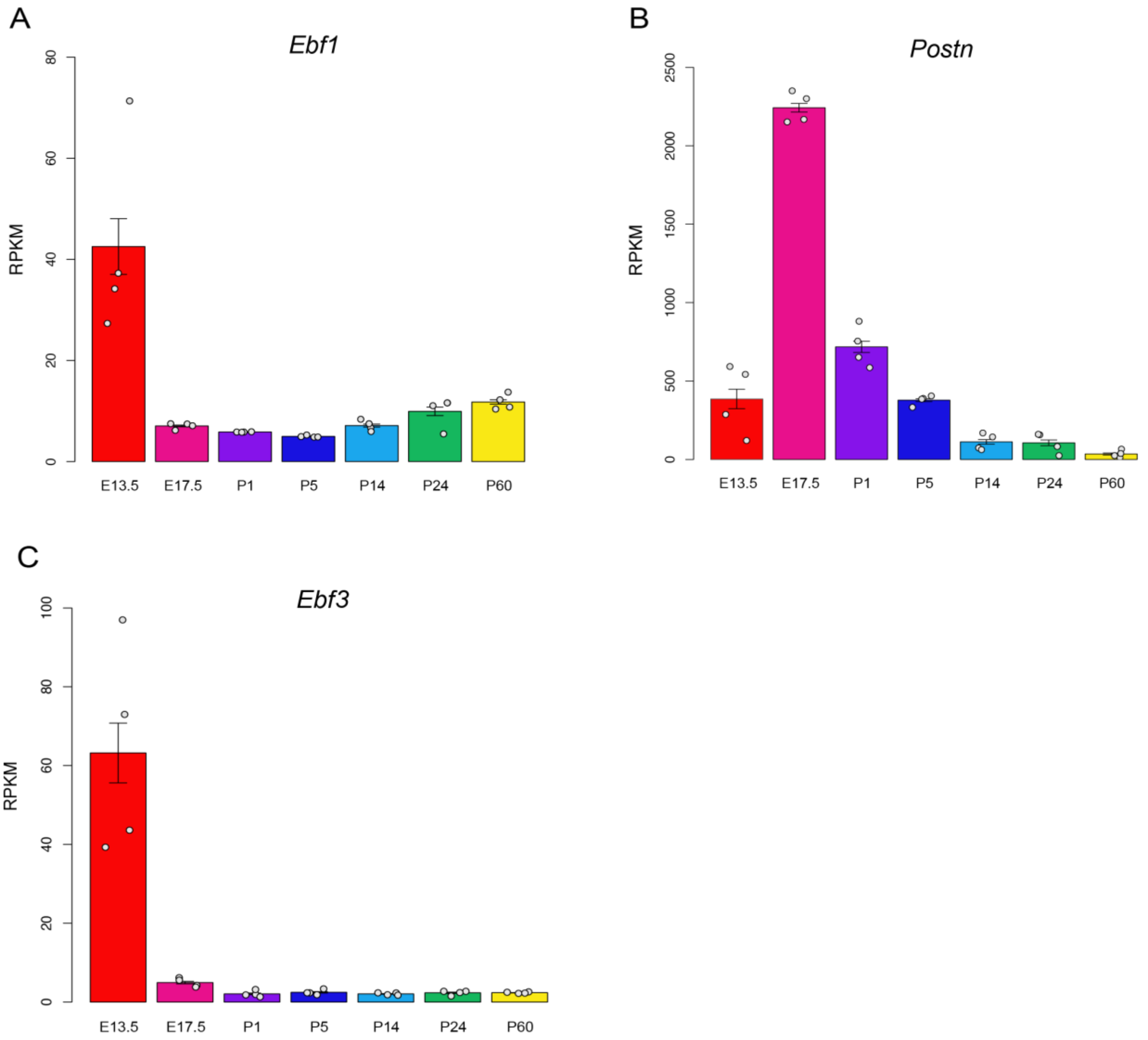


**Supplemental Figure S8**
