## Supplemental Table 1. ZNF423 count by tumor type. for "ZNF423 depletion induces the integrated stress response and represents a potential vulnerability in NF1-associated MPNST"

| ZNF423 | Sample | condition |
| --- | --- | --- |
| 6 912118609 | NF32 | PNF |
| 5 820969148 | S11 | MPNST |
| 5 789177742 | MPNST_2_023 | MPNST |
| 5 847405895 | NF13 | PNF |
| 6 334583118 | M2372 | MPNST |
| 4 708541182 | S6 | MPNST |
| 8 038955728 | S2 | MPNST |
| 3 860594414 | S16 | MPNST |
| 6 400672794 | M803 | MPNST |
| 7 010899194 | G1MPNST3 | MPNST |
| 1 917142329 | S15 | MPNST |
| 7 096782607 | S7 | MPNST |
| 7 092521586 | S13 | MPNST |
| 3 850715789 | M1677 | MPNST |
| 5 719352076 | M3048 | MPNST |
| 6 351972636 | S17 | MPNST |
| 7 592863891 | S10 | MPNST |
| 6 61274266 | G1MPNST2 | MPNST |
| 5 91149164 | G1MPNST1 | MPNST |
| 0 938108496 | M1933 | MPNST |
| 6 664646859 | S5 | MPNST |
| 7 053081135 | S4 | MPNST |
| 5 748278129 | MPNST_2_084 | MPNST |
| 6 305807086 | NF16 | PNF |
| 5 524670838 | PNF_2_084 | PNF |
| 6 597287151 | NF02 | PNF |
| 5 887997719 | S1 | PNF |
| 4 667594993 | PNF_2_004 | PNF |
| 5 072911716 | NF14 | PNF |
| 5 121892372 | PNF_2_006 | PNF |
| 6 254196299 | G2PNF2 | PNF |
| 4 965721215 | PNF_2_013 | PNF |
| 6 447741766 | S8 | MPNST |
| 5 626505933 | NF12 | PNF |
| 1 861342974 | S14 | MPNST |
| 5 453843255 | MPNST_2_015 | MPNST |
| 5 879060022 | G2PNF3 | PNF |
| 5 167078969 | PNF_2_090 | PNF |
| 3 651784362 | CS023066_S0 | PNF |
| 5 010737818 | NF09 | PNF |
| 2 993156429 | CS024097_T1_ | PNF |
| 3 181981238 | CS023066_S0 | PNF |
| 3 082346753 | CS024097_T1_ | PNF |

| | | | |
|---|---|---|---|
| 5 | 37238901 | G2PNF1 | PNF |
| 2 | 913071236 | CS024097_T1_ | PNF |
| 3 | 177764391 | CS024097_T1_ | PNF |
| 4 | 114747756 | MPNST_2_016 | MPNST |
| 6 | 234856697 | PNF_2_021 | PNF |
| 3 | 958254467 | PNF_2_019 | PNF |
| 3 | 981602611 | PNF_2_017 | PNF |
| 3 | 344349631 | NF110 | PNF |
| 3 | 946988966 | PNF_2_026 | PNF |
| 4 | 25438289 | PNF_2_003 | PNF |
| 5 | 889002448 | NF15 | PNF |
| 3 | 387801797 | CS022514_S0[illegible] | PNF |
| 3 | 297383315 | CS022514_S0[illegible] | PNF |
| 3 | 415536607 | CS022514_S0[illegible] | PNF |
| 3 | 263205341 | CS022514_S0[illegible] | PNF |
| 4 | 382421587 | CS023066_S0[illegible] | PNF |
| 3 | 760320881 | CS023066_S0[illegible] | PNF |
| 3 | 409786323 | CS022514_S0[illegible] | PNF |
| 3 | 329068254 | CS024097_T2_ | PNF |
| 3 | 256256271 | CS022514_S0[illegible] | PNF |
| 3 | 516151756 | CS022514_S0[illegible] | PNF |
| 2 | 761645329 | CS024097_T2_ | PNF |
| 3 | 276957435 | CS022514_S0[illegible] | PNF |
| 4 | 354349276 | CS023066_S0[illegible] | PNF |
| 2 | 888917258 | CS024097_T2_ | PNF |
| 3 | 598032542 | CS023066_S0[illegible] | PNF |
| 3 | 623596593 | CS023066_S0[illegible] | PNF |
| 3 | 697735175 | CS023066_S0[illegible] | PNF |
