## Supplemental Table 2. Marker genes. for "ZNF423 depletion induces the integrated stress response and represents a potential vulnerability in NF1-associated MPNST"

| |
|---|
| Cell type |
| Schwann |
| Generic Fibroblast |
| Smooth Muscle Cells (F |
| Endothelial |
| Adipocyte |
| Striated Myocyte |
| Tumor |
| Proliferation |
| Immune_pan |
| Core T cell |
| Exhausted Cytotoxic CD |
| NK |
| B Cells |
| CD4 Treg T cells |
| Plasma Cells |
| Myeloid |
| M2 -like Macrophage (T |
| M1-like inflammatory M |
| MCHII APC DC |
| cDC1 |
| cDC2 |
| pDC |
| Mast Cell |
| Platelet |
| myCAF |
| iCAF |
| apCAF |
| ECM-CAF |
| invCAF |
| pdgfraCAF |
| ABCA-CAF |
| sCAF |
| periCAF |
| prolifCAF |
| angioCAF |
| s_genes |
| g2m genes |

Marker Genes

["SOX10","S100B","ERBB3","NGFR","GFRA1","PLP1","MPZ","PMP22","MBP","EGR2","L1CAM"]

["PDGFRA","FAP","THY1","DCN","LUM","COL1A1","COL1A2","COL3A1","SPARC","FN1","POSTN","MMP2"

["RGS5","CSPG4","MCAM","NOTCH3","ACTA2","TAGLN","MYL9","CNN1","MYH11","PDGFRB","DES"]

["PECAM1","VWF","CDH5","KDR","FLT1","TEK","PLVAP","RAMP2"]

["ADIPOQ","FABP4","PLIN1","PPARG","LPL","CIDEC","LIPE"]

["CKM","TNNT1","TPM2","MYH1","MYH2","MYH7"]

["EGFR","PDGFRA","MET","MYC","MDM2","TERT"]

["MKI67","TOP2A","PCNA","TYMS","MCM5"]

["PTPRC"]

["CD4","IL2RA","FOXP3","CTLA4","IKZF2","TIGIT"]

["CD8A","NKG7","PRF1","GZMB","PDCD1","TOX","LAG3"]

["NKG7","GNLY","KLRD1","FCGR3A","PRF1","GZMB","TRDC"]

["MS4A1","CD79A","CD79B","CD74","HLA-DRA","BANK1"]

["CD4","IL2RA","FOXP3","CTLA4","IKZF2","TIGIT"]

["MZB1","XBP1","JCHAIN","IGHG1","SDC1"]

["LYZ","LST1","TYROBP","FCER1G","CTSS","S100A8","S100A9"]

["CD68","CSF1R","MERTK","C1QA","C1QB","APOE","CD163","MRC1","TREM2"]

["IL1B","TNF","CXCL9","CXCL10","CCL2","NFKBIA"]

["HLA-DRA","HLA-DRB1","CD74","CIITA","ITGAX","FLT3"]

["CLEC9A","XCR1","BATF3","IRF8","CADM1"]

["CD1C","FCER1A","CLEC10A","IRF4","SIRPA"]

["CLEC4C","IL3RA","LILRA4","TCF4","GZMB"]

["TPSAB1","TPSB2","KIT","MS4A2","CPA3"]

["PPBP","PF4","ITGA2B"]

["ACTA2","TAGLN","MYL9","CNN1","MYH11","CALD1","PDGFRB","ITGB1","TGFBI"]

["IL6,"CXCL12","CCL2","CXCL1","LIF","PTGS2","IL1R1","VCAM1","STAT3"]

["HLA-DRA","HLA-DRB1","HLA-DPA1","CD74","CIITA","HLA-DQA1","HLA-DPB1"]

["LAMA2","LAMB1","COL1A1","COL1A2","COL3A1","COL4A1","COL4A2","COL5A2","COL6A1","COL6A3",

["COL11A1","ADAM12","ADAMTS9","ADAMTSL3","LOXL2","TNC","THSD4","POSTN"]

["PDGFRA","COL1A1","LAMA2","PDGFC","IGF1","FAP"]

["ABCA6","ABCA8","ABCA9","ABCA10","IGFBP7","SLC27A2"]

["HSPA5","HSP90AA1","CALR","MT2A","SPP1","TIMP1"]

["MCAM","RGS5","CSPG4","PDGFRB","ACTA2","ANPEP"]

["MKI67","TOP2A"]

["VEGFA","PDGFC","ANGPTL4","CXCL12","HGF"]

"MCM5,"PCNA","TYMS","FEN1","MCM2","MCM4","RRM1","UNG","GINS2","MCM6","CDCA7","DTL","PRIM

"HMGB2,"CDK1","NUSAP1","UBE2C","BIRC5","TPX2","TOP2A","NDC80","CKS2","NUF2","CKS1B","MKI67

1","UHRF1","HELLS","RFC2","RPA2","NASP","RAD51AP1","GMNN","WDR76"
","TMPO","CENPF","TACC3","FAM64A","SMC4","CCNB2","CENPE","CTCF"
