## Supplemental Table 3. Detailed cell type annotation. for "ZNF423 depletion induces the integrated stress response and represents a potential vulnerability in NF1-associated MPNST"

| Cluster | Detailed Annotation |
| --- | --- |
| 0 | Invasive ECM-CAF (quiescent/mature) |
| 1 | Malignant |
| 2 | Stress-responsive CAF |
| 3 | Malignant |
| 4 | Myofibroblastic ECM-CAF (myCAF) |
| 5 | Antigen-presenting CAF (apCAF) |
| 6 | Mixed T cells (exhausted CD8 + Tregs) |
| 7 | Immunosuppressive TAMs (M2-like) |
| 8 | Activated immune-reactive endothelial cells |
| 9 | Malignant |
| 10 | Immune-reactive ABCA-CAF |
| 11 | Malignant |
| 12 | Pericytes / vascular SMCs |
| 13 | Malignant |
| 14 | Malignant |
| 15 | Plasmacytoid dendritic cells (pDCs) |
| 16 | Malignant |
| 17 | Malignant |
| 18 | Myelinating Schwann cells |
| 19 | Malignant |
| 20 | Hybrid myofibroblastic invasive CAF |
| 21 | Inflammatory myofibroblastic CAF (cytokine-dominant) |
| 22 | Inflammatory myCAF (ECM/metabolic-dominant) |
| 23 | Malignant |
| 24 | Highly differentiated myelinating Schwann cells |
| 25 | Inflammatory monocyte-derived TAMs |
| 26 | Malignant |
| 27 | Mast cells |
| 28 | Inflammatory MHC-II+ TAMs |
| 29 | Malignant |
| 30 | Quiescent metabolic ABCA-CAF (baseline) |
| 31 | Malignant |
| 32 | CD8 cytotoxic T cells (partially exhausted) |
| 33 | Malignant |
| 34 | Metabolic FABP4+ endothelial cells |
| 35 | Inflammatory APC-like TAMs (LAMP3+) |
| 36 | Deeply exhausted CD8 T cells |
| 37 | Antigen-presenting DC-like TAMs |
| 38 | Malignant |
| 39 | Classical inflammatory TAMs |
| 40 | Mixed immune cluster (APC–T interface) |
| 41 | Inflammatory APC-like TAMs |
| 42 | Plasmablast-like B cells |

43 Mixed cytotoxic lymphocytes (low-signal)

Key_Features
COL1A1 COL3A1 FN1 POSTN PDGFRA
ECM genes MKI67 TOP2A
UPR mitochondrial stress SPP1
DCN LUM COL1A1 FN1 COL11A1
ACTA2 TAGLN MYH11 PDGFRB
HLA-DRA HLA-DRB1 CD74 CIITA PDGFRA FAP THY1 DCN LUM COL1A1 FN1 SPARC HS[illegible]
CD8A PDCD1 TOX FOXP3
TREM2 APOE CD163 MRC1
PECAM1 VWF MHC-II
High inferCNV
ABCA6-10 PDGFRA MHC-II
COL11A1 POSTN ADAM12 CNV
CSPG4 NOTCH3 PDGFRB ACTA2
ECM LOXL2 EGFR MET
PPARG FABP4 EGFR
CLEC4C LILRA4 TCF4
COL11A1 ADAM12 LOXL2 POSTN ACTA2 HSPA5 MT-CO1 high_CNV
ABCA genes ECM CNV
SOX10 MPZ PMP22 MBP
ABCA genes RTK MYC
ACTA2 COL11A1 ABCA
IL6 CXCL12 CCL2
ECM stress ABCA
COL11A1 ABCA LOXL2
Myelin lipid metabolism
IL1B TNF S100A8/9
SOX10 weak myelin CNV
TPSAB1 KIT CPA3
IL1B MHC-II
ER stress ECM survival
PPARG FABP4 ABCA
SOX10 CAF mimicry ABCA
CD8A NKG7 PRF1
Proteostasis contractility ABCA
PECAM1 FABP4 PPARG
LAMP3 MHC-II IL1B
TOX PDCD1 LAG3
C1QA DC-like genes
ECM COL11A1 ABCA remnants
IL1B S100A8/9
Myeloid + T cell mixing
CD68 CSF1R HLA-DRA HLA-DRB1 IL1B TNF antigen_presentation
MS4A1 CD79A MZB1 XBP1

NKG7 GZMB weak T/NK

| | |
|---|---|
| Annotation_coarse | Overview |
| ECM CAF | CAF |
| Malignant | Malignant |
| Stress CAF | CAF |
| Malignant | Malignant |
| Myofibroblastic CAF | CAF |
| Antigen Presenting CAF | CAF |
| Immune Mixed | Immune Mixed |
| TAM | TAM |
| Endothelial Metabolic | Endothelial |
| Malignant | Malignant |
| ABCA CAF | CAF |
| Malignant | Malignant |
| Perivascular | Perivascular |
| Malignant | Malignant |
| Malignant | Malignant |
| pDC | pDC |
| Malignant | Malignant |
| Malignant | Malignant |
| Schwann | Schwann |
| Malignant | Malignant |
| Myofibroblastic CAF | Myofibroblastic CAF |
| Inflammatory CAF | Inflammatory CAF |
| Inflammatory CAF | Inflammatory CAF |
| Malignant | Malignant |
| Schwann | Schwann |
| TAM | TAM |
| Malignant | Malignant |
| Mast Cells | Mast Cells |
| TAM | TAM |
| Malignant | Malignant |
| ABCA CAF | CAF |
| Malignant | Malignant |
| CD8 T cells | CD8 T cells |
| Malignant | Malignant |
| Endothelial Metabolic | Endothelial |
| APC TAM | TAM |
| T Cells | T Cells |
| APC TAM | TAM |
| Malignant | Malignant |
| TAM | TAM |
| Immune Mixed | Immune Mixed |
| APC TAM | TAM |
| B Cell | B Cell |

Immune Mixed

Immune Mixed
