## Supplemental Table 4. CNV score summary. for "ZNF423 depletion induces the integrated stress response and represents a potential vulnerability in NF1-associated MPNST"

| normal | borderline | tumor | cluster | mean | median |
|---|---|---|---|---|---|
| 0 867633446 | 0 072199938 | 0 060166615 | 0 | 0 010453176 | 0 006910771 |
| 0 218361582 | 0 566525424 | 0 215112994 | 1 | 0 031818585 | 0 034859831 |
| 0 833862492 | 0 072531202 | 0 093606306 | 2 | 0 007013731 | 1 55E-07 |
| 0 224585214 | 0 587584012 | 0 187830774 | 3 | 0 030449058 | 0 034859831 |
| 0 578477444 | 0 389849624 | 0 031672932 | 4 | 0 01819331 | 0 009319422 |
| 0 912805688 | 0 067583405 | 0 019610907 | 5 | 0 005146115 | 0 000571498 |
| 0 939010357 | 0 053771315 | 0 007218328 | 6 | 0 005623885 | 0 002103924 |
| 0 91476387 | 0 071847776 | 0 013388354 | 7 | 0 010887611 | 0 009319422 |
| 0 77481882 | 0 104221788 | 0 120959393 | 8 | 0 015979443 | 0 009319422 |
| 0 04940507 | 0 062726332 | 0 887868598 | 9 | 0 064106515 | 0 04763698 |
| 0 700369738 | 0 239192264 | 0 060437998 | 10 | 0 012835879 | 0 000456638 |
| 0 24768 | 0 295573333 | 0 456746667 | 11 | 0 039355635 | 0 033940173 |
| 0 762036525 | 0 154399557 | 0 083563918 | 12 | 0 015597144 | 0 009319422 |
| 0 247855918 | 0 584476844 | 0 167667238 | 13 | 0 025758749 | 0 025046083 |
| 0 23245614 | 0 75877193 | 0 00877193 | 14 | 0 027828534 | 0 03021837 |
| 0 933510638 | 0 042553191 | 0 02393617 | 15 | 0 008091072 | 0 002601528 |
| 0 03915276 | 0 022464698 | 0 938382542 | 16 | 0 080230026 | 0 084896676 |
| 0 272038293 | 0 232947746 | 0 495013961 | 17 | 0 046011804 | 0 033940173 |
| 0 563719727 | 0 412062074 | 0 024218199 | 18 | 0 020781402 | 0 009319422 |
| 0 23261766 | 0 753637613 | 0 013744727 | 19 | 0 025010798 | 0 03021837 |
| 0 615819209 | 0 363320296 | 0 020860495 | 20 | 0 017842592 | 0 01749079 |
| 0 949503846 | 0 03813634 | 0 012359814 | 21 | 0 014159549 | 0 015318317 |
| 0 605823315 | 0 144700562 | 0 249476122 | 22 | 0 023368256 | 0 009319422 |
| 0 13545638 | 0 654406818 | 0 210136802 | 23 | 0 033198091 | 0 033940173 |
| 0 56375049 | 0 202236171 | 0 234013339 | 24 | 0 024542928 | 0 015318317 |
| 0 973135643 | 0 018349971 | 0 008514386 | 25 | 0 013793972 | 0 01537146 |
| 0 246013289 | 0 267275748 | 0 486710963 | 26 | 0 031328645 | 0 039215805 |
| 0 883561644 | 0 102739726 | 0 01369863 | 27 | 0 007327746 | 0 002103924 |
| 0 993777368 | 0 005070293 | 0 001152339 | 28 | 0 005662555 | 0 005250986 |
| 0 390249647 | 0 587376354 | 0 022373999 | 29 | 0 01710789 | 0 024756152 |
| 0 725559482 | 0 243816254 | 0 030624264 | 30 | 0 014593009 | 0 009319422 |
| 0 078114478 | 0 906060606 | 0 015824916 | 31 | 0 036511758 | 0 039215805 |
| 0 991141219 | 0 006832262 | 0 002026518 | 32 | 0 003157125 | 0 002260983 |
| 0 390804598 | 0 53991923 | 0 069276173 | 33 | 0 023146872 | 0 03021837 |
| 0 639109698 | 0 285373609 | 0 075516693 | 34 | 0 017285218 | 0 009319422 |
| 0 992541467 | 0 006011355 | 0 001447178 | 35 | 0 006110003 | 0 005250986 |
| 0 945023903 | 0 037375054 | 0 017601043 | 36 | 0 007266452 | 0 002260983 |
| 0 988926112 | 0 009259259 | 0 001814629 | 37 | 0 009500422 | 0 015318317 |
| 0 093959732 | 0 82885906 | 0 077181208 | 38 | 0 032030993 | 0 034859831 |
| 0 996275605 | 0 002793296 | 0 000931099 | 39 | 0 001374514 | 0 000456638 |
| 0 993661769 | 0 005762028 | 0 000576203 | 40 | 0 011563632 | 0 01537146 |
| 0 993192404 | 0 004657829 | 0 002149767 | 41 | 0 005007579 | 0 002260983 |
| 0 942895989 | 0 033310673 | 0 023793338 | 42 | 0 012568992 | 0 01537146 |

| | | | | | |
|---|---|---|---|---|---|
| 1 | 43 | 0 | 002366883 | 0 | 00153315 |
