## Supplemental Table 5. Spearman correlation of ZNF423. for "ZNF423 depletion induces the integrated stress response and represents a potential vulnerability in NF1-associated MPNST"

| gene | detect_frac | rho | p | partial_rho | partial_p | fdr |
|---|---|---|---|---|---|---|
| ZNF423 | 0 286516533 | 1 | | 1 | | |
| CDH11 | 0 657440369 | 0 27290395 | 0 | 0 136999645 | 0 | 0 |
| KCNQ1OT1 | 0 633863814 | 0 264293982 | 0 | 0 289206964 | 0 | 0 |
| MIR99AHG | 0 738402086 | 0 263997843 | 0 | 0 166808246 | 0 | 0 |
| EYA4 | 0 360463345 | 0 260605776 | 0 | 0 41605795 | 0 | 0 |
| BMP5 | 0 268708001 | 0 2597314 | 0 | 0 454848301 | 0 | 0 |
| DNM3OS | 0 410805048 | 0 25895336 | 0 | 0 370426335 | 0 | 0 |
| CACNA1C | 0 564228741 | 0 258381262 | 0 | 0 331206336 | 0 | 0 |
| GAS2 | 0 306971122 | 0 257553949 | 0 | 0 439062587 | 0 | 0 |
| SETBP1 | 0 589870743 | 0 25562384 | 0 | 0 102178792 | 8 81E-242 | 0 |
| LDLRAD4 | 0 491281339 | 0 253877577 | 0 | 0 343153994 | 0 | 0 |
| TSHZ2 | 0 799565971 | 0 249535836 | 0 | 0 157374171 | 0 | 0 |
| AUTS2 | 0 930926501 | 0 247125848 | 0 | 0 205349571 | 0 | 0 |
| HIF3A | 0 397727056 | 0 244252515 | 0 | 0 369430897 | 0 | 0 |
| RORA | 0 86811597 | 0 236687463 | 0 | 0 175997109 | 0 | 0 |
| SEPTIN6 | 0 305476766 | 0 234270961 | 0 | 0 436473952 | 0 | 0 |
| CACHD1 | 0 273114923 | 0 23377623 | 0 | 0 42113025 | 0 | 0 |
| BOC | 0 464373418 | 0 233570874 | 0 | -0 00177601 | 0 564847972 | 0 |
| COL25A1 | 0 28164322 | 0 233257216 | 0 | 0 437899525 | 0 | 0 |
| ARHGAP28 | 0 233462146 | 0 225568015 | 0 | 0 450109106 | 0 | 0 |
| NFIB | 0 724058175 | 0 222879298 | 0 | 0 193617991 | 0 | 0 |
| FGFR1 | 0 609287849 | 0 221388508 | 0 | 0 2578664 | 0 | 0 |
| PBX1 | 0 548580838 | 0 220757466 | 0 | 0 049276588 | 1 71E-57 | 0 |
| ZFHX4 | 0 628638328 | 0 219305698 | 0 | 0 200989354 | 0 | 0 |
| EBF3 | 0 399754431 | 0 218369662 | 0 | 0 359685163 | 0 | 0 |
| EBF1 | 0 748034494 | 0 217771606 | 0 | 0 13606983 | 0 | 0 |
| TENM1 | 0 304363138 | 0 217688947 | 0 | 0 419853503 | 0 | 0 |
| EYA1 | 0 381841199 | 0 21613283 | 0 | 0 293109308 | 0 | 0 |
| PLXDC2 | 0 731596581 | 0 2161303 | 0 | 0 137074921 | 0 | 0 |
| PTGIS | 0 236898212 | 0 215750144 | 0 | 0 435237641 | 0 | 0 |
| NFIA | 0 805153148 | 0 21461413 | 0 | 0 19636784 | 0 | 0 |
| ADAMTS9 | 0 370352744 | 0 214497762 | 0 | 0 376684985 | 0 | 0 |
| PREX2 | 0 338285964 | 0 214380005 | 0 | 0 392575653 | 0 | 0 |
| COL11A1 | 0 42388304 | 0 214307742 | 0 | 0 354698277 | 0 | 0 |
| LDB2 | 0 595058156 | 0 213161507 | 0 | 0 084810747 | 5 96E-167 | 0 |
| POSTN | 0 401619996 | 0 211515648 | 0 | 0 343508205 | 0 | 0 |
| FAT3 | 0 198149664 | 0 211190436 | 0 | 0 466636674 | 0 | 0 |
| CYTL1 | 0 151320173 | 0 210202161 | 0 | -0 2936115 | 0 | 0 |
| ADGRL3 | 0 446450667 | 0 209604382 | 0 | 0 016902188 | 4 28E-08 | 0 |
| MEOX2 | 0 604985628 | 0 20947846 | 0 | 0 247962962 | 0 | 0 |
| NRK | 0 200291257 | 0 208035816 | 0 | 0 465256437 | 0 | 0 |
| KALRN | 0 362071919 | 0 207389881 | 0 | 0 384666626 | 0 | 0 |
| EDNRA | 0 327568483 | 0 207212911 | 0 | -0 13203226 | 0 | 0 |

| | | | | | | |
|---|---|---|---|---|---|---|
| THSD4 | 0 527250576 | 0 206635744 | 0 | 0 038333183 | 1 81E-35 | 0 |
| GRIP1 | 0 265224344 | 0 206289033 | 0 | 0 432482128 | 0 | 0 |
| GSE1 | 0 409786602 | 0 203760451 | 0 | 0 343046134 | 0 | 0 |
| ZFHX4-AS1 | 0 202937313 | 0 203271991 | 0 | 0 450691927 | 0 | 0 |
| LINC00632 | 0 277321962 | 0 202222063 | 0 | 0 422341276 | 0 | 0 |
| ADGRL2 | 0 521301708 | 0 198619067 | 0 | 0 291656564 | 0 | 0 |
| CACNA2D3 | 0 194152025 | 0 198221836 | 0 | 0 445766512 | 0 | 0 |
| ZNF704 | 0 45484571 | 0 198214429 | 0 | 0 333266955 | 0 | 0 |
| PLCB4 | 0 381546135 | 0 197238935 | 0 | -0 05943167 | 7 82E-83 | 0 |
| ENOX1 | 0 326102682 | 0 195479815 | 0 | 0 352978003 | 0 | 0 |
| PDE3A | 0 379880452 | 0 19464426 | 0 | 0 3273726 | 0 | 0 |
| SOBP | 0 480735185 | 0 193979243 | 0 | 0 024734907 | 1 07E-15 | 0 |
| SNCAIP | 0 25267937 | 0 192644056 | 0 | 0 424244815 | 0 | 0 |
| TMTC1 | 0 472777979 | 0 191261767 | 0 | -0 02336749 | 3 59E-14 | 0 |
| ASXL3 | 0 13104643 | 0 190933692 | 0 | 0 488572557 | 0 | 0 |
| NEO1 | 0 433372675 | 0 189512804 | 0 | 0 332996355 | 0 | 0 |
| GLI3 | 0 580390626 | 0 189336274 | 0 | 0 221161718 | 0 | 0 |
| CNTN5 | 0 274942415 | 0 18785236 | 0 | 0 371258188 | 0 | 0 |
| COL16A1 | 0 359816109 | 0 187604679 | 0 | -0 07898388 | 5 33E-145 | 0 |
| PIEZO2 | 0 440577944 | 0 187319515 | 0 | 0 321876612 | 0 | 0 |
| ZNF385D | 0 326245455 | 0 187104354 | 0 | 0 354892912 | 0 | 0 |
| UNC5C | 0 454227028 | 0 187092164 | 0 | -0 0145374 | 2 45E-06 | 0 |
| LSAMP | 0 509813253 | 0 186472787 | 0 | 0 311501136 | 0 | 0 |
| TENM2 | 0 573699339 | 0 186297983 | 0 | 0 068673674 | 5 12E-110 | 0 |
| CSRNP3 | 0 421484457 | 0 184963886 | 0 | -0 04166222 | 1 37E-41 | 0 |
| AFF3 | 0 675981801 | 0 183114035 | 0 | 0 15286997 | 0 | 0 |
| MIR100HG | 0 729683425 | 0 182877555 | 0 | 0 07655724 | 2 53E-136 | 0 |
| CADM2 | 0 26040814 | 0 181870001 | 0 | 0 425317471 | 0 | 0 |
| NHSL2 | 0 350107556 | 0 18148848 | 0 | -0 0742732 | 2 09E-128 | 0 |
| IGF2BP2 | 0 537920466 | 0 181193643 | 0 | 0 20690315 | 0 | 0 |
| RBM20 | 0 201138375 | 0 18060924 | 0 | 0 447370898 | 0 | 0 |
| EPHA7 | 0 152462356 | 0 180177143 | 0 | 0 462942169 | 0 | 0 |
| RUNX1T1 | 0 69464697 | 0 18002655 | 0 | 0 096391637 | 2 80E-215 | 0 |
| TCF7L1 | 0 50420704 | 0 17976246 | 0 | 0 00694035 | 0 024474681 | 0 |
| LINC01515 | 0 254583008 | 0 179446405 | 0 | 0 409451788 | 0 | 0 |
| IL17RD | 0 224591194 | 0 179390233 | 0 | 0 425513076 | 0 | 0 |
| CACNB2 | 0 312272753 | 0 178507584 | 0 | 0 379900562 | 0 | 0 |
| MKX | 0 132264758 | 0 178481727 | 0 | 0 478871799 | 0 | 0 |
| COL13A1 | 0 123241515 | 0 17819396 | 0 | 0 454849861 | 0 | 0 |
| AMPH | 0 193866479 | 0 178173118 | 0 | 0 447983309 | 0 | 0 |
| PRRX1 | 0 43655175 | 0 178108734 | 0 | -0 05425446 | 2 53E-69 | 0 |
| PDE10A | 0 415459443 | 0 178095054 | 0 | 0 319006002 | 0 | 0 |
| COL14A1 | 0 289076926 | 0 176614941 | 0 | 0 374534474 | 0 | 0 |
| PLAG1 | 0 297814624 | 0 176520212 | 0 | 0 341505576 | 0 | 0 |

| | | | | | | |
|---|---|---|---|---|---|---|
| TGFB2 | 0 244141555 | 0 175811494 | 0 | 0 412108249 | 0 | 0 |
| COL4A5 | 0 378052959 | 0 173789886 | 0 | 0 330923066 | 0 | 0 |
| TSPAN18 | 0 147351088 | 0 173536616 | 0 | -0 29428662 | 0 | 0 |
| PPP3CA | 0 729930898 | 0 173048489 | 0 | 0 155905629 | 0 | 0 |
| NXPH2 | 0 059707601 | 0 172972893 | 0 | 0 510724123 | 0 | 0 |
| FSTL5 | 0 163332128 | 0 172447111 | 0 | 0 448464183 | 0 | 0 |
| COL1A1 | 0 827968247 | 0 17210485 | 0 | 0 102639623 | 5 85E-244 | 0 |
| COL1A2 | 0 91342255 | 0 171325656 | 0 | 0 136145753 | 0 | 0 |
| MYO15B | 0 333279397 | 0 170751103 | 0 | -0 09818305 | 2 58E-223 | 0 |
| SLC1A3 | 0 353486513 | 0 169886615 | 0 | 0 331458755 | 0 | 0 |
| ANTXR1 | 0 592754754 | 0 169827262 | 0 | 0 030721291 | 2 28E-23 | 0 |
| CPAMD8 | 0 1163789 | 0 169753432 | 0 | 0 483675061 | 0 | 0 |
| ZNF827 | 0 33515448 | 0 169607198 | 0 | 0 341874561 | 0 | 0 |
| ROR2 | 0 395623537 | 0 169560745 | 0 | -0 06404691 | 6 46E-96 | 0 |
| HMGCLL1 | 0 136785898 | 0 168428388 | 0 | -0 31577407 | 0 | 0 |
| IGF2BP3 | 0 427785498 | 0 168332293 | 0 | 0 223256686 | 0 | 0 |
| PHF14 | 0 742018998 | 0 168298951 | 0 | 0 177385248 | 0 | 0 |
| FAT4 | 0 289105481 | 0 168169936 | 0 | 0 378478713 | 0 | 0 |
| ADAMTS10 | 0 162085245 | 0 167795373 | 0 | 0 463383519 | 0 | 0 |
| ELN | 0 306780758 | 0 167360675 | 0 | -0 14911324 | 0 | 0 |
| PROM1 | 0 091422208 | 0 166470915 | 0 | 0 495527921 | 0 | 0 |
| PALLD | 0 587291314 | 0 165850325 | 0 | 0 181762185 | 0 | 0 |
| CACNG8 | 0 085625631 | 0 165438507 | 0 | 0 502127094 | 0 | 0 |
| NCOA2 | 0 706668443 | 0 165312283 | 0 | 0 165820686 | 0 | 0 |
| GLI2 | 0 451133616 | 0 164978802 | 0 | 0 259696573 | 0 | 0 |
| MEST | 0 371713845 | 0 164484356 | 0 | 0 32991368 | 0 | 0 |
| BBS9 | 0 744113 | 0 164459081 | 0 | 0 136304953 | 0 | 0 |
| PRDM6 | 0 117844701 | 0 164426564 | 0 | 0 479935432 | 0 | 0 |
| MPPED2 | 0 203299004 | 0 164139094 | 0 | 0 471016686 | 0 | 0 |
| PDE7B | 0 598446632 | 0 163769814 | 0 | 0 032737171 | 2 57E-26 | 0 |
| PLCE1 | 0 467219356 | 0 163109655 | 0 | 0 264048058 | 0 | 0 |
| DNM3 | 0 489529992 | 0 162833971 | 0 | 0 236250417 | 0 | 0 |
| C14orf39 | 0 218166416 | 0 162475934 | 0 | 0 44992231 | 0 | 0 |
| SLC9A9 | 0 463183644 | 0 162168708 | 0 | -0 03895508 | 1 42E-36 | 0 |
| SNHG14 | 0 686480364 | 0 161249228 | 0 | 0 085117086 | 3 78E-168 | 0 |
| DNMT3A | 0 371799509 | 0 161230518 | 0 | 0 339414063 | 0 | 0 |
| MEIS1 | 0 32448459 | 0 160728269 | 0 | 0 353400047 | 0 | 0 |
| NRG1 | 0 292884202 | 0 160403563 | 0 | 0 39048464 | 0 | 0 |
| STRBP | 0 302507091 | 0 160005335 | 0 | 0 353047064 | 0 | 0 |
| FBN2 | 0 468685157 | 0 159928627 | 0 | -0 01196745 | 0 000104828 | 0 |
| ADAMTS3 | 0 155194076 | 0 159517292 | 0 | 0 431999912 | 0 | 0 |
| OBSCN | 0 189212084 | 0 15902163 | 0 | 0 439120372 | 0 | 0 |
| DLK1 | 0 182530315 | 0 158801532 | 0 | 0 433674058 | 0 | 0 |
| SMAD6 | 0 135967334 | 0 15834609 | 0 | 0 469660864 | 0 | 0 |

| | | | | | | |
|---|---|---|---|---|---|---|
| ZMAT4 | 0 183900935 | 0 158315563 | 0 | -0 25841095 | 0 | 0 |
| COMP | 0 078277588 | 0 15748989 | 0 | 0 498135663 | 0 | 0 |
| UNC5B | 0 105042737 | 0 157333426 | 0 | 0 478266247 | 0 | 0 |
| SEMA6D | 0 272857932 | 0 157095705 | 0 | -0 19579049 | 0 | 0 |
| TOX | 0 271258876 | 0 156897877 | 0 | 0 41377052 | 0 | 0 |
| SLIT2 | 0 481344349 | 0 156247271 | 0 | 0 20532416 | 0 | 0 |
| SOX6 | 0 530553388 | 0 156126134 | 0 | 0 038151356 | 3 78E-35 | 0 |
| TRPS1 | 0 758057147 | 0 155983365 | 0 | 0 094454945 | 9 25E-207 | 0 |
| NPAS3 | 0 666625421 | 0 155546999 | 0 | 0 095301495 | 1 84E-210 | 0 |
| OLFM2 | 0 224657821 | 0 155373058 | 0 | -0 23273717 | 0 | 0 |
| PTH1R | 0 121537759 | 0 155256491 | 0 | 0 473211043 | 0 | 0 |
| FOXN3 | 0 6389846 | 0 154947324 | 0 | 0 179162043 | 0 | 0 |
| COL5A1 | 0 612238488 | 0 154927363 | 0 | 0 200637823 | 0 | 0 |
| REV3L | 0 661942472 | 0 154590281 | 0 | 0 197803421 | 0 | 0 |
| MPDZ | 0 52623213 | 0 154227247 | 0 | 0 033847725 | 5 09E-28 | 0 |
| ZFPM2-AS1 | 0 69007824 | 0 154083018 | 0 | 0 115612337 | 0 | 0 |
| AFF2 | 0 246444956 | 0 153829915 | 0 | 0 413537345 | 0 | 0 |
| ROR1 | 0 635377206 | 0 153435572 | 0 | 0 065192039 | 2 58E-99 | 0 |
| SAMD11 | 0 073889703 | 0 15340147 | 0 | 0 493923573 | 0 | 0 |
| ZNF521 | 0 574032476 | 0 15334507 | 0 | 0 046540906 | 1 79E-51 | 0 |
| ADAMTS9-AS1 | 0 119976776 | 0 15332875 | 0 | 0 470913274 | 0 | 0 |
| SUGCT | 0 323504217 | 0 152968776 | 0 | -0 13223381 | 0 | 0 |
| KCNQ1 | 0 09976966 | 0 152785041 | 0 | 0 487260138 | 0 | 0 |
| TSPAN11 | 0 206240125 | 0 152781069 | 0 | -0 24333058 | 0 | 0 |
| GREB1L | 0 238992214 | 0 152769411 | 0 | 0 396497772 | 0 | 0 |
| TNRC18 | 0 526784185 | 0 152572854 | 0 | 0 25322005 | 0 | 0 |
| CCN4 | 0 140183891 | 0 152439644 | 0 | -0 30981247 | 0 | 0 |
| PRR16 | 0 350897565 | 0 152429484 | 0 | 0 261138253 | 0 | 0 |
| FARP1 | 0 516199958 | 0 151903072 | 0 | 0 256492799 | 0 | 0 |
| RERG | 0 183101407 | 0 151786639 | 0 | -0 26223876 | 0 | 0 |
| TTC28 | 0 827701738 | 0 151314814 | 0 | 0 126616333 | 0 | 0 |
| EPHA6 | 0 117892292 | 0 15130059 | 0 | 0 458047775 | 0 | 0 |
| KIAA1549 | 0 186061564 | 0 151099703 | 0 | 0 434787727 | 0 | 0 |
| LIMCH1 | 0 482353277 | 0 150992743 | 0 | -0 04048056 | 2 33E-39 | 0 |
| PDZRN3 | 0 636471798 | 0 150943224 | 0 | 0 058104883 | 2 97E-79 | 0 |
| ERG | 0 138575317 | 0 150778474 | 0 | 0 45661729 | 0 | 0 |
| KCNMB2 | 0 201985494 | 0 150667099 | 0 | 0 436784616 | 0 | 0 |
| STK32B | 0 29210371 | 0 150612324 | 0 | 0 347332886 | 0 | 0 |
| FLI1 | 0 238611487 | 0 150173968 | 0 | 0 361438092 | 0 | 0 |
| GNAI1 | 0 374930993 | 0 150074814 | 0 | 0 286671252 | 0 | 0 |
| FAT1 | 0 570967619 | 0 149496654 | 0 | 0 218982555 | 0 | 0 |
| FGF14 | 0 364889303 | 0 148894573 | 0 | -0 09272973 | 2 55E-199 | 0 |
| SGCD | 0 503769203 | 0 14876757 | 0 | 0 242030524 | 0 | 0 |
| TENM3 | 0 558336982 | 0 148652307 | 0 | 0 175346834 | 0 | 0 |

| | | | | | | |
|---|---|---|---|---|---|---|
| PLXNA4 | 0 107955303 | 0 148458116 | 0 | 0 481111088 | 0 | 0 |
| FAM110B | 0 484970779 | 0 148257926 | 0 | -0 01889767 | 9 02E-10 | 0 |
| ITGA9 | 0 282709257 | 0 14817614 | 0 | -0 16446358 | 0 | 0 |
| LINC02008 | 0 060383393 | 0 148131631 | 0 | 0 497694267 | 0 | 0 |
| ADAMTS2 | 0 284565304 | 0 147920955 | 0 | -0 16335978 | 0 | 0 |
| DKK2 | 0 115084426 | 0 147507023 | 0 | 0 465506061 | 0 | 0 |
| TCF7L2 | 0 678408939 | 0 147486407 | 0 | 0 149845463 | 0 | 0 |
| ZNF804B | 0 107755421 | 0 147396587 | 0 | 0 47005439 | 0 | 0 |
| TMEM131L | 0 402495669 | 0 147256811 | 0 | 0 275139781 | 0 | 0 |
| SFMBT2 | 0 359197426 | 0 146837579 | 0 | 0 349927376 | 0 | 0 |
| ENAH | 0 502617502 | 0 146759803 | 0 | -0 01300141 | 2 51E-05 | 0 |
| SMAD9 | 0 213112258 | 0 146709133 | 0 | -0 22751031 | 0 | 0 |
| PCSK5 | 0 375949439 | 0 146700711 | 0 | -0 09612598 | 4 23E-214 | 0 |
| GRID1 | 0 227494241 | 0 146618667 | 0 | -0 20995103 | 0 | 0 |
| SLIT3 | 0 588233614 | 0 146592298 | 0 | 0 068696434 | 4 34E-110 | 0 |
| SDK2 | 0 095210447 | 0 146499669 | 0 | -0 35668316 | 0 | 0 |
| MRC2 | 0 528116731 | 0 146214389 | 0 | -0 04764028 | 7 50E-54 | 0 |
| KCNT2 | 0 606356247 | 0 146158491 | 0 | 0 053146629 | 1 36E-66 | 0 |
| EMCN | 0 17330719 | 0 146128286 | 0 | -0 28426028 | 0 | 0 |
| SYCP2L | 0 272572386 | 0 145706199 | 0 | -0 16872284 | 0 | 0 |
| GALNT10 | 0 438997925 | 0 145502592 | 0 | -0 05578167 | 3 52E-73 | 0 |
| LINC01266 | 0 22198321 | 0 145336233 | 0 | 0 417463852 | 0 | 0 |
| PRICKLE1 | 0 379138033 | 0 145207911 | 0 | -0 09872689 | 8 76E-226 | 0 |
| TNS3 | 0 541166169 | 0 145050049 | 0 | 0 226903858 | 0 | 0 |
| LOXL2 | 0 328339457 | 0 144989473 | 0 | 0 322782043 | 0 | 0 |
| MGA | 0 434638594 | 0 144946093 | 0 | -0 04038129 | 3 56E-39 | 0 |
| SVEP1 | 0 377462831 | 0 144941039 | 0 | -0 11379691 | 9 47E-300 | 0 |
| GMDS | 0 575203213 | 0 144706951 | 0 | 0 018944271 | 8 20E-10 | 0 |
| SRGAP3 | 0 272819859 | 0 144238304 | 0 | 0 350954539 | 0 | 0 |
| BRWD1 | 0 667977004 | 0 144018409 | 0 | 0 173554246 | 0 | 0 |
| FLRT2 | 0 52381451 | 0 143845631 | 0 | -0 00354231 | 0 250899656 | 0 |
| TSBP1-AS1 | 0 223924921 | 0 143359158 | 0 | 0 416514279 | 0 | 0 |
| KLHL14 | 0 068407226 | 0 143332603 | 0 | 0 49797209 | 0 | 0 |
| LRFN5 | 0 322295407 | 0 143184278 | 0 | -0 1403731 | 0 | 0 |
| DNAH14 | 0 530686642 | 0 142879077 | 0 | 0 217890289 | 0 | 0 |
| TBX18 | 0 277379071 | 0 142055345 | 0 | -0 14149217 | 0 | 0 |
| CACNA2D1 | 0 630817993 | 0 14194562 | 0 | 0 14333618 | 0 | 0 |
| LMO3 | 0 12967581 | 0 141499448 | 0 | 0 456102653 | 0 | 0 |
| APBA2 | 0 107298548 | 0 141372343 | 0 | 0 488944618 | 0 | 0 |
| ASTN1 | 0 093992119 | 0 14110805 | 0 | 0 484706359 | 0 | 0 |
| EGFR | 0 678256648 | 0 14101446 | 0 | 0 064666539 | 9 53E-98 | 0 |
| SULF1 | 0 46972264 | 0 140969768 | 0 | 0 253955454 | 0 | 0 |
| TLE2 | 0 254516381 | 0 140884265 | 0 | -0 18571486 | 0 | 0 |
| PDE9A | 0 197216881 | 0 140638522 | 0 | -0 23786623 | 0 | 0 |

| | | | | | | |
|---|---|---|---|---|---|---|
| NREP | 0 428061526 | 0 140485287 | 0 | 0 279606518 | 0 | 0 |
| FNBP1L | 0 341236603 | 0 140421263 | 0 | 0 337430742 | 0 | 0 |
| SVIL | 0 657050123 | 0 140027193 | 0 | 0 091686196 | 6 91E-195 | 0 |
| LUC7L | 0 542089433 | 0 139952559 | 0 | 0 228255163 | 0 | 0 |
| ALDH1A2 | 0 186585064 | 0 139675757 | 0 | 0 428764287 | 0 | 0 |
| SPON1 | 0 175439264 | 0 139669647 | 0 | -0 2821421 | 0 | 0 |
| PLCL1 | 0 354286041 | 0 139558275 | 0 | -0 09943509 | 5 08E-229 | 0 |
| FARS2 | 0 601787516 | 0 139353422 | 0 | 0 198492373 | 0 | 0 |
| DOCK11 | 0 317384021 | 0 139316088 | 0 | 0 348339907 | 0 | 0 |
| VLDLR-AS1 | 0 170461251 | 0 138851958 | 0 | -0 29755072 | 0 | 0 |
| WWOX | 0 680798005 | 0 138843565 | 0 | 0 102007949 | 5 62E-241 | 0 |
| MIR181A1HG | 0 44659344 | 0 138750214 | 0 | 0 225899455 | 0 | 0 |
| KCTD8 | 0 192181759 | 0 138661897 | 0 | 0 436424806 | 0 | 0 |
| TNMD | 0 067512516 | 0 138478893 | 0 | 0 492315875 | 0 | 0 |
| LRIG3 | 0 365098704 | 0 138226442 | 0 | 0 321227545 | 0 | 0 |
| GLCCI1 | 0 404694371 | 0 137784891 | 0 | 0 284181334 | 0 | 0 |
| PSD3 | 0 689649921 | 0 137722548 | 0 | 0 110173617 | 5 48E-281 | 0 |
| EYA2 | 0 41106204 | 0 137657029 | 0 | -0 06677767 | 4 05E-104 | 0 |
| NDST4 | 0 133939959 | 0 137628803 | 0 | 0 454602828 | 0 | 0 |
| FGF10 | 0 112971388 | 0 137591396 | 0 | 0 466096891 | 0 | 0 |
| PTPRG | 0 845653043 | 0 137471637 | 0 | 0 070442747 | 1 14E-115 | 0 |
| ABI3BP | 0 47970722 | 0 137382551 | 0 | -0 03086992 | 1 40E-23 | 0 |
| BICC1 | 0 784527232 | 0 137209544 | 0 | 0 105190617 | 3 38E-256 | 0 |
| COL23A1 | 0 144019722 | 0 137170082 | 0 | 0 448895233 | 0 | 0 |
| BNC2 | 0 816432202 | 0 136784131 | 0 | 0 052760926 | 1 18E-65 | 0 |
| TIAM2 | 0 50231292 | 0 136776872 | 0 | -0 02174386 | 1 81E-12 | 0 |
| MGP | 0 473625098 | 0 136717649 | 0 | -0 07025355 | 4 65E-115 | 0 |
| ARHGEF40 | 0 279768137 | 0 136653107 | 0 | 0 373665662 | 0 | 0 |
| FNDC1 | 0 209514382 | 0 136615001 | 0 | -0 24015801 | 0 | 0 |
| BCL11A | 0 096571548 | 0 136558373 | 0 | 0 48093893 | 0 | 0 |
| MIPOL1 | 0 474510289 | 0 136547281 | 0 | 0 265768546 | 0 | 0 |
| SNAP25 | 0 092269327 | 0 136205823 | 0 | 0 480053823 | 0 | 0 |
| DACH1 | 0 132578858 | 0 136183265 | 0 | 0 444868932 | 0 | 0 |
| COL26A1 | 0 080485808 | 0 136074873 | 0 | 0 486446231 | 0 | 0 |
| ZFHX3 | 0 657716396 | 0 135904264 | 0 | 0 163905129 | 0 | 0 |
| LRP1 | 0 650263654 | 0 135761713 | 0 | 0 051037497 | 1 51E-61 | 0 |
| PDS5B | 0 702014049 | 0 134993182 | 0 | 0 162392672 | 0 | 0 |
| CRPPA | 0 295140013 | 0 134969359 | 0 | -0 15884757 | 0 | 0 |
| PLAGL1 | 0 507776361 | 0 134791077 | 0 | -0 02658667 | 6 75E-18 | 0 |
| KCNMB2-AS1 | 0 228845824 | 0 134709675 | 0 | -0 2169866 | 0 | 0 |
| SLC5A4-AS1 | 0 098998686 | 0 134707877 | 0 | 0 476492422 | 0 | 0 |
| TNRC6A | 0 799204279 | 0 134601763 | 0 | 0 090397843 | 1 75E-189 | 0 |
| MID1 | 0 469275285 | 0 134535781 | 0 | 0 25735423 | 0 | 0 |
| INTS6L | 0 217833279 | 0 134176662 | 0 | 0 415791736 | 0 | 0 |

| | | | | | | |
|---|---|---|---|---|---|---|
| FIRRE | 0 281909729 | 0 134049767 | 0 | 0 334954945 | 0 | 0 |
| STRA6 | 0 085530449 | 0 133650993 | 0 | 0 485071747 | 0 | 0 |
| SSBP3 | 0 370866726 | 0 133449226 | 0 | -0 11459155 | 6 00E-304 | 0 |
| TTC6 | 0 060278693 | 0 13336561 | 0 | 0 493411549 | 0 | 0 |
| SMAD7 | 0 100264606 | 0 133287484 | 0 | -0 35493686 | 0 | 0 |
| TNRC6C | 0 590051589 | 0 133174731 | 0 | 0 041492241 | 2 89E-41 | 0 |
| DEPTOR | 0 205326379 | 0 13316641 | 0 | -0 24759337 | 0 | 0 |
| MAGI1 | 0 51277341 | 0 133028308 | 0 | 0 242690527 | 0 | 0 |
| COL12A1 | 0 575041404 | 0 132991948 | 0 | 0 02132316 | 4 77E-12 | 0 |
| FOXP1 | 0 661761626 | 0 132902297 | 0 | 0 152009945 | 0 | 0 |
| KCTD1 | 0 254011917 | 0 13276956 | 0 | 0 406525707 | 0 | 0 |
| PLXNA2 | 0 186546991 | 0 132757438 | 0 | 0 427889523 | 0 | 0 |
| STX18-AS1 | 0 246987493 | 0 132652739 | 0 | 0 384854663 | 0 | 0 |
| CCDC178 | 0 06318174 | 0 132625876 | 0 | 0 492288017 | 0 | 0 |
| DSE | 0 404361234 | 0 132424956 | 0 | -0 07613791 | 7 49E-135 | 0 |
| ZFPM2 | 0 581142563 | 0 132340242 | 0 | 0 116297671 | 0 | 0 |
| MIR503HG | 0 161856808 | 0 13229084 | 0 | 0 427606671 | 0 | 0 |
| CDK13 | 0 734509147 | 0 132140792 | 0 | 0 077718001 | 1 94E-140 | 0 |
| PCDH9 | 0 48612248 | 0 132104058 | 0 | 0 236087497 | 0 | 0 |
| RFLNA | 0 165311911 | 0 132041657 | 0 | -0 28718673 | 0 | 0 |
| HIBADH | 0 546658164 | 0 132026534 | 0 | -0 00805557 | 0 009025561 | 0 |
| CNKSR3 | 0 458786241 | 0 131882037 | 0 | 0 271169347 | 0 | 0 |
| PDGFRA | 0 379956597 | 0 131697809 | 0 | 0 322159055 | 0 | 0 |
| RANBP17 | 0 443024119 | 0 131437745 | 0 | 0 261016349 | 0 | 0 |
| SENP6 | 0 715368068 | 0 131052324 | 0 | 0 104240284 | 1 33E-251 | 0 |
| IQCA1 | 0 103386572 | 0 130744102 | 0 | 0 471448648 | 0 | 0 |
| DLG2 | 0 50394053 | 0 130634336 | 0 | 0 016656015 | 6 70E-08 | 0 |
| ZBTB46 | 0 280967429 | 0 130571974 | 0 | -0 17156025 | 0 | 0 |
| SMARCA1 | 0 473234852 | 0 130490029 | 0 | -0 00793712 | 0 010091144 | 0 |
| ZNF638 | 0 845833889 | 0 12984781 | 0 | 0 112243355 | 1 25E-291 | 0 |
| PELI2 | 0 362690602 | 0 129800736 | 0 | -0 10874963 | 8 77E-274 | 0 |
| TET2 | 0 430012754 | 0 129397933 | 0 | -0 07376734 | 1 10E-126 | 0 |
| CERS6 | 0 570425082 | 0 129278852 | 0 | 0 197210717 | 0 | 0 |
| MXRA5 | 0 180712341 | 0 129253232 | 0 | -0 29290509 | 0 | 0 |
| CCND3 | 0 429060935 | 0 129194345 | 0 | -0 05117291 | 7 27E-62 | 0 |
| MFAP4 | 0 134215987 | 0 129182843 | 0 | 0 444934335 | 0 | 0 |
| MACROD2 | 0 434962213 | 0 128959974 | 0 | -0 02041672 | 3 63E-11 | 0 |
| PTBP2 | 0 664065028 | 0 128800738 | 0 | 0 173817266 | 0 | 0 |
| ROBO1 | 0 614256344 | 0 128573213 | 0 | 0 122939748 | 0 | 0 |
| KCNE4 | 0 209723782 | 0 12850446 | 0 | -0 24601531 | 0 | 0 |
| PTPRS | 0 434962213 | 0 128370971 | 0 | -0 03685699 | 6 46E-33 | 0 |
| NCOA1 | 0 793455293 | 0 128070159 | 0 | 0 095106663 | 1 32E-209 | 0 |
| WNK2 | 0 081608955 | 0 128021703 | 0 | 0 488788633 | 0 | 0 |
| MECOM | 0 335059298 | 0 12799393 | 0 | 0 246682783 | 0 | 0 |

| | | | | | |
|---|---|---|---|---|---|
| CREBBP | 0 62069064 | 0 127990767 | 0 0 035972014 | 1 96E-31 | 0 |
| TRERF1 | 0 431383374 | 0 12795086 | 0 0 237454257 | 0 | 0 |
| SGCZ | 0 131312939 | 0 127919206 | 0 0 434701822 | 0 | 0 |
| FAM13A | 0 592202699 | 0 127810027 | 0 0 001518322 | 0 6226258 | 0 |
| MYH10 | 0 42709067 | 0 127762435 | 0 0 298743614 | 0 | 0 |
| MAGI2 | 0 665483238 | 0 127440737 | 0 0 09823954 | 1 43E-223 | 0 |
| EVL | 0 432535075 | 0 127310334 | 0 -0 04826404 | 3 17E-55 | 0 |
| PCMTD1 | 0 583341265 | 0 127169215 | 0 0 020725487 | 1 84E-11 | 0 |
| LTBP2 | 0 173173935 | 0 127128886 | 0 -0 27112702 | 0 | 0 |
| DOK6 | 0 271087548 | 0 127106918 | 0 0 346944464 | 0 | 0 |
| SDK1 | 0 393681826 | 0 127096517 | 0 0 288188573 | 0 | 0 |
| TBC1D32 | 0 459766614 | 0 127079182 | 0 -0 02762084 | 3 42E-19 | 0 |
| TNS1 | 0 396879938 | 0 127050575 | 0 -0 06057879 | 5 41E-86 | 0 |
| NUAK1 | 0 225019512 | 0 127005666 | 0 -0 22723171 | 0 | 0 |
| INPP4B | 0 415507034 | 0 126953912 | 0 0 272804837 | 0 | 0 |
| PCCA | 0 692248387 | 0 126814935 | 0 0 057850251 | 1 42E-78 | 0 |
| GPR137C | 0 15328092 | 0 126780106 | 0 0 427155037 | 0 | 0 |
| PLEKHA5 | 0 664464792 | 0 126695163 | 0 0 153345416 | 0 | 0 |
| UNC5D | 0 053082942 | 0 126655718 | 0 0 498063985 | 0 | 0 |
| TNFRSF25 | 0 074080067 | 0 126644487 | 0 0 493636835 | 0 | 0 |
| LGR5 | 0 114703699 | 0 126294563 | 0 0 461086632 | 0 | 0 |
| EDIL3 | 0 343130723 | 0 125995898 | 0 -0 11700632 | 0 | 0 |
| DCHS2 | 0 09445851 | 0 125880712 | 0 0 471004476 | 0 | 0 |
| ADARB1 | 0 30025128 | 0 125718879 | 0 -0 15217683 | 0 | 0 |
| NKTR | 0 742837563 | 0 125616037 | 0 0 140893182 | 0 | 0 |
| CHD9 | 0 870019607 | 0 125573176 | 0 0 092449447 | 4 00E-198 | 0 |
| GNA14 | 0 158877615 | 0 125568991 | 0 -0 29929765 | 0 | 0 |
| LINC01524 | 0 128819173 | 0 125405074 | 0 0 458416351 | 0 | 0 |
| THSD7A | 0 348689345 | 0 125399975 | 0 0 332149272 | 0 | 0 |
| MCC | 0 591193771 | 0 125310021 | 0 0 015686534 | 3 68E-07 | 0 |
| FAM20A | 0 346957035 | 0 125309183 | 0 -0 10445841 | 1 18E-252 | 0 |
| PRTG | 0 173478517 | 0 125289749 | 0 -0 28530823 | 0 | 0 |
| MUSK | 0 154147075 | 0 125239443 | 0 -0 31515811 | 0 | 0 |
| TRABD2B | 0 282528412 | 0 125230533 | 0 -0 15072098 | 0 | 0 |
| ADGRA3 | 0 449705888 | 0 125180968 | 0 0 214613258 | 0 | 0 |
| LINC01482 | 0 2911043 | 0 125062988 | 0 -0 1590265 | 0 | 0 |
| COL27A1 | 0 311159125 | 0 125043236 | 0 0 308414192 | 0 | 0 |
| GHR | 0 450524452 | 0 124925531 | 0 -0 0457158 | 1 00E-49 | 0 |
| TBL1X | 0 390626487 | 0 124629054 | 0 0 304574613 | 0 | 0 |
| KSR1 | 0 16847195 | 0 124627433 | 0 -0 28540499 | 0 | 0 |
| ADCY5 | 0 081542327 | 0 12425862 | 0 -0 37160631 | 0 | 0 |
| CDC42BPA | 0 751346824 | 0 124077543 | 0 0 141332888 | 0 | 0 |
| ENPP1 | 0 204412633 | 0 123887481 | 0 0 400559152 | 0 | 0 |
| SMAD3 | 0 507900097 | 0 123841715 | 0 0 178980273 | 0 | 0 |

| | | | | | | |
|---|---|---|---|---|---|---|
| ITGA11 | 0 192115132 | 0 123754768 | 0 | -0 25694062 | 0 | 0 |
| SYNDIG1 | 0 062544022 | 0 123449214 | 0 | 0 480169272 | 0 | 0 |
| CYP7B1 | 0 297510042 | 0 123417304 | 0 | -0 16112308 | 0 | 0 |
| RAD51B | 0 595791057 | 0 12318865 | 0 | 0 16856422 | 0 | 0 |
| FGFR2 | 0 118520493 | 0 122876771 | 0 | 0 473877287 | 0 | 0 |
| RCOR1 | 0 43340123 | 0 122853168 | 0 | 0 249312095 | 0 | 0 |
| TARID | 0 119653157 | 0 122674968 | 0 | -0 33585343 | 0 | 0 |
| ESRRG | 0 243227808 | 0 122603759 | 0 | -0 21979454 | 0 | 0 |
| ZMYM2 | 0 65175801 | 0 122564494 | 0 | 0 06170017 | 3 85E-89 | 0 |
| NFIX | 0 407359464 | 0 122563337 | 0 | 0 23671714 | 0 | 0 |
| CECR2 | 0 170670652 | 0 122369176 | 0 | -0 28917009 | 0 | 0 |
| CCDC102B | 0 27239154 | 0 122260951 | 0 | -0 2036047 | 0 | 0 |
| ADAMTS17 | 0 236165312 | 0 121920483 | 0 | 0 377159422 | 0 | 0 |
| ANOS1 | 0 241771525 | 0 121892763 | 0 | -0 20708344 | 0 | 0 |
| EML4 | 0 569063981 | 0 121880085 | 0 | 0 16940603 | 0 | 0 |
| ST6GALNAC3 | 0 276313034 | 0 121733415 | 0 | 0 374450297 | 0 | 0 |
| POTEF | 0 113932725 | 0 121634311 | 0 | 0 454678238 | 0 | 0 |
| DLEU2 | 0 737021949 | 0 121619112 | 0 | 0 135426267 | 0 | 0 |
| KCNB2 | 0 146884697 | 0 121482287 | 0 | 0 425968378 | 0 | 0 |
| GAREM1 | 0 542346424 | 0 121458986 | 0 | 0 02255851 | 2 62E-13 | 0 |
| SLC24A3 | 0 178941958 | 0 121447307 | 0 | -0 29067067 | 0 | 0 |
| SLC22A3 | 0 226037959 | 0 121293382 | 0 | -0 21668028 | 0 | 0 |
| STARD4-AS1 | 0 276836535 | 0 121260462 | 0 | 0 328649686 | 0 | 0 |
| ZNF292 | 0 664540938 | 0 121227057 | 0 | 0 064787962 | 4 15E-98 | 0 |
| KLF3-AS1 | 0 248329558 | 0 121151221 | 0 | -0 21592643 | 0 | 0 |
| ACAN | 0 032990044 | 0 12108612 | 0 | 0 501124027 | 0 | 0 |
| MEG8 | 0 554653443 | 0 12088512 | 0 | -0 0273639 | 7 24E-19 | 0 |
| TMEM196 | 0 068055053 | 0 120844103 | 0 | 0 486972957 | 0 | 0 |
| TMEM14B | 0 519093488 | 0 12082671 | 0 | -0 00516181 | 0 094307197 | 0 |
| CDYL | 0 48412366 | 0 120790511 | 0 | 0 23005387 | 0 | 0 |
| SPRY4-AS1 | 0 400877577 | 0 120686049 | 0 | 0 270888178 | 0 | 0 |
| WRN | 0 574394167 | 0 120612756 | 0 | 0 19166139 | 0 | 0 |
| EEPD1 | 0 190649331 | 0 120513664 | 0 | -0 26269195 | 0 | 0 |
| MARK1 | 0 308065714 | 0 120298999 | 0 | 0 328679857 | 0 | 0 |
| RCBTB2 | 0 188983648 | 0 120278951 | 0 | 0 426022567 | 0 | 0 |
| BASP1 | 0 28243323 | 0 120246106 | 0 | 0 324888349 | 0 | 0 |
| PAPPA2 | 0 083112829 | 0 120187089 | 0 | 0 475073497 | 0 | 0 |
| FAM135A | 0 394005444 | 0 120138885 | 0 | -0 08781811 | 6 72E-179 | 0 |
| BCKDHB | 0 410471912 | 0 120136283 | 0 | -0 06532273 | 1 05E-99 | 0 |
| PRKG1 | 0 837648246 | 0 120125342 | 0 | 0 1266851 | 0 | 0 |
| PPARG | 0 195094325 | 0 120117841 | 0 | -0 25922853 | 0 | 0 |
| ATP8A2 | 0 116673964 | 0 12003639 | 0 | 0 456910304 | 0 | 0 |
| SENP7 | 0 567683844 | 0 119915653 | 0 | 0 227354831 | 0 | 0 |
| MIR4300HG | 0 049656393 | 0 119701353 | 0 | 0 496660548 | 0 | 0 |

| | | | | | | |
|---|---|---|---|---|---|---|
| ARID1B | 0 828720184 | 0 11967357 | 0 | 0 106837096 | 2 94E-264 | 0 |
| RCOR3 | 0 462184234 | 0 119604012 | 0 | -0 05929375 | 1 86E-82 | 0 |
| ZNF117 | 0 192990805 | 0 119594118 | 0 | 0 419233027 | 0 | 0 |
| CRISPLD1 | 0 338609583 | 0 119191605 | 0 | 0 334444897 | 0 | 0 |
| SPIDR | 0 775780016 | 0 119108326 | 0 | 0 145888107 | 0 | 0 |
| ARHGAP24 | 0 559669528 | 0 119097097 | 0 | 0 144805459 | 0 | 0 |
| DAB2IP | 0 158125678 | 0 118797286 | 0 | 0 454739869 | 0 | 0 |
| OTX2-AS1 | 0 077192515 | 0 118724039 | 0 | 0 482067197 | 0 | 0 |
| ITGA9-AS1 | 0 279435 | 0 118642856 | 0 | -0 18970483 | 0 | 0 |
| COLEC12 | 0 462346043 | 0 118624466 | 0 | -0 05977733 | 8 86E-84 | 0 |
| NHS | 0 349098627 | 0 118608437 | 0 | -0 11363462 | 6 77E-299 | 0 |
| FBXL7 | 0 807646913 | 0 118597974 | 0 | 0 079445488 | 1 11E-146 | 0 |
| PIAS2 | 0 458167558 | 0 118575552 | 0 | 0 25446889 | 0 | 0 |
| SYN2 | 0 129428338 | 0 118534734 | 0 | 0 454400061 | 0 | 0 |
| CCDC88A | 0 565380442 | 0 118503024 | 0 | 0 142036601 | 0 | 0 |
| PLEKHH2 | 0 459166968 | 0 118496595 | 0 | -0 04464915 | 1 64E-47 | 0 |
| CCND2 | 0 418990691 | 0 118375083 | 0 | -0 07600549 | 2 18E-134 | 0 |
| PKNOX2 | 0 247625212 | 0 118057722 | 0 | -0 21013347 | 0 | 0 |
| NTF3 | 0 055291161 | 0 117995346 | 0 | 0 500936794 | 0 | 0 |
| TBC1D4 | 0 352077821 | 0 117905601 | 0 | -0 12581991 | 0 | 0 |
| ADAMTSL3 | 0 275304106 | 0 117889278 | 0 | -0 20046926 | 0 | 0 |
| DDR2 | 0 62379357 | 0 117850949 | 0 | 0 144875811 | 0 | 0 |
| SEMA3F | 0 051245931 | 0 117835656 | 0 | 0 489018977 | 0 | 0 |
| RAI14 | 0 572614266 | 0 117687203 | 0 | 0 157870191 | 0 | 0 |
| SKIL | 0 426329215 | 0 11753187 | 0 | -0 08858514 | 5 16E-182 | 0 |
| MROH9 | 0 204983724 | 0 117336407 | 0 | -0 25613431 | 0 | 0 |
| XKR6 | 0 435066913 | 0 117248209 | 0 | -0 06414629 | 3 29E-96 | 0 |
| NXN | 0 344330015 | 0 11721719 | 0 | 0 336706366 | 0 | 0 |
| PTPRD | 0 406978736 | 0 117197764 | 0 | 0 175938315 | 0 | 0 |
| KDM2B | 0 304781938 | 0 117062475 | 0 | 0 319444524 | 0 | 0 |
| PACRG | 0 131132093 | 0 117056692 | 0 | -0 32524161 | 0 | 0 |
| NCALD | 0 30335421 | 0 116950588 | 0 | -0 18700284 | 0 | 0 |
| ZNF518A | 0 469313358 | 0 116850093 | 0 | 0 268016409 | 0 | 0 |
| CASC15 | 0 544973444 | 0 116826988 | 0 | -0 00108097 | 0 726058447 | 0 |
| PWRN1 | 0 103967181 | 0 116760498 | 0 | 0 477442748 | 0 | 0 |
| JAKMIP2 | 0 223553711 | 0 116721007 | 0 | -0 23358305 | 0 | 0 |
| EMX2OS | 0 0810569 | 0 116699992 | 0 | 0 481168853 | 0 | 0 |
| SHROOM3 | 0 238440159 | 0 116431364 | 0 | 0 377178746 | 0 | 0 |
| CPE | 0 222906474 | 0 116424145 | 0 | 0 403334371 | 0 | 0 |
| NOS1AP | 0 223039729 | 0 116412709 | 0 | -0 2238553 | 0 | 0 |
| CUX1 | 0 709666673 | 0 116333049 | 0 | 0 138483752 | 0 | 0 |
| HACE1 | 0 298033542 | 0 116296775 | 0 | 0 356402281 | 0 | 0 |
| ENTPD1-AS1 | 0 40927262 | 0 116246722 | 0 | 0 275262538 | 0 | 0 |
| MARK4 | 0 27833089 | 0 11619454 | 0 | -0 18547831 | 0 | 0 |

| | | | | | | |
|---|---|---|---|---|---|---|
| BMPR1B | 0 458519731 | 0 116074461 | 0 | 0 234192396 | 0 | 0 |
| PRRX2 | 0 09724734 | 0 116069095 | 0 | 0 470881835 | 0 | 0 |
| ADGRG2 | 0 188488702 | 0 115757935 | | 0 391116853 | 0 | 0 |
| INSR | 0 478269974 | 0 115637764 | 0 | -0 03154724 | 1 49E-24 | 3 17E-308 |
| ARHGEF3 | 0 346899926 | 0 115312743 | 8 76E-308 | -0 16154103 | 0 | 1 72E-306 |
| ZNF618 | 0 361148655 | 0 115261844 | 1 64E-307 | 0 257017405 | 0 | 3 21E-306 |
| PAN3 | 0 701471512 | 0 115143898 | 6 96E-307 | 0 060416664 | 1 53E-85 | 1 36E-305 |
| LINC01239 | 0 068426263 | 0 115096503 | 1 25E-306 | -0 40325045 | 0 | 2 43E-305 |
| WBP2NL | 0 122242105 | 0 115065708 | 1 82E-306 | 0 466078474 | 0 | 3 54E-305 |
| TTC3 | 0 801393463 | 0 115044923 | 2 34E-306 | 0 061472458 | 1 69E-88 | 4 56E-305 |
| STAG1 | 0 835725572 | 0 115043889 | 2 37E-306 | 0 121473659 | 0 | 4 61E-305 |
| MROH7 | 0 084730921 | 0 11480482 | 4 43E-305 | -0 38083319 | 0 | 8 57E-304 |
| LINC01004 | 0 499333727 | 0 114734503 | 1 05E-304 | 0 207191167 | 0 | 2 02E-303 |
| LINC01091 | 0 161837772 | 0 114654102 | 2 80E-304 | 0 432812815 | 0 | 5 38E-303 |
| ZDHHC17 | 0 531676534 | 0 114623402 | 4 07E-304 | -0 03664104 | 1 50E-32 | 7 82E-303 |
| ANGPT1 | 0 191172831 | 0 114606681 | 4 99E-304 | 0 402849385 | 0 | 9 57E-303 |
| MFSD4B | 0 472863642 | 0 114420405 | 4 84E-303 | -0 05933254 | 1 46E-82 | 9 27E-302 |
| SYCP2 | 0 108859531 | 0 11436984 | 8 97E-303 | 0 464145354 | 0 | 1 71E-301 |
| SPRY1 | 0 254516381 | 0 114270456 | 3 01E-302 | -0 23074674 | 0 | 5 73E-301 |
| UNC13C | 0 097009385 | 0 114266745 | 3 15E-302 | 0 455293105 | 0 | 5 99E-301 |
| CHD6 | 0 613247416 | 0 114069727 | 3 46E-301 | 0 0209517 | 1 11E-11 | 6 56E-300 |
| STK33 | 0 255239763 | 0 114036613 | 5 17E-301 | 0 384795304 | 0 | 9 79E-300 |
| COA1 | 0 573927776 | 0 113900417 | 2 70E-300 | 0 17477796 | 0 | 5 10E-299 |
| TBX1 | 0 039243494 | 0 113817482 | 7 38E-300 | 0 499293563 | 0 | 1 39E-298 |
| CCDC171 | 0 342226495 | 0 113816849 | 7 44E-300 | 0 324788502 | 0 | 1 40E-298 |
| ANO1 | 0 106289619 | 0 113770493 | 1 31E-299 | 0 470056617 | 0 | 2 45E-298 |
| LURAP1L | 0 198568464 | 0 113688962 | 3 50E-299 | 0 39777635 | 0 | 6 56E-298 |
| CSMD1 | 0 132169576 | 0 11361841 | 8 23E-299 | -0 33258764 | 0 | 1 54E-297 |
| RNF216 | 0 571795702 | 0 113587317 | 1 20E-298 | 0 197310876 | 0 | 2 24E-297 |
| C5orf17 | 0 07716396 | 0 113555065 | 1 77E-298 | 0 463607252 | 0 | 3 29E-297 |
| SMOC1 | 0 041280387 | 0 113544013 | 2 02E-298 | 0 49463261 | 0 | 3 76E-297 |
| SNAP91 | 0 077221069 | 0 113476355 | 4 59E-298 | 0 481248888 | 0 | 8 50E-297 |
| TRMT11 | 0 499961927 | 0 113466722 | 5 15E-298 | 0 000534775 | 0 862387371 | 9 53E-297 |
| NET1 | 0 162275609 | 0 113465437 | 5 23E-298 | 0 398685357 | 0 | 9 66E-297 |
| MED13L | 0 877063068 | 0 113453452 | 6 05E-298 | 0 070746983 | 1 17E-116 | 1 11E-296 |
| LPAR1 | 0 633892368 | 0 113404357 | 1 09E-297 | 0 045058925 | 2 34E-48 | 2 01E-296 |
| TNKS | 0 71204622 | 0 113402674 | 1 12E-297 | 0 159209299 | 0 | 2 05E-296 |
| LAMA4 | 0 579257962 | 0 113352996 | 2 04E-297 | 0 063509776 | 2 42E-94 | 3 73E-296 |
| RYBP | 0 427062116 | 0 113286079 | 4 57E-297 | -0 07902816 | 3 68E-145 | 8 34E-296 |
| CDON | 0 224286612 | 0 113262448 | 6 07E-297 | 0 407429232 | 0 | 1 11E-295 |
| DMKN | 0 079305553 | 0 113250319 | 7 03E-297 | -0 38918515 | 0 | 1 28E-295 |
| TRAF3IP2-AS1 | 0 427157298 | 0 113108269 | 3 89E-296 | -0 0664068 | 5 51E-103 | 7 08E-295 |
| PRDM5 | 0 360253945 | 0 113006209 | 1 33E-295 | 0 319287925 | 0 | 2 41E-294 |
| RIN2 | 0 570939065 | 0 112974747 | 1 94E-295 | 0 021672945 | 2 13E-12 | 3 52E-294 |

| | | | | | | |
|---|---|---|---|---|---|---|
| RIPOR3 | 0 097485294 | 0 112826881 | 1 15E-294 | 0 450141666 | 0 | 2 07E-293 |
| MAK | 0 308094268 | 0 112805635 | 1 48E-294 | -0 1687454 | 0 | 2 67E-293 |
| CPNE5 | 0 066341779 | 0 112784445 | 1 91E-294 | -0 40335279 | 0 | 3 44E-293 |
| SLC25A21 | 0 191372713 | 0 112770674 | 2 26E-294 | 0 392530361 | 0 | 4 05E-293 |
| THRB | 0 353772058 | 0 112645406 | 1 02E-293 | -0 12582026 | 0 | 1 81E-292 |
| GPC4 | 0 17109897 | 0 112643941 | 1 03E-293 | 0 435255858 | 0 | 1 84E-292 |
| PDSS2 | 0 483857151 | 0 112597806 | 1 80E-293 | -0 03058106 | 3 60E-23 | 3 20E-292 |
| CACNB4 | 0 184529135 | 0 112554858 | 3 01E-293 | 0 41186321 | 0 | 5 34E-292 |
| ADAM22 | 0 150349318 | 0 112282124 | 7 87E-292 | 0 427390461 | 0 | 1 40E-290 |
| AP2A2 | 0 359892254 | 0 112281552 | 7 93E-292 | -0 12073257 | 0 | 1 40E-290 |
| ADAM33 | 0 195056252 | 0 112102198 | 6 76E-291 | -0 26104729 | 0 | 1 19E-289 |
| PCDH9-AS2 | 0 086368049 | 0 1120928 | 7 56E-291 | 0 470250901 | 0 | 1 33E-289 |
| RARB | 0 390359978 | 0 112039021 | 1 44E-290 | 0 264073285 | 0 | 2 53E-289 |
| MGAT4A | 0 159058461 | 0 111937224 | 4 83E-290 | -0 3066483 | 0 | 8 47E-289 |
| SEMA6A | 0 27470446 | 0 111904318 | 7 15E-290 | -0 19685869 | 0 | 1 25E-288 |
| RNF150 | 0 392301689 | 0 111879984 | 9 56E-290 | 0 265354925 | 0 | 1 67E-288 |
| PDCD4 | 0 436827778 | 0 11182162 | 1 92E-289 | -0 08671689 | 1 78E-174 | 3 34E-288 |
| CNTNAP3 | 0 105575755 | 0 111756482 | 4 16E-289 | 0 471271564 | 0 | 7 24E-288 |
| IGSF9B | 0 096181302 | 0 111681421 | 1 02E-288 | -0 36380183 | 0 | 1 76E-287 |
| DENND1B | 0 417077535 | 0 111553026 | 4 67E-288 | 0 25288175 | 0 | 8 10E-287 |
| L3MBTL4 | 0 435038358 | 0 11151196 | 7 61E-288 | 0 271476804 | 0 | 1 32E-286 |
| SLCO5A1 | 0 05703299 | 0 111491344 | 9 72E-288 | 0 478670642 | 0 | 1 68E-286 |
| BICRAL | 0 387989949 | 0 111476718 | 1 16E-287 | -0 09808773 | 6 97E-223 | 1 99E-286 |
| TBCK | 0 587024804 | 0 111459988 | 1 41E-287 | 0 018444324 | 2 25E-09 | 2 43E-286 |
| NOL4L | 0 13530106 | 0 111430088 | 2 01E-287 | 0 451698775 | 0 | 3 45E-286 |
| RYR3 | 0 155574803 | 0 111264191 | 1 44E-286 | -0 29308116 | 0 | 2 46E-285 |
| TBX3 | 0 164816965 | 0 111144875 | 5 90E-286 | 0 420603801 | 0 | 1 01E-284 |
| ITSN1 | 0 514343911 | 0 111140867 | 6 18E-286 | -0 04135768 | 5 22E-41 | 1 06E-284 |
| GOLGA8B | 0 324218081 | 0 111104624 | 9 49E-286 | 0 309754123 | 0 | 1 62E-284 |
| CASD1 | 0 458386477 | 0 111081078 | 1 25E-285 | 0 240762107 | 0 | 2 14E-284 |
| GPR63 | 0 051540995 | 0 110982834 | 4 01E-285 | 0 489064141 | 0 | 6 81E-284 |
| SULT1E1 | 0 11154366 | 0 110971785 | 4 56E-285 | 0 453118847 | 0 | 7 74E-284 |
| TTYH3 | 0 19049704 | 0 110888562 | 1 22E-284 | 0 405856493 | 0 | 2 07E-283 |
| COL22A1 | 0 05882241 | 0 110880619 | 1 34E-284 | 0 491998375 | 0 | 2 26E-283 |
| SLC39A8 | 0 099150978 | 0 110845477 | 2 03E-284 | 0 452386828 | 0 | 3 42E-283 |
| NR3C2 | 0 383525918 | 0 110836497 | 2 25E-284 | -0 09535179 | 1 11E-210 | 3 80E-283 |
| LTBP1 | 0 498762635 | 0 11072014 | 8 89E-284 | 0 17209771 | 0 | 1 49E-282 |
| LMCD1-AS1 | 0 339437665 | 0 110700589 | 1 12E-283 | 0 309921026 | 0 | 1 88E-282 |
| SH3GL3 | 0 0808475 | 0 110608581 | 3 31E-283 | 0 467720234 | 0 | 5 53E-282 |
| RPS6KA6 | 0 144467077 | 0 110569659 | 5 23E-283 | -0 31118751 | 0 | 8 73E-282 |
| KIAA1328 | 0 520654471 | 0 110495992 | 1 24E-282 | 0 219620161 | 0 | 2 07E-281 |
| PHACTR1 | 0 297976433 | 0 110415872 | 3 19E-282 | 0 316206763 | 0 | 5 31E-281 |
| ATP11A | 0 364023148 | 0 110323869 | 9 41E-282 | -0 10408023 | 7 83E-251 | 1 56E-280 |
| TRA2A | 0 680131732 | 0 11010162 | 1 28E-280 | 0 052837755 | 7 70E-66 | 2 11E-279 |

| | | | | | | |
|---|---|---|---|---|---|---|
| LAMA2 | 0 817298357 | 0 109925754 | 1 00E-279 | 0 10087245 | 1 16E-235 | 1 65E-278 |
| HNRNPA2B1 | 0 837981382 | 0 109892635 | 1 47E-279 | 0 128735772 | 0 | 2 43E-278 |
| CBFB | 0 463059907 | 0 109891793 | 1 49E-279 | 0 241756813 | 0 | 2 45E-278 |
| CCDC85A | 0 10176848 | 0 109871493 | 1 89E-279 | -0 36041079 | 0 | 3 10E-278 |
| KIAA1109 | 0 279139936 | 0 109758527 | 7 06E-279 | 0 287317318 | 0 | 1 16E-277 |
| GRIN2B | 0 076564314 | 0 109690175 | 1 57E-278 | -0 3885703 | 0 | 2 57E-277 |
| ZNRF1 | 0 234709029 | 0 109683114 | 1 70E-278 | 0 363485046 | 0 | 2 79E-277 |
| DLGAP1 | 0 415783061 | 0 109581439 | 5 58E-278 | 0 251145933 | 0 | 9 11E-277 |
| LMO7 | 0 314471455 | 0 109396534 | 4 81E-277 | 0 277043825 | 0 | 7 80E-276 |
| MNAT1 | 0 65028269 | 0 109330549 | 1 04E-276 | 0 14565315 | 0 | 1 68E-275 |
| HIVEP1 | 0 346652453 | 0 109092648 | 1 65E-275 | 0 288649909 | 0 | 2 66E-274 |
| EXT1 | 0 769812111 | 0 109086532 | 1 77E-275 | 0 070754212 | 1 11E-116 | 2 85E-274 |
| EPHB1 | 0 109011822 | 0 109051379 | 2 66E-275 | -0 35484143 | 0 | 4 28E-274 |
| PXDN | 0 418667073 | 0 108957831 | 7 87E-275 | 0 225895453 | 0 | 1 26E-273 |
| FKBP10 | 0 391102397 | 0 108799211 | 4 94E-274 | -0 12118947 | 0 | 7 91E-273 |
| TIA1 | 0 420275647 | 0 108676637 | 2 04E-273 | 0 252385688 | 0 | 3 26E-272 |
| HMGA2 | 0 343958805 | 0 108587223 | 5 74E-273 | 0 196788058 | 0 | 9 15E-272 |
| TBL1XR1 | 0 758380766 | 0 108577132 | 6 44E-273 | 0 115771019 | | 1 03E-271 |
| HECW2 | 0 247273039 | 0 108459492 | 2 51E-272 | 0 352331809 | 0 | 3 99E-271 |
| TSBP1 | 0 11033485 | 0 108432531 | 3 42E-272 | -0 36483905 | 0 | 5 43E-271 |
| SAMMSON | 0 082694028 | 0 108358605 | 8 02E-272 | 0 461599809 | 0 | 1 27E-270 |
| ADNP | 0 61343778 | 0 108355562 | 8 31E-272 | 0 166773486 | 0 | 1 32E-270 |
| FANCC | 0 422188803 | 0 108197255 | 5 15E-271 | 0 219009555 | 0 | 8 14E-270 |
| RORA-AS1 | 0 296710514 | 0 108176619 | 6 53E-271 | -0 19562575 | 0 | 1 03E-269 |
| PTPN4 | 0 480411567 | 0 108163263 | 7 61E-271 | 0 208112405 | 0 | 1 20E-269 |
| CALD1 | 0 842521559 | 0 108141491 | 9 78E-271 | 0 094172256 | 1 57E-205 | 1 54E-269 |
| CYP39A1 | 0 106327692 | 0 108040163 | 3 14E-270 | -0 35508666 | 0 | 4 92E-269 |
| CAMK1D | 0 272924559 | 0 108037967 | 3 22E-270 | 0 364107401 | 0 | 5 04E-269 |
| ZNF185 | 0 070853401 | 0 108036383 | 3 28E-270 | 0 474777646 | 0 | 5 12E-269 |
| DPY19L2 | 0 211408502 | 0 107959673 | 7 91E-270 | 0 37205602 | 0 | 1 23E-268 |
| MGAT4C | 0 103986218 | 0 107951051 | 8 73E-270 | 0 454027914 | 0 | 1 36E-268 |
| KAT6B | 0 528763968 | 0 107920355 | 1 24E-269 | -0 01694792 | 3 94E-08 | 1 93E-268 |
| GALNT17 | 0 291075746 | 0 107895868 | 1 65E-269 | -0 18602251 | 0 | 2 55E-268 |
| VWDE | 0 120509794 | 0 107836526 | 3 25E-269 | 0 43606789 | 0 | 5 04E-268 |
| TMEM108 | 0 144600331 | 0 107793491 | 5 32E-269 | 0 412475895 | 0 | 8 24E-268 |
| ROBO2 | 0 308922351 | 0 107675828 | 2 05E-268 | -0 15276957 | 0 | 3 17E-267 |
| MAML2 | 0 786897261 | 0 107586545 | 5 70E-268 | 0 095977025 | 1 93E-213 | 8 79E-267 |
| HOTAIRM1 | 0 145123832 | 0 10756344 | 7 43E-268 | -0 31039303 | 0 | 1 14E-266 |
| RBM26 | 0 658192305 | 0 107510702 | 1 36E-267 | 0 159829006 | 0 | 2 09E-266 |
| MEGF6 | 0 079943272 | 0 107309994 | 1 34E-266 | 0 47337096 | 0 | 2 06E-265 |
| IFT88 | 0 407426091 | 0 10729027 | 1 68E-266 | -0 09486174 | 1 55E-208 | 2 58E-265 |
| AEBP2 | 0 560678457 | 0 107268439 | 2 16E-266 | -0 01668064 | 6 41E-08 | 3 30E-265 |
| RNF152 | 0 145371305 | 0 107153506 | 8 01E-266 | -0 31507624 | 0 | 1 22E-264 |
| DDAH1 | 0 314138318 | 0 107112402 | 1 28E-265 | -0 15381922 | 0 | 1 95E-264 |

| | | | | | | |
|---|---|---|---|---|---|---|
| RMST | 0 134606233 | 0 107089858 | 1 66E-265 | 0 430570729 | 0 | 2 51E-264 |
| ADCY2 | 0 245198074 | 0 107079662 | 1 86E-265 | 0 358952179 | 0 | 2 82E-264 |
| GPCPD1 | 0 470331804 | 0 106984078 | 5 52E-265 | 0 238071283 | 0 | 8 34E-264 |
| SKAP2 | 0 467266947 | 0 106978751 | 5 86E-265 | 0 206472544 | 0 | 8 85E-264 |
| RBFOX2 | 0 792018046 | 0 106978292 | 5 89E-265 | 0 073096568 | 2 02E-124 | 8 88E-264 |
| RALGAPA2 | 0 402990615 | 0 106951233 | 8 02E-265 | -0 09539374 | 7 23E-211 | 1 21E-263 |
| PRPF40B | 0 309017533 | 0 1069175 | 1 18E-264 | 0 309432918 | 0 | 1 77E-263 |
| ARMCX4 | 0 396004264 | 0 106709835 | 1 25E-263 | -0 0793852 | 1 84E-146 | 1 86E-262 |
| DENND5B | 0 476394891 | 0 10661151 | 3 80E-263 | -0 05896543 | 1 45E-81 | 5 67E-262 |
| LINC00877 | 0 051807504 | 0 106579186 | 5 48E-263 | 0 483846483 | 0 | 8 17E-262 |
| GOLGA8A | 0 514305838 | 0 10657628 | 5 67E-263 | 0 217925541 | 0 | 8 43E-262 |
| ATP10A | 0 153062001 | 0 106554498 | 7 25E-263 | 0 434892266 | 0 | 1 08E-261 |
| NLGN4X | 0 241914298 | 0 106383565 | 5 02E-262 | -0 19704885 | 0 | 7 45E-261 |
| NAA16 | 0 374635929 | 0 106382403 | 5 09E-262 | 0 277629607 | 0 | 7 54E-261 |
| SLC5A4 | 0 071053283 | 0 106239654 | 2 56E-261 | 0 482554557 | 0 | 3 78E-260 |
| C11orf58 | 0 510431935 | 0 10623123 | 2 81E-261 | -0 02690861 | 2 70E-18 | 4 15E-260 |
| MEOX1 | 0 08589214 | 0 106087001 | 1 43E-260 | 0 463656209 | 0 | 2 11E-259 |
| IGDCC3 | 0 030819897 | 0 106050982 | 2 15E-260 | 0 502333484 | 0 | 3 17E-259 |
| FTO | 0 691068131 | 0 106009844 | 3 42E-260 | 0 053102711 | 1 75E-66 | 5 03E-259 |
| SORCS2 | 0 359349717 | 0 105988558 | 4 35E-260 | -0 0962932 | 7 67E-215 | 6 38E-259 |
| PTN | 0 312529744 | 0 105940451 | 7 48E-260 | 0 299732709 | 0 | 1 10E-258 |
| N4BP2 | 0 276503398 | 0 105879016 | 1 49E-259 | 0 335030537 | 0 | 2 18E-258 |
| SATB1-AS1 | 0 070967619 | 0 105854235 | 1 97E-259 | -0 40351677 | 0 | 2 88E-258 |
| NSD3 | 0 602901144 | 0 105850946 | 2 05E-259 | 0 181873288 | 0 | 2 99E-258 |
| TOP2B | 0 504654395 | 0 105791864 | 3 98E-259 | 0 231391653 | 0 | 5 80E-258 |
| CDKAL1 | 0 726380613 | 0 10578198 | 4 45E-259 | 0 139069781 | 0 | 6 47E-258 |
| MYCBP2 | 0 650415945 | 0 105769148 | 5 14E-259 | 0 031852823 | 5 32E-25 | 7 46E-258 |
| MN1 | 0 044231025 | 0 105623737 | 2 64E-258 | 0 494704278 | 0 | 3 81E-257 |
| PRSS23 | 0 354352668 | 0 105556556 | 5 60E-258 | -0 11437817 | 8 10E-303 | 8 10E-257 |
| TAFA1 | 0 073166321 | 0 105509665 | 9 48E-258 | 0 486757076 | 0 | 1 37E-256 |
| RBMX | 0 612952352 | 0 105397249 | 3 35E-257 | 0 178258299 | 0 | 4 82E-256 |
| LINC02476 | 0 091860045 | 0 105349973 | 5 68E-257 | 0 464335525 | 0 | 8 17E-256 |
| CDH8 | 0 077068778 | 0 105331865 | 6 96E-257 | 0 472895165 | 0 | 9 99E-256 |
| WIF1 | 0 087681559 | 0 105312684 | 8 63E-257 | 0 457576771 | 0 | 1 24E-255 |
| MKRN3 | 0 081504255 | 0 105301202 | 9 81E-257 | 0 469325849 | 0 | 1 40E-255 |
| TSC22D2 | 0 521368335 | 0 105279288 | 1 25E-256 | 0 205578962 | 0 | 1 79E-255 |
| ATP9B | 0 570187128 | 0 105202846 | 2 95E-256 | 0 021411468 | 3 90E-12 | 4 20E-255 |
| ZNF586 | 0 283565894 | 0 105185197 | 3 59E-256 | -0 18063933 | 0 | 5 11E-255 |
| EPB41L4A | 0 30304011 | 0 10518302 | 3 68E-256 | -0 14739707 | 0 | 5 22E-255 |
| NDUFAF6 | 0 396242219 | 0 105037141 | 1 88E-255 | 0 246619784 | 0 | 2 66E-254 |
| C2CD6 | 0 195398907 | 0 104870719 | 1 20E-254 | 0 392318327 | 0 | 1 70E-253 |
| PDE3B | 0 188374484 | 0 10485892 | 1 37E-254 | 0 422280513 | 0 | 1 93E-253 |
| PREX1 | 0 11112486 | 0 104817898 | 2 17E-254 | -0 34879131 | 0 | 3 05E-253 |
| CASK | 0 615255754 | 0 104784118 | 3 16E-254 | 0 022901392 | 1 14E-13 | 4 43E-253 |

| | | | | | | |
|---|---|---|---|---|---|---|
| EPHB2 | 0 164712265 | 0 104751374 | 4 55E-254 | 0 360652381 | 0 | 6 37E-253 |
| CCDC144NL-A | 0 171660543 | 0 104705405 | 7 59E-254 | 0 3794513 | 0 | 1 06E-252 |
| UBE3D | 0 287753898 | 0 104631277 | 1 73E-253 | -0 17719074 | 0 | 2 41E-252 |
| LINC01844 | 0 050389294 | 0 104574601 | 3 25E-253 | -0 42265439 | 0 | 4 53E-252 |
| POU6F2 | 0 068093126 | 0 104372967 | 3 05E-252 | 0 475669041 | 0 | 4 25E-251 |
| MEIS2 | 0 515210066 | 0 104215461 | 1 75E-251 | 0 205494585 | 0 | 2 43E-250 |
| PDGFRB | 0 269174392 | 0 104198655 | 2 11E-251 | -0 18004699 | 0 | 2 92E-250 |
| CA8 | 0 067712398 | 0 104175009 | 2 74E-251 | 0 491291937 | 0 | 3 79E-250 |
| FGF1 | 0 209200282 | 0 1041489 | 3 66E-251 | -0 25248214 | 0 | 5 06E-250 |
| MBNL3 | 0 13630047 | 0 104139366 | 4 07E-251 | 0 433740857 | 0 | 5 61E-250 |
| COL9A2 | 0 094591765 | 0 104134753 | 4 28E-251 | -0 37209579 | 0 | 5 90E-250 |
| SFRP2 | 0 101482934 | 0 104028203 | 1 39E-250 | 0 453818285 | 0 | 1 91E-249 |
| ANKRD6 | 0 304943748 | 0 10396393 | 2 83E-250 | -0 15325226 | 0 | 3 88E-249 |
| ETV1 | 0 537615884 | 0 103864544 | 8 49E-250 | 0 016502675 | 8 83E-08 | 1 16E-248 |
| JAZF1 | 0 560878339 | 0 103647223 | 9 33E-249 | 0 005277734 | 0 087140641 | 1 27E-247 |
| ASCC3 | 0 633387904 | 0 10364513 | 9 55E-249 | 0 035373738 | 1 88E-30 | 1 30E-247 |
| BCOR | 0 210037882 | 0 103640936 | 1 00E-248 | 0 376695991 | 0 | 1 36E-247 |
| LMX1A | 0 069197236 | 0 103566823 | 2 26E-248 | 0 471874917 | 0 | 3 07E-247 |
| SLC6A16 | 0 133492604 | 0 103503204 | 4 55E-248 | 0 427017645 | 0 | 6 18E-247 |
| IL12A-AS1 | 0 039490967 | 0 103491464 | 5 18E-248 | 0 492558934 | 0 | 7 03E-247 |
| HMBOX1 | 0 56407645 | 0 103462327 | 7 14E-248 | 0 000730529 | 0 812823246 | 9 67E-247 |
| DTNB | 0 427861644 | 0 103435188 | 9 62E-248 | 0 254692199 | 0 | 1 30E-246 |
| CYP1B1-AS1 | 0 252136833 | 0 1034109 | 1 26E-247 | -0 202282 | 0 | 1 70E-246 |
| TCF4 | 0 834545316 | 0 103337619 | 2 81E-247 | 0 051808153 | 2 29E-63 | 3 78E-246 |
| UST | 0 360977328 | 0 103054164 | 6 30E-246 | 0 25926056 | 0 | 8 46E-245 |
| SHANK2 | 0 117673374 | 0 102938639 | 2 23E-245 | 0 451737398 | 0 | 2 99E-244 |
| HAPLN1 | 0 058127582 | 0 102885947 | 3 97E-245 | 0 47821382 | 0 | 5 32E-244 |
| PARD3 | 0 835297253 | 0 10285589 | 5 51E-245 | 0 115594877 | 0 | 7 38E-244 |
| XPO1 | 0 676771811 | 0 102831885 | 7 17E-245 | 0 124026572 | 0 | 9 56E-244 |
| CLMP | 0 454036664 | 0 102745846 | 1 83E-244 | -0 0511177 | 9 79E-62 | 2 44E-243 |
| KIAA1217 | 0 580400145 | 0 102681307 | 3 71E-244 | 0 013205723 | 1 86E-05 | 4 92E-243 |
| EPC1 | 0 591869563 | 0 102642822 | 5 65E-244 | 0 030053402 | 1 97E-22 | 7 48E-243 |
| ICA1L | 0 293988312 | 0 102536349 | 1 80E-243 | -0 17101732 | 0 | 2 38E-242 |
| UBN2 | 0 456301993 | 0 102451235 | 4 56E-243 | -0 06304007 | 5 60E-93 | 6 00E-242 |
| EPB41L2 | 0 776817498 | 0 102441783 | 5 05E-243 | 0 085516575 | 1 02E-169 | 6 65E-242 |
| CBR4 | 0 484847043 | 0 102430487 | 5 71E-243 | -0 05978634 | 8 37E-84 | 7 50E-242 |
| RGS6 | 0 375035693 | 0 102312935 | 2 05E-242 | -0 09439494 | 1 69E-206 | 2 68E-241 |
| DLEU1 | 0 426062706 | 0 102291193 | 2 60E-242 | 0 255372032 | 0 | 3 40E-241 |
| PLCB1 | 0 635472388 | 0 102266018 | 3 42E-242 | 0 130861658 | 0 | 4 46E-241 |
| C19orf38 | 0 050665321 | 0 102228881 | 5 11E-242 | -0 42056356 | 0 | 6 66E-241 |
| PARD3B | 0 761597914 | 0 102209432 | 6 32E-242 | 0 095924466 | 3 30E-213 | 8 21E-241 |
| SPECC1L | 0 552340523 | 0 102182757 | 8 44E-242 | 0 174668182 | 0 | 1 10E-240 |
| TET1 | 0 371599627 | 0 102152758 | 1 17E-241 | 0 270729716 | 0 | 1 52E-240 |
| SCFD2 | 0 53679732 | 0 102099637 | 2 08E-241 | 0 182964636 | 0 | 2 69E-240 |

| | | | | | | |
|---|---|---|---|---|---|---|
| MSC-AS1 | 0 471416878 | 0 101980304 | 7 59E-241 | -0 0759551 | 3 26E-134 | 9 81E-240 |
| HIP1 | 0 488996973 | 0 101893668 | 1 94E-240 | 0 213830168 | 0 | 2 50E-239 |
| ARID3A | 0 154899012 | 0 101892766 | 1 96E-240 | -0 30714348 | 0 | 2 52E-239 |
| RB1CC1 | 0 597951686 | 0 101866462 | 2 60E-240 | 0 012400945 | 5 83E-05 | 3 35E-239 |
| C2CD5 | 0 467914184 | 0 101827182 | 3 98E-240 | 0 243664385 | 0 | 5 12E-239 |
| COG6 | 0 360301536 | 0 101708547 | 1 44E-239 | -0 12039467 | 0 | 1 84E-238 |
| CDK19 | 0 489901201 | 0 101704604 | 1 50E-239 | 0 247083514 | 0 | 1 92E-238 |
| HUNK | 0 120243285 | 0 101689849 | 1 76E-239 | 0 456066602 | 0 | 2 25E-238 |
| PRMT7 | 0 234899393 | 0 101621178 | 3 69E-239 | 0 370458348 | 0 | 4 72E-238 |
| LOXL1 | 0 167605795 | 0 101597867 | 4 75E-239 | 0 394408005 | 0 | 6 06E-238 |
| COL3A1 | 0 784784223 | 0 101589657 | 5 19E-239 | 0 045963256 | 3 02E-50 | 6 62E-238 |
| SLC38A1 | 0 145304677 | 0 101390514 | 4 44E-238 | 0 437516657 | 0 | 5 65E-237 |
| ANKRD11 | 0 624602616 | 0 101322119 | 9 28E-238 | 0 152793963 | 0 | 1 18E-236 |
| TBX15 | 0 32369458 | 0 101285307 | 1 38E-237 | -0 16862164 | 0 | 1 75E-236 |
| DMTF1 | 0 595676838 | 0 101113778 | 8 72E-237 | 0 173301279 | 0 | 1 10E-235 |
| MAP2K6 | 0 254954218 | 0 101076881 | 1 30E-236 | -0 2281002 | 0 | 1 64E-235 |
| EPB41L5 | 0 309417296 | 0 100913519 | 7 48E-236 | -0 16911063 | 0 | 9 45E-235 |
| SNAI2 | 0 111524624 | 0 100869491 | 1 20E-235 | 0 433431179 | 0 | 1 51E-234 |
| LINC01122 | 0 062049076 | 0 100867006 | 1 23E-235 | 0 484423044 | 0 | 1 55E-234 |
| L3MBTL3 | 0 182958634 | 0 10079551 | 2 65E-235 | -0 28158146 | 0 | 3 33E-234 |
| RFTN1 | 0 27491386 | 0 100754577 | 4 10E-235 | 0 306740246 | 0 | 5 16E-234 |
| TSPAN4 | 0 275123261 | 0 100731541 | 5 25E-235 | -0 20022152 | 0 | 6 59E-234 |
| CHD3 | 0 303715901 | 0 100726793 | 5 53E-235 | 0 33290662 | 0 | 6 93E-234 |
| FLG-AS1 | 0 363613866 | 0 100721248 | 5 86E-235 | 0 325230536 | 0 | 7 34E-234 |
| ZRANB2 | 0 741781044 | 0 10066046 | 1 12E-234 | 0 050146261 | 1 78E-59 | 1 40E-233 |
| SNAP25-AS1 | 0 072271611 | 0 100636273 | 1 46E-234 | 0 46593195 | 0 | 1 82E-233 |
| PLOD2 | 0 359521045 | 0 100621547 | 1 70E-234 | 0 264255353 | 0 | 2 12E-233 |
| GRIA1 | 0 176333974 | 0 100592097 | 2 33E-234 | -0 26957983 | 0 | 2 90E-233 |
| ADGRB3 | 0 215501323 | 0 100499061 | 6 30E-234 | 0 411435417 | 0 | 7 84E-233 |
| CNTN3 | 0 176124574 | 0 100477612 | 7 92E-234 | 0 436620062 | 0 | 9 84E-233 |
| TRPC4 | 0 060040738 | 0 100406562 | 1 69E-233 | 0 466591644 | 0 | 2 10E-232 |
| R3HDM2 | 0 665626011 | 0 100405071 | 1 72E-233 | 0 129843469 | 0 | 2 13E-232 |
| SLC8A3 | 0 031714607 | 0 100402922 | 1 76E-233 | 0 498142583 | 0 | 2 17E-232 |
| COL4A6 | 0 079410253 | 0 100199891 | 1 53E-232 | 0 470626281 | 0 | 1 89E-231 |
| CA10 | 0 039833622 | 0 100192916 | 1 65E-232 | 0 487625674 | 0 | 2 03E-231 |
| PDE5A | 0 494631741 | 0 100135339 | 3 04E-232 | -0 0464681 | 2 56E-51 | 3 75E-231 |
| DGKI | 0 171374998 | 0 100106489 | 4 13E-232 | 0 389452976 | 0 | 5 08E-231 |
| RBFOX3 | 0 032085816 | 0 100028097 | 9 51E-232 | 0 495123675 | 0 | 1 17E-230 |
| C1GALT1 | 0 277084008 | 0 100007589 | 1 18E-231 | 0 336395961 | 0 | 1 45E-230 |
| PLEKHG2 | 0 140517028 | 0 100006645 | 1 19E-231 | -0 3309132 | 0 | 1 46E-230 |
| DANT2 | 0 517161295 | 0 100003931 | 1 23E-231 | 0 015698355 | 3 61E-07 | 1 50E-230 |
| SRSF12 | 0 098103977 | 0 099967657 | 1 81E-231 | -0 37107973 | 0 | 2 21E-230 |
| TENT5A | 0 444299556 | 0 099935928 | 2 53E-231 | -0 04365272 | 1 72E-45 | 3 09E-230 |
| RHOBTB3 | 0 532076298 | 0 099927313 | 2 77E-231 | -0 0259769 | 3 72E-17 | 3 38E-230 |

| | | | | | | |
|---|---|---|---|---|---|---|
| UBR5 | 0 75336468 | 0 099884902 | 4 35E-231 | 0 09871408 | 1 00E-225 | 5 29E-230 |
| MRTFB | 0 458748168 | 0 099779128 | 1 33E-230 | 0 225252524 | 0 | 1 62E-229 |
| MAPK10 | 0 583741029 | 0 099743383 | 1 95E-230 | 0 020318682 | 4 50E-11 | 2 36E-229 |
| STXBP5 | 0 541680151 | 0 099667188 | 4 37E-230 | 0 167692472 | 0 | 5 29E-229 |
| MGMT | 0 360549009 | 0 099663316 | 4 55E-230 | -0 12685087 | 0 | 5 51E-229 |
| PIK3R1 | 0 585930213 | 0 099597482 | 9 13E-230 | -0 01417012 | 4 37E-06 | 1 10E-228 |
| HHIP | 0 127239154 | 0 099583338 | 1 06E-229 | -0 34708796 | 0 | 1 28E-228 |
| MON2 | 0 75751461 | 0 099582884 | 1 07E-229 | 0 049747591 | 1 46E-58 | 1 28E-228 |
| NEDD4L | 0 282614075 | 0 099543107 | 1 62E-229 | -0 19272818 | 0 | 1 95E-228 |
| SPRED1 | 0 658135196 | 0 099509223 | 2 32E-229 | 0 02195568 | 1 10E-12 | 2 79E-228 |
| NCAM1 | 0 404342198 | 0 099476459 | 3 28E-229 | 0 254878495 | 0 | 3 93E-228 |
| CDH23 | 0 088642897 | 0 099408695 | 6 72E-229 | 0 467699503 | 0 | 8 03E-228 |
| ITM2A | 0 198006891 | 0 099404509 | 7 02E-229 | 0 413192602 | 0 | 8 39E-228 |
| MEGF10 | 0 15049209 | 0 099398186 | 7 50E-229 | 0 431119222 | 0 | 8 95E-228 |
| DAAM2 | 0 359768518 | 0 099367611 | 1 04E-228 | -0 09651566 | 7 88E-216 | 1 23E-227 |
| PNISR | 0 764938798 | 0 099351451 | 1 23E-228 | 0 093152032 | 3 96E-201 | 1 46E-227 |
| FMNL3 | 0 330804668 | 0 099323674 | 1 65E-228 | 0 259572804 | 0 | 1 96E-227 |
| GUCY1A2 | 0 180493423 | 0 099322202 | 1 67E-228 | 0 43079713 | 0 | 1 99E-227 |
| LINC00174 | 0 428937199 | 0 099284364 | 2 49E-228 | 0 22761339 | 0 | 2 96E-227 |
| LINC01278 | 0 517751423 | 0 09914877 | 1 04E-227 | 0 006381311 | 0 038603547 | 1 23E-226 |
| SYNE1 | 0 614437189 | 0 099128152 | 1 29E-227 | 0 023637486 | 1 82E-14 | 1 53E-226 |
| KMT2C | 0 832984333 | 0 098999763 | 4 99E-227 | 0 097629617 | 8 14E-221 | 5 89E-226 |
| PHF21A | 0 605175991 | 0 098992655 | 5 38E-227 | 0 015504177 | 5 02E-07 | 6 34E-226 |
| SMARCC1 | 0 571519674 | 0 09895403 | 8 07E-227 | 0 174161604 | 0 | 9 50E-226 |
| SRSF6 | 0 359635263 | 0 098854424 | 2 30E-226 | -0 13291639 | 0 | 2 70E-225 |
| GPRC5C | 0 045849118 | 0 098736915 | 7 89E-226 | -0 42967171 | 0 | 9 25E-225 |
| TRPC1 | 0 528744932 | 0 098626298 | 2 51E-225 | -0 02656884 | 7 10E-18 | 2 94E-224 |
| SAP30L-AS1 | 0 221811883 | 0 098566184 | 4 72E-225 | 0 358921989 | 0 | 5 51E-224 |
| AVPR1A | 0 020464107 | 0 09853396 | 6 61E-225 | 0 502014237 | 0 | 7 71E-224 |
| DOP1A | 0 400649141 | 0 098507492 | 8 72E-225 | -0 07516217 | 1 85E-131 | 1 02E-223 |
| TRPC4AP | 0 573261503 | 0 098483358 | 1 12E-224 | 0 172500692 | 0 | 1 31E-223 |
| ADCY9 | 0 341436485 | 0 098428774 | 1 99E-224 | -0 14820906 | 0 | 2 31E-223 |
| TASP1 | 0 490805429 | 0 098376087 | 3 44E-224 | -0 04700026 | 1 84E-52 | 3 99E-223 |
| ASTN2 | 0 383402182 | 0 098364213 | 3 90E-224 | 0 258979211 | 0 | 4 51E-223 |
| LRP6 | 0 515076812 | 0 098190749 | 2 38E-223 | -0 04127199 | 7 59E-41 | 2 76E-222 |
| IL1RAPL2 | 0 218899317 | 0 098162376 | 3 20E-223 | -0 2523918 | 0 | 3 70E-222 |
| CCDC3 | 0 083731511 | 0 09814511 | 3 83E-223 | 0 467286964 | 0 | 4 42E-222 |
| ATXN7L1 | 0 49910529 | 0 098122909 | 4 83E-223 | -0 05361413 | 9 76E-68 | 5 56E-222 |
| KLF12 | 0 746968457 | 0 098109112 | 5 58E-223 | 0 076086924 | 1 13E-134 | 6 41E-222 |
| EDN3 | 0 030324951 | 0 097958859 | 2 67E-222 | 0 494470604 | 0 | 3 06E-221 |
| KLHDC10 | 0 52827854 | 0 097936092 | 3 38E-222 | -0 0329635 | 1 17E-26 | 3 87E-221 |
| FBXL4 | 0 385819802 | 0 097915404 | 4 19E-222 | -0 10263106 | 6 42E-244 | 4 79E-221 |
| PCBP1-AS1 | 0 404656298 | 0 09790244 | 4 79E-222 | -0 09411707 | 2 72E-205 | 5 48E-221 |
| ENOX2 | 0 271230321 | 0 097869739 | 6 73E-222 | -0 20202776 | 0 | 7 68E-221 |

| | | | | | | |
|---|---|---|---|---|---|---|
| CTIF | 0 268393901 | 0 097641066 | 7 23E-221 | -0 18702395 | 0 | 8 23E-220 |
| AHCYL2 | 0 545725381 | 0 097592944 | 1 19E-220 | -0 00878857 | 0 004390068 | 1 35E-219 |
| SLC16A9 | 0 080733281 | 0 097574983 | 1 43E-220 | 0 449551423 | 0 | 1 63E-219 |
| MDM4 | 0 592136072 | 0 097526849 | 2 36E-220 | 0 008086198 | 0 0087669 | 2 68E-219 |
| SLC12A9 | 0 181987779 | 0 097505103 | 2 96E-220 | 0 378185701 | 0 | 3 35E-219 |
| ITCH | 0 674268527 | 0 097441734 | 5 70E-220 | 0 035521176 | 1 08E-30 | 6 45E-219 |
| TAPT1-AS1 | 0 354228931 | 0 09740655 | 8 20E-220 | -0 13275819 | 0 | 9 26E-219 |
| LRRN3 | 0 171755725 | 0 097404579 | 8 36E-220 | -0 29557075 | 0 | 9 44E-219 |
| MMP23B | 0 052216786 | 0 097363251 | 1 28E-219 | -0 42792817 | 0 | 1 45E-218 |
| MLLT10 | 0 608859531 | 0 097324634 | 1 91E-219 | 0 183129409 | 0 | 2 15E-218 |
| FBLN1 | 0 283832404 | 0 09732349 | 1 93E-219 | 0 28942565 | 0 | 2 18E-218 |
| SCMH1 | 0 65963907 | 0 097273011 | 3 26E-219 | 0 123272864 | 0 | 3 66E-218 |
| BACH2 | 0 466086692 | 0 097271437 | 3 31E-219 | -0 03643026 | 3 38E-32 | 3 71E-218 |
| SESN3 | 0 464535227 | 0 09726555 | 3 52E-219 | -0 03550458 | 1 15E-30 | 3 94E-218 |
| KAZN | 0 663998401 | 0 097240756 | 4 55E-219 | 0 086677643 | 2 56E-174 | 5 08E-218 |
| FUT10 | 0 220136681 | 0 097212461 | 6 09E-219 | -0 24352513 | 0 | 6 80E-218 |
| COLQ | 0 14313453 | 0 097097633 | 1 99E-218 | -0 31751479 | 0 | 2 22E-217 |
| FANCL | 0 362519274 | 0 096984705 | 6 37E-218 | 0 249549329 | 0 | 7 09E-217 |
| TPST1 | 0 626096971 | 0 096975013 | 7 04E-218 | 0 13172234 | 0 | 7 82E-217 |
| RBM6 | 0 776236889 | 0 096861774 | 2 26E-217 | 0 114247961 | 3 96E-302 | 2 51E-216 |
| MTA3 | 0 552587996 | 0 096850345 | 2 54E-217 | 0 177849108 | 0 | 2 81E-216 |
| ARHGAP44 | 0 145533114 | 0 096843781 | 2 72E-217 | -0 31466908 | 0 | 3 01E-216 |
| DAPK1 | 0 172279226 | 0 096779621 | 5 25E-217 | 0 42878586 | 0 | 5 81E-216 |
| ZNF280D | 0 57190992 | 0 096766458 | 6 01E-217 | -0 00564553 | 0 067265795 | 6 64E-216 |
| GNA14-AS1 | 0 076916487 | 0 096747879 | 7 27E-217 | -0 40049223 | 0 | 8 03E-216 |
| TMCC1 | 0 567683844 | 0 096703198 | 1 15E-216 | -0 00501529 | 0 104032287 | 1 27E-215 |
| DOK5 | 0 228645942 | 0 096684207 | 1 40E-216 | -0 22558615 | 0 | 1 54E-215 |
| CEP162 | 0 303525537 | 0 096630264 | 2 43E-216 | 0 354050069 | 0 | 2 68E-215 |
| PITPNC1 | 0 33021454 | 0 096618171 | 2 75E-216 | 0 292375505 | 0 | 3 03E-215 |
| NELL2 | 0 077240106 | 0 096585369 | 3 86E-216 | 0 471218014 | 0 | 4 23E-215 |
| FSIP2 | 0 177047838 | 0 096579199 | 4 11E-216 | -0 29541987 | 0 | 4 50E-215 |
| HHAT | 0 247301593 | 0 096546341 | 5 75E-216 | -0 21239596 | 0 | 6 30E-215 |
| IPCEF1 | 0 119557975 | 0 096459761 | 1 40E-215 | -0 348916 | 0 | 1 53E-214 |
| MFAP2 | 0 160048352 | 0 096358962 | 3 92E-215 | 0 393739851 | 0 | 4 28E-214 |
| TRANK1 | 0 258009556 | 0 096358144 | 3 95E-215 | -0 21626209 | 0 | 4 31E-214 |
| BTRC | 0 545744418 | 0 096187215 | 2 27E-214 | -0 0216384 | 2 31E-12 | 2 47E-213 |
| NFIA-AS2 | 0 103453199 | 0 096132128 | 3 98E-214 | 0 453369157 | 0 | 4 33E-213 |
| ATXN1 | 0 68324418 | 0 096131531 | 4 00E-214 | 0 026051226 | 3 03E-17 | 4 35E-213 |
| CHSY3 | 0 354228931 | 0 096063957 | 7 97E-214 | 0 22686114 | 0 | 8 66E-213 |
| U2SURP | 0 641097638 | 0 095989788 | 1 70E-213 | 0 159973738 | 0 | 1 84E-212 |
| CEP57 | 0 327701738 | 0 095949502 | 2 56E-213 | 0 30292194 | 0 | 2 77E-212 |
| IMMP2L | 0 792484438 | 0 095941111 | 2 79E-213 | 0 09972502 | 2 37E-230 | 3 02E-212 |
| POLE | 0 173564181 | 0 095933208 | 3 02E-213 | 0 34608982 | 0 | 3 27E-212 |
| TMLHE | 0 284470122 | 0 095900785 | 4 20E-213 | -0 1884114 | 0 | 4 54E-212 |

| | | | | | | |
|---|---|---|---|---|---|---|
| NFATC3 | 0 437113324 | 0 095876968 | 5 35E-213 | 0 232515833 | 0 | 5 77E-212 |
| COL6A6 | 0 098484704 | 0 095870545 | 5 72E-213 | 0 456186249 | 0 | 6 15E-212 |
| MSI2 | 0 625021416 | 0 095868794 | 5 82E-213 | 0 086767881 | 1 12E-174 | 6 26E-212 |
| CEMIP | 0 189621366 | 0 095853508 | 6 80E-213 | -0 27329192 | 0 | 7 30E-212 |
| SIMC1 | 0 367525842 | 0 095837575 | 7 99E-213 | 0 234699158 | 0 | 8 57E-212 |
| MFSD11 | 0 39718452 | 0 095829948 | 8 64E-213 | 0 242145631 | 0 | 9 26E-212 |
| CNTLN | 0 593021264 | 0 095829557 | 8 67E-213 | 0 127314436 | 0 | 9 28E-212 |
| PLA2G4A | 0 137575908 | 0 095731492 | 2 35E-212 | -0 33351482 | 0 | 2 51E-211 |
| DZIP3 | 0 276960271 | 0 095694766 | 3 41E-212 | 0 356412248 | 0 | 3 64E-211 |
| FXYD6 | 0 089699416 | 0 095595061 | 9 39E-212 | 0 469012307 | 0 | 1 00E-210 |
| ALMS1 | 0 407359464 | 0 095587466 | 1 01E-211 | 0 245615631 | 0 | 1 08E-210 |
| TDRD3 | 0 489796501 | 0 095542717 | 1 60E-211 | -0 0324918 | 6 00E-26 | 1 70E-210 |
| FLT1 | 0 116245645 | 0 09546096 | 3 66E-211 | 0 45158847 | 0 | 3 89E-210 |
| IRAK3 | 0 332832042 | 0 095427454 | 5 14E-211 | 0 296028825 | 0 | 5 45E-210 |
| RIC3 | 0 131293903 | 0 095422743 | 5 39E-211 | -0 33251444 | 0 | 5 71E-210 |
| AGO3 | 0 59666673 | 0 095369688 | 9 22E-211 | -0 00357228 | 0 246912207 | 9 76E-210 |
| BCL2L11 | 0 151681864 | 0 095330846 | 1 37E-210 | 0 385733803 | 0 | 1 44E-209 |
| RANBP3L | 0 094115855 | 0 095280338 | 2 28E-210 | 0 462493297 | 0 | 2 40E-209 |
| PPM1H | 0 115046354 | 0 095250276 | 3 09E-210 | -0 35764469 | 0 | 3 25E-209 |
| WNK1 | 0 651672346 | 0 095165595 | 7 26E-210 | 0 103146957 | 2 28E-246 | 7 65E-209 |
| ASS1 | 0 057680227 | 0 095088522 | 1 58E-209 | -0 42190969 | 0 | 1 66E-208 |
| DENND2A | 0 509223125 | 0 095041489 | 2 54E-209 | -0 04679076 | 5 21E-52 | 2 67E-208 |
| ARID5B | 0 539157831 | 0 095029388 | 2 87E-209 | -0 01401904 | 5 51E-06 | 3 01E-208 |
| HOXD3 | 0 179208467 | 0 095028326 | 2 90E-209 | 0 39358676 | 0 | 3 04E-208 |
| MLLT6 | 0 253869144 | 0 095005091 | 3 67E-209 | -0 21039069 | 0 | 3 84E-208 |
| FSIP2-AS1 | 0 191486932 | 0 095000066 | 3 86E-209 | -0 26987891 | 0 | 4 03E-208 |
| MIR646HG | 0 198263882 | 0 095000055 | 3 86E-209 | -0 23638899 | 0 | 4 03E-208 |
| ARHGAP31 | 0 313034208 | 0 094965307 | 5 48E-209 | -0 15299201 | 0 | 5 72E-208 |
| NOTCH3 | 0 148693153 | 0 094941048 | 6 99E-209 | -0 32581352 | 0 | 7 29E-208 |
| COL5A2 | 0 770050066 | 0 094870393 | 1 42E-208 | 0 028587004 | 1 90E-20 | 1 48E-207 |
| CPVL | 0 148740744 | 0 094807873 | 2 67E-208 | -0 32853927 | 0 | 2 78E-207 |
| KIF9-AS1 | 0 313367345 | 0 094769692 | 3 92E-208 | 0 298811922 | 0 | 4 08E-207 |
| CHODL | 0 145323714 | 0 094715438 | 6 77E-208 | 0 428275225 | 0 | 7 03E-207 |
| ACBD6 | 0 592983191 | 0 094659758 | 1 18E-207 | 0 169119646 | 0 | 1 23E-206 |
| GLIS1 | 0 059974111 | 0 094537099 | 4 06E-207 | 0 471852479 | 0 | 4 21E-206 |
| CDK8 | 0 376625231 | 0 094510987 | 5 27E-207 | 0 286239929 | 0 | 5 46E-206 |
| EPHA3 | 0 412946641 | 0 094462739 | 8 56E-207 | -0 13148579 | 0 | 8 85E-206 |
| HOMER1 | 0 288972226 | 0 094426485 | 1 23E-206 | -0 19391581 | 0 | 1 27E-205 |
| SCFD1 | 0 64613276 | 0 094423112 | 1 27E-206 | 0 030514547 | 4 47E-23 | 1 31E-205 |
| ORC3 | 0 33042394 | 0 094421685 | 1 29E-206 | 0 306970456 | 0 | 1 33E-205 |
| CHRNA7 | 0 052083532 | 0 094350799 | 2 63E-206 | 0 486298157 | 0 | 2 70E-205 |
| POLE2 | 0 167662904 | 0 094340429 | 2 91E-206 | 0 376888919 | 0 | 2 99E-205 |
| STAU2 | 0 450296016 | 0 094338327 | 2 98E-206 | 0 217516146 | 0 | 3 05E-205 |
| VWA8 | 0 546981782 | 0 09431897 | 3 61E-206 | -0 00554303 | 0 072388182 | 3 70E-205 |

| | | | | | | |
|---|---|---|---|---|---|---|
| C8orf34 | 0 088366869 | 0 094194454 | 1 25E-205 | 0 460615751 | 0 | 1 28E-204 |
| PRR5 | 0 126744208 | 0 094193108 | 1 27E-205 | -0 33245336 | 0 | 1 30E-204 |
| ANKRD30BL | 0 091564981 | 0 094188608 | 1 33E-205 | 0 473092691 | 0 | 1 36E-204 |
| BPTF | 0 810940207 | 0 094166943 | 1 65E-205 | 0 07287407 | 1 12E-123 | 1 68E-204 |
| EBF2 | 0 5577183 | 0 094141041 | 2 14E-205 | -0 00558835 | 0 070085363 | 2 18E-204 |
| AKAP8L | 0 320049114 | 0 094084919 | 3 75E-205 | 0 314613271 | 0 | 3 81E-204 |
| PCM1 | 0 693304906 | 0 094072074 | 4 26E-205 | 0 158369664 | 0 | 4 33E-204 |
| HMGB1 | 0 708181835 | 0 094038126 | 5 98E-205 | 0 146984931 | 0 | 6 07E-204 |
| SYBU | 0 072795111 | 0 094016045 | 7 45E-205 | 0 474079381 | 0 | 7 56E-204 |
| PAPPA | 0 222554301 | 0 093961693 | 1 28E-204 | -0 2532135 | 0 | 1 30E-203 |
| CCM2 | 0 426510061 | 0 093885839 | 2 73E-204 | 0 25060619 | 0 | 2 76E-203 |
| TARBP1 | 0 318592831 | 0 093812131 | 5 69E-204 | -0 15074562 | 0 | 5 74E-203 |
| TBC1D5 | 0 792389256 | 0 093756668 | 9 87E-204 | 0 044145371 | 1 74E-46 | 9 96E-203 |
| RAB2A | 0 684405399 | 0 093720289 | 1 42E-203 | 0 128477723 | 0 | 1 43E-202 |
| SULT1C4 | 0 138622908 | 0 093686263 | 1 99E-203 | -0 32338802 | 0 | 2 00E-202 |
| MMS22L | 0 265281453 | 0 093663122 | 2 50E-203 | 0 298017988 | 0 | 2 52E-202 |
| NT5DC1 | 0 227208696 | 0 093624047 | 3 69E-203 | 0 372466267 | 0 | 3 70E-202 |
| AGBL4 | 0 240715006 | 0 093568329 | 6 42E-203 | 0 394610735 | 0 | 6 43E-202 |
| TTN-AS1 | 0 313462527 | 0 093536897 | 8 77E-203 | 0 295457617 | 0 | 8 78E-202 |
| PRKCE | 0 667310731 | 0 093409698 | 3 10E-202 | 0 157931086 | 0 | 3 09E-201 |
| ERBB4 | 0 205145533 | 0 093408156 | 3 14E-202 | 0 422924846 | 0 | 3 13E-201 |
| MAF | 0 151234509 | 0 09339916 | 3 44E-202 | 0 423706147 | 0 | 3 42E-201 |
| ANGPTL6 | 0 018227332 | 0 093393259 | 3 64E-202 | 0 501668689 | 0 | 3 62E-201 |
| SEMA5B | 0 04294607 | 0 093354632 | 5 34E-202 | 0 489984193 | 0 | 5 30E-201 |
| GRIA4 | 0 068740363 | 0 093315157 | 7 90E-202 | -0 40074261 | 0 | 7 83E-201 |
| BGN | 0 194780225 | 0 093306506 | 8 60E-202 | -0 26102837 | 0 | 8 52E-201 |
| COP1 | 0 716167596 | 0 093259027 | 1 38E-201 | 0 132317855 | 0 | 1 36E-200 |
| WDR27 | 0 385572329 | 0 093140585 | 4 44E-201 | -0 11568197 | | 4 39E-200 |
| PTPRT | 0 103748263 | 0 093079716 | 8 10E-201 | 0 431845209 | 0 | 8 00E-200 |
| RAVER2 | 0 182654052 | 0 093000467 | 1 77E-200 | 0 380758439 | 0 | 1 75E-199 |
| B3GLCT | 0 15701205 | 0 092994399 | 1 88E-200 | 0 418139393 | 0 | 1 85E-199 |
| RNF175 | 0 086910586 | 0 09284701 | 8 04E-200 | 0 465643118 | 0 | 7 91E-199 |
| TPT1-AS1 | 0 553111496 | 0 092824613 | 1 00E-199 | 0 006634547 | 0 031517929 | 9 86E-199 |
| CMYA5 | 0 332879633 | 0 092727997 | 2 59E-199 | -0 12978107 | 0 | 2 54E-198 |
| MGAT5 | 0 551103158 | 0 092720305 | 2 80E-199 | -0 01759832 | 1 17E-08 | 2 74E-198 |
| TMEM87B | 0 368782243 | 0 092651571 | 5 50E-199 | 0 267176014 | 0 | 5 38E-198 |
| ELAVL2 | 0 042803297 | 0 092572041 | 1 20E-198 | 0 484008824 | 0 | 1 17E-197 |
| RALGAPA1 | 0 692543451 | 0 092519777 | 2 01E-198 | 0 053398987 | 3 29E-67 | 1 96E-197 |
| TGIF2 | 0 165378538 | 0 092480843 | 2 94E-198 | 0 408327529 | 0 | 2 87E-197 |
| RBM5 | 0 567474444 | 0 092441425 | 4 33E-198 | -0 01358515 | 1 06E-05 | 4 22E-197 |
| COL24A1 | 0 268355828 | 0 092338093 | 1 19E-197 | 0 320633354 | 0 | 1 16E-196 |
| PTK2 | 0 869381889 | 0 092329467 | 1 30E-197 | 0 064999845 | 9 70E-99 | 1 26E-196 |
| DYRK1A | 0 644248158 | 0 092270202 | 2 32E-197 | 0 141379318 | 0 | 2 25E-196 |
| GATAD2B | 0 560269174 | 0 092204258 | 4 42E-197 | 0 170287639 | 0 | 4 29E-196 |

| | | | | | | |
|---|---|---|---|---|---|---|
| HULC | 0 244560355 | 0 092010641 | 2 93E-196 | -0 20013541 | 0 | 2 83E-195 |
| PIBF1 | 0 653871048 | 0 091863406 | 1 23E-195 | 0 021702457 | 1 99E-12 | 1 19E-194 |
| LINC01118 | 0 112010051 | 0 091810404 | 2 06E-195 | -0 34876087 | 0 | 1 99E-194 |
| TUT4 | 0 68981173 | 0 091741383 | 4 04E-195 | 0 026520135 | 8 15E-18 | 3 89E-194 |
| PRKN | 0 540966287 | 0 091677195 | 7 54E-195 | 0 022302771 | 4 83E-13 | 7 26E-194 |
| USP49 | 0 209514382 | 0 091663183 | 8 64E-195 | 0 370026821 | 0 | 8 31E-194 |
| MIA2 | 0 359644781 | 0 091626632 | 1 23E-194 | -0 12836803 | 0 | 1 18E-193 |
| DGKH | 0 382107708 | 0 091580659 | 1 93E-194 | 0 263716612 | 0 | 1 85E-193 |
| ZHX2 | 0 396004264 | 0 091530776 | 3 13E-194 | -0 08639608 | 3 38E-173 | 3 00E-193 |
| EFNA5 | 0 471131332 | 0 091490439 | 4 63E-194 | 0 177636215 | 0 | 4 43E-193 |
| DYNC2H1 | 0 548599874 | 0 091470922 | 5 59E-194 | 0 001131429 | 0 713821179 | 5 34E-193 |
| KAT6A | 0 618453865 | 0 091422113 | 8 98E-194 | 0 163510721 | 0 | 8 57E-193 |
| ARFGEF1 | 0 68487179 | 0 091419485 | 9 21E-194 | 0 123093536 | 0 | 8 78E-193 |
| PHIP | 0 6159601 | 0 091397144 | 1 14E-193 | 0 172378256 | 0 | 1 09E-192 |
| CNOT2 | 0 673373817 | 0 091396182 | 1 15E-193 | 0 12444617 | 0 | 1 10E-192 |
| STK39 | 0 626287335 | 0 091373265 | 1 44E-193 | 0 120766191 | 0 | 1 37E-192 |
| SNX13 | 0 584112238 | 0 091325731 | 2 28E-193 | 0 004928488 | 0 110159919 | 2 17E-192 |
| SCAF8 | 0 596638176 | 0 091293468 | 3 12E-193 | 0 169661563 | 0 | 2 96E-192 |
| SMCHD1 | 0 679855704 | 0 091244802 | 5 00E-193 | 0 136675203 | 0 | 4 74E-192 |
| FTX | 0 920142392 | 0 091222469 | 6 21E-193 | 0 10648977 | 1 51E-262 | 5 87E-192 |
| PDE1C | 0 257876302 | 0 091173224 | 1 00E-192 | -0 21683771 | 0 | 9 44E-192 |
| TMEM135 | 0 476937427 | 0 091127635 | 1 55E-192 | -0 04999393 | 3 98E-59 | 1 46E-191 |
| PCOLCE2 | 0 121947041 | 0 091118584 | 1 70E-192 | 0 449776451 | 0 | 1 60E-191 |
| SPATS2 | 0 541689669 | 0 091098599 | 2 06E-192 | 0 160783063 | 0 | 1 94E-191 |
| TCF25 | 0 621937523 | 0 091095125 | 2 13E-192 | 0 006221785 | 0 043728975 | 2 00E-191 |
| KDM5B | 0 434077021 | 0 091093037 | 2 17E-192 | -0 05126607 | 4 39E-62 | 2 04E-191 |
| LRTM1 | 0 021301708 | 0 091086261 | 2 32E-192 | 0 499157452 | 0 | 2 17E-191 |
| LINC02224 | 0 06107822 | 0 09103435 | 3 82E-192 | -0 4182483 | 0 | 3 59E-191 |
| SH3D21 | 0 080028935 | 0 090952413 | 8 43E-192 | -0 39407724 | 0 | 7 90E-191 |
| UMAD1 | 0 493146904 | 0 090917866 | 1 18E-191 | -0 04869221 | 3 54E-56 | 1 10E-190 |
| PATJ | 0 300593935 | 0 090903634 | 1 35E-191 | -0 17277309 | 0 | 1 26E-190 |
| SLC20A2 | 0 452152063 | 0 090901584 | 1 38E-191 | -0 06562916 | 1 26E-100 | 1 28E-190 |
| SLC27A6 | 0 077792161 | 0 090836482 | 2 58E-191 | -0 39480563 | 0 | 2 40E-190 |
| CCNJL | 0 071005692 | 0 090834828 | 2 62E-191 | 0 466766858 | 0 | 2 44E-190 |
| TMEM260 | 0 319487541 | 0 090832197 | 2 69E-191 | -0 16319602 | 0 | 2 50E-190 |
| DNASE1 | 0 272191658 | 0 090813458 | 3 22E-191 | 0 321935389 | 0 | 2 99E-190 |
| ZNF33A | 0 277312444 | 0 09079491 | 3 85E-191 | -0 19582279 | 0 | 3 57E-190 |
| NPW | 0 080333517 | 0 090668399 | 1 30E-190 | 0 454821324 | 0 | 1 20E-189 |
| POU2F1 | 0 565437551 | 0 090557752 | 3 76E-190 | 0 192512448 | 0 | 3 48E-189 |
| HS6ST2 | 0 08226571 | 0 09049255 | 7 04E-190 | 0 4647322 | 0 | 6 51E-189 |
| CCDC57 | 0 398574175 | 0 090463435 | 9 31E-190 | 0 264938332 | 0 | 8 60E-189 |
| PHYHIPL | 0 043450534 | 0 090335534 | 3 17E-189 | 0 48781131 | 0 | 2 93E-188 |
| MYO10 | 0 404313643 | 0 090329946 | 3 35E-189 | 0 204092845 | 0 | 3 09E-188 |
| MBD5 | 0 828120538 | 0 09028568 | 5 11E-189 | 0 04560499 | 1 71E-49 | 4 71E-188 |

| | | | | | | |
|---|---|---|---|---|---|---|
| ATG5 | 0 393443871 | 0 090267912 | 6 06E-189 | 0 273022989 | 0 | 5 58E-188 |
| STX6 | 0 260446213 | 0 090262349 | 6 39E-189 | -0 22282042 | 0 | 5 88E-188 |
| HMGN1 | 0 408530201 | 0 090181995 | 1 38E-188 | 0 244406089 | 0 | 1 27E-187 |
| GALNTL6 | 0 085930213 | 0 090173453 | 1 50E-188 | -0 38005047 | 0 | 1 37E-187 |
| GRIK1 | 0 167986522 | 0 090131254 | 2 24E-188 | 0 389823385 | 0 | 2 05E-187 |
| FGF10-AS1 | 0 061049666 | 0 090104345 | 2 90E-188 | -0 4198207 | 0 | 2 65E-187 |
| GABRG3 | 0 069539891 | 0 090101911 | 2 97E-188 | 0 456589939 | 0 | 2 71E-187 |
| BCL2L14 | 0 317288839 | 0 090092657 | 3 24E-188 | 0 277748917 | 0 | 2 96E-187 |
| ZNF821 | 0 125563953 | 0 089916237 | 1 74E-187 | -0 34213811 | 0 | 1 59E-186 |
| SHOC1 | 0 325779064 | 0 089908152 | 1 88E-187 | -0 18298462 | 0 | 1 72E-186 |
| LHFPL6 | 0 651843673 | 0 089855738 | 3 11E-187 | 0 030837865 | 1 56E-23 | 2 83E-186 |
| MYLIP | 0 192924178 | 0 089843985 | 3 47E-187 | -0 27196298 | 0 | 3 16E-186 |
| CALCRL | 0 297633778 | 0 089843737 | 3 48E-187 | -0 18082192 | 0 | 3 17E-186 |
| ASH2L | 0 208172317 | 0 089788859 | 5 87E-187 | 0 374151029 | 0 | 5 34E-186 |
| ENOSF1 | 0 395614018 | 0 089751828 | 8 35E-187 | 0 275983293 | 0 | 7 58E-186 |
| CADM1 | 0 339199711 | 0 089733325 | 9 96E-187 | -0 14843569 | 0 | 9 04E-186 |
| NINJ2 | 0 085130685 | 0 089700506 | 1 36E-186 | -0 39855769 | 0 | 1 23E-185 |
| LINC01301 | 0 116788182 | 0 089679168 | 1 67E-186 | 0 446125257 | 0 | 1 51E-185 |
| PAX3 | 0 243846491 | 0 089670987 | 1 80E-186 | 0 279682563 | 0 | 1 63E-185 |
| THADA | 0 620062439 | 0 08965648 | 2 07E-186 | 0 000297379 | 0 923211543 | 1 87E-185 |
| LRRC17 | 0 237012431 | 0 089644941 | 2 31E-186 | 0 320764458 | 0 | 2 08E-185 |
| MAST4 | 0 510641336 | 0 089622871 | 2 85E-186 | 0 129288072 | 0 | 2 56E-185 |
| ZNF148 | 0 610287259 | 0 089588886 | 3 93E-186 | 0 163952113 | 0 | 3 54E-185 |
| CLEC16A | 0 431116864 | 0 089463634 | 1 29E-185 | -0 09400122 | 8 64E-205 | 1 16E-184 |
| SPATA16 | 0 027878776 | 0 089453853 | 1 42E-185 | 0 501275466 | 0 | 1 27E-184 |
| KLHL23 | 0 202194894 | 0 089435346 | 1 69E-185 | 0 359859537 | 0 | 1 51E-184 |
| CEP126 | 0 285126877 | 0 089431982 | 1 74E-185 | -0 18199359 | 0 | 1 56E-184 |
| PHF21B | 0 111800651 | 0 089430319 | 1 77E-185 | -0 35614354 | 0 | 1 58E-184 |
| FGF2 | 0 278387999 | 0 089380174 | 2 85E-185 | 0 325390522 | 0 | 2 54E-184 |
| ROBO3 | 0 077135406 | 0 089328679 | 4 64E-185 | 0 470055058 | 0 | 4 14E-184 |
| ZNF721 | 0 497382498 | 0 089324853 | 4 81E-185 | 0 22355181 | 0 | 4 29E-184 |
| SMARCA2 | 0 504007158 | 0 089318978 | 5 09E-185 | 0 192799929 | 0 | 4 53E-184 |
| KIRREL1 | 0 427404771 | 0 089238779 | 1 09E-184 | -0 08954817 | 5 79E-186 | 9 67E-184 |
| BAZ2B | 0 723905884 | 0 089229721 | 1 18E-184 | 0 050618765 | 1 43E-60 | 1 05E-183 |
| NSUN6 | 0 335106889 | 0 089199936 | 1 57E-184 | 0 299732698 | 0 | 1 39E-183 |
| SNTB1 | 0 300184653 | 0 089172662 | 2 03E-184 | -0 16773528 | 0 | 1 80E-183 |
| WLS | 0 449001542 | 0 089155737 | 2 38E-184 | -0 04827402 | 3 02E-55 | 2 11E-183 |
| RHOJ | 0 409472502 | 0 088918676 | 2 24E-183 | -0 06333625 | 7 74E-94 | 1 97E-182 |
| KCTD12 | 0 225343131 | 0 088885188 | 3 06E-183 | -0 25867585 | 0 | 2 70E-182 |
| SRGAP2C | 0 359416345 | 0 088676377 | 2 19E-182 | 0 276910252 | 0 | 1 92E-181 |
| CNTN6 | 0 039576631 | 0 088669883 | 2 33E-182 | 0 489080662 | 0 | 2 04E-181 |
| TTC7B | 0 350079001 | 0 088665226 | 2 43E-182 | 0 262557859 | 0 | 2 13E-181 |
| PTH2R | 0 098722659 | 0 088663635 | 2 47E-182 | 0 449626251 | 0 | 2 16E-181 |
| ZMYM4 | 0 623374769 | 0 088585832 | 5 12E-182 | 0 006872957 | 0 025897192 | 4 48E-181 |

| | | | | | | |
|---|---|---|---|---|---|---|
| ZSWIM5 | 0 253022025 | 0 088486262 | 1 30E-181 | -0 20672567 | 0 | 1 14E-180 |
| MBD6 | 0 164436238 | 0 088459414 | 1 68E-181 | 0 405965901 | 0 | 1 46E-180 |
| PDZD2 | 0 457710685 | 0 088450837 | 1 82E-181 | -0 07068189 | 1 91E-116 | 1 58E-180 |
| MCPH1 | 0 386162456 | 0 088443632 | 1 95E-181 | 0 278236613 | 0 | 1 69E-180 |
| ARL17B | 0 346290762 | 0 088420125 | 2 43E-181 | 0 221178377 | 0 | 2 10E-180 |
| TSC22D1 | 0 528573604 | 0 08841278 | 2 60E-181 | -0 01980808 | 1 35E-10 | 2 25E-180 |
| HS3ST3A1 | 0 157849651 | 0 088372168 | 3 80E-181 | 0 445964983 | 0 | 3 29E-180 |
| C12orf60 | 0 069197236 | 0 08832008 | 6 19E-181 | -0 40584881 | 0 | 5 35E-180 |
| LRRN2 | 0 048495174 | 0 088306376 | 7 04E-181 | -0 42956441 | 0 | 6 07E-180 |
| PDE7A | 0 465506082 | 0 088290358 | 8 18E-181 | 0 191797759 | 0 | 7 04E-180 |
| PHTF2 | 0 426871752 | 0 088218383 | 1 60E-180 | 0 204357129 | 0 | 1 38E-179 |
| GTDC1 | 0 463526299 | 0 088202898 | 1 85E-180 | -0 06339587 | 5 19E-94 | 1 59E-179 |
| BMPR2 | 0 666197103 | 0 088084519 | 5 60E-180 | -0 00162211 | 0 599045812 | 4 81E-179 |
| SLC40A1 | 0 156983495 | 0 088057484 | 7 21E-180 | -0 31666838 | 0 | 6 19E-179 |
| ZNHIT6 | 0 340732139 | 0 088057273 | 7 22E-180 | -0 14729669 | 0 | 6 20E-179 |
| CENPP | 0 520311816 | 0 088025907 | 9 68E-180 | 0 125095101 | 0 | 8 30E-179 |
| EEF2K | 0 235794103 | 0 087970525 | 1 62E-179 | -0 22935465 | 0 | 1 39E-178 |
| KCND2 | 0 080761836 | 0 087852132 | 4 89E-179 | 0 46558145 | 0 | 4 18E-178 |
| YAP1 | 0 633663932 | 0 08783064 | 5 98E-179 | 0 028065769 | 9 14E-20 | 5 11E-178 |
| ID1 | 0 109278331 | 0 087803465 | 7 70E-179 | 0 440755978 | 0 | 6 57E-178 |
| PUM2 | 0 711484647 | 0 08772142 | 1 65E-178 | 0 108345261 | 9 36E-272 | 1 41E-177 |
| ATRN | 0 429498772 | 0 087710026 | 1 84E-178 | -0 09227322 | 2 25E-197 | 1 56E-177 |
| WNT5A | 0 057775409 | 0 087663806 | 2 82E-178 | 0 479756376 | 0 | 2 40E-177 |
| CYP46A1 | 0 113218861 | 0 087601732 | 5 02E-178 | -0 35843148 | 0 | 4 26E-177 |
| ADAMTS5 | 0 091098589 | 0 087596069 | 5 29E-178 | -0 38690993 | 0 | 4 49E-177 |
| COL8A2 | 0 071548229 | 0 087406757 | 3 06E-177 | -0 39661625 | 0 | 2 60E-176 |
| LINC01473 | 0 678903885 | 0 087394962 | 3 42E-177 | 0 044727795 | 1 13E-47 | 2 89E-176 |
| HECW1 | 0 129580629 | 0 087392656 | 3 49E-177 | -0 34707652 | 0 | 2 95E-176 |
| LINC00240 | 0 133663932 | 0 087368149 | 4 38E-177 | -0 34096541 | 0 | 3 70E-176 |
| PLCL2 | 0 37147589 | 0 087334126 | 6 00E-177 | 0 224800175 | 0 | 5 06E-176 |
| RFLNB | 0 083940911 | 0 087330057 | 6 23E-177 | -0 39353263 | 0 | 5 25E-176 |
| TMEM132C | 0 083112829 | 0 087300471 | 8 20E-177 | -0 37842525 | 0 | 6 90E-176 |
| TSPAN9 | 0 290428509 | 0 087264289 | 1 15E-176 | 0 289526771 | 0 | 9 63E-176 |
| SEPTIN11 | 0 574660677 | 0 0872525 | 1 28E-176 | 0 131793963 | 0 | 1 07E-175 |
| LINC00960 | 0 191686814 | 0 087205204 | 1 98E-176 | -0 28822187 | 0 | 1 66E-175 |
| RBPMS | 0 477908283 | 0 087056728 | 7 79E-176 | -0 0391271 | 6 97E-37 | 6 54E-175 |
| RUNX2 | 0 610182559 | 0 087034945 | 9 53E-176 | -0 01495575 | 1 25E-06 | 7 98E-175 |
| PRKDC | 0 610087377 | 0 087031165 | 9 87E-176 | 0 142300862 | 0 | 8 26E-175 |
| ST3GAL5 | 0 187375074 | 0 087015877 | 1 14E-175 | -0 25845304 | 0 | 9 50E-175 |
| SMC4 | 0 340456112 | 0 086918828 | 2 78E-175 | 0 195590319 | 0 | 2 32E-174 |
| DDX60L | 0 300898517 | 0 086904801 | 3 16E-175 | 0 305567052 | 0 | 2 64E-174 |
| ZEB1 | 0 804106147 | 0 086820511 | 6 87E-175 | 0 07752613 | 9 36E-140 | 5 71E-174 |
| HELLS | 0 203299004 | 0 086817553 | 7 06E-175 | 0 2800453 | 0 | 5 86E-174 |
| GNG12-AS1 | 0 314595191 | 0 086785993 | 9 44E-175 | -0 1782204 | 0 | 7 83E-174 |

| | | | | | | |
|---|---|---|---|---|---|---|
| RTTN | 0 29158021 | 0 086773381 | 1 06E-174 | 0 288834767 | 0 | 8 78E-174 |
| NPY1R | 0 030724715 | 0 086764807 | 1 15E-174 | 0 492829044 | 0 | 9 49E-174 |
| HDAC8 | 0 57059641 | 0 086751022 | 1 30E-174 | 0 187445336 | 0 | 1 08E-173 |
| PPARD | 0 266499781 | 0 086711283 | 1 88E-174 | -0 21551755 | 0 | 1 55E-173 |
| CPXM1 | 0 208762445 | 0 086706593 | 1 96E-174 | -0 26266997 | 0 | 1 62E-173 |
| RNGTT | 0 468589975 | 0 086705858 | 1 97E-174 | 0 240243076 | 0 | 1 63E-173 |
| ISYNA1 | 0 110553768 | 0 08664328 | 3 51E-174 | 0 449089474 | 0 | 2 89E-173 |
| STAC | 0 077468542 | 0 086598289 | 5 30E-174 | 0 4542159 | 0 | 4 36E-173 |
| ITGA10 | 0 320115741 | 0 086563607 | 7 28E-174 | -0 15464152 | 0 | 5 98E-173 |
| FAM214A | 0 186889646 | 0 086551282 | 8 15E-174 | 0 356167917 | 0 | 6 69E-173 |
| GUCY1A1 | 0 101797034 | 0 086524354 | 1 04E-173 | -0 37973151 | 0 | 8 56E-173 |
| SGK1 | 0 192714778 | 0 086522117 | 1 07E-173 | 0 40707747 | 0 | 8 73E-173 |
| NFAT5 | 0 733366964 | 0 086521298 | 1 07E-173 | 0 046384154 | 3 86E-51 | 8 79E-173 |
| VPS54 | 0 604052845 | 0 086513536 | 1 15E-173 | -0 01354142 | 1 14E-05 | 9 43E-173 |
| SLX4IP | 0 307180522 | 0 086494716 | 1 37E-173 | 0 322335382 | 0 | 1 12E-172 |
| SRPK2 | 0 746997011 | 0 086494234 | 1 38E-173 | 0 087316519 | 7 07E-177 | 1 12E-172 |
| ADAM32 | 0 154784794 | 0 086490352 | 1 43E-173 | -0 32603836 | 0 | 1 16E-172 |
| OVCH1 | 0 041651596 | 0 086431974 | 2 43E-173 | -0 4386612 | 0 | 1 98E-172 |
| PID1 | 0 386257638 | 0 086415499 | 2 83E-173 | -0 09336021 | 5 05E-202 | 2 31E-172 |
| LINC02641 | 0 10381489 | 0 086384276 | 3 77E-173 | 0 455739185 | 0 | 3 07E-172 |
| SLC14A2 | 0 063115113 | 0 086338461 | 5 73E-173 | -0 41299678 | 0 | 4 66E-172 |
| EXOC4 | 0 80477242 | 0 08633103 | 6 13E-173 | 0 100064558 | 6 45E-232 | 4 98E-172 |
| SDCCAG8 | 0 621889932 | 0 086314904 | 7 11E-173 | 0 006313958 | 0 040701669 | 5 76E-172 |
| NFATC1 | 0 126829872 | 0 086269659 | 1 08E-172 | -0 33445622 | 0 | 8 71E-172 |
| EFR3B | 0 094610801 | 0 086201122 | 2 01E-172 | 0 451625474 | 0 | 1 63E-171 |
| FRMD4A | 0 700234147 | 0 086184065 | 2 35E-172 | 0 030967651 | 1 02E-23 | 1 90E-171 |
| CEP83 | 0 389760332 | 0 086171178 | 2 64E-172 | 0 240048801 | 0 | 2 13E-171 |
| TMLHE-AS1 | 0 247006529 | 0 086147759 | 3 27E-172 | 0 343373045 | 0 | 2 64E-171 |
| EIF4E3 | 0 117873256 | 0 086097473 | 5 18E-172 | -0 35420534 | 0 | 4 17E-171 |
| LEF1 | 0 254278426 | 0 086067508 | 6 81E-172 | -0 20401298 | 0 | 5 47E-171 |
| MDN1 | 0 410909749 | 0 086034654 | 9 19E-172 | -0 09369182 | 1 88E-203 | 7 38E-171 |
| PLAC1 | 0 028630713 | 0 085823863 | 6 27E-171 | 0 49337062 | 0 | 5 02E-170 |
| UBE2E2 | 0 776922198 | 0 085750048 | 1 23E-170 | 0 042949207 | 4 31E-44 | 9 82E-170 |
| CBFA2T2 | 0 594448992 | 0 085704892 | 1 85E-170 | -0 00788619 | 0 010582807 | 1 48E-169 |
| ZFAND3 | 0 829262721 | 0 085651661 | 3 00E-170 | 0 040268813 | 5 75E-39 | 2 40E-169 |
| SETD2 | 0 67572481 | 0 085618714 | 4 04E-170 | 0 128567984 | 0 | 3 23E-169 |
| HAPLN3 | 0 069378082 | 0 085592519 | 5 12E-170 | -0 40365904 | 0 | 4 09E-169 |
| SYNPO | 0 083940911 | 0 085480077 | 1 42E-169 | -0 38640699 | 0 | 1 13E-168 |
| NTRK3 | 0 210142582 | 0 085469183 | 1 57E-169 | -0 22582876 | 0 | 1 25E-168 |
| PKDCC | 0 278483181 | 0 085435501 | 2 13E-169 | -0 19886619 | 0 | 1 69E-168 |
| PTPRK | 0 571157983 | 0 085402462 | 2 87E-169 | 0 131935483 | 0 | 2 28E-168 |
| PLEKHG5 | 0 043650416 | 0 085297916 | 7 38E-169 | 0 493177002 | 0 | 5 86E-168 |
| FAM107B | 0 176048429 | 0 085288238 | 8 05E-169 | 0 387465657 | 0 | 6 39E-168 |
| RPS6KC1 | 0 579010489 | 0 085220382 | 1 49E-168 | -0 00193908 | 0 529667967 | 1 18E-167 |

| | | | | | | |
|---|---|---|---|---|---|---|
| OXCT1 | 0 279520664 | 0 085190765 | 1 94E-168 | 0 342988087 | 0 | 1 54E-167 |
| GATA6 | 0 08389332 | 0 08514424 | 2 96E-168 | 0 446030233 | 0 | 2 34E-167 |
| ATAD2B | 0 569102054 | 0 085089118 | 4 86E-168 | -0 02015179 | 6 47E-11 | 3 84E-167 |
| COPG2 | 0 270963812 | 0 084996038 | 1 12E-167 | 0 347764345 | 0 | 8 88E-167 |
| EXOC6B | 0 77791209 | 0 084962225 | 1 52E-167 | 0 071471818 | 5 03E-119 | 1 20E-166 |
| ASB3 | 0 596514439 | 0 084942559 | 1 82E-167 | 0 142547419 | 0 | 1 43E-166 |
| RYR2 | 0 093820792 | 0 084882432 | 3 13E-167 | 0 447430263 | 0 | 2 46E-166 |
| PHF3 | 0 53952904 | 0 084826789 | 5 16E-167 | -0 01585888 | 2 74E-07 | 4 06E-166 |
| RGS12 | 0 293569511 | 0 084817932 | 5 58E-167 | 0 292788146 | 0 | 4 39E-166 |
| DUOX1 | 0 178627858 | 0 084808145 | 6 10E-167 | 0 39336822 | 0 | 4 79E-166 |
| WNT5B | 0 218975462 | 0 084763024 | 9 14E-167 | 0 323616467 | 0 | 7 16E-166 |
| PDE8B | 0 157640251 | 0 084680079 | 1 93E-166 | -0 31952586 | 0 | 1 51E-165 |
| SUCO | 0 42788068 | 0 084674634 | 2 02E-166 | 0 224417313 | 0 | 1 58E-165 |
| EPHB3 | 0 086377568 | 0 084643698 | 2 67E-166 | -0 3854911 | 0 | 2 08E-165 |
| ADCY6 | 0 170023415 | 0 084544049 | 6 52E-166 | -0 28878216 | 0 | 5 09E-165 |
| STAC3 | 0 042451124 | 0 084538699 | 6 84E-166 | 0 482672916 | 0 | 5 33E-165 |
| DIP2A | 0 406093545 | 0 084523498 | 7 84E-166 | 0 215054999 | 0 | 6 10E-165 |
| FHIT | 0 499952409 | 0 084463844 | 1 34E-165 | -0 03806957 | 5 25E-35 | 1 04E-164 |
| PBX4 | 0 122613314 | 0 084413062 | 2 11E-165 | 0 408799162 | 0 | 1 64E-164 |
| SMAP1 | 0 382850127 | 0 084406324 | 2 24E-165 | -0 11057574 | 4 87E-283 | 1 74E-164 |
| CABLES1 | 0 128866764 | 0 084402547 | 2 31E-165 | -0 34624004 | 0 | 1 79E-164 |
| PARP16 | 0 146713369 | 0 084362282 | 3 32E-165 | 0 415748749 | 0 | 2 57E-164 |
| TYW1 | 0 475747654 | 0 084299894 | 5 79E-165 | 0 207141503 | 0 | 4 48E-164 |
| GMDS-DT | 0 561477984 | 0 084258912 | 8 35E-165 | 0 004934454 | 0 109729833 | 6 46E-164 |
| SYN3 | 0 21325503 | 0 084223623 | 1 14E-164 | 0 339882856 | 0 | 8 84E-164 |
| GNAL | 0 119557975 | 0 084213888 | 1 25E-164 | 0 44882559 | 0 | 9 64E-164 |
| RRBP1 | 0 512040509 | 0 08416165 | 1 99E-164 | -0 05222558 | 2 31E-64 | 1 53E-163 |
| HNMT | 0 130208829 | 0 084149802 | 2 21E-164 | 0 400924404 | 0 | 1 70E-163 |
| HDGFL3 | 0 355808951 | 0 084136361 | 2 49E-164 | 0 245487209 | 0 | 1 92E-163 |
| UBE2W | 0 466838629 | 0 084095201 | 3 59E-164 | 0 207108676 | 0 | 2 76E-163 |
| FBXW4 | 0 397336811 | 0 084056175 | 5 09E-164 | -0 10521424 | 2 60E-256 | 3 91E-163 |
| SSBP4 | 0 25115646 | 0 084048647 | 5 44E-164 | -0 23014438 | 0 | 4 18E-163 |
| ZNF713 | 0 167244103 | 0 084042643 | 5 74E-164 | 0 422066355 | 0 | 4 40E-163 |
| HECTD2 | 0 435209686 | 0 083993969 | 8 85E-164 | 0 236973813 | 0 | 6 78E-163 |
| ERICH1 | 0 349070073 | 0 083965383 | 1 14E-163 | -0 15015725 | 0 | 8 73E-163 |
| NEBL | 0 203736841 | 0 083963849 | 1 16E-163 | -0 2741985 | 0 | 8 84E-163 |
| HLF | 0 114370562 | 0 083926501 | 1 61E-163 | -0 35115728 | 0 | 1 23E-162 |
| EFCAB11 | 0 18234947 | 0 083904156 | 1 97E-163 | 0 358682515 | 0 | 1 50E-162 |
| HCG18 | 0 266366526 | 0 083898022 | 2 08E-163 | -0 19481878 | 0 | 1 58E-162 |
| CCDC125 | 0 212379357 | 0 083853254 | 3 09E-163 | -0 25885269 | 0 | 2 35E-162 |
| EPHA4 | 0 223534675 | 0 083832602 | 3 72E-163 | 0 30365938 | 0 | 2 82E-162 |
| LINC00882 | 0 404523044 | 0 083786181 | 5 61E-163 | -0 10468995 | 9 02E-254 | 4 26E-162 |
| GTF2IRD1 | 0 355218823 | 0 083771536 | 6 39E-163 | 0 254186308 | 0 | 4 85E-162 |
| RALY | 0 427347661 | 0 08367883 | 1 45E-162 | 0 222846454 | 0 | 1 10E-161 |

| | | | | | | |
|---|---|---|---|---|---|---|
| DIRC3 | 0 277474253 | 0 083668521 | 1 59E-162 | 0 332482825 | 0 | 1 21E-161 |
| TRMT9B | 0 210485237 | 0 083634411 | 2 16E-162 | -0 25521778 | 0 | 1 63E-161 |
| DEF6 | 0 058689155 | 0 083554175 | 4 39E-162 | 0 474611049 | 0 | 3 32E-161 |
| SESN1 | 0 37941406 | 0 083546536 | 4 70E-162 | -0 10990343 | 1 30E-279 | 3 55E-161 |
| ROR1-AS1 | 0 130246902 | 0 083526866 | 5 59E-162 | 0 428452177 | 0 | 4 22E-161 |
| MYO9A | 0 74101007 | 0 083435821 | 1 25E-161 | 0 046305989 | 5 67E-51 | 9 43E-161 |
| ADAM23 | 0 322447698 | 0 083428898 | 1 33E-161 | -0 1218301 | 0 | 1 00E-160 |
| PARVA | 0 507338524 | 0 083404129 | 1 65E-161 | -0 03126837 | 3 77E-24 | 1 24E-160 |
| VSNL1 | 0 04736251 | 0 083399279 | 1 73E-161 | 0 484049109 | 0 | 1 30E-160 |
| CAMSAP2 | 0 572423902 | 0 083340683 | 2 90E-161 | -0 02387861 | 9 88E-15 | 2 17E-160 |
| SHE | 0 083208011 | 0 083316222 | 3 60E-161 | -0 38779879 | 0 | 2 70E-160 |
| FNTB | 0 242333099 | 0 083277069 | 5 08E-161 | 0 363780332 | 0 | 3 81E-160 |
| ANKAR | 0 385991129 | 0 083226507 | 7 93E-161 | -0 13433922 | 0 | 5 94E-160 |
| RBM39 | 0 809493442 | 0 0832265 | 7 93E-161 | 0 041203177 | 1 02E-40 | 5 94E-160 |
| SH3BP2 | 0 129371228 | 0 083209048 | 9 25E-161 | 0 424820172 | 0 | 6 92E-160 |
| SEZ6L | 0 020111934 | 0 083189608 | 1 10E-160 | 0 500344795 | 0 | 8 20E-160 |
| NKD1 | 0 050493994 | 0 083160423 | 1 42E-160 | -0 4203467 | 0 | 1 06E-159 |
| MAD1L1 | 0 29378843 | 0 08315601 | 1 48E-160 | 0 312136982 | 0 | 1 10E-159 |
| TUBGCP3 | 0 316660638 | 0 083149892 | 1 56E-160 | 0 277336898 | 0 | 1 16E-159 |
| EDARADD | 0 068921208 | 0 083041533 | 4 05E-160 | 0 470507652 | 0 | 3 01E-159 |
| PPA2 | 0 397441511 | 0 082987306 | 6 52E-160 | -0 11931727 | 0 | 4 85E-159 |
| ADCY1 | 0 146399269 | 0 082987211 | 6 52E-160 | 0 429691491 | 0 | 4 85E-159 |
| NOTCH2 | 0 566008643 | 0 082964914 | 7 93E-160 | 0 128197498 | 0 | 5 89E-159 |
| ACSS3 | 0 531010261 | 0 082940071 | 9 87E-160 | 0 157275399 | 0 | 7 33E-159 |
| NARF | 0 254697226 | 0 082934734 | 1 03E-159 | 0 327098287 | 0 | 7 67E-159 |
| FZD4 | 0 070862919 | 0 082881759 | 1 65E-159 | -0 40689098 | 0 | 1 22E-158 |
| CFAP161 | 0 113999353 | 0 082863119 | 1 94E-159 | 0 442202996 | 0 | 1 44E-158 |
| PCNX2 | 0 441596391 | 0 082856052 | 2 06E-159 | 0 212794069 | 0 | 1 53E-158 |
| FRY | 0 320810569 | 0 082854258 | 2 10E-159 | -0 16488373 | 0 | 1 55E-158 |
| HIPK2 | 0 465087282 | 0 082852264 | 2 13E-159 | 0 174827057 | 0 | 1 58E-158 |
| XPO4 | 0 404285089 | 0 082830695 | 2 58E-159 | 0 247167464 | 0 | 1 90E-158 |
| PWWP3B | 0 101835107 | 0 082741929 | 5 62E-159 | -0 37028779 | 0 | 4 14E-158 |
| LIPE-AS1 | 0 208619672 | 0 082679669 | 9 69E-159 | 0 345126555 | 0 | 7 14E-158 |
| TACR1 | 0 096857094 | 0 082467409 | 6 20E-158 | 0 428596343 | 0 | 4 56E-157 |
| CHTF18 | 0 12169005 | 0 082418542 | 9 51E-158 | 0 375201201 | 0 | 6 99E-157 |
| TMC3-AS1 | 0 087767223 | 0 082395863 | 1 16E-157 | 0 450971728 | 0 | 8 51E-157 |
| MTF2 | 0 333298433 | 0 082356522 | 1 63E-157 | 0 268760542 | 0 | 1 20E-156 |
| PPHLN1 | 0 515162475 | 0 082343823 | 1 82E-157 | 0 166965832 | 0 | 1 34E-156 |
| VPS50 | 0 438226952 | 0 082259049 | 3 82E-157 | 0 228729035 | 0 | 2 80E-156 |
| NBEA | 0 625173707 | 0 082242399 | 4 42E-157 | 0 046140452 | 1 27E-50 | 3 23E-156 |
| CDH10 | 0 069349527 | 0 082234205 | 4 74E-157 | 0 458917293 | 0 | 3 47E-156 |
| TBC1D2B | 0 33010984 | 0 082179807 | 7 62E-157 | -0 16992588 | 0 | 5 56E-156 |
| ANKRD26 | 0 393377244 | 0 082163503 | 8 78E-157 | 0 287393155 | 0 | 6 41E-156 |
| PHYKPL | 0 409672384 | 0 082158704 | 9 15E-157 | -0 09245104 | 3 94E-198 | 6 68E-156 |

| | | | | | | |
|---|---|---|---|---|---|---|
| EFCAB2 | 0 336934382 | 0 08215815 | 9 20E-157 | 0 317064541 | 0 | 6 70E-156 |
| TFDP2 | 0 473863052 | 0 08211543 | 1 33E-156 | 0 210409818 | 0 | 9 71E-156 |
| FRAS1 | 0 219556072 | 0 082114339 | 1 35E-156 | -0 24615886 | 0 | 9 80E-156 |
| BMPR1A | 0 605794674 | 0 082046419 | 2 43E-156 | -0 00396794 | 0 198398206 | 1 77E-155 |
| KLHL7 | 0 289276808 | 0 081966948 | 4 85E-156 | 0 317794628 | 0 | 3 52E-155 |
| LRRC3B | 0 040347604 | 0 081862365 | 1 20E-155 | -0 44081629 | 0 | 8 70E-155 |
| LPCAT1 | 0 187060974 | 0 081820532 | 1 73E-155 | 0 383447213 | 0 | 1 25E-154 |
| EHMT1 | 0 60039786 | 0 081798693 | 2 09E-155 | 0 159339796 | 0 | 1 51E-154 |
| TPP2 | 0 42025661 | 0 081773962 | 2 58E-155 | 0 249719949 | 0 | 1 87E-154 |
| TCF12 | 0 877358131 | 0 081742129 | 3 40E-155 | 0 045692756 | 1 12E-49 | 2 46E-154 |
| ARL15 | 0 739772706 | 0 081674766 | 6 10E-155 | 0 042482576 | 3 55E-43 | 4 40E-154 |
| LCLAT1 | 0 381222516 | 0 081671646 | 6 26E-155 | 0 219676343 | 0 | 4 51E-154 |
| SFXN1 | 0 318564276 | 0 081666409 | 6 56E-155 | 0 299127022 | 0 | 4 72E-154 |
| SLC7A6 | 0 268698483 | 0 081635349 | 8 57E-155 | -0 20920649 | 0 | 6 17E-154 |
| DHRS3 | 0 302278655 | 0 081625344 | 9 35E-155 | -0 18554743 | 0 | 6 72E-154 |
| GRM7 | 0 086748777 | 0 081598014 | 1 18E-154 | -0 39654435 | 0 | 8 50E-154 |
| PRIM2 | 0 405598599 | 0 081532347 | 2 09E-154 | 0 18966909 | 0 | 1 50E-153 |
| NOL4 | 0 025565856 | 0 08146374 | 3 78E-154 | 0 493403378 | 0 | 2 71E-153 |
| CHRM3 | 0 057299499 | 0 081447518 | 4 34E-154 | -0 42102907 | 0 | 3 11E-153 |
| ARHGAP6 | 0 407882964 | 0 081428278 | 5 13E-154 | 0 232179727 | 0 | 3 67E-153 |
| VASH1 | 0 105309246 | 0 081268218 | 2 04E-153 | -0 36319788 | 0 | 1 46E-152 |
| FUBP1 | 0 485618016 | 0 081211052 | 3 33E-153 | 0 200064942 | 0 | 2 38E-152 |
| NEDD9 | 0 259313548 | 0 081158514 | 5 23E-153 | -0 20392718 | 0 | 3 73E-152 |
| AP1G2 | 0 078677352 | 0 081158202 | 5 24E-153 | 0 461313505 | 0 | 3 74E-152 |
| MBTD1 | 0 444527993 | 0 081106425 | 8 18E-153 | -0 1039565 | 3 07E-250 | 5 82E-152 |
| RGL2 | 0 158182787 | 0 081104096 | 8 35E-153 | -0 30779806 | 0 | 5 94E-152 |
| CACNA1B | 0 055662371 | 0 081096763 | 8 89E-153 | 0 476549598 | 0 | 6 32E-152 |
| ZNF569 | 0 114218271 | 0 081090298 | 9 40E-153 | 0 452883941 | 0 | 6 67E-152 |
| SBF2 | 0 792846129 | 0 081073996 | 1 08E-152 | 0 037972561 | 7 76E-35 | 7 67E-152 |
| SRBD1 | 0 459014677 | 0 081055882 | 1 26E-152 | 0 202885707 | 0 | 8 95E-152 |
| ELMO1 | 0 546953228 | 0 080944075 | 3 30E-152 | 0 016385115 | 1 09E-07 | 2 33E-151 |
| PPFIBP2 | 0 129780511 | 0 080935095 | 3 56E-152 | -0 32282538 | 0 | 2 52E-151 |
| SLC25A37 | 0 37868116 | 0 080878367 | 5 79E-152 | -0 13178558 | 0 | 4 09E-151 |
| ZNF141 | 0 265167235 | 0 080850401 | 7 35E-152 | -0 21586419 | 0 | 5 19E-151 |
| INTS6 | 0 424301841 | 0 080832597 | 8 56E-152 | -0 09975356 | 1 75E-230 | 6 04E-151 |
| SEPTIN4-AS1 | 0 042755706 | 0 080822535 | 9 33E-152 | 0 485782831 | 0 | 6 58E-151 |
| FMOD | 0 153766347 | 0 080819451 | 9 58E-152 | -0 32566872 | 0 | 6 75E-151 |
| OXR1 | 0 604347909 | 0 080780332 | 1 34E-151 | 0 008950255 | 0 003718625 | 9 43E-151 |
| TSPAN8 | 0 070720146 | 0 080756356 | 1 64E-151 | 0 45853054 | 0 | 1 16E-150 |
| GGA2 | 0 273619387 | 0 080749063 | 1 75E-151 | 0 317382906 | 0 | 1 23E-150 |
| DMXL2 | 0 406388609 | 0 080715379 | 2 33E-151 | 0 224641683 | 0 | 1 64E-150 |
| GAB3 | 0 099579296 | 0 080699131 | 2 68E-151 | 0 442598966 | 0 | 1 88E-150 |
| PKP4 | 0 584169348 | 0 080665954 | 3 56E-151 | 0 132734345 | 0 | 2 50E-150 |
| TTN | 0 192533932 | 0 080653494 | 3 96E-151 | 0 354431072 | 0 | 2 77E-150 |

| | | | | | | |
|---|---|---|---|---|---|---|
| LYN | 0 07974339 | 0 080647212 | 4 18E-151 | 0 464230686 | 0 | 2 93E-150 |
| MIR181A2HG | 0 229502579 | 0 080641886 | 4 37E-151 | 0 320510017 | 0 | 3 06E-150 |
| MAP3K13 | 0 37851935 | 0 080615788 | 5 47E-151 | 0 265578135 | 0 | 3 82E-150 |
| LINC02227 | 0 018788906 | 0 080584003 | 7 17E-151 | 0 498232077 | 0 | 5 01E-150 |
| COL9A1 | 0 036273819 | 0 080545857 | 9 93E-151 | 0 481595678 | 0 | 6 93E-150 |
| TWIST1 | 0 23785955 | 0 080543652 | 1 01E-150 | 0 321487003 | 0 | 7 05E-150 |
| NELL1 | 0 041775333 | 0 080530759 | 1 13E-150 | -0 43673643 | 0 | 7 87E-150 |
| ZNF75D | 0 244646019 | 0 08051615 | 1 28E-150 | 0 353876816 | 0 | 8 91E-150 |
| LINC01422 | 0 061354248 | 0 080448548 | 2 28E-150 | -0 4085367 | 0 | 1 58E-149 |
| EPB41L3 | 0 338143192 | 0 080437457 | 2 50E-150 | 0 292495254 | 0 | 1 74E-149 |
| PIGN | 0 394624127 | 0 080430547 | 2 65E-150 | 0 254784128 | 0 | 1 84E-149 |
| ZNF519 | 0 166606385 | 0 080316783 | 6 98E-150 | 0 357474321 | 0 | 4 84E-149 |
| GRK3 | 0 227189659 | 0 080296966 | 8 27E-150 | -0 24800195 | 0 | 5 72E-149 |
| PARP4 | 0 433848585 | 0 080268863 | 1 05E-149 | -0 09841051 | 2 40E-224 | 7 26E-149 |
| AFG1L | 0 229778607 | 0 080235697 | 1 39E-149 | -0 23122678 | 0 | 9 62E-149 |
| MEF2A | 0 719936799 | 0 08023245 | 1 43E-149 | 0 081507541 | 2 59E-154 | 9 88E-149 |
| GAS1 | 0 05966001 | 0 080218771 | 1 61E-149 | 0 472153067 | 0 | 1 11E-148 |
| FLNB | 0 380061297 | 0 080135514 | 3 26E-149 | 0 219673214 | 0 | 2 25E-148 |
| JAK3 | 0 061773048 | 0 080129206 | 3 44E-149 | 0 476973531 | 0 | 2 37E-148 |
| FIP1L1 | 0 441834345 | 0 080117132 | 3 81E-149 | 0 2151535 | 0 | 2 62E-148 |
| ARHGAP35 | 0 447916468 | 0 080109965 | 4 05E-149 | 0 201981871 | 0 | 2 79E-148 |
| BCLAF1 | 0 491262302 | 0 080107496 | 4 13E-149 | -0 0400845 | 1 26E-38 | 2 84E-148 |
| GASK1A | 0 077944452 | 0 080104169 | 4 25E-149 | 0 454962541 | 0 | 2 92E-148 |
| HGSNAT | 0 4423864 | 0 080102351 | 4 32E-149 | -0 07921031 | 7 99E-146 | 2 96E-148 |
| SLC16A7 | 0 149711599 | 0 080060321 | 6 16E-149 | 0 426869133 | 0 | 4 23E-148 |
| NUP214 | 0 427242961 | 0 080057878 | 6 29E-149 | 0 22734741 | 0 | 4 31E-148 |
| HMG20A | 0 334069407 | 0 079978975 | 1 23E-148 | -0 16084305 | 0 | 8 41E-148 |
| KCNN3 | 0 060126402 | 0 079975984 | 1 26E-148 | -0 4157846 | 0 | 8 62E-148 |
| MAX | 0 392834707 | 0 079954481 | 1 51E-148 | -0 11943607 | 0 | 1 03E-147 |
| RGS22 | 0 129323637 | 0 079932607 | 1 82E-148 | -0 32727479 | 0 | 1 24E-147 |
| PTPRM | 0 577563724 | 0 079904904 | 2 30E-148 | 0 016704843 | 6 13E-08 | 1 57E-147 |
| KIFAP3 | 0 486074889 | 0 079867057 | 3 17E-148 | -0 05737172 | 2 61E-77 | 2 16E-147 |
| SPEF2 | 0 271782376 | 0 079864629 | 3 23E-148 | -0 20026679 | 0 | 2 20E-147 |
| MRPS31 | 0 28847728 | 0 079861973 | 3 31E-148 | -0 20268607 | 0 | 2 25E-147 |
| ZSCAN5A | 0 224895776 | 0 079831205 | 4 29E-148 | -0 25023677 | 0 | 2 92E-147 |
| AQR | 0 403875807 | 0 079770664 | 7 15E-148 | 0 242529479 | 0 | 4 86E-147 |
| CARD11 | 0 029458796 | 0 079761099 | 7 75E-148 | 0 491742911 | 0 | 5 26E-147 |
| TTYH2 | 0 15228151 | 0 079758618 | 7 92E-148 | -0 30811266 | 0 | 5 37E-147 |
| CNTN4 | 0 559326874 | 0 079757152 | 8 02E-148 | 0 01014681 | 0 001005589 | 5 43E-147 |
| MCF2L | 0 064771278 | 0 079663583 | 1 77E-147 | 0 470117515 | 0 | 1 20E-146 |
| SOS2 | 0 53711142 | 0 079655367 | 1 89E-147 | -0 02833323 | 4 10E-20 | 1 28E-146 |
| MAP2K5 | 0 542203651 | 0 079642193 | 2 12E-147 | -0 02378021 | 1 27E-14 | 1 43E-146 |
| PRMT8 | 0 030905561 | 0 079570271 | 3 88E-147 | -0 44864432 | 0 | 2 62E-146 |
| C20orf194 | 0 255154099 | 0 079542184 | 4 91E-147 | 0 280642764 | 0 | 3 32E-146 |

| | | | | | | |
|---|---|---|---|---|---|---|
| F11-AS1 | 0 037720584 | 0 079427411 | 1 29E-146 | 0 490177082 | 0 | 8 70E-146 |
| MSH2 | 0 268222573 | 0 079411384 | 1 48E-146 | 0 308391551 | 0 | 9 95E-146 |
| ZNF532 | 0 419818774 | 0 079309883 | 3 47E-146 | -0 08807272 | 6 25E-180 | 2 33E-145 |
| TLK1 | 0 555119834 | 0 079305216 | 3 60E-146 | 0 151063109 | 0 | 2 42E-145 |
| AHI1 | 0 67220308 | 0 079260544 | 5 25E-146 | 0 028492675 | 2 53E-20 | 3 52E-145 |
| PIAS1 | 0 710913556 | 0 079231569 | 6 69E-146 | 0 027955047 | 1 27E-19 | 4 49E-145 |
| SLC44A2 | 0 186480364 | 0 079222295 | 7 23E-146 | -0 27519046 | 0 | 4 85E-145 |
| CTCF | 0 302069254 | 0 079185589 | 9 84E-146 | 0 243487398 | 0 | 6 59E-145 |
| BRINP1 | 0 042489197 | 0 079182119 | 1 01E-145 | -0 44197687 | 0 | 6 78E-145 |
| CRABP1 | 0 05209305 | 0 079096837 | 2 07E-145 | 0 471767175 | 0 | 1 38E-144 |
| TAF1 | 0 461384706 | 0 079089781 | 2 20E-145 | 0 215707027 | 0 | 1 47E-144 |
| LAMB1 | 0 79825246 | 0 079087751 | 2 23E-145 | 0 065298924 | 1 24E-99 | 1 49E-144 |
| GALNT1 | 0 362290838 | 0 079081365 | 2 36E-145 | 0 273631438 | 0 | 1 57E-144 |
| SUMF1 | 0 51802745 | 0 079066713 | 2 66E-145 | -0 04670664 | 7 90E-52 | 1 78E-144 |
| OXNAD1 | 0 234632883 | 0 078971282 | 5 92E-145 | 0 342185957 | 0 | 3 95E-144 |
| KCNK1 | 0 1167977 | 0 078968099 | 6 08E-145 | 0 432681702 | 0 | 4 05E-144 |
| PRPF4B | 0 584074166 | 0 078946434 | 7 28E-145 | 0 151789389 | 0 | 4 85E-144 |
| ELP4 | 0 416458853 | 0 078889469 | 1 17E-144 | 0 223937436 | 0 | 7 79E-144 |
| IQSEC2 | 0 108297957 | 0 078874528 | 1 33E-144 | -0 36274363 | 0 | 8 82E-144 |
| TENM4 | 0 485522834 | 0 07886855 | 1 40E-144 | -0 07470234 | 7 10E-130 | 9 26E-144 |
| UGGT2 | 0 636290952 | 0 078860043 | 1 50E-144 | 0 160099994 | 0 | 9 94E-144 |
| NUP107 | 0 395328473 | 0 078855559 | 1 56E-144 | 0 202032806 | 0 | 1 03E-143 |
| SLC35F4 | 0 048104929 | 0 078805383 | 2 37E-144 | 0 479101206 | 0 | 1 57E-143 |
| PRLR | 0 081618473 | 0 078773269 | 3 09E-144 | 0 451901469 | 0 | 2 04E-143 |
| LINC02086 | 0 057946736 | 0 07876554 | 3 30E-144 | 0 46349123 | 0 | 2 18E-143 |
| CFDP1 | 0 552607032 | 0 078736909 | 4 19E-144 | -0 02603068 | 3 21E-17 | 2 77E-143 |
| PCDH15 | 0 087795778 | 0 07868952 | 6 21E-144 | 0 427238882 | 0 | 4 10E-143 |
| ARID2 | 0 597589994 | 0 078646603 | 8 89E-144 | 0 120309235 | 0 | 5 86E-143 |
| WDR70 | 0 661295235 | 0 078635911 | 9 71E-144 | 0 123426182 | 0 | 6 40E-143 |
| TRPM7 | 0 640440883 | 0 07863133 | 1 01E-143 | 0 125792015 | 0 | 6 64E-143 |
| NAP1L4 | 0 458015267 | 0 078620411 | 1 11E-143 | 0 216409341 | 0 | 7 27E-143 |
| ZDHHC20 | 0 331908778 | 0 078589019 | 1 43E-143 | 0 285871837 | 0 | 9 44E-143 |
| ARMC9 | 0 35234433 | 0 078576971 | 1 59E-143 | -0 13492162 | 0 | 1 04E-142 |
| BABAM2 | 0 62384116 | 0 078537875 | 2 20E-143 | 0 010988585 | 0 000368286 | 1 44E-142 |
| ZNF652 | 0 521901353 | 0 078526842 | 2 41E-143 | 0 170039183 | 0 | 1 58E-142 |
| AASS | 0 471730978 | 0 078494723 | 3 14E-143 | -0 05755897 | 8 37E-78 | 2 06E-142 |
| PPP6R2 | 0 549285184 | 0 078466885 | 3 96E-143 | 0 148160036 | 0 | 2 59E-142 |
| RMDN2 | 0 214263958 | 0 078464683 | 4 03E-143 | -0 24803713 | 0 | 2 64E-142 |
| LURAP1L-AS1 | 0 12683939 | 0 078445453 | 4 73E-143 | 0 410487909 | 0 | 3 09E-142 |
| ZNF678 | 0 252955398 | 0 078429435 | 5 41E-143 | 0 327973537 | 0 | 3 53E-142 |
| AEBP1 | 0 297709924 | 0 078349153 | 1 05E-142 | -0 17892973 | 0 | 6 86E-142 |
| TFCP2 | 0 483723896 | 0 078306414 | 1 50E-142 | -0 07522652 | 1 11E-131 | 9 78E-142 |
| UACA | 0 710114028 | 0 078268963 | 2 05E-142 | 0 033934449 | 3 72E-28 | 1 33E-141 |
| TNFAIP2 | 0 1281529 | 0 078242967 | 2 54E-142 | -0 34189599 | 0 | 1 65E-141 |

| | | | | | | |
|---|---|---|---|---|---|---|
| ZNF44 | 0 294645067 | 0 078219718 | 3 08E-142 | -0 19102401 | 0 | 2 00E-141 |
| TDRP | 0 12736289 | 0 078187154 | 4 03E-142 | -0 3370141 | 0 | 2 62E-141 |
| CCNL2 | 0 535093564 | 0 078179791 | 4 28E-142 | 0 161715879 | 0 | 2 78E-141 |
| TAFA5 | 0 09009918 | 0 078151127 | 5 43E-142 | -0 374238 | 0 | 3 52E-141 |
| ENSA | 0 462774362 | 0 078124995 | 6 74E-142 | -0 08610091 | 5 02E-172 | 4 37E-141 |
| SYT6 | 0 018950715 | 0 078063482 | 1 12E-141 | -0 46574574 | 0 | 7 26E-141 |
| ITGBL1 | 0 200652948 | 0 078050332 | 1 25E-141 | -0 25965624 | 0 | 8 09E-141 |
| CDK5RAP2 | 0 370571662 | 0 078026794 | 1 52E-141 | 0 235756392 | 0 | 9 82E-141 |
| UBE2E1 | 0 550893758 | 0 078021461 | 1 59E-141 | -0 03752342 | 4 68E-34 | 1 03E-140 |
| NEK11 | 0 273962041 | 0 078013413 | 1 69E-141 | -0 1976319 | 0 | 1 09E-140 |
| TTLL5 | 0 42625307 | 0 078006095 | 1 80E-141 | 0 230218715 | 0 | 1 16E-140 |
| STEAP2 | 0 171898498 | 0 077954573 | 2 75E-141 | -0 30926978 | 0 | 1 78E-140 |
| SASH1 | 0 674592146 | 0 077946944 | 2 93E-141 | 0 017240665 | 2 29E-08 | 1 89E-140 |
| ARHGEF10 | 0 471150368 | 0 077931789 | 3 32E-141 | 0 235537994 | 0 | 2 14E-140 |
| DIPK2A | 0 193723706 | 0 077909485 | 4 00E-141 | 0 389908496 | 0 | 2 57E-140 |
| LINC00954 | 0 03873903 | 0 077907199 | 4 07E-141 | 0 485595888 | 0 | 2 62E-140 |
| ZBTB7C | 0 182853934 | 0 077872722 | 5 41E-141 | -0 26076714 | 0 | 3 47E-140 |
| SLCO3A1 | 0 276322552 | 0 077841324 | 7 01E-141 | 0 369839711 | 0 | 4 50E-140 |
| TTC7A | 0 276617616 | 0 077830367 | 7 67E-141 | 0 290962343 | 0 | 4 92E-140 |
| NR2C2 | 0 382050599 | 0 077781072 | 1 15E-140 | 0 2495154 | 0 | 7 38E-140 |
| LINC02438 | 0 065418515 | 0 07776514 | 1 31E-140 | 0 469375974 | 0 | 8 41E-140 |
| ZNF248 | 0 304163256 | 0 077738574 | 1 63E-140 | -0 16739884 | 0 | 1 05E-139 |
| WDR59 | 0 360920219 | 0 077687898 | 2 48E-140 | -0 14008149 | 0 | 1 58E-139 |
| IQCN | 0 093297291 | 0 077658798 | 3 15E-140 | -0 36705326 | 0 | 2 01E-139 |
| CEP78 | 0 222706592 | 0 077640909 | 3 65E-140 | 0 304605801 | 0 | 2 33E-139 |
| WDPCP | 0 656707468 | 0 077633497 | 3 88E-140 | 0 119831138 | 0 | 2 47E-139 |
| ST6GAL2 | 0 033237517 | 0 077631026 | 3 96E-140 | 0 488117658 | 0 | 2 52E-139 |
| FRMD4B | 0 364565685 | 0 077623407 | 4 21E-140 | -0 15216407 | 0 | 2 68E-139 |
| ASH1L | 0 782252384 | 0 07759411 | 5 36E-140 | 0 094849449 | 1 76E-208 | 3 41E-139 |
| SMAD2 | 0 496839961 | 0 07756276 | 6 93E-140 | -0 04591247 | 3 86E-50 | 4 41E-139 |
| ARMC8 | 0 458376958 | 0 077551884 | 7 58E-140 | 0 214417611 | 0 | 4 82E-139 |
| CCNY | 0 57332813 | 0 077540191 | 8 34E-140 | -0 01241713 | 5 70E-05 | 5 30E-139 |
| PLSCR4 | 0 497496716 | 0 077533897 | 8 79E-140 | -0 04542068 | 4 15E-49 | 5 58E-139 |
| EPG5 | 0 35465725 | 0 077516731 | 1 01E-139 | 0 274405022 | 0 | 6 41E-139 |
| CALN1 | 0 105347319 | 0 077427143 | 2 11E-139 | 0 446824637 | 0 | 1 34E-138 |
| MRPS28 | 0 411385658 | 0 077416806 | 2 30E-139 | -0 12337467 | 0 | 1 45E-138 |
| STXBP4 | 0 455407283 | 0 077415079 | 2 33E-139 | -0 06803513 | 5 17E-108 | 1 47E-138 |
| SSC5D | 0 083550665 | 0 077404336 | 2 54E-139 | -0 39272146 | 0 | 1 61E-138 |
| TRPC3 | 0 043279207 | 0 077395436 | 2 73E-139 | 0 486561176 | 0 | 1 73E-138 |
| PLXNC1 | 0 191534522 | 0 077355425 | 3 80E-139 | 0 366651853 | 0 | 2 40E-138 |
| ALPK3 | 0 053863433 | 0 077325744 | 4 84E-139 | -0 422888 | 0 | 3 05E-138 |
| NOD1 | 0 231596581 | 0 077321781 | 5 00E-139 | 0 34320323 | 0 | 3 15E-138 |
| TRAPPC9 | 0 535322 | 0 07730813 | 5 59E-139 | 0 175307382 | 0 | 3 52E-138 |
| ANKRD36B | 0 364556167 | 0 077252439 | 8 81E-139 | 0 228703685 | 0 | 5 55E-138 |

| | | | | | | |
|---|---|---|---|---|---|---|
| DCLK1 | 0 384772801 | 0 077147543 | 2 08E-138 | -0 12302147 | 0 | 1 31E-137 |
| CCDC144NL | 0 112923797 | 0 07712868 | 2 42E-138 | 0 423299553 | 0 | 1 52E-137 |
| LBR | 0 194837334 | 0 077107504 | 2 88E-138 | 0 357999001 | 0 | 1 81E-137 |
| FYN | 0 528449868 | 0 077036658 | 5 13E-138 | -0 01073812 | 0 00050023 | 3 22E-137 |
| RREB1 | 0 428366108 | 0 077029561 | 5 44E-138 | -0 09980478 | 1 02E-230 | 3 41E-137 |
| GREM2 | 0 040138204 | 0 077028689 | 5 48E-138 | 0 484498747 | 0 | 3 43E-137 |
| LINC02245 | 0 135862633 | 0 077028208 | 5 50E-138 | 0 41898356 | 0 | 3 44E-137 |
| TXNDC16 | 0 189821248 | 0 076987076 | 7 69E-138 | 0 38346388 | 0 | 4 81E-137 |
| GRIK4 | 0 069815918 | 0 076958354 | 9 72E-138 | 0 480519204 | 0 | 6 07E-137 |
| ZFP36L1 | 0 479859512 | 0 07694669 | 1 07E-137 | 0 180837593 | 0 | 6 67E-137 |
| ZNF407 | 0 528126249 | 0 076931854 | 1 21E-137 | -0 03837851 | 1 50E-35 | 7 52E-137 |
| GLT1D1 | 0 045278026 | 0 076898079 | 1 59E-137 | 0 481795414 | 0 | 9 89E-137 |
| TMPO | 0 294273857 | 0 076849174 | 2 36E-137 | 0 218776794 | 0 | 1 47E-136 |
| CLOCK | 0 420932402 | 0 076755299 | 5 07E-137 | 0 210899395 | 0 | 3 15E-136 |
| LINC01036 | 0 135443833 | 0 07673832 | 5 82E-137 | 0 408920091 | 0 | 3 62E-136 |
| ZBTB10 | 0 228950524 | 0 076671292 | 1 00E-136 | -0 24032502 | 0 | 6 23E-136 |
| MTERF1 | 0 355628105 | 0 076649997 | 1 19E-136 | -0 1362856 | 0 | 7 40E-136 |
| NMD3 | 0 358607299 | 0 076623679 | 1 48E-136 | 0 260394199 | 0 | 9 15E-136 |
| THUMPD2 | 0 32869163 | 0 07659237 | 1 90E-136 | 0 277707259 | 0 | 1 18E-135 |
| STAT6 | 0 298861625 | 0 076469906 | 5 13E-136 | -0 19665536 | 0 | 3 18E-135 |
| ANKRD36C | 0 579867126 | 0 076447522 | 6 15E-136 | 0 137371012 | 0 | 3 80E-135 |
| TLE1 | 0 363880375 | 0 076432836 | 6 93E-136 | -0 11993273 | 0 | 4 28E-135 |
| LYPD6 | 0 079105671 | 0 076426063 | 7 32E-136 | 0 452628198 | 0 | 4 52E-135 |
| SCX | 0 070567855 | 0 076378426 | 1 08E-135 | 0 462378298 | 0 | 6 63E-135 |
| HRH2 | 0 043774152 | 0 076365705 | 1 19E-135 | -0 4332294 | 0 | 7 34E-135 |
| RANBP9 | 0 474101007 | 0 076320543 | 1 72E-135 | 0 221581185 | 0 | 1 06E-134 |
| SLC38A6 | 0 401153605 | 0 07630619 | 1 93E-135 | 0 235354795 | 0 | 1 19E-134 |
| CDK5RAP3 | 0 31623232 | 0 076228423 | 3 61E-135 | 0 28373586 | 0 | 2 22E-134 |
| ARMC2 | 0 242161771 | 0 076189422 | 4 95E-135 | -0 20640797 | 0 | 3 04E-134 |
| SRGAP2 | 0 514686566 | 0 076172304 | 5 68E-135 | 0 210068622 | 0 | 3 49E-134 |
| PLD3 | 0 412156631 | 0 07616348 | 6 10E-135 | -0 08573702 | 1 38E-170 | 3 74E-134 |
| FZD1 | 0 156707468 | 0 076133344 | 7 78E-135 | -0 31623048 | 0 | 4 77E-134 |
| ZDBF2 | 0 283737222 | 0 076109554 | 9 42E-135 | -0 21211406 | 0 | 5 77E-134 |
| TUSC3 | 0 522367745 | 0 076096096 | 1 05E-134 | 0 152898939 | 0 | 6 42E-134 |
| KANTR | 0 345634007 | 0 076055516 | 1 46E-134 | 0 290889979 | 0 | 8 90E-134 |
| ITGA8 | 0 193162133 | 0 07603433 | 1 73E-134 | -0 26191713 | 0 | 1 05E-133 |
| RFC3 | 0 233109973 | 0 075994033 | 2 39E-134 | 0 309634432 | 0 | 1 46E-133 |
| ZNF718 | 0 201100303 | 0 07598424 | 2 58E-134 | 0 403730294 | 0 | 1 58E-133 |
| HERPUD2 | 0 286449906 | 0 075973553 | 2 81E-134 | -0 19238291 | 0 | 1 72E-133 |
| MEG3 | 0 708838591 | 0 075935945 | 3 81E-134 | 0 030884111 | 1 34E-23 | 2 32E-133 |
| LINC02307 | 0 083560184 | 0 07592144 | 4 28E-134 | -0 38864891 | 0 | 2 60E-133 |
| STK38L | 0 278778245 | 0 075845826 | 7 85E-134 | 0 329305295 | 0 | 4 77E-133 |
| LRRN1 | 0 088214578 | 0 075742919 | 1 79E-133 | 0 444899524 | 0 | 1 09E-132 |
| LTBP3 | 0 387980431 | 0 075737357 | 1 87E-133 | -0 10799462 | 5 29E-270 | 1 14E-132 |

| | | | | | | |
|---|---|---|---|---|---|---|
| NEMF | 0 546734309 | 0 075732788 | 1 94E-133 | -0 0248638 | 7 61E-16 | 1 18E-132 |
| RAB3GAP2 | 0 550750985 | 0 075685806 | 2 83E-133 | -0 02659556 | 6 58E-18 | 1 72E-132 |
| TAFA2 | 0 197464354 | 0 075674285 | 3 11E-133 | -0 28646381 | 0 | 1 88E-132 |
| ANKRD44 | 0 460727951 | 0 075668438 | 3 26E-133 | 0 232115974 | 0 | 1 97E-132 |
| ZHX3 | 0 505768023 | 0 075619438 | 4 82E-133 | 0 185665581 | 0 | 2 91E-132 |
| USP3 | 0 504026194 | 0 075618913 | 4 84E-133 | 0 176923869 | 0 | 2 92E-132 |
| MCCC1 | 0 28012031 | 0 075610337 | 5 18E-133 | -0 19768783 | 0 | 3 13E-132 |
| KLHL29 | 0 557937218 | 0 075588928 | 6 15E-133 | -0 04785754 | 2 51E-54 | 3 71E-132 |
| FGF7 | 0 182530315 | 0 07554853 | 8 50E-133 | -0 27457587 | 0 | 5 12E-132 |
| UFL1-AS1 | 0 081846909 | 0 07554285 | 8 90E-133 | -0 39563735 | 0 | 5 36E-132 |
| CACNA1H | 0 017580096 | 0 075509163 | 1 16E-132 | 0 496964163 | 0 | 7 01E-132 |
| EPN2 | 0 499095772 | 0 075462892 | 1 69E-132 | -0 03531918 | 2 31E-30 | 1 01E-131 |
| NAA38 | 0 302859264 | 0 075434528 | 2 11E-132 | 0 32114383 | 0 | 1 27E-131 |
| NBAS | 0 651834155 | 0 07540361 | 2 71E-132 | 0 001622085 | 0 599051555 | 1 63E-131 |
| TTC17 | 0 66095258 | 0 075394393 | 2 91E-132 | 0 011415395 | 0 000215447 | 1 75E-131 |
| RIMKLB | 0 397679465 | 0 075387232 | 3 08E-132 | 0 245044996 | 0 | 1 85E-131 |
| NRXN3 | 0 24873884 | 0 07538502 | 3 14E-132 | -0 23381566 | 0 | 1 88E-131 |
| HELB | 0 203460814 | 0 075332116 | 4 78E-132 | 0 348968322 | 0 | 2 87E-131 |
| GLMN | 0 269707411 | 0 075298194 | 6 27E-132 | 0 324355322 | 0 | 3 76E-131 |
| STAG2 | 0 569320972 | 0 075254017 | 8 92E-132 | 0 182329063 | 0 | 5 33E-131 |
| SH3BGRL2 | 0 100435933 | 0 075221149 | 1 16E-131 | -0 36232312 | 0 | 6 92E-131 |
| MALRD1 | 0 235451448 | 0 07517342 | 1 69E-131 | -0 23094613 | 0 | 1 01E-130 |
| CNTNAP2 | 0 191258495 | 0 075148563 | 2 06E-131 | -0 27807296 | 0 | 1 23E-130 |
| CAPS2 | 0 282595039 | 0 075136336 | 2 27E-131 | -0 19313264 | 0 | 1 36E-130 |
| ANKRD36 | 0 613104643 | 0 075127336 | 2 44E-131 | 0 117046308 | 0 | 1 46E-130 |
| ZNF577 | 0 146646742 | 0 075077558 | 3 63E-131 | 0 416731821 | 0 | 2 16E-130 |
| CENPJ | 0 170327997 | 0 075061522 | 4 12E-131 | 0 342348259 | 0 | 2 45E-130 |
| PCOLCE | 0 443519065 | 0 07501025 | 6 19E-131 | 0 191476088 | 0 | 3 69E-130 |
| PLPP3 | 0 391406979 | 0 074907735 | 1 40E-130 | -0 10028681 | 6 06E-233 | 8 30E-130 |
| LITAF | 0 304172774 | 0 074867346 | 1 92E-130 | -0 163833 | 0 | 1 14E-129 |
| CDH3 | 0 041956178 | 0 074826758 | 2 65E-130 | 0 485766272 | 0 | 1 57E-129 |
| HS3ST3B1 | 0 067426853 | 0 07481332 | 2 95E-130 | 0 477799965 | 0 | 1 75E-129 |
| OVCH1-AS1 | 0 034808018 | 0 074794652 | 3 42E-130 | -0 45172019 | 0 | 2 03E-129 |
| XRN1 | 0 643058385 | 0 074790023 | 3 55E-130 | 0 135738567 | 0 | 2 10E-129 |
| VPS13B | 0 864127848 | 0 074764918 | 4 33E-130 | 0 038687731 | 4 26E-36 | 2 56E-129 |
| LANCL2 | 0 194447088 | 0 0747578 | 4 58E-130 | 0 380514707 | 0 | 2 70E-129 |
| ATOH8 | 0 036863947 | 0 074745627 | 5 04E-130 | 0 486130976 | 0 | 2 98E-129 |
| ATG4B | 0 312367935 | 0 074732804 | 5 58E-130 | 0 282573413 | 0 | 3 29E-129 |
| PHACTR3 | 0 05061773 | 0 074721562 | 6 10E-130 | 0 480378823 | 0 | 3 60E-129 |
| ELAVL4 | 0 012126173 | 0 074698374 | 7 33E-130 | 0 501926095 | 0 | 4 32E-129 |
| SERPINI1 | 0 087110468 | 0 074679322 | 8 52E-130 | -0 39049186 | 0 | 5 02E-129 |
| CCDC144A | 0 247035084 | 0 074660336 | 9 90E-130 | 0 367444345 | 0 | 5 82E-129 |
| IFT80 | 0 366650168 | 0 074656879 | 1 02E-129 | 0 251927605 | 0 | 5 98E-129 |
| RGL1 | 0 554415488 | 0 074652357 | 1 05E-129 | -0 01746338 | 1 51E-08 | 6 19E-129 |

| | | | | | | |
|---|---|---|---|---|---|---|
| CAMK4 | 0 180483905 | 0 07460298 | 1 56E-129 | 0 321013946 | 0 | 9 14E-129 |
| NSMCE2 | 0 691534522 | 0 074493961 | 3 68E-129 | 0 095402451 | 6 62E-211 | 2 16E-128 |
| JCAD | 0 181530906 | 0 074492669 | 3 72E-129 | -0 28511792 | 0 | 2 18E-128 |
| L3MBTL1 | 0 146161314 | 0 074469786 | 4 45E-129 | 0 420344502 | 0 | 2 61E-128 |
| LINC00351 | 0 016970931 | 0 074468249 | 4 51E-129 | 0 498439481 | 0 | 2 64E-128 |
| HS6ST1 | 0 117416383 | 0 074447601 | 5 30E-129 | -0 34465336 | 0 | 3 10E-128 |
| CEP290 | 0 483133769 | 0 074406126 | 7 35E-129 | 0 194515852 | 0 | 4 30E-128 |
| MYRFL | 0 050598694 | 0 074401458 | 7 63E-129 | 0 478356234 | 0 | 4 46E-128 |
| FBXL13 | 0 340589366 | 0 074391359 | 8 26E-129 | 0 246102725 | 0 | 4 82E-128 |
| ANKRD17 | 0 768051246 | 0 074288428 | 1 86E-128 | 0 082309594 | 2 46E-157 | 1 08E-127 |
| TMEM241 | 0 306323885 | 0 074260949 | 2 30E-128 | -0 19816978 | 0 | 1 34E-127 |
| NLRP1 | 0 150891854 | 0 074258371 | 2 35E-128 | 0 387536793 | 0 | 1 37E-127 |
| CDC14A | 0 25872342 | 0 074254916 | 2 41E-128 | -0 22799542 | 0 | 1 41E-127 |
| ABCA5 | 0 480839885 | 0 074222093 | 3 12E-128 | -0 07870316 | 5 55E-144 | 1 82E-127 |
| JPX | 0 682701643 | 0 074193493 | 3 91E-128 | 0 011556487 | 0 000179737 | 2 27E-127 |
| ARSB | 0 200633911 | 0 074141654 | 5 88E-128 | 0 354639157 | 0 | 3 41E-127 |
| DDX10 | 0 35134492 | 0 074107897 | 7 66E-128 | -0 14595795 | 0 | 4 44E-127 |
| LTC4S | 0 030981706 | 0 074099985 | 8 15E-128 | -0 45233974 | 0 | 4 73E-127 |
| EIF1B-AS1 | 0 187850983 | 0 074097203 | 8 33E-128 | -0 27500739 | 0 | 4 83E-127 |
| ANKRD50 | 0 332308542 | 0 074079432 | 9 57E-128 | 0 291575028 | 0 | 5 55E-127 |
| KCNIP4 | 0 158496888 | 0 074044782 | 1 26E-127 | 0 387360585 | 0 | 7 27E-127 |
| OGT | 0 609963641 | 0 074008754 | 1 67E-127 | -0 00076107 | 0 805152891 | 9 64E-127 |
| TLE3 | 0 146732406 | 0 073974603 | 2 18E-127 | 0 405768848 | 0 | 1 26E-126 |
| TSPOAP1 | 0 037092384 | 0 073948358 | 2 67E-127 | 0 494040907 | 0 | 1 54E-126 |
| UBR2 | 0 617311683 | 0 073946357 | 2 71E-127 | 0 00573998 | 0 062813632 | 1 57E-126 |
| SCML4 | 0 025603929 | 0 07393194 | 3 04E-127 | 0 49399937 | 0 | 1 75E-126 |
| RBMS3-AS2 | 0 359349717 | 0 073917239 | 3 41E-127 | -0 14757128 | 0 | 1 97E-126 |
| LRRTM3 | 0 046448764 | 0 073901857 | 3 84E-127 | 0 484047144 | 0 | 2 22E-126 |
| GIGYF1 | 0 282252384 | 0 073892003 | 4 15E-127 | 0 317237205 | 0 | 2 39E-126 |
| NID2 | 0 207610744 | 0 073881533 | 4 51E-127 | -0 25127729 | 0 | 2 59E-126 |
| CTNNA3 | 0 144114904 | 0 073865212 | 5 12E-127 | 0 411588601 | 0 | 2 94E-126 |
| LRRC1 | 0 245550247 | 0 073859948 | 5 33E-127 | -0 22102625 | 0 | 3 07E-126 |
| ZBTB20-AS5 | 0 359130799 | 0 073827913 | 6 85E-127 | -0 17018371 | 0 | 3 94E-126 |
| PSPC1 | 0 510850736 | 0 073777087 | 1 02E-126 | 0 188873344 | 0 | 5 85E-126 |
| SGCG | 0 055167425 | 0 073745134 | 1 31E-126 | 0 465444479 | 0 | 7 50E-126 |
| SMAD1 | 0 225828558 | 0 073727104 | 1 51E-126 | 0 35915195 | 0 | 8 63E-126 |
| ASXL1 | 0 591564981 | 0 073722081 | 1 57E-126 | 0 156842856 | 0 | 8 97E-126 |
| LYRM4 | 0 290523691 | 0 073698406 | 1 88E-126 | 0 307382833 | 0 | 1 08E-125 |
| CPEB3 | 0 319715977 | 0 073686442 | 2 07E-126 | -0 18059688 | 0 | 1 18E-125 |
| SRGAP2B | 0 343244941 | 0 073672253 | 2 31E-126 | 0 242641908 | 0 | 1 32E-125 |
| HERC1 | 0 684300699 | 0 073652835 | 2 69E-126 | 0 124201796 | 0 | 1 53E-125 |
| SEMA3E | 0 354847614 | 0 073626176 | 3 31E-126 | -0 12233134 | 0 | 1 89E-125 |
| FAM227B | 0 28232853 | 0 073590469 | 4 37E-126 | -0 21407456 | 0 | 2 49E-125 |
| ARHGEF9 | 0 215034932 | 0 073439538 | 1 41E-125 | -0 24454002 | 0 | 8 05E-125 |

| | | | | | | |
|---|---|---|---|---|---|---|
| CHST15 | 0 06218233 | 0 073426214 | 1 57E-125 | 0 468704593 | 0 | 8 92E-125 |
| CARD8 | 0 22834136 | 0 073423683 | 1 60E-125 | 0 334211666 | 0 | 9 10E-125 |
| IRAK1BP1 | 0 214606613 | 0 073403502 | 1 87E-125 | -0 24392049 | 0 | 1 06E-124 |
| ANP32A | 0 212227066 | 0 073401256 | 1 90E-125 | 0 379074731 | 0 | 1 08E-124 |
| SPRY4 | 0 281871657 | 0 073343449 | 2 98E-125 | -0 18200573 | 0 | 1 69E-124 |
| UBAC2 | 0 442986046 | 0 073333747 | 3 21E-125 | -0 0936412 | 3 11E-203 | 1 82E-124 |
| NVL | 0 351040338 | 0 073328317 | 3 35E-125 | 0 256745248 | 0 | 1 90E-124 |
| LFNG | 0 051598104 | 0 073300164 | 4 17E-125 | -0 42937838 | 0 | 2 36E-124 |
| SLCO2A1 | 0 07422284 | 0 073280694 | 4 85E-125 | -0 40495143 | 0 | 2 75E-124 |
| NASP | 0 380042261 | 0 073279337 | 4 90E-125 | 0 165025733 | 0 | 2 78E-124 |
| ZBED3-AS1 | 0 097018903 | 0 0732404 | 6 63E-125 | 0 451815306 | 0 | 3 75E-124 |
| PRKD2 | 0 167672422 | 0 073224798 | 7 48E-125 | 0 380366703 | 0 | 4 23E-124 |
| GAB1 | 0 536102492 | 0 07321104 | 8 32E-125 | -0 02412293 | 5 28E-15 | 4 70E-124 |
| CD36 | 0 067512516 | 0 073208369 | 8 49E-125 | 0 462716265 | 0 | 4 80E-124 |
| MYO3A | 0 069663627 | 0 073204709 | 8 74E-125 | 0 454109758 | 0 | 4 93E-124 |
| TG | 0 07732577 | 0 073160616 | 1 23E-124 | 0 463703297 | 0 | 6 93E-124 |
| TERF2IP | 0 388599113 | 0 073125734 | 1 61E-124 | -0 08775886 | 1 17E-178 | 9 07E-124 |
| ITPR1 | 0 423207249 | 0 073102328 | 1 93E-124 | -0 1236442 | 0 | 1 09E-123 |
| IPO9-AS1 | 0 312034798 | 0 073041605 | 3 09E-124 | 0 245035719 | 0 | 1 74E-123 |
| SNTG1 | 0 045344654 | 0 072998588 | 4 30E-124 | 0 478738544 | 0 | 2 42E-123 |
| LTB4R | 0 087329386 | 0 072985976 | 4 74E-124 | 0 455765074 | 0 | 2 66E-123 |
| DYM | 0 639317736 | 0 072977242 | 5 07E-124 | 0 013381306 | 1 44E-05 | 2 85E-123 |
| ISM1 | 0 061677866 | 0 072945275 | 6 49E-124 | 0 473962115 | 0 | 3 64E-123 |
| NEK7 | 0 566008643 | 0 072917645 | 8 04E-124 | -0 02893252 | 6 61E-21 | 4 51E-123 |
| LRP1B | 0 336125336 | 0 072836009 | 1 51E-123 | -0 14924808 | 0 | 8 45E-123 |
| PXDNL | 0 092583427 | 0 072819655 | 1 71E-123 | -0 38936323 | 0 | 9 58E-123 |
| DENND1A | 0 600121833 | 0 072743384 | 3 08E-123 | 0 149987769 | 0 | 1 72E-122 |
| OLFM3 | 0 023976319 | 0 072714264 | 3 85E-123 | 0 496151323 | 0 | 2 15E-122 |
| STAT5B | 0 375949439 | 0 072655836 | 6 04E-123 | -0 09817706 | 2 75E-223 | 3 37E-122 |
| TSHZ3 | 0 3948716 | 0 072655353 | 6 06E-123 | -0 11953894 | 0 | 3 38E-122 |
| SUPT3H | 0 554862843 | 0 072638488 | 6 90E-123 | -0 01540503 | 5 93E-07 | 3 84E-122 |
| NAB2 | 0 10250138 | 0 072608187 | 8 70E-123 | -0 37898616 | 0 | 4 85E-122 |
| ZNF235 | 0 282937694 | 0 072595591 | 9 59E-123 | -0 21346402 | 0 | 5 34E-122 |
| SEMA5A | 0 383678209 | 0 072554438 | 1 32E-122 | -0 10015641 | 2 43E-232 | 7 31E-122 |
| APBA1 | 0 220441263 | 0 072539663 | 1 47E-122 | -0 23479692 | 0 | 8 18E-122 |
| ZBTB20 | 0 925815233 | 0 072520912 | 1 70E-122 | 0 063811295 | 3 18E-95 | 9 45E-122 |
| LINC00922 | 0 1052331 | 0 072506865 | 1 89E-122 | -0 36798488 | 0 | 1 05E-121 |
| ZNF254 | 0 344120614 | 0 072496706 | 2 05E-122 | 0 273250219 | 0 | 1 14E-121 |
| PRSS35 | 0 027954922 | 0 072479932 | 2 33E-122 | 0 488021226 | 0 | 1 29E-121 |
| ANXA6 | 0 255515791 | 0 072389469 | 4 66E-122 | -0 20973548 | 0 | 2 58E-121 |
| TBCD | 0 41563077 | 0 072375915 | 5 17E-122 | 0 200799894 | 0 | 2 86E-121 |
| SLC25A29 | 0 161542708 | 0 072352959 | 6 16E-122 | -0 29692913 | 0 | 3 41E-121 |
| ANO4 | 0 129161828 | 0 072346571 | 6 47E-122 | 0 432146208 | 0 | 3 58E-121 |
| STEAP4 | 0 06565647 | 0 072296613 | 9 49E-122 | 0 482228894 | 0 | 5 24E-121 |

| | | | | | | |
|---|---|---|---|---|---|---|
| SLC13A3 | 0 161361863 | 0 072289544 | 1 00E-121 | -0 2838498 | 0 | 5 53E-121 |
| LINC00649 | 0 049190002 | 0 072281383 | 1 07E-121 | -0 4338309 | 0 | 5 88E-121 |
| ZC3H13 | 0 424539796 | 0 072217898 | 1 73E-121 | -0 10092268 | 6 78E-236 | 9 54E-121 |
| SNHG31 | 0 102634635 | 0 072216931 | 1 74E-121 | 0 389306912 | 0 | 9 61E-121 |
| TES | 0 260227294 | 0 072206988 | 1 88E-121 | 0 28516942 | 0 | 1 04E-120 |
| RPAP2 | 0 43660886 | 0 072056142 | 5 95E-121 | -0 09754426 | 1 97E-220 | 3 27E-120 |
| KCNMB4 | 0 22387733 | 0 072030724 | 7 22E-121 | 0 385429524 | 0 | 3 97E-120 |
| PPP1R13B | 0 22592374 | 0 072000965 | 9 06E-121 | -0 24056362 | 0 | 4 97E-120 |
| TTC21B | 0 470074813 | 0 0719685 | 1 16E-120 | 0 196887391 | 0 | 6 36E-120 |
| MYCBP2-AS1 | 0 186956273 | 0 071945778 | 1 38E-120 | 0 360863312 | 0 | 7 56E-120 |
| ITFG1 | 0 660819326 | 0 07194181 | 1 42E-120 | 0 022678651 | 1 96E-13 | 7 79E-120 |
| AVL9 | 0 567207934 | 0 071931954 | 1 53E-120 | -0 03382423 | 5 53E-28 | 8 39E-120 |
| CSMD3 | 0 200205593 | 0 071927845 | 1 58E-120 | -0 2736889 | 0 | 8 65E-120 |
| BRAF | 0 706611334 | 0 071915144 | 1 74E-120 | 0 026233837 | 1 82E-17 | 9 52E-120 |
| ST7 | 0 273962041 | 0 071911811 | 1 79E-120 | 0 297451841 | 0 | 9 76E-120 |
| TAX1BP1 | 0 576250214 | 0 071907696 | 1 84E-120 | -0 01109081 | 0 000324439 | 1 01E-119 |
| SLC18B1 | 0 098475186 | 0 07190373 | 1 90E-120 | -0 37769504 | 0 | 1 04E-119 |
| CHIC2 | 0 381850717 | 0 071896751 | 2 00E-120 | -0 1313653 | 0 | 1 09E-119 |
| DDHD1 | 0 393177362 | 0 071886303 | 2 17E-120 | -0 10937191 | 6 41E-277 | 1 18E-119 |
| LINC02408 | 0 056128762 | 0 071881198 | 2 25E-120 | 0 462642953 | 0 | 1 23E-119 |
| TMEM63A | 0 173078753 | 0 071880817 | 2 26E-120 | -0 29606079 | 0 | 1 23E-119 |
| NEK1 | 0 496640079 | 0 071873595 | 2 39E-120 | 0 199687796 | 0 | 1 30E-119 |
| TCF20 | 0 273162514 | 0 071867476 | 2 50E-120 | 0 285692709 | 0 | 1 36E-119 |
| SLC35E3 | 0 328091984 | 0 071841077 | 3 06E-120 | 0 260089992 | 0 | 1 66E-119 |
| CEMIP2 | 0 216738688 | 0 071835456 | 3 19E-120 | -0 26029261 | 0 | 1 73E-119 |
| ABHD18 | 0 279139936 | 0 071713292 | 8 07E-120 | -0 20016299 | 0 | 4 38E-119 |
| PROK2 | 0 008233234 | 0 071656397 | 1 24E-119 | 0 500600022 | 0 | 6 75E-119 |
| SGIP1 | 0 194846852 | 0 071653543 | 1 27E-119 | -0 29135819 | 0 | 6 89E-119 |
| UBASH3B | 0 310825988 | 0 071649604 | 1 31E-119 | 0 277516318 | 0 | 7 09E-119 |
| LEF1-AS1 | 0 090346652 | 0 071563214 | 2 52E-119 | -0 37231048 | 0 | 1 36E-118 |
| RAB27A | 0 238440159 | 0 071540825 | 2 98E-119 | 0 318037229 | 0 | 1 61E-118 |
| DISP1 | 0 480877958 | 0 071538188 | 3 04E-119 | -0 05301717 | 2 82E-66 | 1 65E-118 |
| PRKD3 | 0 475890427 | 0 071508195 | 3 82E-119 | 0 196912193 | 0 | 2 06E-118 |
| HS6ST3 | 0 039367231 | 0 071484754 | 4 56E-119 | 0 475610942 | 0 | 2 46E-118 |
| BCAS3 | 0 753945289 | 0 071482088 | 4 65E-119 | 0 022973152 | 9 53E-14 | 2 51E-118 |
| FILIP1 | 0 109354476 | 0 071477614 | 4 81E-119 | -0 37525428 | 0 | 2 60E-118 |
| METTL16 | 0 323094934 | 0 071462641 | 5 39E-119 | -0 17144744 | 0 | 2 91E-118 |
| UNKL | 0 150139917 | 0 071449114 | 5 97E-119 | -0 32571569 | 0 | 3 22E-118 |
| FSTL1 | 0 607736384 | 0 071400148 | 8 64E-119 | 0 014406111 | 3 02E-06 | 4 66E-118 |
| UHRF1BP1 | 0 101444861 | 0 071399649 | 8 67E-119 | 0 419268678 | 0 | 4 67E-118 |
| FRMPD3 | 0 042013287 | 0 071370367 | 1 08E-118 | 0 484201099 | 0 | 5 82E-118 |
| CYTH1 | 0 370485999 | 0 071344203 | 1 32E-118 | 0 23674778 | 0 | 7 09E-118 |
| ATP6V1H | 0 353572176 | 0 071323831 | 1 54E-118 | 0 283279053 | 0 | 8 26E-118 |
| KIDINS220 | 0 560992557 | 0 071237455 | 2 95E-118 | -0 03132826 | 3 09E-24 | 1 58E-117 |

| | | | | | | |
|---|---|---|---|---|---|---|
| PSTPIP2 | 0 129285565 | 0 071235316 | 3 00E-118 | -0 3481219 | 0 | 1 61E-117 |
| SSBP2 | 0 782966249 | 0 071184491 | 4 39E-118 | 0 03829455 | 2 12E-35 | 2 36E-117 |
| LONP2 | 0 538539148 | 0 071174903 | 4 72E-118 | -0 01901393 | 7 11E-10 | 2 53E-117 |
| CRLF3 | 0 376511013 | 0 071165098 | 5 09E-118 | 0 216395656 | 0 | 2 73E-117 |
| LINC00499 | 0 043126916 | 0 071164276 | 5 12E-118 | 0 478858763 | 0 | 2 74E-117 |
| SPATA5 | 0 419019246 | 0 071109944 | 7 70E-118 | 0 180929997 | 0 | 4 12E-117 |
| ZNF804A | 0 288762826 | 0 071064778 | 1 08E-117 | 0 191918248 | 0 | 5 78E-117 |
| TYW1B | 0 347642345 | 0 071056594 | 1 15E-117 | -0 1543558 | 0 | 6 15E-117 |
| MYO1F | 0 113723325 | 0 07101456 | 1 58E-117 | -0 35712224 | 0 | 8 42E-117 |
| KAT2A | 0 149321353 | 0 070985786 | 1 96E-117 | -0 32001133 | 0 | 1 04E-116 |
| LINC02301 | 0 051712322 | 0 070980539 | 2 04E-117 | 0 474663841 | 0 | 1 09E-116 |
| GARNL3 | 0 306447621 | 0 070977072 | 2 09E-117 | -0 18585728 | 0 | 1 11E-116 |
| MTBP | 0 212588757 | 0 070977067 | 2 09E-117 | 0 309330876 | 0 | 1 11E-116 |
| NEXN | 0 219337153 | 0 070971023 | 2 19E-117 | -0 26548892 | 0 | 1 16E-116 |
| FAM81A | 0 093611391 | 0 070934414 | 2 88E-117 | 0 436686771 | 0 | 1 53E-116 |
| TAF15 | 0 577392397 | 0 070917817 | 3 26E-117 | 0 159136851 | 0 | 1 73E-116 |
| RBBP6 | 0 425167996 | 0 070898711 | 3 76E-117 | -0 08936406 | 3 32E-185 | 2 00E-116 |
| POLB | 0 149121471 | 0 070835928 | 6 03E-117 | 0 408503413 | 0 | 3 20E-116 |
| C3orf62 | 0 092050408 | 0 070834263 | 6 10E-117 | 0 446510418 | 0 | 3 23E-116 |
| GAS6 | 0 187632065 | 0 070832817 | 6 17E-117 | -0 2947783 | 0 | 3 27E-116 |
| SON | 0 786002551 | 0 070802318 | 7 75E-117 | 0 074072271 | 1 01E-127 | 4 10E-116 |
| ARID4B | 0 718290152 | 0 070801583 | 7 80E-117 | 0 026705545 | 4 82E-18 | 4 12E-116 |
| SPAG16 | 0 361481792 | 0 070777931 | 9 31E-117 | -0 1330695 | 0 | 4 92E-116 |
| RPRD2 | 0 511288572 | 0 070763166 | 1 04E-116 | 0 172370567 | 0 | 5 49E-116 |
| RORB | 0 11485599 | 0 070722122 | 1 41E-116 | 0 397591654 | 0 | 7 46E-116 |
| CRTAC1 | 0 045201881 | 0 070657378 | 2 29E-116 | 0 480926393 | 0 | 1 21E-115 |
| LINC00937 | 0 12504997 | 0 070654962 | 2 34E-116 | 0 386656385 | 0 | 1 23E-115 |
| SH3PXD2B | 0 455683311 | 0 070626102 | 2 90E-116 | -0 08835336 | 4 53E-181 | 1 53E-115 |
| GSK3B | 0 60791723 | 0 070602947 | 3 44E-116 | 0 130512315 | 0 | 1 81E-115 |
| DPH6 | 0 450648189 | 0 07056065 | 4 72E-116 | -0 07772298 | 1 86E-140 | 2 49E-115 |
| ORC2 | 0 361481792 | 0 070549644 | 5 13E-116 | 0 258173842 | 0 | 2 70E-115 |
| PAXBP1 | 0 515647903 | 0 070530598 | 5 91E-116 | 0 158971285 | 0 | 3 11E-115 |
| TLL1 | 0 116226609 | 0 070516722 | 6 56E-116 | 0 425647556 | 0 | 3 44E-115 |
| USP34 | 0 824941463 | 0 070493763 | 7 78E-116 | 0 05416053 | 4 34E-69 | 4 08E-115 |
| BBX | 0 656726504 | 0 070492684 | 7 84E-116 | -0 01144592 | 0 000207199 | 4 11E-115 |
| HNRNPLL | 0 346623898 | 0 070478055 | 8 75E-116 | -0 14567931 | 0 | 4 59E-115 |
| PRKAR1B | 0 170832461 | 0 070464814 | 9 65E-116 | 0 403995171 | 0 | 5 06E-115 |
| SPAST | 0 290114409 | 0 070412839 | 1 42E-115 | -0 20519023 | 0 | 7 44E-115 |
| FSCN1 | 0 186118673 | 0 070399792 | 1 57E-115 | -0 29719733 | 0 | 8 20E-115 |
| NSD1 | 0 62683939 | 0 070365753 | 2 02E-115 | -0 00829534 | 0 00717085 | 1 06E-114 |
| G2E3 | 0 340437075 | 0 070343679 | 2 38E-115 | 0 249601167 | 0 | 1 24E-114 |
| DCAF6 | 0 570872437 | 0 070296311 | 3 38E-115 | -0 02143255 | 3 71E-12 | 1 77E-114 |
| CLDN15 | 0 07726866 | 0 070235488 | 5 32E-115 | 0 454684624 | 0 | 2 77E-114 |
| SLF2 | 0 505844168 | 0 070235383 | 5 32E-115 | 0 184194074 | 0 | 2 77E-114 |

| | | | | | | |
|---|---|---|---|---|---|---|
| TMEM67 | 0 281843102 | 0 070231255 | 5 49E-115 | -0 20237902 | 0 | 2 86E-114 |
| PMS1 | 0 375740039 | 0 070218845 | 6 02E-115 | 0 23315876 | 0 | 3 13E-114 |
| SLIT1 | 0 042070397 | 0 070200465 | 6 90E-115 | 0 488068485 | 0 | 3 59E-114 |
| LINC00377 | 0 257961965 | 0 070187652 | 7 59E-115 | -0 24495948 | 0 | 3 94E-114 |
| MALT1 | 0 2785974 | 0 070179766 | 8 05E-115 | 0 2849305 | 0 | 4 18E-114 |
| HIBCH | 0 58068569 | 0 070146388 | 1 03E-114 | -0 01938054 | 3 33E-10 | 5 35E-114 |
| THBS4 | 0 447040795 | 0 070119928 | 1 25E-114 | -0 09523345 | 3 66E-210 | 6 51E-114 |
| PPP2R5C | 0 451809408 | 0 070061172 | 1 94E-114 | 0 21493066 | 0 | 1 00E-113 |
| SOS1 | 0 638080372 | 0 070058499 | 1 98E-114 | -0 00624299 | 0 043016176 | 1 02E-113 |
| KPNA5 | 0 284365422 | 0 070055609 | 2 02E-114 | -0 18378966 | 0 | 1 05E-113 |
| D2HGDH | 0 222687556 | 0 070033552 | 2 38E-114 | -0 2572615 | 0 | 1 23E-113 |
| RIC8B | 0 366259923 | 0 070032141 | 2 40E-114 | 0 247923983 | 0 | 1 24E-113 |
| PRKCQ | 0 037882393 | 0 070017085 | 2 69E-114 | 0 487235732 | 0 | 1 39E-113 |
| COMMD10 | 0 533189926 | 0 069968578 | 3 85E-114 | 0 177086012 | 0 | 1 99E-113 |
| PALD1 | 0 063410177 | 0 069962636 | 4 02E-114 | -0 41310595 | 0 | 2 07E-113 |
| RHOXF1-AS1 | 0 044507053 | 0 069918256 | 5 59E-114 | -0 43983738 | 0 | 2 88E-113 |
| CDCA7L | 0 076697569 | 0 069911437 | 5 88E-114 | 0 427979211 | 0 | 3 02E-113 |
| CEP95 | 0 522986427 | 0 069907573 | 6 05E-114 | 0 157529754 | 0 | 3 11E-113 |
| CCDC68 | 0 099284232 | 0 069896236 | 6 57E-114 | -0 37279515 | 0 | 3 38E-113 |
| CCDC14 | 0 458148522 | 0 069894913 | 6 64E-114 | 0 208079709 | 0 | 3 41E-113 |
| DIS3L2 | 0 508690107 | 0 069883631 | 7 22E-114 | 0 181559923 | 0 | 3 70E-113 |
| LINC00923 | 0 083607774 | 0 069867465 | 8 13E-114 | 0 457612112 | 0 | 4 17E-113 |
| TLN2 | 0 580885572 | 0 069860719 | 8 55E-114 | -0 01151402 | 0 000189853 | 4 38E-113 |
| BANP | 0 223496602 | 0 069828986 | 1 08E-113 | 0 349585699 | 0 | 5 54E-113 |
| RAD18 | 0 225885668 | 0 069798468 | 1 35E-113 | 0 302340099 | 0 | 6 93E-113 |
| RGS20 | 0 158772915 | 0 069796207 | 1 38E-113 | 0 387344339 | 0 | 7 04E-113 |
| PCBP3 | 0 226551941 | 0 06975327 | 1 89E-113 | 0 355801404 | 0 | 9 66E-113 |
| IL4R | 0 130180275 | 0 069695211 | 2 90E-113 | -0 34896699 | 0 | 1 48E-112 |
| SAP25 | 0 086158649 | 0 069669824 | 3 50E-113 | 0 451061228 | 0 | 1 79E-112 |
| CCDC7 | 0 419238164 | 0 069664859 | 3 63E-113 | -0 1000035 | 1 23E-231 | 1 85E-112 |
| CCDC66 | 0 405512935 | 0 069609677 | 5 45E-113 | 0 240260652 | 0 | 2 78E-112 |
| QPRT | 0 081713655 | 0 069606485 | 5 58E-113 | -0 39904409 | 0 | 2 84E-112 |
| LINC02607 | 0 025584893 | 0 069600222 | 5 84E-113 | -0 46204788 | 0 | 2 97E-112 |
| SPON2 | 0 360396718 | 0 069593082 | 6 15E-113 | -0 17520537 | 0 | 3 13E-112 |
| ST3GAL1 | 0 297119796 | 0 069537277 | 9 28E-113 | -0 1673525 | 0 | 4 72E-112 |
| IRS1 | 0 150501609 | 0 069524749 | 1 02E-112 | 0 402030545 | 0 | 5 17E-112 |
| SDC2 | 0 426995488 | 0 069490791 | 1 31E-112 | 0 22825596 | 0 | 6 64E-112 |
| ATP1B1 | 0 183072852 | 0 069468363 | 1 54E-112 | -0 29059802 | 0 | 7 82E-112 |
| CELF4 | 0 0468866 | 0 06946767 | 1 55E-112 | -0 43632907 | 0 | 7 86E-112 |
| RBPJ | 0 576469133 | 0 06944991 | 1 76E-112 | 0 143167612 | 0 | 8 95E-112 |
| TCEA1 | 0 428718281 | 0 0693975 | 2 59E-112 | 0 224698657 | 0 | 1 31E-111 |
| RBFOX1 | 0 202023567 | 0 069374933 | 3 06E-112 | 0 357586965 | 0 | 1 55E-111 |
| FAM184B | 0 123013078 | 0 06933898 | 3 98E-112 | -0 35690458 | 0 | 2 02E-111 |
| PHKA2 | 0 224657821 | 0 069306814 | 5 04E-112 | 0 33497179 | 0 | 2 55E-111 |

| | | | | | | |
|---|---|---|---|---|---|---|
| MSRB3 | 0 448649369 | 0 069285131 | 5 91E-112 | 0 16383328 | 0 | 2 99E-111 |
| CUL5 | 0 457710685 | 0 069277914 | 6 23E-112 | -0 06952941 | 9 83E-113 | 3 15E-111 |
| SEC24B-AS1 | 0 102482344 | 0 069250205 | 7 63E-112 | -0 36553152 | 0 | 3 86E-111 |
| GABRA2 | 0 054938988 | 0 069243108 | 8 04E-112 | 0 473175483 | 0 | 4 06E-111 |
| GPHN | 0 751242124 | 0 069239556 | 8 25E-112 | 0 065622091 | 1 32E-100 | 4 17E-111 |
| CDC73 | 0 527954922 | 0 069232705 | 8 68E-112 | -0 0413689 | 4 97E-41 | 4 38E-111 |
| PPP3R1 | 0 464497154 | 0 069196986 | 1 13E-111 | 0 194925925 | 0 | 5 68E-111 |
| NR2F2-AS1 | 0 636462279 | 0 06915211 | 1 56E-111 | 0 017737818 | 8 94E-09 | 7 88E-111 |
| GLG1 | 0 636443243 | 0 069150535 | 1 58E-111 | 0 007655806 | 0 013082932 | 7 97E-111 |
| ALDH2 | 0 175905656 | 0 069128434 | 1 86E-111 | -0 29583816 | 0 | 9 36E-111 |
| ASIC2 | 0 078943862 | 0 069071792 | 2 81E-111 | -0 39014309 | 0 | 1 42E-110 |
| KCNIP1 | 0 205116979 | 0 069025328 | 3 95E-111 | -0 25505639 | 0 | 1 99E-110 |
| TASOR2 | 0 540490377 | 0 068973366 | 5 77E-111 | -0 03217915 | 1 75E-25 | 2 90E-110 |
| EXOC3L2 | 0 027022139 | 0 068936401 | 7 56E-111 | -0 45878562 | 0 | 3 79E-110 |
| IGF2 | 0 15801146 | 0 06886218 | 1 30E-110 | -0 32469105 | 0 | 6 51E-110 |
| ARIH1 | 0 670489806 | 0 068854788 | 1 37E-110 | 0 015919915 | 2 46E-07 | 6 87E-110 |
| CARF | 0 229502579 | 0 068837903 | 1 55E-110 | 0 368275784 | 0 | 7 76E-110 |
| CRACD | 0 120757267 | 0 068759073 | 2 75E-110 | 0 429621445 | 0 | 1 38E-109 |
| MIOS | 0 269164874 | 0 068752916 | 2 88E-110 | 0 319505131 | 0 | 1 44E-109 |
| SLC7A11 | 0 061696903 | 0 06873385 | 3 30E-110 | 0 46417851 | 0 | 1 65E-109 |
| SEMA6A-AS1 | 0 16920485 | 0 068712283 | 3 86E-110 | -0 31245935 | 0 | 1 93E-109 |
| JDP2 | 0 133920923 | 0 068701294 | 4 19E-110 | -0 33800063 | 0 | 2 09E-109 |
| YTHDF3 | 0 448097314 | 0 068684462 | 4 73E-110 | 0 186619891 | 0 | 2 36E-109 |
| JAZF1-AS1 | 0 023690773 | 0 068656193 | 5 81E-110 | -0 46216256 | 0 | 2 90E-109 |
| CTDSPL | 0 516970931 | 0 068645139 | 6 29E-110 | -0 03085621 | 1 47E-23 | 3 14E-109 |
| NCK2 | 0 399402258 | 0 06858842 | 9 50E-110 | -0 12520262 | 0 | 4 73E-109 |
| NKAIN3 | 0 190154385 | 0 068549463 | 1 26E-109 | 0 416022026 | 0 | 6 27E-109 |
| KPNA3 | 0 379328397 | 0 068505712 | 1 73E-109 | -0 13283944 | 0 | 8 60E-109 |
| RGPD2 | 0 03127677 | 0 06847957 | 2 09E-109 | 0 488504713 | 0 | 1 04E-108 |
| RNF213-AS1 | 0 293074565 | 0 06847024 | 2 24E-109 | 0 291203484 | 0 | 1 11E-108 |
| ATG7 | 0 586634559 | 0 06846161 | 2 38E-109 | -0 04030656 | 4 90E-39 | 1 18E-108 |
| CCDC148 | 0 113723325 | 0 06846046 | 2 40E-109 | 0 423538498 | 0 | 1 19E-108 |
| FBXL17 | 0 691448859 | 0 068455498 | 2 49E-109 | 0 038091007 | 4 82E-35 | 1 23E-108 |
| SRRM2 | 0 689250157 | 0 068438009 | 2 83E-109 | 0 117677484 | 0 | 1 40E-108 |
| CCDC200 | 0 121842341 | 0 068426925 | 3 06E-109 | 0 416238897 | 0 | 1 52E-108 |
| ATP11C | 0 332851078 | 0 0684223 | 3 17E-109 | -0 1501658 | 0 | 1 57E-108 |
| GNAS | 0 591698235 | 0 068412299 | 3 40E-109 | -0 01356103 | 1 10E-05 | 1 68E-108 |
| RBM41 | 0 246207002 | 0 06841216 | 3 41E-109 | 0 347264851 | 0 | 1 68E-108 |
| SLC12A7 | 0 09362091 | 0 068400457 | 3 71E-109 | 0 454536381 | 0 | 1 83E-108 |
| SLC12A8 | 0 051483886 | 0 06839666 | 3 81E-109 | 0 472403707 | 0 | 1 88E-108 |
| DCAF5 | 0 446564885 | 0 068376347 | 4 41E-109 | -0 10208501 | 2 44E-241 | 2 18E-108 |
| NEDD4 | 0 380670461 | 0 068373304 | 4 51E-109 | 0 208475432 | 0 | 2 23E-108 |
| NPAS2 | 0 338219337 | 0 068368112 | 4 69E-109 | -0 17231081 | 0 | 2 31E-108 |
| STRN3 | 0 624678761 | 0 068352921 | 5 23E-109 | 0 000879402 | 0 775613815 | 2 58E-108 |

| | | | | | | |
|---|---|---|---|---|---|---|
| LUC7L3 | 0 839932611 | 0 068307939 | 7 24E-109 | 0 054912684 | 5 68E-71 | 3 56E-108 |
| PDLIM5 | 0 488825646 | 0 068254061 | 1 07E-108 | 0 193760389 | 0 | 5 26E-108 |
| DACT1 | 0 042403533 | 0 068247637 | 1 12E-108 | 0 480888972 | 0 | 5 50E-108 |
| CDH2 | 0 12000533 | 0 068233019 | 1 24E-108 | 0 446897 | 0 | 6 11E-108 |
| ZNF131 | 0 490339038 | 0 068222596 | 1 34E-108 | 0 179502707 | 0 | 6 59E-108 |
| NISCH | 0 341807695 | 0 068195595 | 1 63E-108 | -0 15804964 | 0 | 8 00E-108 |
| NGF | 0 148207725 | 0 068188702 | 1 71E-108 | -0 30561975 | 0 | 8 40E-108 |
| SFRP1 | 0 106832156 | 0 06818703 | 1 73E-108 | -0 36285464 | 0 | 8 50E-108 |
| RNF2 | 0 18518589 | 0 0681178 | 2 85E-108 | 0 361930519 | 0 | 1 40E-107 |
| GOLIM4 | 0 474738726 | 0 068102049 | 3 20E-108 | 0 188507705 | 0 | 1 56E-107 |
| NR6A1 | 0 299318498 | 0 068081758 | 3 70E-108 | 0 269112168 | 0 | 1 81E-107 |
| MRPS6 | 0 348965373 | 0 068055054 | 4 48E-108 | -0 17044394 | 0 | 2 19E-107 |
| PHLDB2 | 0 371295045 | 0 068024178 | 5 60E-108 | -0 13673877 | 0 | 2 73E-107 |
| CNIH3 | 0 32700691 | 0 068023733 | 5 62E-108 | -0 16034919 | 0 | 2 74E-107 |
| LINC02798 | 0 087376977 | 0 068010745 | 6 17E-108 | 0 45090409 | 0 | 3 01E-107 |
| TMSB15B-AS1 | 0 103338981 | 0 06800729 | 6 32E-108 | 0 448071823 | 0 | 3 08E-107 |
| MTSS1 | 0 396727647 | 0 067960905 | 8 82E-108 | -0 09923764 | 4 08E-228 | 4 30E-107 |
| ADCY3 | 0 287830043 | 0 067918533 | 1 20E-107 | 0 283973709 | 0 | 5 82E-107 |
| LINC01500 | 0 057156727 | 0 067915141 | 1 23E-107 | -0 42726362 | 0 | 5 96E-107 |
| HDAC2-AS2 | 0 135443833 | 0 067902817 | 1 34E-107 | 0 424321113 | 0 | 6 51E-107 |
| FBXO11 | 0 683786716 | 0 067888716 | 1 48E-107 | 0 097141954 | 1 26E-218 | 7 20E-107 |
| HFM1 | 0 097637585 | 0 06787115 | 1 68E-107 | 0 424588505 | 0 | 8 16E-107 |
| RYR1 | 0 062058594 | 0 067866739 | 1 74E-107 | -0 40836954 | 0 | 8 41E-107 |
| CD200 | 0 115436599 | 0 067861634 | 1 80E-107 | -0 35368512 | 0 | 8 72E-107 |
| PI4KA | 0 446336449 | 0 067859099 | 1 83E-107 | 0 199641644 | 0 | 8 88E-107 |
| NSL1 | 0 264900725 | 0 067855245 | 1 88E-107 | 0 3325999 | 0 | 9 12E-107 |
| RNF122 | 0 04152786 | 0 067848735 | 1 97E-107 | 0 479457095 | 0 | 9 56E-107 |
| BMP7 | 0 130313529 | 0 067802433 | 2 75E-107 | -0 34470655 | 0 | 1 33E-106 |
| CHD2 | 0 696503017 | 0 067778121 | 3 28E-107 | 0 10248676 | 3 09E-243 | 1 58E-106 |
| CC2D2B | 0 106613238 | 0 067672058 | 7 01E-107 | 0 432551976 | 0 | 3 38E-106 |
| CCDC141 | 0 02050218 | 0 067651809 | 8 10E-107 | 0 497927963 | 0 | 3 90E-106 |
| ZMYND11 | 0 44349051 | 0 067647196 | 8 37E-107 | -0 07443809 | 5 71E-129 | 4 03E-106 |
| MTUS1 | 0 417353563 | 0 067645343 | 8 48E-107 | -0 08187983 | 1 03E-155 | 4 08E-106 |
| MAP4K3 | 0 663094173 | 0 06762155 | 1 01E-106 | 0 012753633 | 3 57E-05 | 4 84E-106 |
| UBR1 | 0 58899507 | 0 06757806 | 1 37E-106 | -0 00826458 | 0 007387917 | 6 60E-106 |
| XPR1 | 0 447992614 | 0 067572356 | 1 43E-106 | 0 201173035 | 0 | 6 87E-106 |
| HAND2-AS1 | 0 033542099 | 0 067555368 | 1 61E-106 | 0 489035483 | 0 | 7 75E-106 |
| TRIM13 | 0 399240448 | 0 067540053 | 1 80E-106 | 0 210956342 | 0 | 8 64E-106 |
| TMEM182 | 0 082046791 | 0 067489612 | 2 58E-106 | -0 39330096 | 0 | 1 24E-105 |
| SCML2 | 0 141078601 | 0 067470274 | 2 96E-106 | 0 35477531 | 0 | 1 42E-105 |
| IGLON5 | 0 015866821 | 0 067442765 | 3 61E-106 | 0 496605904 | 0 | 1 73E-105 |
| LRCH1 | 0 50525404 | 0 067431038 | 3 92E-106 | -0 06349612 | 2 65E-94 | 1 88E-105 |
| KIAA0825 | 0 37105709 | 0 067390802 | 5 22E-106 | -0 13330735 | 0 | 2 50E-105 |
| TMEM91 | 0 215053968 | 0 067367975 | 6 14E-106 | 0 343754226 | 0 | 2 94E-105 |

| | | | | | | |
|---|---|---|---|---|---|---|
| UBE2H | 0 658182787 | 0 067365394 | 6 26E-106 | 0 017429029 | 1 61E-08 | 2 99E-105 |
| PDE4DIP | 0 357845843 | 0 06735721 | 6 63E-106 | -0 15351916 | 0 | 3 17E-105 |
| GFOD1 | 0 089927852 | 0 067341078 | 7 44E-106 | -0 38572176 | 0 | 3 55E-105 |
| PTP4A3 | 0 03679732 | 0 067340013 | 7 50E-106 | 0 487140812 | 0 | 3 58E-105 |
| CMKLR1 | 0 066960461 | 0 067323317 | 8 44E-106 | -0 41346467 | 0 | 4 03E-105 |
| CTNNB1 | 0 578553616 | 0 067311857 | 9 16E-106 | -0 01420268 | 4 15E-06 | 4 37E-105 |
| TRIQK | 0 320782014 | 0 067290202 | 1 07E-105 | -0 18232661 | 0 | 5 09E-105 |
| TNPO1 | 0 597885058 | 0 067287073 | 1 09E-105 | 0 122451335 | 0 | 5 20E-105 |
| SGO1-AS1 | 0 206725553 | 0 067278866 | 1 16E-105 | 0 360634625 | 0 | 5 51E-105 |
| GATA2 | 0 024851992 | 0 067240139 | 1 53E-105 | 0 489366599 | 0 | 7 26E-105 |
| ZRANB3 | 0 404237498 | 0 06718461 | 2 26E-105 | 0 181240146 | 0 | 1 08E-104 |
| SPOPL | 0 29137081 | 0 067172798 | 2 46E-105 | 0 286574537 | 0 | 1 17E-104 |
| DHX57 | 0 310721288 | 0 067152402 | 2 85E-105 | 0 291678842 | 0 | 1 35E-104 |
| E2F3 | 0 239458605 | 0 067151996 | 2 86E-105 | 0 324485158 | 0 | 1 36E-104 |
| PNN | 0 494412823 | 0 067129973 | 3 34E-105 | 0 185169051 | 0 | 1 58E-104 |
| COL21A1 | 0 34219794 | 0 067124741 | 3 47E-105 | -0 12381335 | 0 | 1 64E-104 |
| RAB28 | 0 44469932 | 0 067117179 | 3 66E-105 | 0 205190332 | 0 | 1 73E-104 |
| RUFY3 | 0 620633531 | 0 067061821 | 5 42E-105 | 0 001534944 | 0 618822373 | 2 56E-104 |
| DIO3OS | 0 1051284 | 0 067011837 | 7 72E-105 | -0 35095615 | 0 | 3 65E-104 |
| HMGA2-AS1 | 0 184690944 | 0 066958518 | 1 13E-104 | 0 294116078 | 0 | 5 32E-104 |
| DNMT1 | 0 307742095 | 0 066907763 | 1 61E-104 | 0 23702581 | 0 | 7 62E-104 |
| GSR | 0 170004378 | 0 066877751 | 1 99E-104 | 0 384855191 | 0 | 9 41E-104 |
| ZEB1-AS1 | 0 231082599 | 0 066874151 | 2 05E-104 | -0 22204305 | 0 | 9 65E-104 |
| DDB2 | 0 329824294 | 0 066861221 | 2 24E-104 | 0 262043423 | 0 | 1 06E-103 |
| VIPR2 | 0 072500048 | 0 066856872 | 2 31E-104 | -0 41262486 | 0 | 1 09E-103 |
| DSCAML1 | 0 043193543 | 0 066775689 | 4 10E-104 | -0 43511401 | 0 | 1 93E-103 |
| HNRNPH1 | 0 704983724 | 0 066744342 | 5 12E-104 | 0 097239011 | 4 63E-219 | 2 41E-103 |
| ABCG2 | 0 074698749 | 0 066731418 | 5 61E-104 | -0 40345584 | 0 | 2 64E-103 |
| MVB12B | 0 267813291 | 0 066710084 | 6 52E-104 | 0 333360313 | 0 | 3 06E-103 |
| MSX1 | 0 054834288 | 0 066702987 | 6 85E-104 | -0 43037653 | 0 | 3 22E-103 |
| PI15 | 0 065037787 | 0 066679206 | 8 10E-104 | 0 462182685 | 0 | 3 81E-103 |
| H19 | 0 133026213 | 0 066671016 | 8 59E-104 | -0 36029196 | 0 | 4 03E-103 |
| COMMD1 | 0 544725971 | 0 066669384 | 8 69E-104 | 0 166275262 | 0 | 4 08E-103 |
| NGF-AS1 | 0 077906379 | 0 066627899 | 1 16E-103 | -0 39182117 | 0 | 5 46E-103 |
| CEP85L | 0 559326874 | 0 066627671 | 1 17E-103 | 0 016886146 | 4 41E-08 | 5 46E-103 |
| DNAH3 | 0 046125145 | 0 066601494 | 1 40E-103 | 0 475535757 | 0 | 6 56E-103 |
| RAPGEF3 | 0 03317089 | 0 066600155 | 1 41E-103 | 0 486704064 | 0 | 6 62E-103 |
| LPGAT1 | 0 36905827 | 0 066586995 | 1 55E-103 | 0 25021064 | 0 | 7 26E-103 |
| FOXP2 | 0 706068797 | 0 066574018 | 1 70E-103 | 0 017323679 | 1 96E-08 | 7 95E-103 |
| NEXMIF | 0 098341931 | 0 066557702 | 1 91E-103 | 0 454751549 | 0 | 8 92E-103 |
| ARHGEF17 | 0 190792104 | 0 066552662 | 1 98E-103 | -0 29224352 | 0 | 9 23E-103 |
| ARGLU1 | 0 731244408 | 0 066552397 | 1 98E-103 | 0 039121827 | 7 13E-37 | 9 24E-103 |
| DLC1 | 0 84230264 | 0 066530584 | 2 31E-103 | 0 084881845 | 3 14E-167 | 1 08E-102 |
| PREP | 0 241305134 | 0 066508896 | 2 69E-103 | -0 24091009 | 0 | 1 25E-102 |

| | | | | | | |
|---|---|---|---|---|---|---|
| C9orf85 | 0 278854391 | 0 066503558 | 2 79E-103 | 0 316941444 | 0 | 1 30E-102 |
| NLGN1 | 0 731910681 | 0 066478893 | 3 32E-103 | 0 077615079 | 4 51E-140 | 1 55E-102 |
| CDH7 | 0 036330928 | 0 066473301 | 3 45E-103 | 0 489631994 | 0 | 1 61E-102 |
| PIK3C2B | 0 079591099 | 0 066434462 | 4 54E-103 | -0 39466422 | 0 | 2 11E-102 |
| AKAP9 | 0 772058404 | 0 066428291 | 4 74E-103 | 0 021880628 | 1 31E-12 | 2 20E-102 |
| CDH20 | 0 042603415 | 0 066391944 | 6 12E-103 | 0 479883195 | 0 | 2 84E-102 |
| DIP2C | 0 622165959 | 0 06638733 | 6 32E-103 | 0 177332225 | 0 | 2 93E-102 |
| REPS2 | 0 19491348 | 0 066372101 | 7 03E-103 | -0 26887463 | 0 | 3 26E-102 |
| SOX4 | 0 30629533 | 0 06632142 | 1 00E-102 | 0 278408992 | 0 | 4 65E-102 |
| ARHGEF7 | 0 465277646 | 0 066279475 | 1 35E-102 | -0 06953017 | 9 78E-113 | 6 24E-102 |
| MEF2C-AS1 | 0 186851573 | 0 066244238 | 1 72E-102 | -0 28151076 | 0 | 7 98E-102 |
| LRGUK | 0 092050408 | 0 066235315 | 1 83E-102 | -0 38708748 | 0 | 8 49E-102 |
| RNF111 | 0 520730616 | 0 066190858 | 2 50E-102 | 0 152651799 | 0 | 1 16E-101 |
| CEP135 | 0 210808856 | 0 066166552 | 2 97E-102 | 0 312317283 | 0 | 1 37E-101 |
| GLIPR1L1 | 0 105366355 | 0 066129149 | 3 86E-102 | -0 37728965 | 0 | 1 78E-101 |
| UGP2 | 0 364127848 | 0 066127274 | 3 91E-102 | 0 253137543 | 0 | 1 80E-101 |
| MFAP3 | 0 188355447 | 0 066104901 | 4 57E-102 | -0 28714074 | 0 | 2 11E-101 |
| ATXN2L | 0 296405932 | 0 066096085 | 4 86E-102 | 0 290065404 | 0 | 2 24E-101 |
| CLTCL1 | 0 149797263 | 0 066090058 | 5 07E-102 | 0 387128341 | 0 | 2 34E-101 |
| ATG10 | 0 383107118 | 0 066032333 | 7 59E-102 | 0 253912636 | 0 | 3 50E-101 |
| DEGS2 | 0 040081095 | 0 066027474 | 7 85E-102 | -0 44469684 | 0 | 3 61E-101 |
| AMOTL1 | 0 37757705 | 0 066012957 | 8 69E-102 | -0 11628706 | 0 | 4 00E-101 |
| SPG7 | 0 46961794 | 0 06593436 | 1 50E-101 | 0 187497198 | 0 | 6 91E-101 |
| SERP1 | 0 416040053 | 0 065931442 | 1 53E-101 | -0 11955566 | 0 | 7 05E-101 |
| MFSD8 | 0 18708001 | 0 065922706 | 1 63E-101 | -0 27839951 | 0 | 7 49E-101 |
| CD22 | 0 043583789 | 0 06590118 | 1 89E-101 | -0 43735624 | 0 | 8 69E-101 |
| ODF2L | 0 420170947 | 0 065888372 | 2 07E-101 | -0 0962102 | 1 79E-214 | 9 50E-101 |
| OLFML2B | 0 332993851 | 0 065843531 | 2 83E-101 | -0 14656835 | 0 | 1 30E-100 |
| WWC2 | 0 497401534 | 0 065830503 | 3 10E-101 | -0 07606476 | 1 35E-134 | 1 42E-100 |
| PICALM | 0 660895471 | 0 065795467 | 3 95E-101 | 0 029240809 | 2 55E-21 | 1 81E-100 |
| CYGB | 0 147360606 | 0 065782943 | 4 31E-101 | -0 32960932 | 0 | 1 97E-100 |
| LINC02055 | 0 042917515 | 0 065773814 | 4 60E-101 | -0 42950981 | 0 | 2 10E-100 |
| PPP2R5E | 0 501075555 | 0 065770154 | 4 72E-101 | 0 181941084 | 0 | 2 16E-100 |
| DACH2 | 0 02596562 | 0 065750682 | 5 40E-101 | 0 489889102 | 0 | 2 47E-100 |
| TSEN54 | 0 102548971 | 0 065739224 | 5 85E-101 | 0 421006531 | 0 | 2 67E-100 |
| PKIG | 0 202185376 | 0 065667815 | 9 60E-101 | -0 26530126 | 0 | 4 38E-100 |
| EOGT | 0 126982163 | 0 065620353 | 1 34E-100 | 0 416127651 | 0 | 6 09E-100 |
| SH3GL2 | 0 039881213 | 0 06561979 | 1 34E-100 | 0 479891475 | 0 | 6 11E-100 |
| CSPP1 | 0 501884601 | 0 065567237 | 1 93E-100 | 0 166582617 | 0 | 8 79E-100 |
| LINC00598 | 0 153128629 | 0 065554685 | 2 11E-100 | -0 30097298 | 0 | 9 58E-100 |
| B4GALNT3 | 0 080285926 | 0 065545791 | 2 24E-100 | -0 39417116 | 0 | 1 02E-99 |
| ZFR2 | 0 014220175 | 0 065538703 | 2 35E-100 | -0 47500491 | 0 | 1 07E-99 |
| MYO5C | 0 052207268 | 0 065511474 | 2 84E-100 | 0 471818441 | 0 | 1 29E-99 |
| DOCK6 | 0 195084807 | 0 065495585 | 3 17E-100 | -0 2709355 | 0 | 1 44E-99 |

| | | | | | | |
|---|---|---|---|---|---|---|
| LINC00159 | 0 04373608 | 0 065491566 | 3 26E-100 | 0 476844664 | 0 | 1 48E-99 |
| FIBIN | 0 088976033 | 0 065464787 | 3 92E-100 | 0 44912831 | 0 | 1 78E-99 |
| TMEM200A | 0 082751137 | 0 065431321 | 4 95E-100 | -0 37728435 | 0 | 2 24E-99 |
| ATM | 0 524461746 | 0 065429958 | 4 99E-100 | -0 0373805 | 8 24E-34 | 2 26E-99 |
| ZNF562 | 0 244569873 | 0 065425881 | 5 14E-100 | 0 340137875 | 0 | 2 33E-99 |
| TESK2 | 0 155165521 | 0 065423702 | 5 22E-100 | 0 381714189 | 0 | 2 36E-99 |
| LIG1 | 0 134225505 | 0 065406528 | 5 87E-100 | 0 35041553 | 0 | 2 66E-99 |
| MORC3 | 0 375464012 | 0 065362083 | 7 99E-100 | 0 241320389 | 0 | 3 61E-99 |
| COL9A3 | 0 137461689 | 0 065324202 | 1 04E-99 | -0 33198371 | 0 | 4 69E-99 |
| ABCC5 | 0 303220955 | 0 065309058 | 1 15E-99 | -0 17681185 | 0 | 5 20E-99 |
| RIMS1 | 0 195218062 | 0 065308575 | 1 16E-99 | -0 29704767 | 0 | 5 21E-99 |
| KDM4B | 0 360139727 | 0 06530749 | 1 16E-99 | -0 14085748 | 0 | 5 25E-99 |
| LAMA1 | 0 121394986 | 0 06526184 | 1 60E-99 | 0 434320397 | 0 | 7 19E-99 |
| STK3 | 0 7911043 | 0 06525499 | 1 67E-99 | 0 059980559 | 2 45E-84 | 7 54E-99 |
| ZNF385B | 0 110553768 | 0 065217543 | 2 17E-99 | 0 388540024 | 0 | 9 75E-99 |
| PTPRO | 0 062496431 | 0 065204846 | 2 36E-99 | 0 464351102 | 0 | 1 06E-98 |
| ARNTL | 0 038615294 | 0 065204132 | 2 38E-99 | 0 476718328 | 0 | 1 07E-98 |
| CLEC4F | 0 016637795 | 0 06519625 | 2 51E-99 | 0 496277747 | 0 | 1 13E-98 |
| CHIC1 | 0 307675468 | 0 065187283 | 2 67E-99 | -0 16857413 | 0 | 1 20E-98 |
| RMDN2-AS1 | 0 120757267 | 0 065130397 | 3 95E-99 | -0 34817642 | 0 | 1 77E-98 |
| DENND5A | 0 496107061 | 0 065130265 | 3 95E-99 | 0 199143077 | 0 | 1 77E-98 |
| SIRPA | 0 052292932 | 0 065105272 | 4 70E-99 | 0 471280904 | 0 | 2 11E-98 |
| ZNF544 | 0 338143192 | 0 065076864 | 5 71E-99 | 0 238755489 | 0 | 2 56E-98 |
| MSH6 | 0 226789895 | 0 065066718 | 6 12E-99 | 0 3047868 | 0 | 2 74E-98 |
| PTPN13 | 0 655308294 | 0 065064311 | 6 22E-99 | 0 047041789 | 1 50E-52 | 2 79E-98 |
| EEFSEC | 0 280272601 | 0 065051353 | 6 80E-99 | 0 311558146 | 0 | 3 04E-98 |
| CREM | 0 246997011 | 0 065001423 | 9 59E-99 | -0 23393605 | 0 | 4 28E-98 |
| PPM1E | 0 142677657 | 0 064965136 | 1 23E-98 | 0 401911553 | 0 | 5 49E-98 |
| EPS8 | 0 729683425 | 0 064961161 | 1 26E-98 | 0 010268409 | 0 000873542 | 5 64E-98 |
| CAMTA1 | 0 517037559 | 0 06491346 | 1 76E-98 | -0 04128278 | 7 24E-41 | 7 81E-98 |
| SFPQ | 0 66805315 | 0 064903414 | 1 88E-98 | 0 130985429 | 0 | 8 37E-98 |
| LAMA3 | 0 157230968 | 0 064878269 | 2 23E-98 | 0 366270202 | 0 | 9 93E-98 |
| FCHSD2 | 0 508756734 | 0 064861735 | 2 50E-98 | 0 130897606 | 0 | 1 11E-97 |
| PTPRR | 0 072414384 | 0 064829865 | 3 11E-98 | 0 447359426 | 0 | 1 38E-97 |
| KDM6A | 0 639498582 | 0 064811542 | 3 53E-98 | -0 00753785 | 0 014554818 | 1 57E-97 |
| LYPD6B | 0 022967391 | 0 064763977 | 4 89E-98 | 0 493181749 | 0 | 2 17E-97 |
| ZNF75A | 0 204726733 | 0 064751947 | 5 31E-98 | 0 363609718 | 0 | 2 36E-97 |
| ARHGEF2 | 0 283527822 | 0 064751852 | 5 31E-98 | 0 310574709 | 0 | 2 36E-97 |
| GREM1 | 0 02391921 | 0 06475048 | 5 36E-98 | 0 492307788 | 0 | 2 38E-97 |
| RNASEH2B | 0 262540214 | 0 064693741 | 7 91E-98 | 0 309840998 | 0 | 3 50E-97 |
| NCOR2 | 0 618625193 | 0 064682673 | 8 53E-98 | -0 00384716 | 0 212404563 | 3 78E-97 |
| SEC31A | 0 746892311 | 0 064659908 | 9 97E-98 | 0 011876873 | 0 000118228 | 4 41E-97 |
| TOPBP1 | 0 251651406 | 0 064631293 | 1 21E-97 | 0 283635472 | 0 | 5 36E-97 |
| NSUN7 | 0 076250214 | 0 064572824 | 1 81E-97 | 0 452153286 | 0 | 7 99E-97 |

| | | | | | | |
|---|---|---|---|---|---|---|
| LINC02284 | 0 033180408 | 0 06457158 | 1 82E-97 | -0 45179854 | 0 | 8 05E-97 |
| RBM33 | 0 622965487 | 0 064567913 | 1 87E-97 | 0 119441449 | 0 | 8 25E-97 |
| ZNF337-AS1 | 0 228055815 | 0 064543928 | 2 20E-97 | 0 355140165 | 0 | 9 72E-97 |
| PAX8-AS1 | 0 224867221 | 0 064537514 | 2 30E-97 | -0 22986101 | 0 | 1 01E-96 |
| OPHN1 | 0 46993204 | 0 06450438 | 2 88E-97 | -0 04967288 | 2 16E-58 | 1 27E-96 |
| IMMP1L | 0 285564714 | 0 064455792 | 4 02E-97 | 0 287746321 | 0 | 1 77E-96 |
| USP25 | 0 50273172 | 0 064434319 | 4 65E-97 | 0 158527993 | 0 | 2 05E-96 |
| LINC00910 | 0 118282538 | 0 064372104 | 7 10E-97 | 0 433061835 | 0 | 3 12E-96 |
| STX16 | 0 485237288 | 0 064345823 | 8 49E-97 | -0 07068901 | 1 81E-116 | 3 73E-96 |
| AFMID | 0 11385658 | 0 064335768 | 9 09E-97 | -0 3670444 | 0 | 3 99E-96 |
| MYOM1 | 0 072347757 | 0 064334291 | 9 18E-97 | -0 40895428 | 0 | 4 03E-96 |
| UBOX5-AS1 | 0 136652643 | 0 064326087 | 9 71E-97 | 0 40608843 | 0 | 4 26E-96 |
| GREB1 | 0 133635377 | 0 064321729 | 1 00E-96 | -0 32945392 | 0 | 4 39E-96 |
| LEMD3 | 0 36895357 | 0 064310663 | 1 08E-96 | 0 231003287 | 0 | 4 73E-96 |
| HLCS | 0 407949592 | 0 064272894 | 1 39E-96 | 0 212669225 | 0 | 6 11E-96 |
| NFYC | 0 241209952 | 0 064270279 | 1 42E-96 | 0 335872586 | 0 | 6 22E-96 |
| ICA1 | 0 149407017 | 0 064238776 | 1 76E-96 | -0 32878331 | 0 | 7 70E-96 |
| RNF24 | 0 493156422 | 0 064220369 | 1 99E-96 | -0 0909718 | 7 00E-192 | 8 72E-96 |
| EPC2 | 0 45726333 | 0 064212461 | 2 10E-96 | 0 209440869 | 0 | 9 20E-96 |
| INHBA | 0 13204584 | 0 064172593 | 2 76E-96 | -0 34233622 | 0 | 1 20E-95 |
| KLHL20 | 0 318431022 | 0 06415711 | 3 06E-96 | 0 289101386 | 0 | 1 34E-95 |
| MAP2 | 0 259494394 | 0 064139171 | 3 46E-96 | -0 21002531 | 0 | 1 51E-95 |
| ACYP2 | 0 570501228 | 0 064131558 | 3 64E-96 | -0 02485333 | 7 82E-16 | 1 59E-95 |
| CDHR3 | 0 30466772 | 0 064116668 | 4 03E-96 | 0 28836446 | 0 | 1 76E-95 |
| SLC4A4 | 0 215092041 | 0 064116362 | 4 03E-96 | -0 26410212 | 0 | 1 76E-95 |
| MYOZ3 | 0 024071501 | 0 064110521 | 4 20E-96 | 0 491982843 | 0 | 1 83E-95 |
| SPATA6 | 0 563848014 | 0 064080426 | 5 15E-96 | -0 0218416 | 1 44E-12 | 2 24E-95 |
| LINC01640 | 0 027907331 | 0 06407799 | 5 23E-96 | -0 46032482 | 0 | 2 28E-95 |
| IBTK | 0 468875521 | 0 06405503 | 6 11E-96 | -0 06095436 | 4 84E-87 | 2 66E-95 |
| TNFRSF10D | 0 106194438 | 0 064050801 | 6 29E-96 | -0 36682592 | 0 | 2 73E-95 |
| GRAMD1B | 0 126411072 | 0 064048541 | 6 39E-96 | 0 433091073 | 0 | 2 78E-95 |
| BAIAP2L1 | 0 087434087 | 0 064034156 | 7 04E-96 | 0 449135903 | 0 | 3 06E-95 |
| LINC01776 | 0 076269251 | 0 0639944 | 9 22E-96 | 0 429594176 | 0 | 4 00E-95 |
| ARNT2 | 0 218708953 | 0 063936711 | 1 36E-95 | -0 24426289 | 0 | 5 91E-95 |
| WAKMAR2 | 0 042955588 | 0 063913306 | 1 59E-95 | 0 482633831 | 0 | 6 92E-95 |
| EML1 | 0 263292151 | 0 063909506 | 1 64E-95 | -0 21631586 | 0 | 7 09E-95 |
| BRWD3 | 0 386343302 | 0 063887436 | 1 90E-95 | -0 13191725 | 0 | 8 23E-95 |
| LINC02328 | 0 375797148 | 0 063882102 | 1 97E-95 | -0 1420628 | 0 | 8 52E-95 |
| ZNF266 | 0 179903295 | 0 06386106 | 2 27E-95 | 0 370264837 | 0 | 9 82E-95 |
| DUSP2 | 0 053101978 | 0 063844113 | 2 54E-95 | -0 42889302 | 0 | 1 10E-94 |
| BCL9L | 0 080761836 | 0 063738572 | 5 19E-95 | -0 39275452 | 0 | 2 24E-94 |
| RAB1A | 0 679969923 | 0 063687125 | 7 33E-95 | 0 008767176 | 0 00448668 | 3 16E-94 |
| TOGARAM1 | 0 384401591 | 0 063620601 | 1 15E-94 | -0 11805071 | 0 | 4 95E-94 |
| MTO1 | 0 139603282 | 0 063601747 | 1 30E-94 | -0 33379371 | 0 | 5 61E-94 |

| | | | | | | |
|---|---|---|---|---|---|---|
| SLC39A11 | 0 520797244 | 0 063586601 | 1 44E-94 | 0 135478328 | 0 | 6 21E-94 |
| JAM3 | 0 406464754 | 0 063561058 | 1 71E-94 | -0 08788374 | 3 64E-179 | 7 37E-94 |
| CD37 | 0 045477908 | 0 063554016 | 1 80E-94 | 0 473077835 | 0 | 7 72E-94 |
| CYP26A1 | 0 017570577 | 0 063516724 | 2 31E-94 | 0 493686067 | 0 | 9 91E-94 |
| CACNA2D2 | 0 019540843 | 0 063492591 | 2 71E-94 | 0 495791358 | 0 | 1 16E-93 |
| MORC4 | 0 290076336 | 0 06348313 | 2 89E-94 | 0 311081362 | 0 | 1 24E-93 |
| GNAQ | 0 674125754 | 0 063469485 | 3 17E-94 | 0 099118332 | 1 43E-227 | 1 36E-93 |
| MAST3 | 0 064419105 | 0 06343095 | 4 10E-94 | -0 41379554 | 0 | 1 76E-93 |
| PRKACB | 0 384239782 | 0 063421576 | 4 37E-94 | 0 244672407 | 0 | 1 87E-93 |
| LINC02432 | 0 07958158 | 0 063372318 | 6 08E-94 | 0 445891519 | 0 | 2 60E-93 |
| ENGASE | 0 104452609 | 0 063343352 | 7 38E-94 | -0 37316031 | 0 | 3 16E-93 |
| ANKUB1 | 0 425929451 | 0 063341893 | 7 45E-94 | 0 19883902 | 0 | 3 18E-93 |
| CYTH3 | 0 563495841 | 0 063328895 | 8 13E-94 | -0 00971663 | 0 001635413 | 3 47E-93 |
| PURPL | 0 112086197 | 0 063295884 | 1 01E-93 | -0 35972584 | 0 | 4 33E-93 |
| AGL | 0 182920561 | 0 063235178 | 1 52E-93 | -0 28454041 | 0 | 6 50E-93 |
| ZNF7 | 0 154232739 | 0 063221558 | 1 67E-93 | 0 394007606 | 0 | 7 11E-93 |
| NME7 | 0 38572462 | 0 063208595 | 1 82E-93 | -0 11849168 | 0 | 7 75E-93 |
| MIR3681HG | 0 163141764 | 0 06317817 | 2 23E-93 | 0 335005255 | 0 | 9 49E-93 |
| AFF1 | 0 588604824 | 0 063075114 | 4 43E-93 | -0 00456136 | 0 139280868 | 1 89E-92 |
| ULK2 | 0 297338714 | 0 063074444 | 4 45E-93 | 0 320930396 | 0 | 1 90E-92 |
| MFGE8 | 0 374712075 | 0 063052107 | 5 17E-93 | -0 11627262 | 0 | 2 20E-92 |
| FAM193A | 0 544278616 | 0 063051217 | 5 20E-93 | 0 177918916 | 0 | 2 21E-92 |
| CNTNAP3B | 0 209561973 | 0 063046345 | 5 37E-93 | 0 359692125 | 0 | 2 28E-92 |
| CACNA1D | 0 133368868 | 0 0630292 | 6 02E-93 | -0 33065568 | 0 | 2 56E-92 |
| RFPL1S | 0 042184615 | 0 063017554 | 6 50E-93 | -0 43960514 | 0 | 2 76E-92 |
| CPS1 | 0 094287183 | 0 06301025 | 6 83E-93 | 0 40437098 | 0 | 2 90E-92 |
| PPP4R3A | 0 414812206 | 0 062964328 | 9 27E-93 | -0 11372448 | 2 28E-299 | 3 93E-92 |
| NUBPL | 0 47103615 | 0 062935464 | 1 12E-92 | 0 205430164 | 0 | 4 76E-92 |
| FOXK1 | 0 326987874 | 0 0629311 | 1 16E-92 | -0 179128 | 0 | 4 90E-92 |
| RASA4B | 0 034788982 | 0 062928725 | 1 17E-92 | 0 481091371 | 0 | 4 97E-92 |
| GOLGA4 | 0 72619025 | 0 062926098 | 1 20E-92 | 0 016845647 | 4 75E-08 | 5 06E-92 |
| PLEKHA4 | 0 288087034 | 0 062922929 | 1 22E-92 | -0 18910783 | 0 | 5 17E-92 |
| FGFR1OP2 | 0 32385639 | 0 062865595 | 1 79E-92 | -0 17356354 | 0 | 7 56E-92 |
| SLC47A1 | 0 044288135 | 0 062863591 | 1 81E-92 | 0 48454413 | 0 | 7 65E-92 |
| ZNF609 | 0 634929851 | 0 062847057 | 2 02E-92 | 0 121471821 | 0 | 8 54E-92 |
| AGBL3 | 0 213350212 | 0 062839088 | 2 13E-92 | 0 353031789 | 0 | 9 00E-92 |
| MCM6 | 0 121747159 | 0 062838054 | 2 15E-92 | 0 355276252 | 0 | 9 06E-92 |
| TRAF5 | 0 230844644 | 0 062820324 | 2 42E-92 | -0 25152548 | 0 | 1 02E-91 |
| AGPAT3 | 0 330871295 | 0 062771041 | 3 35E-92 | 0 257834035 | 0 | 1 41E-91 |
| SLC27A3 | 0 054139461 | 0 062760401 | 3 60E-92 | 0 477821469 | 0 | 1 51E-91 |
| C7orf31 | 0 074965259 | 0 062738897 | 4 15E-92 | -0 40173926 | 0 | 1 74E-91 |
| ATR | 0 50893758 | 0 062732396 | 4 33E-92 | 0 174887405 | 0 | 1 82E-91 |
| FAM102B | 0 10812663 | 0 062711349 | 4 98E-92 | 0 385735978 | 0 | 2 09E-91 |
| TP53I11 | 0 079334107 | 0 062694751 | 5 56E-92 | 0 455797965 | 0 | 2 33E-91 |

| | | | | | | |
|---|---|---|---|---|---|---|
| ANKS1B | 0 38199349 | 0 062660057 | 6 99E-92 | -0 11197602 | 3 04E-290 | 2 94E-91 |
| MOB3B | 0 136024443 | 0 062633901 | 8 32E-92 | -0 3369561 | 0 | 3 49E-91 |
| PDE1A | 0 242361653 | 0 06261369 | 9 51E-92 | -0 26019085 | 0 | 3 99E-91 |
| ZCWPW2 | 0 221964174 | 0 062591474 | 1 10E-91 | -0 25981566 | 0 | 4 61E-91 |
| TOP1 | 0 560173992 | 0 062580187 | 1 19E-91 | -0 03867973 | 4 40E-36 | 4 97E-91 |
| CAPN7 | 0 470398431 | 0 062562629 | 1 33E-91 | -0 08068712 | 2 97E-151 | 5 58E-91 |
| LINC01376 | 0 256343873 | 0 062556078 | 1 39E-91 | -0 23229386 | 0 | 5 82E-91 |
| ZC4H2 | 0 213902267 | 0 062545738 | 1 49E-91 | 0 36849506 | 0 | 6 23E-91 |
| BRINP3 | 0 070434601 | 0 062542239 | 1 53E-91 | 0 456406167 | 0 | 6 37E-91 |
| SNHG5 | 0 251127905 | 0 062472713 | 2 41E-91 | 0 33998543 | 0 | 1 01E-90 |
| NNT | 0 392044697 | 0 062406617 | 3 74E-91 | 0 22699977 | 0 | 1 56E-90 |
| NDFIP2 | 0 203308523 | 0 062404629 | 3 79E-91 | -0 27086294 | 0 | 1 58E-90 |
| CXXC4-AS1 | 0 023728846 | 0 062385833 | 4 28E-91 | 0 494215591 | 0 | 1 78E-90 |
| NIPAL2 | 0 2092098 | 0 062369566 | 4 77E-91 | -0 26262035 | 0 | 1 98E-90 |
| FGGY | 0 340627439 | 0 062342948 | 5 69E-91 | 0 275755552 | 0 | 2 36E-90 |
| CBX3 | 0 349983819 | 0 062326367 | 6 34E-91 | 0 263912302 | 0 | 2 64E-90 |
| ZNF789 | 0 232377073 | 0 062314021 | 6 88E-91 | 0 317992989 | 0 | 2 86E-90 |
| SRGAP2-AS1 | 0 082874874 | 0 062303063 | 7 39E-91 | 0 437122174 | 0 | 3 07E-90 |
| LRBA | 0 654213702 | 0 062281542 | 8 52E-91 | 0 118104575 | 0 | 3 54E-90 |
| TNFSF13B | 0 111676915 | 0 062271571 | 9 10E-91 | -0 36042617 | 0 | 3 78E-90 |
| UPF2 | 0 44407112 | 0 062244524 | 1 09E-90 | 0 221423247 | 0 | 4 51E-90 |
| LINC01572 | 0 253221907 | 0 062236288 | 1 15E-90 | 0 269210564 | 0 | 4 75E-90 |
| VPS8 | 0 514153547 | 0 062205758 | 1 40E-90 | -0 06210573 | 2 71E-90 | 5 81E-90 |
| LYPLA1 | 0 2902667 | 0 062204289 | 1 42E-90 | 0 282206064 | 0 | 5 86E-90 |
| TEX9 | 0 224429385 | 0 062197355 | 1 48E-90 | 0 336468402 | 0 | 6 13E-90 |
| BTBD9 | 0 575555386 | 0 062166039 | 1 82E-90 | -0 00741858 | 0 016189811 | 7 53E-90 |
| LTBP4 | 0 41037673 | 0 062140353 | 2 16E-90 | -0 12099762 | 0 | 8 91E-90 |
| LYPLAL1 | 0 476823209 | 0 062085507 | 3 09E-90 | -0 06899019 | 5 11E-111 | 1 28E-89 |
| PRPF3 | 0 342921323 | 0 062075156 | 3 31E-90 | 0 243722723 | 0 | 1 36E-89 |
| PLSCR1 | 0 349412728 | 0 062074036 | 3 33E-90 | 0 253944594 | 0 | 1 37E-89 |
| ATRX | 0 760341513 | 0 061969164 | 6 63E-90 | 0 08834682 | 4 82E-181 | 2 73E-89 |
| SP4 | 0 154651539 | 0 061961686 | 6 96E-90 | 0 411228572 | 0 | 2 87E-89 |
| MAPKBP1 | 0 224467457 | 0 061938172 | 8 12E-90 | -0 25500177 | 0 | 3 34E-89 |
| ITIH4 | 0 039881213 | 0 061932468 | 8 43E-90 | -0 4443789 | 0 | 3 47E-89 |
| TSPAN2 | 0 020330852 | 0 061881916 | 1 17E-89 | 0 491781383 | 0 | 4 82E-89 |
| CLASRP | 0 307637395 | 0 061881209 | 1 18E-89 | 0 275743492 | 0 | 4 84E-89 |
| NOL10 | 0 403085797 | 0 061880929 | 1 18E-89 | 0 215865871 | 0 | 4 85E-89 |
| CCDC80 | 0 256600864 | 0 061874002 | 1 24E-89 | -0 23248281 | 0 | 5 07E-89 |
| SCAPER | 0 561192439 | 0 061812183 | 1 85E-89 | 0 160045146 | 0 | 7 60E-89 |
| FKBP5 | 0 539148312 | 0 061801687 | 1 98E-89 | -0 070103 | 1 42E-114 | 8 13E-89 |
| DPP4 | 0 057689745 | 0 061778658 | 2 31E-89 | 0 465495958 | 0 | 9 45E-89 |
| TLL2 | 0 032304734 | 0 061748064 | 2 81E-89 | 0 484879778 | 0 | 1 15E-88 |
| LETM2 | 0 050551103 | 0 061696824 | 3 93E-89 | 0 468974271 | 0 | 1 61E-88 |
| RASA4 | 0 028430831 | 0 061690552 | 4 10E-89 | 0 483532085 | 0 | 1 68E-88 |

| | | | | | | |
|---|---|---|---|---|---|---|
| CDK6 | 0 580495327 | 0 061689818 | 4 12E-89 | 0 073737955 | 1 38E-126 | 1 68E-88 |
| NIPBL | 0 767251718 | 0 061672472 | 4 61E-89 | 0 088883009 | 3 13E-183 | 1 88E-88 |
| HNF4G | 0 177533266 | 0 061653925 | 5 20E-89 | 0 366368186 | 0 | 2 13E-88 |
| HLX-AS1 | 0 043098361 | 0 061642375 | 5 61E-89 | -0 44116528 | 0 | 2 29E-88 |
| CFH | 0 63861339 | 0 061628897 | 6 12E-89 | 0 014660064 | 2 01E-06 | 2 50E-88 |
| ARID1A | 0 475681026 | 0 061621672 | 6 42E-89 | 0 164525769 | 0 | 2 62E-88 |
| LMBR1 | 0 471502541 | 0 061531599 | 1 15E-88 | 0 181188383 | 0 | 4 70E-88 |
| IGFBP2 | 0 227265805 | 0 061526672 | 1 19E-88 | 0 360116445 | 0 | 4 85E-88 |
| SRGAP1 | 0 520578325 | 0 061471507 | 1 71E-88 | 0 159235385 | 0 | 6 94E-88 |
| ARHGAP26 | 0 415840171 | 0 061471327 | 1 71E-88 | -0 07713024 | 2 39E-138 | 6 94E-88 |
| EPB41 | 0 262597323 | 0 061465051 | 1 78E-88 | 0 296527945 | 0 | 7 23E-88 |
| ZNF467 | 0 025642002 | 0 061435951 | 2 15E-88 | 0 489900367 | 0 | 8 73E-88 |
| WWP1 | 0 481011212 | 0 061426026 | 2 29E-88 | -0 08535971 | 4 22E-169 | 9 31E-88 |
| LRPPRC | 0 532894862 | 0 061402224 | 2 67E-88 | 0 147808355 | 0 | 1 09E-87 |
| PEBP4 | 0 062677276 | 0 061393485 | 2 83E-88 | -0 4181158 | 0 | 1 15E-87 |
| TBC1D30 | 0 075631532 | 0 061370621 | 3 28E-88 | -0 39535392 | 0 | 1 33E-87 |
| PTGS2 | 0 030363024 | 0 061359116 | 3 54E-88 | -0 45437827 | 0 | 1 43E-87 |
| SPTLC3 | 0 232948164 | 0 061328826 | 4 31E-88 | -0 24672378 | 0 | 1 74E-87 |
| ST8SIA1 | 0 165816375 | 0 061307107 | 4 96E-88 | -0 31273777 | 0 | 2 01E-87 |
| C1QTNF7 | 0 139984009 | 0 061301391 | 5 14E-88 | -0 33160791 | 0 | 2 08E-87 |
| INTS7 | 0 204231787 | 0 061268748 | 6 36E-88 | 0 318987446 | 0 | 2 57E-87 |
| MAPK8 | 0 571424492 | 0 061256921 | 6 86E-88 | 0 154707282 | 0 | 2 77E-87 |
| PPEF1 | 0 058041918 | 0 061254329 | 6 98E-88 | 0 466963155 | 0 | 2 82E-87 |
| LMCD1 | 0 153376102 | 0 06123654 | 7 83E-88 | 0 376515022 | 0 | 3 16E-87 |
| ZNF43 | 0 25678171 | 0 06117965 | 1 13E-87 | 0 266285929 | 0 | 4 57E-87 |
| LIN9 | 0 115817327 | 0 061178871 | 1 14E-87 | 0 367178244 | 0 | 4 59E-87 |
| HECTD1 | 0 609897013 | 0 061164712 | 1 25E-87 | 0 153980516 | 0 | 5 03E-87 |
| GLRX | 0 227008814 | 0 061161859 | 1 27E-87 | -0 26782136 | 0 | 5 12E-87 |
| LINC00342 | 0 47455788 | 0 061151999 | 1 35E-87 | 0 168656628 | 0 | 5 45E-87 |
| RBL2 | 0 3371533 | 0 061131122 | 1 55E-87 | -0 16359898 | 0 | 6 24E-87 |
| PDZD4 | 0 048247701 | 0 061102001 | 1 87E-87 | 0 478733854 | 0 | 7 53E-87 |
| ADAR | 0 40937732 | 0 061096795 | 1 93E-87 | -0 12358718 | 0 | 7 78E-87 |
| AP3B1 | 0 655955531 | 0 061079919 | 2 16E-87 | 0 10284505 | 6 21E-245 | 8 67E-87 |
| ORC5 | 0 340303821 | 0 061058736 | 2 47E-87 | 0 244790817 | 0 | 9 94E-87 |
| DNAJC27-AS1 | 0 109306885 | 0 061049437 | 2 62E-87 | -0 36803918 | 0 | 1 06E-86 |
| HACL1 | 0 208124726 | 0 061042335 | 2 75E-87 | -0 27724836 | 0 | 1 10E-86 |
| MTUS2 | 0 041565932 | 0 061037572 | 2 83E-87 | 0 479031401 | 0 | 1 14E-86 |
| P4HA3 | 0 215710723 | 0 061010992 | 3 36E-87 | 0 310801856 | 0 | 1 35E-86 |
| R3HDM1 | 0 523985837 | 0 060997789 | 3 66E-87 | 0 126168695 | 0 | 1 47E-86 |
| TNRC6B | 0 755116027 | 0 060954381 | 4 84E-87 | 0 028657125 | 1 54E-20 | 1 94E-86 |
| VGLL4 | 0 568711808 | 0 060947854 | 5 05E-87 | -0 02689562 | 2 80E-18 | 2 03E-86 |
| NARS2 | 0 299880071 | 0 060929839 | 5 67E-87 | 0 250683957 | 0 | 2 27E-86 |
| MYO5B | 0 062924749 | 0 060924512 | 5 87E-87 | 0 463485939 | 0 | 2 35E-86 |
| CEP112 | 0 7446365 | 0 060864947 | 8 62E-87 | 0 070160823 | 9 26E-115 | 3 45E-86 |

| | | | | | | |
|---|---|---|---|---|---|---|
| SUFU | 0 224990958 | 0 06086281 | 8 74E-87 | 0 324141577 | 0 | 3 50E-86 |
| POLH | 0 217690507 | 0 060845938 | 9 74E-87 | 0 331083741 | 0 | 3 90E-86 |
| NAA50 | 0 344539415 | 0 060844193 | 9 85E-87 | 0 243549198 | 0 | 3 94E-86 |
| PCNT | 0 191239459 | 0 06083171 | 1 07E-86 | 0 318557681 | 0 | 4 27E-86 |
| ERV3-1 | 0 105214064 | 0 060804036 | 1 27E-86 | 0 433038925 | 0 | 5 09E-86 |
| NUTM2B-AS1 | 0 514134511 | 0 060804028 | 1 27E-86 | -0 05232606 | 1 32E-64 | 5 09E-86 |
| MIR4713HG | 0 067645771 | 0 06069601 | 2 55E-86 | -0 41947201 | 0 | 1 02E-85 |
| RNASEH2B-AS | 0 036302374 | 0 060679368 | 2 84E-86 | -0 44871164 | 0 | 1 13E-85 |
| SPG11 | 0 506596105 | 0 060658525 | 3 24E-86 | 0 180429231 | 0 | 1 29E-85 |
| SLC39A10 | 0 337629209 | 0 060655054 | 3 32E-86 | 0 260604571 | 0 | 1 32E-85 |
| DPP10 | 0 081380518 | 0 060647434 | 3 48E-86 | 0 453423458 | 0 | 1 39E-85 |
| PTGIR | 0 021349299 | 0 060643073 | 3 58E-86 | -0 46700079 | 0 | 1 43E-85 |
| C21orf62-AS1 | 0 142753803 | 0 060620408 | 4 14E-86 | -0 33630127 | 0 | 1 65E-85 |
| RIPOR2 | 0 130960766 | 0 060598986 | 4 75E-86 | -0 33025193 | 0 | 1 89E-85 |
| RNF44 | 0 099465078 | 0 060576954 | 5 47E-86 | -0 3793581 | 0 | 2 17E-85 |
| WAC | 0 660895471 | 0 060563442 | 5 97E-86 | 0 017013519 | 3 49E-08 | 2 37E-85 |
| TBC1D19 | 0 428975272 | 0 060560668 | 6 07E-86 | -0 1004743 | 8 21E-234 | 2 41E-85 |
| RHBDL3 | 0 042574861 | 0 060547609 | 6 60E-86 | 0 475323524 | 0 | 2 62E-85 |
| THOC1 | 0 339294893 | 0 060537113 | 7 06E-86 | 0 261268661 | 0 | 2 80E-85 |
| MIR924HG | 0 170556433 | 0 060534838 | 7 17E-86 | 0 305366705 | 0 | 2 84E-85 |
| CNBD1 | 0 064000305 | 0 060520074 | 7 88E-86 | -0 41870441 | 0 | 3 12E-85 |
| CPEB4 | 0 272010813 | 0 060516194 | 8 07E-86 | -0 21919767 | 0 | 3 20E-85 |
| LINC02511 | 0 099788696 | 0 060514796 | 8 15E-86 | 0 433191827 | 0 | 3 23E-85 |
| CWF19L2 | 0 341626849 | 0 060503574 | 8 75E-86 | 0 291705964 | 0 | 3 47E-85 |
| THOC2 | 0 642001866 | 0 060496635 | 9 15E-86 | 0 148495837 | 0 | 3 62E-85 |
| YTHDC1 | 0 483685824 | 0 060495966 | 9 19E-86 | -0 07920429 | 8 41E-146 | 3 63E-85 |
| DLGAP1-AS4 | 0 044668862 | 0 060480549 | 1 01E-85 | 0 474811676 | 0 | 4 01E-85 |
| SLC2A4RG | 0 180074623 | 0 060475446 | 1 05E-85 | 0 359505513 | 0 | 4 14E-85 |
| CUX2 | 0 0238621 | 0 060470517 | 1 08E-85 | 0 487569589 | 0 | 4 27E-85 |
| GNA12 | 0 338181264 | 0 060452338 | 1 21E-85 | 0 27135149 | 0 | 4 79E-85 |
| ASCC1 | 0 276817498 | 0 060374794 | 1 99E-85 | -0 20801294 | 0 | 7 86E-85 |
| FAM43A | 0 086387086 | 0 060333569 | 2 59E-85 | -0 39429109 | 0 | 1 02E-84 |
| P3H2 | 0 245702538 | 0 06032578 | 2 72E-85 | -0 22526519 | 0 | 1 07E-84 |
| TENT2 | 0 659581961 | 0 060261031 | 4 12E-85 | -0 00853478 | 0 005667581 | 1 62E-84 |
| AZIN2 | 0 137709162 | 0 060255201 | 4 27E-85 | -0 34821938 | 0 | 1 68E-84 |
| CAMKMT | 0 496249833 | 0 060195519 | 6 25E-85 | -0 07853527 | 2 24E-143 | 2 46E-84 |
| ANKIB1 | 0 6514915 | 0 060190443 | 6 45E-85 | 0 102095272 | 2 18E-241 | 2 54E-84 |
| ATF1 | 0 241866707 | 0 060168196 | 7 43E-85 | -0 24932181 | 0 | 2 92E-84 |
| LNX1 | 0 063667168 | 0 060102253 | 1 13E-84 | -0 42052005 | 0 | 4 44E-84 |
| ASAP1 | 0 778978127 | 0 059989975 | 2 31E-84 | 0 044707583 | 1 24E-47 | 9 05E-84 |
| CRABP2 | 0 089870743 | 0 059979174 | 2 47E-84 | -0 3977183 | 0 | 9 69E-84 |
| SNED1 | 0 506215378 | 0 059946865 | 3 03E-84 | -0 03465262 | 2 74E-29 | 1 19E-83 |
| DGKB | 0 14182102 | 0 059925305 | 3 47E-84 | -0 33939385 | 0 | 1 36E-83 |
| MROH1 | 0 231092117 | 0 059899217 | 4 10E-84 | 0 358087391 | 0 | 1 61E-83 |

| | | | | | | |
|---|---|---|---|---|---|---|
| CMTM4 | 0 206030725 | 0 059888795 | 4 38E-84 | 0 360581627 | 0 | 1 72E-83 |
| ATG4C | 0 236241457 | 0 059848956 | 5 63E-84 | -0 24895946 | 0 | 2 21E-83 |
| FBN1 | 0 677695075 | 0 059835942 | 6 12E-84 | -0 01696347 | 3 83E-08 | 2 39E-83 |
| DNAJC1 | 0 510117835 | 0 059824303 | 6 58E-84 | 0 190698402 | 0 | 2 58E-83 |
| CWC27 | 0 41563077 | 0 05980576 | 7 40E-84 | 0 214510193 | 0 | 2 90E-83 |
| LAMC3 | 0 027298167 | 0 059798806 | 7 74E-84 | 0 488627323 | 0 | 3 02E-83 |
| SETX | 0 571405456 | 0 059767395 | 9 43E-84 | 0 16478459 | 0 | 3 69E-83 |
| SLC6A3 | 0 053958615 | 0 05976057 | 9 85E-84 | -0 44148753 | 0 | 3 85E-83 |
| ARMH3 | 0 427281034 | 0 059752502 | 1 04E-83 | -0 11206901 | 1 00E-290 | 4 05E-83 |
| ZNF516 | 0 433429784 | 0 059751829 | 1 04E-83 | -0 05305905 | 2 23E-66 | 4 06E-83 |
| MIB1 | 0 525089947 | 0 059747488 | 1 07E-83 | 0 17717241 | 0 | 4 17E-83 |
| MED14 | 0 39287278 | 0 059746072 | 1 08E-83 | 0 240774894 | 0 | 4 21E-83 |
| MAPK14 | 0 391673488 | 0 05974494 | 1 09E-83 | 0 216420272 | 0 | 4 24E-83 |
| CARMIL1 | 0 487902381 | 0 059724955 | 1 23E-83 | -0 05195991 | 9 96E-64 | 4 80E-83 |
| GABRB2 | 0 044754526 | 0 059708304 | 1 37E-83 | 0 476918639 | 0 | 5 33E-83 |
| DGKE | 0 128352782 | 0 059704301 | 1 41E-83 | -0 34644289 | 0 | 5 47E-83 |
| MICU1 | 0 553482705 | 0 059700069 | 1 44E-83 | 0 149600617 | 0 | 5 61E-83 |
| FAM126B | 0 167653386 | 0 059683058 | 1 61E-83 | 0 327316337 | 0 | 6 25E-83 |
| EGFL6 | 0 025937066 | 0 059672267 | 1 72E-83 | 0 486082814 | 0 | 6 68E-83 |
| IQGAP2 | 0 181331024 | 0 059664992 | 1 80E-83 | 0 395556306 | 0 | 6 99E-83 |
| ERBIN | 0 72592374 | 0 059658278 | 1 88E-83 | 0 103999873 | 1 90E-250 | 7 29E-83 |
| FREM1 | 0 347451981 | 0 059646121 | 2 03E-83 | -0 15526159 | 0 | 7 87E-83 |
| CAMK2D | 0 688993166 | 0 059591726 | 2 86E-83 | 0 078474692 | 3 71E-143 | 1 11E-82 |
| SLMAP | 0 573347166 | 0 059567461 | 3 33E-83 | -0 03677698 | 8 82E-33 | 1 29E-82 |
| OSBPL8 | 0 640707392 | 0 059553891 | 3 63E-83 | 0 090046027 | 5 06E-188 | 1 41E-82 |
| PUDP | 0 226513868 | 0 059532254 | 4 16E-83 | -0 25193213 | 0 | 1 61E-82 |
| TRIB2 | 0 282604557 | 0 059523326 | 4 40E-83 | -0 21655519 | 0 | 1 70E-82 |
| CRTC3 | 0 428832499 | 0 059518237 | 4 54E-83 | -0 10813764 | 1 02E-270 | 1 76E-82 |
| EMILIN1 | 0 244522282 | 0 059516654 | 4 58E-83 | -0 23902654 | 0 | 1 77E-82 |
| FGD5 | 0 220650663 | 0 059497806 | 5 16E-83 | -0 26300297 | 0 | 2 00E-82 |
| ANKRD13C | 0 416868135 | 0 059488307 | 5 48E-83 | -0 11699284 | 0 | 2 12E-82 |
| SEC24B | 0 521682435 | 0 059486803 | 5 53E-83 | -0 06210432 | 2 73E-90 | 2 14E-82 |
| REEP1 | 0 023490891 | 0 059481892 | 5 70E-83 | 0 489804702 | 0 | 2 20E-82 |
| FAR2 | 0 278930536 | 0 059476359 | 5 91E-83 | 0 284227153 | 0 | 2 28E-82 |
| INPPL1 | 0 180798005 | 0 059447897 | 7 06E-83 | 0 349108687 | 0 | 2 73E-82 |
| KIAA0586 | 0 311463707 | 0 059443734 | 7 25E-83 | 0 256382386 | 0 | 2 80E-82 |
| KDM3B | 0 391816261 | 0 05943797 | 7 52E-83 | 0 23683801 | 0 | 2 90E-82 |
| TSNARE1 | 0 120033885 | 0 059407262 | 9 12E-83 | -0 35100502 | 0 | 3 52E-82 |
| PTPRA | 0 668747977 | 0 059331595 | 1 47E-82 | 0 016218687 | 1 46E-07 | 5 65E-82 |
| ABAT | 0 125659135 | 0 05931513 | 1 63E-82 | -0 33815664 | 0 | 6 26E-82 |
| GLT8D2 | 0 326911728 | 0 059310028 | 1 68E-82 | -0 16594635 | 0 | 6 46E-82 |
| OGDH | 0 347994518 | 0 059292045 | 1 88E-82 | -0 15974231 | 0 | 7 23E-82 |
| RBKS | 0 121632941 | 0 059253067 | 2 40E-82 | -0 3588417 | 0 | 9 22E-82 |
| CNOT1 | 0 455692829 | 0 059223164 | 2 89E-82 | 0 191573067 | 0 | 1 11E-81 |

| | | | | | | |
|---|---|---|---|---|---|---|
| ZBTB1 | 0 350183701 | 0 059216901 | 3 01E-82 | -0 15325141 | 0 | 1 16E-81 |
| CXXC4 | 0 02848794 | 0 059195905 | 3 43E-82 | 0 492240158 | 0 | 1 32E-81 |
| OGN | 0 340056348 | 0 059190611 | 3 55E-82 | 0 236413302 | 0 | 1 36E-81 |
| ARHGAP18 | 0 309436333 | 0 059180289 | 3 78E-82 | -0 19826674 | 0 | 1 45E-81 |
| ACSS1 | 0 141706802 | 0 059160231 | 4 29E-82 | -0 33224243 | 0 | 1 64E-81 |
| LPIN2 | 0 304591574 | 0 059134124 | 5 05E-82 | 0 302899721 | 0 | 1 93E-81 |
| WIPF3 | 0 068302526 | 0 059126523 | 5 29E-82 | -0 39893963 | 0 | 2 03E-81 |
| SYT17 | 0 082484628 | 0 059125871 | 5 32E-82 | -0 38747884 | 0 | 2 04E-81 |
| ABI2 | 0 542708115 | 0 059124242 | 5 37E-82 | 0 156166528 | 0 | 2 06E-81 |
| UBXN2B | 0 185509509 | 0 059086218 | 6 81E-82 | -0 2871062 | 0 | 2 60E-81 |
| AKAP7 | 0 228084369 | 0 059084667 | 6 88E-82 | 0 366540718 | 0 | 2 63E-81 |
| EIF2B3 | 0 375406903 | 0 059061366 | 7 95E-82 | -0 15885994 | 0 | 3 04E-81 |
| CSNK1G3 | 0 438721898 | 0 059059618 | 8 04E-82 | -0 09816175 | 3 22E-223 | 3 07E-81 |
| NBDY | 0 33573509 | 0 059040549 | 9 06E-82 | -0 16846498 | 0 | 3 46E-81 |
| LINC01133 | 0 048799756 | 0 05903931 | 9 13E-82 | 0 464356627 | 0 | 3 48E-81 |
| ATP2B1 | 0 511583636 | 0 059035864 | 9 33E-82 | 0 135798376 | 0 | 3 56E-81 |
| LINC00276 | 0 040785441 | 0 058995238 | 1 20E-81 | -0 4438747 | 0 | 4 58E-81 |
| CYYR1 | 0 033637281 | 0 058949436 | 1 60E-81 | 0 484743085 | 0 | 6 09E-81 |
| CYP20A1 | 0 342102758 | 0 058944627 | 1 65E-81 | -0 16002975 | 0 | 6 27E-81 |
| USPL1 | 0 254973254 | 0 05891178 | 2 02E-81 | -0 23506809 | 0 | 7 69E-81 |
| UBTD2 | 0 326312082 | 0 05889991 | 2 18E-81 | 0 263466882 | 0 | 8 28E-81 |
| NGLY1 | 0 304858084 | 0 058891598 | 2 29E-81 | 0 286231078 | 0 | 8 71E-81 |
| LINC01357 | 0 022910282 | 0 058861669 | 2 76E-81 | 0 491036483 | 0 | 1 05E-80 |
| RBMS3 | 0 897536693 | 0 05883985 | 3 16E-81 | 0 090421252 | 1 40E-189 | 1 20E-80 |
| PPIP5K2 | 0 481582304 | 0 058836734 | 3 22E-81 | 0 215790633 | 0 | 1 22E-80 |
| LINC01138 | 0 357227161 | 0 058835509 | 3 25E-81 | -0 17428707 | 0 | 1 23E-80 |
| ZNF726 | 0 039919286 | 0 058750189 | 5 52E-81 | 0 466051941 | 0 | 2 09E-80 |
| FRG1-DT | 0 097542404 | 0 05874573 | 5 68E-81 | -0 3810031 | 0 | 2 15E-80 |
| FBXW11 | 0 552949687 | 0 058742341 | 5 80E-81 | 0 14918223 | 0 | 2 19E-80 |
| PDS5A | 0 631741258 | 0 058730494 | 6 24E-81 | 0 10409663 | 6 53E-251 | 2 36E-80 |
| OPRM1 | 0 030125069 | 0 058726912 | 6 38E-81 | 0 48281913 | 0 | 2 41E-80 |
| LINC00871 | 0 024556928 | 0 058700079 | 7 54E-81 | 0 489774933 | 0 | 2 85E-80 |
| ATXN3 | 0 388047058 | 0 058690424 | 8 00E-81 | -0 11211326 | 5 92E-291 | 3 02E-80 |
| NIBAN1 | 0 441948564 | 0 058689251 | 8 06E-81 | 0 156133999 | 0 | 3 04E-80 |
| USP42 | 0 297433896 | 0 058688575 | 8 10E-81 | 0 309189695 | 0 | 3 05E-80 |
| SLC25A42 | 0 096838058 | 0 058688042 | 8 12E-81 | -0 37661131 | 0 | 3 06E-80 |
| TMEM116 | 0 173059717 | 0 058685754 | 8 24E-81 | 0 353070516 | 0 | 3 10E-80 |
| PPP2R5A | 0 222411528 | 0 058674689 | 8 82E-81 | -0 24893655 | 0 | 3 32E-80 |
| SOCS2-AS1 | 0 048875902 | 0 058581874 | 1 57E-80 | 0 480241415 | 0 | 5 90E-80 |
| MROH8 | 0 07401344 | 0 058572398 | 1 66E-80 | 0 459639026 | 0 | 6 25E-80 |
| ULK3 | 0 108202776 | 0 058564803 | 1 74E-80 | 0 431042634 | 0 | 6 55E-80 |
| ANGPTL1 | 0 164988293 | 0 058547744 | 1 94E-80 | 0 382695006 | 0 | 7 28E-80 |
| ABCD3 | 0 373607965 | 0 058531824 | 2 14E-80 | -0 14724053 | 0 | 8 03E-80 |
| TMEM243 | 0 166377948 | 0 058528093 | 2 19E-80 | 0 374861704 | 0 | 8 21E-80 |

| | | | | | | |
|---|---|---|---|---|---|---|
| EPB41L1 | 0 165645048 | 0 058505657 | 2 51E-80 | -0 29203637 | 0 | 9 43E-80 |
| YPEL2 | 0 222440083 | 0 058488633 | 2 79E-80 | -0 23252064 | 0 | 1 05E-79 |
| TMEM119 | 0 049999048 | 0 058463604 | 3 26E-80 | -0 4337177 | 0 | 1 22E-79 |
| PARN | 0 423473758 | 0 058386721 | 5 24E-80 | 0 196349609 | 0 | 1 96E-79 |
| PILRA | 0 081028345 | 0 058374697 | 5 65E-80 | 0 438356039 | 0 | 2 12E-79 |
| CDR2 | 0 354181341 | 0 058371684 | 5 75E-80 | -0 16104205 | 0 | 2 15E-79 |
| SPRY2 | 0 268232092 | 0 058311385 | 8 34E-80 | -0 22537908 | 0 | 3 12E-79 |
| RHOU | 0 127619882 | 0 058282985 | 9 94E-80 | -0 33962791 | 0 | 3 72E-79 |
| WHRN | 0 15922027 | 0 058269529 | 1 08E-79 | -0 32294566 | 0 | 4 04E-79 |
| TAZ | 0 071900402 | 0 058259054 | 1 15E-79 | 0 452001679 | 0 | 4 30E-79 |
| DAB1 | 0 277321962 | 0 058207667 | 1 58E-79 | -0 16928399 | 0 | 5 90E-79 |
| PRKAR2B | 0 129514001 | 0 058206687 | 1 59E-79 | -0 34089052 | 0 | 5 93E-79 |
| CKMT2-AS1 | 0 32479869 | 0 058191603 | 1 74E-79 | 0 269981522 | 0 | 6 51E-79 |
| INPP5B | 0 259380176 | 0 058189313 | 1 77E-79 | -0 23161686 | 0 | 6 60E-79 |
| TRIM2 | 0 367868497 | 0 058177657 | 1 90E-79 | -0 11721298 | 0 | 7 08E-79 |
| RALYL | 0 061906303 | 0 058170287 | 1 99E-79 | -0 42040699 | 0 | 7 41E-79 |
| LACTB2 | 0 123926824 | 0 058146806 | 2 30E-79 | 0 404841584 | 0 | 8 56E-79 |
| SLC45A4 | 0 076069369 | 0 058127919 | 2 58E-79 | -0 40062467 | 0 | 9 60E-79 |
| DOCK9 | 0 511612191 | 0 058118492 | 2 73E-79 | -0 07101487 | 1 57E-117 | 1 02E-78 |
| PLB1 | 0 073404276 | 0 058109442 | 2 89E-79 | -0 40111134 | 0 | 1 07E-78 |
| DDX27 | 0 186861092 | 0 058085174 | 3 35E-79 | 0 371351827 | 0 | 1 25E-78 |
| TTC14 | 0 476985018 | 0 058056564 | 4 00E-79 | -0 06204024 | 4 16E-90 | 1 49E-78 |
| CSRP2 | 0 095343702 | 0 058054378 | 4 05E-79 | 0 406862915 | 0 | 1 50E-78 |
| CACNA1I | 0 013611011 | 0 058039871 | 4 43E-79 | 0 495204717 | 0 | 1 64E-78 |
| CEP68 | 0 201176448 | 0 058028717 | 4 74E-79 | -0 28252611 | 0 | 1 76E-78 |
| TMEM132D | 0 032856789 | 0 058027523 | 4 78E-79 | 0 483873257 | 0 | 1 77E-78 |
| RNF180 | 0 204983724 | 0 058008391 | 5 37E-79 | -0 25981369 | 0 | 1 99E-78 |
| FAM149A | 0 087767223 | 0 057994199 | 5 86E-79 | 0 435001207 | 0 | 2 17E-78 |
| CEP192 | 0 457234776 | 0 057991135 | 5 97E-79 | 0 177269024 | 0 | 2 21E-78 |
| LANCL3 | 0 026603339 | 0 057987636 | 6 10E-79 | 0 483975674 | 0 | 2 26E-78 |
| CERCAM | 0 16111439 | 0 057982628 | 6 29E-79 | -0 31713877 | 0 | 2 33E-78 |
| XG | 0 065418515 | 0 057954042 | 7 50E-79 | -0 41691489 | 0 | 2 77E-78 |
| ATP1A2 | 0 070986655 | 0 057944389 | 7 95E-79 | -0 40065016 | 0 | 2 94E-78 |
| DGKD | 0 25267937 | 0 057941907 | 8 08E-79 | 0 292471531 | 0 | 2 99E-78 |
| ARHGEF12 | 0 584036093 | 0 057920518 | 9 21E-79 | -0 00942123 | 0 002259925 | 3 40E-78 |
| RFX7 | 0 50552055 | 0 057919202 | 9 28E-79 | 0 140152369 | 0 | 3 43E-78 |
| GLCE | 0 317155584 | 0 057868318 | 1 27E-78 | -0 19061329 | 0 | 4 68E-78 |
| NDUFA4L2 | 0 033865717 | 0 05786755 | 1 27E-78 | 0 487720531 | 0 | 4 70E-78 |
| OAF | 0 390398051 | 0 05785464 | 1 38E-78 | -0 10374491 | 3 18E-249 | 5 08E-78 |
| DPP8 | 0 347480535 | 0 057845315 | 1 46E-78 | 0 258595871 | 0 | 5 38E-78 |
| SLC37A1 | 0 078467952 | 0 057832544 | 1 58E-78 | -0 39840065 | 0 | 5 81E-78 |
| POLM | 0 108431212 | 0 057818309 | 1 72E-78 | -0 36740747 | 0 | 6 34E-78 |
| SLC25A48 | 0 036235747 | 0 057813179 | 1 78E-78 | -0 45104793 | 0 | 6 54E-78 |
| STARD8 | 0 078801089 | 0 057806821 | 1 85E-78 | 0 428548547 | 0 | 6 79E-78 |

| | | | | | | |
|---|---|---|---|---|---|---|
| TRMT13 | 0 255039881 | 0 057711451 | 3 30E-78 | -0 23124315 | 0 | 1 21E-77 |
| ZFYVE9 | 0 425824751 | 0 057695523 | 3 64E-78 | -0 12165572 | 0 | 1 34E-77 |
| C1orf21 | 0 651501018 | 0 057683614 | 3 92E-78 | -0 00732629 | 0 017563504 | 1 44E-77 |
| IGF1R | 0 691877177 | 0 057666925 | 4 33E-78 | 0 090225714 | 9 08E-189 | 1 59E-77 |
| TNK2 | 0 145285641 | 0 05765314 | 4 71E-78 | 0 39915746 | 0 | 1 73E-77 |
| HMGN5 | 0 113180788 | 0 057639282 | 5 13E-78 | 0 429568834 | 0 | 1 88E-77 |
| TPR | 0 534969827 | 0 057602209 | 6 43E-78 | 0 166355002 | 0 | 2 36E-77 |
| POT1 | 0 336924863 | 0 057588369 | 7 00E-78 | 0 252339164 | 0 | 2 56E-77 |
| STARD9 | 0 432649293 | 0 057543696 | 9 18E-78 | 0 195361152 | 0 | 3 36E-77 |
| BDNF-AS | 0 200386438 | 0 057524547 | 1 03E-77 | -0 28186355 | 0 | 3 78E-77 |
| NPTX2 | 0 089366279 | 0 057518312 | 1 07E-77 | 0 424822963 | 0 | 3 92E-77 |
| UBE2D3 | 0 596809503 | 0 057505836 | 1 16E-77 | -0 00989502 | 0 001339683 | 4 23E-77 |
| ATP8B1 | 0 317774267 | 0 057494764 | 1 24E-77 | -0 17419925 | 0 | 4 52E-77 |
| LCOR | 0 526812739 | 0 057477109 | 1 38E-77 | 0 177404465 | 0 | 5 03E-77 |
| RAP2C-AS1 | 0 20731568 | 0 057443168 | 1 69E-77 | 0 373453483 | 0 | 6 18E-77 |
| ARHGAP25 | 0 071310274 | 0 057385623 | 2 40E-77 | -0 41224494 | 0 | 8 76E-77 |
| ETV6 | 0 590194361 | 0 057365936 | 2 70E-77 | -0 04551656 | 2 62E-49 | 9 87E-77 |
| MIS18BP1 | 0 308836687 | 0 057350347 | 2 97E-77 | 0 266950156 | 0 | 1 08E-76 |
| L2HGDH | 0 151158364 | 0 057340983 | 3 14E-77 | 0 389249061 | 0 | 1 15E-76 |
| BIRC6 | 0 817031848 | 0 057339583 | 3 17E-77 | 0 057081716 | 1 51E-76 | 1 16E-76 |
| EPHB6 | 0 031495688 | 0 057334684 | 3 27E-77 | 0 486325163 | 0 | 1 19E-76 |
| LINC01208 | 0 034018008 | 0 057334315 | 3 27E-77 | 0 48280417 | 0 | 1 19E-76 |
| PARVG | 0 043678971 | 0 057305024 | 3 91E-77 | -0 44091174 | 0 | 1 42E-76 |
| UPP2 | 0 136757343 | 0 057154695 | 9 71E-77 | -0 32936707 | 0 | 3 53E-76 |
| CLIP2 | 0 272515277 | 0 057135026 | 1 09E-76 | 0 314506295 | 0 | 3 97E-76 |
| STARD5 | 0 038034684 | 0 057134997 | 1 09E-76 | -0 45114246 | 0 | 3 97E-76 |
| CDYL2 | 0 116036245 | 0 057103967 | 1 32E-76 | 0 407866242 | 0 | 4 79E-76 |
| LRP4 | 0 139232072 | 0 057099139 | 1 36E-76 | -0 31219685 | 0 | 4 93E-76 |
| TMEM65 | 0 362043365 | 0 057085213 | 1 48E-76 | 0 207165593 | 0 | 5 36E-76 |
| NUP160 | 0 387304639 | 0 057064345 | 1 67E-76 | 0 224058692 | 0 | 6 07E-76 |
| ZNF710 | 0 182539834 | 0 057055292 | 1 77E-76 | 0 345039478 | 0 | 6 41E-76 |
| FAM174B | 0 147750852 | 0 057052951 | 1 79E-76 | -0 3305288 | 0 | 6 50E-76 |
| SEPTIN4 | 0 033066189 | 0 05704078 | 1 93E-76 | -0 45322791 | 0 | 6 99E-76 |
| ANGPT2 | 0 180483905 | 0 057037346 | 1 97E-76 | 0 364765553 | 0 | 7 14E-76 |
| BCR | 0 232662618 | 0 057033099 | 2 02E-76 | -0 20046193 | 0 | 7 32E-76 |
| ILDR2 | 0 022167863 | 0 057018842 | 2 20E-76 | 0 489669587 | 0 | 7 97E-76 |
| ANKRD18A | 0 041251832 | 0 057013655 | 2 27E-76 | 0 478932624 | 0 | 8 22E-76 |
| SYT16 | 0 053558851 | 0 057010318 | 2 32E-76 | -0 42736502 | 0 | 8 38E-76 |
| MAP1LC3B2 | 0 017846605 | 0 056998622 | 2 49E-76 | -0 47234158 | 0 | 8 98E-76 |
| EZH2 | 0 27670328 | 0 056987995 | 2 65E-76 | 0 225249595 | 0 | 9 57E-76 |
| PIKFYVE | 0 426919343 | 0 056969898 | 2 96E-76 | 0 211780914 | 0 | 1 07E-75 |
| CPLANE1 | 0 608145666 | 0 056963525 | 3 07E-76 | 0 132560927 | 0 | 1 11E-75 |
| ZNF236 | 0 281871657 | 0 056898624 | 4 54E-76 | 0 317337062 | 0 | 1 64E-75 |
| PIGL | 0 376444385 | 0 056862202 | 5 66E-76 | -0 14479292 | 0 | 2 04E-75 |

| | | | | | | |
|---|---|---|---|---|---|---|
| SULF2 | 0 194942034 | 0 056847862 | 6 17E-76 | -0 28856608 | 0 | 2 22E-75 |
| HLX | 0 038367821 | 0 056847197 | 6 19E-76 | 0 477967839 | 0 | 2 23E-75 |
| SLC16A14 | 0 041223278 | 0 056798614 | 8 29E-76 | 0 47915159 | 0 | 2 98E-75 |
| DSTYK | 0 233871428 | 0 05679493 | 8 47E-76 | -0 23646947 | 0 | 3 04E-75 |
| TAL1 | 0 015695494 | 0 056785804 | 8 95E-76 | -0 47500245 | 0 | 3 21E-75 |
| PPP1CB | 0 510964954 | 0 056782761 | 9 12E-76 | -0 06182667 | 1 68E-89 | 3 27E-75 |
| PCNX4 | 0 486979117 | 0 056775853 | 9 50E-76 | 0 196995125 | 0 | 3 41E-75 |
| TBC1D15 | 0 448639851 | 0 056768812 | 9 91E-76 | 0 195956511 | 0 | 3 56E-75 |
| TTTY14 | 0 237640631 | 0 056765684 | 1 01E-75 | -0 24492724 | 0 | 3 62E-75 |
| STOX2 | 0 310302488 | 0 056759834 | 1 05E-75 | -0 14054551 | 0 | 3 75E-75 |
| RAPGEF1 | 0 317307875 | 0 05672583 | 1 28E-75 | 0 271415003 | 0 | 4 60E-75 |
| ATF7IP2 | 0 315994365 | 0 05671374 | 1 38E-75 | 0 238371834 | 0 | 4 94E-75 |
| APC | 0 455569093 | 0 056663072 | 1 87E-75 | 0 205458872 | 0 | 6 69E-75 |
| SP3 | 0 488245036 | 0 056630773 | 2 27E-75 | 0 140848878 | 0 | 8 11E-75 |
| WDR7 | 0 481982068 | 0 056557161 | 3 52E-75 | -0 07112812 | 6 72E-118 | 1 26E-74 |
| IDH2 | 0 102206316 | 0 056545249 | 3 78E-75 | 0 425938596 | 0 | 1 35E-74 |
| CLK4 | 0 280681883 | 0 056537098 | 3 97E-75 | -0 20063212 | 0 | 1 42E-74 |
| GLDN | 0 035702728 | 0 056527264 | 4 21E-75 | -0 44828407 | 0 | 1 50E-74 |
| SNTB2 | 0 339942129 | 0 056419092 | 8 03E-75 | 0 237220467 | 0 | 2 87E-74 |
| N4BP1 | 0 311435153 | 0 056344664 | 1 25E-74 | -0 17319641 | 0 | 4 47E-74 |
| SYNE3 | 0 176924102 | 0 056339045 | 1 29E-74 | -0 29379307 | 0 | 4 61E-74 |
| DRAXIN | 0 01046049 | 0 056299633 | 1 64E-74 | 0 496355219 | 0 | 5 83E-74 |
| SYNRG | 0 400839504 | 0 056290293 | 1 73E-74 | -0 12149125 | 0 | 6 16E-74 |
| C5orf63 | 0 061963412 | 0 056288361 | 1 75E-74 | -0 41467957 | 0 | 6 23E-74 |
| COX19 | 0 168709905 | 0 05618634 | 3 21E-74 | 0 392565414 | 0 | 1 14E-73 |
| ARHGAP17 | 0 349536464 | 0 056166187 | 3 62E-74 | -0 15226472 | 0 | 1 29E-73 |
| ARHGAP8 | 0 050065676 | 0 056163697 | 3 67E-74 | -0 42464706 | 0 | 1 31E-73 |
| OR3A2 | 0 046962746 | 0 056150678 | 3 97E-74 | 0 475318777 | 0 | 1 41E-73 |
| MAFB | 0 072842702 | 0 056147445 | 4 04E-74 | -0 40673034 | 0 | 1 44E-73 |
| MYO9B | 0 395690164 | 0 05614062 | 4 21E-74 | 0 21263059 | 0 | 1 50E-73 |
| ITM2C | 0 191030058 | 0 056133408 | 4 39E-74 | 0 380967113 | 0 | 1 56E-73 |
| ETNK1 | 0 578363252 | 0 056120924 | 4 73E-74 | -0 04256048 | 2 50E-43 | 1 68E-73 |
| ECM2 | 0 352106375 | 0 056115513 | 4 89E-74 | 0 236020737 | 0 | 1 73E-73 |
| ZNF608 | 0 321514915 | 0 056114538 | 4 91E-74 | -0 1770721 | 0 | 1 74E-73 |
| GSTCD | 0 222915992 | 0 056078994 | 6 07E-74 | 0 308386382 | 0 | 2 15E-73 |
| YEATS2 | 0 459442996 | 0 056057418 | 6 89E-74 | 0 18107275 | 0 | 2 44E-73 |
| IFNGR2 | 0 372132645 | 0 056056161 | 6 95E-74 | -0 15114293 | 0 | 2 46E-73 |
| RCN3 | 0 151453427 | 0 056039704 | 7 66E-74 | -0 31870895 | 0 | 2 71E-73 |
| MPPED1 | 0 037577811 | 0 056035986 | 7 83E-74 | -0 4462724 | 0 | 2 77E-73 |
| USP8 | 0 47145495 | 0 056032599 | 7 99E-74 | -0 08593114 | 2 36E-171 | 2 83E-73 |
| KLHDC1 | 0 260960195 | 0 056016654 | 8 78E-74 | -0 21274481 | 0 | 3 10E-73 |
| PPP1R3C | 0 033780054 | 0 056003721 | 9 48E-74 | -0 45129838 | 0 | 3 35E-73 |
| GRB10 | 0 367887533 | 0 055997664 | 9 82E-74 | -0 14683763 | 0 | 3 47E-73 |
| DDX19B | 0 170261369 | 0 055978339 | 1 10E-73 | 0 380018639 | 0 | 3 89E-73 |

| | | | | | | |
|---|---|---|---|---|---|---|
| MTHFR | 0 134177914 | 0 055977816 | 1 10E-73 | -0 32918351 | 0 | 3 90E-73 |
| CYP27C1 | 0 104947555 | 0 055952637 | 1 28E-73 | -0 37004732 | 0 | 4 53E-73 |
| PRDM1 | 0 090118216 | 0 055950318 | 1 30E-73 | 0 430738112 | 0 | 4 59E-73 |
| EED | 0 165530829 | 0 055941557 | 1 37E-73 | 0 392448479 | 0 | 4 83E-73 |
| CHSY1 | 0 404104243 | 0 055914008 | 1 61E-73 | 0 204351392 | 0 | 5 68E-73 |
| TNFRSF21 | 0 104528754 | 0 055904873 | 1 70E-73 | -0 37233913 | 0 | 5 99E-73 |
| PRKAR2A | 0 378357541 | 0 055889635 | 1 86E-73 | -0 11773335 | 0 | 6 55E-73 |
| NRF1 | 0 490177229 | 0 055880745 | 1 96E-73 | 0 180854192 | 0 | 6 90E-73 |
| TBX2 | 0 152976338 | 0 055878381 | 1 99E-73 | -0 30868992 | 0 | 6 99E-73 |
| CRY1 | 0 320801051 | 0 055873634 | 2 05E-73 | 0 27166354 | 0 | 7 19E-73 |
| AVEN | 0 217823761 | 0 055839589 | 2 50E-73 | 0 336133722 | 0 | 8 78E-73 |
| ABTB2 | 0 293369629 | 0 055814127 | 2 91E-73 | -0 17518333 | 0 | 1 02E-72 |
| TNFAIP8L3 | 0 039043612 | 0 055809598 | 2 98E-73 | 0 476503853 | 0 | 1 05E-72 |
| RNF165 | 0 093382955 | 0 055776978 | 3 62E-73 | -0 37752505 | 0 | 1 27E-72 |
| ADAMTSL4-AS: | 0 111714987 | 0 055767604 | 3 82E-73 | 0 377140458 | 0 | 1 34E-72 |
| PRKCH | 0 140688355 | 0 055763713 | 3 91E-73 | -0 33763216 | 0 | 1 37E-72 |
| NRG4 | 0 046077554 | 0 05575371 | 4 15E-73 | 0 468568285 | 0 | 1 45E-72 |
| DSP | 0 048875902 | 0 055747129 | 4 31E-73 | 0 475087149 | 0 | 1 51E-72 |
| KRT18 | 0 027421903 | 0 055746285 | 4 34E-73 | -0 45697888 | 0 | 1 52E-72 |
| MAGED1 | 0 467495384 | 0 055743627 | 4 40E-73 | -0 07869592 | 5 89E-144 | 1 54E-72 |
| C2orf88 | 0 184072262 | 0 055739026 | 4 52E-73 | -0 3056745 | 0 | 1 58E-72 |
| SCN3A | 0 035274409 | 0 055710688 | 5 35E-73 | 0 479295768 | 0 | 1 87E-72 |
| HMGA1P4 | 0 076202623 | 0 055701818 | 5 63E-73 | -0 40297235 | 0 | 1 97E-72 |
| PLD2 | 0 074051512 | 0 055657327 | 7 32E-73 | 0 457009052 | 0 | 2 56E-72 |
| NFYA | 0 133292722 | 0 055635554 | 8 32E-73 | -0 34513695 | 0 | 2 91E-72 |
| ITPRID1 | 0 019759761 | 0 055598217 | 1 04E-72 | -0 46913374 | 0 | 3 62E-72 |
| PRDM2 | 0 461270488 | 0 05559662 | 1 05E-72 | -0 0963966 | 2 67E-215 | 3 65E-72 |
| ZNF394 | 0 189878358 | 0 055583218 | 1 13E-72 | 0 358487691 | 0 | 3 95E-72 |
| UBR3 | 0 735984466 | 0 055539217 | 1 47E-72 | 0 008641207 | 0 005095767 | 5 11E-72 |
| SYCP1 | 0 020692543 | 0 055502052 | 1 82E-72 | 0 486826429 | 0 | 6 35E-72 |
| PTGER4 | 0 092478727 | 0 055486898 | 1 99E-72 | -0 39483364 | 0 | 6 93E-72 |
| MGST1 | 0 129295083 | 0 055471855 | 2 18E-72 | 0 380118386 | 0 | 7 57E-72 |
| CEP120 | 0 219755954 | 0 05546483 | 2 27E-72 | 0 326828193 | 0 | 7 89E-72 |
| MADD | 0 179408349 | 0 055456363 | 2 38E-72 | -0 30681018 | 0 | 8 28E-72 |
| AARSD1 | 0 172460071 | 0 055445143 | 2 54E-72 | -0 30502096 | 0 | 8 84E-72 |
| TPRG1 | 0 17603891 | 0 055430114 | 2 78E-72 | -0 30287274 | 0 | 9 65E-72 |
| STS | 0 106004074 | 0 055421455 | 2 92E-72 | 0 444216008 | 0 | 1 02E-71 |
| DNM2 | 0 331927814 | 0 055408983 | 3 14E-72 | 0 264220819 | 0 | 1 09E-71 |
| WDR37 | 0 298404751 | 0 055405545 | 3 21E-72 | -0 18084827 | 0 | 1 11E-71 |
| RALGPS2 | 0 373712665 | 0 055386507 | 3 59E-72 | 0 24815621 | 0 | 1 24E-71 |
| SGCE | 0 567036607 | 0 055358893 | 4 22E-72 | 0 130223459 | 0 | 1 46E-71 |
| CD24 | 0 029325541 | 0 055345757 | 4 55E-72 | 0 488019613 | 0 | 1 58E-71 |
| IL7 | 0 313947955 | 0 055336811 | 4 80E-72 | 0 210678141 | 0 | 1 66E-71 |
| NTRK1 | 0 025489711 | 0 055320538 | 5 28E-72 | -0 46246819 | 0 | 1 83E-71 |

| | | | | | | |
|---|---|---|---|---|---|---|
| SNX14 | 0 464858845 | 0 055318041 | 5 35E-72 | -0 06506657 | 6 13E-99 | 1 85E-71 |
| VPS53 | 0 442938455 | 0 055311935 | 5 55E-72 | -0 09333919 | 6 22E-202 | 1 92E-71 |
| SNRNP70 | 0 56334355 | 0 055311723 | 5 56E-72 | -0 04975443 | 1 41E-58 | 1 92E-71 |
| SNX9 | 0 532200034 | 0 055292844 | 6 20E-72 | -0 06654863 | 2 03E-103 | 2 14E-71 |
| SYT9 | 0 030810379 | 0 055285217 | 6 49E-72 | 0 491832868 | 0 | 2 24E-71 |
| TEAD1 | 0 646722887 | 0 055256161 | 7 69E-72 | 0 083589441 | 3 21E-162 | 2 66E-71 |
| PHACTR2 | 0 471683387 | 0 055248531 | 8 04E-72 | -0 06466967 | 9 33E-98 | 2 78E-71 |
| SLITRK5 | 0 032990044 | 0 055239496 | 8 47E-72 | 0 480140481 | 0 | 2 92E-71 |
| FUS | 0 680674269 | 0 055236421 | 8 63E-72 | 0 09826855 | 1 06E-223 | 2 98E-71 |
| CCAR1 | 0 476413927 | 0 055235338 | 8 68E-72 | 0 192443609 | 0 | 2 99E-71 |
| ULK4 | 0 380603834 | 0 055214558 | 9 80E-72 | -0 14136225 | 0 | 3 38E-71 |
| TMEM178A | 0 036054901 | 0 055193578 | 1 11E-71 | -0 44586742 | 0 | 3 82E-71 |
| ABL1 | 0 565789724 | 0 055175629 | 1 23E-71 | -0 03376394 | 6 87E-28 | 4 24E-71 |
| ABCG1 | 0 086291904 | 0 055156336 | 1 38E-71 | -0 38587084 | 0 | 4 74E-71 |
| PBRM1 | 0 560507129 | 0 05513187 | 1 59E-71 | 0 174402502 | 0 | 5 46E-71 |
| ZNF160 | 0 240429461 | 0 055129014 | 1 61E-71 | -0 24396938 | 0 | 5 55E-71 |
| CUL2 | 0 225409758 | 0 055117 | 1 73E-71 | -0 25920781 | 0 | 5 95E-71 |
| GABPB1-AS1 | 0 467476347 | 0 055104459 | 1 86E-71 | -0 1026227 | 7 03E-244 | 6 40E-71 |
| AMT | 0 117720965 | 0 055100013 | 1 91E-71 | -0 35838422 | 0 | 6 57E-71 |
| IREB2 | 0 427385734 | 0 055089828 | 2 03E-71 | 0 199779209 | 0 | 6 97E-71 |
| NCAPD3 | 0 129552074 | 0 055038632 | 2 73E-71 | 0 379697825 | 0 | 9 38E-71 |
| FOXK2 | 0 485589461 | 0 054909824 | 5 78E-71 | 0 161244756 | 0 | 1 98E-70 |
| MPZL1 | 0 364232548 | 0 054885209 | 6 66E-71 | -0 15750404 | 0 | 2 28E-70 |
| SLC7A6OS | 0 195513126 | 0 054869709 | 7 29E-71 | -0 29269163 | 0 | 2 50E-70 |
| SINHCAF | 0 159562925 | 0 054853451 | 8 01E-71 | 0 357020239 | 0 | 2 74E-70 |
| LINC01250 | 0 021073271 | 0 054846211 | 8 35E-71 | 0 490822766 | 0 | 2 86E-70 |
| GATA3 | 0 037377929 | 0 054837693 | 8 78E-71 | 0 46818264 | 0 | 3 00E-70 |
| VOPP1 | 0 391616379 | 0 054835368 | 8 90E-71 | 0 220884675 | 0 | 3 04E-70 |
| TET3 | 0 244969637 | 0 054832229 | 9 06E-71 | 0 298080427 | 0 | 3 10E-70 |
| LINC02253 | 0 058793855 | 0 054823954 | 9 50E-71 | 0 457469399 | 0 | 3 25E-70 |
| LINC01791 | 0 033475472 | 0 054807605 | 1 04E-70 | -0 45516425 | 0 | 3 57E-70 |
| STAT2 | 0 47545259 | 0 054794223 | 1 13E-70 | -0 10935803 | 7 53E-277 | 3 85E-70 |
| CREB1 | 0 428803944 | 0 054715954 | 1 78E-70 | 0 187072527 | 0 | 6 06E-70 |
| HNRNPM | 0 510679408 | 0 054695509 | 2 00E-70 | 0 17215889 | 0 | 6 81E-70 |
| IL33 | 0 052521368 | 0 054688918 | 2 08E-70 | -0 4372446 | 0 | 7 08E-70 |
| TP53 | 0 117321201 | 0 054678135 | 2 21E-70 | 0 430077854 | 0 | 7 53E-70 |
| SHANK3 | 0 052311968 | 0 054672474 | 2 28E-70 | -0 42617844 | 0 | 7 78E-70 |
| TPTE2 | 0 039062649 | 0 054668686 | 2 33E-70 | 0 479943742 | 0 | 7 95E-70 |
| GJA5 | 0 011840627 | 0 054627672 | 2 96E-70 | 0 49575682 | 0 | 1 01E-69 |
| LRRK1 | 0 270821039 | 0 054580013 | 3 90E-70 | 0 268427503 | 0 | 1 32E-69 |
| SBK1 | 0 016647313 | 0 054576727 | 3 97E-70 | 0 491671465 | 0 | 1 35E-69 |
| EPHX1 | 0 177162057 | 0 05455103 | 4 60E-70 | -0 2878281 | 0 | 1 56E-69 |
| OLFML3 | 0 163037064 | 0 054486042 | 6 70E-70 | -0 31818421 | 0 | 2 27E-69 |
| SMC5 | 0 636757343 | 0 054457375 | 7 90E-70 | 0 128001718 | 0 | 2 68E-69 |

| | | | | | | |
|---|---|---|---|---|---|---|
| DCP2 | 0 340713103 | 0 054435656 | 8 95E-70 | 0 231409986 | 0 | 3 04E-69 |
| MICU2 | 0 42788068 | 0 054433541 | 9 06E-70 | -0 10329211 | 4 63E-247 | 3 07E-69 |
| MTHFS | 0 138518208 | 0 054410143 | 1 04E-69 | 0 399822551 | 0 | 3 51E-69 |
| C1orf112 | 0 153185738 | 0 054404647 | 1 07E-69 | 0 351190732 | 0 | 3 62E-69 |
| CCDC6 | 0 281976357 | 0 054394994 | 1 13E-69 | -0 2153448 | 0 | 3 83E-69 |
| LINC00536 | 0 032999562 | 0 054352557 | 1 44E-69 | -0 455729 | 0 | 4 88E-69 |
| JPH1 | 0 098446632 | 0 054345245 | 1 50E-69 | 0 434921435 | 0 | 5 09E-69 |
| PASK | 0 052454741 | 0 054285726 | 2 12E-69 | 0 464285299 | 0 | 7 16E-69 |
| DAB2 | 0 291408882 | 0 054273097 | 2 28E-69 | -0 19636522 | 0 | 7 70E-69 |
| SMC6 | 0 386381375 | 0 054271782 | 2 29E-69 | 0 215139435 | 0 | 7 75E-69 |
| TMEM45A | 0 189545221 | 0 05426726 | 2 35E-69 | -0 27640188 | 0 | 7 96E-69 |
| SMOC2 | 0 243446727 | 0 054263854 | 2 40E-69 | -0 25103875 | 0 | 8 11E-69 |
| PER2 | 0 185338181 | 0 054255052 | 2 53E-69 | -0 2815086 | 0 | 8 53E-69 |
| ANKS1A | 0 473910643 | 0 054214332 | 3 19E-69 | -0 0554536 | 2 42E-72 | 1 08E-68 |
| RECQL4 | 0 044354762 | 0 054189233 | 3 68E-69 | 0 446815255 | 0 | 1 24E-68 |
| ZNF655 | 0 436742114 | 0 054174925 | 4 00E-69 | -0 11756609 | 0 | 1 35E-68 |
| CERK | 0 156878795 | 0 054135583 | 5 01E-69 | 0 362192399 | 0 | 1 69E-68 |
| PIPOX | 0 031210143 | 0 054104602 | 5 98E-69 | -0 45435315 | 0 | 2 01E-68 |
| THY1 | 0 092773791 | 0 054059909 | 7 72E-69 | 0 435008537 | 0 | 2 60E-68 |
| ACAD11 | 0 347242581 | 0 05404831 | 8 25E-69 | -0 14484957 | 0 | 2 77E-68 |
| TTF1 | 0 184957454 | 0 05404682 | 8 32E-69 | 0 359221287 | 0 | 2 80E-68 |
| TRAK1 | 0 424654014 | 0 054046689 | 8 32E-69 | -0 09932261 | 1 67E-228 | 2 80E-68 |
| SCUBE2 | 0 096152748 | 0 053993079 | 1 13E-68 | -0 37886883 | 0 | 3 80E-68 |
| IMPG2 | 0 055176943 | 0 053973013 | 1 27E-68 | 0 46582385 | 0 | 4 26E-68 |
| WDHD1 | 0 154109002 | 0 053953841 | 1 41E-68 | 0 312719114 | 0 | 4 74E-68 |
| GKAP1 | 0 137766271 | 0 053935009 | 1 57E-68 | 0 386721782 | 0 | 5 28E-68 |
| SNX8 | 0 23154899 | 0 053933442 | 1 59E-68 | 0 345828322 | 0 | 5 33E-68 |
| CDC25B | 0 166358912 | 0 053924313 | 1 67E-68 | 0 360906659 | 0 | 5 61E-68 |
| CDK14 | 0 812910472 | 0 053919574 | 1 72E-68 | 0 061667383 | 4 76E-89 | 5 76E-68 |
| TTC39C | 0 347689935 | 0 053905933 | 1 86E-68 | 0 228378359 | 0 | 6 22E-68 |
| PRMT9 | 0 213845158 | 0 053902002 | 1 90E-68 | -0 2755327 | 0 | 6 36E-68 |
| DOCK4 | 0 539833622 | 0 053895163 | 1 98E-68 | -0 04705544 | 1 40E-52 | 6 61E-68 |
| CIP2A | 0 129352192 | 0 05387126 | 2 26E-68 | 0 347580093 | 0 | 7 57E-68 |
| ZNF433-AS1 | 0 190097276 | 0 053867596 | 2 31E-68 | 0 339077388 | 0 | 7 73E-68 |
| ZDHHC21 | 0 453046772 | 0 053839518 | 2 71E-68 | -0 08448655 | 1 09E-165 | 9 06E-68 |
| CNN2 | 0 099465078 | 0 053825543 | 2 94E-68 | -0 37163662 | 0 | 9 81E-68 |
| AKAP10 | 0 429860463 | 0 053820533 | 3 02E-68 | -0 11006707 | 1 91E-280 | 1 01E-67 |
| M1AP | 0 015495612 | 0 053807225 | 3 26E-68 | 0 494051386 | 0 | 1 09E-67 |
| EYS | 0 118101692 | 0 053777695 | 3 86E-68 | -0 36423581 | 0 | 1 29E-67 |
| DOCK8 | 0 124202852 | 0 053756179 | 4 36E-68 | -0 35592286 | 0 | 1 45E-67 |
| LINC00472 | 0 318326322 | 0 053729091 | 5 08E-68 | -0 17227587 | 0 | 1 69E-67 |
| MINK1 | 0 20700158 | 0 053726361 | 5 16E-68 | -0 27737755 | 0 | 1 72E-67 |
| USF3 | 0 179484495 | 0 053719363 | 5 37E-68 | -0 2947788 | 0 | 1 79E-67 |
| LYRM2 | 0 179217986 | 0 053699499 | 6 01E-68 | -0 29364797 | 0 | 2 00E-67 |

| | | | | | | |
|---|---|---|---|---|---|---|
| SGF29 | 0 070815328 | 0 0536762 | 6 86E-68 | -0 4116018 | 0 | 2 28E-67 |
| CDK12 | 0 411366622 | 0 053565097 | 1 29E-67 | -0 11812341 | 0 | 4 28E-67 |
| STAG3 | 0 089632788 | 0 053504452 | 1 82E-67 | 0 41876577 | 0 | 6 03E-67 |
| ZNF528-AS1 | 0 127086863 | 0 053486936 | 2 00E-67 | 0 398915294 | 0 | 6 65E-67 |
| NT5C3A | 0 36680246 | 0 053459454 | 2 34E-67 | 0 237304893 | 0 | 7 77E-67 |
| CHFR | 0 284879405 | 0 053454777 | 2 40E-67 | -0 19985343 | 0 | 7 97E-67 |
| ZNF10 | 0 204365042 | 0 0534534 | 2 42E-67 | -0 27088549 | 0 | 8 03E-67 |
| CDC7 | 0 088899888 | 0 053430648 | 2 76E-67 | 0 412597196 | 0 | 9 13E-67 |
| LIN7A | 0 235698921 | 0 053411292 | 3 07E-67 | 0 280740302 | 0 | 1 02E-66 |
| CTTNBP2 | 0 235118311 | 0 053388147 | 3 50E-67 | 0 273592881 | 0 | 1 16E-66 |
| C18orf25 | 0 282918658 | 0 053351787 | 4 30E-67 | -0 21942617 | 0 | 1 42E-66 |
| TAPT1 | 0 278997164 | 0 053307603 | 5 52E-67 | 0 304434443 | 0 | 1 82E-66 |
| STAT5A | 0 07863928 | 0 053285491 | 6 25E-67 | -0 39247117 | 0 | 2 06E-66 |
| SLTM | 0 533665835 | 0 053281362 | 6 39E-67 | 0 156352061 | 0 | 2 11E-66 |
| LINC02254 | 0 046010927 | 0 053271804 | 6 75E-67 | 0 468695773 | 0 | 2 23E-66 |
| APBB2 | 0 76555748 | 0 053267531 | 6 91E-67 | -0 00078726 | 0 798588194 | 2 28E-66 |
| CNOT4 | 0 589642306 | 0 053262478 | 7 11E-67 | 0 131415152 | 0 | 2 35E-66 |
| GINS1 | 0 09130799 | 0 053173029 | 1 18E-66 | 0 36896845 | 0 | 3 88E-66 |
| CDS1 | 0 035036455 | 0 053157519 | 1 28E-66 | 0 4849429 | 0 | 4 23E-66 |
| ZNF571 | 0 085121167 | 0 053150468 | 1 34E-66 | -0 39304033 | 0 | 4 40E-66 |
| PTCHD1 | 0 017208886 | 0 053140215 | 1 41E-66 | -0 47229841 | 0 | 4 66E-66 |
| TMEM106A | 0 078334698 | 0 05312336 | 1 55E-66 | 0 44317393 | 0 | 5 12E-66 |
| GRAMD1A | 0 181416687 | 0 053076505 | 2 02E-66 | -0 29579107 | 0 | 6 65E-66 |
| HECA | 0 16021968 | 0 053034922 | 2 55E-66 | -0 31905218 | 0 | 8 40E-66 |
| SLC7A8 | 0 101920771 | 0 053021188 | 2 76E-66 | -0 36829922 | 0 | 9 06E-66 |
| EDEM3 | 0 356808361 | 0 053019176 | 2 79E-66 | -0 12908758 | 0 | 9 16E-66 |
| GSG1L | 0 022224972 | 0 053016542 | 2 83E-66 | -0 46514191 | 0 | 9 30E-66 |
| ZBTB20-AS2 | 0 094677429 | 0 053008538 | 2 96E-66 | -0 39573123 | 0 | 9 72E-66 |
| ILRUN | 0 318097885 | 0 052981434 | 3 45E-66 | 0 26189545 | 0 | 1 13E-65 |
| ZNF846 | 0 151443909 | 0 052979301 | 3 49E-66 | -0 31724381 | 0 | 1 14E-65 |
| NUP153 | 0 445127639 | 0 052977109 | 3 53E-66 | 0 190034009 | 0 | 1 16E-65 |
| B4GALT6 | 0 101911252 | 0 052912374 | 5 07E-66 | 0 460432045 | 0 | 1 66E-65 |
| MTMR9 | 0 220108127 | 0 052899381 | 5 46E-66 | 0 343272747 | 0 | 1 79E-65 |
| DISC1FP1 | 0 073099694 | 0 052857886 | 6 88E-66 | -0 40907629 | 0 | 2 25E-65 |
| KDM1B | 0 175601074 | 0 052849367 | 7 21E-66 | -0 30234308 | 0 | 2 36E-65 |
| SEC24D | 0 546229845 | 0 052845831 | 7 36E-66 | -0 04877882 | 2 26E-56 | 2 41E-65 |
| FSD2 | 0 059936038 | 0 052833724 | 7 87E-66 | -0 42878424 | 0 | 2 57E-65 |
| ATP2C1 | 0 490767356 | 0 052813502 | 8 81E-66 | 0 170765228 | 0 | 2 88E-65 |
| FAM13C | 0 079591099 | 0 052765139 | 1 15E-65 | -0 4042647 | 0 | 3 77E-65 |
| SLC9A8 | 0 232624546 | 0 052755932 | 1 22E-65 | 0 322207309 | 0 | 3 97E-65 |
| BCL2 | 0 50152291 | 0 052741375 | 1 32E-65 | -0 01207196 | 9 12E-05 | 4 30E-65 |
| HEATR5B | 0 427909235 | 0 052737961 | 1 34E-65 | -0 11624786 | 0 | 4 38E-65 |
| FBXO41 | 0 028012031 | 0 052737746 | 1 35E-65 | 0 489064187 | 0 | 4 39E-65 |
| SUPT5H | 0 262978051 | 0 052713174 | 1 54E-65 | 0 298341751 | 0 | 5 03E-65 |

| | | | | | | |
|---|---|---|---|---|---|---|
| DLGAP2 | 0 046477318 | 0 052713022 | 1 54E-65 | 0 47130397 | 0 | 5 03E-65 |
| PCDH19 | 0 111058232 | 0 05269767 | 1 68E-65 | -0 37309807 | 0 | 5 48E-65 |
| MEI1 | 0 032571244 | 0 052686605 | 1 79E-65 | 0 477889554 | 0 | 5 82E-65 |
| ANK3 | 0 395633055 | 0 052672318 | 1 94E-65 | 0 14915187 | 0 | 6 30E-65 |
| PEAK1 | 0 799708743 | 0 052634227 | 2 39E-65 | 0 036050549 | 1 45E-31 | 7 79E-65 |
| MYSM1 | 0 421474939 | 0 052626541 | 2 50E-65 | 0 206096805 | 0 | 8 12E-65 |
| ATXN2 | 0 690268603 | 0 052619378 | 2 60E-65 | 0 079161756 | 1 20E-145 | 8 45E-65 |
| PCF11 | 0 283737222 | 0 052616079 | 2 65E-65 | 0 321324347 | 0 | 8 60E-65 |
| MAGI3 | 0 424663532 | 0 052586702 | 3 12E-65 | 0 151049417 | 0 | 1 01E-64 |
| MTMR1 | 0 267108945 | 0 052585875 | 3 13E-65 | -0 22162935 | 0 | 1 02E-64 |
| CHRM2 | 0 014915003 | 0 052572731 | 3 37E-65 | 0 493533463 | 0 | 1 09E-64 |
| INTS9 | 0 251984542 | 0 052563009 | 3 56E-65 | -0 23783562 | 0 | 1 15E-64 |
| ERCC6 | 0 17372599 | 0 052544482 | 3 94E-65 | -0 29818493 | 0 | 1 28E-64 |
| POLA1 | 0 398735984 | 0 052530477 | 4 26E-65 | 0 191329146 | 0 | 1 38E-64 |
| FAM172A | 0 741400316 | 0 052519721 | 4 52E-65 | 0 029192843 | 2 96E-21 | 1 47E-64 |
| ELF4 | 0 060697493 | 0 052483185 | 5 54E-65 | 0 447803 | 0 | 1 80E-64 |
| ERCC5 | 0 233052864 | 0 052461058 | 6 26E-65 | -0 25057296 | 0 | 2 03E-64 |
| ZFP36L2 | 0 393091698 | 0 052440441 | 7 02E-65 | -0 1352514 | 0 | 2 27E-64 |
| LMBR1L | 0 207325198 | 0 052435954 | 7 20E-65 | -0 27028724 | 0 | 2 33E-64 |
| TAF3 | 0 329852849 | 0 05242451 | 7 67E-65 | 0 297342463 | 0 | 2 48E-64 |
| GK5 | 0 347204508 | 0 052415446 | 8 07E-65 | -0 16720334 | 0 | 2 61E-64 |
| SPOCK3 | 0 079695799 | 0 052411397 | 8 25E-65 | 0 44438821 | 0 | 2 67E-64 |
| SLC10A7 | 0 356894024 | 0 052375287 | 1 01E-64 | 0 241931984 | 0 | 3 26E-64 |
| B3GALNT2 | 0 272772268 | 0 052336998 | 1 25E-64 | -0 21867526 | 0 | 4 02E-64 |
| ACSF3 | 0 13788049 | 0 052334801 | 1 26E-64 | 0 412219816 | 0 | 4 07E-64 |
| PLEKHA2 | 0 418809846 | 0 052305241 | 1 48E-64 | -0 09302608 | 1 38E-200 | 4 78E-64 |
| GPR173 | 0 332765415 | 0 052299701 | 1 53E-64 | -0 16529067 | 0 | 4 93E-64 |
| PDE4B | 0 391026251 | 0 052264412 | 1 86E-64 | 0 183676261 | 0 | 5 99E-64 |
| USP12 | 0 346890408 | 0 052259662 | 1 91E-64 | 0 260545208 | 0 | 6 15E-64 |
| TPM2 | 0 224943367 | 0 052245315 | 2 07E-64 | -0 26159426 | 0 | 6 65E-64 |
| LINC02388 | 0 055262607 | 0 05222731 | 2 28E-64 | 0 461836198 | 0 | 7 35E-64 |
| SAFB2 | 0 296929432 | 0 052150377 | 3 49E-64 | -0 19918076 | 0 | 1 12E-63 |
| SLC25A26 | 0 319487541 | 0 052132866 | 3 84E-64 | -0 19399889 | 0 | 1 24E-63 |
| DENND6A | 0 45673983 | 0 052117511 | 4 18E-64 | -0 10196449 | 9 01E-241 | 1 34E-63 |
| SLC12A6 | 0 173373817 | 0 052095409 | 4 73E-64 | -0 31029303 | 0 | 1 52E-63 |
| RSPO2 | 0 021987017 | 0 05208685 | 4 95E-64 | 0 490916257 | 0 | 1 59E-63 |
| LINC02250 | 0 057147208 | 0 052076396 | 5 25E-64 | -0 42752001 | 0 | 1 68E-63 |
| EFNA2 | 0 012868592 | 0 052068564 | 5 48E-64 | 0 495655575 | 0 | 1 76E-63 |
| PPM1L | 0 146047096 | 0 052039967 | 6 41E-64 | -0 33567211 | 0 | 2 05E-63 |
| TTC27 | 0 346176543 | 0 052027146 | 6 88E-64 | -0 17097242 | 0 | 2 20E-63 |
| PRELID2 | 0 331413832 | 0 052021395 | 7 10E-64 | 0 262459377 | 0 | 2 27E-63 |
| STEAP2-AS1 | 0 072585711 | 0 051968858 | 9 48E-64 | 0 431899984 | 0 | 3 03E-63 |
| RTN1 | 0 116017209 | 0 051909543 | 1 31E-63 | -0 35278919 | 0 | 4 20E-63 |
| SACM1L | 0 313624336 | 0 051893464 | 1 43E-63 | -0 19428313 | 0 | 4 58E-63 |

| | | | | | | |
|---|---|---|---|---|---|---|
| AOX1 | 0 052578477 | 0 051892523 | 1 44E-63 | -0 43445104 | 0 | 4 60E-63 |
| DNAH1 | 0 078962898 | 0 051870904 | 1 62E-63 | -0 4000511 | 0 | 5 18E-63 |
| SIK2 | 0 409520093 | 0 051861212 | 1 71E-63 | -0 12193793 | 0 | 5 46E-63 |
| CCDC34 | 0 077144924 | 0 051857849 | 1 74E-63 | 0 423718568 | 0 | 5 56E-63 |
| ZFC3H1 | 0 579543508 | 0 051824415 | 2 09E-63 | 0 11452646 | 1 33E-303 | 6 67E-63 |
| TANK | 0 467076583 | 0 051818756 | 2 16E-63 | 0 166662956 | 0 | 6 88E-63 |
| YAF2 | 0 37810055 | 0 051817864 | 2 17E-63 | 0 215750608 | 0 | 6 91E-63 |
| MDM2 | 0 525108983 | 0 051800921 | 2 38E-63 | 0 145206617 | 0 | 7 58E-63 |
| RAD50 | 0 425853306 | 0 051800733 | 2 38E-63 | -0 11329845 | 3 93E-297 | 7 59E-63 |
| UBXN2A | 0 38472521 | 0 051791838 | 2 50E-63 | 0 225966477 | 0 | 7 96E-63 |
| AKNA | 0 100074242 | 0 051761294 | 2 96E-63 | -0 38151947 | 0 | 9 41E-63 |
| FGF13 | 0 256800746 | 0 051736321 | 3 39E-63 | -0 19786827 | 0 | 1 08E-62 |
| ALPK1 | 0 332137214 | 0 051730537 | 3 50E-63 | 0 265036035 | 0 | 1 11E-62 |
| ME1 | 0 162913327 | 0 051727135 | 3 57E-63 | -0 31355456 | 0 | 1 13E-62 |
| ASGR1 | 0 018360587 | 0 051726946 | 3 57E-63 | 0 492274618 | 0 | 1 13E-62 |
| VPS13D | 0 551046049 | 0 051725464 | 3 60E-63 | -0 0520199 | 7 16E-64 | 1 14E-62 |
| SMG7 | 0 400249377 | 0 051719559 | 3 72E-63 | 0 230464672 | 0 | 1 18E-62 |
| NAF1 | 0 235137347 | 0 051690098 | 4 37E-63 | 0 329466275 | 0 | 1 38E-62 |
| NCOR1 | 0 689402448 | 0 051686189 | 4 46E-63 | 0 020188099 | 5 98E-11 | 1 41E-62 |
| FHL1 | 0 246987493 | 0 051650254 | 5 43E-63 | -0 22509126 | 0 | 1 72E-62 |
| MPHOSPH8 | 0 452647008 | 0 051630525 | 6 05E-63 | -0 09434756 | 2 71E-206 | 1 91E-62 |
| ZC3H6 | 0 345053397 | 0 051614244 | 6 61E-63 | 0 256920022 | 0 | 2 09E-62 |
| TBC1D31 | 0 143857912 | 0 051594449 | 7 36E-63 | 0 344505637 | 0 | 2 33E-62 |
| APAF1 | 0 256772192 | 0 051566073 | 8 59E-63 | 0 274907695 | 0 | 2 72E-62 |
| SSTR2 | 0 030001333 | 0 051502384 | 1 22E-62 | 0 480248359 | 0 | 3 84E-62 |
| ARFGEF2 | 0 469313358 | 0 051500493 | 1 23E-62 | -0 08393261 | 1 53E-163 | 3 88E-62 |
| STIM2 | 0 471531096 | 0 051461577 | 1 52E-62 | -0 08261912 | 1 65E-158 | 4 79E-62 |
| ASPH | 0 629923283 | 0 051436135 | 1 74E-62 | 0 078195184 | 3 77E-142 | 5 50E-62 |
| ANKHD1 | 0 644152976 | 0 051435903 | 1 75E-62 | 0 115607555 | 0 | 5 51E-62 |
| EP300 | 0 451057471 | 0 051434715 | 1 76E-62 | -0 09976836 | 1 50E-230 | 5 54E-62 |
| MIATNB | 0 133787668 | 0 051420949 | 1 89E-62 | -0 3541006 | 0 | 5 97E-62 |
| RABGAP1 | 0 623974415 | 0 051417469 | 1 93E-62 | 0 131466506 | 0 | 6 08E-62 |
| CDKL3 | 0 260065485 | 0 051400916 | 2 11E-62 | -0 23964881 | 0 | 6 64E-62 |
| ZBTB25 | 0 176676629 | 0 051373822 | 2 45E-62 | 0 368453699 | 0 | 7 69E-62 |
| IRF2BP2 | 0 278454627 | 0 051369605 | 2 50E-62 | 0 309332618 | 0 | 7 87E-62 |
| POU2F2 | 0 036540329 | 0 051368742 | 2 51E-62 | 0 477028916 | 0 | 7 90E-62 |
| TEX14 | 0 069616036 | 0 051364219 | 2 58E-62 | -0 414268 | 0 | 8 10E-62 |
| USP3-AS1 | 0 25094706 | 0 051316083 | 3 35E-62 | 0 285186352 | 0 | 1 05E-61 |
| SUSD6 | 0 311930098 | 0 051270967 | 4 27E-62 | -0 16146741 | 0 | 1 34E-61 |
| ZCCHC14 | 0 28264263 | 0 05124356 | 4 96E-62 | -0 19283635 | 0 | 1 56E-61 |
| SIDT1 | 0 022767509 | 0 051241953 | 5 00E-62 | 0 490867044 | 0 | 1 57E-61 |
| SNX2 | 0 350069483 | 0 051238314 | 5 10E-62 | -0 16184171 | 0 | 1 60E-61 |
| BTN3A1 | 0 112723915 | 0 051232086 | 5 28E-62 | -0 36427638 | 0 | 1 65E-61 |
| ALOX12-AS1 | 0 41742019 | 0 051194944 | 6 45E-62 | 0 20113444 | 0 | 2 02E-61 |

| | | | | | | |
|---|---|---|---|---|---|---|
| GALNT8 | 0 067931317 | 0 051185894 | 6 77E-62 | -0 4153961 | 0 | 2 12E-61 |
| KIAA1755 | 0 348736936 | 0 051174693 | 7 20E-62 | -0 12953368 | 0 | 2 25E-61 |
| GNAO1 | 0 095714911 | 0 051172586 | 7 28E-62 | -0 37889469 | 0 | 2 28E-61 |
| PHLPP1 | 0 442805201 | 0 051171694 | 7 31E-62 | 0 127598447 | 0 | 2 29E-61 |
| MSR1 | 0 148664598 | 0 051134602 | 8 94E-62 | 0 352500625 | 0 | 2 80E-61 |
| RAP1B | 0 70710628 | 0 051120595 | 9 64E-62 | -0 00975568 | 0 001565963 | 3 02E-61 |
| TPK1 | 0 226532904 | 0 051120011 | 9 67E-62 | -0 25589608 | 0 | 3 02E-61 |
| DIS3L | 0 18266357 | 0 051081592 | 1 19E-61 | 0 357470805 | 0 | 3 72E-61 |
| SLC5A3 | 0 330918886 | 0 051079256 | 1 21E-61 | -0 19954929 | 0 | 3 77E-61 |
| SLC38A9 | 0 333326988 | 0 05102236 | 1 64E-61 | -0 18241648 | 0 | 5 12E-61 |
| CMIP | 0 381850717 | 0 051018534 | 1 67E-61 | 0 234956686 | 0 | 5 22E-61 |
| MAP4K5 | 0 611629324 | 0 051017864 | 1 68E-61 | 0 131664453 | 0 | 5 24E-61 |
| MYRIP | 0 123679351 | 0 051017116 | 1 69E-61 | 0 423771823 | 0 | 5 26E-61 |
| PPP2R2A | 0 523662218 | 0 050994444 | 1 91E-61 | -0 05666773 | 1 82E-75 | 5 94E-61 |
| VCAN-AS1 | 0 13215054 | 0 050980525 | 2 05E-61 | 0 379105042 | 0 | 6 40E-61 |
| HSD17B14 | 0 093744646 | 0 050971016 | 2 16E-61 | -0 37962252 | 0 | 6 73E-61 |
| GTSF1 | 0 015743085 | 0 050966199 | 2 22E-61 | 0 494046268 | 0 | 6 91E-61 |
| SCAI | 0 34309265 | 0 050958657 | 2 31E-61 | 0 233128594 | 0 | 7 19E-61 |
| CDH4 | 0 092107517 | 0 050899954 | 3 17E-61 | 0 428269039 | 0 | 9 86E-61 |
| RAB12 | 0 263377815 | 0 050879449 | 3 54E-61 | 0 315831014 | 0 | 1 10E-60 |
| SLC4A5 | 0 154032857 | 0 050857777 | 3 98E-61 | -0 32006373 | 0 | 1 24E-60 |
| DCP1B | 0 236517485 | 0 050771336 | 6 33E-61 | -0 26377804 | 0 | 1 97E-60 |
| SECISBP2 | 0 487969009 | 0 05077079 | 6 34E-61 | 0 173369607 | 0 | 1 97E-60 |
| KCNK2 | 0 196084217 | 0 05076091 | 6 69E-61 | -0 27711717 | 0 | 2 08E-60 |
| REL | 0 200434029 | 0 050739003 | 7 52E-61 | -0 28712079 | 0 | 2 34E-60 |
| ABHD17A | 0 16495022 | 0 050712355 | 8 68E-61 | -0 31615586 | 0 | 2 69E-60 |
| BDP1 | 0 552607032 | 0 050694186 | 9 57E-61 | 0 145237253 | 0 | 2 97E-60 |
| ACOXL | 0 017456359 | 0 050691146 | 9 72E-61 | 0 490932548 | 0 | 3 02E-60 |
| ERCC8 | 0 253459862 | 0 050671134 | 1 08E-60 | 0 308072935 | 0 | 3 36E-60 |
| RNF212B | 0 044716453 | 0 050666399 | 1 11E-60 | -0 44187489 | 0 | 3 44E-60 |
| MAP3K2 | 0 584435857 | 0 050636231 | 1 31E-60 | -0 03693928 | 4 68E-33 | 4 04E-60 |
| VAV3 | 0 107269993 | 0 050635891 | 1 31E-60 | -0 37550821 | 0 | 4 05E-60 |
| HRC | 0 006815024 | 0 050602695 | 1 56E-60 | -0 48514877 | 0 | 4 83E-60 |
| YPEL1 | 0 081589918 | 0 050595948 | 1 62E-60 | 0 434444537 | 0 | 5 01E-60 |
| EFEMP1 | 0 113485371 | 0 050595079 | 1 63E-60 | -0 35755634 | 0 | 5 03E-60 |
| CGNL1 | 0 221364528 | 0 0505903 | 1 67E-60 | -0 25444963 | 0 | 5 16E-60 |
| PRDM16 | 0 094258628 | 0 050590263 | 1 67E-60 | 0 430045028 | 0 | 5 16E-60 |
| NIFK-AS1 | 0 202080676 | 0 050582306 | 1 74E-60 | 0 325617674 | 0 | 5 38E-60 |
| HOXD10 | 0 059412537 | 0 050578811 | 1 77E-60 | 0 443444049 | 0 | 5 48E-60 |
| B4GALT5 | 0 378119587 | 0 050552508 | 2 04E-60 | 0 220665799 | 0 | 6 30E-60 |
| SCAF4 | 0 31570882 | 0 050539556 | 2 19E-60 | 0 257791558 | 0 | 6 75E-60 |
| ACAP2 | 0 610096895 | 0 050485162 | 2 93E-60 | -0 02356122 | 2 21E-14 | 9 02E-60 |
| ERI1 | 0 146456378 | 0 0504776 | 3 05E-60 | 0 370843481 | 0 | 9 39E-60 |
| TAB2 | 0 447078868 | 0 050472235 | 3 14E-60 | 0 190814529 | 0 | 9 66E-60 |

| | | | | | | |
|---|---|---|---|---|---|---|
| KIAA2026 | 0 468913594 | 0 05047091 | 3 16E-60 | -0 09145272 | 6 67E-194 | 9 72E-60 |
| SGK3 | 0 409786602 | 0 050450857 | 3 51E-60 | 0 199516774 | 0 | 1 08E-59 |
| MCU | 0 390331423 | 0 05044024 | 3 72E-60 | 0 220868752 | 0 | 1 14E-59 |
| MAP2K4 | 0 523538482 | 0 05038497 | 4 99E-60 | -0 05339374 | 3 39E-67 | 1 54E-59 |
| NSMAF | 0 350259847 | 0 05036212 | 5 64E-60 | -0 16568082 | 0 | 1 73E-59 |
| AMBRA1 | 0 498952999 | 0 050336216 | 6 47E-60 | -0 07036538 | 2 02E-115 | 1 99E-59 |
| SLC2A9 | 0 074022958 | 0 050327059 | 6 80E-60 | 0 458898846 | 0 | 2 09E-59 |
| MB21D2 | 0 137851935 | 0 050325076 | 6 87E-60 | 0 375759401 | 0 | 2 11E-59 |
| ASPN | 0 240077288 | 0 050315316 | 7 23E-60 | 0 261519586 | 0 | 2 22E-59 |
| MXRA8 | 0 344834479 | 0 050288042 | 8 36E-60 | -0 16492695 | 0 | 2 57E-59 |
| ANKDD1A | 0 218756544 | 0 050270502 | 9 18E-60 | -0 25939998 | 0 | 2 82E-59 |
| MARCHF3 | 0 15890617 | 0 050258266 | 9 80E-60 | -0 31156838 | 0 | 3 00E-59 |
| MXD1 | 0 165911557 | 0 050249585 | 1 03E-59 | -0 32376061 | 0 | 3 15E-59 |
| KCNQ5 | 0 186347109 | 0 050223253 | 1 18E-59 | 0 297421175 | 0 | 3 62E-59 |
| LINC02652 | 0 035997792 | 0 050202439 | 1 32E-59 | -0 45016646 | 0 | 4 04E-59 |
| PPM1K | 0 20642097 | 0 050197489 | 1 35E-59 | -0 26758414 | 0 | 4 14E-59 |
| SCN11A | 0 037035274 | 0 050189781 | 1 41E-59 | 0 481640834 | 0 | 4 32E-59 |
| PEG10 | 0 119729303 | 0 050171474 | 1 55E-59 | 0 387785361 | 0 | 4 75E-59 |
| RAB11FIP1 | 0 091660163 | 0 050152839 | 1 71E-59 | 0 437660181 | 0 | 5 24E-59 |
| GBF1 | 0 554624888 | 0 050148079 | 1 76E-59 | 0 150762003 | 0 | 5 38E-59 |
| UNC13D | 0 046924673 | 0 050143768 | 1 80E-59 | -0 43153824 | 0 | 5 50E-59 |
| DLL1 | 0 048181074 | 0 050101115 | 2 25E-59 | 0 470594971 | 0 | 6 89E-59 |
| FAR1 | 0 472711351 | 0 050098323 | 2 29E-59 | 0 201221785 | 0 | 6 99E-59 |
| USP16 | 0 316927148 | 0 050080393 | 2 52E-59 | 0 278587758 | 0 | 7 68E-59 |
| DCN | 0 669614133 | 0 050072782 | 2 62E-59 | -0 02214196 | 7 09E-13 | 8 00E-59 |
| CRACDL | 0 049075784 | 0 050070181 | 2 66E-59 | -0 42826821 | 0 | 8 10E-59 |
| NFATC2IP | 0 285022177 | 0 050049073 | 2 97E-59 | 0 263261645 | 0 | 9 06E-59 |
| ATP6V1A | 0 26845101 | 0 050044295 | 3 05E-59 | -0 22810337 | 0 | 9 29E-59 |
| CRB1 | 0 052131123 | 0 050029744 | 3 29E-59 | -0 43033725 | 0 | 1 00E-58 |
| BMP2 | 0 027574194 | 0 050026915 | 3 34E-59 | -0 46050371 | 0 | 1 02E-58 |
| TGFBR1 | 0 279720546 | 0 050023188 | 3 41E-59 | 0 291551959 | 0 | 1 04E-58 |
| SGPP1 | 0 166739639 | 0 050021645 | 3 43E-59 | 0 36532749 | 0 | 1 05E-58 |
| MYO1E | 0 45636862 | 0 050016643 | 3 53E-59 | 0 156712414 | 0 | 1 07E-58 |
| PBX3 | 0 503426548 | 0 050016072 | 3 54E-59 | 0 109998728 | 4 26E-280 | 1 08E-58 |
| FANCM | 0 149978108 | 0 050000238 | 3 85E-59 | 0 348473309 | 0 | 1 17E-58 |
| FHOD1 | 0 086510822 | 0 049964506 | 4 64E-59 | -0 39292277 | 0 | 1 41E-58 |
| DCUN1D4 | 0 289762236 | 0 049945889 | 5 12E-59 | 0 293122449 | 0 | 1 56E-58 |
| LINC01013 | 0 181940188 | 0 049940546 | 5 27E-59 | -0 26983499 | 0 | 1 60E-58 |
| INPP4A | 0 364936894 | 0 049938526 | 5 33E-59 | 0 236414339 | 0 | 1 62E-58 |
| SAMD3 | 0 062153776 | 0 049929134 | 5 60E-59 | -0 41875204 | 0 | 1 70E-58 |
| ARPP21 | 0 021520626 | 0 049906024 | 6 32E-59 | -0 46833794 | 0 | 1 92E-58 |
| SHLD2 | 0 469884449 | 0 049900541 | 6 51E-59 | -0 09246893 | 3 31E-198 | 1 98E-58 |
| EVI5 | 0 610982087 | 0 049898098 | 6 59E-59 | -0 00219104 | 0 47759267 | 2 00E-58 |
| BEND7 | 0 136519389 | 0 04988521 | 7 06E-59 | -0 33439229 | 0 | 2 14E-58 |

| | | | | | | |
|---|---|---|---|---|---|---|
| TMED3 | 0 246730502 | 0 049872697 | 7 54E-59 | -0 25377919 | 0 | 2 29E-58 |
| LINC00683 | 0 059165064 | 0 049865847 | 7 82E-59 | -0 41248334 | 0 | 2 37E-58 |
| RRP12 | 0 141497402 | 0 049858742 | 8 12E-59 | -0 33798707 | 0 | 2 46E-58 |
| DLX2 | 0 007100569 | 0 049857589 | 8 16E-59 | 0 497173606 | 0 | 2 47E-58 |
| C21orf91 | 0 099683996 | 0 049853446 | 8 35E-59 | -0 38151769 | 0 | 2 53E-58 |
| NBL1 | 0 100121833 | 0 049843576 | 8 79E-59 | -0 37576892 | 0 | 2 66E-58 |
| URI1 | 0 556319126 | 0 049840033 | 8 96E-59 | 0 131232109 | 0 | 2 71E-58 |
| NAIP | 0 266652072 | 0 049822247 | 9 84E-59 | 0 292324951 | 0 | 2 97E-58 |
| ATXN7 | 0 410985894 | 0 049797749 | 1 12E-58 | 0 224794073 | 0 | 3 38E-58 |
| FANCA | 0 145257086 | 0 049762852 | 1 34E-58 | 0 313758851 | 0 | 4 06E-58 |
| RSBN1L | 0 438065142 | 0 049762653 | 1 35E-58 | 0 177604062 | 0 | 4 07E-58 |
| FAM117A | 0 096133712 | 0 049740585 | 1 51E-58 | -0 37635031 | 0 | 4 56E-58 |
| TSHZ1 | 0 174354191 | 0 049734511 | 1 56E-58 | 0 383553323 | 0 | 4 71E-58 |
| ITSN2 | 0 489691801 | 0 04972679 | 1 63E-58 | -0 06415417 | 3 12E-96 | 4 90E-58 |
| FBXO28 | 0 292541547 | 0 049701974 | 1 85E-58 | -0 20311602 | 0 | 5 58E-58 |
| MPHOSPH9 | 0 320801051 | 0 049689348 | 1 98E-58 | 0 22619525 | 0 | 5 96E-58 |
| TFEB | 0 074613086 | 0 049665844 | 2 24E-58 | -0 39912644 | 0 | 6 74E-58 |
| FAM171B | 0 212779121 | 0 049658784 | 2 32E-58 | -0 28933463 | 0 | 7 00E-58 |
| REPS1 | 0 426928861 | 0 049653024 | 2 39E-58 | -0 10769538 | 1 64E-268 | 7 21E-58 |
| LPAR2 | 0 030448687 | 0 049629636 | 2 71E-58 | 0 478349678 | 0 | 8 15E-58 |
| WDR26 | 0 445860539 | 0 049618224 | 2 87E-58 | -0 08887133 | 3 49E-183 | 8 65E-58 |
| PPP1R12B | 0 559707601 | 0 049597624 | 3 20E-58 | -0 01650292 | 8 82E-08 | 9 63E-58 |
| GPBP1 | 0 672012716 | 0 049595066 | 3 25E-58 | 0 11115246 | 5 39E-286 | 9 76E-58 |
| CDK11B | 0 171698616 | 0 049586678 | 3 39E-58 | 0 353100372 | 0 | 1 02E-57 |
| SMG6 | 0 556414308 | 0 049581127 | 3 49E-58 | -0 05448679 | 6 67E-70 | 1 05E-57 |
| GNB5 | 0 169347623 | 0 04957347 | 3 63E-58 | 0 370918758 | 0 | 1 09E-57 |
| IGF2R | 0 48191544 | 0 04955765 | 3 95E-58 | -0 08529625 | 7 49E-169 | 1 18E-57 |
| ZNF217 | 0 246397365 | 0 049531208 | 4 54E-58 | 0 303706917 | 0 | 1 36E-57 |
| TENM3-AS1 | 0 142430184 | 0 049528161 | 4 61E-58 | 0 375585688 | 0 | 1 38E-57 |
| MLXIPL | 0 024909101 | 0 049515134 | 4 93E-58 | 0 487381524 | 0 | 1 48E-57 |
| GPATCH2 | 0 465591746 | 0 049500758 | 5 32E-58 | -0 08151015 | 2 53E-154 | 1 59E-57 |
| SPATC1 | 0 021587253 | 0 049452731 | 6 84E-58 | -0 46885798 | 0 | 2 05E-57 |
| PGM5 | 0 103424644 | 0 049448789 | 6 98E-58 | -0 38160012 | 0 | 2 09E-57 |
| HDAC4 | 0 469294321 | 0 049445985 | 7 08E-58 | -0 08866455 | 2 45E-182 | 2 12E-57 |
| CYB5R4 | 0 212541166 | 0 049443927 | 7 16E-58 | -0 2607356 | 0 | 2 14E-57 |
| ZBTB44 | 0 266518817 | 0 049431567 | 7 64E-58 | -0 22534902 | 0 | 2 28E-57 |
| SRSF1 | 0 446650549 | 0 049426623 | 7 84E-58 | 0 159918234 | 0 | 2 34E-57 |
| RBM23 | 0 207782072 | 0 049416688 | 8 26E-58 | 0 362962014 | 0 | 2 46E-57 |
| CERS4 | 0 236393748 | 0 049391796 | 9 40E-58 | -0 2319903 | 0 | 2 80E-57 |
| BTBD10 | 0 267327864 | 0 049388484 | 9 57E-58 | -0 22393062 | 0 | 2 85E-57 |
| PLEKHM3 | 0 281862139 | 0 049377274 | 1 01E-57 | -0 21824203 | 0 | 3 02E-57 |
| PCDH10 | 0 016380804 | 0 049340103 | 1 23E-57 | 0 48892003 | 0 | 3 67E-57 |
| PGBD5 | 0 047495764 | 0 049333013 | 1 28E-57 | 0 478205682 | 0 | 3 80E-57 |
| CPNE4 | 0 050113266 | 0 049330781 | 1 29E-57 | 0 462556384 | 0 | 3 85E-57 |

| | | | | | | |
|---|---|---|---|---|---|---|
| PLCB2 | 0 020749653 | 0 049312127 | 1 42E-57 | 0 488569954 | 0 | 4 24E-57 |
| SYNJ2 | 0 204241305 | 0 049262638 | 1 84E-57 | 0 298807824 | 0 | 5 49E-57 |
| NUCKS1 | 0 580133635 | 0 049259478 | 1 87E-57 | 0 140706538 | 0 | 5 57E-57 |
| CCSER1 | 0 150920409 | 0 049251743 | 1 95E-57 | 0 367124357 | 0 | 5 80E-57 |
| CATSPERE | 0 140241001 | 0 049244242 | 2 03E-57 | 0 400354499 | 0 | 6 03E-57 |
| FMO5 | 0 113932725 | 0 04924133 | 2 06E-57 | 0 395931736 | 0 | 6 12E-57 |
| PRTFDC1 | 0 164093583 | 0 049214649 | 2 37E-57 | 0 388581204 | 0 | 7 03E-57 |
| IL34 | 0 092878491 | 0 04918707 | 2 73E-57 | -0 39377049 | 0 | 8 11E-57 |
| NAV1 | 0 492023757 | 0 049140546 | 3 48E-57 | 0 125049432 | 0 | 1 03E-56 |
| NKAIN2 | 0 191553559 | 0 049133623 | 3 61E-57 | -0 2599433 | 0 | 1 07E-56 |
| RFTN2 | 0 335506653 | 0 04910714 | 4 14E-57 | -0 14269197 | 0 | 1 23E-56 |
| TTC23L | 0 072185947 | 0 049095208 | 4 40E-57 | -0 41207932 | 0 | 1 30E-56 |
| MAP3K1 | 0 29925187 | 0 049072727 | 4 95E-57 | 0 292761795 | 0 | 1 47E-56 |
| SKP2 | 0 118672784 | 0 049041079 | 5 83E-57 | 0 395927603 | 0 | 1 73E-56 |
| STEAP1B | 0 203670214 | 0 049012196 | 6 77E-57 | 0 297785873 | 0 | 2 00E-56 |
| MBOAT2 | 0 350221774 | 0 049003812 | 7 07E-57 | 0 260273813 | 0 | 2 09E-56 |
| VEGFA | 0 301897927 | 0 048994022 | 7 44E-57 | -0 21701272 | 0 | 2 20E-56 |
| ATF6B | 0 200243666 | 0 048986225 | 7 75E-57 | -0 28393688 | 0 | 2 29E-56 |
| TAOK1 | 0 603272353 | 0 048982401 | 7 90E-57 | -0 00910868 | 0 003152669 | 2 34E-56 |
| CD164 | 0 337381736 | 0 048979814 | 8 01E-57 | -0 16582406 | 0 | 2 37E-56 |
| TSPYL2 | 0 239820297 | 0 048949817 | 9 36E-57 | -0 22231158 | 0 | 2 76E-56 |
| RERE | 0 822533361 | 0 048929156 | 1 04E-56 | -0 00447292 | 0 147112086 | 3 07E-56 |
| OSBPL1A | 0 437408387 | 0 048901784 | 1 20E-56 | 0 209467061 | 0 | 3 54E-56 |
| TOX2 | 0 084150311 | 0 048888897 | 1 28E-56 | 0 422998117 | 0 | 3 78E-56 |
| TCAIM | 0 177447602 | 0 048872826 | 1 39E-56 | -0 30365431 | 0 | 4 11E-56 |
| AMPD2 | 0 099998096 | 0 048872445 | 1 40E-56 | -0 37980796 | 0 | 4 11E-56 |
| STK24 | 0 387580667 | 0 048830394 | 1 73E-56 | 0 242003715 | 0 | 5 11E-56 |
| PHF20 | 0 438417315 | 0 048785497 | 2 19E-56 | 0 187504543 | 0 | 6 44E-56 |
| RECK | 0 305029411 | 0 048768218 | 2 39E-56 | 0 275385867 | 0 | 7 04E-56 |
| MGST2 | 0 158763397 | 0 048740184 | 2 76E-56 | -0 31992198 | 0 | 8 13E-56 |
| LRRC9 | 0 037825284 | 0 048736562 | 2 81E-56 | 0 476235989 | 0 | 8 28E-56 |
| IPO11 | 0 344872551 | 0 048726269 | 2 97E-56 | 0 259533437 | 0 | 8 73E-56 |
| LATS2 | 0 326064609 | 0 048709429 | 3 24E-56 | -0 17095136 | 0 | 9 51E-56 |
| B3GAT2 | 0 043164988 | 0 048695098 | 3 48E-56 | 0 479763568 | 0 | 1 02E-55 |
| CYP26B1 | 0 05030363 | 0 048691534 | 3 55E-56 | -0 43617466 | 0 | 1 04E-55 |
| MSL2 | 0 257067256 | 0 048678235 | 3 80E-56 | 0 300689838 | 0 | 1 12E-55 |
| IL18R1 | 0 057537454 | 0 048668589 | 3 99E-56 | -0 42496765 | 0 | 1 17E-55 |
| RERGL | 0 073861149 | 0 048665652 | 4 05E-56 | 0 450765186 | 0 | 1 19E-55 |
| BASP1-AS1 | 0 058060954 | 0 048662069 | 4 13E-56 | 0 469710429 | 0 | 1 21E-55 |
| MYH15 | 0 033427881 | 0 04865601 | 4 26E-56 | 0 47219196 | 0 | 1 25E-55 |
| SMCR5 | 0 091479317 | 0 048647679 | 4 45E-56 | 0 433798014 | 0 | 1 30E-55 |
| CCNH | 0 627124936 | 0 048637067 | 4 70E-56 | -0 0268889 | 2 86E-18 | 1 38E-55 |
| CIRBP | 0 519674097 | 0 048592444 | 5 91E-56 | -0 05092111 | 2 83E-61 | 1 73E-55 |
| ARHGAP19 | 0 204879024 | 0 048585784 | 6 11E-56 | 0 369577045 | 0 | 1 79E-55 |

| | | | | | | |
|---|---|---|---|---|---|---|
| TTC33 | 0 248900649 | 0 048566081 | 6 76E-56 | -0 24458988 | 0 | 1 98E-55 |
| BMF | 0 033542099 | 0 048560829 | 6 95E-56 | -0 45166732 | 0 | 2 03E-55 |
| RHBDD1 | 0 301517199 | 0 048546696 | 7 47E-56 | -0 20831538 | 0 | 2 19E-55 |
| VCAM1 | 0 068235899 | 0 048537506 | 7 83E-56 | 0 438978088 | 0 | 2 29E-55 |
| LINC00470 | 0 010717481 | 0 048537254 | 7 84E-56 | -0 48086099 | 0 | 2 29E-55 |
| FMO3 | 0 108593021 | 0 048518271 | 8 65E-56 | -0 37379274 | 0 | 2 53E-55 |
| KLHL8 | 0 219241971 | 0 048385619 | 1 71E-55 | 0 340465186 | 0 | 4 98E-55 |
| MSH4 | 0 059945556 | 0 048384661 | 1 71E-55 | -0 41965554 | 0 | 5 00E-55 |
| MMP25-AS1 | 0 089509052 | 0 048361133 | 1 93E-55 | -0 3841216 | 0 | 5 64E-55 |
| CFAP70 | 0 141640174 | 0 048347573 | 2 07E-55 | -0 31996805 | 0 | 6 04E-55 |
| HECTD4 | 0 616093354 | 0 048336061 | 2 20E-55 | 0 103717407 | 4 30E-249 | 6 40E-55 |
| TRABD | 0 141050047 | 0 048330315 | 2 26E-55 | 0 38329732 | 0 | 6 59E-55 |
| BBOX1 | 0 046715273 | 0 048316395 | 2 43E-55 | 0 46143539 | 0 | 7 08E-55 |
| SLC22A15 | 0 184881308 | 0 048295039 | 2 71E-55 | 0 368033958 | 0 | 7 89E-55 |
| CFAP47 | 0 022082199 | 0 048292641 | 2 74E-55 | 0 487842757 | 0 | 7 98E-55 |
| CARHSP1 | 0 247415812 | 0 04824628 | 3 48E-55 | 0 307627494 | 0 | 1 01E-54 |
| METTL15 | 0 546582018 | 0 048188517 | 4 67E-55 | -0 03839472 | 1 41E-35 | 1 36E-54 |
| SNX5 | 0 234423483 | 0 048186916 | 4 70E-55 | 0 282119944 | 0 | 1 37E-54 |
| SLC24A1 | 0 086491786 | 0 04816348 | 5 30E-55 | 0 43436145 | 0 | 1 54E-54 |
| ATAD2 | 0 226637604 | 0 048159369 | 5 41E-55 | 0 244732415 | 0 | 1 57E-54 |
| JAG1 | 0 282880585 | 0 048158167 | 5 45E-55 | -0 2310089 | 0 | 1 58E-54 |
| DPH6-DT | 0 246359293 | 0 048115029 | 6 78E-55 | -0 23796456 | 0 | 1 97E-54 |
| C5AR1 | 0 079362662 | 0 048098943 | 7 36E-55 | -0 40358408 | 0 | 2 14E-54 |
| ARID4A | 0 342797586 | 0 048079343 | 8 13E-55 | 0 268033588 | 0 | 2 36E-54 |
| AGPAT5 | 0 200224629 | 0 048073861 | 8 36E-55 | 0 328056778 | 0 | 2 42E-54 |
| CDK5RAP1 | 0 184148408 | 0 048016006 | 1 12E-54 | 0 347753298 | 0 | 3 25E-54 |
| CTPS2 | 0 328234757 | 0 047984692 | 1 32E-54 | 0 246881503 | 0 | 3 81E-54 |
| DDX6 | 0 380137443 | 0 047983643 | 1 32E-54 | -0 12243256 | 0 | 3 83E-54 |
| PKD2 | 0 551550513 | 0 047950956 | 1 56E-54 | -0 03997172 | 2 03E-38 | 4 51E-54 |
| PKIA-AS1 | 0 122346805 | 0 047917196 | 1 85E-54 | 0 36281509 | 0 | 5 36E-54 |
| PCSK6 | 0 108469285 | 0 04788324 | 2 20E-54 | -0 36304092 | 0 | 6 36E-54 |
| LINC02100 | 0 134349241 | 0 047877086 | 2 27E-54 | 0 368176826 | 0 | 6 56E-54 |
| CNKSR2 | 0 167034703 | 0 047859566 | 2 48E-54 | -0 31615397 | 0 | 7 16E-54 |
| NMT2 | 0 206906398 | 0 047836415 | 2 79E-54 | 0 36313919 | 0 | 8 05E-54 |
| LINC02542 | 0 047667092 | 0 047829001 | 2 89E-54 | -0 43897051 | 0 | 8 35E-54 |
| AMOTL2 | 0 123983933 | 0 047773043 | 3 84E-54 | -0 34383725 | 0 | 1 11E-53 |
| GALM | 0 154584912 | 0 047770712 | 3 89E-54 | 0 367469128 | 0 | 1 12E-53 |
| ACACB | 0 216224705 | 0 047749143 | 4 33E-54 | -0 2690947 | 0 | 1 25E-53 |
| ZFYVE28 | 0 083550665 | 0 047744448 | 4 44E-54 | -0 3888455 | 0 | 1 28E-53 |
| VEZT | 0 479126611 | 0 047738225 | 4 58E-54 | 0 159563467 | 0 | 1 32E-53 |
| GAK | 0 322561916 | 0 047731105 | 4 75E-54 | 0 276056186 | 0 | 1 37E-53 |
| RABGEF1 | 0 628714473 | 0 047728246 | 4 81E-54 | -0 02629365 | 1 54E-17 | 1 39E-53 |
| ARHGEF18 | 0 216424587 | 0 047699158 | 5 58E-54 | 0 333318727 | 0 | 1 60E-53 |
| SMCO4 | 0 077925415 | 0 047696337 | 5 66E-54 | -0 40587716 | 0 | 1 63E-53 |

| | | | | | | |
|---|---|---|---|---|---|---|
| MYH7B | 0 033332699 | 0 047683606 | 6 03E-54 | -0 45630355 | 0 | 1 73E-53 |
| SETDB2 | 0 20999981 | 0 047658277 | 6 85E-54 | -0 26820103 | 0 | 1 97E-53 |
| ABCC10 | 0 113456816 | 0 047650826 | 7 11E-54 | -0 36047101 | 0 | 2 04E-53 |
| DRAIC | 0 038919876 | 0 047622146 | 8 22E-54 | 0 48012785 | 0 | 2 36E-53 |
| TSHR | 0 081018827 | 0 047611932 | 8 65E-54 | -0 40352342 | 0 | 2 48E-53 |
| FXYD2 | 0 007690697 | 0 047610807 | 8 70E-54 | -0 48513424 | 0 | 2 50E-53 |
| ARSJ | 0 214635168 | 0 047591646 | 9 58E-54 | 0 301124131 | 0 | 2 75E-53 |
| HES4 | 0 125716244 | 0 047591233 | 9 60E-54 | 0 418007487 | 0 | 2 75E-53 |
| SMYD2 | 0 116302755 | 0 047544151 | 1 22E-53 | 0 412416288 | 0 | 3 49E-53 |
| UTRN | 0 738982696 | 0 047537264 | 1 26E-53 | 0 089102013 | 3 96E-184 | 3 61E-53 |
| TATDN3 | 0 121033295 | 0 047526773 | 1 33E-53 | -0 35348534 | 0 | 3 80E-53 |
| TNFRSF13C | 0 012402201 | 0 047515634 | 1 40E-53 | 0 489084581 | 0 | 4 02E-53 |
| SCN2A | 0 020692543 | 0 04750977 | 1 45E-53 | 0 487581615 | 0 | 4 14E-53 |
| CYRIA | 0 086301422 | 0 047495378 | 1 55E-53 | 0 447208132 | 0 | 4 45E-53 |
| SLC22A23 | 0 204945651 | 0 047482968 | 1 65E-53 | -0 27151371 | 0 | 4 73E-53 |
| CREB5 | 0 413365441 | 0 04747374 | 1 73E-53 | -0 11652272 | 0 | 4 96E-53 |
| SLC49A4 | 0 182863452 | 0 047463912 | 1 82E-53 | -0 29605241 | 0 | 5 21E-53 |
| COL4A2 | 0 599046277 | 0 047457805 | 1 88E-53 | -0 00934982 | 0 002440574 | 5 37E-53 |
| MCM3AP-AS1 | 0 113237898 | 0 047434437 | 2 11E-53 | 0 398605804 | 0 | 6 03E-53 |
| PPM1D | 0 322533361 | 0 047422715 | 2 24E-53 | 0 235208886 | 0 | 6 39E-53 |
| EMILIN3 | 0 104319354 | 0 047416234 | 2 31E-53 | -0 37946089 | 0 | 6 60E-53 |
| VKORC1L1 | 0 360834555 | 0 047386974 | 2 68E-53 | 0 224544226 | 0 | 7 64E-53 |
| GXYLT1 | 0 217100379 | 0 047382941 | 2 73E-53 | 0 321615351 | 0 | 7 79E-53 |
| LRFN2 | 0 013572938 | 0 047367366 | 2 95E-53 | -0 47583015 | 0 | 8 42E-53 |
| ELMO1-AS1 | 0 0927262 | 0 047355658 | 3 13E-53 | -0 3840824 | 0 | 8 93E-53 |
| CARMN | 0 073090175 | 0 047313823 | 3 86E-53 | -0 39413014 | 0 | 1 10E-52 |
| SHLD1 | 0 125611544 | 0 047304766 | 4 04E-53 | -0 3597577 | 0 | 1 15E-52 |
| MED27 | 0 365488949 | 0 047285619 | 4 45E-53 | 0 242057224 | 0 | 1 27E-52 |
| KLHL3 | 0 152538501 | 0 047267998 | 4 86E-53 | 0 382483673 | 0 | 1 38E-52 |
| ANO2 | 0 096447812 | 0 047266281 | 4 90E-53 | 0 404657028 | 0 | 1 39E-52 |
| AOPEP | 0 721098018 | 0 047246307 | 5 41E-53 | 0 003111889 | 0 31313992 | 1 54E-52 |
| SHPRH | 0 481277722 | 0 047225119 | 6 02E-53 | -0 08580606 | 7 37E-171 | 1 71E-52 |
| AP3S2 | 0 321971788 | 0 047219854 | 6 18E-53 | 0 255874484 | 0 | 1 76E-52 |
| SNRNP200 | 0 324227599 | 0 047212047 | 6 42E-53 | 0 231278478 | 0 | 1 83E-52 |
| CEP295 | 0 215349032 | 0 047203187 | 6 71E-53 | 0 319332396 | 0 | 1 91E-52 |
| NUP62CL | 0 02465211 | 0 047180116 | 7 53E-53 | 0 484283964 | 0 | 2 14E-52 |
| MATN3 | 0 048552283 | 0 047178928 | 7 58E-53 | -0 43408418 | 0 | 2 15E-52 |
| LENG8 | 0 435999695 | 0 047164613 | 8 14E-53 | -0 09768161 | 4 75E-221 | 2 31E-52 |
| CDK11A | 0 241067179 | 0 047137628 | 9 31E-53 | 0 290537277 | 0 | 2 64E-52 |
| DNAJC9 | 0 255030363 | 0 047119559 | 1 02E-52 | 0 280106655 | 0 | 2 89E-52 |
| MMP9 | 0 012287982 | 0 04709993 | 1 12E-52 | 0 492808321 | 0 | 3 18E-52 |
| HSDL2 | 0 268622337 | 0 047070803 | 1 30E-52 | 0 289932081 | 0 | 3 68E-52 |
| RASSF7 | 0 021120862 | 0 047066783 | 1 33E-52 | 0 487385582 | 0 | 3 75E-52 |
| TOX3 | 0 048999638 | 0 047049209 | 1 45E-52 | -0 43589123 | 0 | 4 10E-52 |

| | | | | | | |
|---|---|---|---|---|---|---|
| DIAPH1 | 0 297586187 | 0 047039351 | 1 52E-52 | 0 24839507 | 0 | 4 30E-52 |
| ENPP2 | 0 176000838 | 0 047029335 | 1 60E-52 | -0 29954516 | 0 | 4 52E-52 |
| YES1 | 0 339199711 | 0 047016442 | 1 70E-52 | 0 227322224 | 0 | 4 82E-52 |
| ACSF2 | 0 106422874 | 0 047013782 | 1 72E-52 | 0 410271055 | 0 | 4 88E-52 |
| CMTM2 | 0 012821001 | 0 046999006 | 1 86E-52 | 0 49598157 | 0 | 5 25E-52 |
| EGLN3 | 0 175877101 | 0 046992569 | 1 92E-52 | -0 30666211 | 0 | 5 42E-52 |
| DPT | 0 030448687 | 0 046982256 | 2 02E-52 | 0 476677043 | 0 | 5 70E-52 |
| SLC44A5 | 0 080400145 | 0 046965279 | 2 20E-52 | -0 40181639 | 0 | 6 20E-52 |
| GABRB3 | 0 059383983 | 0 046948225 | 2 39E-52 | 0 45906921 | 0 | 6 74E-52 |
| FAF1 | 0 739096914 | 0 046944774 | 2 43E-52 | 0 061933851 | 8 35E-90 | 6 86E-52 |
| SUSD4 | 0 040328568 | 0 046917823 | 2 78E-52 | 0 475521446 | 0 | 7 84E-52 |
| BRD2 | 0 302973482 | 0 046885533 | 3 26E-52 | 0 289685102 | 0 | 9 20E-52 |
| NMNAT3 | 0 124850089 | 0 04688256 | 3 31E-52 | -0 34093844 | 0 | 9 33E-52 |
| PAPOLG | 0 179865222 | 0 046870563 | 3 51E-52 | 0 362336961 | 0 | 9 90E-52 |
| DDX60 | 0 178447012 | 0 046854717 | 3 80E-52 | -0 31135922 | 0 | 1 07E-51 |
| GJA1 | 0 160686071 | 0 046853611 | 3 82E-52 | 0 366625913 | 0 | 1 08E-51 |
| LRRC28 | 0 285545678 | 0 046853609 | 3 82E-52 | 0 275222401 | 0 | 1 08E-51 |
| HMCN2 | 0 042603415 | 0 046850796 | 3 87E-52 | -0 44893439 | 0 | 1 09E-51 |
| ST3GAL3 | 0 442310255 | 0 046848978 | 3 91E-52 | 0 18614387 | 0 | 1 10E-51 |
| ATRNL1 | 0 144857322 | 0 04682943 | 4 31E-52 | 0 338255607 | 0 | 1 21E-51 |
| ENPP3 | 0 019493252 | 0 046822754 | 4 45E-52 | -0 46997029 | 0 | 1 25E-51 |
| ZBTB16 | 0 465058727 | 0 046809521 | 4 75E-52 | -0 06170778 | 3 66E-89 | 1 33E-51 |
| MERTK | 0 158887133 | 0 04677746 | 5 57E-52 | -0 31062325 | 0 | 1 56E-51 |
| SH3BP1 | 0 050760503 | 0 046752595 | 6 30E-52 | -0 43180285 | 0 | 1 77E-51 |
| EMILIN2 | 0 086263349 | 0 046752518 | 6 30E-52 | -0 39619394 | 0 | 1 77E-51 |
| HAT1 | 0 267251718 | 0 046748762 | 6 42E-52 | 0 261380094 | 0 | 1 80E-51 |
| LINC02246 | 0 089766043 | 0 046733155 | 6 93E-52 | -0 39347429 | 0 | 1 94E-51 |
| CCDC138 | 0 227513278 | 0 046711084 | 7 73E-52 | 0 26113719 | 0 | 2 17E-51 |
| RSRC1 | 0 601625707 | 0 046665158 | 9 70E-52 | 0 102525539 | 2 03E-243 | 2 72E-51 |
| MAP3K5 | 0 369562734 | 0 046660168 | 9 94E-52 | -0 14869553 | 0 | 2 78E-51 |
| DENND4A | 0 499171918 | 0 04664131 | 1 09E-51 | -0 07679881 | 3 56E-137 | 3 06E-51 |
| KLHL24 | 0 35423845 | 0 046627163 | 1 17E-51 | -0 14335184 | 0 | 3 28E-51 |
| GPC5 | 0 118530011 | 0 046612785 | 1 26E-51 | -0 35982013 | 0 | 3 51E-51 |
| TTTY10 | 0 113637662 | 0 046608351 | 1 28E-51 | -0 36474604 | 0 | 3 59E-51 |
| C1orf54 | 0 114113571 | 0 046595552 | 1 37E-51 | 0 414234799 | 0 | 3 82E-51 |
| LEFTY1 | 0 013487274 | 0 046567545 | 1 57E-51 | -0 47719192 | 0 | 4 39E-51 |
| TNFRSF10B | 0 267204127 | 0 046545068 | 1 75E-51 | 0 283926192 | 0 | 4 90E-51 |
| VPS45 | 0 396927529 | 0 046511405 | 2 07E-51 | 0 207612423 | 0 | 5 78E-51 |
| RAP2A | 0 16957606 | 0 046503134 | 2 15E-51 | -0 30688367 | 0 | 6 02E-51 |
| TCP10L | 0 120890522 | 0 046481018 | 2 40E-51 | 0 423748637 | 0 | 6 71E-51 |
| HNRNPC | 0 69128705 | 0 046471838 | 2 51E-51 | 0 005197489 | 0 092052766 | 7 01E-51 |
| USP33 | 0 509156498 | 0 046470811 | 2 53E-51 | -0 07459073 | 1 71E-129 | 7 05E-51 |
| CCNT2-AS1 | 0 082655956 | 0 046399163 | 3 59E-51 | 0 432809734 | 0 | 1 00E-50 |
| LDAH | 0 325769546 | 0 046389912 | 3 76E-51 | 0 263518861 | 0 | 1 05E-50 |

| | | | | | | |
|---|---|---|---|---|---|---|
| KCNB1 | 0 121918486 | 0 046387582 | 3 80E-51 | -0 33034998 | 0 | 1 06E-50 |
| CCN1 | 0 292018046 | 0 046386669 | 3 82E-51 | -0 19035485 | 0 | 1 06E-50 |
| ZNF331 | 0 184995526 | 0 046354907 | 4 46E-51 | 0 358980209 | 0 | 1 24E-50 |
| PODN | 0 151501018 | 0 046348981 | 4 59E-51 | -0 30421133 | 0 | 1 28E-50 |
| ATF6 | 0 451933144 | 0 046311296 | 5 52E-51 | -0 08525868 | 1 05E-168 | 1 54E-50 |
| ERC1 | 0 70521216 | 0 046309083 | 5 58E-51 | 0 048257137 | 3 29E-55 | 1 55E-50 |
| SERTAD4-AS1 | 0 036597438 | 0 046296644 | 5 93E-51 | 0 477184274 | 0 | 1 65E-50 |
| OAZ2 | 0 15863966 | 0 046291863 | 6 07E-51 | -0 32237363 | 0 | 1 69E-50 |
| MMP11 | 0 039871695 | 0 046291629 | 6 08E-51 | -0 44482481 | 0 | 1 69E-50 |
| FBXO42 | 0 438398279 | 0 046278633 | 6 48E-51 | -0 11926727 | 0 | 1 80E-50 |
| ACACA | 0 538225048 | 0 046267027 | 6 86E-51 | 0 132488777 | 0 | 1 90E-50 |
| GTF2A1 | 0 198016409 | 0 046259338 | 7 12E-51 | 0 355141375 | 0 | 1 98E-50 |
| RAB7B | 0 059650492 | 0 046257418 | 7 19E-51 | -0 42371463 | 0 | 1 99E-50 |
| EFR3A | 0 463516781 | 0 046251304 | 7 41E-51 | 0 173140291 | 0 | 2 05E-50 |
| CDRT4 | 0 170832461 | 0 046230302 | 8 21E-51 | -0 31428519 | 0 | 2 27E-50 |
| CLSTN3 | 0 131084502 | 0 046215694 | 8 82E-51 | -0 35743298 | 0 | 2 44E-50 |
| RGS4 | 0 022577145 | 0 046213611 | 8 91E-51 | -0 46485893 | 0 | 2 47E-50 |
| HIP1R | 0 064895014 | 0 046208575 | 9 13E-51 | 0 453112107 | 0 | 2 53E-50 |
| IL11 | 0 012992328 | 0 046190513 | 9 97E-51 | -0 47767464 | 0 | 2 76E-50 |
| CD72 | 0 039348194 | 0 046185848 | 1 02E-50 | -0 44699719 | 0 | 2 82E-50 |
| TMEM120B | 0 264805543 | 0 046179771 | 1 05E-50 | 0 280602616 | 0 | 2 90E-50 |
| LZTS1 | 0 154832385 | 0 046152679 | 1 20E-50 | -0 31097666 | 0 | 3 31E-50 |
| ADPGK | 0 254735299 | 0 04614543 | 1 24E-50 | 0 309352266 | 0 | 3 43E-50 |
| SEC22A | 0 337343664 | 0 046118213 | 1 42E-50 | -0 17645177 | 0 | 3 92E-50 |
| PCLO | 0 179541604 | 0 046118058 | 1 42E-50 | 0 354351767 | 0 | 3 92E-50 |
| ABCA12 | 0 040813996 | 0 046111432 | 1 47E-50 | 0 451343576 | 0 | 4 05E-50 |
| GSDMB | 0 097723249 | 0 046097039 | 1 57E-50 | 0 424832919 | 0 | 4 34E-50 |
| SCLT1 | 0 473234852 | 0 046073281 | 1 77E-50 | 0 152713081 | 0 | 4 87E-50 |
| LIX1-AS1 | 0 154851421 | 0 046060478 | 1 88E-50 | 0 370728848 | 0 | 5 18E-50 |
| MCM9 | 0 257952447 | 0 046053784 | 1 94E-50 | 0 31713735 | 0 | 5 35E-50 |
| ANKRD7 | 0 081237745 | 0 046035761 | 2 12E-50 | 0 448337958 | 0 | 5 84E-50 |
| MTMR6 | 0 181902115 | 0 046021868 | 2 27E-50 | 0 370397717 | 0 | 6 25E-50 |
| MYH13 | 0 029982296 | 0 046012734 | 2 37E-50 | -0 45496741 | 0 | 6 53E-50 |
| RFX3 | 0 590489425 | 0 04600688 | 2 44E-50 | -0 04640978 | 3 41E-51 | 6 72E-50 |
| KMT2E | 0 776874607 | 0 046003621 | 2 48E-50 | 0 00740158 | 0 016435498 | 6 82E-50 |
| NEFM | 0 023205345 | 0 045957877 | 3 10E-50 | 0 487875811 | 0 | 8 52E-50 |
| TPM1 | 0 452856409 | 0 04595216 | 3 18E-50 | -0 10158072 | 5 71E-239 | 8 75E-50 |
| MAGED2 | 0 274647351 | 0 045948296 | 3 25E-50 | -0 21115652 | 0 | 8 92E-50 |
| TMEM41B | 0 320696351 | 0 045925288 | 3 63E-50 | 0 259021784 | 0 | 9 97E-50 |
| CCDC30 | 0 182930079 | 0 045924748 | 3 64E-50 | -0 28803508 | 0 | 9 99E-50 |
| MAN1C1 | 0 140364737 | 0 045889392 | 4 32E-50 | -0 33797213 | 0 | 1 19E-49 |
| PTCHD4 | 0 155403476 | 0 045878891 | 4 55E-50 | -0 32245228 | 0 | 1 25E-49 |
| SWT1 | 0 195398907 | 0 045870751 | 4 73E-50 | -0 27755781 | 0 | 1 30E-49 |
| CRIM1 | 0 442243628 | 0 045868478 | 4 78E-50 | -0 0904363 | 1 21E-189 | 1 31E-49 |

| | | | | | | |
|---|---|---|---|---|---|---|
| PITPNB | 0 491623993 | 0 045861649 | 4 94E-50 | 0 151768508 | 0 | 1 36E-49 |
| FAM241A | 0 085701776 | 0 04585034 | 5 22E-50 | 0 435086685 | 0 | 1 43E-49 |
| DUSP22 | 0 149587862 | 0 045822577 | 5 97E-50 | -0 32273714 | 0 | 1 64E-49 |
| RPH3AL | 0 101568598 | 0 045809017 | 6 38E-50 | -0 36784244 | 0 | 1 75E-49 |
| NLK | 0 438626716 | 0 045803796 | 6 54E-50 | 0 181246123 | 0 | 1 79E-49 |
| HSD17B11 | 0 241552607 | 0 045769674 | 7 72E-50 | 0 334036782 | 0 | 2 11E-49 |
| GMCL1 | 0 282033466 | 0 04574387 | 8 74E-50 | 0 300872383 | 0 | 2 39E-49 |
| ANAPC4 | 0 24242828 | 0 045740908 | 8 87E-50 | -0 24990652 | 0 | 2 43E-49 |
| EDA | 0 263882279 | 0 045735626 | 9 10E-50 | -0 22465979 | 0 | 2 49E-49 |
| REST | 0 241143325 | 0 04573066 | 9 32E-50 | -0 25298823 | 0 | 2 55E-49 |
| RAB2B | 0 157230968 | 0 04570682 | 1 05E-49 | -0 31920551 | 0 | 2 86E-49 |
| LRCH3 | 0 635872152 | 0 045699384 | 1 08E-49 | -0 01751448 | 1 37E-08 | 2 96E-49 |
| LINC01135 | 0 066436961 | 0 045675957 | 1 21E-49 | 0 44685826 | 0 | 3 31E-49 |
| GPR156 | 0 083046201 | 0 045653621 | 1 35E-49 | 0 418164225 | 0 | 3 69E-49 |
| NPFFR2 | 0 05913651 | 0 045629162 | 1 52E-49 | -0 42573361 | 0 | 4 15E-49 |
| MSH3 | 0 423292913 | 0 045603173 | 1 73E-49 | 0 204319851 | 0 | 4 70E-49 |
| SYNPO2 | 0 329281757 | 0 045601418 | 1 74E-49 | -0 16607207 | 0 | 4 74E-49 |
| RHOBTB1 | 0 167082294 | 0 045593015 | 1 81E-49 | -0 31317153 | 0 | 4 94E-49 |
| C9orf153 | 0 076240696 | 0 045591108 | 1 83E-49 | -0 40615407 | 0 | 4 98E-49 |
| NPPC | 0 020559289 | 0 04558535 | 1 88E-49 | 0 484976385 | 0 | 5 12E-49 |
| SRL | 0 007738288 | 0 045572323 | 2 00E-49 | 0 496508478 | 0 | 5 45E-49 |
| LINC01568 | 0 010232053 | 0 045569521 | 2 03E-49 | 0 495408179 | 0 | 5 52E-49 |
| KCNAB2 | 0 071814738 | 0 045558915 | 2 14E-49 | 0 44981988 | 0 | 5 81E-49 |
| CCNG2 | 0 143172603 | 0 045546838 | 2 26E-49 | 0 406685998 | 0 | 6 15E-49 |
| AZI2 | 0 295720622 | 0 045519364 | 2 58E-49 | -0 20174295 | 0 | 7 02E-49 |
| VRK1 | 0 210647047 | 0 045490015 | 2 98E-49 | 0 28417361 | 0 | 8 08E-49 |
| SNAI3 | 0 042651006 | 0 045487568 | 3 01E-49 | -0 44045979 | 0 | 8 18E-49 |
| ARHGAP31-AS | 0 027469494 | 0 045472874 | 3 23E-49 | 0 4820638 | 0 | 8 77E-49 |
| KANK2 | 0 294245303 | 0 045468936 | 3 29E-49 | -0 18130256 | 0 | 8 94E-49 |
| KIZ-AS1 | 0 150082808 | 0 04546406 | 3 37E-49 | 0 370900771 | 0 | 9 15E-49 |
| DDX3Y | 0 142201748 | 0 045463502 | 3 38E-49 | -0 34223957 | 0 | 9 17E-49 |
| SND1 | 0 676733738 | 0 045460721 | 3 43E-49 | 0 07428359 | 1 93E-128 | 9 29E-49 |
| SIRT1 | 0 183034779 | 0 045458309 | 3 47E-49 | -0 3009042 | 0 | 9 40E-49 |
| CES4A | 0 02171099 | 0 04543273 | 3 92E-49 | -0 46697355 | 0 | 1 06E-48 |
| EGFL7 | 0 040604595 | 0 045425631 | 4 05E-49 | -0 44790265 | 0 | 1 10E-48 |
| DNAJC5 | 0 249604995 | 0 045420185 | 4 16E-49 | 0 308818028 | 0 | 1 13E-48 |
| ZFP14 | 0 180893187 | 0 04541697 | 4 23E-49 | -0 30060404 | 0 | 1 14E-48 |
| ZNF451 | 0 471540614 | 0 045406814 | 4 44E-49 | 0 188197737 | 0 | 1 20E-48 |
| ULBP1 | 0 012021473 | 0 045404344 | 4 49E-49 | -0 478304 | 0 | 1 22E-48 |
| ZGRF1 | 0 13377815 | 0 045403202 | 4 52E-49 | 0 333910645 | 0 | 1 22E-48 |
| IFNG-AS1 | 0 038386857 | 0 045401294 | 4 56E-49 | -0 44820043 | 0 | 1 23E-48 |
| USP13 | 0 302678418 | 0 045363949 | 5 45E-49 | 0 239340762 | 0 | 1 47E-48 |
| RIN3 | 0 205754697 | 0 045356735 | 5 64E-49 | -0 28833786 | 0 | 1 53E-48 |
| ARID3B | 0 081932573 | 0 045346048 | 5 94E-49 | 0 442565 | 0 | 1 61E-48 |

| | | | | | | |
|---|---|---|---|---|---|---|
| CAMK2B | 0 031038815 | 0 04533947 | 6 13E-49 | -0 45610771 | 0 | 1 66E-48 |
| ESCO2 | 0 053187642 | 0 045330587 | 6 40E-49 | 0 41967413 | 0 | 1 73E-48 |
| EIF2AK2 | 0 417363081 | 0 045327692 | 6 49E-49 | 0 193543704 | 0 | 1 75E-48 |
| ZNF675 | 0 101673298 | 0 045302646 | 7 31E-49 | 0 405636073 | 0 | 1 97E-48 |
| CDH18 | 0 050712912 | 0 045221885 | 1 08E-48 | 0 467358998 | 0 | 2 90E-48 |
| TBC1D12 | 0 341798176 | 0 045206303 | 1 16E-48 | -0 17102532 | 0 | 3 13E-48 |
| LINC02664 | 0 150463536 | 0 045202 | 1 18E-48 | -0 31677554 | 0 | 3 19E-48 |
| LARP4B | 0 482762559 | 0 045196614 | 1 21E-48 | -0 07359943 | 4 07E-126 | 3 27E-48 |
| RNF19A | 0 489691801 | 0 045187374 | 1 27E-48 | 0 157880732 | 0 | 3 42E-48 |
| ALPL | 0 018750833 | 0 045156988 | 1 47E-48 | 0 488596792 | 0 | 3 95E-48 |
| MCM8 | 0 111734024 | 0 045111967 | 1 82E-48 | 0 356899182 | 0 | 4 90E-48 |
| ADAM12 | 0 332803487 | 0 04510359 | 1 89E-48 | 0 195576172 | 0 | 5 10E-48 |
| DCLRE1C | 0 173269117 | 0 045096464 | 1 96E-48 | 0 373277978 | 0 | 5 27E-48 |
| DCLK2 | 0 310845025 | 0 045067319 | 2 25E-48 | 0 264791681 | 0 | 6 05E-48 |
| TCF4-AS1 | 0 135224915 | 0 045061407 | 2 32E-48 | -0 36286934 | 0 | 6 23E-48 |
| ABCB10 | 0 172802726 | 0 045050083 | 2 44E-48 | 0 377735446 | 0 | 6 57E-48 |
| MTR | 0 447469113 | 0 045041058 | 2 55E-48 | -0 09048888 | 7 29E-190 | 6 86E-48 |
| ALDH1A1 | 0 159382079 | 0 04504049 | 2 56E-48 | -0 3171403 | 0 | 6 87E-48 |
| ABCC6 | 0 02623213 | 0 045035869 | 2 62E-48 | 0 480601126 | 0 | 7 02E-48 |
| ANKRD33B | 0 100626297 | 0 045034832 | 2 63E-48 | -0 38777535 | 0 | 7 06E-48 |
| CMSS1 | 0 542194133 | 0 045012809 | 2 92E-48 | 0 127818012 | 0 | 7 83E-48 |
| SLC6A11 | 0 023909691 | 0 045010454 | 2 95E-48 | -0 46602267 | 0 | 7 92E-48 |
| SLC35E1 | 0 134863224 | 0 044978686 | 3 43E-48 | -0 3434591 | 0 | 9 21E-48 |
| FAM117B | 0 144666959 | 0 044952292 | 3 89E-48 | -0 32352537 | 0 | 1 04E-47 |
| TNFSF15 | 0 012487864 | 0 044905304 | 4 87E-48 | -0 47719269 | 0 | 1 30E-47 |
| SEC31B | 0 148350498 | 0 04490355 | 4 91E-48 | -0 32325007 | 0 | 1 31E-47 |
| SH2D3C | 0 014362948 | 0 044864002 | 5 92E-48 | 0 492276065 | 0 | 1 58E-47 |
| RNF214 | 0 171394034 | 0 044820665 | 7 27E-48 | -0 31075499 | 0 | 1 95E-47 |
| LINC02694 | 0 037168529 | 0 044812125 | 7 57E-48 | 0 473532412 | 0 | 2 02E-47 |
| PDLIM7 | 0 32106756 | 0 044793749 | 8 26E-48 | -0 18347233 | 0 | 2 21E-47 |
| RBBP5 | 0 160676553 | 0 044791384 | 8 36E-48 | 0 384250815 | 0 | 2 23E-47 |
| PPP1R21 | 0 355704251 | 0 044780872 | 8 78E-48 | -0 14607969 | 0 | 2 34E-47 |
| CNNM2 | 0 301126954 | 0 044763545 | 9 53E-48 | -0 20786603 | 0 | 2 54E-47 |
| SMURF2 | 0 589252061 | 0 044760782 | 9 66E-48 | 0 073041508 | 3 09E-124 | 2 58E-47 |
| TCTN3 | 0 222868402 | 0 044742395 | 1 05E-47 | -0 25788856 | 0 | 2 81E-47 |
| PDE4A | 0 068883136 | 0 044686957 | 1 37E-47 | 0 444365742 | 0 | 3 65E-47 |
| SUCLG2 | 0 517418286 | 0 044685263 | 1 38E-47 | 0 148601363 | 0 | 3 68E-47 |
| MAPK8IP3 | 0 405008471 | 0 044683553 | 1 39E-47 | 0 202030696 | 0 | 3 71E-47 |
| TMEM181 | 0 308627287 | 0 044672247 | 1 47E-47 | 0 260844743 | 0 | 3 91E-47 |
| PNPT1 | 0 301507681 | 0 044656066 | 1 59E-47 | 0 264144858 | 0 | 4 22E-47 |
| PCNX1 | 0 602234871 | 0 04463788 | 1 73E-47 | 0 093336492 | 6 39E-202 | 4 60E-47 |
| LINC01229 | 0 024433192 | 0 04461442 | 1 93E-47 | 0 482794285 | 0 | 5 13E-47 |
| MICAL1 | 0 069425672 | 0 044606477 | 2 00E-47 | -0 41203571 | 0 | 5 33E-47 |
| TTPAL | 0 090927262 | 0 044578599 | 2 28E-47 | -0 39174534 | 0 | 6 07E-47 |

| | | | | | | |
|---|---|---|---|---|---|---|
| HGF | 0 210608974 | 0 044546044 | 2 66E-47 | 0 315831446 | 0 | 7 08E-47 |
| TULP4 | 0 394785936 | 0 044533604 | 2 82E-47 | 0 180323326 | 0 | 7 50E-47 |
| RASL10A | 0 012230873 | 0 04452388 | 2 96E-47 | -0 47794967 | 0 | 7 85E-47 |
| C21orf58 | 0 111876797 | 0 044509318 | 3 17E-47 | 0 356276235 | 0 | 8 40E-47 |
| BEX2 | 0 061839676 | 0 044490022 | 3 47E-47 | -0 41550419 | 0 | 9 20E-47 |
| MED6 | 0 233623955 | 0 04447012 | 3 81E-47 | 0 321663358 | 0 | 1 01E-46 |
| HCN1 | 0 038815176 | 0 044452353 | 4 14E-47 | -0 45131307 | 0 | 1 10E-46 |
| PTPN2 | 0 274780606 | 0 044431003 | 4 58E-47 | -0 22635684 | 0 | 1 21E-46 |
| MYO1B | 0 366964269 | 0 044394573 | 5 43E-47 | 0 189908139 | 0 | 1 44E-46 |
| LOX | 0 132664522 | 0 044368572 | 6 14E-47 | 0 382033721 | 0 | 1 62E-46 |
| WWTR1 | 0 672717062 | 0 044348547 | 6 74E-47 | -0 0296172 | 7 85E-22 | 1 78E-46 |
| MIPEP | 0 293426738 | 0 044347002 | 6 79E-47 | -0 20789179 | 0 | 1 80E-46 |
| DTWD2 | 0 135738897 | 0 0443433 | 6 91E-47 | -0 34719486 | 0 | 1 83E-46 |
| ZNF276 | 0 146494451 | 0 044321882 | 7 64E-47 | 0 38475347 | 0 | 2 02E-46 |
| PILRB | 0 370400335 | 0 044317903 | 7 78E-47 | 0 202976868 | 0 | 2 06E-46 |
| TMC6 | 0 042355942 | 0 044309269 | 8 10E-47 | -0 4417317 | 0 | 2 14E-46 |
| IL1RAPL1 | 0 471150368 | 0 044284795 | 9 09E-47 | 0 161338714 | 0 | 2 40E-46 |
| LINC02197 | 0 021882317 | 0 04422419 | 1 21E-46 | 0 484044996 | 0 | 3 19E-46 |
| EIF4G3 | 0 732976719 | 0 044222963 | 1 21E-46 | -0 0085249 | 0 005723463 | 3 20E-46 |
| INVS | 0 589566161 | 0 044196385 | 1 37E-46 | 0 09671916 | 9 77E-217 | 3 63E-46 |
| ZFX | 0 343968323 | 0 044161004 | 1 62E-46 | 0 256608687 | 0 | 4 28E-46 |
| DES | 0 016180922 | 0 044158403 | 1 64E-46 | 0 49032678 | 0 | 4 33E-46 |
| MFHAS1 | 0 178408939 | 0 044155873 | 1 66E-46 | 0 337784959 | 0 | 4 38E-46 |
| TSC22D3 | 0 363347357 | 0 044136328 | 1 82E-46 | -0 12463804 | 0 | 4 79E-46 |
| GFPT1 | 0 454274619 | 0 044103987 | 2 12E-46 | 0 170287849 | 0 | 5 57E-46 |
| CTNNBL1 | 0 297814624 | 0 044087921 | 2 28E-46 | 0 263534779 | 0 | 6 00E-46 |
| RGS14 | 0 023995355 | 0 04408788 | 2 28E-46 | -0 46424689 | 0 | 6 00E-46 |
| ESR1 | 0 154937085 | 0 044080451 | 2 36E-46 | -0 32946497 | 0 | 6 21E-46 |
| DBNL | 0 194370943 | 0 044063826 | 2 55E-46 | -0 29187515 | 0 | 6 71E-46 |
| EGFLAM | 0 112248006 | 0 04404903 | 2 73E-46 | -0 36492783 | 0 | 7 19E-46 |
| DHRS12 | 0 101644743 | 0 044029591 | 2 99E-46 | -0 37333603 | 0 | 7 87E-46 |
| EIF2AK3 | 0 372218309 | 0 043984646 | 3 69E-46 | -0 14218408 | 0 | 9 70E-46 |
| WARS2-AS1 | 0 331356723 | 0 043968299 | 3 98E-46 | -0 1966045 | 0 | 1 05E-45 |
| ALG13 | 0 465049209 | 0 043966335 | 4 02E-46 | -0 08647678 | 1 61E-173 | 1 06E-45 |
| LINC00621 | 0 0236527 | 0 043966007 | 4 02E-46 | 0 488335998 | 0 | 1 06E-45 |
| CABIN1 | 0 314757001 | 0 043952573 | 4 28E-46 | 0 254475236 | 0 | 1 12E-45 |
| ESRP1 | 0 026365384 | 0 043939134 | 4 56E-46 | 0 485607545 | 0 | 1 20E-45 |
| APPL2 | 0 461251452 | 0 043930041 | 4 76E-46 | 0 184694495 | 0 | 1 25E-45 |
| H2BC5 | 0 049637357 | 0 043929067 | 4 78E-46 | -0 43671356 | 0 | 1 25E-45 |
| ARHGEF6 | 0 199158592 | 0 043914525 | 5 11E-46 | -0 27570372 | 0 | 1 34E-45 |
| TRIM33 | 0 468447203 | 0 043902177 | 5 41E-46 | 0 168203351 | 0 | 1 42E-45 |
| PRAM1 | 0 018655651 | 0 043896281 | 5 56E-46 | -0 47112388 | 0 | 1 46E-45 |
| FRRS1 | 0 074594049 | 0 04389599 | 5 57E-46 | -0 40996408 | 0 | 1 46E-45 |
| ZNF14 | 0 110077859 | 0 043885483 | 5 85E-46 | -0 37268297 | 0 | 1 53E-45 |

| | | | | | | |
|---|---|---|---|---|---|---|
| SNAI3-AS1 | 0 052540405 | 0 043833297 | 7 45E-46 | -0 42563299 | 0 | 1 95E-45 |
| RIPK1 | 0 264196379 | 0 043815909 | 8 08E-46 | -0 22197225 | 0 | 2 11E-45 |
| STC1 | 0 038596257 | 0 043794655 | 8 91E-46 | 0 46923121 | 0 | 2 33E-45 |
| CDC42EP3 | 0 177057357 | 0 043760768 | 1 04E-45 | 0 357962378 | 0 | 2 73E-45 |
| KLHL28 | 0 298709334 | 0 043736871 | 1 16E-45 | -0 20587762 | 0 | 3 04E-45 |
| OARD1 | 0 10875483 | 0 043734389 | 1 18E-45 | -0 37423988 | 0 | 3 08E-45 |
| RPA3 | 0 221678628 | 0 043734149 | 1 18E-45 | 0 328836581 | 0 | 3 08E-45 |
| RFX2 | 0 238373532 | 0 043722273 | 1 25E-45 | -0 25208879 | 0 | 3 25E-45 |
| MRPS9 | 0 212065257 | 0 0437185 | 1 27E-45 | -0 28279605 | 0 | 3 31E-45 |
| CEP72 | 0 052749805 | 0 0436956 | 1 41E-45 | 0 441094268 | 0 | 3 68E-45 |
| ST7L | 0 362623974 | 0 043630705 | 1 90E-45 | 0 216323215 | 0 | 4 96E-45 |
| HOMER2 | 0 180997887 | 0 043623336 | 1 97E-45 | -0 28867824 | 0 | 5 13E-45 |
| GEN1 | 0 180645714 | 0 043622751 | 1 97E-45 | 0 299928556 | 0 | 5 14E-45 |
| ANKRD10 | 0 593754164 | 0 043622026 | 1 98E-45 | -0 02448324 | 2 07E-15 | 5 16E-45 |
| ARL5B | 0 134177914 | 0 043602281 | 2 17E-45 | -0 34634821 | 0 | 5 65E-45 |
| POGZ | 0 654442139 | 0 043602203 | 2 17E-45 | -0 00592128 | 0 054949474 | 5 65E-45 |
| RALGDS | 0 130208829 | 0 043598127 | 2 21E-45 | -0 34626454 | 0 | 5 76E-45 |
| C2orf27A | 0 251346824 | 0 043586268 | 2 34E-45 | 0 268067112 | 0 | 6 08E-45 |
| GTPBP3 | 0 069368563 | 0 043583211 | 2 37E-45 | 0 44524523 | 0 | 6 16E-45 |
| GPR89B | 0 193590451 | 0 043577055 | 2 44E-45 | 0 336600608 | 0 | 6 34E-45 |
| STK26 | 0 086272867 | 0 043572266 | 2 49E-45 | 0 435887741 | 0 | 6 48E-45 |
| FAM133B | 0 525518265 | 0 043566283 | 2 56E-45 | 0 12264014 | 0 | 6 66E-45 |
| SCIMP | 0 072871257 | 0 043509418 | 3 33E-45 | -0 41548867 | 0 | 8 65E-45 |
| HENMT1 | 0 023824028 | 0 04349813 | 3 50E-45 | -0 46468499 | 0 | 9 11E-45 |
| PKN2 | 0 660248234 | 0 043465779 | 4 07E-45 | 0 099447148 | 4 47E-229 | 1 06E-44 |
| TTC29 | 0 018312996 | 0 043461632 | 4 14E-45 | 0 490058589 | 0 | 1 08E-44 |
| WDR20 | 0 363023738 | 0 043458418 | 4 21E-45 | 0 235781278 | 0 | 1 09E-44 |
| UBE2O | 0 303516019 | 0 043445909 | 4 46E-45 | 0 250960101 | 0 | 1 16E-44 |
| TM9SF2 | 0 394195808 | 0 043428672 | 4 82E-45 | -0 12666175 | 0 | 1 25E-44 |
| IGSF1 | 0 038729512 | 0 043399909 | 5 50E-45 | -0 45430328 | 0 | 1 43E-44 |
| CCDC26 | 0 046296473 | 0 043397117 | 5 57E-45 | 0 468759058 | 0 | 1 45E-44 |
| APLP1 | 0 021187489 | 0 043390906 | 5 74E-45 | 0 483839703 | 0 | 1 49E-44 |
| SLC44A4 | 0 007024424 | 0 043347698 | 6 99E-45 | -0 48600254 | 0 | 1 81E-44 |
| ADD2 | 0 061601721 | 0 043294255 | 8 93E-45 | 0 422425161 | 0 | 2 31E-44 |
| WNT10B | 0 018065523 | 0 043254076 | 1 07E-44 | -0 47513151 | 0 | 2 78E-44 |
| VWCE | 0 088814224 | 0 043250938 | 1 09E-44 | -0 37640523 | 0 | 2 82E-44 |
| JUP | 0 06045002 | 0 043244466 | 1 12E-44 | -0 42545153 | 0 | 2 90E-44 |
| PPP2R2C | 0 018436733 | 0 043238435 | 1 15E-44 | 0 49029312 | 0 | 2 98E-44 |
| RFWD3 | 0 175810474 | 0 043226274 | 1 22E-44 | 0 313279385 | 0 | 3 15E-44 |
| PIK3C2G | 0 093297291 | 0 043221167 | 1 25E-44 | 0 428795093 | 0 | 3 23E-44 |
| NHLRC3 | 0 255344463 | 0 043215955 | 1 28E-44 | 0 332699399 | 0 | 3 30E-44 |
| AMZ1 | 0 174744437 | 0 043209113 | 1 32E-44 | 0 345599004 | 0 | 3 41E-44 |
| GAD1 | 0 049561211 | 0 043208819 | 1 32E-44 | -0 43621857 | 0 | 3 41E-44 |
| CCN2 | 0 376273058 | 0 043174267 | 1 55E-44 | -0 12431927 | 0 | 3 99E-44 |

| | | | | | | |
|---|---|---|---|---|---|---|
| DENND6A-AS1 | 0 252289125 | 0 043158888 | 1 66E-44 | 0 30041113 | 0 | 4 28E-44 |
| JPH2 | 0 017732387 | 0 043145898 | 1 76E-44 | 0 483797749 | 0 | 4 54E-44 |
| RIOK1 | 0 12725819 | 0 043125148 | 1 94E-44 | -0 34941789 | 0 | 4 99E-44 |
| DNM1L | 0 43660886 | 0 043067332 | 2 52E-44 | 0 179896498 | 0 | 6 49E-44 |
| TTBK2 | 0 378538387 | 0 043056439 | 2 65E-44 | 0 217534684 | 0 | 6 82E-44 |
| BORCS5 | 0 263158897 | 0 043051075 | 2 71E-44 | -0 22768734 | 0 | 6 99E-44 |
| NAA40 | 0 098960614 | 0 043021241 | 3 11E-44 | 0 399582364 | 0 | 8 00E-44 |
| PIGU | 0 257057737 | 0 043013545 | 3 22E-44 | 0 306935485 | 0 | 8 28E-44 |
| ISG20L2 | 0 078734462 | 0 042999928 | 3 42E-44 | -0 40617093 | 0 | 8 81E-44 |
| KCNC3 | 0 024290419 | 0 042983504 | 3 69E-44 | -0 46051703 | 0 | 9 49E-44 |
| THAP2 | 0 090470389 | 0 042983281 | 3 69E-44 | -0 39491717 | 0 | 9 50E-44 |
| SSPN | 0 252279606 | 0 042970328 | 3 92E-44 | -0 23530687 | 0 | 1 01E-43 |
| LEKR1 | 0 099712551 | 0 042968495 | 3 95E-44 | 0 42925605 | 0 | 1 02E-43 |
| LINC00571 | 0 107869639 | 0 042967116 | 3 97E-44 | -0 37251618 | 0 | 1 02E-43 |
| TRIM38 | 0 247577621 | 0 042966648 | 3 98E-44 | -0 24309533 | 0 | 1 02E-43 |
| ZCCHC24 | 0 270316575 | 0 042961173 | 4 08E-44 | -0 19488975 | 0 | 1 05E-43 |
| LTA4H | 0 18082656 | 0 042923637 | 4 84E-44 | 0 338614647 | 0 | 1 24E-43 |
| PRR5L | 0 147351088 | 0 042918315 | 4 96E-44 | -0 34029806 | 0 | 1 27E-43 |
| NOP58 | 0 42115132 | 0 042916423 | 5 00E-44 | 0 185184447 | 0 | 1 28E-43 |
| BTBD7 | 0 401934096 | 0 042904775 | 5 28E-44 | -0 13678915 | 0 | 1 35E-43 |
| OGA | 0 639241591 | 0 042898196 | 5 44E-44 | -0 02208908 | 8 03E-13 | 1 39E-43 |
| RAP2C | 0 094629838 | 0 042869149 | 6 20E-44 | -0 38660293 | 0 | 1 59E-43 |
| SBNO1 | 0 374236165 | 0 042860532 | 6 45E-44 | 0 214385117 | 0 | 1 65E-43 |
| EGR3 | 0 100302678 | 0 042856261 | 6 57E-44 | -0 36166731 | 0 | 1 68E-43 |
| TCP11L2 | 0 103852963 | 0 04284975 | 6 77E-44 | -0 36849984 | 0 | 1 73E-43 |
| ISLR | 0 256810264 | 0 042828182 | 7 47E-44 | -0 2169492 | 0 | 1 91E-43 |
| UBE4B | 0 497392016 | 0 042827908 | 7 48E-44 | -0 0797602 | 7 81E-148 | 1 91E-43 |
| PVT1 | 0 482667377 | 0 042827796 | 7 48E-44 | 0 093207557 | 2 29E-201 | 1 91E-43 |
| SARNP | 0 426767052 | 0 042808139 | 8 18E-44 | 0 184001631 | 0 | 2 09E-43 |
| SPANXA2-OT1 | 0 045201881 | 0 042806263 | 8 25E-44 | 0 464411826 | 0 | 2 11E-43 |
| GRM3 | 0 011659782 | 0 042772236 | 9 62E-44 | 0 493860094 | 0 | 2 46E-43 |
| ITGB3BP | 0 327197274 | 0 042766539 | 9 87E-44 | 0 219647261 | 0 | 2 52E-43 |
| CCDC15 | 0 0584512 | 0 042730424 | 1 16E-43 | 0 436651152 | 0 | 2 97E-43 |
| ZNF124 | 0 200786202 | 0 042709988 | 1 27E-43 | 0 342594239 | 0 | 3 25E-43 |
| TBX19 | 0 076649978 | 0 042694318 | 1 37E-43 | -0 40293747 | 0 | 3 49E-43 |
| FOXJ3 | 0 569063981 | 0 042677022 | 1 48E-43 | 0 12879215 | 0 | 3 77E-43 |
| STT3B | 0 493679922 | 0 04264766 | 1 69E-43 | 0 182604794 | 0 | 4 30E-43 |
| RNF114 | 0 234214083 | 0 042633433 | 1 80E-43 | 0 323329234 | 0 | 4 59E-43 |
| DICER1 | 0 528706859 | 0 042627899 | 1 85E-43 | -0 04191637 | 4 45E-42 | 4 70E-43 |
| USP6NL | 0 242847081 | 0 042626063 | 1 86E-43 | 0 322046721 | 0 | 4 74E-43 |
| STK10 | 0 214968304 | 0 042626004 | 1 86E-43 | -0 26741772 | 0 | 4 74E-43 |
| NOP56 | 0 204031905 | 0 042625616 | 1 87E-43 | 0 313423717 | 0 | 4 75E-43 |
| C1QL1 | 0 018351069 | 0 042617177 | 1 94E-43 | 0 485740318 | 0 | 4 93E-43 |
| AKIRIN2 | 0 170623061 | 0 042616044 | 1 95E-43 | -0 31168347 | 0 | 4 95E-43 |

| | | | | | | |
|---|---|---|---|---|---|---|
| NOSIP | 0 16132379 | 0 042585739 | 2 23E-43 | 0 371159566 | 0 | 5 68E-43 |
| LUZP2 | 0 092992709 | 0 042578816 | 2 30E-43 | 0 430837033 | 0 | 5 86E-43 |
| AGK | 0 303677828 | 0 042555461 | 2 56E-43 | -0 19710407 | 0 | 6 50E-43 |
| DAPK2 | 0 141982829 | 0 042541674 | 2 72E-43 | -0 33672215 | 0 | 6 92E-43 |
| KRI1 | 0 110896423 | 0 042524852 | 2 94E-43 | 0 401438655 | 0 | 7 46E-43 |
| UBE2K | 0 636252879 | 0 042516968 | 3 04E-43 | 0 093055515 | 1 03E-200 | 7 73E-43 |
| F8 | 0 073109212 | 0 042511158 | 3 12E-43 | -0 4104116 | 0 | 7 93E-43 |
| BMT2 | 0 310007424 | 0 04247322 | 3 70E-43 | -0 19628924 | 0 | 9 40E-43 |
| ALDH1L2 | 0 167872304 | 0 042472752 | 3 71E-43 | 0 364449178 | 0 | 9 41E-43 |
| DUBR | 0 276779425 | 0 04245181 | 4 08E-43 | -0 2308893 | 0 | 1 03E-42 |
| CHD7 | 0 298195351 | 0 042435624 | 4 39E-43 | 0 251173419 | 0 | 1 11E-42 |
| LINC00578 | 0 054187051 | 0 042424518 | 4 61E-43 | 0 468501737 | 0 | 1 17E-42 |
| GNG4 | 0 052549923 | 0 042406994 | 4 99E-43 | 0 459880712 | 0 | 1 26E-42 |
| HIF1A | 0 464144981 | 0 042392923 | 5 31E-43 | -0 1113294 | 6 63E-287 | 1 34E-42 |
| ADAMTS18 | 0 016238031 | 0 042375878 | 5 73E-43 | 0 484888171 | 0 | 1 45E-42 |
| STYX | 0 132226685 | 0 042374652 | 5 77E-43 | 0 393894605 | 0 | 1 46E-42 |
| METAP2 | 0 404903771 | 0 042362606 | 6 09E-43 | 0 208184678 | 0 | 1 54E-42 |
| CPLX1 | 0 027164912 | 0 042358958 | 6 19E-43 | -0 45724195 | 0 | 1 56E-42 |
| CBX4 | 0 068645181 | 0 042343866 | 6 62E-43 | -0 41410842 | 0 | 1 67E-42 |
| CISH | 0 010993509 | 0 04233813 | 6 79E-43 | 0 493607531 | 0 | 1 72E-42 |
| PIK3C2A | 0 464963545 | 0 042335289 | 6 88E-43 | -0 07108854 | 9 05E-118 | 1 74E-42 |
| PPP6R1 | 0 146104205 | 0 042335285 | 6 88E-43 | 0 378022391 | 0 | 1 74E-42 |
| HIVEP2 | 0 441900973 | 0 042323475 | 7 25E-43 | -0 12444873 | 0 | 1 83E-42 |
| GLB1 | 0 237973768 | 0 042267243 | 9 33E-43 | -0 25343417 | 0 | 2 35E-42 |
| NAA25 | 0 422436276 | 0 042252806 | 9 95E-43 | 0 172655583 | 0 | 2 51E-42 |
| LINC01707 | 0 021691953 | 0 042232897 | 1 09E-42 | -0 46852469 | 0 | 2 74E-42 |
| NAALADL2 | 0 74994765 | 0 042208867 | 1 21E-42 | 0 030888346 | 1 32E-23 | 3 05E-42 |
| POLG2 | 0 251965506 | 0 042183787 | 1 35E-42 | 0 287822987 | 0 | 3 41E-42 |
| PIWIL4 | 0 104614418 | 0 042179126 | 1 38E-42 | -0 3686899 | 0 | 3 48E-42 |
| CBLB | 0 767308827 | 0 042167639 | 1 46E-42 | 0 047502773 | 1 50E-53 | 3 66E-42 |
| SEPTIN7 | 0 654204184 | 0 042161909 | 1 49E-42 | 0 146744341 | 0 | 3 76E-42 |
| SEC23B | 0 171241743 | 0 042143097 | 1 62E-42 | -0 31155859 | 0 | 4 08E-42 |
| LINC01550 | 0 043345834 | 0 042128473 | 1 73E-42 | 0 468887925 | 0 | 4 36E-42 |
| SLA | 0 039938322 | 0 04210876 | 1 89E-42 | -0 4474313 | 0 | 4 76E-42 |
| CAV3 | 0 057899145 | 0 042105497 | 1 92E-42 | 0 446621201 | 0 | 4 83E-42 |
| SGMS2 | 0 16536902 | 0 04209219 | 2 04E-42 | -0 32432512 | 0 | 5 12E-42 |
| WDR89 | 0 097989758 | 0 042046404 | 2 50E-42 | 0 424101798 | 0 | 6 27E-42 |
| PRKCB | 0 115179608 | 0 042040429 | 2 56E-42 | -0 34872585 | 0 | 6 44E-42 |
| LAPTM4B | 0 237878586 | 0 042037438 | 2 60E-42 | -0 25755192 | 0 | 6 52E-42 |
| PPCDC | 0 251660924 | 0 042031999 | 2 66E-42 | 0 324208232 | 0 | 6 68E-42 |
| NR3C1 | 0 636157697 | 0 042021733 | 2 79E-42 | -0 02299502 | 9 03E-14 | 6 99E-42 |
| TBC1D9 | 0 234946984 | 0 042017199 | 2 84E-42 | 0 297572803 | 0 | 7 13E-42 |
| SEL1L3 | 0 076659496 | 0 041968252 | 3 53E-42 | 0 45669394 | 0 | 8 86E-42 |
| SH3RF3 | 0 492385449 | 0 041962857 | 3 62E-42 | 0 132763142 | 0 | 9 07E-42 |

| | | | | | | |
|---|---|---|---|---|---|---|
| ZC3H7A | 0 43855057 | 0 041937445 | 4 05E-42 | 0 17866112 | 0 | 1 02E-41 |
| SNTG2 | 0 159724734 | 0 041926919 | 4 25E-42 | -0 30515924 | 0 | 1 06E-41 |
| LRRC56 | 0 012126173 | 0 041926111 | 4 26E-42 | -0 47953923 | 0 | 1 07E-41 |
| LINC01362 | 0 060878339 | 0 041921692 | 4 35E-42 | 0 45298631 | 0 | 1 09E-41 |
| EAPP | 0 230949344 | 0 041916114 | 4 46E-42 | -0 25400097 | 0 | 1 11E-41 |
| FBLN5 | 0 145818659 | 0 041904975 | 4 68E-42 | -0 31969037 | 0 | 1 17E-41 |
| SLC12A1 | 0 044554644 | 0 041904123 | 4 70E-42 | 0 473515942 | 0 | 1 18E-41 |
| TANGO6 | 0 285916887 | 0 04186647 | 5 55E-42 | 0 284237048 | 0 | 1 39E-41 |
| ZNF83 | 0 482743523 | 0 041861673 | 5 67E-42 | 0 121929939 | 0 | 1 42E-41 |
| METTL21A | 0 161685481 | 0 041852063 | 5 92E-42 | 0 357103869 | 0 | 1 48E-41 |
| SYNJ1 | 0 19181055 | 0 041831446 | 6 48E-42 | 0 337244261 | 0 | 1 62E-41 |
| HESX1 | 0 042060878 | 0 041828625 | 6 56E-42 | 0 468519586 | 0 | 1 64E-41 |
| PXDC1 | 0 168738459 | 0 04178195 | 8 07E-42 | -0 29793243 | 0 | 2 01E-41 |
| BCL11B | 0 035835983 | 0 041772644 | 8 41E-42 | 0 466137134 | 0 | 2 10E-41 |
| RBM24 | 0 016333213 | 0 041760409 | 8 88E-42 | 0 491223484 | 0 | 2 21E-41 |
| MAEL | 0 009641926 | 0 041759392 | 8 92E-42 | 0 494087352 | 0 | 2 22E-41 |
| JKAMP | 0 196322172 | 0 041740355 | 9 70E-42 | -0 28940714 | 0 | 2 42E-41 |
| SCUBE3 | 0 128305191 | 0 041687361 | 1 23E-41 | -0 34427241 | 0 | 3 05E-41 |
| TTC26 | 0 177590375 | 0 041687254 | 1 23E-41 | 0 3686667 | 0 | 3 06E-41 |
| OCA2 | 0 052254859 | 0 041670214 | 1 32E-41 | 0 451666415 | 0 | 3 29E-41 |
| CMTR2 | 0 130094611 | 0 041662612 | 1 37E-41 | -0 35054945 | 0 | 3 40E-41 |
| PLEKHG1 | 0 407997183 | 0 04165794 | 1 40E-41 | -0 13180938 | 0 | 3 47E-41 |
| NUP205 | 0 269050656 | 0 041641335 | 1 50E-41 | 0 261213492 | 0 | 3 74E-41 |
| PROCR | 0 176667111 | 0 041641267 | 1 50E-41 | -0 32670148 | 0 | 3 74E-41 |
| GGA3 | 0 221078982 | 0 041633678 | 1 55E-41 | 0 311866926 | 0 | 3 86E-41 |
| WSCD2 | 0 021406408 | 0 041612639 | 1 70E-41 | -0 46873905 | 0 | 4 24E-41 |
| POLR3E | 0 142325484 | 0 041594841 | 1 84E-41 | -0 33845349 | 0 | 4 58E-41 |
| LBH | 0 109154594 | 0 041592385 | 1 86E-41 | 0 397579118 | 0 | 4 63E-41 |
| FBXO33 | 0 166930003 | 0 041588869 | 1 89E-41 | -0 30914993 | 0 | 4 70E-41 |
| RGS13 | 0 021672917 | 0 041576965 | 1 99E-41 | 0 486699077 | 0 | 4 95E-41 |
| PTPDC1 | 0 339266338 | 0 041547774 | 2 27E-41 | 0 265896677 | 0 | 5 63E-41 |
| LINC00393 | 0 031714607 | 0 041534291 | 2 40E-41 | -0 46013869 | 0 | 5 97E-41 |
| SPATA9 | 0 053016314 | 0 041516716 | 2 60E-41 | -0 43211295 | 0 | 6 45E-41 |
| ZNF335 | 0 145057204 | 0 041515591 | 2 61E-41 | 0 380822379 | 0 | 6 48E-41 |
| NDUFAF5 | 0 175734328 | 0 041474415 | 3 13E-41 | 0 369427686 | 0 | 7 75E-41 |
| MTMR3 | 0 391149988 | 0 041405963 | 4 22E-41 | -0 15165856 | 0 | 1 05E-40 |
| CKLF | 0 194342388 | 0 041401595 | 4 30E-41 | 0 344575381 | 0 | 1 07E-40 |
| PPP1R3E | 0 097513849 | 0 041397849 | 4 38E-41 | -0 38187752 | 0 | 1 08E-40 |
| CAT | 0 172964535 | 0 041390784 | 4 51E-41 | -0 29953562 | 0 | 1 12E-40 |
| CLEC4A | 0 063952714 | 0 04138941 | 4 54E-41 | 0 445260734 | 0 | 1 12E-40 |
| CLEC1A | 0 059022292 | 0 041378158 | 4 77E-41 | 0 460276296 | 0 | 1 18E-40 |
| PRKD1 | 0 596362148 | 0 041377733 | 4 78E-41 | -0 00314941 | 0 30734088 | 1 18E-40 |
| SH2D4B | 0 013896556 | 0 041349019 | 5 42E-41 | -0 47681405 | 0 | 1 34E-40 |
| JMJD1C | 0 890731187 | 0 041344488 | 5 53E-41 | 0 011297431 | 0 000250313 | 1 37E-40 |

| | | | | | | |
|---|---|---|---|---|---|---|
| CCNE2 | 0 057632636 | 0 041325456 | 6 01E-41 | 0 428954565 | 0 | 1 48E-40 |
| SPIRE2 | 0 074946222 | 0 041318716 | 6 19E-41 | 0 43415949 | 0 | 1 53E-40 |
| MLLT3 | 0 372332527 | 0 041310154 | 6 42E-41 | 0 195927021 | 0 | 1 59E-40 |
| BCAT1 | 0 269269574 | 0 041304129 | 6 59E-41 | 0 241621054 | 0 | 1 63E-40 |
| PIEZO1 | 0 146951324 | 0 041300301 | 6 70E-41 | 0 362848356 | 0 | 1 65E-40 |
| RESF1 | 0 235460966 | 0 041296023 | 6 83E-41 | 0 297507228 | 0 | 1 69E-40 |
| MAB21L1 | 0 038101312 | 0 041291612 | 6 96E-41 | 0 47409344 | 0 | 1 72E-40 |
| LMBRD1 | 0 334059888 | 0 041291134 | 6 98E-41 | -0 13914201 | 0 | 1 72E-40 |
| HVCN1 | 0 124593097 | 0 041265123 | 7 82E-41 | 0 391800069 | 0 | 1 93E-40 |
| METTL25 | 0 202147303 | 0 041255806 | 8 14E-41 | -0 28031396 | 0 | 2 01E-40 |
| ICAM5 | 0 007928652 | 0 041238075 | 8 80E-41 | -0 4850753 | 0 | 2 17E-40 |
| TRIL | 0 098798805 | 0 041232091 | 9 03E-41 | -0 38264746 | 0 | 2 22E-40 |
| MBD4 | 0 191477413 | 0 0412213 | 9 47E-41 | 0 352434771 | 0 | 2 33E-40 |
| SRSF10 | 0 550627249 | 0 041206717 | 1 01E-40 | 0 09441249 | 1 42E-206 | 2 48E-40 |
| DSC2 | 0 033570654 | 0 041178028 | 1 14E-40 | -0 45099173 | 0 | 2 81E-40 |
| ACTN1 | 0 327720774 | 0 041176463 | 1 15E-40 | 0 202654839 | 0 | 2 83E-40 |
| PHC3 | 0 600093278 | 0 041157748 | 1 25E-40 | -0 03653887 | 2 22E-32 | 3 07E-40 |
| SPPL3 | 0 593183073 | 0 041119201 | 1 48E-40 | -0 04991641 | 5 99E-59 | 3 63E-40 |
| MAPK13 | 0 02665093 | 0 041104448 | 1 57E-40 | -0 46158452 | 0 | 3 87E-40 |
| SHISAL2A | 0 03227618 | 0 041092717 | 1 66E-40 | -0 44872851 | 0 | 4 07E-40 |
| RSRC2 | 0 469808304 | 0 041085844 | 1 71E-40 | 0 170298225 | 0 | 4 19E-40 |
| TSPAN12 | 0 028230949 | 0 04106828 | 1 84E-40 | 0 472851136 | 0 | 4 52E-40 |
| ENTPD1 | 0 119101102 | 0 041061116 | 1 90E-40 | -0 36592036 | 0 | 4 66E-40 |
| CLASP1 | 0 739353905 | 0 041040513 | 2 08E-40 | 0 074806452 | 3 12E-130 | 5 10E-40 |
| NFIC | 0 493860768 | 0 041040333 | 2 08E-40 | 0 186963813 | 0 | 5 10E-40 |
| CMC1 | 0 313205536 | 0 041010718 | 2 37E-40 | 0 263661279 | 0 | 5 80E-40 |
| KLF5 | 0 168481468 | 0 040981103 | 2 69E-40 | -0 3224259 | 0 | 6 59E-40 |
| ZDHHC13 | 0 157897242 | 0 040973529 | 2 78E-40 | 0 350768779 | 0 | 6 81E-40 |
| PRG4 | 0 038148903 | 0 040925655 | 3 42E-40 | -0 45445611 | 0 | 8 38E-40 |
| KLRD1 | 0 085482858 | 0 040905246 | 3 74E-40 | -0 39378357 | 0 | 9 15E-40 |
| FOXO3 | 0 622727532 | 0 040867741 | 4 40E-40 | 0 002701979 | 0 38114366 | 1 08E-39 |
| PLXND1 | 0 16032438 | 0 040866862 | 4 41E-40 | -0 32174836 | 0 | 1 08E-39 |
| XKR9 | 0 096723839 | 0 040861069 | 4 53E-40 | 0 394525681 | 0 | 1 11E-39 |
| BTN2A1 | 0 137223735 | 0 040851927 | 4 71E-40 | 0 393019282 | 0 | 1 15E-39 |
| WDFY2 | 0 396927529 | 0 040842585 | 4 90E-40 | 0 192987835 | 0 | 1 20E-39 |
| EMP1 | 0 485427652 | 0 040837707 | 5 01E-40 | -0 08627744 | 1 00E-172 | 1 22E-39 |
| TPCN1 | 0 306419067 | 0 040832392 | 5 12E-40 | -0 20989251 | 0 | 1 25E-39 |
| VMO1 | 0 014705602 | 0 040805714 | 5 75E-40 | 0 492302945 | 0 | 1 40E-39 |
| ROCK2 | 0 614132607 | 0 040804577 | 5 78E-40 | 0 096530394 | 6 77E-216 | 1 41E-39 |
| GDPD3 | 0 035531401 | 0 040795799 | 6 00E-40 | -0 45213033 | 0 | 1 46E-39 |
| PPP1R2 | 0 185233481 | 0 040776773 | 6 51E-40 | -0 29479757 | 0 | 1 59E-39 |
| TNNT1 | 0 053082942 | 0 040771308 | 6 67E-40 | 0 448047625 | 0 | 1 63E-39 |
| LRRC7 | 0 184652872 | 0 040756177 | 7 12E-40 | -0 31703379 | 0 | 1 73E-39 |
| ANKZF1 | 0 123822124 | 0 040750795 | 7 29E-40 | -0 35649721 | 0 | 1 77E-39 |

| | | | | | | |
|---|---|---|---|---|---|---|
| TYMS | 0 077792161 | 0 04074772 | 7 38E-40 | 0 390759639 | 0 | 1 80E-39 |
| PPP3CC | 0 398060193 | 0 04073882 | 7 67E-40 | -0 13118136 | 0 | 1 87E-39 |
| LINC02384 | 0 031029297 | 0 040703022 | 8 95E-40 | -0 46014739 | 0 | 2 18E-39 |
| ATP8B3 | 0 021463517 | 0 040682477 | 9 78E-40 | 0 477415513 | 0 | 2 38E-39 |
| POC1B | 0 248596067 | 0 040665593 | 1 05E-39 | 0 293412241 | 0 | 2 56E-39 |
| MTHFD1L | 0 388865622 | 0 040653633 | 1 11E-39 | 0 180650735 | 0 | 2 69E-39 |
| CLCN5 | 0 353714949 | 0 040647456 | 1 14E-39 | -0 16845912 | 0 | 2 76E-39 |
| TRIM36 | 0 051693286 | 0 040591243 | 1 45E-39 | 0 444300114 | 0 | 3 51E-39 |
| LIN52 | 0 267156536 | 0 040576914 | 1 54E-39 | 0 258104181 | 0 | 3 74E-39 |
| VRK2 | 0 387371266 | 0 040568201 | 1 60E-39 | 0 195821708 | 0 | 3 88E-39 |
| P4HA1 | 0 481572786 | 0 040567435 | 1 60E-39 | -0 10611765 | 1 01E-260 | 3 89E-39 |
| TRAPPC10 | 0 350726238 | 0 040560598 | 1 65E-39 | 0 228865938 | 0 | 4 00E-39 |
| SPDYA | 0 146180351 | 0 040533443 | 1 86E-39 | 0 37265512 | 0 | 4 50E-39 |
| PRIM1 | 0 0817898 | 0 040486052 | 2 27E-39 | 0 382411342 | 0 | 5 50E-39 |
| RICTOR | 0 589147361 | 0 040480416 | 2 33E-39 | 0 11216328 | 3 26E-291 | 5 64E-39 |
| BTAF1 | 0 491395557 | 0 040470636 | 2 43E-39 | 0 179645723 | 0 | 5 88E-39 |
| SFXN5 | 0 307151967 | 0 040470367 | 2 43E-39 | -0 18438954 | 0 | 5 88E-39 |
| MEMO1 | 0 102958253 | 0 040460806 | 2 53E-39 | 0 409120881 | 0 | 6 13E-39 |
| FUT8 | 0 588262169 | 0 040453888 | 2 61E-39 | 0 118261641 | 0 | 6 31E-39 |
| MGAM | 0 031609907 | 0 040447797 | 2 68E-39 | -0 45490016 | 0 | 6 47E-39 |
| TWIST2 | 0 226628086 | 0 04042783 | 2 92E-39 | -0 2375279 | 0 | 7 05E-39 |
| LINC02698 | 0 012497382 | 0 040412014 | 3 12E-39 | 0 494570597 | 0 | 7 54E-39 |
| HAND2 | 0 015238621 | 0 040406449 | 3 20E-39 | 0 492525687 | 0 | 7 72E-39 |
| ASXL2 | 0 404818107 | 0 040400317 | 3 28E-39 | 0 212153874 | 0 | 7 92E-39 |
| CUL3 | 0 613837544 | 0 040395211 | 3 35E-39 | 0 120052034 | 0 | 8 10E-39 |
| DDX11 | 0 166339876 | 0 040387495 | 3 47E-39 | 0 29148417 | 0 | 8 37E-39 |
| CENPK | 0 14392454 | 0 040367524 | 3 78E-39 | 0 287992299 | 0 | 9 11E-39 |
| ERF | 0 115360454 | 0 040366932 | 3 78E-39 | -0 36395624 | 0 | 9 13E-39 |
| LPAR6 | 0 231425254 | 0 040321019 | 4 60E-39 | -0 26601092 | 0 | 1 11E-38 |
| PRKCD | 0 039081685 | 0 040313591 | 4 75E-39 | 0 470581344 | 0 | 1 14E-38 |
| KIF15 | 0 088195542 | 0 040310379 | 4 82E-39 | 0 382784428 | 0 | 1 16E-38 |
| PLSCR2 | 0 034960309 | 0 040300818 | 5 02E-39 | 0 47203365 | 0 | 1 21E-38 |
| PRRC2C | 0 712579239 | 0 040281727 | 5 44E-39 | 0 08692331 | 2 67E-175 | 1 31E-38 |
| ID3 | 0 179541604 | 0 04026478 | 5 85E-39 | -0 30949095 | 0 | 1 41E-38 |
| UBAP1 | 0 340579848 | 0 040262039 | 5 92E-39 | -0 17009204 | 0 | 1 42E-38 |
| KRT17 | 0 013763302 | 0 040241474 | 6 46E-39 | -0 47633325 | 0 | 1 55E-38 |
| YME1L1 | 0 399744913 | 0 040233665 | 6 68E-39 | -0 11183608 | 1 61E-289 | 1 61E-38 |
| FIG4 | 0 18334888 | 0 040228696 | 6 82E-39 | -0 28491243 | 0 | 1 64E-38 |
| ZMYND8 | 0 48764539 | 0 040227969 | 6 85E-39 | 0 177484846 | 0 | 1 65E-38 |
| SCAF11 | 0 654109002 | 0 040219784 | 7 09E-39 | 0 082876028 | 1 73E-159 | 1 70E-38 |
| ELL | 0 141202338 | 0 040216943 | 7 18E-39 | -0 33683047 | 0 | 1 72E-38 |
| NCAPG2 | 0 16405551 | 0 040207516 | 7 47E-39 | 0 291224121 | 0 | 1 79E-38 |
| CA3 | 0 050646285 | 0 040191735 | 7 99E-39 | -0 43652991 | 0 | 1 92E-38 |
| PRICKLE2 | 0 513791856 | 0 04018538 | 8 21E-39 | -0 04054958 | 1 73E-39 | 1 97E-38 |

| | | | | | | |
|---|---|---|---|---|---|---|
| GINS4 | 0 037092384 | 0 040170653 | 8 74E-39 | 0 449349226 | 0 | 2 10E-38 |
| PPP1R10 | 0 342902286 | 0 040127795 | 1 05E-38 | -0 16951896 | 0 | 2 51E-38 |
| NOSTRIN | 0 059964592 | 0 040096494 | 1 20E-38 | 0 410382357 | 0 | 2 87E-38 |
| ZNF462 | 0 480002284 | 0 040092261 | 1 22E-38 | -0 08916855 | 2 11E-184 | 2 92E-38 |
| WDR91 | 0 1276294 | 0 040068822 | 1 35E-38 | -0 34745199 | 0 | 3 23E-38 |
| TRA2B | 0 504321258 | 0 040048837 | 1 47E-38 | 0 161163917 | 0 | 3 51E-38 |
| DLGAP3 | 0 009451562 | 0 040020082 | 1 66E-38 | 0 496010211 | 0 | 3 96E-38 |
| CCDC91 | 0 6281529 | 0 039994338 | 1 85E-38 | -0 02701698 | 1 98E-18 | 4 42E-38 |
| LINC02234 | 0 0238621 | 0 039984671 | 1 92E-38 | 0 478579486 | 0 | 4 60E-38 |
| SGPL1 | 0 213064667 | 0 039961855 | 2 12E-38 | -0 28021047 | 0 | 5 06E-38 |
| UBE2J1 | 0 213578649 | 0 039958741 | 2 15E-38 | -0 26061117 | 0 | 5 13E-38 |
| DDX17 | 0 823932535 | 0 039950992 | 2 22E-38 | 0 024773041 | 9 68E-16 | 5 30E-38 |
| FBXO34 | 0 334859416 | 0 039936956 | 2 35E-38 | 0 237898511 | 0 | 5 62E-38 |
| CORIN | 0 077944452 | 0 039931199 | 2 41E-38 | -0 40795503 | 0 | 5 76E-38 |
| ADCK1 | 0 084730921 | 0 039910477 | 2 63E-38 | -0 39562274 | 0 | 6 28E-38 |
| OPA1 | 0 45826274 | 0 039902399 | 2 72E-38 | 0 190602063 | 0 | 6 50E-38 |
| AGAP2 | 0 05724239 | 0 03989594 | 2 80E-38 | -0 427896 | 0 | 6 67E-38 |
| CCNYL1 | 0 156440959 | 0 039883924 | 2 94E-38 | 0 382235132 | 0 | 7 02E-38 |
| KCNQ3 | 0 065732615 | 0 039863719 | 3 21E-38 | 0 458554292 | 0 | 7 64E-38 |
| CBX6 | 0 166549276 | 0 039861564 | 3 24E-38 | -0 30225865 | 0 | 7 71E-38 |
| IL1R1 | 0 173021644 | 0 039851979 | 3 37E-38 | -0 31875228 | 0 | 8 02E-38 |
| PRORP | 0 298547524 | 0 039832717 | 3 65E-38 | 0 268481039 | 0 | 8 70E-38 |
| HEATR1 | 0 300317908 | 0 039826096 | 3 76E-38 | 0 272920272 | 0 | 8 94E-38 |
| EPSTI1 | 0 123755497 | 0 039821781 | 3 83E-38 | 0 380285699 | 0 | 9 11E-38 |
| ARAP1 | 0 154546839 | 0 039813867 | 3 96E-38 | 0 376986197 | 0 | 9 41E-38 |
| DHX36 | 0 518217814 | 0 039775201 | 4 66E-38 | -0 06578872 | 4 14E-101 | 1 11E-37 |
| NOVA1 | 0 438997925 | 0 039739565 | 5 41E-38 | -0 06978337 | 1 51E-113 | 1 29E-37 |
| ADAM10 | 0 512506901 | 0 039725094 | 5 75E-38 | -0 07109578 | 8 57E-118 | 1 37E-37 |
| KDM5A | 0 557861073 | 0 039721769 | 5 83E-38 | 0 108575276 | 6 58E-273 | 1 39E-37 |
| NBN | 0 294759285 | 0 039711958 | 6 07E-38 | 0 274718542 | 0 | 1 44E-37 |
| PLA2R1 | 0 329843331 | 0 039685512 | 6 79E-38 | -0 1586315 | 0 | 1 61E-37 |
| ZNF385C | 0 012392682 | 0 039678415 | 6 99E-38 | 0 492925036 | 0 | 1 66E-37 |
| PRAG1 | 0 15680265 | 0 039671011 | 7 22E-38 | 0 364960212 | 0 | 1 71E-37 |
| S100PBP | 0 302021663 | 0 039660095 | 7 55E-38 | 0 267980522 | 0 | 1 79E-37 |
| GALNT16 | 0 112485961 | 0 039659931 | 7 56E-38 | -0 36234586 | 0 | 1 79E-37 |
| PROX1 | 0 058736746 | 0 039622514 | 8 84E-38 | -0 42100972 | 0 | 2 10E-37 |
| XYLT1 | 0 407188137 | 0 039607968 | 9 40E-38 | -0 11295221 | 2 55E-295 | 2 23E-37 |
| PRMT3 | 0 277084008 | 0 039595774 | 9 89E-38 | 0 286890915 | 0 | 2 34E-37 |
| RHD | 0 054967543 | 0 039548283 | 1 21E-37 | 0 444108834 | 0 | 2 86E-37 |
| LIMD2 | 0 086815404 | 0 039548013 | 1 21E-37 | -0 39766879 | 0 | 2 86E-37 |
| COL4A1 | 0 543479089 | 0 039541211 | 1 24E-37 | -0 04107424 | 1 80E-40 | 2 94E-37 |
| APBB1IP | 0 052978241 | 0 039530134 | 1 30E-37 | 0 456974905 | 0 | 3 08E-37 |
| KATNAL1 | 0 288658126 | 0 039529582 | 1 31E-37 | 0 29164942 | 0 | 3 09E-37 |
| ABCB7 | 0 269307647 | 0 039523723 | 1 34E-37 | 0 311867285 | 0 | 3 17E-37 |

| | | | | | | |
|---|---|---|---|---|---|---|
| HERC3 | 0 245331328 | 0 039507209 | 1 43E-37 | -0 24649215 | 0 | 3 39E-37 |
| EMSY | 0 367782833 | 0 039471823 | 1 66E-37 | 0 177905474 | 0 | 3 93E-37 |
| GSDMD | 0 130656184 | 0 039454805 | 1 78E-37 | 0 392153624 | 0 | 4 22E-37 |
| WDR17 | 0 116997582 | 0 039426684 | 2 01E-37 | 0 418844738 | 0 | 4 74E-37 |
| CYP19A1 | 0 035617064 | 0 039419633 | 2 07E-37 | -0 45987406 | 0 | 4 88E-37 |
| NUP98 | 0 463393044 | 0 039391131 | 2 33E-37 | 0 168048569 | 0 | 5 50E-37 |
| USP37 | 0 33657269 | 0 03938261 | 2 41E-37 | 0 216700804 | 0 | 5 69E-37 |
| SH3TC1 | 0 041670633 | 0 039382059 | 2 42E-37 | 0 468316929 | 0 | 5 71E-37 |
| SLC25A13 | 0 523005463 | 0 039343309 | 2 84E-37 | 0 135394376 | 0 | 6 70E-37 |
| FRMPD1 | 0 026869848 | 0 039341682 | 2 86E-37 | 0 482886445 | 0 | 6 75E-37 |
| RPS6KA2 | 0 260227294 | 0 039326473 | 3 05E-37 | -0 22291174 | 0 | 7 19E-37 |
| CD40 | 0 058127582 | 0 039271485 | 3 83E-37 | 0 45240057 | 0 | 9 03E-37 |
| HEATR4 | 0 080019417 | 0 039262532 | 3 98E-37 | 0 427162656 | 0 | 9 37E-37 |
| ROBO4 | 0 011107727 | 0 039243355 | 4 31E-37 | 0 492034937 | 0 | 1 01E-36 |
| BIN1 | 0 133892368 | 0 039231488 | 4 52E-37 | -0 34722362 | 0 | 1 07E-36 |
| CABP1 | 0 016133331 | 0 039230215 | 4 55E-37 | 0 489799056 | 0 | 1 07E-36 |
| ZBTB37 | 0 179256058 | 0 039227614 | 4 60E-37 | -0 29661949 | 0 | 1 08E-36 |
| AGMO | 0 410957339 | 0 039219583 | 4 75E-37 | -0 14777886 | 0 | 1 12E-36 |
| CCDC146 | 0 374797738 | 0 039196424 | 5 23E-37 | 0 19183284 | 0 | 1 23E-36 |
| SPRED2 | 0 594525138 | 0 039185473 | 5 48E-37 | -0 04159873 | 1 81E-41 | 1 29E-36 |
| CEACAM1 | 0 062115703 | 0 039169925 | 5 84E-37 | 0 443845383 | 0 | 1 37E-36 |
| SATB1 | 0 093078373 | 0 039164949 | 5 96E-37 | -0 38115038 | 0 | 1 40E-36 |
| GPC6 | 0 711322838 | 0 039152216 | 6 28E-37 | 0 040172673 | 8 66E-39 | 1 48E-36 |
| LINC00278 | 0 127286745 | 0 039125524 | 7 02E-37 | -0 34939376 | 0 | 1 65E-36 |
| FAP | 0 300698635 | 0 039120007 | 7 18E-37 | 0 188881179 | 0 | 1 69E-36 |
| SPATS2L | 0 641649693 | 0 039114854 | 7 34E-37 | -0 04250303 | 3 24E-43 | 1 72E-36 |
| DNA2 | 0 111600769 | 0 03910599 | 7 61E-37 | 0 356745756 | 0 | 1 79E-36 |
| RASGEF1B | 0 099427005 | 0 039105126 | 7 64E-37 | -0 38769782 | 0 | 1 79E-36 |
| SFMBT1 | 0 28080562 | 0 03907123 | 8 79E-37 | 0 285902569 | 0 | 2 06E-36 |
| CCDC192 | 0 111581733 | 0 039062211 | 9 12E-37 | -0 36626004 | 0 | 2 14E-36 |
| AR | 0 11333308 | 0 039034409 | 1 02E-36 | 0 391184214 | 0 | 2 40E-36 |
| RNF144A | 0 387799585 | 0 039023266 | 1 07E-36 | 0 193717778 | 0 | 2 51E-36 |
| RARRES1 | 0 221250309 | 0 039017951 | 1 10E-36 | -0 2719914 | 0 | 2 57E-36 |
| CDH26 | 0 020121452 | 0 039013647 | 1 12E-36 | -0 47172747 | 0 | 2 61E-36 |
| BMPER | 0 206116388 | 0 038994588 | 1 21E-36 | -0 25012714 | 0 | 2 83E-36 |
| ACAP1 | 0 049837239 | 0 03895671 | 1 41E-36 | -0 43376655 | 0 | 3 30E-36 |
| PPP2R3C | 0 251308751 | 0 038953856 | 1 43E-36 | 0 314185943 | 0 | 3 34E-36 |
| DNAH11 | 0 073737412 | 0 038951905 | 1 44E-36 | 0 45399358 | 0 | 3 37E-36 |
| TUBGCP6 | 0 100074242 | 0 038949263 | 1 45E-36 | 0 42122181 | 0 | 3 40E-36 |
| RC3H2 | 0 392853744 | 0 038943224 | 1 49E-36 | 0 205236459 | 0 | 3 49E-36 |
| COL19A1 | 0 057756372 | 0 038942191 | 1 50E-36 | 0 446720072 | 0 | 3 50E-36 |
| EAF2 | 0 061696903 | 0 038940827 | 1 51E-36 | -0 42614754 | 0 | 3 52E-36 |
| MAP4K2 | 0 057175763 | 0 038927862 | 1 59E-36 | 0 454960261 | 0 | 3 71E-36 |
| ANAPC16 | 0 325579182 | 0 038913834 | 1 68E-36 | -0 18301856 | 0 | 3 93E-36 |

| | | | | | | |
|---|---|---|---|---|---|---|
| ZNF33B | 0 120728713 | 0 038902438 | 1 76E-36 | -0 36208193 | 0 | 4 12E-36 |
| NEDD1 | 0 247330148 | 0 038899787 | 1 78E-36 | 0 262855114 | 0 | 4 17E-36 |
| SHROOM2 | 0 077649388 | 0 038894862 | 1 82E-36 | -0 40493505 | 0 | 4 25E-36 |
| CYREN | 0 170718243 | 0 038877834 | 1 95E-36 | 0 374790237 | 0 | 4 55E-36 |
| TSC22D4 | 0 170166188 | 0 038867211 | 2 04E-36 | -0 30184404 | 0 | 4 76E-36 |
| TCHH | 0 007528888 | 0 038861948 | 2 08E-36 | 0 495041226 | 0 | 4 86E-36 |
| SLFN13 | 0 025594411 | 0 038847445 | 2 21E-36 | 0 491513361 | 0 | 5 16E-36 |
| ADAT1 | 0 097694695 | 0 038835059 | 2 33E-36 | 0 420874223 | 0 | 5 42E-36 |
| AFAP1 | 0 421027584 | 0 038833706 | 2 34E-36 | -0 12223499 | 0 | 5 45E-36 |
| LINC00968 | 0 020978089 | 0 038816802 | 2 51E-36 | 0 484287944 | 0 | 5 84E-36 |
| OSBPL6 | 0 232586473 | 0 038813993 | 2 54E-36 | 0 219926838 | 0 | 5 91E-36 |
| ADAM28 | 0 04031905 | 0 038784393 | 2 87E-36 | 0 470359744 | 0 | 6 67E-36 |
| ZNF16 | 0 067502998 | 0 038775827 | 2 97E-36 | 0 449711698 | 0 | 6 91E-36 |
| GMIP | 0 020311816 | 0 038745819 | 3 36E-36 | 0 48463184 | 0 | 7 81E-36 |
| BOD1L1 | 0 460185414 | 0 038732402 | 3 55E-36 | 0 196246556 | 0 | 8 25E-36 |
| RNF115 | 0 630503893 | 0 038715374 | 3 80E-36 | -0 0190582 | 6 50E-10 | 8 85E-36 |
| GPR158 | 0 160809808 | 0 038678583 | 4 42E-36 | -0 31390017 | 0 | 1 03E-35 |
| FGL1 | 0 042298833 | 0 038673481 | 4 52E-36 | -0 44725543 | 0 | 1 05E-35 |
| DHTKD1 | 0 15270031 | 0 038651565 | 4 94E-36 | 0 392233306 | 0 | 1 15E-35 |
| LINC01807 | 0 031324361 | 0 038636539 | 5 25E-36 | -0 45847393 | 0 | 1 22E-35 |
| SLC12A5 | 0 012107137 | 0 038616978 | 5 69E-36 | 0 491952501 | 0 | 1 32E-35 |
| IQCB1 | 0 330642859 | 0 038519083 | 8 49E-36 | 0 260594385 | 0 | 1 97E-35 |
| PDE6B | 0 020073861 | 0 038501557 | 9 12E-36 | -0 46855483 | 0 | 2 12E-35 |
| EPAS1 | 0 149264244 | 0 038485223 | 9 75E-36 | -0 33600601 | 0 | 2 26E-35 |
| LINC02398 | 0 036454665 | 0 038475507 | 1 01E-35 | -0 44498253 | 0 | 2 35E-35 |
| PNMA3 | 0 008556852 | 0 038472176 | 1 03E-35 | -0 48533408 | 0 | 2 38E-35 |
| LRRC37B | 0 19191525 | 0 038445479 | 1 15E-35 | 0 352981255 | 0 | 2 66E-35 |
| CSGALNACT2 | 0 305676648 | 0 038441389 | 1 16E-35 | -0 18284698 | 0 | 2 70E-35 |
| ADORA1 | 0 014305838 | 0 038432272 | 1 21E-35 | -0 47762364 | 0 | 2 80E-35 |
| EVA1A | 0 135824561 | 0 038428553 | 1 23E-35 | -0 35351658 | 0 | 2 84E-35 |
| MECP2 | 0 409919857 | 0 038425076 | 1 24E-35 | -0 10989701 | 1 40E-279 | 2 88E-35 |
| CHRDL1 | 0 057832518 | 0 038395401 | 1 40E-35 | -0 43576895 | 0 | 3 25E-35 |
| HEY1 | 0 078201443 | 0 038386051 | 1 46E-35 | -0 40832058 | 0 | 3 38E-35 |
| KANSL1 | 0 796596296 | 0 038380733 | 1 49E-35 | 0 011578943 | 0 000174595 | 3 45E-35 |
| CDH12 | 0 174477927 | 0 038365252 | 1 59E-35 | 0 342324389 | 0 | 3 67E-35 |
| WBP4 | 0 240791152 | 0 038343226 | 1 74E-35 | -0 23994219 | 0 | 4 01E-35 |
| HSPA4L | 0 172012716 | 0 03833858 | 1 77E-35 | 0 358296392 | 0 | 4 09E-35 |
| N4BP2L2 | 0 826645219 | 0 038321784 | 1 89E-35 | 0 051143598 | 8 52E-62 | 4 38E-35 |
| SLC1A4 | 0 092478727 | 0 038307993 | 2 00E-35 | 0 433162983 | 0 | 4 63E-35 |
| NWD1 | 0 020654471 | 0 038296351 | 2 10E-35 | -0 46987381 | 0 | 4 85E-35 |
| BAIAP2 | 0 1511774 | 0 038290691 | 2 15E-35 | -0 31491232 | 0 | 4 96E-35 |
| TM7SF3 | 0 290637909 | 0 038281247 | 2 23E-35 | -0 20504908 | 0 | 5 15E-35 |
| SLC23A2 | 0 371047572 | 0 038270096 | 2 34E-35 | -0 13247704 | 0 | 5 39E-35 |
| GPR174 | 0 019788315 | 0 038263228 | 2 40E-35 | 0 48627466 | 0 | 5 54E-35 |

| | | | | | | |
|---|---|---|---|---|---|---|
| MANBA | 0 299232834 | 0 038250655 | 2 53E-35 | -0 21186257 | 0 | 5 83E-35 |
| PIGX | 0 133787668 | 0 038248257 | 2 55E-35 | 0 399558567 | 0 | 5 88E-35 |
| BTBD8 | 0 132521749 | 0 038248079 | 2 55E-35 | -0 35036982 | 0 | 5 88E-35 |
| MAILR | 0 19364756 | 0 038245084 | 2 58E-35 | -0 27040962 | 0 | 5 96E-35 |
| ADRB2 | 0 024614037 | 0 038239957 | 2 64E-35 | 0 482190634 | 0 | 6 08E-35 |
| RASSF1 | 0 136043479 | 0 038223599 | 2 82E-35 | 0 397522965 | 0 | 6 49E-35 |
| PFKP | 0 12742 | 0 038221263 | 2 85E-35 | 0 412910973 | 0 | 6 55E-35 |
| LRRC4C | 0 205402524 | 0 038215011 | 2 92E-35 | -0 24022649 | 0 | 6 72E-35 |
| ADCY4 | 0 018541433 | 0 038196835 | 3 14E-35 | 0 487880112 | 0 | 7 23E-35 |
| ANGPT4 | 0 003931012 | 0 038188385 | 3 25E-35 | 0 496686717 | 0 | 7 48E-35 |
| PHACTR4 | 0 518094078 | 0 03816159 | 3 62E-35 | 0 148905989 | 0 | 8 33E-35 |
| LINC01340 | 0 036359483 | 0 03814942 | 3 81E-35 | 0 466891539 | 0 | 8 75E-35 |
| FAM222B | 0 38482991 | 0 038147258 | 3 84E-35 | -0 15164249 | 0 | 8 83E-35 |
| SCARA5 | 0 123717424 | 0 038144257 | 3 89E-35 | -0 34364486 | 0 | 8 93E-35 |
| MKLN1 | 0 825598218 | 0 03813172 | 4 09E-35 | 0 035929748 | 2 30E-31 | 9 39E-35 |
| ARHGEF26-AS: | 0 071643411 | 0 038102581 | 4 60E-35 | -0 41025728 | 0 | 1 06E-34 |
| RNLS | 0 216843388 | 0 038097125 | 4 70E-35 | -0 26720019 | 0 | 1 08E-34 |
| COL4A2-AS2 | 0 011621709 | 0 038061614 | 5 42E-35 | -0 48085254 | 0 | 1 25E-34 |
| IP6K1 | 0 341227085 | 0 038057759 | 5 51E-35 | -0 18239926 | 0 | 1 26E-34 |
| FAM135B | 0 009737108 | 0 038046381 | 5 77E-35 | -0 48355022 | 0 | 1 32E-34 |
| DPYSL5 | 0 021368335 | 0 03804378 | 5 83E-35 | 0 490129682 | 0 | 1 34E-34 |
| HMGN3 | 0 338438256 | 0 03804053 | 5 91E-35 | -0 17069122 | 0 | 1 35E-34 |
| STPG4 | 0 073632712 | 0 038034112 | 6 06E-35 | 0 415915358 | 0 | 1 39E-34 |
| SLC25A53 | 0 146494451 | 0 038012232 | 6 62E-35 | -0 32753255 | 0 | 1 52E-34 |
| TIPIN | 0 12289886 | 0 038011059 | 6 65E-35 | 0 39684574 | 0 | 1 52E-34 |
| ORAI2 | 0 126353962 | 0 037999117 | 6 98E-35 | 0 398525887 | 0 | 1 60E-34 |
| PUS10 | 0 139707982 | 0 037995743 | 7 07E-35 | -0 340061 | 0 | 1 62E-34 |
| RHOQ | 0 373360492 | 0 037981968 | 7 48E-35 | -0 13297259 | 0 | 1 71E-34 |
| LUZP1 | 0 331699378 | 0 037968946 | 7 88E-35 | 0 200298448 | 0 | 1 80E-34 |
| TRPM6 | 0 012516419 | 0 037963313 | 8 06E-35 | 0 492154426 | 0 | 1 84E-34 |
| PXYLP1 | 0 186861092 | 0 037950209 | 8 49E-35 | -0 29546132 | 0 | 1 94E-34 |
| STAM | 0 254373608 | 0 037944366 | 8 70E-35 | -0 23997521 | 0 | 1 99E-34 |
| CHST2 | 0 026184539 | 0 037911739 | 9 91E-35 | -0 46200832 | 0 | 2 27E-34 |
| SPATA17 | 0 105756601 | 0 037902774 | 1 03E-34 | -0 36646435 | 0 | 2 35E-34 |
| LGALSL-DT | 0 059326874 | 0 037892824 | 1 07E-34 | -0 41634515 | 0 | 2 44E-34 |
| DMD | 0 554310788 | 0 03788992 | 1 08E-34 | -0 04604971 | 1 98E-50 | 2 47E-34 |
| WDFY3 | 0 680798005 | 0 03788267 | 1 11E-34 | 0 08431757 | 4 94E-165 | 2 54E-34 |
| RAD21 | 0 395376064 | 0 037877643 | 1 14E-34 | 0 171232637 | 0 | 2 59E-34 |
| TRIM52 | 0 159067979 | 0 037851869 | 1 26E-34 | 0 377855221 | 0 | 2 88E-34 |
| PHF12 | 0 208077135 | 0 037837156 | 1 34E-34 | 0 334169581 | 0 | 3 05E-34 |
| SOAT2 | 0 039881213 | 0 03783574 | 1 34E-34 | -0 4491072 | 0 | 3 07E-34 |
| TDO2 | 0 017998896 | 0 037805965 | 1 51E-34 | -0 47228067 | 0 | 3 45E-34 |
| IKBKB | 0 285450496 | 0 037784307 | 1 65E-34 | 0 28583727 | 0 | 3 76E-34 |
| TRIM39 | 0 053739697 | 0 03775021 | 1 89E-34 | -0 42892778 | 0 | 4 31E-34 |

| | | | | | | |
|---|---|---|---|---|---|---|
| NDEL1 | 0 297491005 | 0 037746849 | 1 92E-34 | -0 20604191 | 0 | 4 37E-34 |
| KCNH7 | 0 012069064 | 0 03773548 | 2 01E-34 | 0 492005433 | 0 | 4 57E-34 |
| FBXL20 | 0 501275437 | 0 037722716 | 2 11E-34 | -0 08282984 | 2 60E-159 | 4 81E-34 |
| LIMK2 | 0 12310826 | 0 037722286 | 2 12E-34 | 0 40395791 | 0 | 4 82E-34 |
| G3BP2 | 0 281462375 | 0 037716696 | 2 16E-34 | -0 2191435 | 0 | 4 92E-34 |
| FNBP4 | 0 619814966 | 0 037696139 | 2 35E-34 | 0 108982245 | 5 93E-275 | 5 34E-34 |
| CHN2 | 0 118130247 | 0 037694349 | 2 37E-34 | -0 36675459 | 0 | 5 38E-34 |
| F3 | 0 241723934 | 0 037669624 | 2 61E-34 | -0 23404001 | 0 | 5 94E-34 |
| PRKCZ | 0 071386419 | 0 037665294 | 2 66E-34 | -0 40585075 | 0 | 6 04E-34 |
| INTS6-AS1 | 0 07826807 | 0 037647949 | 2 85E-34 | -0 40555542 | 0 | 6 47E-34 |
| NOS1 | 0 041090023 | 0 037624097 | 3 13E-34 | -0 44054394 | 0 | 7 11E-34 |
| MED13 | 0 713074185 | 0 037623148 | 3 14E-34 | 0 054644045 | 2 69E-70 | 7 14E-34 |
| SLFN11 | 0 193885515 | 0 037610347 | 3 31E-34 | 0 344363522 | 0 | 7 51E-34 |
| CSGALNACT1 | 0 216167596 | 0 037608238 | 3 34E-34 | -0 24196098 | 0 | 7 57E-34 |
| PGAP1 | 0 42404485 | 0 037590689 | 3 58E-34 | 0 189838682 | 0 | 8 12E-34 |
| HACD4 | 0 058251318 | 0 037557637 | 4 08E-34 | 0 45915463 | 0 | 9 25E-34 |
| LNX2 | 0 090013516 | 0 037553236 | 4 15E-34 | 0 426262867 | 0 | 9 41E-34 |
| SEMA3C | 0 458538768 | 0 037462637 | 5 95E-34 | 0 171174624 | 0 | 1 35E-33 |
| MTIF3 | 0 219422817 | 0 037456192 | 6 11E-34 | -0 26839254 | 0 | 1 38E-33 |
| CLASP2 | 0 597894577 | 0 037453874 | 6 16E-34 | -0 02853334 | 2 24E-20 | 1 40E-33 |
| NUF2 | 0 144609849 | 0 037432764 | 6 70E-34 | 0 307393405 | 0 | 1 52E-33 |
| SALL1 | 0 0232339 | 0 037414257 | 7 21E-34 | 0 489269795 | 0 | 1 63E-33 |
| ANAPC10 | 0 299423198 | 0 037388233 | 7 99E-34 | 0 263825728 | 0 | 1 81E-33 |
| DDX43 | 0 026945994 | 0 037378823 | 8 30E-34 | -0 46511415 | 0 | 1 88E-33 |
| USF2 | 0 148721707 | 0 037362873 | 8 84E-34 | -0 33447265 | 0 | 2 00E-33 |
| ZNRF2 | 0 271620567 | 0 037327316 | 1 02E-33 | 0 287915981 | 0 | 2 30E-33 |
| PPM1B | 0 424606423 | 0 037316453 | 1 06E-33 | 0 215076025 | 0 | 2 40E-33 |
| PSD4 | 0 021806172 | 0 037301878 | 1 12E-33 | 0 48845702 | 0 | 2 54E-33 |
| NPDC1 | 0 203736841 | 0 037278993 | 1 23E-33 | -0 2654659 | 0 | 2 78E-33 |
| TBC1D8 | 0 316384611 | 0 037270481 | 1 27E-33 | -0 19643062 | 0 | 2 87E-33 |
| GDAP1 | 0 176257829 | 0 037269314 | 1 28E-33 | -0 29638592 | 0 | 2 89E-33 |
| SLC36A4 | 0 213140812 | 0 037261095 | 1 32E-33 | 0 344163778 | 0 | 2 98E-33 |
| CDC42EP5 | 0 200633911 | 0 037253265 | 1 36E-33 | -0 25769923 | 0 | 3 07E-33 |
| ADA | 0 059983629 | 0 037237612 | 1 45E-33 | 0 44891153 | 0 | 3 27E-33 |
| LCORL | 0 445870058 | 0 037236131 | 1 46E-33 | 0 165285176 | 0 | 3 29E-33 |
| DNAJB14 | 0 402457596 | 0 037218062 | 1 57E-33 | -0 12333834 | 0 | 3 53E-33 |
| SAMD4A | 0 568740363 | 0 03721679 | 1 57E-33 | -0 02301299 | 8 64E-14 | 3 55E-33 |
| EGR1 | 0 351497211 | 0 037208484 | 1 63E-33 | -0 12778115 | 0 | 3 66E-33 |
| WWP2 | 0 284070358 | 0 037206045 | 1 64E-33 | -0 21741471 | 0 | 3 70E-33 |
| PER1 | 0 369781653 | 0 037204171 | 1 65E-33 | -0 14284092 | 0 | 3 72E-33 |
| PLA2G5 | 0 013953665 | 0 037200775 | 1 68E-33 | -0 47845007 | 0 | 3 77E-33 |
| ZNF211 | 0 050855685 | 0 037196535 | 1 70E-33 | -0 4331926 | 0 | 3 84E-33 |
| GPR85 | 0 019007824 | 0 037082817 | 2 67E-33 | -0 47145128 | 0 | 6 00E-33 |
| FMN1 | 0 267784737 | 0 037081583 | 2 68E-33 | 0 278525792 | 0 | 6 02E-33 |

| | | | | | | |
|---|---|---|---|---|---|---|
| ZNF184 | 0 046534427 | 0 037068984 | 2 81E-33 | 0 459870128 | 0 | 6 33E-33 |
| MCM4 | 0 088033732 | 0 037054694 | 2 98E-33 | 0 369403912 | 0 | 6 69E-33 |
| NECTIN3-AS1 | 0 060126402 | 0 037033712 | 3 23E-33 | 0 443943722 | 0 | 7 26E-33 |
| PTCH1 | 0 349898155 | 0 037030691 | 3 27E-33 | -0 14183 | 0 | 7 34E-33 |
| ABCC9 | 0 075108031 | 0 037002613 | 3 65E-33 | 0 42168582 | 0 | 8 20E-33 |
| DNAJC27 | 0 121547277 | 0 036963447 | 4 26E-33 | -0 35831453 | 0 | 9 55E-33 |
| ORC4 | 0 366564505 | 0 036944101 | 4 59E-33 | 0 208483214 | 0 | 1 03E-32 |
| HELZ | 0 660971617 | 0 036933335 | 4 79E-33 | -0 03295694 | 1 19E-26 | 1 07E-32 |
| ITGB7 | 0 017313586 | 0 03692463 | 4 96E-33 | -0 47445297 | 0 | 1 11E-32 |
| LHCGR | 0 058593973 | 0 03690487 | 5 35E-33 | -0 43780451 | 0 | 1 20E-32 |
| PSIP1 | 0 321429251 | 0 036897847 | 5 50E-33 | 0 206293683 | 0 | 1 23E-32 |
| SLC15A2 | 0 051369667 | 0 03688404 | 5 81E-33 | -0 43340591 | 0 | 1 30E-32 |
| ZNF350 | 0 056852144 | 0 03687039 | 6 13E-33 | -0 42858142 | 0 | 1 37E-32 |
| H2AC6 | 0 135976852 | 0 036861348 | 6 35E-33 | -0 3544169 | 0 | 1 42E-32 |
| FRMD3 | 0 184519617 | 0 036861256 | 6 35E-33 | -0 29327818 | 0 | 1 42E-32 |
| PPP4R4 | 0 043526679 | 0 036836105 | 7 00E-33 | 0 468346675 | 0 | 1 57E-32 |
| NAA15 | 0 444394738 | 0 036752824 | 9 69E-33 | 0 172547619 | 0 | 2 17E-32 |
| SFXN2 | 0 079962308 | 0 036750435 | 9 78E-33 | 0 424831199 | 0 | 2 19E-32 |
| HTT | 0 444299556 | 0 036749799 | 9 80E-33 | 0 188942151 | 0 | 2 19E-32 |
| PAK3 | 0 306799794 | 0 036719711 | 1 10E-32 | -0 1584319 | 0 | 2 46E-32 |
| GCHFR | 0 013449201 | 0 03671585 | 1 12E-32 | 0 488999804 | 0 | 2 50E-32 |
| ALPK2 | 0 027469494 | 0 036699009 | 1 19E-32 | 0 475243335 | 0 | 2 67E-32 |
| EMC10 | 0 156412404 | 0 036686695 | 1 25E-32 | -0 32108713 | 0 | 2 80E-32 |
| STOM | 0 281338638 | 0 036679857 | 1 29E-32 | -0 23170005 | 0 | 2 87E-32 |
| ENPEP | 0 10964954 | 0 036679832 | 1 29E-32 | -0 36643532 | 0 | 2 87E-32 |
| SYTL3 | 0 054444043 | 0 036662305 | 1 38E-32 | -0 43287227 | 0 | 3 07E-32 |
| COG5 | 0 643543812 | 0 036660689 | 1 39E-32 | 0 095246686 | 3 20E-210 | 3 09E-32 |
| ARHGEF10L | 0 191011022 | 0 036641464 | 1 49E-32 | -0 30286358 | 0 | 3 33E-32 |
| ZGLP1 | 0 02123508 | 0 03661879 | 1 63E-32 | 0 486584937 | 0 | 3 63E-32 |
| CIR1 | 0 349764901 | 0 036616944 | 1 64E-32 | -0 1765401 | 0 | 3 66E-32 |
| ADAMTS6 | 0 334393025 | 0 036597694 | 1 77E-32 | 0 184408172 | 0 | 3 94E-32 |
| STK38 | 0 439530944 | 0 036571518 | 1 96E-32 | -0 09783666 | 9 49E-222 | 4 36E-32 |
| CCDC126 | 0 129666292 | 0 036558845 | 2 06E-32 | -0 34663315 | 0 | 4 58E-32 |
| SKI | 0 348441872 | 0 036500856 | 2 58E-32 | -0 15626211 | 0 | 5 73E-32 |
| SPTB | 0 011507491 | 0 036494065 | 2 64E-32 | 0 491931284 | 0 | 5 88E-32 |
| DNHD1 | 0 278292818 | 0 036483775 | 2 75E-32 | 0 264247788 | 0 | 6 11E-32 |
| SYTL1 | 0 030772306 | 0 036465701 | 2 95E-32 | 0 478096927 | 0 | 6 55E-32 |
| BRCA1 | 0 186194818 | 0 036462808 | 2 98E-32 | 0 30138372 | 0 | 6 63E-32 |
| COL6A1 | 0 663855628 | 0 036438722 | 3 27E-32 | -0 04879563 | 2 08E-56 | 7 27E-32 |
| MIR3659HG | 0 037863357 | 0 0364333 | 3 34E-32 | -0 44562915 | 0 | 7 42E-32 |
| NRP1 | 0 449077687 | 0 03642389 | 3 47E-32 | -0 12368026 | 0 | 7 69E-32 |
| LRRC49 | 0 209409682 | 0 036399081 | 3 82E-32 | -0 28467811 | 0 | 8 46E-32 |
| FANCI | 0 161038244 | 0 036390333 | 3 95E-32 | 0 301189138 | 0 | 8 75E-32 |
| KCNK6 | 0 028230949 | 0 03638161 | 4 08E-32 | -0 45976818 | 0 | 9 05E-32 |

| | | | | | | |
|---|---|---|---|---|---|---|
| GABPB2 | 0 234709029 | 0 0363812 | 4 09E-32 | -0 24604851 | 0 | 9 06E-32 |
| COL10A1 | 0 055157907 | 0 036355696 | 4 51E-32 | 0 449964605 | 0 | 9 99E-32 |
| CHMP3 | 0 278692582 | 0 03635366 | 4 55E-32 | 0 289939465 | 0 | 1 01E-31 |
| SLC7A2 | 0 190420894 | 0 036339478 | 4 80E-32 | -0 26048571 | 0 | 1 06E-31 |
| VCPKMT | 0 117987474 | 0 036308782 | 5 40E-32 | 0 408976485 | 0 | 1 20E-31 |
| AP4B1 | 0 077040224 | 0 036306969 | 5 44E-32 | 0 431884639 | 0 | 1 20E-31 |
| CD1D | 0 014248729 | 0 036290531 | 5 80E-32 | -0 47859094 | 0 | 1 28E-31 |
| VTI1A | 0 625287925 | 0 036285167 | 5 92E-32 | 0 088733405 | 1 28E-182 | 1 31E-31 |
| RAMP1 | 0 049085302 | 0 036275918 | 6 13E-32 | -0 43461785 | 0 | 1 36E-31 |
| POLG | 0 158925206 | 0 036255092 | 6 64E-32 | 0 35714169 | 0 | 1 47E-31 |
| TSGA10 | 0 247729912 | 0 036244888 | 6 91E-32 | -0 23970955 | 0 | 1 53E-31 |
| EFCAB13 | 0 380670461 | 0 036233099 | 7 23E-32 | -0 14825343 | 0 | 1 60E-31 |
| AHCTF1 | 0 435723668 | 0 036231401 | 7 27E-32 | 0 198903331 | 0 | 1 61E-31 |
| PDE1B | 0 010831699 | 0 036208568 | 7 94E-32 | 0 494269316 | 0 | 1 75E-31 |
| NREP-AS1 | 0 026051284 | 0 036204368 | 8 07E-32 | 0 485760407 | 0 | 1 78E-31 |
| TMEM170A | 0 115655518 | 0 03615854 | 9 62E-32 | -0 36841758 | 0 | 2 12E-31 |
| BMP6 | 0 081627991 | 0 036156611 | 9 69E-32 | -0 38909292 | 0 | 2 14E-31 |
| ATF5 | 0 053654033 | 0 036141377 | 1 03E-31 | 0 455050213 | 0 | 2 26E-31 |
| RAB31 | 0 281567075 | 0 036112735 | 1 15E-31 | -0 22504543 | 0 | 2 53E-31 |
| COX6B2 | 0 008404561 | 0 036112549 | 1 15E-31 | 0 493662368 | 0 | 2 53E-31 |
| GRM5 | 0 02160629 | 0 036087532 | 1 26E-31 | 0 488716433 | 0 | 2 78E-31 |
| SPRY3 | 0 058527346 | 0 036083557 | 1 28E-31 | -0 42075351 | 0 | 2 82E-31 |
| KIAA1671 | 0 302021663 | 0 036072444 | 1 34E-31 | -0 18274949 | 0 | 2 94E-31 |
| NFKBID | 0 034417772 | 0 036060981 | 1 40E-31 | 0 474651941 | 0 | 3 07E-31 |
| IQCH-AS1 | 0 176905066 | 0 036041068 | 1 51E-31 | -0 31230195 | 0 | 3 32E-31 |
| WTAP | 0 402800251 | 0 036011516 | 1 69E-31 | 0 218444126 | 0 | 3 71E-31 |
| CBFA2T3 | 0 021054235 | 0 036007645 | 1 71E-31 | -0 46791428 | 0 | 3 76E-31 |
| LINC01478 | 0 012725819 | 0 036003885 | 1 74E-31 | 0 492319922 | 0 | 3 82E-31 |
| PCDH11X | 0 05361596 | 0 035999948 | 1 76E-31 | 0 461204844 | 0 | 3 88E-31 |
| RNF145 | 0 454150882 | 0 035992731 | 1 81E-31 | 0 162992696 | 0 | 3 98E-31 |
| CREBL2 | 0 13588167 | 0 03595057 | 2 13E-31 | -0 34628223 | 0 | 4 68E-31 |
| BCO2 | 0 096838058 | 0 035946216 | 2 16E-31 | -0 37526472 | 0 | 4 75E-31 |
| ADGRF5 | 0 026412975 | 0 035939215 | 2 22E-31 | 0 481299615 | 0 | 4 88E-31 |
| SMYD4 | 0 142658621 | 0 035880506 | 2 78E-31 | 0 384670253 | 0 | 6 10E-31 |
| CLCN7 | 0 190430413 | 0 035876429 | 2 82E-31 | -0 30048406 | 0 | 6 19E-31 |
| ZMIZ1 | 0 391435533 | 0 035874433 | 2 84E-31 | -0 11630339 | 0 | 6 24E-31 |
| LINC01163 | 0 005387295 | 0 035870436 | 2 89E-31 | 0 497095201 | 0 | 6 33E-31 |
| SLC27A1 | 0 224343721 | 0 035858817 | 3 02E-31 | -0 25787032 | 0 | 6 61E-31 |
| ZFAND2B | 0 081694618 | 0 03579139 | 3 90E-31 | 0 438684672 | 0 | 8 54E-31 |
| LINC02325 | 0 011688336 | 0 035773016 | 4 18E-31 | -0 48477254 | 0 | 9 15E-31 |
| RUNX3 | 0 034922236 | 0 035766515 | 4 28E-31 | 0 459181825 | 0 | 9 38E-31 |
| UPK2 | 0 010155908 | 0 035766327 | 4 29E-31 | -0 48210098 | 0 | 9 38E-31 |
| RAB3GAP1 | 0 552340523 | 0 035764913 | 4 31E-31 | 0 130470043 | 0 | 9 43E-31 |
| DOCK3 | 0 160695589 | 0 035653732 | 6 56E-31 | 0 351501477 | 0 | 1 43E-30 |

| | | | | | | |
|---|---|---|---|---|---|---|
| CENPU | 0 078763016 | 0 035643594 | 6 82E-31 | 0 377825445 | 0 | 1 49E-30 |
| RPP38 | 0 182092479 | 0 035641479 | 6 88E-31 | -0 30768907 | 0 | 1 50E-30 |
| ERO1B | 0 250690069 | 0 035618136 | 7 51E-31 | -0 22450577 | 0 | 1 64E-30 |
| PARPBP | 0 117235537 | 0 035564313 | 9 20E-31 | 0 32781987 | 0 | 2 01E-30 |
| SLC9B1 | 0 05219775 | 0 035561388 | 9 30E-31 | -0 4334235 | 0 | 2 03E-30 |
| ALG14 | 0 199177628 | 0 035551324 | 9 66E-31 | -0 29290073 | 0 | 2 11E-30 |
| GCH1 | 0 04515429 | 0 035538645 | 1 01E-30 | -0 44029792 | 0 | 2 21E-30 |
| LINC01725 | 0 109107003 | 0 035518077 | 1 09E-30 | -0 36843924 | 0 | 2 39E-30 |
| TRPM3 | 0 229873789 | 0 035497681 | 1 18E-30 | 0 310621338 | 0 | 2 57E-30 |
| ADD1 | 0 524318974 | 0 035484749 | 1 24E-30 | -0 0606609 | 3 20E-86 | 2 70E-30 |
| PDZRN4 | 0 129514001 | 0 035484349 | 1 24E-30 | -0 35078819 | 0 | 2 71E-30 |
| BCAM | 0 062191849 | 0 035473098 | 1 30E-30 | -0 42445657 | 0 | 2 82E-30 |
| RTN4RL1 | 0 029125659 | 0 035471551 | 1 30E-30 | -0 45716871 | 0 | 2 84E-30 |
| POLQ | 0 104595382 | 0 035442121 | 1 46E-30 | 0 333809747 | 0 | 3 17E-30 |
| SETD5 | 0 754916145 | 0 03543523 | 1 50E-30 | 0 025192982 | 3 16E-16 | 3 25E-30 |
| LIPE | 0 064342959 | 0 035433858 | 1 50E-30 | 0 438491825 | 0 | 3 27E-30 |
| LRRC36 | 0 019264815 | 0 035433706 | 1 50E-30 | 0 485671579 | 0 | 3 27E-30 |
| UBL3 | 0 343454341 | 0 035416356 | 1 60E-30 | -0 15058755 | 0 | 3 49E-30 |
| RDH10 | 0 106032628 | 0 035410845 | 1 64E-30 | -0 37330781 | 0 | 3 56E-30 |
| AZIN1 | 0 372694219 | 0 035395807 | 1 73E-30 | 0 208950238 | 0 | 3 77E-30 |
| SRPK1 | 0 290647427 | 0 035387377 | 1 79E-30 | 0 229439794 | 0 | 3 89E-30 |
| MAN1A2 | 0 457615503 | 0 035371366 | 1 90E-30 | -0 09946579 | 3 67E-229 | 4 12E-30 |
| CNTNAP5 | 0 026831776 | 0 035366359 | 1 94E-30 | -0 46196583 | 0 | 4 20E-30 |
| ADRA1B | 0 064066932 | 0 035360575 | 1 98E-30 | 0 436402792 | 0 | 4 29E-30 |
| USP47 | 0 627991091 | 0 035343222 | 2 11E-30 | -0 00187981 | 0 54232533 | 4 58E-30 |
| TGFA | 0 025765738 | 0 035329084 | 2 23E-30 | 0 481645101 | 0 | 4 82E-30 |
| CDIN1 | 0 1855571 | 0 035317395 | 2 33E-30 | -0 28006709 | 0 | 5 04E-30 |
| ADAMTS12 | 0 382488435 | 0 035310497 | 2 39E-30 | -0 15419374 | 0 | 5 17E-30 |
| CD226 | 0 042936552 | 0 03530248 | 2 46E-30 | 0 458066649 | 0 | 5 32E-30 |
| MAP3K4 | 0 423054958 | 0 035302074 | 2 46E-30 | -0 11871637 | 0 | 5 33E-30 |
| CYP2R1 | 0 078410843 | 0 035293505 | 2 54E-30 | -0 40735194 | 0 | 5 50E-30 |
| TOM1 | 0 118758447 | 0 035249981 | 2 99E-30 | -0 36027558 | 0 | 6 47E-30 |
| ABCC4 | 0 142525366 | 0 035242662 | 3 08E-30 | 0 354326939 | 0 | 6 65E-30 |
| HNRNPU | 0 653014411 | 0 035233055 | 3 19E-30 | 0 122861193 | 0 | 6 89E-30 |
| CFAP69 | 0 217471588 | 0 035226693 | 3 26E-30 | -0 24719214 | 0 | 7 05E-30 |
| SOX9-AS1 | 0 068131199 | 0 03522028 | 3 34E-30 | -0 41219506 | 0 | 7 22E-30 |
| VWA3A | 0 023186309 | 0 035192832 | 3 70E-30 | -0 46428016 | 0 | 8 00E-30 |
| KIF26B | 0 274932897 | 0 035188201 | 3 77E-30 | 0 273632914 | 0 | 8 14E-30 |
| CEP350 | 0 604167063 | 0 035174288 | 3 97E-30 | 0 138556548 | 0 | 8 57E-30 |
| CDC37 | 0 311016352 | 0 035170617 | 4 02E-30 | 0 26736805 | 0 | 8 68E-30 |
| ABR | 0 315451828 | 0 035152294 | 4 31E-30 | -0 19376628 | 0 | 9 29E-30 |
| HCG20 | 0 007871543 | 0 035143061 | 4 46E-30 | -0 48444366 | 0 | 9 62E-30 |
| CNTRL | 0 3135006 | 0 035133138 | 4 63E-30 | 0 246308618 | 0 | 9 98E-30 |
| SLAIN1 | 0 038567703 | 0 035120046 | 4 86E-30 | 0 479236367 | 0 | 1 05E-29 |

| | | | | | | |
|---|---|---|---|---|---|---|
| CCDC175 | 0 016009594 | 0 035110941 | 5 03E-30 | 0 488539358 | 0 | 1 08E-29 |
| FAM91A1 | 0 318450058 | 0 035109274 | 5 06E-30 | 0 235023483 | 0 | 1 09E-29 |
| TSPAN32 | 0 013144619 | 0 035108435 | 5 07E-30 | -0 4783894 | 0 | 1 09E-29 |
| TMEM161B | 0 313329272 | 0 035101002 | 5 21E-30 | 0 259008816 | 0 | 1 12E-29 |
| CPQ | 0 622099332 | 0 035099171 | 5 25E-30 | -0 01802288 | 5 15E-09 | 1 13E-29 |
| COL6A3 | 0 656583732 | 0 035067328 | 5 91E-30 | -0 04852103 | 8 52E-56 | 1 27E-29 |
| LINC01588 | 0 061192439 | 0 035059776 | 6 08E-30 | 0 453448733 | 0 | 1 31E-29 |
| LTK | 0 024737774 | 0 035058064 | 6 12E-30 | 0 47804574 | 0 | 1 32E-29 |
| STC2 | 0 017427805 | 0 035057288 | 6 14E-30 | -0 47320175 | 0 | 1 32E-29 |
| PIM1 | 0 048599874 | 0 035055044 | 6 19E-30 | -0 4356044 | 0 | 1 33E-29 |
| HNRNPUL1 | 0 307494622 | 0 035034569 | 6 68E-30 | 0 242429009 | 0 | 1 43E-29 |
| VWC2L | 0 011897737 | 0 035017489 | 7 11E-30 | -0 480621 | 0 | 1 53E-29 |
| LINC02755 | 0 02744094 | 0 035004686 | 7 46E-30 | 0 475879451 | 0 | 1 60E-29 |
| TMEM164 | 0 168176886 | 0 034998616 | 7 63E-30 | 0 371161975 | 0 | 1 64E-29 |
| PRSS3 | 0 015781158 | 0 034998429 | 7 63E-30 | 0 488430397 | 0 | 1 64E-29 |
| DCTN4 | 0 34598618 | 0 03498931 | 7 90E-30 | -0 17346539 | 0 | 1 70E-29 |
| GPR35 | 0 02223449 | 0 03498183 | 8 12E-30 | -0 46628949 | 0 | 1 74E-29 |
| ITFG2 | 0 229835716 | 0 034965027 | 8 64E-30 | 0 296155882 | 0 | 1 85E-29 |
| NRL | 0 027050694 | 0 034957278 | 8 89E-30 | 0 480273052 | 0 | 1 91E-29 |
| SEC14L1 | 0 347337762 | 0 034941514 | 9 43E-30 | -0 16691763 | 0 | 2 02E-29 |
| MIDN | 0 155527213 | 0 034937185 | 9 58E-30 | -0 32527443 | 0 | 2 05E-29 |
| ITPKB | 0 211551274 | 0 034911311 | 1 05E-29 | -0 2540378 | 0 | 2 26E-29 |
| FBLN7 | 0 095020083 | 0 034898025 | 1 11E-29 | 0 377763008 | 0 | 2 37E-29 |
| CNIH2 | 0 007766842 | 0 034894945 | 1 12E-29 | 0 493496545 | 0 | 2 40E-29 |
| HSPB6 | 0 027526603 | 0 034883816 | 1 17E-29 | -0 461908 | 0 | 2 50E-29 |
| CLIC2 | 0 07364223 | 0 03487033 | 1 23E-29 | -0 41778119 | 0 | 2 63E-29 |
| FAM89A | 0 080447736 | 0 0348507 | 1 32E-29 | -0 40090261 | 0 | 2 82E-29 |
| EGR4 | 0 002170147 | 0 034847032 | 1 34E-29 | -0 49304015 | 0 | 2 86E-29 |
| KRT8 | 0 055395861 | 0 034845873 | 1 34E-29 | -0 42950269 | 0 | 2 87E-29 |
| PLD5 | 0 027383831 | 0 034827237 | 1 44E-29 | 0 482036247 | 0 | 3 08E-29 |
| INO80D | 0 345015324 | 0 03482063 | 1 47E-29 | -0 17617957 | 0 | 3 15E-29 |
| CHST12 | 0 141373665 | 0 034803343 | 1 57E-29 | -0 33759543 | 0 | 3 36E-29 |
| USP20 | 0 12263235 | 0 03478922 | 1 65E-29 | -0 36092304 | 0 | 3 54E-29 |
| ANKRD13A | 0 181502351 | 0 034772456 | 1 76E-29 | 0 340015306 | 0 | 3 76E-29 |
| ZNF274 | 0 143524776 | 0 034764967 | 1 81E-29 | -0 33800448 | 0 | 3 87E-29 |
| TM7SF2 | 0 027345758 | 0 034755579 | 1 87E-29 | -0 45919275 | 0 | 4 00E-29 |
| ELMOD1 | 0 021254117 | 0 034751345 | 1 90E-29 | -0 47008541 | 0 | 4 06E-29 |
| CXCR5 | 0 004140412 | 0 034736919 | 2 01E-29 | 0 497097981 | 0 | 4 28E-29 |
| COQ7 | 0 081028345 | 0 034708233 | 2 23E-29 | -0 40193927 | 0 | 4 76E-29 |
| CXCR4 | 0 046715273 | 0 034691414 | 2 37E-29 | 0 465226812 | 0 | 5 06E-29 |
| ZNF813 | 0 035150673 | 0 034688951 | 2 39E-29 | 0 478078704 | 0 | 5 11E-29 |
| CTH | 0 030324951 | 0 034629419 | 2 98E-29 | -0 45742663 | 0 | 6 35E-29 |
| GUCY1B1 | 0 123127296 | 0 034626977 | 3 01E-29 | -0 36069049 | 0 | 6 41E-29 |
| GAN | 0 181959224 | 0 034625412 | 3 02E-29 | -0 29810251 | 0 | 6 44E-29 |

| | | | | | | |
|---|---|---|---|---|---|---|
| EXOC2 | 0 432363747 | 0 034612557 | 3 17E-29 | -0 10923454 | 3 17E-276 | 6 75E-29 |
| FLRT3 | 0 075517314 | 0 034606693 | 3 24E-29 | 0 439768329 | 0 | 6 89E-29 |
| EIF2S2 | 0 337191373 | 0 03460368 | 3 27E-29 | 0 239098472 | 0 | 6 97E-29 |
| LINC00862 | 0 072747521 | 0 03460329 | 3 28E-29 | -0 4087035 | 0 | 6 98E-29 |
| HBP1 | 0 339742247 | 0 034599617 | 3 32E-29 | -0 16825338 | 0 | 7 07E-29 |
| AGO4 | 0 337334146 | 0 034575625 | 3 63E-29 | 0 256999462 | 0 | 7 72E-29 |
| CDKL5 | 0 323884944 | 0 034548604 | 4 01E-29 | 0 219799006 | 0 | 8 52E-29 |
| TIGD1 | 0 047562392 | 0 034537771 | 4 17E-29 | 0 465577415 | 0 | 8 86E-29 |
| PLAC8 | 0 012497382 | 0 034534285 | 4 22E-29 | 0 49122906 | 0 | 8 97E-29 |
| RNF138 | 0 211760675 | 0 034527899 | 4 32E-29 | 0 31658499 | 0 | 9 18E-29 |
| DOC2A | 0 016057185 | 0 034492247 | 4 92E-29 | 0 488790096 | 0 | 1 05E-28 |
| MRPS27 | 0 314376273 | 0 034430664 | 6 17E-29 | 0 245019578 | 0 | 1 31E-28 |
| TSPEAR | 0 011793036 | 0 034426558 | 6 26E-29 | 0 491090364 | 0 | 1 33E-28 |
| CYBRD1 | 0 401324932 | 0 034413128 | 6 57E-29 | -0 14496869 | 0 | 1 39E-28 |
| CNBD2 | 0 03642611 | 0 034395248 | 7 02E-29 | -0 45180584 | 0 | 1 49E-28 |
| CARNMT1 | 0 207325198 | 0 0343785 | 7 46E-29 | 0 30998665 | 0 | 1 58E-28 |
| PCBD2 | 0 224753003 | 0 03436747 | 7 77E-29 | 0 295469457 | 0 | 1 64E-28 |
| MOB1B | 0 343930251 | 0 034350834 | 8 25E-29 | -0 18071505 | 0 | 1 75E-28 |
| ACVR2A | 0 308322705 | 0 034337101 | 8 67E-29 | 0 240453051 | 0 | 1 84E-28 |
| HDAC10 | 0 062039558 | 0 034336751 | 8 69E-29 | 0 445500967 | 0 | 1 84E-28 |
| DAGLB | 0 128571701 | 0 034330108 | 8 90E-29 | 0 402779323 | 0 | 1 88E-28 |
| ADAMTS9-AS2 | 0 447678514 | 0 034319772 | 9 24E-29 | -0 06872653 | 3 48E-110 | 1 95E-28 |
| EIF3E | 0 51077459 | 0 034287026 | 1 04E-28 | 0 140886801 | 0 | 2 20E-28 |
| ZNF671 | 0 030772306 | 0 03427391 | 1 09E-28 | 0 480916737 | 0 | 2 31E-28 |
| LIN54 | 0 207410862 | 0 034232815 | 1 27E-28 | 0 285229932 | 0 | 2 68E-28 |
| BANK1 | 0 040233386 | 0 034208155 | 1 39E-28 | 0 458839881 | 0 | 2 93E-28 |
| TBC1D14 | 0 202194894 | 0 034205647 | 1 40E-28 | -0 28448184 | 0 | 2 95E-28 |
| ZNF836 | 0 065513697 | 0 034204685 | 1 40E-28 | -0 42170515 | 0 | 2 96E-28 |
| DRD4 | 0 018265405 | 0 034197679 | 1 44E-28 | 0 48790274 | 0 | 3 04E-28 |
| ZSCAN31 | 0 079048562 | 0 034186785 | 1 50E-28 | -0 40365054 | 0 | 3 16E-28 |
| TMEM100 | 0 055215016 | 0 034178712 | 1 54E-28 | -0 43405169 | 0 | 3 25E-28 |
| SCOC-AS1 | 0 034931755 | 0 034162865 | 1 63E-28 | 0 474720905 | 0 | 3 44E-28 |
| ECD | 0 229845234 | 0 034128014 | 1 85E-28 | -0 26518333 | 0 | 3 90E-28 |
| FCHO2 | 0 483657269 | 0 034121337 | 1 90E-28 | -0 10739456 | 5 12E-267 | 4 00E-28 |
| PEX14 | 0 278340409 | 0 034112577 | 1 96E-28 | 0 271741183 | 0 | 4 12E-28 |
| CORO7 | 0 117759038 | 0 034109734 | 1 98E-28 | -0 36217306 | 0 | 4 16E-28 |
| DNAH8 | 0 013068474 | 0 034101999 | 2 04E-28 | 0 49298511 | 0 | 4 28E-28 |
| CRYBG3 | 0 455902229 | 0 034075463 | 2 24E-28 | -0 09912958 | 1 27E-227 | 4 71E-28 |
| NTN1 | 0 185157336 | 0 034063125 | 2 34E-28 | -0 28232904 | 0 | 4 93E-28 |
| HS3ST2 | 0 059003255 | 0 034056747 | 2 40E-28 | -0 41942959 | 0 | 5 04E-28 |
| LGALS2 | 0 012592564 | 0 034045383 | 2 50E-28 | 0 489360157 | 0 | 5 25E-28 |
| POLD1 | 0 061915821 | 0 034031551 | 2 63E-28 | 0 414403956 | 0 | 5 51E-28 |
| SPC24 | 0 04578249 | 0 034030693 | 2 63E-28 | 0 425324796 | 0 | 5 53E-28 |
| TRIM24 | 0 291761055 | 0 034029351 | 2 65E-28 | 0 247144445 | 0 | 5 55E-28 |

| | | | | | | |
|---|---|---|---|---|---|---|
| AGPAT4 | 0 311492262 | 0 034027763 | 2 66E-28 | 0 271266469 | 0 | 5 59E-28 |
| IL17RA | 0 151139327 | 0 034023575 | 2 70E-28 | 0 363421598 | 0 | 5 67E-28 |
| CHAF1A | 0 091764863 | 0 034010313 | 2 83E-28 | 0 359505053 | 0 | 5 95E-28 |
| AUH | 0 48754069 | 0 033968669 | 3 29E-28 | -0 09552176 | 1 98E-211 | 6 91E-28 |
| SPEN | 0 406093545 | 0 033955555 | 3 45E-28 | 0 198341257 | 0 | 7 24E-28 |
| LIME1 | 0 066122861 | 0 033944171 | 3 60E-28 | 0 441335511 | 0 | 7 54E-28 |
| EXOC1 | 0 412851459 | 0 033928574 | 3 80E-28 | 0 206936588 | 0 | 7 97E-28 |
| SEMA4A | 0 048019265 | 0 033927 | 3 83E-28 | -0 43566776 | 0 | 8 02E-28 |
| MCM7 | 0 12694409 | 0 033915939 | 3 98E-28 | 0 346286183 | 0 | 8 34E-28 |
| PDE4C | 0 024642592 | 0 0339156 | 3 99E-28 | -0 4626446 | 0 | 8 35E-28 |
| NPIPB4 | 0 170565952 | 0 033903552 | 4 16E-28 | 0 354542353 | 0 | 8 71E-28 |
| SPC25 | 0 082094382 | 0 033902679 | 4 18E-28 | 0 371451176 | 0 | 8 74E-28 |
| ZCCHC10 | 0 187679656 | 0 033864995 | 4 78E-28 | 0 344142016 | 0 | 1 00E-27 |
| ZWILCH | 0 107355657 | 0 033864899 | 4 78E-28 | 0 361070796 | 0 | 1 00E-27 |
| ZBTB11 | 0 184472026 | 0 033862462 | 4 83E-28 | 0 351918333 | 0 | 1 01E-27 |
| KIF2A | 0 413022787 | 0 033857413 | 4 91E-28 | 0 171693509 | 0 | 1 03E-27 |
| DNAJA4 | 0 103177172 | 0 033832009 | 5 38E-28 | -0 38159557 | 0 | 1 13E-27 |
| OSBPL7 | 0 058460718 | 0 033829484 | 5 43E-28 | 0 450120349 | 0 | 1 14E-27 |
| TAC1 | 0 044621271 | 0 033823101 | 5 56E-28 | -0 44229382 | 0 | 1 16E-27 |
| PCDH17 | 0 048913975 | 0 033813861 | 5 74E-28 | 0 445820666 | 0 | 1 20E-27 |
| GAA | 0 147636634 | 0 033797309 | 6 10E-28 | -0 33265229 | 0 | 1 27E-27 |
| WIPF1 | 0 519379033 | 0 033776359 | 6 57E-28 | 0 129840017 | 0 | 1 37E-27 |
| DEK | 0 35938779 | 0 03374907 | 7 25E-28 | 0 208844846 | 0 | 1 51E-27 |
| GGT5 | 0 128743028 | 0 033729947 | 7 76E-28 | -0 33106757 | 0 | 1 62E-27 |
| PLIN2 | 0 099436523 | 0 033722534 | 7 97E-28 | -0 38666204 | 0 | 1 66E-27 |
| GCLC | 0 137242771 | 0 033720092 | 8 04E-28 | -0 3369949 | 0 | 1 68E-27 |
| ZNF670 | 0 09808494 | 0 033712801 | 8 25E-28 | 0 418662242 | 0 | 1 72E-27 |
| ZBED4 | 0 178799185 | 0 033708051 | 8 39E-28 | 0 327564094 | 0 | 1 75E-27 |
| SLC35F2 | 0 057994327 | 0 033705244 | 8 47E-28 | 0 461496535 | 0 | 1 77E-27 |
| ARMCX6 | 0 034808018 | 0 033701294 | 8 59E-28 | 0 47582854 | 0 | 1 79E-27 |
| TMEM209 | 0 183891416 | 0 033699611 | 8 65E-28 | 0 33644012 | 0 | 1 80E-27 |
| UBE2R2 | 0 506405741 | 0 033689394 | 8 97E-28 | 0 152498593 | 0 | 1 87E-27 |
| SIAH2 | 0 099531705 | 0 033683505 | 9 16E-28 | 0 414892637 | 0 | 1 91E-27 |
| NRGN | 0 017047077 | 0 0336766 | 9 39E-28 | 0 481633599 | 0 | 1 95E-27 |
| USP2-AS1 | 0 031400506 | 0 033675356 | 9 43E-28 | 0 481412791 | 0 | 1 96E-27 |
| SKA2 | 0 306038339 | 0 033674153 | 9 47E-28 | 0 187432382 | 0 | 1 97E-27 |
| MND1 | 0 072871257 | 0 033673072 | 9 51E-28 | 0 385901821 | 0 | 1 98E-27 |
| ELOVL6 | 0 193885515 | 0 033646808 | 1 04E-27 | 0 305705922 | 0 | 2 17E-27 |
| TCEANC2 | 0 070472673 | 0 033627016 | 1 12E-27 | 0 43743262 | 0 | 2 33E-27 |
| WDR44 | 0 298081133 | 0 033593113 | 1 26E-27 | -0 19848581 | 0 | 2 63E-27 |
| EME2 | 0 050760503 | 0 033573338 | 1 36E-27 | -0 43764412 | 0 | 2 82E-27 |
| EIF4B | 0 420808665 | 0 033565269 | 1 40E-27 | 0 178653062 | 0 | 2 90E-27 |
| TNFSF11 | 0 004007158 | 0 033565057 | 1 40E-27 | -0 49078728 | 0 | 2 90E-27 |
| CREBRF | 0 342740477 | 0 033552767 | 1 46E-27 | -0 16327305 | 0 | 3 03E-27 |

| | | | | | | |
|---|---|---|---|---|---|---|
| MAP3K21 | 0 0343797 | 0 033534736 | 1 56E-27 | 0 466622529 | 0 | 3 23E-27 |
| RSAD2 | 0 05571948 | 0 033531211 | 1 58E-27 | 0 445621766 | 0 | 3 27E-27 |
| SMIM14 | 0 336782091 | 0 033506451 | 1 72E-27 | -0 17391641 | 0 | 3 57E-27 |
| RFFL | 0 224286612 | 0 033497611 | 1 78E-27 | -0 27605251 | 0 | 3 68E-27 |
| TRIM41 | 0 099074832 | 0 033483552 | 1 87E-27 | 0 421366845 | 0 | 3 87E-27 |
| SLC1A2 | 0 081351964 | 0 033474468 | 1 93E-27 | -0 3905009 | 0 | 3 99E-27 |
| KIAA0232 | 0 493698959 | 0 033454516 | 2 07E-27 | -0 07565209 | 3 71E-133 | 4 28E-27 |
| SLF1 | 0 257971484 | 0 03345163 | 2 09E-27 | 0 243722378 | 0 | 4 33E-27 |
| KDM6B | 0 089642306 | 0 033450956 | 2 10E-27 | -0 38577741 | 0 | 4 34E-27 |
| AP5Z1 | 0 147113133 | 0 033439261 | 2 18E-27 | -0 33683528 | 0 | 4 52E-27 |
| LAG3 | 0 022253526 | 0 03335497 | 2 94E-27 | 0 479258607 | 0 | 6 09E-27 |
| RB1 | 0 548761684 | 0 033340684 | 3 10E-27 | 0 130835911 | 0 | 6 40E-27 |
| NUFIP2 | 0 424092441 | 0 033339257 | 3 11E-27 | -0 12043429 | 0 | 6 43E-27 |
| C11orf65 | 0 111353296 | 0 033335364 | 3 16E-27 | -0 37735555 | 0 | 6 52E-27 |
| CHORDC1 | 0 202937313 | 0 033316108 | 3 38E-27 | -0 27970842 | 0 | 6 98E-27 |
| APLF | 0 241933335 | 0 033271333 | 3 96E-27 | 0 30294391 | 0 | 8 17E-27 |
| PAIP2B | 0 036235747 | 0 033254072 | 4 21E-27 | -0 45083789 | 0 | 8 68E-27 |
| MYBL2 | 0 044821153 | 0 03324062 | 4 41E-27 | 0 419390359 | 0 | 9 10E-27 |
| PPIG | 0 503150521 | 0 033234289 | 4 51E-27 | 0 141967181 | 0 | 9 30E-27 |
| SLC29A3 | 0 02544212 | 0 033221895 | 4 71E-27 | -0 46461699 | 0 | 9 72E-27 |
| KANK3 | 0 026527193 | 0 033194626 | 5 19E-27 | -0 46405744 | 0 | 1 07E-26 |
| LINC00907 | 0 063115113 | 0 033176217 | 5 53E-27 | -0 42083931 | 0 | 1 14E-26 |
| SUB1 | 0 408977556 | 0 033153122 | 6 00E-27 | -0 15360688 | 0 | 1 24E-26 |
| LIMK1 | 0 15059679 | 0 033148096 | 6 11E-27 | 0 360115475 | 0 | 1 26E-26 |
| LINC01630 | 0 035331519 | 0 033145447 | 6 17E-27 | 0 476258003 | 0 | 1 27E-26 |
| LMTK2 | 0 277512326 | 0 033130617 | 6 50E-27 | 0 279912605 | 0 | 1 34E-26 |
| LRRK2 | 0 265766881 | 0 033124842 | 6 63E-27 | -0 23010653 | 0 | 1 36E-26 |
| RTN4IP1 | 0 080209781 | 0 033110399 | 6 97E-27 | 0 442291083 | 0 | 1 44E-26 |
| CD248 | 0 082103901 | 0 033098571 | 7 27E-27 | -0 39540593 | 0 | 1 50E-26 |
| LIMS2 | 0 034027527 | 0 0330902 | 7 49E-27 | -0 45511512 | 0 | 1 54E-26 |
| PXN-AS1 | 0 090632198 | 0 033062415 | 8 25E-27 | -0 40063052 | 0 | 1 70E-26 |
| CCNF | 0 069606518 | 0 033036754 | 9 03E-27 | 0 404868383 | 0 | 1 85E-26 |
| UPF3B | 0 190382822 | 0 033030273 | 9 24E-27 | 0 339675008 | 0 | 1 90E-26 |
| ANO6 | 0 545239954 | 0 033027997 | 9 31E-27 | 0 110693897 | 1 21E-283 | 1 91E-26 |
| ZNF763 | 0 037063829 | 0 033005601 | 1 01E-26 | 0 473451 | 0 | 2 07E-26 |
| ZMAT3 | 0 334802307 | 0 033000414 | 1 03E-26 | -0 16634394 | 0 | 2 10E-26 |
| AMZ2 | 0 188859911 | 0 033000378 | 1 03E-26 | -0 29011793 | 0 | 2 10E-26 |
| UNC79 | 0 060535684 | 0 032977091 | 1 11E-26 | -0 42280085 | 0 | 2 28E-26 |
| PTCSC2 | 0 013658602 | 0 032973379 | 1 13E-26 | 0 490851158 | 0 | 2 31E-26 |
| OFD1 | 0 355095087 | 0 032970882 | 1 14E-26 | -0 17136898 | 0 | 2 33E-26 |
| CHST11 | 0 300632008 | 0 032960769 | 1 18E-26 | 0 251400547 | 0 | 2 42E-26 |
| CXXC5 | 0 112904761 | 0 032948141 | 1 23E-26 | -0 35138912 | 0 | 2 52E-26 |
| LRRC39 | 0 050770022 | 0 032947092 | 1 24E-26 | -0 4373595 | 0 | 2 53E-26 |
| MSLN | 0 001770383 | 0 032934336 | 1 29E-26 | 0 497795903 | 0 | 2 65E-26 |

| | | | | | | |
|---|---|---|---|---|---|---|
| MR1 | 0 10912604 | 0 032913979 | 1 39E-26 | -0 37424694 | 0 | 2 84E-26 |
| NKRF | 0 057470827 | 0 032913349 | 1 39E-26 | -0 4299298 | 0 | 2 85E-26 |
| RBMS3-AS3 | 0 209780891 | 0 032898813 | 1 46E-26 | -0 26912321 | 0 | 2 99E-26 |
| KCNH1 | 0 063828977 | 0 032893239 | 1 49E-26 | 0 445979582 | 0 | 3 05E-26 |
| PAPSS2 | 0 168747977 | 0 032885542 | 1 53E-26 | -0 33000556 | 0 | 3 14E-26 |
| ATP11B | 0 489739392 | 0 032885207 | 1 53E-26 | 0 146459684 | 0 | 3 14E-26 |
| PCSK7 | 0 25167996 | 0 03288336 | 1 54E-26 | -0 22508582 | 0 | 3 16E-26 |
| EPM2A | 0 125820944 | 0 032869689 | 1 62E-26 | -0 35803864 | 0 | 3 31E-26 |
| BLOC1S6 | 0 243246845 | 0 03284789 | 1 75E-26 | 0 317705224 | 0 | 3 57E-26 |
| SH3BP4 | 0 194418534 | 0 032838759 | 1 80E-26 | -0 29377662 | 0 | 3 68E-26 |
| CDKL1 | 0 245673983 | 0 032816662 | 1 95E-26 | 0 289871217 | 0 | 3 98E-26 |
| KCNJ3 | 0 020035788 | 0 032815154 | 1 96E-26 | 0 486647775 | 0 | 4 00E-26 |
| KPNB1 | 0 497249243 | 0 032809039 | 2 00E-26 | 0 131028782 | 0 | 4 08E-26 |
| PUS7 | 0 240496088 | 0 032808057 | 2 01E-26 | 0 277395845 | 0 | 4 09E-26 |
| USP18 | 0 053263787 | 0 032802676 | 2 04E-26 | 0 450117112 | 0 | 4 17E-26 |
| SREK1 | 0 520835316 | 0 032775355 | 2 25E-26 | 0 145434226 | 0 | 4 58E-26 |
| LRRC2 | 0 031809789 | 0 032774987 | 2 25E-26 | -0 45649111 | 0 | 4 59E-26 |
| HEYL | 0 089785079 | 0 032770639 | 2 29E-26 | -0 39625363 | 0 | 4 66E-26 |
| PLPPR4 | 0 059307837 | 0 032764929 | 2 33E-26 | -0 4236053 | 0 | 4 75E-26 |
| MAPKAPK2 | 0 235394339 | 0 032755703 | 2 41E-26 | -0 25034975 | 0 | 4 90E-26 |
| CPEB2-DT | 0 034979346 | 0 032755521 | 2 41E-26 | -0 45000966 | 0 | 4 91E-26 |
| EHMT2-AS1 | 0 023414746 | 0 032740865 | 2 54E-26 | 0 480309634 | 0 | 5 16E-26 |
| TMEM165 | 0 49668767 | 0 032738 | 2 56E-26 | 0 145145306 | 0 | 5 21E-26 |
| IMPDH1 | 0 080638099 | 0 032724639 | 2 68E-26 | 0 430818815 | 0 | 5 46E-26 |
| GDF15 | 0 078467952 | 0 032712866 | 2 79E-26 | -0 40358688 | 0 | 5 68E-26 |
| TNNT3 | 0 039776513 | 0 032697267 | 2 95E-26 | -0 4510613 | 0 | 6 00E-26 |
| SMC1A | 0 269812111 | 0 032688207 | 3 04E-26 | 0 260306179 | 0 | 6 19E-26 |
| AOAH | 0 07380404 | 0 032663574 | 3 31E-26 | -0 40754175 | 0 | 6 74E-26 |
| PIGB | 0 156507586 | 0 032618588 | 3 87E-26 | -0 33156342 | 0 | 7 87E-26 |
| WARS2 | 0 23565133 | 0 032591862 | 4 25E-26 | -0 25718103 | 0 | 8 63E-26 |
| LINC02196 | 0 038415412 | 0 032587091 | 4 32E-26 | 0 468979501 | 0 | 8 77E-26 |
| ANPEP | 0 088252651 | 0 032583741 | 4 37E-26 | -0 39356366 | 0 | 8 87E-26 |
| DAP3 | 0 361215282 | 0 032576053 | 4 49E-26 | 0 212184326 | 0 | 9 11E-26 |
| RELCH | 0 391416497 | 0 032555895 | 4 81E-26 | -0 15116326 | 0 | 9 76E-26 |
| SAV1 | 0 192048505 | 0 032503812 | 5 76E-26 | -0 28858967 | 0 | 1 17E-25 |
| WDFY4 | 0 027973958 | 0 032495494 | 5 93E-26 | -0 46274548 | 0 | 1 20E-25 |
| KBTBD12 | 0 017494432 | 0 032479661 | 6 26E-26 | 0 48860514 | 0 | 1 27E-25 |
| HSPBAP1 | 0 14755097 | 0 032469119 | 6 49E-26 | -0 33476687 | 0 | 1 31E-25 |
| LARP7 | 0 219241971 | 0 032456528 | 6 78E-26 | 0 335006222 | 0 | 1 37E-25 |
| CLK1 | 0 439216843 | 0 032456274 | 6 78E-26 | -0 10896639 | 7 12E-275 | 1 37E-25 |
| PDK4 | 0 224048657 | 0 032453793 | 6 84E-26 | -0 26918336 | 0 | 1 38E-25 |
| TCF19 | 0 048590356 | 0 032442867 | 7 10E-26 | 0 426051205 | 0 | 1 44E-25 |
| PTER | 0 116673964 | 0 032436612 | 7 26E-26 | 0 411523005 | 0 | 1 47E-25 |
| CACNA1C-AS2 | 0 005387295 | 0 032422477 | 7 62E-26 | 0 49678266 | 0 | 1 54E-25 |

| | | | | | | |
|---|---|---|---|---|---|---|
| DAAM1 | 0 439435762 | 0 032413324 | 7 86E-26 | -0 08554996 | 7 54E-170 | 1 59E-25 |
| PKN2-AS1 | 0 095743466 | 0 032407901 | 8 01E-26 | -0 39193086 | 0 | 1 62E-25 |
| TCIM | 0 081551846 | 0 032389051 | 8 55E-26 | -0 40881074 | 0 | 1 73E-25 |
| TMC4 | 0 036721174 | 0 032383487 | 8 71E-26 | 0 464056223 | 0 | 1 76E-25 |
| LY6H | 0 012659192 | 0 032334047 | 1 03E-25 | 0 49062112 | 0 | 2 09E-25 |
| TMEM86B | 0 032637871 | 0 03232288 | 1 07E-25 | -0 45708284 | 0 | 2 17E-25 |
| ARL5A | 0 302726009 | 0 032316681 | 1 10E-25 | -0 21924928 | 0 | 2 21E-25 |
| DNAJC13 | 0 421417829 | 0 032314007 | 1 11E-25 | 0 196015308 | 0 | 2 23E-25 |
| KDM4C | 0 534284518 | 0 032303734 | 1 15E-25 | -0 06344445 | 3 75E-94 | 2 31E-25 |
| PDE6D | 0 168024595 | 0 03226773 | 1 30E-25 | 0 340955509 | 0 | 2 61E-25 |
| TOM1L2 | 0 413698578 | 0 032267401 | 1 30E-25 | -0 10343814 | 9 31E-248 | 2 62E-25 |
| ESCO1 | 0 316974739 | 0 032253271 | 1 36E-25 | -0 19672255 | 0 | 2 75E-25 |
| NRN1 | 0 091612572 | 0 03225241 | 1 37E-25 | -0 3909553 | 0 | 2 75E-25 |
| CYP1B1 | 0 260331994 | 0 032239874 | 1 43E-25 | -0 21941298 | 0 | 2 87E-25 |
| BEND6 | 0 089908816 | 0 032237598 | 1 44E-25 | -0 39828428 | 0 | 2 89E-25 |
| UBAP2 | 0 388085131 | 0 032218446 | 1 53E-25 | 0 20386889 | 0 | 3 09E-25 |
| GPBP1L1 | 0 426110297 | 0 032209796 | 1 58E-25 | -0 13191521 | 0 | 3 18E-25 |
| PGF | 0 099940987 | 0 032205229 | 1 61E-25 | -0 36929631 | 0 | 3 23E-25 |
| TRIM5 | 0 275684834 | 0 032197947 | 1 65E-25 | -0 2204564 | 0 | 3 31E-25 |
| NPIPB5 | 0 205050351 | 0 032191899 | 1 68E-25 | 0 322696173 | 0 | 3 38E-25 |
| LEPROTL1 | 0 154404066 | 0 032173616 | 1 79E-25 | -0 32799425 | 0 | 3 60E-25 |
| SLC35F3 | 0 052159677 | 0 032172402 | 1 80E-25 | -0 43153493 | 0 | 3 61E-25 |
| PTX3 | 0 08531153 | 0 032168286 | 1 82E-25 | -0 39449598 | 0 | 3 66E-25 |
| MYH3 | 0 028630713 | 0 032145376 | 1 97E-25 | -0 46208699 | 0 | 3 96E-25 |
| METTL7A | 0 142868021 | 0 032141237 | 2 00E-25 | -0 34193778 | 0 | 4 01E-25 |
| USP4 | 0 294150121 | 0 032131851 | 2 06E-25 | 0 286503853 | 0 | 4 14E-25 |
| ACVRL1 | 0 051065085 | 0 032112945 | 2 20E-25 | -0 4406513 | 0 | 4 42E-25 |
| LINC01284 | 0 012640155 | 0 032090326 | 2 38E-25 | -0 4791804 | 0 | 4 77E-25 |
| DTNBP1 | 0 278673545 | 0 032089482 | 2 38E-25 | -0 22778549 | 0 | 4 78E-25 |
| PART1 | 0 035369591 | 0 032087803 | 2 40E-25 | 0 481793266 | 0 | 4 81E-25 |
| SLC28A1 | 0 010726999 | 0 032070372 | 2 54E-25 | -0 48259038 | 0 | 5 10E-25 |
| SLC6A1 | 0 039119758 | 0 032060524 | 2 63E-25 | -0 44069009 | 0 | 5 27E-25 |
| WSB1 | 0 761835868 | 0 032047112 | 2 75E-25 | 0 014130779 | 4 64E-06 | 5 51E-25 |
| ZNF169 | 0 130846548 | 0 032042404 | 2 80E-25 | 0 3823622 | 0 | 5 60E-25 |
| STK17B | 0 26303516 | 0 032040028 | 2 82E-25 | 0 236660712 | 0 | 5 64E-25 |
| ACRBP | 0 014610421 | 0 032023069 | 2 99E-25 | -0 47952161 | 0 | 5 98E-25 |
| LINC00662 | 0 222487674 | 0 032013606 | 3 08E-25 | 0 332816857 | 0 | 6 17E-25 |
| SUSD3 | 0 006262969 | 0 03200807 | 3 14E-25 | 0 496452576 | 0 | 6 29E-25 |
| LINC01090 | 0 157735432 | 0 032005989 | 3 16E-25 | 0 356587312 | 0 | 6 33E-25 |
| RAB30 | 0 124145743 | 0 032002733 | 3 20E-25 | -0 34419284 | 0 | 6 40E-25 |
| NUDT14 | 0 026432012 | 0 031981482 | 3 44E-25 | -0 46457505 | 0 | 6 88E-25 |
| ADGB | 0 006158268 | 0 031945547 | 3 88E-25 | 0 496735304 | 0 | 7 77E-25 |
| PRUNE2 | 0 203774914 | 0 031944185 | 3 90E-25 | -0 29142434 | 0 | 7 80E-25 |
| ZFAT | 0 155565285 | 0 031939599 | 3 96E-25 | -0 33230906 | 0 | 7 92E-25 |

| | | | | | | |
|---|---|---|---|---|---|---|
| SUSD1 | 0 21315033 | 0 031937178 | 4 00E-25 | -0 27993217 | 0 | 7 98E-25 |
| CPT1A | 0 235127829 | 0 031904781 | 4 46E-25 | 0 273339309 | 0 | 8 91E-25 |
| TBK1 | 0 385914984 | 0 031894807 | 4 61E-25 | 0 205842778 | 0 | 9 21E-25 |
| FAM169A | 0 087119986 | 0 031889101 | 4 70E-25 | 0 418625435 | 0 | 9 39E-25 |
| DTL | 0 085844549 | 0 031850665 | 5 36E-25 | 0 346786398 | 0 | 1 07E-24 |
| KRT86 | 0 005815614 | 0 031786678 | 6 65E-25 | 0 49523409 | 0 | 1 32E-24 |
| BTNL9 | 0 019198188 | 0 031786139 | 6 66E-25 | 0 486229071 | 0 | 1 33E-24 |
| ZNF281 | 0 088766633 | 0 031775118 | 6 91E-25 | -0 39031247 | 0 | 1 38E-24 |
| JAM2 | 0 176886029 | 0 031763072 | 7 20E-25 | -0 30396203 | 0 | 1 43E-24 |
| DIO2 | 0 096885648 | 0 031758152 | 7 32E-25 | -0 3909114 | 0 | 1 46E-24 |
| CNOT6L | 0 333298433 | 0 031752585 | 7 46E-25 | 0 233451342 | 0 | 1 48E-24 |
| PDCD6IP | 0 513782338 | 0 031748029 | 7 57E-25 | -0 07203482 | 7 00E-121 | 1 51E-24 |
| CASS4 | 0 019616988 | 0 031725296 | 8 18E-25 | -0 47025432 | 0 | 1 63E-24 |
| ITPR2 | 0 508613961 | 0 031723905 | 8 22E-25 | -0 06316798 | 2 38E-93 | 1 63E-24 |
| MMP14 | 0 322419143 | 0 03170956 | 8 62E-25 | -0 20420348 | 0 | 1 71E-24 |
| TANK-AS1 | 0 006596105 | 0 031705636 | 8 74E-25 | 0 497052432 | 0 | 1 74E-24 |
| RPS6KA3 | 0 468247321 | 0 031695016 | 9 05E-25 | -0 10344744 | 8 41E-248 | 1 80E-24 |
| MPP7 | 0 076992633 | 0 031687274 | 9 29E-25 | 0 433719419 | 0 | 1 85E-24 |
| NPHS1 | 0 007100569 | 0 031685704 | 9 34E-25 | 0 49449354 | 0 | 1 85E-24 |
| ALKBH8 | 0 08872856 | 0 031683383 | 9 42E-25 | -0 39387873 | 0 | 1 87E-24 |
| TEP1 | 0 115141535 | 0 031667939 | 9 92E-25 | -0 36191015 | 0 | 1 97E-24 |
| USP28 | 0 147684225 | 0 03163401 | 1 11E-24 | 0 376390122 | 0 | 2 20E-24 |
| LPCAT4 | 0 066427443 | 0 031605048 | 1 22E-24 | 0 437830447 | 0 | 2 43E-24 |
| CFAP54 | 0 153823457 | 0 03157739 | 1 34E-24 | -0 32461014 | 0 | 2 66E-24 |
| SUMF2 | 0 251480078 | 0 031565689 | 1 40E-24 | -0 24830427 | 0 | 2 77E-24 |
| SLC4A10 | 0 028802041 | 0 031550732 | 1 47E-24 | 0 481418111 | 0 | 2 91E-24 |
| BHLHE40-AS1 | 0 042708115 | 0 031545194 | 1 50E-24 | 0 46388681 | 0 | 2 97E-24 |
| CCDC170 | 0 067084198 | 0 031532166 | 1 56E-24 | -0 40904057 | 0 | 3 10E-24 |
| IRAG1 | 0 007405151 | 0 03151056 | 1 68E-24 | -0 48565693 | 0 | 3 33E-24 |
| CEACAM21 | 0 005035122 | 0 031490244 | 1 80E-24 | 0 495281391 | 0 | 3 56E-24 |
| SLC8A1 | 0 318459576 | 0 031486719 | 1 82E-24 | 0 169143138 | 0 | 3 60E-24 |
| STARD3 | 0 133997068 | 0 031481334 | 1 85E-24 | 0 400011698 | 0 | 3 67E-24 |
| DIAPH3 | 0 1401363 | 0 031475107 | 1 89E-24 | 0 314838011 | 0 | 3 74E-24 |
| CTSV | 0 0231292 | 0 031464982 | 1 96E-24 | 0 482290119 | 0 | 3 87E-24 |
| ERC2 | 0 113466334 | 0 031457459 | 2 01E-24 | -0 3627179 | 0 | 3 97E-24 |
| LNPEP | 0 390093469 | 0 031423699 | 2 25E-24 | 0 199484738 | 0 | 4 44E-24 |
| SLC11A2 | 0 290085854 | 0 031411563 | 2 34E-24 | -0 22071275 | 0 | 4 62E-24 |
| CHRM3-AS2 | 0 008918543 | 0 031397152 | 2 45E-24 | -0 48567425 | 0 | 4 85E-24 |
| SPTY2D1 | 0 124164779 | 0 031392181 | 2 50E-24 | -0 35876955 | 0 | 4 93E-24 |
| FGD6 | 0 282166721 | 0 031374152 | 2 65E-24 | -0 2334768 | 0 | 5 23E-24 |
| P2RY10 | 0 005358741 | 0 03134635 | 2 91E-24 | 0 495840687 | 0 | 5 73E-24 |
| ATP13A1 | 0 080837981 | 0 031324487 | 3 13E-24 | -0 40599745 | 0 | 6 17E-24 |
| SLC29A2 | 0 013449201 | 0 031288937 | 3 52E-24 | 0 492394816 | 0 | 6 94E-24 |
| DPYSL2 | 0 363166511 | 0 031283405 | 3 58E-24 | -0 13507281 | 0 | 7 06E-24 |

| | | | | | | |
|---|---|---|---|---|---|---|
| ZBTB8OS | 0 336230036 | 0 031275536 | 3 68E-24 | 0 23238357 | 0 | 7 25E-24 |
| PCK2 | 0 029039995 | 0 031271737 | 3 73E-24 | 0 477544752 | 0 | 7 34E-24 |
| PDK3 | 0 197768936 | 0 031261109 | 3 86E-24 | -0 28630178 | 0 | 7 60E-24 |
| CAPN6 | 0 14129752 | 0 031257538 | 3 91E-24 | -0 34699413 | 0 | 7 69E-24 |
| SREK1IP1 | 0 249538368 | 0 031246958 | 4 05E-24 | 0 298954122 | 0 | 7 96E-24 |
| CHCHD3 | 0 56486646 | 0 031227497 | 4 31E-24 | 0 127585649 | 0 | 8 49E-24 |
| KIF19 | 0 003512212 | 0 031216228 | 4 48E-24 | 0 498331388 | 0 | 8 81E-24 |
| DNAJC24 | 0 217576288 | 0 031205856 | 4 64E-24 | -0 27880493 | 0 | 9 12E-24 |
| RHOXF1 | 0 116293236 | 0 031201818 | 4 70E-24 | -0 35105318 | 0 | 9 24E-24 |
| NFASC | 0 111619805 | 0 031193903 | 4 82E-24 | -0 34817622 | 0 | 9 48E-24 |
| ZNF844 | 0 070025318 | 0 031193666 | 4 83E-24 | -0 40895182 | 0 | 9 49E-24 |
| SEMA6B | 0 042003769 | 0 031159724 | 5 40E-24 | -0 44061955 | 0 | 1 06E-23 |
| MAFK | 0 110391959 | 0 031159271 | 5 41E-24 | -0 36426054 | 0 | 1 06E-23 |
| ACTR6 | 0 156945423 | 0 031154218 | 5 50E-24 | 0 368256316 | 0 | 1 08E-23 |
| KCNJ2 | 0 018322514 | 0 031148336 | 5 61E-24 | 0 481458977 | 0 | 1 10E-23 |
| KLHL18 | 0 152053073 | 0 031133984 | 5 88E-24 | -0 32892311 | 0 | 1 15E-23 |
| HS2ST1 | 0 391254688 | 0 0311231 | 6 10E-24 | 0 185082116 | 0 | 1 20E-23 |
| HOXD9 | 0 019902534 | 0 031115262 | 6 26E-24 | 0 481763553 | 0 | 1 23E-23 |
| KIT | 0 032114371 | 0 031062161 | 7 45E-24 | 0 473716318 | 0 | 1 46E-23 |
| MAOA | 0 053949097 | 0 031059124 | 7 53E-24 | -0 42928428 | 0 | 1 48E-23 |
| SLC16A10 | 0 078839162 | 0 031054356 | 7 65E-24 | -0 40846242 | 0 | 1 50E-23 |
| NIN | 0 366516914 | 0 031045411 | 7 88E-24 | 0 248895614 | 0 | 1 54E-23 |
| CHD4 | 0 342797586 | 0 031044549 | 7 90E-24 | 0 22385203 | 0 | 1 55E-23 |
| UVSSA | 0 203708287 | 0 031043348 | 7 93E-24 | -0 27511245 | 0 | 1 55E-23 |
| AATK | 0 117559156 | 0 031025125 | 8 42E-24 | -0 36055057 | 0 | 1 65E-23 |
| PPM1M | 0 044497535 | 0 031003294 | 9 05E-24 | -0 44818285 | 0 | 1 77E-23 |
| HPSE2 | 0 054187051 | 0 030988368 | 9 51E-24 | 0 455992644 | 0 | 1 86E-23 |
| GDI2 | 0 392577716 | 0 030979955 | 9 77E-24 | -0 14253895 | 0 | 1 91E-23 |
| SF3B1 | 0 631017875 | 0 030979006 | 9 80E-24 | -0 03406696 | 2 31E-28 | 1 92E-23 |
| HAS2-AS1 | 0 070891474 | 0 030976436 | 9 89E-24 | -0 41981487 | 0 | 1 93E-23 |
| ZNF654 | 0 398260075 | 0 030976043 | 9 90E-24 | 0 205951405 | 0 | 1 94E-23 |
| FBXO43 | 0 030972188 | 0 030970686 | 1 01E-23 | 0 445407854 | 0 | 1 97E-23 |
| ZNF707 | 0 042860406 | 0 030957267 | 1 05E-23 | 0 463249166 | 0 | 2 06E-23 |
| ABCC3 | 0 034855609 | 0 030928253 | 1 16E-23 | 0 470890136 | 0 | 2 26E-23 |
| CTHRC1 | 0 293598066 | 0 030906164 | 1 25E-23 | 0 207954619 | 0 | 2 43E-23 |
| ADCK5 | 0 054843807 | 0 0308942 | 1 30E-23 | 0 455904258 | 0 | 2 53E-23 |
| NECTIN3 | 0 320401287 | 0 030891236 | 1 31E-23 | 0 222507814 | 0 | 2 55E-23 |
| LINC01470 | 0 042936552 | 0 030880874 | 1 35E-23 | 0 465138668 | 0 | 2 64E-23 |
| SRSF5 | 0 595943348 | 0 030879078 | 1 36E-23 | 0 13911862 | 0 | 2 66E-23 |
| BARD1 | 0 206620852 | 0 030865071 | 1 43E-23 | 0 250458343 | 0 | 2 78E-23 |
| CHMP4A | 0 152643201 | 0 030862838 | 1 44E-23 | -0 3321695 | 0 | 2 80E-23 |
| GAL3ST4 | 0 046800937 | 0 030836484 | 1 56E-23 | -0 44136405 | 0 | 3 05E-23 |
| TPO | 0 020092898 | 0 030821194 | 1 65E-23 | 0 485576543 | 0 | 3 21E-23 |
| SNX25 | 0 301859854 | 0 030810285 | 1 71E-23 | -0 21317411 | 0 | 3 32E-23 |

| | | | | | | |
|---|---|---|---|---|---|---|
| LSM5 | 0 158592069 | 0 030808837 | 1 71E-23 | 0 366291803 | 0 | 3 34E-23 |
| MSN | 0 354800023 | 0 030802304 | 1 75E-23 | -0 15811714 | 0 | 3 41E-23 |
| TP53INP1 | 0 161314272 | 0 030793484 | 1 80E-23 | 0 368499298 | 0 | 3 51E-23 |
| SLC16A1-AS1 | 0 472435324 | 0 03078752 | 1 84E-23 | 0 141292344 | 0 | 3 58E-23 |
| PAK6 | 0 015248139 | 0 030787082 | 1 84E-23 | 0 477856412 | 0 | 3 58E-23 |
| PMFBP1 | 0 028468904 | 0 030774979 | 1 91E-23 | 0 471424545 | 0 | 3 73E-23 |
| ABCC1 | 0 296805696 | 0 030771092 | 1 94E-23 | 0 242587905 | 0 | 3 77E-23 |
| ARHGEF1 | 0 180731378 | 0 030753064 | 2 06E-23 | -0 30880415 | 0 | 4 00E-23 |
| ZNF438 | 0 329700558 | 0 030745134 | 2 11E-23 | -0 16761837 | 0 | 4 10E-23 |
| SEZ6 | 0 018151187 | 0 030700047 | 2 44E-23 | 0 485332029 | 0 | 4 75E-23 |
| LIPG | 0 020387961 | 0 030699023 | 2 45E-23 | 0 479164519 | 0 | 4 77E-23 |
| LLGL2 | 0 026945994 | 0 03068543 | 2 56E-23 | 0 479984318 | 0 | 4 98E-23 |
| DHX34 | 0 116131427 | 0 030682471 | 2 59E-23 | 0 388728506 | 0 | 5 03E-23 |
| ZBTB18 | 0 043507643 | 0 030658404 | 2 80E-23 | 0 458290745 | 0 | 5 44E-23 |
| LXN | 0 105861301 | 0 030654032 | 2 84E-23 | -0 37990065 | 0 | 5 51E-23 |
| KNTC1 | 0 212912376 | 0 030648022 | 2 90E-23 | 0 225362116 | 0 | 5 62E-23 |
| METTL8 | 0 264053606 | 0 0306343 | 3 03E-23 | 0 277636421 | 0 | 5 88E-23 |
| ENPP6 | 0 052321486 | 0 030624894 | 3 12E-23 | -0 43009036 | 0 | 6 06E-23 |
| MFAP1 | 0 115122499 | 0 030621622 | 3 16E-23 | 0 400172494 | 0 | 6 12E-23 |
| LINC01344 | 0 02896385 | 0 030605755 | 3 32E-23 | -0 45763389 | 0 | 6 44E-23 |
| SIPA1L2 | 0 279606328 | 0 030604495 | 3 34E-23 | -0 19742124 | 0 | 6 47E-23 |
| KLF10 | 0 104224172 | 0 030577339 | 3 64E-23 | 0 394092809 | 0 | 7 06E-23 |
| RPA1 | 0 198682683 | 0 03056583 | 3 78E-23 | 0 329295428 | 0 | 7 33E-23 |
| ELK3 | 0 30345891 | 0 030516005 | 4 45E-23 | 0 198485002 | 0 | 8 61E-23 |
| OSBPL11 | 0 145628296 | 0 030495434 | 4 75E-23 | 0 378647356 | 0 | 9 20E-23 |
| NECTIN2 | 0 256505682 | 0 030492781 | 4 79E-23 | -0 24101754 | 0 | 9 28E-23 |
| MEF2C-AS2 | 0 054520188 | 0 030487661 | 4 87E-23 | 0 447780582 | 0 | 9 43E-23 |
| TACC3 | 0 088224096 | 0 03046555 | 5 23E-23 | 0 371039155 | 0 | 1 01E-22 |
| CYFIP1 | 0 393110735 | 0 030463484 | 5 27E-23 | -0 14559247 | 0 | 1 02E-22 |
| PPIH | 0 126896499 | 0 030456141 | 5 40E-23 | 0 372182554 | 0 | 1 04E-22 |
| CA2 | 0 011022063 | 0 030441946 | 5 65E-23 | 0 492447657 | 0 | 1 09E-22 |
| CCDC28A | 0 101425825 | 0 030440169 | 5 68E-23 | -0 38342405 | 0 | 1 10E-22 |
| ST3GAL4 | 0 207867735 | 0 030426834 | 5 93E-23 | -0 28214074 | 0 | 1 15E-22 |
| NPM2 | 0 011260018 | 0 030426444 | 5 94E-23 | -0 48120797 | 0 | 1 15E-22 |
| TTC19 | 0 424720641 | 0 030424486 | 5 98E-23 | -0 12946329 | 0 | 1 15E-22 |
| CSNK1G1 | 0 52402391 | 0 03042018 | 6 06E-23 | 0 144673857 | 0 | 1 17E-22 |
| DUSP16 | 0 360292018 | 0 03039408 | 6 60E-23 | -0 15528504 | 0 | 1 27E-22 |
| KANSL2 | 0 166549276 | 0 030381249 | 6 87E-23 | 0 335245067 | 0 | 1 33E-22 |
| ANKDD1B | 0 128685919 | 0 030358605 | 7 39E-23 | -0 33954125 | 0 | 1 43E-22 |
| RNF217 | 0 356922579 | 0 030358252 | 7 40E-23 | -0 14804969 | 0 | 1 43E-22 |
| DXO | 0 042717633 | 0 03032434 | 8 26E-23 | -0 44256995 | 0 | 1 59E-22 |
| SLC25A44 | 0 068892654 | 0 030317579 | 8 44E-23 | 0 44781398 | 0 | 1 63E-22 |
| C6orf62 | 0 342502522 | 0 030282548 | 9 45E-23 | -0 16149951 | 0 | 1 82E-22 |
| ACBD5 | 0 141478365 | 0 03027414 | 9 71E-23 | -0 34328339 | 0 | 1 87E-22 |

| | | | | | | |
|---|---|---|---|---|---|---|
| YTHDC2 | 0 49595477 | 0 030269759 | 9 84E-23 | -0 08138244 | 7 61E-154 | 1 90E-22 |
| ABLIM1 | 0 451600008 | 0 030268025 | 9 90E-23 | -0 08189629 | 8 95E-156 | 1 91E-22 |
| MAP3K3 | 0 368867907 | 0 03026238 | 1 01E-22 | 0 198976056 | 0 | 1 94E-22 |
| ZWINT | 0 046239363 | 0 030228967 | 1 12E-22 | 0 423394252 | 0 | 2 16E-22 |
| BTN2A2 | 0 068102644 | 0 030202547 | 1 22E-22 | 0 447625818 | 0 | 2 35E-22 |
| PCDH12 | 0 030305915 | 0 030192812 | 1 26E-22 | -0 4623929 | 0 | 2 42E-22 |
| LINC00378 | 0 017599132 | 0 030185183 | 1 29E-22 | 0 486357079 | 0 | 2 48E-22 |
| TTK | 0 044240544 | 0 030182746 | 1 30E-22 | 0 423032581 | 0 | 2 50E-22 |
| DPYD | 0 722421047 | 0 030171964 | 1 35E-22 | 0 016359076 | 1 14E-07 | 2 59E-22 |
| HOOK1 | 0 033837163 | 0 030170993 | 1 35E-22 | 0 483271635 | 0 | 2 60E-22 |
| KCNJ8 | 0 012392682 | 0 030168805 | 1 36E-22 | 0 489218802 | 0 | 2 62E-22 |
| PIK3CD-AS2 | 0 015276694 | 0 030164319 | 1 38E-22 | -0 47786714 | 0 | 2 65E-22 |
| ADAMTSL2 | 0 035331519 | 0 030160917 | 1 40E-22 | -0 45670149 | 0 | 2 68E-22 |
| DNAJC15 | 0 178542194 | 0 030149833 | 1 45E-22 | -0 31217204 | 0 | 2 78E-22 |
| SOCS6 | 0 153328511 | 0 030134807 | 1 52E-22 | -0 33039752 | 0 | 2 91E-22 |
| ATP2A3 | 0 022700881 | 0 030130449 | 1 54E-22 | -0 4685563 | 0 | 2 95E-22 |
| AIG1 | 0 254107099 | 0 030120603 | 1 59E-22 | -0 23741923 | 0 | 3 05E-22 |
| FGF9 | 0 016066703 | 0 030113646 | 1 62E-22 | 0 489548865 | 0 | 3 11E-22 |
| TMEM59L | 0 08447393 | 0 030108983 | 1 65E-22 | -0 40600383 | 0 | 3 16E-22 |
| GALNT3 | 0 033827645 | 0 030062107 | 1 92E-22 | -0 45488372 | 0 | 3 67E-22 |
| PIK3CA | 0 51419162 | 0 030022541 | 2 17E-22 | -0 05816556 | 2 05E-79 | 4 16E-22 |
| NNMT | 0 2331766 | 0 030015213 | 2 22E-22 | -0 24973032 | 0 | 4 26E-22 |
| KLRK1-AS1 | 0 052892578 | 0 029999742 | 2 34E-22 | -0 42857151 | 0 | 4 47E-22 |
| SYNPO2L | 0 021225562 | 0 029998776 | 2 34E-22 | -0 47175323 | 0 | 4 49E-22 |
| GDPD1 | 0 056814072 | 0 029992379 | 2 39E-22 | -0 42861825 | 0 | 4 58E-22 |
| RASGRP1 | 0 061725457 | 0 029985132 | 2 45E-22 | 0 446779888 | 0 | 4 68E-22 |
| ESF1 | 0 369524662 | 0 029985048 | 2 45E-22 | 0 228484095 | 0 | 4 68E-22 |
| SH3BGRL | 0 391092878 | 0 029967379 | 2 59E-22 | -0 13077232 | 0 | 4 95E-22 |
| LRRC61 | 0 014638975 | 0 029955444 | 2 69E-22 | -0 47750386 | 0 | 5 14E-22 |
| SCG2 | 0 033998972 | 0 029952864 | 2 71E-22 | -0 45004543 | 0 | 5 18E-22 |
| LINC02389 | 0 016514058 | 0 029951963 | 2 72E-22 | -0 47432847 | 0 | 5 20E-22 |
| MEF2C | 0 372932173 | 0 029946737 | 2 77E-22 | -0 15592587 | 0 | 5 28E-22 |
| ARL4A | 0 159648588 | 0 029943056 | 2 80E-22 | 0 356738179 | 0 | 5 35E-22 |
| PSMA1 | 0 321924197 | 0 029916862 | 3 04E-22 | 0 264227937 | 0 | 5 81E-22 |
| FKBP11 | 0 131693667 | 0 029900931 | 3 20E-22 | 0 379556412 | 0 | 6 11E-22 |
| SIGLEC14 | 0 003864385 | 0 029892567 | 3 29E-22 | -0 48988905 | 0 | 6 27E-22 |
| LINC00158 | 0 01014639 | 0 029885625 | 3 36E-22 | -0 48314477 | 0 | 6 41E-22 |
| ACTC1 | 0 005539586 | 0 02987704 | 3 45E-22 | 0 495274974 | 0 | 6 58E-22 |
| RBM44 | 0 021035198 | 0 029868096 | 3 55E-22 | -0 46858537 | 0 | 6 77E-22 |
| NPAT | 0 168900268 | 0 0298545 | 3 71E-22 | 0 358095668 | 0 | 7 07E-22 |
| CCDC12 | 0 20637338 | 0 02985294 | 3 73E-22 | -0 28945112 | 0 | 7 10E-22 |
| GIT2 | 0 318830786 | 0 029843001 | 3 85E-22 | 0 241579567 | 0 | 7 32E-22 |
| R3HCC1L | 0 294673621 | 0 029840251 | 3 88E-22 | 0 270802474 | 0 | 7 39E-22 |
| ABHD17B | 0 114998763 | 0 029825024 | 4 07E-22 | 0 401540374 | 0 | 7 75E-22 |

| | | | | | | |
|---|---|---|---|---|---|---|
| CALY | 0 011260018 | 0 029823829 | 4 09E-22 | -0 48126826 | 0 | 7 78E-22 |
| CDO1 | 0 111410405 | 0 029820464 | 4 13E-22 | -0 36077803 | 0 | 7 86E-22 |
| LINC02770 | 0 006738878 | 0 029817063 | 4 18E-22 | -0 48805415 | 0 | 7 94E-22 |
| SPATA2L | 0 038434448 | 0 02979776 | 4 44E-22 | -0 44851058 | 0 | 8 44E-22 |
| LINC02149 | 0 012782928 | 0 029777912 | 4 73E-22 | 0 488950966 | 0 | 8 99E-22 |
| RTF1 | 0 33079515 | 0 029772403 | 4 81E-22 | 0 23731532 | 0 | 9 14E-22 |
| RPS4Y1 | 0 172079344 | 0 029763509 | 4 95E-22 | -0 32957813 | 0 | 9 40E-22 |
| DDX46 | 0 528126249 | 0 029757474 | 5 04E-22 | 0 120712946 | 0 | 9 58E-22 |
| CCDC150 | 0 090765453 | 0 029751095 | 5 14E-22 | 0 356241904 | 0 | 9 77E-22 |
| USP43 | 0 024157164 | 0 029738125 | 5 36E-22 | 0 482909418 | 0 | 1 02E-21 |
| MTDH | 0 634577678 | 0 029732387 | 5 46E-22 | 0 091758426 | 3 42E-195 | 1 04E-21 |
| ALKBH1 | 0 151777046 | 0 029729056 | 5 52E-22 | 0 369263719 | 0 | 1 05E-21 |
| SNAP23 | 0 274609278 | 0 029725673 | 5 58E-22 | -0 20705871 | 0 | 1 06E-21 |
| NR4A3 | 0 049561211 | 0 029720865 | 5 66E-22 | -0 43318171 | 0 | 1 07E-21 |
| HOPX | 0 018008414 | 0 029699042 | 6 06E-22 | 0 486024922 | 0 | 1 15E-21 |
| CAP2 | 0 134787078 | 0 029690243 | 6 24E-22 | 0 34464364 | 0 | 1 18E-21 |
| SCT | 0 005387295 | 0 029686487 | 6 31E-22 | -0 48810308 | 0 | 1 20E-21 |
| ZNF143 | 0 158363633 | 0 029666159 | 6 73E-22 | 0 361749902 | 0 | 1 27E-21 |
| MEF2D | 0 193238278 | 0 029656854 | 6 93E-22 | -0 28000483 | 0 | 1 31E-21 |
| HSD11B1 | 0 035341037 | 0 029647079 | 7 14E-22 | -0 45442243 | 0 | 1 35E-21 |
| TEX26-AS1 | 0 026165502 | 0 029639367 | 7 32E-22 | -0 46333238 | 0 | 1 39E-21 |
| FES | 0 05892711 | 0 029637703 | 7 36E-22 | -0 42892616 | 0 | 1 39E-21 |
| PLCH1 | 0 050246521 | 0 029618646 | 7 81E-22 | 0 455286682 | 0 | 1 48E-21 |
| SACS | 0 259446803 | 0 029618428 | 7 82E-22 | 0 266898534 | 0 | 1 48E-21 |
| GPC3 | 0 371628181 | 0 029611548 | 7 99E-22 | -0 15247928 | 0 | 1 51E-21 |
| CASC8 | 0 018617578 | 0 029596591 | 8 37E-22 | 0 480235932 | 0 | 1 58E-21 |
| MED26 | 0 106470465 | 0 029594533 | 8 43E-22 | 0 409911481 | 0 | 1 59E-21 |
| LYPLAL1-DT | 0 032142925 | 0 02956101 | 9 37E-22 | 0 477510104 | 0 | 1 77E-21 |
| FBXL14 | 0 103367535 | 0 029537805 | 1 01E-21 | 0 390519239 | 0 | 1 90E-21 |
| TGFB1 | 0 196455426 | 0 029516819 | 1 08E-21 | 0 309489011 | 0 | 2 03E-21 |
| ZFY | 0 096885648 | 0 029514874 | 1 08E-21 | -0 38870914 | 0 | 2 04E-21 |
| TMEM192 | 0 124060079 | 0 029513826 | 1 09E-21 | -0 3565052 | 0 | 2 05E-21 |
| PKD1L1 | 0 034351145 | 0 029478617 | 1 21E-21 | -0 45598594 | 0 | 2 29E-21 |
| TAF4B | 0 145171423 | 0 029471575 | 1 24E-21 | 0 388374679 | 0 | 2 34E-21 |
| LINC01950 | 0 024499819 | 0 029466095 | 1 26E-21 | -0 46656704 | 0 | 2 38E-21 |
| ZNF101 | 0 036483219 | 0 029465151 | 1 27E-21 | 0 468981376 | 0 | 2 39E-21 |
| GPATCH2L | 0 446812358 | 0 029464756 | 1 27E-21 | 0 169204278 | 0 | 2 39E-21 |
| TMEM204 | 0 082589328 | 0 029458422 | 1 29E-21 | -0 40271402 | 0 | 2 44E-21 |
| SLC25A24 | 0 223239611 | 0 029457184 | 1 30E-21 | 0 294754111 | 0 | 2 44E-21 |
| ARMCX3 | 0 214758904 | 0 029450854 | 1 32E-21 | -0 25974537 | 0 | 2 49E-21 |
| DLEC1 | 0 025489711 | 0 029438702 | 1 37E-21 | -0 46326099 | 0 | 2 59E-21 |
| NLGN4Y | 0 270259466 | 0 029421368 | 1 45E-21 | -0 24108268 | 0 | 2 73E-21 |
| LDB3 | 0 022405818 | 0 029416991 | 1 47E-21 | 0 48180608 | 0 | 2 77E-21 |
| MAP3K7CL | 0 094582247 | 0 029416422 | 1 47E-21 | 0 400501258 | 0 | 2 77E-21 |

| | | | | | | |
|---|---|---|---|---|---|---|
| LINC00504 | 0 037416002 | 0 029391638 | 1 59E-21 | -0 45187014 | 0 | 3 00E-21 |
| ERRFI1 | 0 230568617 | 0 029381098 | 1 65E-21 | -0 27943166 | 0 | 3 09E-21 |
| ST7-AS2 | 0 166025775 | 0 029379639 | 1 65E-21 | 0 3352634 | 0 | 3 11E-21 |
| PUM3 | 0 524851992 | 0 029366784 | 1 72E-21 | 0 11864073 | 0 | 3 24E-21 |
| ATAD5 | 0 201109821 | 0 029340525 | 1 87E-21 | 0 258808218 | 0 | 3 51E-21 |
| IVNS1ABP | 0 252108279 | 0 029317759 | 2 00E-21 | 0 312846953 | 0 | 3 77E-21 |
| GABRB1 | 0 022643772 | 0 02931277 | 2 04E-21 | 0 477251095 | 0 | 3 82E-21 |
| MARCKS | 0 38109878 | 0 029275677 | 2 29E-21 | -0 15498713 | 0 | 4 29E-21 |
| HNRNPH3 | 0 411452285 | 0 02926994 | 2 33E-21 | 0 211335153 | 0 | 4 37E-21 |
| AKIRIN1 | 0 180103177 | 0 029259064 | 2 41E-21 | -0 3045362 | 0 | 4 52E-21 |
| C3orf38 | 0 130732329 | 0 02924955 | 2 48E-21 | -0 35554543 | 0 | 4 65E-21 |
| COLGALT2 | 0 178770631 | 0 029248899 | 2 48E-21 | -0 31108275 | 0 | 4 66E-21 |
| CDH5 | 0 020378443 | 0 02923309 | 2 61E-21 | -0 471873 | 0 | 4 90E-21 |
| PSMD7-DT | 0 038872285 | 0 029223831 | 2 69E-21 | 0 474429753 | 0 | 5 04E-21 |
| THBS1 | 0 286430869 | 0 029210243 | 2 80E-21 | -0 1963453 | 0 | 5 25E-21 |
| TFAP2E | 0 05019893 | 0 029208674 | 2 81E-21 | -0 43904061 | 0 | 5 28E-21 |
| SULT1A3 | 0 037187565 | 0 02920407 | 2 86E-21 | 0 46926069 | 0 | 5 35E-21 |
| DUSP10 | 0 111343778 | 0 02916543 | 3 22E-21 | -0 37688665 | 0 | 6 03E-21 |
| WHAMM | 0 168690868 | 0 029150614 | 3 37E-21 | -0 32260668 | 0 | 6 31E-21 |
| ALOX12P2 | 0 125763835 | 0 029124193 | 3 66E-21 | 0 418255731 | 0 | 6 85E-21 |
| WNT2B | 0 117854219 | 0 029107147 | 3 86E-21 | 0 377565859 | 0 | 7 22E-21 |
| LMO2 | 0 023985837 | 0 029103011 | 3 91E-21 | 0 480169152 | 0 | 7 31E-21 |
| LINC02576 | 0 024880547 | 0 02909155 | 4 05E-21 | 0 481433876 | 0 | 7 57E-21 |
| OSR2 | 0 09114618 | 0 029084036 | 4 14E-21 | -0 37946914 | 0 | 7 75E-21 |
| HIC1 | 0 096390703 | 0 029078568 | 4 21E-21 | -0 38917249 | 0 | 7 88E-21 |
| WDR45B | 0 326702328 | 0 029069023 | 4 34E-21 | 0 244630642 | 0 | 8 11E-21 |
| EVA1C | 0 276741353 | 0 029048956 | 4 62E-21 | -0 21838533 | 0 | 8 63E-21 |
| TENT4B | 0 234461556 | 0 029040908 | 4 73E-21 | -0 26551 | 0 | 8 85E-21 |
| DNAAF2 | 0 056842626 | 0 029036587 | 4 80E-21 | -0 42820987 | 0 | 8 96E-21 |
| C21orf62 | 0 043859816 | 0 029030077 | 4 89E-21 | -0 44973751 | 0 | 9 14E-21 |
| AP3M2 | 0 160000761 | 0 029023557 | 4 99E-21 | 0 36146789 | 0 | 9 32E-21 |
| XRCC2 | 0 081989682 | 0 029023377 | 4 99E-21 | 0 356209475 | 0 | 9 33E-21 |
| LINC01411 | 0 015486094 | 0 029006929 | 5 25E-21 | 0 486817779 | 0 | 9 81E-21 |
| SCN3B | 0 01056519 | 0 028999325 | 5 38E-21 | -0 48196584 | 0 | 1 00E-20 |
| IER3 | 0 077934934 | 0 028989989 | 5 54E-21 | -0 40115842 | 0 | 1 03E-20 |
| NUMB | 0 640935828 | 0 028973733 | 5 82E-21 | -0 04071739 | 8 41E-40 | 1 09E-20 |
| PPP1R14A | 0 023224382 | 0 028971491 | 5 86E-21 | -0 46710235 | 0 | 1 09E-20 |
| USP24 | 0 49016771 | 0 028963407 | 6 01E-21 | -0 0799738 | 1 28E-148 | 1 12E-20 |
| RGS7BP | 0 047533837 | 0 028953398 | 6 20E-21 | -0 43994961 | 0 | 1 16E-20 |
| C4orf47 | 0 083512593 | 0 0289478 | 6 30E-21 | -0 38801703 | 0 | 1 17E-20 |
| CACNA1A | 0 240705488 | 0 028920067 | 6 87E-21 | 0 224547326 | 0 | 1 28E-20 |
| ENO3 | 0 021530144 | 0 028899292 | 7 32E-21 | 0 481637483 | 0 | 1 36E-20 |
| RNF217-AS1 | 0 093630428 | 0 028893023 | 7 46E-21 | -0 38689139 | 0 | 1 39E-20 |
| CCNT1 | 0 239915478 | 0 028892592 | 7 47E-21 | 0 291215344 | 0 | 1 39E-20 |

| | | | | | | |
|---|---|---|---|---|---|---|
| PRRG1 | 0 223934439 | 0 028869479 | 8 02E-21 | 0 291280052 | 0 | 1 49E-20 |
| LIFR | 0 453265691 | 0 028858653 | 8 29E-21 | -0 09386194 | 3 46E-204 | 1 54E-20 |
| HELQ | 0 148921589 | 0 028852376 | 8 45E-21 | 0 366832951 | 0 | 1 57E-20 |
| LINC00243 | 0 00572995 | 0 028820933 | 9 31E-21 | -0 48871953 | 0 | 1 73E-20 |
| ASTL | 0 006443814 | 0 028815952 | 9 45E-21 | -0 48740866 | 0 | 1 76E-20 |
| ITPRIPL1 | 0 016866231 | 0 028803223 | 9 83E-21 | 0 485315256 | 0 | 1 83E-20 |
| LAT2 | 0 069539891 | 0 028797736 | 9 99E-21 | 0 415042829 | 0 | 1 86E-20 |
| GABPB1 | 0 195341798 | 0 028787993 | 1 03E-20 | 0 307448086 | 0 | 1 91E-20 |
| LGI1 | 0 009423007 | 0 028775161 | 1 07E-20 | 0 493050054 | 0 | 1 99E-20 |
| CILP | 0 046382136 | 0 028765381 | 1 10E-20 | -0 44383334 | 0 | 2 05E-20 |
| ACVR1 | 0 41342255 | 0 02875921 | 1 12E-20 | -0 1453016 | 0 | 2 09E-20 |
| NDUFV2-AS1 | 0 064885496 | 0 028751208 | 1 15E-20 | 0 435655789 | 0 | 2 14E-20 |
| REEP4 | 0 03900554 | 0 02874292 | 1 18E-20 | 0 45755245 | 0 | 2 19E-20 |
| SH2B2 | 0 022196417 | 0 028737567 | 1 20E-20 | 0 482610586 | 0 | 2 23E-20 |
| GPRIN3 | 0 025889475 | 0 028726174 | 1 24E-20 | 0 479582964 | 0 | 2 30E-20 |
| TRIM25 | 0 169376178 | 0 028724158 | 1 25E-20 | 0 336398157 | 0 | 2 32E-20 |
| RAD9A | 0 360929737 | 0 028718833 | 1 27E-20 | 0 206466983 | 0 | 2 36E-20 |
| LINC01031 | 0 018265405 | 0 028714466 | 1 29E-20 | 0 484578799 | 0 | 2 39E-20 |
| GALNT14 | 0 079771944 | 0 028696498 | 1 36E-20 | 0 409027006 | 0 | 2 52E-20 |
| FAM13A-AS1 | 0 079048562 | 0 028691045 | 1 38E-20 | -0 40471943 | 0 | 2 56E-20 |
| CD163L1 | 0 037787211 | 0 028647574 | 1 58E-20 | -0 45591378 | 0 | 2 92E-20 |
| ANP32E | 0 135700824 | 0 0286399 | 1 62E-20 | 0 372316994 | 0 | 2 99E-20 |
| SNX30 | 0 183777198 | 0 028628777 | 1 67E-20 | -0 30446345 | 0 | 3 09E-20 |
| LRCH4 | 0 095943348 | 0 028620018 | 1 72E-20 | 0 423468666 | 0 | 3 18E-20 |
| RASA3 | 0 240981516 | 0 028610448 | 1 77E-20 | 0 284065812 | 0 | 3 27E-20 |
| FKBP15 | 0 243979745 | 0 02859432 | 1 86E-20 | 0 295224012 | 0 | 3 43E-20 |
| ZCCHC7 | 0 720431745 | 0 02858601 | 1 91E-20 | -0 0102555 | 0 000886757 | 3 52E-20 |
| ZCCHC2 | 0 164131656 | 0 0285662 | 2 02E-20 | -0 32395589 | 0 | 3 74E-20 |
| IRF4 | 0 010707963 | 0 028551035 | 2 12E-20 | -0 4821844 | 0 | 3 91E-20 |
| KLRC4 | 0 007909615 | 0 028548851 | 2 13E-20 | -0 48649009 | 0 | 3 94E-20 |
| OSR1 | 0 013611011 | 0 028545339 | 2 16E-20 | -0 48099077 | 0 | 3 98E-20 |
| ADAMTS14 | 0 104271763 | 0 02851973 | 2 33E-20 | 0 397613403 | 0 | 4 30E-20 |
| LINC02201 | 0 008642516 | 0 028506837 | 2 42E-20 | 0 493771964 | 0 | 4 47E-20 |
| TTC39B | 0 235223011 | 0 028489505 | 2 56E-20 | 0 285336575 | 0 | 4 71E-20 |
| TRMT10B | 0 199263292 | 0 028479046 | 2 64E-20 | -0 29441168 | 0 | 4 86E-20 |
| LINC00958 | 0 009784699 | 0 028472391 | 2 69E-20 | 0 493432387 | 0 | 4 95E-20 |
| HLA-DMA | 0 086882032 | 0 028455756 | 2 83E-20 | -0 40255779 | 0 | 5 21E-20 |
| CCL28 | 0 059298319 | 0 028453262 | 2 85E-20 | 0 471882157 | 0 | 5 25E-20 |
| OSGIN2 | 0 124155261 | 0 028433776 | 3 02E-20 | 0 380110814 | 0 | 5 57E-20 |
| ABCD2 | 0 013315947 | 0 028432565 | 3 04E-20 | 0 489800674 | 0 | 5 59E-20 |
| DGKA | 0 100531115 | 0 028395853 | 3 39E-20 | 0 414462001 | 0 | 6 24E-20 |
| MYOM2 | 0 019083969 | 0 028346611 | 3 94E-20 | -0 47237171 | 0 | 7 24E-20 |
| GRK7 | 0 019569397 | 0 028346079 | 3 94E-20 | -0 47097745 | 0 | 7 25E-20 |
| IDO2 | 0 010079762 | 0 02833365 | 4 09E-20 | 0 491629539 | 0 | 7 52E-20 |

| | | | | | | |
|---|---|---|---|---|---|---|
| ZNF93 | 0 067084198 | 0 028325521 | 4 19E-20 | 0 409670952 | 0 | 7 71E-20 |
| CORO2B | 0 141440292 | 0 028311537 | 4 37E-20 | -0 33993791 | 0 | 8 04E-20 |
| PICK1 | 0 056633226 | 0 028308635 | 4 41E-20 | 0 452638861 | 0 | 8 11E-20 |
| FGF12 | 0 152367174 | 0 028281695 | 4 79E-20 | -0 32705072 | 0 | 8 79E-20 |
| ABCA9-AS1 | 0 312196608 | 0 028268752 | 4 98E-20 | -0 20088379 | 0 | 9 14E-20 |
| CFLAR | 0 440996745 | 0 028266776 | 5 01E-20 | -0 10228613 | 2 75E-242 | 9 19E-20 |
| LINC00534 | 0 095990939 | 0 028257466 | 5 15E-20 | 0 411391536 | 0 | 9 45E-20 |
| SEMA4D | 0 109573395 | 0 028248428 | 5 29E-20 | -0 36343179 | 0 | 9 70E-20 |
| MCM3 | 0 119748339 | 0 028235962 | 5 49E-20 | 0 35454692 | 0 | 1 01E-19 |
| LINC02789 | 0 00504464 | 0 028222141 | 5 72E-20 | 0 495440621 | 0 | 1 05E-19 |
| THSD7B | 0 105804192 | 0 028204819 | 6 03E-20 | -0 37807676 | 0 | 1 10E-19 |
| KCNK13 | 0 011536045 | 0 028198229 | 6 15E-20 | 0 492476056 | 0 | 1 13E-19 |
| WEE2-AS1 | 0 049190002 | 0 028179803 | 6 50E-20 | 0 46173551 | 0 | 1 19E-19 |
| MAPKAPK3 | 0 044887781 | 0 028178832 | 6 52E-20 | 0 457127287 | 0 | 1 19E-19 |
| ZNF81 | 0 233738174 | 0 028167305 | 6 75E-20 | 0 300582045 | 0 | 1 24E-19 |
| NUP88 | 0 24311359 | 0 028158084 | 6 94E-20 | 0 305836111 | 0 | 1 27E-19 |
| CRCP | 0 254240353 | 0 02812739 | 7 60E-20 | 0 277928409 | 0 | 1 39E-19 |
| ZNRF3 | 0 285212541 | 0 028119034 | 7 80E-20 | 0 250366388 | 0 | 1 43E-19 |
| RUSC2 | 0 283603967 | 0 02811783 | 7 82E-20 | -0 20560313 | 0 | 1 43E-19 |
| LINC01967 | 0 026308275 | 0 028099557 | 8 26E-20 | 0 476384147 | 0 | 1 51E-19 |
| ARHGDIB | 0 050855685 | 0 028094203 | 8 40E-20 | 0 45207166 | 0 | 1 54E-19 |
| ARHGAP42 | 0 510650854 | 0 02809241 | 8 44E-20 | -0 05147331 | 1 42E-62 | 1 54E-19 |
| SERTAD2 | 0 273152995 | 0 028079158 | 8 78E-20 | -0 23328438 | 0 | 1 61E-19 |
| C2 | 0 077344806 | 0 028077042 | 8 84E-20 | -0 40973693 | 0 | 1 62E-19 |
| SLC14A1 | 0 06838819 | 0 028071845 | 8 98E-20 | 0 430926519 | 0 | 1 64E-19 |
| FBXL16 | 0 006891169 | 0 028070869 | 9 00E-20 | -0 48684454 | 0 | 1 64E-19 |
| CENPI | 0 09666673 | 0 028070457 | 9 01E-20 | 0 362267275 | 0 | 1 65E-19 |
| ATP7A | 0 118491938 | 0 028050478 | 9 57E-20 | -0 35928473 | 0 | 1 75E-19 |
| HTRA3 | 0 087224686 | 0 028040141 | 9 87E-20 | -0 40089096 | 0 | 1 80E-19 |
| TCEAL7 | 0 072871257 | 0 028038251 | 9 92E-20 | 0 447615345 | 0 | 1 81E-19 |
| LINC00869 | 0 076231178 | 0 028028544 | 1 02E-19 | -0 39438151 | 0 | 1 86E-19 |
| PAXBP1-AS1 | 0 094020674 | 0 028022661 | 1 04E-19 | 0 40424075 | 0 | 1 90E-19 |
| TRIM56 | 0 287506425 | 0 028015075 | 1 06E-19 | -0 22034294 | 0 | 1 94E-19 |
| MLF1 | 0 184224553 | 0 028011528 | 1 07E-19 | -0 28340627 | 0 | 1 96E-19 |
| DERA | 0 321867088 | 0 028010118 | 1 08E-19 | 0 23674188 | 0 | 1 97E-19 |
| BRIP1 | 0 18991643 | 0 02798545 | 1 16E-19 | 0 225431562 | 0 | 2 12E-19 |
| AGBL1 | 0 130132684 | 0 027984199 | 1 17E-19 | -0 32934385 | 0 | 2 12E-19 |
| JAK1 | 0 656250595 | 0 027983748 | 1 17E-19 | 0 002836952 | 0 35781325 | 2 13E-19 |
| ARL5C | 0 004559213 | 0 027970217 | 1 22E-19 | 0 496064973 | 0 | 2 21E-19 |
| TJP2 | 0 154784794 | 0 027945629 | 1 31E-19 | 0 360183847 | 0 | 2 38E-19 |
| TECPR1 | 0 138832309 | 0 027939003 | 1 33E-19 | -0 34570364 | 0 | 2 43E-19 |
| BLVRA | 0 121842341 | 0 027936506 | 1 34E-19 | 0 397805979 | 0 | 2 44E-19 |
| SMPD3 | 0 027269612 | 0 027935954 | 1 35E-19 | -0 45873839 | 0 | 2 45E-19 |
| RAB11A | 0 443195446 | 0 027919336 | 1 41E-19 | -0 13001497 | 0 | 2 57E-19 |

| | | | | | | |
|---|---|---|---|---|---|---|
| E2F1 | 0 03721612 | 0 02789348 | 1 53E-19 | 0 435156348 | 0 | 2 78E-19 |
| IQSEC1 | 0 265814471 | 0 027858424 | 1 69E-19 | -0 21761232 | 0 | 3 08E-19 |
| SEC61A2 | 0 140831128 | 0 027857555 | 1 70E-19 | 0 387906682 | 0 | 3 09E-19 |
| TNFRSF10C | 0 008928062 | 0 027818311 | 1 91E-19 | -0 48297161 | 0 | 3 46E-19 |
| PPARGC1B | 0 086320458 | 0 027814865 | 1 93E-19 | -0 39489488 | 0 | 3 50E-19 |
| RAPGEF5 | 0 171022825 | 0 027799014 | 2 02E-19 | -0 30840058 | 0 | 3 67E-19 |
| FBXL5 | 0 449296606 | 0 027771732 | 2 19E-19 | 0 169811803 | 0 | 3 97E-19 |
| SPAG17 | 0 098018313 | 0 027748249 | 2 35E-19 | -0 38662039 | 0 | 4 26E-19 |
| USP51 | 0 037225638 | 0 027728241 | 2 49E-19 | -0 45037265 | 0 | 4 52E-19 |
| LINC01237 | 0 173659363 | 0 027727975 | 2 49E-19 | -0 3113962 | 0 | 4 52E-19 |
| ZMYM5 | 0 254706745 | 0 027727634 | 2 49E-19 | -0 24387581 | 0 | 4 52E-19 |
| ZNF786 | 0 081133045 | 0 027716622 | 2 58E-19 | -0 40485569 | 0 | 4 67E-19 |
| MITF | 0 255801336 | 0 02771104 | 2 62E-19 | -0 22227164 | 0 | 4 75E-19 |
| GPR82 | 0 041128096 | 0 027704041 | 2 67E-19 | 0 463967785 | 0 | 4 85E-19 |
| IDI2-AS1 | 0 015428985 | 0 02768461 | 2 83E-19 | -0 47708325 | 0 | 5 13E-19 |
| EPS8L2 | 0 052473777 | 0 027683816 | 2 84E-19 | -0 43141775 | 0 | 5 14E-19 |
| PTK2B | 0 087986142 | 0 027672489 | 2 93E-19 | -0 39789875 | 0 | 5 31E-19 |
| MORF4L2-AS1 | 0 018912642 | 0 027670962 | 2 95E-19 | -0 47295003 | 0 | 5 34E-19 |
| PARP12 | 0 150063772 | 0 027665254 | 3 00E-19 | -0 34263292 | 0 | 5 42E-19 |
| RBM4 | 0 365898232 | 0 02766246 | 3 02E-19 | 0 198920413 | 0 | 5 47E-19 |
| AATF | 0 284527232 | 0 027654825 | 3 09E-19 | 0 278645696 | 0 | 5 59E-19 |
| FMO1 | 0 112124269 | 0 027630915 | 3 32E-19 | -0 36493224 | 0 | 6 00E-19 |
| SCLY | 0 163503455 | 0 027627189 | 3 35E-19 | 0 343857767 | 0 | 6 06E-19 |
| AKAP13 | 0 857246197 | 0 027625162 | 3 37E-19 | 0 012031154 | 9 63E-05 | 6 10E-19 |
| TPRN | 0 03069616 | 0 027624694 | 3 38E-19 | -0 45991845 | 0 | 6 10E-19 |
| DNAH17 | 0 043126916 | 0 027612321 | 3 50E-19 | 0 462153542 | 0 | 6 33E-19 |
| CDA | 0 00893758 | 0 027597037 | 3 66E-19 | 0 491250103 | 0 | 6 62E-19 |
| MBOAT7 | 0 121861377 | 0 027594538 | 3 69E-19 | 0 394812714 | 0 | 6 67E-19 |
| CHCHD7 | 0 09687613 | 0 027566997 | 4 00E-19 | 0 406290721 | 0 | 7 23E-19 |
| FAM163A | 0 029059032 | 0 027553509 | 4 16E-19 | 0 474477957 | 0 | 7 51E-19 |
| ANKMY1 | 0 215644096 | 0 027549061 | 4 22E-19 | 0 306259269 | 0 | 7 61E-19 |
| TIGAR | 0 150292208 | 0 027545393 | 4 26E-19 | 0 357611514 | 0 | 7 69E-19 |
| LINC01980 | 0 014448611 | 0 02754063 | 4 32E-19 | -0 48298172 | 0 | 7 80E-19 |
| NPR3 | 0 037672993 | 0 027524542 | 4 53E-19 | -0 45539651 | 0 | 8 17E-19 |
| TGFBR3L | 0 007995279 | 0 027523388 | 4 55E-19 | -0 48480855 | 0 | 8 20E-19 |
| PLK5 | 0 077820715 | 0 027520321 | 4 59E-19 | -0 38903848 | 0 | 8 27E-19 |
| SNHG15 | 0 097656622 | 0 027504429 | 4 81E-19 | -0 39119423 | 0 | 8 66E-19 |
| EIF3H | 0 463183644 | 0 027499005 | 4 88E-19 | 0 141010166 | 0 | 8 79E-19 |
| FGD3 | 0 013753783 | 0 027492796 | 4 97E-19 | 0 489791692 | 0 | 8 95E-19 |
| PELI1 | 0 325864727 | 0 027470741 | 5 30E-19 | -0 17722443 | 0 | 9 55E-19 |
| VPS33A | 0 130066056 | 0 027464409 | 5 40E-19 | -0 36090069 | 0 | 9 73E-19 |
| PLIN5 | 0 018303478 | 0 027457052 | 5 52E-19 | -0 47221564 | 0 | 9 94E-19 |
| MAT2B | 0 104262245 | 0 027455673 | 5 54E-19 | 0 402329426 | 0 | 9 97E-19 |
| C2CD2 | 0 226894596 | 0 027447895 | 5 67E-19 | -0 25810318 | 0 | 1 02E-18 |

| | | | | | | |
|---|---|---|---|---|---|---|
| PNPLA7 | 0 097266376 | 0 027445124 | 5 72E-19 | -0 38341889 | 0 | 1 03E-18 |
| DENND4B | 0 154794312 | 0 027423356 | 6 09E-19 | 0 354685861 | 0 | 1 09E-18 |
| MRPL1 | 0 261064895 | 0 027409524 | 6 34E-19 | 0 286386935 | 0 | 1 14E-18 |
| CGGBP1 | 0 428118635 | 0 027386776 | 6 78E-19 | 0 184785709 | 0 | 1 22E-18 |
| C9 | 0 018655651 | 0 027384884 | 6 81E-19 | -0 47494025 | 0 | 1 22E-18 |
| PLCXD3 | 0 033399326 | 0 027374379 | 7 03E-19 | -0 4508277 | 0 | 1 26E-18 |
| VCAN | 0 617911329 | 0 027366847 | 7 18E-19 | -0 04533899 | 6 15E-49 | 1 29E-18 |
| FBLIM1 | 0 105870819 | 0 027349134 | 7 56E-19 | -0 37468632 | 0 | 1 36E-18 |
| MYO1D | 0 279358855 | 0 027311947 | 8 43E-19 | -0 1977957 | 0 | 1 51E-18 |
| SEZ6L2 | 0 060964002 | 0 027305151 | 8 59E-19 | -0 42572971 | 0 | 1 54E-18 |
| NFX1 | 0 407035846 | 0 027274696 | 9 39E-19 | -0 14116765 | 0 | 1 68E-18 |
| ZIK1 | 0 015952485 | 0 02726441 | 9 67E-19 | 0 490098684 | 0 | 1 73E-18 |
| PDGFC | 0 348556091 | 0 02725219 | 1 00E-18 | 0 164033652 | 0 | 1 79E-18 |
| CENPW | 0 028554568 | 0 027249605 | 1 01E-18 | 0 463225936 | 0 | 1 81E-18 |
| ARMC5 | 0 031781234 | 0 027238922 | 1 04E-18 | -0 45684545 | 0 | 1 86E-18 |
| HAS1 | 0 006139232 | 0 027233551 | 1 06E-18 | 0 493794598 | 0 | 1 89E-18 |
| TNFRSF11A | 0 01797986 | 0 027229654 | 1 07E-18 | 0 48433351 | 0 | 1 91E-18 |
| TVP23A | 0 039195903 | 0 027221306 | 1 10E-18 | 0 455251501 | 0 | 1 96E-18 |
| SLC5A6 | 0 0577183 | 0 0272197 | 1 10E-18 | 0 446966489 | 0 | 1 97E-18 |
| RDH10-AS1 | 0 056595153 | 0 027208894 | 1 14E-18 | -0 43186966 | 0 | 2 03E-18 |
| ATP2A2 | 0 50704346 | 0 027206966 | 1 14E-18 | 0 141331715 | 0 | 2 04E-18 |
| OSER1 | 0 117644819 | 0 027203352 | 1 15E-18 | 0 405128554 | 0 | 2 06E-18 |
| SLC30A7 | 0 354942796 | 0 027189744 | 1 20E-18 | 0 225260778 | 0 | 2 15E-18 |
| LGALS9 | 0 024537892 | 0 027161926 | 1 30E-18 | -0 46594518 | 0 | 2 32E-18 |
| CORO2A | 0 037130456 | 0 027157459 | 1 32E-18 | 0 46831011 | 0 | 2 35E-18 |
| ADRA1A | 0 044069216 | 0 027147186 | 1 36E-18 | -0 44925546 | 0 | 2 43E-18 |
| MYB | 0 005710914 | 0 027141715 | 1 38E-18 | 0 493650492 | 0 | 2 46E-18 |
| NFIL3 | 0 116873846 | 0 027140585 | 1 38E-18 | -0 35952743 | 0 | 2 47E-18 |
| TCF21 | 0 014686566 | 0 027138813 | 1 39E-18 | -0 47721359 | 0 | 2 48E-18 |
| ARRDC2 | 0 044706935 | 0 027123259 | 1 46E-18 | -0 4430923 | 0 | 2 60E-18 |
| NDC80 | 0 08778626 | 0 027095512 | 1 58E-18 | 0 372898097 | 0 | 2 81E-18 |
| CD2AP | 0 425263178 | 0 027094994 | 1 58E-18 | 0 188418228 | 0 | 2 82E-18 |
| DNAJB6 | 0 428185262 | 0 027079191 | 1 65E-18 | -0 12311508 | 0 | 2 95E-18 |
| ATE1 | 0 454503055 | 0 02707722 | 1 66E-18 | 0 171544253 | 0 | 2 96E-18 |
| RASA2 | 0 542203651 | 0 027073314 | 1 68E-18 | 0 137728854 | 0 | 3 00E-18 |
| SLC6A6 | 0 109935086 | 0 027072424 | 1 69E-18 | -0 35604482 | 0 | 3 00E-18 |
| ZNF92 | 0 11869182 | 0 027068776 | 1 70E-18 | 0 359229411 | 0 | 3 03E-18 |
| CABCOCO1 | 0 035045973 | 0 027053639 | 1 78E-18 | -0 45100103 | 0 | 3 17E-18 |
| LINC02785 | 0 003340884 | 0 027044177 | 1 83E-18 | 0 496017449 | 0 | 3 25E-18 |
| NIPSNAP3B | 0 044716453 | 0 027035339 | 1 88E-18 | 0 467607798 | 0 | 3 34E-18 |
| CABYR | 0 05603358 | 0 027013525 | 2 00E-18 | 0 445306411 | 0 | 3 55E-18 |
| USP9Y | 0 307389922 | 0 0270057 | 2 04E-18 | -0 21979489 | 0 | 3 63E-18 |
| KCTD16 | 0 056223944 | 0 026982028 | 2 19E-18 | -0 42608199 | 0 | 3 89E-18 |
| EDIL3-DT | 0 042917515 | 0 026963652 | 2 30E-18 | -0 44225433 | 0 | 4 09E-18 |

| | | | | | | |
|---|---|---|---|---|---|---|
| PYROXD1 | 0 151624755 | 0 026963522 | 2 31E-18 | -0 33793397 | 0 | 4 10E-18 |
| HLA-F | 0 104547791 | 0 02695374 | 2 37E-18 | 0 40110367 | 0 | 4 21E-18 |
| ATG9B | 0 009346862 | 0 026950374 | 2 39E-18 | -0 48384542 | 0 | 4 25E-18 |
| LGMN | 0 141878129 | 0 026949166 | 2 40E-18 | -0 33841804 | 0 | 4 27E-18 |
| THUMPD1 | 0 17178428 | 0 026929919 | 2 54E-18 | 0 35700812 | 0 | 4 51E-18 |
| LINC00326 | 0 00420704 | 0 026914629 | 2 65E-18 | 0 496686274 | 0 | 4 71E-18 |
| CLU | 0 2324437 | 0 026914515 | 2 65E-18 | -0 21658917 | 0 | 4 71E-18 |
| FAM110C | 0 006262969 | 0 026903481 | 2 74E-18 | 0 495084932 | 0 | 4 86E-18 |
| CTDP1 | 0 081142563 | 0 026892025 | 2 83E-18 | -0 40193594 | 0 | 5 02E-18 |
| CD207 | 0 002607984 | 0 026884416 | 2 89E-18 | 0 496760691 | 0 | 5 13E-18 |
| CASP6 | 0 072795111 | 0 026842537 | 3 26E-18 | 0 434449829 | 0 | 5 78E-18 |
| GATAD2A | 0 319830196 | 0 026836444 | 3 32E-18 | 0 254086325 | 0 | 5 88E-18 |
| UPB1 | 0 010831699 | 0 026824376 | 3 43E-18 | -0 48151871 | 0 | 6 08E-18 |
| DDX3X | 0 415373779 | 0 026811003 | 3 57E-18 | -0 11348882 | 3 95E-298 | 6 32E-18 |
| FER | 0 71252213 | 0 02680811 | 3 60E-18 | 0 016482891 | 9 14E-08 | 6 37E-18 |
| TTYH1 | 0 044583198 | 0 026805451 | 3 63E-18 | -0 44540063 | 0 | 6 42E-18 |
| DISC1 | 0 344587006 | 0 026777294 | 3 93E-18 | -0 1653422 | 0 | 6 95E-18 |
| LONRF3 | 0 070767737 | 0 026768118 | 4 03E-18 | -0 40459703 | 0 | 7 13E-18 |
| ITGB5 | 0 272943595 | 0 026755816 | 4 18E-18 | -0 21408225 | 0 | 7 39E-18 |
| SLC2A14 | 0 013858484 | 0 026751601 | 4 23E-18 | 0 486926105 | 0 | 7 47E-18 |
| CCNI | 0 498543717 | 0 026741678 | 4 35E-18 | -0 09988259 | 4 46E-231 | 7 69E-18 |
| HTR2C | 0 015600312 | 0 026733856 | 4 45E-18 | 0 4858654 | 0 | 7 86E-18 |
| CLEC2D | 0 367068969 | 0 026729568 | 4 50E-18 | 0 153615796 | 0 | 7 95E-18 |
| SH2D3A | 0 015476576 | 0 026711915 | 4 73E-18 | 0 486844385 | 0 | 8 36E-18 |
| MAN2B2 | 0 106479983 | 0 026702949 | 4 85E-18 | -0 37601391 | 0 | 8 58E-18 |
| ST6GALNAC5 | 0 141649693 | 0 026696696 | 4 94E-18 | 0 422281145 | 0 | 8 73E-18 |
| TNNI3K | 0 134549123 | 0 026689387 | 5 05E-18 | -0 33610785 | 0 | 8 91E-18 |
| LRRC8D | 0 205221679 | 0 026670159 | 5 33E-18 | -0 28284751 | 0 | 9 41E-18 |
| CGAS | 0 08641564 | 0 026664276 | 5 42E-18 | 0 418309691 | 0 | 9 57E-18 |
| BUB1B | 0 069263863 | 0 026630619 | 5 96E-18 | 0 377888195 | 0 | 1 05E-17 |
| BHLHE41 | 0 092107517 | 0 026617083 | 6 19E-18 | -0 38468771 | 0 | 1 09E-17 |
| ZC3H8 | 0 281062611 | 0 0266002 | 6 50E-18 | 0 237878321 | 0 | 1 15E-17 |
| PGGHG | 0 056204908 | 0 026596205 | 6 57E-18 | -0 43037368 | 0 | 1 16E-17 |
| DEPDC1B | 0 076278769 | 0 026559664 | 7 29E-18 | 0 376479454 | 0 | 1 29E-17 |
| KDM7A | 0 32858693 | 0 026548875 | 7 51E-18 | -0 14197264 | 0 | 1 32E-17 |
| GCSAML | 0 002293884 | 0 026538973 | 7 73E-18 | -0 49340563 | 0 | 1 36E-17 |
| APCDD1 | 0 110810759 | 0 026537183 | 7 77E-18 | -0 36946147 | 0 | 1 37E-17 |
| SELE | 0 006700805 | 0 026523698 | 8 07E-18 | 0 494444105 | 0 | 1 42E-17 |
| KIF5B | 0 391092878 | 0 026508517 | 8 42E-18 | 0 228657783 | 0 | 1 48E-17 |
| GALNT15 | 0 034522472 | 0 026476163 | 9 23E-18 | -0 45708004 | 0 | 1 62E-17 |
| ELOC | 0 226561459 | 0 026471836 | 9 34E-18 | 0 304294594 | 0 | 1 64E-17 |
| CMC2 | 0 221754773 | 0 026450114 | 9 93E-18 | 0 306900335 | 0 | 1 75E-17 |
| PDE8A | 0 588157469 | 0 026433026 | 1 04E-17 | 0 122855333 | 0 | 1 83E-17 |
| PTGS1 | 0 018436733 | 0 026423801 | 1 07E-17 | 0 483897848 | 0 | 1 88E-17 |

| | | | | | | |
|---|---|---|---|---|---|---|
| RINL | 0 016437913 | 0 02642251 | 1 07E-17 | -0 47584989 | 0 | 1 89E-17 |
| IL10RB | 0 173507072 | 0 02641668 | 1 09E-17 | -0 3203364 | 0 | 1 92E-17 |
| SLC25A40 | 0 184776608 | 0 026393874 | 1 16E-17 | 0 312238152 | 0 | 2 04E-17 |
| CD160 | 0 03779673 | 0 02638858 | 1 18E-17 | -0 45526237 | 0 | 2 07E-17 |
| JPT1 | 0 130475338 | 0 026376536 | 1 22E-17 | 0 34959494 | 0 | 2 14E-17 |
| ADCYAP1 | 0 007909615 | 0 026376389 | 1 22E-17 | 0 492774504 | 0 | 2 15E-17 |
| DHRS7B | 0 174639737 | 0 026372202 | 1 24E-17 | 0 362347778 | 0 | 2 17E-17 |
| MAP3K20 | 0 381603244 | 0 02623677 | 1 81E-17 | 0 139796979 | 0 | 3 17E-17 |
| FLT4 | 0 010327235 | 0 026206242 | 1 97E-17 | 0 489157722 | 0 | 3 45E-17 |
| PCSK1N | 0 044716453 | 0 026203023 | 1 99E-17 | -0 4409232 | 0 | 3 48E-17 |
| PPP1R12A | 0 732567436 | 0 026197772 | 2 02E-17 | 0 052169143 | 3 15E-64 | 3 53E-17 |
| PODXL | 0 043897889 | 0 026178625 | 2 13E-17 | -0 44911623 | 0 | 3 73E-17 |
| TOR1B | 0 074660677 | 0 026138643 | 2 38E-17 | 0 438230232 | 0 | 4 16E-17 |
| PTPRN2 | 0 192429232 | 0 026120898 | 2 50E-17 | 0 369105605 | 0 | 4 37E-17 |
| ALKAL2 | 0 018569987 | 0 026095155 | 2 68E-17 | -0 47534629 | 0 | 4 70E-17 |
| CELF1 | 0 60055967 | 0 026084562 | 2 76E-17 | 0 104633808 | 1 68E-253 | 4 84E-17 |
| HSPH1 | 0 267946546 | 0 026080757 | 2 79E-17 | -0 23385038 | 0 | 4 89E-17 |
| LY86-AS1 | 0 008756734 | 0 02607845 | 2 81E-17 | 0 49275006 | 0 | 4 92E-17 |
| DNAH12 | 0 063714759 | 0 026076155 | 2 83E-17 | 0 451374962 | 0 | 4 95E-17 |
| DMXL1 | 0 521625326 | 0 026065265 | 2 91E-17 | 0 155484568 | 0 | 5 10E-17 |
| USP15 | 0 572661857 | 0 026054696 | 3 00E-17 | 0 108998637 | 4 90E-275 | 5 25E-17 |
| MMP25 | 0 019578915 | 0 026040069 | 3 13E-17 | -0 47127789 | 0 | 5 47E-17 |
| KCNN2 | 0 041347014 | 0 026018635 | 3 32E-17 | 0 479047667 | 0 | 5 80E-17 |
| SV2C | 0 010098799 | 0 02598156 | 3 68E-17 | 0 491432991 | 0 | 6 42E-17 |
| HMCN1 | 0 492337858 | 0 025977762 | 3 71E-17 | -0 08198316 | 4 21E-156 | 6 49E-17 |
| PTPN18 | 0 119529421 | 0 025969971 | 3 80E-17 | 0 407267502 | 0 | 6 63E-17 |
| C1QTNF7-AS1 | 0 121594868 | 0 025919518 | 4 36E-17 | -0 34276639 | 0 | 7 62E-17 |
| FRG1 | 0 169985342 | 0 025917106 | 4 39E-17 | 0 370122548 | 0 | 7 67E-17 |
| CDK15 | 0 067674326 | 0 025907175 | 4 52E-17 | -0 41539422 | 0 | 7 88E-17 |
| H1-3 | 0 071757629 | 0 025904802 | 4 55E-17 | 0 422790226 | 0 | 7 93E-17 |
| TANC2 | 0 645856732 | 0 025892525 | 4 70E-17 | 0 032703478 | 2 89E-26 | 8 20E-17 |
| EFHD1 | 0 091317508 | 0 025892188 | 4 71E-17 | -0 37630522 | 0 | 8 21E-17 |
| TMX4 | 0 218823171 | 0 025879352 | 4 88E-17 | -0 27202784 | 0 | 8 50E-17 |
| LINC02355 | 0 006967315 | 0 025865798 | 5 06E-17 | 0 494781022 | 0 | 8 82E-17 |
| CTR9 | 0 154337439 | 0 025827136 | 5 63E-17 | -0 32739877 | 0 | 9 81E-17 |
| KLHL5 | 0 293293484 | 0 0258096 | 5 91E-17 | 0 236831836 | 0 | 1 03E-16 |
| NUSAP1 | 0 110115932 | 0 025801929 | 6 03E-17 | 0 330048576 | 0 | 1 05E-16 |
| HMGB3 | 0 033703908 | 0 025798798 | 6 09E-17 | 0 460733551 | 0 | 1 06E-16 |
| DKC1 | 0 133283204 | 0 025796298 | 6 13E-17 | 0 378083967 | 0 | 1 07E-16 |
| LHFPL3 | 0 043621861 | 0 025749626 | 6 97E-17 | 0 468572751 | 0 | 1 21E-16 |
| UBE2G1 | 0 327520892 | 0 025749218 | 6 98E-17 | 0 249467883 | 0 | 1 21E-16 |
| CATSPER2 | 0 11537949 | 0 025716882 | 7 62E-17 | -0 36976421 | 0 | 1 33E-16 |
| DCAF10 | 0 433448821 | 0 025710333 | 7 76E-17 | 0 182338709 | 0 | 1 35E-16 |
| COL4A3 | 0 035445737 | 0 025704645 | 7 88E-17 | -0 4569168 | 0 | 1 37E-16 |

| | | | | | | |
|---|---|---|---|---|---|---|
| LSP1 | 0 139450991 | 0 025704538 | 7 88E-17 | -0 35457969 | 0 | 1 37E-16 |
| RBM25 | 0 628819173 | 0 025699167 | 8 00E-17 | 0 097594057 | 1 18E-220 | 1 39E-16 |
| TRIM71 | 0 038044202 | 0 025685896 | 8 30E-17 | -0 44300148 | 0 | 1 44E-16 |
| TARS3 | 0 153261883 | 0 025646128 | 9 25E-17 | -0 3076895 | 0 | 1 61E-16 |
| GPR132 | 0 011612191 | 0 025611854 | 1 02E-16 | -0 48137097 | 0 | 1 76E-16 |
| STK32A | 0 08447393 | 0 025610191 | 1 02E-16 | -0 38377073 | 0 | 1 77E-16 |
| TNK1 | 0 011231463 | 0 025607591 | 1 03E-16 | 0 491473699 | 0 | 1 78E-16 |
| MIR4435-2HG | 0 47145495 | 0 025600252 | 1 05E-16 | 0 108052319 | 2 73E-270 | 1 82E-16 |
| CENPH | 0 050750985 | 0 025587601 | 1 09E-16 | 0 415901372 | 0 | 1 88E-16 |
| JARID2 | 0 566693952 | 0 025542441 | 1 23E-16 | -0 03003696 | 2 08E-22 | 2 13E-16 |
| SCNN1A | 0 017932269 | 0 025533225 | 1 26E-16 | 0 48333816 | 0 | 2 18E-16 |
| PHKB | 0 477203937 | 0 025524785 | 1 29E-16 | -0 09908811 | 1 97E-227 | 2 23E-16 |
| CEPT1 | 0 258019074 | 0 025498319 | 1 38E-16 | 0 28644679 | 0 | 2 40E-16 |
| MPHOSPH6 | 0 128371819 | 0 025486538 | 1 43E-16 | 0 376355678 | 0 | 2 48E-16 |
| RAI2 | 0 042108469 | 0 025473553 | 1 48E-16 | -0 43768438 | 0 | 2 56E-16 |
| GALC | 0 141744874 | 0 025471431 | 1 49E-16 | -0 33626571 | 0 | 2 58E-16 |
| COQ2 | 0 044821153 | 0 025465935 | 1 51E-16 | 0 443574506 | 0 | 2 62E-16 |
| KLHDC4 | 0 173002608 | 0 025464048 | 1 52E-16 | 0 351926679 | 0 | 2 63E-16 |
| DIAPH2 | 0 571291238 | 0 025451308 | 1 57E-16 | 0 118256519 | 0 | 2 72E-16 |
| ZSWIM3 | 0 033722945 | 0 025441494 | 1 62E-16 | -0 45556255 | 0 | 2 79E-16 |
| TNFRSF4 | 0 012792446 | 0 02543898 | 1 63E-16 | -0 48157295 | 0 | 2 81E-16 |
| OLFML2A | 0 110096895 | 0 025436645 | 1 64E-16 | -0 36798331 | 0 | 2 83E-16 |
| PLEKHB2 | 0 256696046 | 0 025426964 | 1 68E-16 | 0 269105912 | 0 | 2 90E-16 |
| TRDMT1 | 0 14555215 | 0 025406938 | 1 77E-16 | -0 33413409 | 0 | 3 06E-16 |
| MAP2K1 | 0 297795587 | 0 025397731 | 1 82E-16 | 0 250032287 | 0 | 3 14E-16 |
| PDLIM1 | 0 115027317 | 0 02538927 | 1 86E-16 | -0 37231552 | 0 | 3 21E-16 |
| KIFC1 | 0 049342293 | 0 025376668 | 1 93E-16 | 0 405877498 | 0 | 3 32E-16 |
| H1-2 | 0 061601721 | 0 025357167 | 2 03E-16 | 0 446603972 | 0 | 3 50E-16 |
| LIPA | 0 234813729 | 0 025354111 | 2 05E-16 | -0 25856919 | 0 | 3 53E-16 |
| CPEB1 | 0 205393006 | 0 025351517 | 2 06E-16 | -0 28895719 | 0 | 3 56E-16 |
| RHOF | 0 042546306 | 0 025324788 | 2 22E-16 | 0 459315874 | 0 | 3 82E-16 |
| KCNJ6 | 0 013468238 | 0 025320514 | 2 24E-16 | -0 47857241 | 0 | 3 86E-16 |
| TRAPPC6A | 0 062201367 | 0 025313831 | 2 28E-16 | -0 42917166 | 0 | 3 93E-16 |
| FAM20C | 0 326978356 | 0 025311395 | 2 30E-16 | -0 18917929 | 0 | 3 96E-16 |
| DYNC1LI1 | 0 218927871 | 0 025263927 | 2 61E-16 | 0 316445785 | 0 | 4 50E-16 |
| ADGRV1 | 0 093792237 | 0 025259296 | 2 64E-16 | 0 421172898 | 0 | 4 55E-16 |
| EDN1 | 0 011678818 | 0 025246645 | 2 74E-16 | -0 48119671 | 0 | 4 71E-16 |
| RABGAP1L | 0 804515429 | 0 025241647 | 2 77E-16 | 0 030258228 | 1 02E-22 | 4 77E-16 |
| METTL24 | 0 099150978 | 0 025231773 | 2 85E-16 | -0 36863396 | 0 | 4 90E-16 |
| ARHGAP29 | 0 339399593 | 0 025213631 | 2 99E-16 | -0 15891348 | 0 | 5 14E-16 |
| SDR42E2 | 0 015571758 | 0 025208619 | 3 03E-16 | -0 47580306 | 0 | 5 21E-16 |
| TEX41 | 0 101092688 | 0 025205329 | 3 06E-16 | 0 395903242 | 0 | 5 26E-16 |
| WASHC4 | 0 499514572 | 0 025198736 | 3 11E-16 | 0 135998194 | 0 | 5 35E-16 |
| KLRK1 | 0 036749729 | 0 025168409 | 3 38E-16 | 0 463039094 | 0 | 5 80E-16 |

| | | | | | | |
|---|---|---|---|---|---|---|
| GPD2 | 0 434895585 | 0 025167678 | 3 38E-16 | 0 173163121 | 0 | 5 81E-16 |
| LINC01235 | 0 010089281 | 0 025162609 | 3 43E-16 | 0 490821583 | 0 | 5 89E-16 |
| GOLGB1 | 0 555100798 | 0 025148783 | 3 56E-16 | -0 04259669 | 2 13E-43 | 6 11E-16 |
| HSF5 | 0 010841218 | 0 025140565 | 3 64E-16 | -0 48300469 | 0 | 6 25E-16 |
| ID4 | 0 101787516 | 0 025138634 | 3 66E-16 | -0 36059024 | 0 | 6 28E-16 |
| LINC02397 | 0 006662732 | 0 02511856 | 3 86E-16 | 0 494526485 | 0 | 6 62E-16 |
| SLC25A45 | 0 056052617 | 0 025099827 | 4 06E-16 | -0 42355086 | 0 | 6 96E-16 |
| CDC14B | 0 355932687 | 0 025093978 | 4 12E-16 | -0 16195674 | 0 | 7 07E-16 |
| ARMC7 | 0 039700367 | 0 025090155 | 4 16E-16 | -0 45182962 | 0 | 7 14E-16 |
| LASP1 | 0 238249795 | 0 025083144 | 4 24E-16 | -0 23898614 | 0 | 7 28E-16 |
| CCDC18-AS1 | 0 537625402 | 0 025074766 | 4 34E-16 | -0 02714946 | 1 35E-18 | 7 44E-16 |
| RNF168 | 0 273552759 | 0 025073066 | 4 36E-16 | 0 240378 | 0 | 7 47E-16 |
| BAIAP3 | 0 010213017 | 0 025069605 | 4 40E-16 | 0 492134833 | 0 | 7 54E-16 |
| GPS2 | 0 201528621 | 0 02504516 | 4 70E-16 | -0 29455926 | 0 | 8 05E-16 |
| RTN4R | 0 004406922 | 0 025044495 | 4 70E-16 | 0 495672439 | 0 | 8 06E-16 |
| ARL8B | 0 348736936 | 0 025041417 | 4 74E-16 | -0 1757109 | 0 | 8 12E-16 |
| WDR43 | 0 392596752 | 0 025031165 | 4 87E-16 | 0 192862592 | 0 | 8 35E-16 |
| RNF13 | 0 515019703 | 0 025019478 | 5 03E-16 | -0 07247203 | 2 48E-122 | 8 61E-16 |
| IRGM | 0 015590794 | 0 025011074 | 5 14E-16 | 0 486187111 | 0 | 8 80E-16 |
| PROCA1 | 0 032200034 | 0 025002255 | 5 27E-16 | 0 474519402 | 0 | 9 01E-16 |
| LRR1 | 0 058546382 | 0 024982649 | 5 55E-16 | 0 418031233 | 0 | 9 49E-16 |
| KCNJ16 | 0 010660372 | 0 024957318 | 5 93E-16 | -0 48505183 | 0 | 1 02E-15 |
| CENPQ | 0 086111058 | 0 024950867 | 6 04E-16 | 0 397527315 | 0 | 1 03E-15 |
| TRAF3IP3 | 0 012621119 | 0 024934409 | 6 31E-16 | 0 488686965 | 0 | 1 08E-15 |
| RNF213 | 0 5698064 | 0 024927533 | 6 42E-16 | 0 10906703 | 2 22E-275 | 1 10E-15 |
| TBRG1 | 0 209952219 | 0 02486956 | 7 49E-16 | -0 27165465 | 0 | 1 28E-15 |
| TRPV4 | 0 073556567 | 0 02486482 | 7 59E-16 | -0 40726806 | 0 | 1 30E-15 |
| NUBP1 | 0 075069959 | 0 024853584 | 7 82E-16 | 0 424936878 | 0 | 1 34E-15 |
| REXO4 | 0 111448478 | 0 024847809 | 7 94E-16 | 0 40685015 | 0 | 1 36E-15 |
| MMP24OS | 0 248091603 | 0 024841535 | 8 07E-16 | -0 24822226 | 0 | 1 38E-15 |
| TMEM51-AS1 | 0 01598104 | 0 024832155 | 8 28E-16 | 0 486659109 | 0 | 1 41E-15 |
| MOK | 0 284869886 | 0 024830828 | 8 30E-16 | -0 20704508 | 0 | 1 42E-15 |
| H4C3 | 0 080990272 | 0 024802902 | 8 94E-16 | 0 432126951 | 0 | 1 53E-15 |
| F2RL3 | 0 003559803 | 0 024797771 | 9 06E-16 | 0 495998973 | 0 | 1 55E-15 |
| CYYR1-AS1 | 0 008851916 | 0 024783363 | 9 42E-16 | -0 48614275 | 0 | 1 61E-15 |
| CDC42EP4 | 0 159972207 | 0 024781968 | 9 45E-16 | 0 349475449 | 0 | 1 61E-15 |
| DPYS | 0 015619349 | 0 024779794 | 9 51E-16 | -0 47792878 | 0 | 1 62E-15 |
| NAGK | 0 176286383 | 0 024771903 | 9 71E-16 | -0 3168006 | 0 | 1 65E-15 |
| C12orf65 | 0 046420209 | 0 024771671 | 9 71E-16 | 0 441613124 | 0 | 1 65E-15 |
| JAG2 | 0 018484324 | 0 024769901 | 9 76E-16 | 0 483736714 | 0 | 1 66E-15 |
| ILF3-DT | 0 104376463 | 0 0247698 | 9 76E-16 | -0 3757671 | 0 | 1 66E-15 |
| IER5L | 0 123127296 | 0 024736917 | 1 06E-15 | -0 36051268 | 0 | 1 81E-15 |
| CA1 | 0 023690773 | 0 024727986 | 1 09E-15 | -0 46693331 | 0 | 1 85E-15 |
| SH3BP5 | 0 335954008 | 0 02472269 | 1 11E-15 | -0 17480016 | 0 | 1 88E-15 |

| | | | | | | |
|---|---|---|---|---|---|---|
| SEPTIN1 | 0 059117473 | 0 024712765 | 1 13E-15 | -0 43278717 | 0 | 1 93E-15 |
| ATP2B1-AS1 | 0 044649826 | 0 02468888 | 1 21E-15 | 0 458630619 | 0 | 2 06E-15 |
| LINC01358 | 0 083750547 | 0 024686316 | 1 22E-15 | 0 399589873 | 0 | 2 07E-15 |
| PSME4 | 0 543174506 | 0 02468079 | 1 23E-15 | 0 112937941 | 3 03E-295 | 2 10E-15 |
| ACTA2 | 0 179408349 | 0 024674461 | 1 26E-15 | 0 343097851 | 0 | 2 13E-15 |
| OXSM | 0 043031734 | 0 024664686 | 1 29E-15 | -0 44558713 | 0 | 2 19E-15 |
| MAP3K14 | 0 260075003 | 0 024650684 | 1 34E-15 | 0 269912037 | 0 | 2 27E-15 |
| ELMO3 | 0 019959643 | 0 024649383 | 1 34E-15 | 0 479770689 | 0 | 2 28E-15 |
| ADD3-AS1 | 0 084407302 | 0 024648853 | 1 34E-15 | -0 39789761 | 0 | 2 28E-15 |
| SLC10A1 | 0 041670633 | 0 024619032 | 1 45E-15 | 0 460640405 | 0 | 2 47E-15 |
| NPTN | 0 409329729 | 0 024599131 | 1 53E-15 | -0 12901579 | 0 | 2 60E-15 |
| MMP16 | 0 504711504 | 0 024567569 | 1 66E-15 | 0 108857913 | 2 50E-274 | 2 82E-15 |
| P2RY11 | 0 027260094 | 0 024516309 | 1 90E-15 | -0 46451148 | 0 | 3 22E-15 |
| MPZL3 | 0 013068474 | 0 024516014 | 1 90E-15 | 0 489793144 | 0 | 3 23E-15 |
| PARD6G | 0 075945632 | 0 024508998 | 1 94E-15 | -0 40895889 | 0 | 3 29E-15 |
| H2AZ1-DT | 0 033780054 | 0 024498815 | 1 99E-15 | -0 45293629 | 0 | 3 37E-15 |
| KCNG3 | 0 095315147 | 0 024498012 | 1 99E-15 | -0 3800001 | 0 | 3 38E-15 |
| STMN2 | 0 022539072 | 0 024496798 | 2 00E-15 | 0 480967521 | 0 | 3 39E-15 |
| RBM22 | 0 116045763 | 0 024485637 | 2 06E-15 | -0 37287244 | 0 | 3 49E-15 |
| CACUL1 | 0 37568293 | 0 024485292 | 2 06E-15 | 0 202058158 | 0 | 3 49E-15 |
| OLMALINC | 0 141754393 | 0 024485122 | 2 06E-15 | -0 34180104 | 0 | 3 49E-15 |
| NR4A1 | 0 227142068 | 0 024456855 | 2 22E-15 | -0 22938602 | 0 | 3 76E-15 |
| CCNA2 | 0 047724201 | 0 024453273 | 2 24E-15 | 0 430418347 | 0 | 3 79E-15 |
| LINC00298 | 0 001256401 | 0 024452347 | 2 25E-15 | 0 497783147 | 0 | 3 80E-15 |
| SESN2 | 0 08079039 | 0 024429339 | 2 39E-15 | 0 427423363 | 0 | 4 04E-15 |
| ZYX | 0 161818736 | 0 024405789 | 2 54E-15 | -0 31467812 | 0 | 4 29E-15 |
| HEXA | 0 326921247 | 0 024386715 | 2 67E-15 | -0 1885987 | 0 | 4 51E-15 |
| FAM162B | 0 078068188 | 0 024373981 | 2 76E-15 | -0 4042999 | 0 | 4 66E-15 |
| NHSL1 | 0 448297196 | 0 024371335 | 2 78E-15 | -0 11805036 | 0 | 4 69E-15 |
| ANO10 | 0 357988616 | 0 024360862 | 2 85E-15 | 0 227651715 | 0 | 4 82E-15 |
| RAB37 | 0 006538996 | 0 024352696 | 2 91E-15 | -0 48756397 | 0 | 4 92E-15 |
| FNBP1 | 0 468532866 | 0 024348214 | 2 95E-15 | -0 084188 | 1 57E-164 | 4 98E-15 |
| ATP6V1C1 | 0 324027717 | 0 024341004 | 3 00E-15 | -0 2013485 | 0 | 5 07E-15 |
| SARM1 | 0 128210009 | 0 024332382 | 3 07E-15 | -0 36063961 | 0 | 5 18E-15 |
| LINC01001 | 0 044402353 | 0 024329826 | 3 09E-15 | 0 436915875 | 0 | 5 22E-15 |
| CBL | 0 205326379 | 0 024316342 | 3 20E-15 | 0 336072935 | 0 | 5 40E-15 |
| NSUN3 | 0 200595839 | 0 024314006 | 3 22E-15 | -0 29483955 | 0 | 5 43E-15 |
| LINC02506 | 0 030296396 | 0 024305294 | 3 29E-15 | 0 467427341 | 0 | 5 56E-15 |
| ADM5 | 0 011317127 | 0 024300261 | 3 34E-15 | 0 4878743 | 0 | 5 63E-15 |
| BCAS4 | 0 064961642 | 0 024289486 | 3 43E-15 | 0 435700835 | 0 | 5 79E-15 |
| IFI44 | 0 273429023 | 0 024278285 | 3 53E-15 | 0 254180708 | 0 | 5 96E-15 |
| PROSER2 | 0 058298909 | 0 024264058 | 3 67E-15 | -0 41823804 | 0 | 6 18E-15 |
| FAAH2 | 0 072880775 | 0 024221711 | 4 09E-15 | 0 448220281 | 0 | 6 89E-15 |
| HAS2 | 0 157840132 | 0 02419433 | 4 39E-15 | 0 356545462 | 0 | 7 39E-15 |

| | | | | | | |
|---|---|---|---|---|---|---|
| RALGPS1 | 0 163769964 | 0 024177293 | 4 59E-15 | -0 31615447 | 0 | 7 72E-15 |
| RNMT | 0 430850355 | 0 024176327 | 4 60E-15 | -0 12873353 | 0 | 7 74E-15 |
| UBA6 | 0 41069083 | 0 024161298 | 4 78E-15 | 0 176784895 | 0 | 8 04E-15 |
| LPIN1 | 0 296301232 | 0 024159532 | 4 80E-15 | -0 21381492 | 0 | 8 08E-15 |
| RLF | 0 441196627 | 0 024119914 | 5 32E-15 | -0 12233515 | 0 | 8 94E-15 |
| MRPL33 | 0 283156612 | 0 024119651 | 5 32E-15 | -0 22213063 | 0 | 8 95E-15 |
| KL | 0 030734233 | 0 024107941 | 5 49E-15 | -0 45401556 | 0 | 9 22E-15 |
| SYNPR | 0 015324285 | 0 024103131 | 5 56E-15 | 0 488206003 | 0 | 9 33E-15 |
| ADAM9 | 0 379480688 | 0 024084205 | 5 83E-15 | 0 202261637 | 0 | 9 79E-15 |
| UXS1 | 0 220260418 | 0 024060299 | 6 20E-15 | 0 322127439 | 0 | 1 04E-14 |
| PIK3CB | 0 312910472 | 0 02404476 | 6 45E-15 | -0 19612562 | 0 | 1 08E-14 |
| MYO18A | 0 128609773 | 0 024039531 | 6 54E-15 | -0 35012207 | 0 | 1 10E-14 |
| USP36 | 0 342845177 | 0 024025266 | 6 79E-15 | -0 19206069 | 0 | 1 14E-14 |
| PDGFD | 0 184681426 | 0 02400296 | 7 19E-15 | -0 29077147 | 0 | 1 20E-14 |
| PMS2P4 | 0 269707411 | 0 02400245 | 7 19E-15 | -0 20147301 | 0 | 1 21E-14 |
| SLC35D1 | 0 220841027 | 0 023979764 | 7 63E-15 | 0 309832823 | 0 | 1 28E-14 |
| CD8A | 0 010470008 | 0 023970484 | 7 81E-15 | 0 490309664 | 0 | 1 31E-14 |
| SCARB1 | 0 17120367 | 0 023963582 | 7 95E-15 | -0 31754893 | 0 | 1 33E-14 |
| CELSR1 | 0 052331005 | 0 023963191 | 7 96E-15 | 0 44348374 | 0 | 1 33E-14 |
| LYVE1 | 0 021882317 | 0 023961445 | 7 99E-15 | 0 478607288 | 0 | 1 34E-14 |
| UBE2A | 0 191353677 | 0 023951221 | 8 20E-15 | -0 29320191 | 0 | 1 37E-14 |
| LPP | 0 88582932 | 0 023940474 | 8 43E-15 | 0 011831412 | 0 000125546 | 1 41E-14 |
| HEATR5A | 0 356142088 | 0 023911379 | 9 08E-15 | -0 15514163 | 0 | 1 52E-14 |
| TCF7 | 0 096999867 | 0 023897522 | 9 41E-15 | 0 412039605 | 0 | 1 57E-14 |
| NDC1 | 0 144705031 | 0 023885229 | 9 71E-15 | 0 339341611 | 0 | 1 62E-14 |
| UBE2E3 | 0 536054901 | 0 023877774 | 9 90E-15 | 0 088592458 | 4 81E-182 | 1 65E-14 |
| LRRC32 | 0 023081609 | 0 023873766 | 1 00E-14 | -0 4661065 | 0 | 1 67E-14 |
| MYO16 | 0 031667016 | 0 023866802 | 1 02E-14 | 0 480414554 | 0 | 1 70E-14 |
| OOEP | 0 025918029 | 0 023853442 | 1 05E-14 | 0 474833218 | 0 | 1 76E-14 |
| KNL1 | 0 088281205 | 0 023834399 | 1 11E-14 | 0 35643425 | 0 | 1 84E-14 |
| SLC38A4 | 0 067959871 | 0 023801279 | 1 20E-14 | 0 432380117 | 0 | 2 01E-14 |
| PDLIM3 | 0 118625193 | 0 023795148 | 1 22E-14 | -0 36692612 | 0 | 2 04E-14 |
| SOX7 | 0 009546744 | 0 023782914 | 1 26E-14 | 0 492967582 | 0 | 2 10E-14 |
| FZD6 | 0 257048219 | 0 0237802 | 1 27E-14 | 0 248426064 | 0 | 2 12E-14 |
| SLC19A2 | 0 225771449 | 0 02377867 | 1 27E-14 | -0 23881844 | 0 | 2 12E-14 |
| PKP2 | 0 078391807 | 0 023778559 | 1 27E-14 | -0 38495963 | 0 | 2 12E-14 |
| MMP24 | 0 173192972 | 0 023766259 | 1 31E-14 | -0 33231201 | 0 | 2 19E-14 |
| HMOX2 | 0 144038758 | 0 023751042 | 1 37E-14 | -0 33018945 | 0 | 2 28E-14 |
| KLRB1 | 0 026298757 | 0 023749596 | 1 37E-14 | 0 467607252 | 0 | 2 28E-14 |
| ARHGAP12 | 0 440473244 | 0 023746847 | 1 38E-14 | -0 09203784 | 2 25E-196 | 2 30E-14 |
| FOXC1 | 0 053339933 | 0 023744344 | 1 39E-14 | 0 45733517 | 0 | 2 31E-14 |
| TLR4 | 0 062020521 | 0 023724489 | 1 46E-14 | -0 42091924 | 0 | 2 43E-14 |
| ZNF106 | 0 439778417 | 0 023722598 | 1 47E-14 | -0 10195754 | 9 71E-241 | 2 44E-14 |
| PCNA | 0 082503665 | 0 023686244 | 1 61E-14 | 0 39587723 | 0 | 2 68E-14 |

| | | | | | | |
|---|---|---|---|---|---|---|
| UNC119 | 0 042860406 | 0 02368456 | 1 62E-14 | 0 460636695 | 0 | 2 69E-14 |
| CES3 | 0 006282005 | 0 02367512 | 1 66E-14 | 0 493157737 | 0 | 2 75E-14 |
| RNF166 | 0 055015134 | 0 023668282 | 1 68E-14 | 0 453903138 | 0 | 2 80E-14 |
| PABPC4 | 0 288382098 | 0 023665185 | 1 70E-14 | 0 260784344 | 0 | 2 82E-14 |
| GPM6A | 0 050703394 | 0 02364816 | 1 77E-14 | 0 465114115 | 0 | 2 95E-14 |
| GXYLT2 | 0 281091165 | 0 023639415 | 1 81E-14 | -0 24147887 | 0 | 3 01E-14 |
| PTPN1 | 0 352601321 | 0 023636362 | 1 83E-14 | 0 22095375 | 0 | 3 03E-14 |
| EPHA1-AS1 | 0 031219661 | 0 023627159 | 1 87E-14 | -0 4590509 | 0 | 3 10E-14 |
| NCK1 | 0 290733091 | 0 023611951 | 1 94E-14 | -0 21956825 | 0 | 3 22E-14 |
| CDC25C | 0 060630866 | 0 023610183 | 1 95E-14 | 0 388215018 | 0 | 3 24E-14 |
| PCP4 | 0 027431421 | 0 023586636 | 2 07E-14 | -0 46376843 | 0 | 3 43E-14 |
| PCSK4 | 0 015847785 | 0 023583745 | 2 09E-14 | 0 487973276 | 0 | 3 46E-14 |
| PISD | 0 160705107 | 0 023580353 | 2 10E-14 | 0 355095039 | 0 | 3 49E-14 |
| IRAK1 | 0 09929375 | 0 023572598 | 2 15E-14 | 0 410751104 | 0 | 3 56E-14 |
| ESAM | 0 017522986 | 0 023566058 | 2 18E-14 | -0 47658872 | 0 | 3 61E-14 |
| HERC2P3 | 0 084378748 | 0 023564825 | 2 19E-14 | -0 38178688 | 0 | 3 63E-14 |
| CEP170 | 0 533208962 | 0 023559682 | 2 22E-14 | 0 134325567 | 0 | 3 67E-14 |
| STXBP6 | 0 274095296 | 0 023536975 | 2 35E-14 | -0 22779226 | 0 | 3 89E-14 |
| NECAP1 | 0 151548609 | 0 023510904 | 2 51E-14 | 0 363766589 | 0 | 4 15E-14 |
| ITGAE | 0 1163789 | 0 023505097 | 2 54E-14 | 0 395266594 | 0 | 4 21E-14 |
| SCGB2B2 | 0 182082961 | 0 023496426 | 2 60E-14 | 0 31659136 | 0 | 4 30E-14 |
| PCDH11Y | 0 018541433 | 0 023488468 | 2 65E-14 | 0 484413278 | 0 | 4 38E-14 |
| GMNN | 0 088776151 | 0 023474569 | 2 74E-14 | 0 37437481 | 0 | 4 54E-14 |
| CAPZA2 | 0 422855076 | 0 023468962 | 2 78E-14 | 0 185662483 | 0 | 4 60E-14 |
| EIF1B | 0 151358246 | 0 023434628 | 3 03E-14 | -0 34605783 | 0 | 5 01E-14 |
| CENPC | 0 434181721 | 0 023423648 | 3 12E-14 | 0 18707831 | 0 | 5 15E-14 |
| ZNF41 | 0 113723325 | 0 02340415 | 3 27E-14 | 0 406859603 | 0 | 5 41E-14 |
| ASCC2 | 0 178190021 | 0 023380227 | 3 48E-14 | 0 334682008 | 0 | 5 74E-14 |
| TPD52L1 | 0 164417201 | 0 023379832 | 3 48E-14 | -0 291915 | 0 | 5 75E-14 |
| PIR | 0 153614056 | 0 023363069 | 3 63E-14 | -0 33431166 | 0 | 5 99E-14 |
| STX8 | 0 510746036 | 0 023359395 | 3 66E-14 | -0 07345167 | 1 29E-125 | 6 04E-14 |
| RGMA | 0 133578268 | 0 023357364 | 3 68E-14 | -0 35106289 | 0 | 6 07E-14 |
| NETO1 | 0 041956178 | 0 02335091 | 3 74E-14 | 0 472944662 | 0 | 6 17E-14 |
| RGL4 | 0 01671394 | 0 023342965 | 3 81E-14 | -0 47757919 | 0 | 6 29E-14 |
| MTCP1 | 0 112428852 | 0 0233393 | 3 85E-14 | 0 396973385 | 0 | 6 35E-14 |
| MMP2 | 0 322914089 | 0 023331803 | 3 92E-14 | -0 19944022 | 0 | 6 47E-14 |
| MEDAG | 0 045173326 | 0 023300557 | 4 24E-14 | -0 43716327 | 0 | 6 99E-14 |
| HAUS8 | 0 050370258 | 0 023294566 | 4 30E-14 | 0 425304891 | 0 | 7 10E-14 |
| PCDH7 | 0 303925301 | 0 023292814 | 4 32E-14 | 0 255359612 | 0 | 7 13E-14 |
| UNC5CL | 0 006748396 | 0 023280816 | 4 45E-14 | -0 48778175 | 0 | 7 34E-14 |
| DNAI2 | 0 006558032 | 0 023274982 | 4 52E-14 | -0 48830098 | 0 | 7 45E-14 |
| ID2 | 0 095162856 | 0 023262172 | 4 67E-14 | 0 391593954 | 0 | 7 69E-14 |
| SRPX2 | 0 383916164 | 0 023260994 | 4 68E-14 | -0 11715871 | 0 | 7 71E-14 |
| SAR1A | 0 280662847 | 0 023259055 | 4 70E-14 | -0 23035639 | 0 | 7 74E-14 |

| | | | | | | |
|---|---|---|---|---|---|---|
| SUSD5 | 0 037396966 | 0 02325229 | 4 78E-14 | 0 468935954 | 0 | 7 87E-14 |
| MKNK1 | 0 202499476 | 0 023239971 | 4 93E-14 | -0 28093185 | 0 | 8 11E-14 |
| KANSL1L | 0 49910529 | 0 023227824 | 5 08E-14 | -0 06714251 | 3 05E-105 | 8 36E-14 |
| ALOX5 | 0 021691953 | 0 023219172 | 5 19E-14 | 0 481802279 | 0 | 8 54E-14 |
| SNW1 | 0 276274961 | 0 023174151 | 5 80E-14 | 0 272565421 | 0 | 9 55E-14 |
| RNF130 | 0 510612781 | 0 023157377 | 6 05E-14 | 0 162309961 | 0 | 9 95E-14 |
| GCNA | 0 086434677 | 0 023151251 | 6 14E-14 | -0 39802139 | 0 | 1 01E-13 |
| KCNJ15 | 0 032038225 | 0 023147221 | 6 20E-14 | 0 475222586 | 0 | 1 02E-13 |
| SLC39A4 | 0 015428985 | 0 023122504 | 6 60E-14 | 0 487630933 | 0 | 1 08E-13 |
| SCNM1 | 0 10066437 | 0 023113381 | 6 75E-14 | 0 402119492 | 0 | 1 11E-13 |
| CUBN | 0 122461023 | 0 023110939 | 6 79E-14 | -0 36089205 | 0 | 1 11E-13 |
| OXSR1 | 0 292075156 | 0 023107336 | 6 85E-14 | 0 256131574 | 0 | 1 12E-13 |
| ZG16B | 0 002922084 | 0 023087621 | 7 19E-14 | 0 496373432 | 0 | 1 18E-13 |
| CCDC137 | 0 115227199 | 0 02307394 | 7 44E-14 | 0 386304242 | 0 | 1 22E-13 |
| ACE | 0 020997126 | 0 023055759 | 7 78E-14 | -0 46974981 | 0 | 1 28E-13 |
| SLC25A33 | 0 160305344 | 0 023049468 | 7 90E-14 | -0 3235626 | 0 | 1 30E-13 |
| TTC9 | 0 010003617 | 0 02298485 | 9 26E-14 | 0 493248753 | 0 | 1 52E-13 |
| LLPH | 0 145609259 | 0 022979367 | 9 39E-14 | 0 361792568 | 0 | 1 54E-13 |
| OMA1 | 0 060545202 | 0 022956979 | 9 92E-14 | -0 42223242 | 0 | 1 63E-13 |
| PRKCQ-AS1 | 0 009166016 | 0 022944158 | 1 02E-13 | 0 490092223 | 0 | 1 68E-13 |
| EML6 | 0 104890446 | 0 022931085 | 1 06E-13 | 0 373749153 | 0 | 1 73E-13 |
| XAF1 | 0 254116617 | 0 022928986 | 1 06E-13 | -0 25232474 | 0 | 1 74E-13 |
| DSCR9 | 0 011926291 | 0 02288882 | 1 17E-13 | -0 48279493 | 0 | 1 92E-13 |
| LINC01924 | 0 00641526 | 0 022873987 | 1 22E-13 | -0 48802713 | 0 | 1 99E-13 |
| GCC2 | 0 537349375 | 0 022868907 | 1 23E-13 | -0 08085793 | 6 89E-152 | 2 01E-13 |
| ARHGAP45 | 0 015781158 | 0 022848575 | 1 29E-13 | 0 486249537 | 0 | 2 11E-13 |
| BTG3 | 0 218537625 | 0 022830615 | 1 35E-13 | 0 320892106 | 0 | 2 21E-13 |
| LINC01483 | 0 011240982 | 0 02282186 | 1 38E-13 | -0 48303615 | 0 | 2 26E-13 |
| NEIL3 | 0 04940892 | 0 022805045 | 1 44E-13 | 0 401795972 | 0 | 2 35E-13 |
| SYK | 0 045230435 | 0 022798526 | 1 46E-13 | 0 459655466 | 0 | 2 39E-13 |
| KCNE3 | 0 008309379 | 0 022791168 | 1 49E-13 | 0 492085526 | 0 | 2 43E-13 |
| PARVB | 0 158154233 | 0 022781478 | 1 52E-13 | 0 346063305 | 0 | 2 49E-13 |
| CCK | 0 00310293 | 0 022778252 | 1 54E-13 | 0 496138734 | 0 | 2 51E-13 |
| MSRA | 0 223429975 | 0 022768489 | 1 57E-13 | -0 26583659 | 0 | 2 57E-13 |
| THOC3 | 0 028630713 | 0 022745023 | 1 66E-13 | 0 470670454 | 0 | 2 72E-13 |
| CHRNA1 | 0 040014468 | 0 022744229 | 1 67E-13 | 0 447310578 | 0 | 2 72E-13 |
| LMNTD2 | 0 007138642 | 0 022738041 | 1 69E-13 | 0 493527318 | 0 | 2 76E-13 |
| DSEL-AS1 | 0 02806914 | 0 022737855 | 1 69E-13 | -0 46011417 | 0 | 2 76E-13 |
| ALDH1L1 | 0 025822847 | 0 022735791 | 1 70E-13 | -0 46355497 | 0 | 2 78E-13 |
| ZBTB24 | 0 081770764 | 0 022704161 | 1 84E-13 | -0 40201484 | 0 | 3 00E-13 |
| KIAA1958 | 0 193885515 | 0 022686098 | 1 92E-13 | 0 324335604 | 0 | 3 13E-13 |
| AKAP5 | 0 01750395 | 0 022672573 | 1 99E-13 | 0 483399352 | 0 | 3 23E-13 |
| ACER2 | 0 04820011 | 0 022672545 | 1 99E-13 | 0 456085029 | 0 | 3 23E-13 |
| PDE6A | 0 013753783 | 0 022660971 | 2 04E-13 | -0 4804046 | 0 | 3 32E-13 |

| | | | | | | |
|---|---|---|---|---|---|---|
| NUDT19 | 0 079324589 | 0 022656225 | 2 07E-13 | 0 423681827 | 0 | 3 36E-13 |
| PDZK1 | 0 110077859 | 0 022654462 | 2 07E-13 | 0 376665379 | 0 | 3 38E-13 |
| PIK3R3 | 0 166520721 | 0 022630668 | 2 20E-13 | 0 294381265 | 0 | 3 57E-13 |
| ATP10D | 0 394424245 | 0 022626664 | 2 22E-13 | -0 14862683 | 0 | 3 61E-13 |
| PDCD4-AS1 | 0 01014639 | 0 022611345 | 2 30E-13 | 0 49223397 | 0 | 3 74E-13 |
| BAZ1A | 0 3262169 | 0 022602646 | 2 35E-13 | 0 225374728 | 0 | 3 82E-13 |
| SNAPC1 | 0 160943062 | 0 02257214 | 2 53E-13 | -0 32453354 | 0 | 4 11E-13 |
| HAL | 0 006215378 | 0 022565907 | 2 57E-13 | 0 493298831 | 0 | 4 18E-13 |
| TNFAIP3 | 0 084826103 | 0 022556477 | 2 63E-13 | -0 40321083 | 0 | 4 27E-13 |
| DLG1 | 0 740971997 | 0 022556094 | 2 63E-13 | 0 043428092 | 4 84E-45 | 4 27E-13 |
| LOXHD1 | 0 002779311 | 0 022551748 | 2 66E-13 | 0 496391515 | 0 | 4 32E-13 |
| RUNDC3B | 0 034265481 | 0 022530489 | 2 80E-13 | 0 477075777 | 0 | 4 54E-13 |
| HORMAD1 | 0 067303116 | 0 02250905 | 2 95E-13 | -0 41533615 | 0 | 4 78E-13 |
| ATP2B2 | 0 015048257 | 0 022492709 | 3 07E-13 | 0 487286894 | 0 | 4 97E-13 |
| ARMH1 | 0 072185947 | 0 022488772 | 3 10E-13 | -0 41199679 | 0 | 5 02E-13 |
| PTCH2 | 0 317602939 | 0 022488314 | 3 10E-13 | -0 17495309 | 0 | 5 02E-13 |
| LEPR | 0 222525747 | 0 022473143 | 3 21E-13 | -0 2526518 | 0 | 5 21E-13 |
| CLASP1-AS1 | 0 035283928 | 0 022458755 | 3 33E-13 | -0 45127413 | 0 | 5 39E-13 |
| SYNC | 0 219137271 | 0 022447534 | 3 42E-13 | 0 301051001 | 0 | 5 54E-13 |
| CTNND2 | 0 11222897 | 0 022441616 | 3 47E-13 | 0 400632808 | 0 | 5 61E-13 |
| SDE2 | 0 066817689 | 0 022437743 | 3 50E-13 | -0 4199437 | 0 | 5 67E-13 |
| LINC01374 | 0 024214273 | 0 022428864 | 3 57E-13 | -0 46809725 | 0 | 5 79E-13 |
| ZNF230 | 0 029230359 | 0 022422166 | 3 63E-13 | 0 471484367 | 0 | 5 88E-13 |
| MPZL2 | 0 017285032 | 0 022415347 | 3 69E-13 | 0 488496773 | 0 | 5 98E-13 |
| CYB5B | 0 22729436 | 0 022399153 | 3 84E-13 | 0 311091538 | 0 | 6 21E-13 |
| ATP6V0D1 | 0 119329539 | 0 022391525 | 3 91E-13 | -0 36681934 | 0 | 6 33E-13 |
| KBTBD11 | 0 014077402 | 0 022378162 | 4 04E-13 | -0 47795522 | 0 | 6 53E-13 |
| GBGT1 | 0 024709219 | 0 022366465 | 4 15E-13 | -0 46624985 | 0 | 6 71E-13 |
| ETV3 | 0 079724353 | 0 022365522 | 4 16E-13 | -0 40964253 | 0 | 6 73E-13 |
| RAPGEF2 | 0 62090004 | 0 022357708 | 4 24E-13 | -0 01156275 | 0 00017829 | 6 85E-13 |
| GRM8 | 0 051169785 | 0 022355043 | 4 27E-13 | 0 458860276 | 0 | 6 90E-13 |
| OSBPL10 | 0 529896632 | 0 022352972 | 4 29E-13 | -0 08845597 | 1 73E-181 | 6 93E-13 |
| WIPF2 | 0 309350669 | 0 022344398 | 4 38E-13 | -0 20579024 | 0 | 7 07E-13 |
| NRIP3 | 0 049180484 | 0 022339936 | 4 42E-13 | -0 4359319 | 0 | 7 15E-13 |
| CARMIL2 | 0 004007158 | 0 022312334 | 4 73E-13 | 0 496020959 | 0 | 7 63E-13 |
| ABHD3 | 0 173478517 | 0 022309273 | 4 76E-13 | 0 335674339 | 0 | 7 69E-13 |
| KLHDC7B | 0 00514934 | 0 022300081 | 4 87E-13 | -0 48995064 | 0 | 7 85E-13 |
| NDST2 | 0 056081171 | 0 022259401 | 5 36E-13 | -0 42959169 | 0 | 8 65E-13 |
| PLA2G15 | 0 05592888 | 0 022237925 | 5 64E-13 | 0 453891486 | 0 | 9 11E-13 |
| HAVCR2 | 0 06738878 | 0 022188913 | 6 34E-13 | 0 444098162 | 0 | 1 02E-12 |
| MAOB | 0 051464849 | 0 022182403 | 6 44E-13 | -0 43004347 | 0 | 1 04E-12 |
| NUP210 | 0 052492814 | 0 022168092 | 6 66E-13 | 0 458185737 | 0 | 1 07E-12 |
| SMC2 | 0 137033371 | 0 022129529 | 7 30E-13 | 0 315479462 | 0 | 1 18E-12 |
| TMEM163 | 0 150473054 | 0 022117025 | 7 52E-13 | -0 33599653 | 0 | 1 21E-12 |

| | | | | | | |
|---|---|---|---|---|---|---|
| WEE1 | 0 169166778 | 0 022110637 | 7 63E-13 | 0 314506139 | 0 | 1 23E-12 |
| C16orf74 | 0 046524909 | 0 02209554 | 7 91E-13 | 0 436415476 | 0 | 1 27E-12 |
| ELOVL2 | 0 1740972 | 0 02209508 | 7 92E-13 | -0 29689271 | 0 | 1 27E-12 |
| IQCH | 0 190744513 | 0 022092376 | 7 97E-13 | 0 342518305 | 0 | 1 28E-12 |
| TCAP | 0 015238621 | 0 022090616 | 8 00E-13 | 0 486199775 | 0 | 1 29E-12 |
| PURB | 0 217785688 | 0 022083481 | 8 14E-13 | -0 27560501 | 0 | 1 31E-12 |
| C11orf80 | 0 304020483 | 0 022073611 | 8 33E-13 | 0 243693112 | 0 | 1 34E-12 |
| PIGK | 0 166063848 | 0 022065148 | 8 50E-13 | 0 366847223 | 0 | 1 37E-12 |
| PPARA | 0 216795797 | 0 02205798 | 8 65E-13 | -0 27664746 | 0 | 1 39E-12 |
| PRMT5-AS1 | 0 015371876 | 0 022049015 | 8 83E-13 | 0 485210213 | 0 | 1 42E-12 |
| PPP1R3A | 0 00504464 | 0 022031199 | 9 21E-13 | 0 495155949 | 0 | 1 48E-12 |
| CCDC167 | 0 051607622 | 0 022023817 | 9 37E-13 | 0 45820528 | 0 | 1 51E-12 |
| FZD5 | 0 016199958 | 0 022022395 | 9 40E-13 | 0 483764595 | 0 | 1 51E-12 |
| SHH | 0 007576479 | 0 022016618 | 9 53E-13 | -0 48840533 | 0 | 1 53E-12 |
| MFAP5 | 0 180893187 | 0 02201614 | 9 54E-13 | -0 31624016 | 0 | 1 53E-12 |
| CRNKL1 | 0 095714911 | 0 022001713 | 9 87E-13 | 0 424939452 | 0 | 1 59E-12 |
| IFNGR1 | 0 266204717 | 0 021979599 | 1 04E-12 | -0 20683039 | 0 | 1 67E-12 |
| PACS1 | 0 431706992 | 0 021957707 | 1 10E-12 | 0 142349366 | 0 | 1 76E-12 |
| LINC02642 | 0 003493175 | 0 021954845 | 1 10E-12 | 0 495378696 | 0 | 1 77E-12 |
| C17orf99 | 0 007157678 | 0 021942928 | 1 13E-12 | 0 492640582 | 0 | 1 82E-12 |
| LAT | 0 048561802 | 0 021937389 | 1 15E-12 | 0 455944155 | 0 | 1 84E-12 |
| ENTPD3-AS1 | 0 141383183 | 0 021936965 | 1 15E-12 | -0 34494931 | 0 | 1 84E-12 |
| C5 | 0 165045402 | 0 021926962 | 1 18E-12 | 0 350704971 | 0 | 1 89E-12 |
| HCLS1 | 0 031552797 | 0 021922817 | 1 19E-12 | 0 472807384 | 0 | 1 91E-12 |
| CXCL13 | 0 016266585 | 0 021920162 | 1 20E-12 | -0 47723941 | 0 | 1 92E-12 |
| LMNB1 | 0 102320535 | 0 021919343 | 1 20E-12 | 0 322192716 | 0 | 1 92E-12 |
| SLA2 | 0 011164836 | 0 021907241 | 1 23E-12 | 0 490076363 | 0 | 1 98E-12 |
| TXNIP | 0 394738345 | 0 021906391 | 1 24E-12 | -0 16290395 | 0 | 1 98E-12 |
| TGFBR3 | 0 445632103 | 0 021905274 | 1 24E-12 | -0 07041985 | 1 35E-115 | 1 98E-12 |
| FAM13B | 0 506957796 | 0 021902448 | 1 25E-12 | 0 145443384 | 0 | 2 00E-12 |
| MTMR10 | 0 213321658 | 0 021886479 | 1 29E-12 | -0 26808511 | 0 | 2 07E-12 |
| SECISBP2L | 0 461793988 | 0 021874261 | 1 33E-12 | -0 09350665 | 1 18E-202 | 2 13E-12 |
| SERTAD3 | 0 041794369 | 0 021862022 | 1 37E-12 | -0 44564438 | 0 | 2 19E-12 |
| RBMS1 | 0 778102454 | 0 021861691 | 1 37E-12 | -0 03621621 | 7 71E-32 | 2 19E-12 |
| RRM1 | 0 145437932 | 0 021842726 | 1 43E-12 | 0 301189489 | 0 | 2 29E-12 |
| ARHGAP32 | 0 546001409 | 0 021831742 | 1 47E-12 | -0 05621133 | 2 77E-74 | 2 35E-12 |
| BCL6 | 0 237440749 | 0 021811314 | 1 54E-12 | -0 23824033 | 0 | 2 47E-12 |
| LINC01151 | 0 049228075 | 0 021809021 | 1 55E-12 | -0 43733494 | 0 | 2 48E-12 |
| SPINT1 | 0 007605033 | 0 021796791 | 1 60E-12 | -0 48685804 | 0 | 2 55E-12 |
| PARP1 | 0 176134092 | 0 021790385 | 1 62E-12 | 0 357556095 | 0 | 2 59E-12 |
| MYO18B | 0 008851916 | 0 021786045 | 1 64E-12 | 0 490407303 | 0 | 2 62E-12 |
| EDEM1 | 0 163988883 | 0 021775796 | 1 68E-12 | -0 32656243 | 0 | 2 68E-12 |
| HEY2 | 0 020597362 | 0 02176435 | 1 72E-12 | 0 482581349 | 0 | 2 75E-12 |
| REC8 | 0 050731949 | 0 021757027 | 1 75E-12 | 0 450143187 | 0 | 2 80E-12 |

| | | | | | | |
|---|---|---|---|---|---|---|
| RC3H1 | 0 585835031 | 0 021753893 | 1 76E-12 | -0 04961459 | 2 93E-58 | 2 82E-12 |
| SPOCK1 | 0 267070872 | 0 021736035 | 1 84E-12 | -0 24424276 | 0 | 2 94E-12 |
| GRK2 | 0 101340161 | 0 0217314 | 1 86E-12 | 0 406707858 | 0 | 2 97E-12 |
| SCUBE1 | 0 011574118 | 0 021719671 | 1 91E-12 | -0 48038286 | 0 | 3 05E-12 |
| HBZ | 0 004778131 | 0 021714772 | 1 93E-12 | -0 49045615 | 0 | 3 08E-12 |
| HINT3 | 0 072576193 | 0 021711519 | 1 95E-12 | -0 40737818 | 0 | 3 11E-12 |
| CARNS1 | 0 003759685 | 0 021709739 | 1 96E-12 | 0 495821542 | 0 | 3 12E-12 |
| DHDH | 0 005558623 | 0 021708194 | 1 96E-12 | 0 494956763 | 0 | 3 13E-12 |
| PCLAF | 0 075298395 | 0 02170731 | 1 97E-12 | 0 366186509 | 0 | 3 13E-12 |
| NOVA2 | 0 008157088 | 0 021702246 | 1 99E-12 | -0 48570056 | 0 | 3 17E-12 |
| WNT5A-AS1 | 0 002179665 | 0 021700701 | 2 00E-12 | 0 496494978 | 0 | 3 18E-12 |
| SSUH2 | 0 008223716 | 0 021696644 | 2 02E-12 | 0 493869244 | 0 | 3 21E-12 |
| FUT9 | 0 013411129 | 0 021678911 | 2 10E-12 | 0 487677301 | 0 | 3 35E-12 |
| NCOA7 | 0 263872761 | 0 02165784 | 2 21E-12 | -0 21579879 | 0 | 3 51E-12 |
| PRDX6-AS1 | 0 134901296 | 0 021654236 | 2 23E-12 | -0 33495854 | 0 | 3 54E-12 |
| LINC01320 | 0 040595077 | 0 021651612 | 2 24E-12 | 0 464445349 | 0 | 3 56E-12 |
| PRR22 | 0 015305248 | 0 021646684 | 2 26E-12 | 0 484291027 | 0 | 3 60E-12 |
| LINC01446 | 0 025299347 | 0 021645863 | 2 27E-12 | 0 482326808 | 0 | 3 61E-12 |
| DCAF7 | 0 385715102 | 0 021642715 | 2 29E-12 | 0 17199044 | 0 | 3 63E-12 |
| ABHD12 | 0 250080905 | 0 021640225 | 2 30E-12 | 0 297385181 | 0 | 3 65E-12 |
| SSR1 | 0 277169671 | 0 021625719 | 2 38E-12 | -0 22514537 | 0 | 3 78E-12 |
| TMUB1 | 0 061201957 | 0 021618759 | 2 42E-12 | 0 445654176 | 0 | 3 84E-12 |
| GPR137B | 0 227256287 | 0 021618217 | 2 42E-12 | -0 23669828 | 0 | 3 84E-12 |
| TTC13 | 0 176048429 | 0 021611544 | 2 46E-12 | -0 31530223 | 0 | 3 90E-12 |
| HS3ST4 | 0 012973292 | 0 021605553 | 2 49E-12 | -0 48081065 | 0 | 3 96E-12 |
| PIP5K1B | 0 056052617 | 0 021600178 | 2 52E-12 | -0 42791616 | 0 | 4 00E-12 |
| GRIPAP1 | 0 174696846 | 0 021573467 | 2 68E-12 | -0 31497181 | 0 | 4 26E-12 |
| LINC01099 | 0 005882241 | 0 02156697 | 2 72E-12 | 0 494352018 | 0 | 4 32E-12 |
| LINC02752 | 0 006072605 | 0 021541984 | 2 89E-12 | -0 48951097 | 0 | 4 58E-12 |
| CBR3 | 0 09866555 | 0 021515581 | 3 07E-12 | 0 408496344 | 0 | 4 86E-12 |
| ZNF100 | 0 17109897 | 0 021515492 | 3 07E-12 | 0 300833934 | 0 | 4 86E-12 |
| RALA | 0 392967962 | 0 021512651 | 3 09E-12 | -0 15202918 | 0 | 4 90E-12 |
| CHKA | 0 260731758 | 0 02150937 | 3 11E-12 | 0 279815237 | 0 | 4 93E-12 |
| ETS2 | 0 191658259 | 0 021503857 | 3 15E-12 | 0 287194958 | 0 | 4 99E-12 |
| LINC01480 | 0 006529478 | 0 021503142 | 3 16E-12 | 0 492965895 | 0 | 5 00E-12 |
| PUS7L | 0 243875045 | 0 021499812 | 3 18E-12 | 0 275941435 | 0 | 5 04E-12 |
| AGTPBP1 | 0 323351926 | 0 021496897 | 3 20E-12 | 0 206836351 | 0 | 5 07E-12 |
| CERNA1 | 0 012649674 | 0 021480828 | 3 32E-12 | 0 488127893 | 0 | 5 26E-12 |
| GGNBP2 | 0 376511013 | 0 021468329 | 3 42E-12 | 0 227401391 | 0 | 5 41E-12 |
| BBOX1-AS1 | 0 088138433 | 0 021463958 | 3 45E-12 | 0 40452525 | 0 | 5 47E-12 |
| RAB3C | 0 0232339 | 0 021458993 | 3 49E-12 | -0 46837863 | 0 | 5 53E-12 |
| MEGF11 | 0 041232796 | 0 021435896 | 3 68E-12 | -0 45009357 | 0 | 5 83E-12 |
| WDFY1 | 0 307989568 | 0 02142628 | 3 77E-12 | -0 21116352 | 0 | 5 96E-12 |
| UAP1 | 0 238087986 | 0 021417223 | 3 85E-12 | 0 289456539 | 0 | 6 08E-12 |

| | | | | | | |
|---|---|---|---|---|---|---|
| ELAVL3 | 0 035950201 | 0 021415048 | 3 87E-12 | -0 44934684 | 0 | 6 11E-12 |
| CBLN3 | 0 012030991 | 0 021374187 | 4 24E-12 | -0 48091382 | 0 | 6 71E-12 |
| NRIR | 0 008623479 | 0 021331785 | 4 68E-12 | 0 491081012 | 0 | 7 39E-12 |
| RASGRP4 | 0 004968495 | 0 021324235 | 4 76E-12 | -0 49047022 | 0 | 7 52E-12 |
| ABCD1 | 0 038377339 | 0 021303755 | 4 99E-12 | -0 44790555 | 0 | 7 88E-12 |
| UPRT | 0 105090328 | 0 021290001 | 5 15E-12 | -0 37633587 | 0 | 8 13E-12 |
| RASA1 | 0 619519903 | 0 021285313 | 5 20E-12 | 0 061308383 | 4 92E-88 | 8 21E-12 |
| TIMELESS | 0 065142487 | 0 021245993 | 5 69E-12 | 0 387225749 | 0 | 8 98E-12 |
| ICAM4 | 0 00120881 | 0 02123555 | 5 83E-12 | -0 49505505 | 0 | 9 19E-12 |
| SOCS1 | 0 033313662 | 0 021231322 | 5 88E-12 | -0 45780562 | 0 | 9 28E-12 |
| TM6SF1 | 0 034627173 | 0 021227028 | 5 94E-12 | 0 464222363 | 0 | 9 37E-12 |
| ITGAL | 0 014210657 | 0 021200913 | 6 30E-12 | 0 487038062 | 0 | 9 94E-12 |
| COPS3 | 0 126801317 | 0 021179149 | 6 62E-12 | 0 391175836 | 0 | 1 04E-11 |
| RABEP1 | 0 497925035 | 0 021173489 | 6 71E-12 | -0 08342905 | 1 33E-161 | 1 06E-11 |
| DBF4 | 0 151453427 | 0 021134361 | 7 33E-12 | 0 316296852 | 0 | 1 16E-11 |
| FITM1 | 0 003807276 | 0 021116707 | 7 63E-12 | 0 495857633 | 0 | 1 20E-11 |
| CAMK2G | 0 15659325 | 0 02109579 | 8 00E-12 | 0 354114469 | 0 | 1 26E-11 |
| AMN1 | 0 124935752 | 0 02109273 | 8 06E-12 | -0 36746185 | 0 | 1 27E-11 |
| RASGRP2 | 0 014096438 | 0 021091624 | 8 08E-12 | -0 4785464 | 0 | 1 27E-11 |
| FAM107A | 0 072500048 | 0 021088179 | 8 14E-12 | -0 41680384 | 0 | 1 28E-11 |
| SERPINF2 | 0 015809712 | 0 021085641 | 8 19E-12 | -0 4758193 | 0 | 1 29E-11 |
| ULBP2 | 0 012602083 | 0 021074787 | 8 39E-12 | 0 488826387 | 0 | 1 32E-11 |
| RBL1 | 0 219384744 | 0 021052457 | 8 83E-12 | 0 247922287 | 0 | 1 39E-11 |
| LGALS12 | 0 002341475 | 0 021035857 | 9 16E-12 | -0 4935749 | 0 | 1 44E-11 |
| TRMT2B | 0 107403248 | 0 021035181 | 9 18E-12 | 0 419152396 | 0 | 1 44E-11 |
| LINGO2 | 0 132236203 | 0 021027506 | 9 34E-12 | -0 35064967 | 0 | 1 47E-11 |
| ERP27 | 0 006881651 | 0 021018265 | 9 53E-12 | 0 493260618 | 0 | 1 50E-11 |
| PTHLH | 0 037396966 | 0 021017666 | 9 55E-12 | 0 462393657 | 0 | 1 50E-11 |
| PTGFR | 0 133549714 | 0 020996147 | 1 00E-11 | 0 338293504 | 0 | 1 57E-11 |
| RCHY1 | 0 165759266 | 0 02099084 | 1 01E-11 | -0 32789131 | 0 | 1 59E-11 |
| TEX29 | 0 010098799 | 0 020958687 | 1 09E-11 | 0 490199912 | 0 | 1 71E-11 |
| NR5A2 | 0 01377282 | 0 020956952 | 1 09E-11 | 0 486577081 | 0 | 1 72E-11 |
| ZNF703 | 0 049332775 | 0 020947122 | 1 12E-11 | -0 4351659 | 0 | 1 76E-11 |
| MAP7 | 0 084283566 | 0 020942713 | 1 13E-11 | -0 3957647 | 0 | 1 77E-11 |
| FAM53C | 0 12148065 | 0 020941756 | 1 13E-11 | -0 3603681 | 0 | 1 78E-11 |
| AGT | 0 011602673 | 0 0209216 | 1 18E-11 | -0 48148858 | 0 | 1 86E-11 |
| FSD1 | 0 006386705 | 0 02090313 | 1 24E-11 | 0 493679915 | 0 | 1 94E-11 |
| GPR19 | 0 015990558 | 0 020882468 | 1 29E-11 | 0 479157421 | 0 | 2 03E-11 |
| CFI | 0 165483238 | 0 020881919 | 1 30E-11 | -0 30428691 | 0 | 2 03E-11 |
| CKAP5 | 0 328577411 | 0 020879276 | 1 30E-11 | 0 20719614 | 0 | 2 04E-11 |
| SHD | 0 002036893 | 0 020853052 | 1 38E-11 | 0 496972744 | 0 | 2 17E-11 |
| TAF1D | 0 489549028 | 0 020842001 | 1 42E-11 | -0 08070977 | 2 45E-151 | 2 22E-11 |
| PBXIP1 | 0 084197902 | 0 020840893 | 1 42E-11 | -0 39398419 | 0 | 2 22E-11 |
| SERPINB2 | 0 004997049 | 0 020823539 | 1 48E-11 | 0 494677907 | 0 | 2 31E-11 |

| | | | | | | |
|---|---|---|---|---|---|---|
| FSTL4 | 0 020188079 | 0 020806678 | 1 53E-11 | -0 47144627 | 0 | 2 40E-11 |
| CR1 | 0 012925701 | 0 020803646 | 1 54E-11 | 0 485971728 | 0 | 2 42E-11 |
| TTC28-AS1 | 0 17099427 | 0 020802276 | 1 55E-11 | 0 336953735 | 0 | 2 42E-11 |
| IGFBP3 | 0 132007767 | 0 020761065 | 1 70E-11 | -0 36528843 | 0 | 2 65E-11 |
| ZNF780A | 0 079514953 | 0 020760798 | 1 70E-11 | -0 41149962 | 0 | 2 66E-11 |
| ZNF277 | 0 284926995 | 0 02073677 | 1 79E-11 | 0 257518269 | 0 | 2 80E-11 |
| ADAMTS16 | 0 075412614 | 0 020717702 | 1 87E-11 | 0 408469362 | 0 | 2 92E-11 |
| EAF1-AS1 | 0 013144619 | 0 020714398 | 1 88E-11 | -0 48102784 | 0 | 2 94E-11 |
| ZC3H12B | 0 362966629 | 0 020707338 | 1 91E-11 | -0 11617438 | 0 | 2 99E-11 |
| C6 | 0 0121928 | 0 020693853 | 1 97E-11 | -0 48217941 | 0 | 3 08E-11 |
| KC6 | 0 099950505 | 0 020676558 | 2 05E-11 | -0 37880618 | 0 | 3 20E-11 |
| HSD17B6 | 0 010098799 | 0 020674092 | 2 06E-11 | 0 487950492 | 0 | 3 21E-11 |
| PTGER3 | 0 028373722 | 0 020654455 | 2 15E-11 | -0 46591099 | 0 | 3 35E-11 |
| TCEAL2 | 0 019093488 | 0 020653881 | 2 15E-11 | 0 48624227 | 0 | 3 36E-11 |
| EIF2S3 | 0 375302203 | 0 020653106 | 2 16E-11 | 0 218224103 | 0 | 3 36E-11 |
| SNHG7 | 0 170756315 | 0 020640212 | 2 22E-11 | -0 33478043 | 0 | 3 46E-11 |
| SLC26A7 | 0 035578991 | 0 020634695 | 2 25E-11 | -0 45379232 | 0 | 3 50E-11 |
| SHISA5 | 0 165407093 | 0 020621426 | 2 31E-11 | -0 31998184 | 0 | 3 61E-11 |
| OPA3 | 0 128447964 | 0 020620252 | 2 32E-11 | 0 376196728 | 0 | 3 61E-11 |
| MOB3A | 0 12069064 | 0 020614529 | 2 35E-11 | -0 36374154 | 0 | 3 66E-11 |
| ING1 | 0 06003122 | 0 020603112 | 2 41E-11 | -0 42548702 | 0 | 3 75E-11 |
| NFKBIZ | 0 288382098 | 0 020600224 | 2 42E-11 | -0 22970004 | 0 | 3 78E-11 |
| SNX18 | 0 085187794 | 0 020591261 | 2 47E-11 | -0 40095048 | 0 | 3 85E-11 |
| RHEX | 0 016437913 | 0 020588293 | 2 49E-11 | 0 486265948 | 0 | 3 87E-11 |
| RGS19 | 0 022006054 | 0 020586112 | 2 50E-11 | 0 479735403 | 0 | 3 89E-11 |
| MYO3B | 0 04015724 | 0 020562382 | 2 64E-11 | -0 45203868 | 0 | 4 10E-11 |
| MAPRE2 | 0 272315395 | 0 02054605 | 2 73E-11 | -0 22500698 | 0 | 4 25E-11 |
| LNX1-AS1 | 0 012801965 | 0 020512882 | 2 94E-11 | -0 48268981 | 0 | 4 57E-11 |
| PAWR | 0 310016942 | 0 020505542 | 2 99E-11 | 0 223315404 | 0 | 4 64E-11 |
| BLM | 0 079514953 | 0 020498548 | 3 03E-11 | 0 357847659 | 0 | 4 72E-11 |
| RDM1 | 0 005558623 | 0 020485674 | 3 12E-11 | 0 490955792 | 0 | 4 85E-11 |
| AMIGO2 | 0 012954256 | 0 020481059 | 3 15E-11 | 0 486943322 | 0 | 4 90E-11 |
| CNNM4 | 0 087595896 | 0 020474486 | 3 20E-11 | -0 40366847 | 0 | 4 97E-11 |
| RGS18 | 0 007214787 | 0 020474313 | 3 20E-11 | 0 491842378 | 0 | 4 97E-11 |
| COLGALT1 | 0 179008585 | 0 02046879 | 3 24E-11 | 0 336210522 | 0 | 5 03E-11 |
| CLIC5 | 0 025080429 | 0 020461467 | 3 29E-11 | -0 46545026 | 0 | 5 11E-11 |
| CYP3A5 | 0 083445965 | 0 020460467 | 3 30E-11 | 0 423745987 | 0 | 5 12E-11 |
| NCKAP1 | 0 642563439 | 0 020460277 | 3 30E-11 | -0 04505624 | 2 37E-48 | 5 12E-11 |
| TSPOAP1-AS1 | 0 073290057 | 0 020448222 | 3 39E-11 | 0 412872968 | 0 | 5 26E-11 |
| SNRK | 0 279530182 | 0 020435475 | 3 49E-11 | -0 24019897 | 0 | 5 41E-11 |
| DYNC1I1 | 0 407321391 | 0 020428332 | 3 54E-11 | -0 12541479 | 0 | 5 49E-11 |
| G0S2 | 0 020016752 | 0 020403334 | 3 74E-11 | 0 48043191 | 0 | 5 80E-11 |
| SMC1B | 0 041918105 | 0 020392979 | 3 83E-11 | 0 452554732 | 0 | 5 93E-11 |
| LINC01098 | 0 008813843 | 0 020390451 | 3 85E-11 | 0 492195814 | 0 | 5 96E-11 |

| | | | | | | |
|---|---|---|---|---|---|---|
| IK | 0 169480878 | 0 02037992 | 3 94E-11 | -0 32522438 | 0 | 6 10E-11 |
| CYP4A22-AS1 | 0 029630123 | 0 020365132 | 4 07E-11 | -0 46206217 | 0 | 6 30E-11 |
| TAF5L | 0 090299062 | 0 020362809 | 4 09E-11 | 0 420439242 | 0 | 6 33E-11 |
| SF1 | 0 485979707 | 0 020362264 | 4 09E-11 | -0 08301179 | 5 25E-160 | 6 34E-11 |
| MTHFD2 | 0 104871409 | 0 020344257 | 4 26E-11 | 0 375948293 | 0 | 6 59E-11 |
| HNRNPA3 | 0 601844625 | 0 020338425 | 4 31E-11 | 0 089583851 | 4 13E-186 | 6 67E-11 |
| TRIB3 | 0 035131637 | 0 020335685 | 4 34E-11 | 0 465584497 | 0 | 6 71E-11 |
| COX20 | 0 219327635 | 0 020327994 | 4 41E-11 | 0 335537253 | 0 | 6 82E-11 |
| ROCK1 | 0 634844187 | 0 020308403 | 4 60E-11 | -0 04431841 | 7 76E-47 | 7 12E-11 |
| MEI4 | 0 010108317 | 0 020289885 | 4 79E-11 | 0 49040452 | 0 | 7 41E-11 |
| HTR4 | 0 00651996 | 0 020286727 | 4 83E-11 | 0 492539882 | 0 | 7 46E-11 |
| GPR34 | 0 037863357 | 0 020284462 | 4 85E-11 | -0 45787071 | 0 | 7 50E-11 |
| JMY | 0 233995165 | 0 020279032 | 4 91E-11 | -0 25143352 | 0 | 7 59E-11 |
| MTMR8 | 0 089756525 | 0 020265585 | 5 05E-11 | -0 39094212 | 0 | 7 81E-11 |
| LINC02112 | 0 018465287 | 0 020249328 | 5 24E-11 | -0 47201286 | 0 | 8 09E-11 |
| PLA2G6 | 0 093078373 | 0 020240501 | 5 34E-11 | -0 38966949 | 0 | 8 25E-11 |
| GLIPR2 | 0 084968876 | 0 020239331 | 5 35E-11 | -0 40182136 | 0 | 8 27E-11 |
| MIR2052HG | 0 060012183 | 0 020234893 | 5 40E-11 | -0 42887394 | 0 | 8 34E-11 |
| TBXA2R | 0 011231463 | 0 020222933 | 5 55E-11 | -0 48359542 | 0 | 8 56E-11 |
| ABCA10 | 0 600578706 | 0 020220238 | 5 58E-11 | -0 04076058 | 6 98E-40 | 8 61E-11 |
| CDKN1B | 0 095638766 | 0 020216972 | 5 62E-11 | 0 414740965 | 0 | 8 67E-11 |
| HLA-DOA | 0 015238621 | 0 020212585 | 5 67E-11 | -0 47821791 | 0 | 8 75E-11 |
| OLFM1 | 0 031076888 | 0 020209203 | 5 71E-11 | 0 47617745 | 0 | 8 81E-11 |
| SNX3 | 0 287325579 | 0 020201404 | 5 81E-11 | -0 21489677 | 0 | 8 96E-11 |
| HPS4 | 0 194561307 | 0 02018736 | 5 99E-11 | 0 31700325 | 0 | 9 24E-11 |
| DTD1 | 0 233433592 | 0 020155035 | 6 43E-11 | -0 26947884 | 0 | 9 90E-11 |
| KRTCAP3 | 0 03290438 | 0 020150235 | 6 49E-11 | 0 462826863 | 0 | 1 00E-10 |
| MCF2 | 0 011440863 | 0 020118758 | 6 95E-11 | -0 48180515 | 0 | 1 07E-10 |
| NF1 | 0 726095068 | 0 020102555 | 7 20E-11 | -0 0245741 | 1 63E-15 | 1 11E-10 |
| HSD17B8 | 0 019683615 | 0 020090745 | 7 38E-11 | -0 47215219 | 0 | 1 14E-10 |
| SEPTIN9 | 0 528602159 | 0 020067157 | 7 77E-11 | -0 09155264 | 2 53E-194 | 1 20E-10 |
| CAMKK1 | 0 058965182 | 0 020065303 | 7 80E-11 | 0 444738448 | 0 | 1 20E-10 |
| ARHGAP39 | 0 125516362 | 0 020061567 | 7 86E-11 | -0 34761842 | 0 | 1 21E-10 |
| LINC02208 | 0 045753936 | 0 020061476 | 7 87E-11 | -0 44041328 | 0 | 1 21E-10 |
| NIPAL1 | 0 022719918 | 0 020061182 | 7 87E-11 | 0 480248631 | 0 | 1 21E-10 |
| DOK3 | 0 020426034 | 0 020057824 | 7 93E-11 | -0 47242254 | 0 | 1 22E-10 |
| FERMT2 | 0 465315718 | 0 020050921 | 8 05E-11 | -0 08499845 | 1 10E-167 | 1 24E-10 |
| HMGN2 | 0 380242143 | 0 020046155 | 8 13E-11 | 0 178151014 | 0 | 1 25E-10 |
| TMC1 | 0 015124403 | 0 020028257 | 8 45E-11 | -0 47731402 | 0 | 1 30E-10 |
| MARK3 | 0 655765167 | 0 020028049 | 8 45E-11 | -0 03466058 | 2 66E-29 | 1 30E-10 |
| MYBPC2 | 0 004768613 | 0 020021736 | 8 57E-11 | 0 494521052 | 0 | 1 32E-10 |
| RAB3IP | 0 143981649 | 0 019999711 | 8 98E-11 | -0 33972058 | 0 | 1 38E-10 |
| GGA1 | 0 225409758 | 0 019984921 | 9 27E-11 | -0 27699514 | 0 | 1 42E-10 |
| HPS3 | 0 216795797 | 0 019979108 | 9 39E-11 | 0 328647223 | 0 | 1 44E-10 |

| | | | | | | |
|---|---|---|---|---|---|---|
| IKZF1 | 0 022329672 | 0 01997035 | 9 57E-11 | 0 478367282 | 0 | 1 47E-10 |
| TOGARAM2 | 0 040699777 | 0 019954251 | 9 91E-11 | -0 4416379 | 0 | 1 52E-10 |
| MAP7D2 | 0 019588433 | 0 019949299 | 1 00E-10 | 0 484739055 | 0 | 1 54E-10 |
| ANXA11 | 0 273267214 | 0 019912992 | 1 08E-10 | -0 21559023 | 0 | 1 66E-10 |
| PRR14L | 0 318297767 | 0 019906555 | 1 10E-10 | 0 220588845 | 0 | 1 68E-10 |
| MIR137HG | 0 127895909 | 0 019893989 | 1 13E-10 | 0 370225721 | 0 | 1 73E-10 |
| UBA7 | 0 062896195 | 0 019892853 | 1 13E-10 | -0 42839085 | 0 | 1 73E-10 |
| AMMECR1 | 0 208876663 | 0 019890891 | 1 13E-10 | 0 309131428 | 0 | 1 74E-10 |
| TRPC6 | 0 048181074 | 0 019883786 | 1 15E-10 | -0 43194811 | 0 | 1 76E-10 |
| POLR3GL | 0 088661933 | 0 01987844 | 1 17E-10 | 0 425348997 | 0 | 1 78E-10 |
| ENG | 0 2091051 | 0 019872248 | 1 18E-10 | -0 28225359 | 0 | 1 81E-10 |
| DIP2B | 0 586929623 | 0 019867041 | 1 19E-10 | -0 05926442 | 2 23E-82 | 1 83E-10 |
| SCGB3A1 | 0 003769203 | 0 019855666 | 1 22E-10 | 0 496011741 | 0 | 1 87E-10 |
| LINC02015 | 0 006025014 | 0 019833741 | 1 28E-10 | -0 48903609 | 0 | 1 96E-10 |
| ADD3 | 0 613466334 | 0 019832472 | 1 29E-10 | -0 0233884 | 3 41E-14 | 1 97E-10 |
| LINC00623 | 0 098418077 | 0 019827192 | 1 30E-10 | 0 40549966 | 0 | 1 99E-10 |
| FMN2 | 0 230730426 | 0 019820155 | 1 32E-10 | 0 282552754 | 0 | 2 02E-10 |
| ST8SIA5 | 0 023243418 | 0 019803873 | 1 37E-10 | 0 477372412 | 0 | 2 09E-10 |
| LONRF2 | 0 09193619 | 0 019803127 | 1 37E-10 | -0 37766385 | 0 | 2 09E-10 |
| EHBP1L1 | 0 214340104 | 0 019799625 | 1 38E-10 | -0 28426979 | 0 | 2 11E-10 |
| DNAJC3 | 0 359682854 | 0 019767573 | 1 48E-10 | -0 15767819 | 0 | 2 25E-10 |
| SLC30A8 | 0 015448021 | 0 019763995 | 1 49E-10 | -0 47910058 | 0 | 2 27E-10 |
| ZFAND2A | 0 07401344 | 0 01975738 | 1 51E-10 | -0 41252535 | 0 | 2 30E-10 |
| MGLL | 0 268860292 | 0 01975532 | 1 51E-10 | -0 23242872 | 0 | 2 31E-10 |
| NCAPG | 0 086006358 | 0 019746353 | 1 54E-10 | 0 359444701 | 0 | 2 36E-10 |
| SEMA7A | 0 019683615 | 0 019740801 | 1 56E-10 | -0 47057313 | 0 | 2 38E-10 |
| TNNI2 | 0 012202319 | 0 019739907 | 1 57E-10 | 0 487232399 | 0 | 2 39E-10 |
| COL18A1 | 0 362880966 | 0 01973257 | 1 59E-10 | -0 19629013 | 0 | 2 42E-10 |
| TNR | 0 005292113 | 0 019720837 | 1 63E-10 | -0 48974603 | 0 | 2 49E-10 |
| MAP4K1 | 0 006919724 | 0 01970139 | 1 70E-10 | 0 492364196 | 0 | 2 59E-10 |
| HERC4 | 0 672707544 | 0 019690333 | 1 74E-10 | -0 02944735 | 1 34E-21 | 2 65E-10 |
| FANCD2 | 0 104671527 | 0 019669935 | 1 82E-10 | 0 31190923 | 0 | 2 77E-10 |
| LINC02341 | 0 008632998 | 0 019659152 | 1 86E-10 | -0 48664313 | 0 | 2 83E-10 |
| ZKSCAN4 | 0 035026936 | 0 019636672 | 1 95E-10 | -0 45777966 | 0 | 2 97E-10 |
| CKAP2 | 0 08731035 | 0 019631861 | 1 97E-10 | 0 371666412 | 0 | 3 00E-10 |
| LINC02458 | 0 019893016 | 0 019582781 | 2 18E-10 | 0 479032762 | 0 | 3 32E-10 |
| LINC02613 | 0 03169557 | 0 019580222 | 2 19E-10 | -0 45869586 | 0 | 3 34E-10 |
| ADRB1 | 0 005777541 | 0 019572892 | 2 23E-10 | 0 494400916 | 0 | 3 39E-10 |
| SLC26A5 | 0 158392187 | 0 019568279 | 2 25E-10 | -0 33674709 | 0 | 3 42E-10 |
| MYCT1 | 0 00614875 | 0 019549938 | 2 34E-10 | -0 48978011 | 0 | 3 56E-10 |
| RSF1 | 0 553254269 | 0 019530869 | 2 43E-10 | 0 100560971 | 3 25E-234 | 3 70E-10 |
| TNC | 0 283194685 | 0 019523946 | 2 47E-10 | 0 219996112 | 0 | 3 75E-10 |
| IFNLR1 | 0 01408692 | 0 01950526 | 2 57E-10 | 0 487306872 | 0 | 3 90E-10 |
| MYOZ2 | 0 014610421 | 0 019500225 | 2 60E-10 | -0 4774681 | 0 | 3 94E-10 |

| | | | | | | |
|---|---|---|---|---|---|---|
| UBE2Q2 | 0 366355105 | 0 019494733 | 2 63E-10 | 0 192202802 | 0 | 3 99E-10 |
| TEK | 0 067902762 | 0 019491553 | 2 64E-10 | 0 423519066 | 0 | 4 01E-10 |
| LINC00865 | 0 014734157 | 0 019451902 | 2 87E-10 | -0 47827244 | 0 | 4 36E-10 |
| CCDC62 | 0 03784432 | 0 019450089 | 2 88E-10 | -0 45447993 | 0 | 4 38E-10 |
| ZSWIM7 | 0 198121109 | 0 019410633 | 3 13E-10 | -0 29414967 | 0 | 4 75E-10 |
| C2CD4B | 0 00362643 | 0 019387234 | 3 29E-10 | -0 49181996 | 0 | 4 99E-10 |
| TIMP3 | 0 52023567 | 0 019380846 | 3 33E-10 | -0 09751178 | 2 76E-220 | 5 05E-10 |
| KLHL15 | 0 076107441 | 0 019378124 | 3 35E-10 | 0 419028773 | 0 | 5 08E-10 |
| CD47 | 0 466058137 | 0 019371532 | 3 40E-10 | 0 156360596 | 0 | 5 15E-10 |
| UBE2T | 0 051379186 | 0 019367728 | 3 42E-10 | 0 403083973 | 0 | 5 19E-10 |
| ATG2A | 0 077401915 | 0 019364018 | 3 45E-10 | 0 427281486 | 0 | 5 23E-10 |
| PIN4 | 0 23890655 | 0 019355905 | 3 51E-10 | 0 31097536 | 0 | 5 32E-10 |
| SLC7A7 | 0 034189336 | 0 019331483 | 3 69E-10 | -0 45925767 | 0 | 5 59E-10 |
| KIF26B-AS1 | 0 043479089 | 0 019331001 | 3 70E-10 | 0 44397551 | 0 | 5 60E-10 |
| GZF1 | 0 039291085 | 0 019307463 | 3 88E-10 | 0 462559868 | 0 | 5 88E-10 |
| RELN | 0 052730768 | 0 019304902 | 3 90E-10 | -0 43683819 | 0 | 5 91E-10 |
| ADAT2 | 0 10681312 | 0 019302991 | 3 92E-10 | -0 38381696 | 0 | 5 93E-10 |
| SLC30A10 | 0 033361253 | 0 019301628 | 3 93E-10 | 0 467471026 | 0 | 5 95E-10 |
| PABPC1 | 0 595619729 | 0 019260961 | 4 28E-10 | 0 064943906 | 1 42E-98 | 6 47E-10 |
| ZNF525 | 0 062248958 | 0 019239237 | 4 47E-10 | 0 437521333 | 0 | 6 76E-10 |
| MICAL2 | 0 167148922 | 0 019230714 | 4 55E-10 | -0 32858517 | 0 | 6 88E-10 |
| NIBAN3 | 0 009975062 | 0 019220904 | 4 65E-10 | -0 48477223 | 0 | 7 02E-10 |
| CDKN1C | 0 08589214 | 0 019218373 | 4 67E-10 | -0 39473458 | 0 | 7 06E-10 |
| SENP3 | 0 078220479 | 0 019186567 | 4 99E-10 | 0 4339982 | 0 | 7 53E-10 |
| IQCM | 0 010822181 | 0 019160648 | 5 26E-10 | 0 489193811 | 0 | 7 94E-10 |
| SHROOM4 | 0 362852411 | 0 01915653 | 5 31E-10 | -0 16902178 | 0 | 8 01E-10 |
| KIF11 | 0 098161086 | 0 019154764 | 5 33E-10 | 0 354412073 | 0 | 8 04E-10 |
| HS3ST5 | 0 082713065 | 0 019142179 | 5 47E-10 | -0 40444842 | 0 | 8 25E-10 |
| RAPH1 | 0 486236698 | 0 019128659 | 5 62E-10 | 0 128709991 | 0 | 8 48E-10 |
| TPH1 | 0 02002627 | 0 019115003 | 5 78E-10 | 0 483537192 | 0 | 8 72E-10 |
| HAUS2 | 0 095553102 | 0 019114326 | 5 79E-10 | 0 408221746 | 0 | 8 73E-10 |
| IFI44L | 0 167443985 | 0 019103225 | 5 92E-10 | -0 3395715 | 0 | 8 93E-10 |
| ING3 | 0 121309322 | 0 019102022 | 5 94E-10 | 0 389832822 | 0 | 8 95E-10 |
| GALNT18 | 0 112657288 | 0 019096221 | 6 01E-10 | 0 401989133 | 0 | 9 06E-10 |
| ZNF317 | 0 085511412 | 0 019080672 | 6 20E-10 | 0 419519653 | 0 | 9 35E-10 |
| PPP1R15B | 0 109877977 | 0 019077933 | 6 24E-10 | -0 37352082 | 0 | 9 40E-10 |
| CTSH | 0 085635149 | 0 019076626 | 6 26E-10 | -0 41060438 | 0 | 9 43E-10 |
| L3MBTL2 | 0 076354914 | 0 019053631 | 6 56E-10 | 0 428311143 | 0 | 9 88E-10 |
| NSRP1 | 0 567236489 | 0 019003997 | 7 26E-10 | 0 103799348 | 1 74E-249 | 1 09E-09 |
| PGR | 0 032733053 | 0 019002661 | 7 28E-10 | -0 45102963 | 0 | 1 10E-09 |
| SPPL2A | 0 417448745 | 0 019002608 | 7 28E-10 | -0 12439279 | 0 | 1 10E-09 |
| PGPEP1 | 0 083322229 | 0 019000261 | 7 32E-10 | -0 3894523 | 0 | 1 10E-09 |
| ZFAND6 | 0 500713864 | 0 018995207 | 7 39E-10 | 0 142745681 | 0 | 1 11E-09 |
| KRAS | 0 249014867 | 0 018952436 | 8 07E-10 | 0 307879893 | 0 | 1 21E-09 |

| | | | | | | |
|---|---|---|---|---|---|---|
| PPP1R9A-AS1 | 0 044383317 | 0 018944246 | 8 20E-10 | -0 44752518 | 0 | 1 23E-09 |
| MTM1 | 0 172536217 | 0 01892966 | 8 45E-10 | -0 30824947 | 0 | 1 27E-09 |
| MARCO | 0 013411129 | 0 018928464 | 8 47E-10 | 0 484124525 | 0 | 1 27E-09 |
| FAM184A | 0 160143534 | 0 018913194 | 8 74E-10 | 0 381291713 | 0 | 1 31E-09 |
| MBP | 0 126534808 | 0 018894006 | 9 09E-10 | -0 36751147 | 0 | 1 37E-09 |
| ABHD15-AS1 | 0 200081856 | 0 018877572 | 9 40E-10 | 0 309932477 | 0 | 1 41E-09 |
| LY6G5C | 0 005215968 | 0 018870826 | 9 53E-10 | -0 49010664 | 0 | 1 43E-09 |
| ADAMTS4 | 0 037682511 | 0 018869375 | 9 56E-10 | 0 44802731 | 0 | 1 43E-09 |
| C1orf35 | 0 100645333 | 0 01886172 | 9 71E-10 | 0 409423371 | 0 | 1 46E-09 |
| HPRT1 | 0 193009842 | 0 018859334 | 9 75E-10 | 0 333867812 | 0 | 1 46E-09 |
| ZNF257 | 0 018436733 | 0 018858432 | 9 77E-10 | 0 482986198 | 0 | 1 47E-09 |
| SLC16A6 | 0 021739544 | 0 018831527 | 1 03E-09 | 0 476024599 | 0 | 1 55E-09 |
| LGR6 | 0 010736517 | 0 018824296 | 1 05E-09 | -0 48241078 | 0 | 1 57E-09 |
| S1PR5 | 0 002864975 | 0 018824129 | 1 05E-09 | 0 495865577 | 0 | 1 57E-09 |
| AADACL2-AS1 | 0 069844473 | 0 018809474 | 1 08E-09 | -0 42576995 | 0 | 1 62E-09 |
| TIGIT | 0 008652034 | 0 018807764 | 1 08E-09 | 0 490578891 | 0 | 1 62E-09 |
| MVD | 0 082151492 | 0 018797787 | 1 11E-09 | -0 40010652 | 0 | 1 66E-09 |
| PRKAR1A | 0 43282062 | 0 018787565 | 1 13E-09 | -0 12482822 | 0 | 1 69E-09 |
| CWC15 | 0 137975672 | 0 018785574 | 1 13E-09 | -0 35220162 | 0 | 1 70E-09 |
| TOB2 | 0 138327845 | 0 018767568 | 1 17E-09 | -0 340166 | 0 | 1 76E-09 |
| KCNA3 | 0 005596695 | 0 018756396 | 1 20E-09 | 0 494475552 | 0 | 1 80E-09 |
| HNRNPDL | 0 661000171 | 0 018672041 | 1 42E-09 | 0 080673614 | 3 34E-151 | 2 13E-09 |
| AIF1L | 0 041480269 | 0 018651954 | 1 48E-09 | -0 44750628 | 0 | 2 22E-09 |
| CHRNA5 | 0 040566523 | 0 018615806 | 1 60E-09 | 0 444186365 | 0 | 2 39E-09 |
| SPARC | 0 674211418 | 0 018615034 | 1 60E-09 | -0 02947447 | 1 23E-21 | 2 39E-09 |
| ZNF628 | 0 012973292 | 0 018612097 | 1 61E-09 | -0 4807532 | 0 | 2 40E-09 |
| AXDND1 | 0 020302298 | 0 018607245 | 1 62E-09 | -0 47266215 | 0 | 2 43E-09 |
| KCNJ1 | 0 002189184 | 0 018577673 | 1 72E-09 | 0 496177737 | 0 | 2 57E-09 |
| COL4A4 | 0 01729455 | 0 018540782 | 1 85E-09 | 0 484938956 | 0 | 2 77E-09 |
| SSH1 | 0 343768441 | 0 018536996 | 1 87E-09 | -0 18783625 | 0 | 2 79E-09 |
| AAK1 | 0 576107441 | 0 018534197 | 1 88E-09 | 0 117183169 | 0 | 2 80E-09 |
| AFTPH | 0 361624565 | 0 018533714 | 1 88E-09 | -0 17142325 | 0 | 2 81E-09 |
| SCN8A | 0 167072776 | 0 018519501 | 1 93E-09 | 0 283758527 | 0 | 2 89E-09 |
| SLC38A11 | 0 038263121 | 0 018506237 | 1 99E-09 | -0 44888505 | 0 | 2 96E-09 |
| MAPKAP1 | 0 505196931 | 0 018491307 | 2 05E-09 | 0 132276585 | 0 | 3 05E-09 |
| FAM114A1 | 0 437979479 | 0 018487885 | 2 06E-09 | -0 13371949 | 0 | 3 07E-09 |
| VWF | 0 057090099 | 0 018486889 | 2 06E-09 | 0 442459646 | 0 | 3 08E-09 |
| IRF2BPL | 0 06344825 | 0 018465509 | 2 15E-09 | -0 4259143 | 0 | 3 21E-09 |
| LINC02762 | 0 064895014 | 0 018443842 | 2 25E-09 | -0 4222932 | 0 | 3 35E-09 |
| SLC12A2 | 0 438569607 | 0 01842512 | 2 33E-09 | -0 08126694 | 2 06E-153 | 3 48E-09 |
| STIL | 0 094068264 | 0 018419463 | 2 36E-09 | 0 331439499 | 0 | 3 52E-09 |
| APOLD1 | 0 151681864 | 0 018419277 | 2 36E-09 | 0 292006 | 0 | 3 52E-09 |
| PWP2 | 0 03811083 | 0 018374143 | 2 58E-09 | -0 45015583 | 0 | 3 84E-09 |
| SLC37A4 | 0 02444271 | 0 018365586 | 2 63E-09 | 0 478908604 | 0 | 3 91E-09 |

| | | | | | | |
|---|---|---|---|---|---|---|
| CEP128 | 0 242494908 | 0 018358861 | 2 66E-09 | 0 22189172 | 0 | 3 96E-09 |
| CCDC134 | 0 069263863 | 0 018357968 | 2 67E-09 | 0 421595261 | 0 | 3 97E-09 |
| JSRP1 | 0 007509851 | 0 018338575 | 2 77E-09 | 0 489623629 | 0 | 4 12E-09 |
| RAB11FIP4 | 0 028097695 | 0 018322511 | 2 86E-09 | -0 46511945 | 0 | 4 25E-09 |
| FER1L6-AS2 | 0 004911386 | 0 01832125 | 2 87E-09 | 0 494121082 | 0 | 4 26E-09 |
| LINC02465 | 0 005225486 | 0 018320532 | 2 87E-09 | 0 494534965 | 0 | 4 27E-09 |
| FILIP1L | 0 526298757 | 0 018300434 | 2 99E-09 | 0 118059007 | 0 | 4 44E-09 |
| ASTE1 | 0 053587406 | 0 018298815 | 3 00E-09 | -0 43747714 | 0 | 4 45E-09 |
| CNST | 0 270402239 | 0 018276786 | 3 13E-09 | -0 22683512 | 0 | 4 65E-09 |
| TMOD3 | 0 385163047 | 0 018268736 | 3 18E-09 | 0 180854297 | 0 | 4 72E-09 |
| OPRL1 | 0 006767433 | 0 018247552 | 3 32E-09 | 0 492348719 | 0 | 4 93E-09 |
| BMP2K | 0 436018732 | 0 018238753 | 3 38E-09 | 0 151001442 | 0 | 5 01E-09 |
| BACH1 | 0 537558775 | 0 018225146 | 3 47E-09 | 0 105880962 | 1 46E-259 | 5 15E-09 |
| ZNF480 | 0 084550075 | 0 018221461 | 3 49E-09 | 0 417104243 | 0 | 5 18E-09 |
| RSBN1 | 0 201738021 | 0 018206294 | 3 60E-09 | -0 29430852 | 0 | 5 34E-09 |
| SOX17 | 0 007414669 | 0 01820601 | 3 60E-09 | -0 48795977 | 0 | 5 34E-09 |
| MMD | 0 089194952 | 0 01820153 | 3 63E-09 | 0 41833621 | 0 | 5 39E-09 |
| ARHGAP27 | 0 050770022 | 0 01818307 | 3 77E-09 | -0 43396673 | 0 | 5 58E-09 |
| SENCR | 0 003255221 | 0 018180166 | 3 79E-09 | 0 495006744 | 0 | 5 61E-09 |
| LINC02073 | 0 005530068 | 0 018132798 | 4 16E-09 | 0 494301229 | 0 | 6 16E-09 |
| SPINK2 | 0 00331233 | 0 018130317 | 4 18E-09 | 0 495368879 | 0 | 6 19E-09 |
| OTOA | 0 00846167 | 0 018127213 | 4 20E-09 | 0 490932169 | 0 | 6 22E-09 |
| LINC00894 | 0 155965049 | 0 018102024 | 4 41E-09 | -0 35305667 | 0 | 6 54E-09 |
| IL18BP | 0 029411205 | 0 018088883 | 4 53E-09 | 0 465389523 | 0 | 6 70E-09 |
| COMMD7 | 0 11716891 | 0 018088163 | 4 54E-09 | -0 3743972 | 0 | 6 71E-09 |
| FXYD7 | 0 014705602 | 0 018078588 | 4 62E-09 | 0 488793934 | 0 | 6 84E-09 |
| STK17A | 0 295558813 | 0 018075965 | 4 64E-09 | 0 227378915 | 0 | 6 87E-09 |
| SLC9A6 | 0 096790467 | 0 018069561 | 4 70E-09 | 0 418730582 | 0 | 6 96E-09 |
| PHF19 | 0 098218195 | 0 018062008 | 4 77E-09 | 0 378417655 | 0 | 7 06E-09 |
| ANKRD37 | 0 051950277 | 0 018049061 | 4 90E-09 | -0 435659 | 0 | 7 24E-09 |
| SPIRE1 | 0 436685005 | 0 018046616 | 4 92E-09 | 0 178204455 | 0 | 7 27E-09 |
| CMTM3 | 0 124754907 | 0 018039849 | 4 98E-09 | -0 35643377 | 0 | 7 37E-09 |
| MIR4500HG | 0 015276694 | 0 018023358 | 5 15E-09 | 0 483962243 | 0 | 7 61E-09 |
| GGH | 0 07805867 | 0 017973328 | 5 67E-09 | 0 397547337 | 0 | 8 38E-09 |
| ADK | 0 674839619 | 0 017962626 | 5 79E-09 | -0 01655704 | 8 01E-08 | 8 55E-09 |
| GULP1 | 0 59645733 | 0 0179595 | 5 83E-09 | -0 04676602 | 5 89E-52 | 8 60E-09 |
| VAPA | 0 483152805 | 0 017903681 | 6 49E-09 | -0 09816874 | 3 00E-223 | 9 58E-09 |
| RHOT1 | 0 35807428 | 0 017875119 | 6 86E-09 | -0 18299421 | 0 | 1 01E-08 |
| GANC | 0 220660182 | 0 017859171 | 7 08E-09 | -0 26977169 | 0 | 1 04E-08 |
| LINGO3 | 0 004958977 | 0 017856246 | 7 12E-09 | 0 495439944 | 0 | 1 05E-08 |
| CD96 | 0 023605109 | 0 017845788 | 7 26E-09 | 0 478781097 | 0 | 1 07E-08 |
| ZNF107 | 0 17230778 | 0 017844445 | 7 28E-09 | 0 283120254 | 0 | 1 07E-08 |
| LINC02884 | 0 082808247 | 0 017841781 | 7 32E-09 | -0 39188406 | 0 | 1 08E-08 |
| OTUD4 | 0 288762826 | 0 017836149 | 7 40E-09 | 0 247156611 | 0 | 1 09E-08 |

| | | | | | | |
|---|---|---|---|---|---|---|
| TMEM255A | 0 017484914 | 0 017823526 | 7 58E-09 | 0 486680928 | 0 | 1 12E-08 |
| ALAS1 | 0 053987169 | 0 017804917 | 7 85E-09 | -0 43533393 | 0 | 1 16E-08 |
| ANKEF1 | 0 005463441 | 0 017796893 | 7 98E-09 | 0 494026059 | 0 | 1 17E-08 |
| ADAMTS15 | 0 040233386 | 0 017784927 | 8 16E-09 | -0 44653903 | 0 | 1 20E-08 |
| LINC01226 | 0 005139822 | 0 017749939 | 8 73E-09 | -0 49001737 | 0 | 1 28E-08 |
| CASP10 | 0 043935962 | 0 017739392 | 8 91E-09 | 0 455499862 | 0 | 1 31E-08 |
| FABP3 | 0 024119092 | 0 017734092 | 9 00E-09 | -0 47012429 | 0 | 1 32E-08 |
| ARL13B | 0 284546268 | 0 017732386 | 9 03E-09 | -0 22499686 | 0 | 1 33E-08 |
| DNAJC4 | 0 057699263 | 0 01771851 | 9 27E-09 | -0 42675114 | 0 | 1 36E-08 |
| USP9X | 0 619948221 | 0 017696141 | 9 68E-09 | -0 0365758 | 1 93E-32 | 1 42E-08 |
| OSTN | 0 028040586 | 0 017692673 | 9 74E-09 | 0 474986029 | 0 | 1 43E-08 |
| CTLA4 | 0 012402201 | 0 017692286 | 9 75E-09 | 0 488785417 | 0 | 1 43E-08 |
| NRIP1 | 0 544250062 | 0 017689259 | 9 81E-09 | -0 08505044 | 6 89E-168 | 1 44E-08 |
| FAM110A | 0 017009004 | 0 017687314 | 9 84E-09 | 0 470770417 | 0 | 1 44E-08 |
| SPIB | 0 003578839 | 0 017685328 | 9 88E-09 | 0 495959625 | 0 | 1 45E-08 |
| AGFG1 | 0 435438122 | 0 017684612 | 9 89E-09 | 0 175458552 | 0 | 1 45E-08 |
| KCNC1 | 0 056994917 | 0 017680864 | 9 96E-09 | 0 452137226 | 0 | 1 46E-08 |
| TGM2 | 0 035950201 | 0 017679416 | 9 99E-09 | -0 44977172 | 0 | 1 47E-08 |
| SLC8B1 | 0 100416897 | 0 017665601 | 1 03E-08 | -0 39502022 | 0 | 1 50E-08 |
| SEL1L2 | 0 006900687 | 0 017647676 | 1 06E-08 | -0 48815694 | 0 | 1 56E-08 |
| CMPK1 | 0 261940568 | 0 017641588 | 1 07E-08 | -0 24620593 | 0 | 1 57E-08 |
| DUSP8 | 0 022596181 | 0 017628519 | 1 10E-08 | -0 46909011 | 0 | 1 61E-08 |
| CRTAM | 0 004759095 | 0 017610421 | 1 14E-08 | -0 49055596 | 0 | 1 67E-08 |
| LYPD1 | 0 045792009 | 0 017604699 | 1 15E-08 | 0 454328076 | 0 | 1 69E-08 |
| AP1S3 | 0 093173555 | 0 0175895 | 1 19E-08 | 0 420878095 | 0 | 1 74E-08 |
| MGC4859 | 0 00872818 | 0 017585025 | 1 20E-08 | -0 4856593 | 0 | 1 75E-08 |
| PLA2G7 | 0 032723535 | 0 017584758 | 1 20E-08 | -0 45387597 | 0 | 1 75E-08 |
| CLYBL | 0 155565285 | 0 017549242 | 1 28E-08 | 0 368035983 | 0 | 1 87E-08 |
| MORF4L2 | 0 497553825 | 0 017545116 | 1 29E-08 | -0 06772024 | 4 96E-107 | 1 89E-08 |
| PENK | 0 016495022 | 0 017519991 | 1 35E-08 | -0 47632156 | 0 | 1 98E-08 |
| LINC00467 | 0 114161162 | 0 01751798 | 1 36E-08 | 0 396925918 | 0 | 1 99E-08 |
| SNX29 | 0 482229541 | 0 017513307 | 1 37E-08 | -0 11706457 | 0 | 2 00E-08 |
| TMEM88 | 0 009651444 | 0 017509866 | 1 38E-08 | -0 48435904 | 0 | 2 02E-08 |
| ZNF831 | 0 006481887 | 0 017507855 | 1 38E-08 | 0 492061017 | 0 | 2 02E-08 |
| GPR27 | 0 010803145 | 0 017500139 | 1 41E-08 | -0 48162792 | 0 | 2 05E-08 |
| B3GNTL1 | 0 146875178 | 0 017497471 | 1 41E-08 | -0 35128232 | 0 | 2 06E-08 |
| MAN1A1 | 0 388389713 | 0 017491793 | 1 43E-08 | -0 12969023 | 0 | 2 09E-08 |
| SORCS1 | 0 19112524 | 0 017491747 | 1 43E-08 | -0 29042709 | 0 | 2 09E-08 |
| BARX2 | 0 010298681 | 0 017480963 | 1 46E-08 | -0 48494222 | 0 | 2 13E-08 |
| PRKG2 | 0 022006054 | 0 017466053 | 1 50E-08 | -0 46858971 | 0 | 2 19E-08 |
| ZBTB43 | 0 19170585 | 0 017456075 | 1 53E-08 | -0 30613363 | 0 | 2 23E-08 |
| FHL5 | 0 002151111 | 0 017450012 | 1 55E-08 | 0 49658694 | 0 | 2 25E-08 |
| FAM171A1 | 0 281376711 | 0 017447759 | 1 55E-08 | -0 21453766 | 0 | 2 26E-08 |
| INPP5A | 0 276113152 | 0 017432101 | 1 60E-08 | -0 2265212 | 0 | 2 33E-08 |

| | | | | | | |
|---|---|---|---|---|---|---|
| HTRA4 | 0 016809122 | 0 01742725 | 1 61E-08 | -0 47514514 | 0 | 2 35E-08 |
| LINC01393 | 0 022862691 | 0 017425666 | 1 62E-08 | 0 477184113 | 0 | 2 36E-08 |
| INSL3 | 0 001856047 | 0 01737726 | 1 77E-08 | 0 496533464 | 0 | 2 58E-08 |
| GPR4 | 0 005349222 | 0 017357024 | 1 84E-08 | 0 493658968 | 0 | 2 68E-08 |
| RABGGTA | 0 027041176 | 0 017349593 | 1 87E-08 | -0 46250889 | 0 | 2 72E-08 |
| PPP1R9A | 0 335906417 | 0 017348789 | 1 87E-08 | 0 208912061 | 0 | 2 72E-08 |
| SLPI | 0 012002437 | 0 017333493 | 1 92E-08 | -0 4794478 | 0 | 2 80E-08 |
| LY6G6C | 0 002293884 | 0 017317374 | 1 98E-08 | 0 496129198 | 0 | 2 89E-08 |
| DUT | 0 26413927 | 0 017309915 | 2 01E-08 | 0 275265117 | 0 | 2 93E-08 |
| STAT3 | 0 590394243 | 0 017303342 | 2 04E-08 | -0 00710965 | 0 021196276 | 2 96E-08 |
| CCDC97 | 0 084654775 | 0 017279035 | 2 13E-08 | 0 407148272 | 0 | 3 10E-08 |
| TTC38 | 0 086396604 | 0 017260952 | 2 20E-08 | -0 40182412 | 0 | 3 21E-08 |
| POLR2A | 0 264815062 | 0 017252102 | 2 24E-08 | -0 2272458 | 0 | 3 26E-08 |
| LPAR3 | 0 008185643 | 0 017237334 | 2 30E-08 | 0 492042934 | 0 | 3 35E-08 |
| GADL1 | 0 007110087 | 0 017237262 | 2 30E-08 | 0 493681559 | 0 | 3 35E-08 |
| RWDD1 | 0 160143534 | 0 017235103 | 2 31E-08 | -0 33311736 | 0 | 3 36E-08 |
| TLE4 | 0 364870267 | 0 017222506 | 2 37E-08 | 0 19956654 | 0 | 3 44E-08 |
| SLCO4C1 | 0 006558032 | 0 017218155 | 2 39E-08 | 0 493912251 | 0 | 3 47E-08 |
| STK31 | 0 025175611 | 0 01721366 | 2 41E-08 | 0 481472582 | 0 | 3 50E-08 |
| GPATCH8 | 0 766366526 | 0 017207655 | 2 43E-08 | 0 032875469 | 1 59E-26 | 3 54E-08 |
| SCD5 | 0 217157488 | 0 017206423 | 2 44E-08 | -0 27726516 | 0 | 3 54E-08 |
| THEM4 | 0 139003636 | 0 017188428 | 2 52E-08 | -0 339811 | 0 | 3 66E-08 |
| RND1 | 0 008442634 | 0 017188033 | 2 52E-08 | -0 48486767 | 0 | 3 67E-08 |
| DLL4 | 0 005511032 | 0 017177862 | 2 57E-08 | -0 48972707 | 0 | 3 73E-08 |
| CKM | 0 01319221 | 0 01715746 | 2 67E-08 | -0 48366478 | 0 | 3 88E-08 |
| GTF3A | 0 152195846 | 0 017151979 | 2 70E-08 | -0 34200608 | 0 | 3 92E-08 |
| NALCN-AS1 | 0 017075632 | 0 017141758 | 2 75E-08 | 0 486677464 | 0 | 3 99E-08 |
| SERPINB9 | 0 092602463 | 0 017133791 | 2 79E-08 | -0 39140167 | 0 | 4 05E-08 |
| MCTP2 | 0 270630675 | 0 017131299 | 2 81E-08 | -0 23259704 | 0 | 4 07E-08 |
| TMEM117 | 0 557128172 | 0 017119478 | 2 87E-08 | 0 095736717 | 2 23E-212 | 4 16E-08 |
| CAVIN2-AS1 | 0 027136358 | 0 017096632 | 2 99E-08 | -0 45540271 | 0 | 4 33E-08 |
| SP110 | 0 225533495 | 0 017082532 | 3 07E-08 | -0 28182309 | 0 | 4 45E-08 |
| DEPDC5 | 0 213369249 | 0 017070972 | 3 14E-08 | 0 278522534 | 0 | 4 54E-08 |
| FCHO1 | 0 008652034 | 0 017065453 | 3 17E-08 | 0 491345917 | 0 | 4 59E-08 |
| PAIP2 | 0 321257924 | 0 017030789 | 3 38E-08 | -0 2047589 | 0 | 4 89E-08 |
| TC2N | 0 053539815 | 0 017025874 | 3 41E-08 | -0 43073584 | 0 | 4 93E-08 |
| ZBTB2 | 0 052768841 | 0 017025344 | 3 41E-08 | 0 446534167 | 0 | 4 94E-08 |
| NR2F1-AS1 | 0 365393767 | 0 017018118 | 3 46E-08 | -0 18218759 | 0 | 5 00E-08 |
| ZNF689 | 0 02465211 | 0 017008308 | 3 52E-08 | 0 473829818 | 0 | 5 09E-08 |
| EIF3F | 0 191477413 | 0 016972491 | 3 76E-08 | -0 30854452 | 0 | 5 44E-08 |
| GABARAPL1 | 0 143296339 | 0 016961231 | 3 84E-08 | -0 33851612 | 0 | 5 55E-08 |
| CALCR | 0 011536045 | 0 016917593 | 4 16E-08 | -0 48166162 | 0 | 6 01E-08 |
| ACTR2 | 0 444356666 | 0 016911558 | 4 21E-08 | 0 169331838 | 0 | 6 08E-08 |
| LINC01982 | 0 031371952 | 0 016908711 | 4 23E-08 | -0 45772381 | 0 | 6 11E-08 |

| | | | | | | |
|---|---|---|---|---|---|---|
| WBP1L | 0 288772344 | 0 016895583 | 4 33E-08 | -0 22875809 | 0 | 6 25E-08 |
| KIF21B | 0 015067294 | 0 016891187 | 4 37E-08 | 0 485354648 | 0 | 6 30E-08 |
| POLR2B | 0 375901848 | 0 016889729 | 4 38E-08 | 0 19737191 | 0 | 6 32E-08 |
| LINC01934 | 0 046724791 | 0 01686147 | 4 61E-08 | 0 43814804 | 0 | 6 65E-08 |
| FEZ2 | 0 352506139 | 0 016841584 | 4 78E-08 | 0 221442053 | 0 | 6 90E-08 |
| FAM209A | 0 011393273 | 0 016828286 | 4 90E-08 | 0 488370651 | 0 | 7 07E-08 |
| TBC1D22A | 0 518341551 | 0 016827391 | 4 91E-08 | 0 119136173 | 0 | 7 08E-08 |
| TMTC2 | 0 455245474 | 0 016813303 | 5 04E-08 | 0 133758617 | 0 | 7 26E-08 |
| LINC01814 | 0 005273077 | 0 016812506 | 5 04E-08 | 0 493594173 | 0 | 7 27E-08 |
| HSPA6 | 0 021977499 | 0 016797124 | 5 19E-08 | -0 47010614 | 0 | 7 47E-08 |
| LINC00900 | 0 009213607 | 0 01679201 | 5 23E-08 | -0 48487141 | 0 | 7 54E-08 |
| MYO6 | 0 425558242 | 0 016789126 | 5 26E-08 | -0 09869626 | 1 21E-225 | 7 58E-08 |
| RPS6KB2 | 0 096819021 | 0 01678859 | 5 27E-08 | 0 398082296 | 0 | 7 59E-08 |
| CHI3L1 | 0 028621195 | 0 016777342 | 5 38E-08 | -0 45934626 | 0 | 7 74E-08 |
| CASZ1 | 0 024633074 | 0 016754779 | 5 60E-08 | -0 46879036 | 0 | 8 07E-08 |
| PSEN1 | 0 361186728 | 0 016747715 | 5 67E-08 | -0 18142384 | 0 | 8 17E-08 |
| EPB41L4A-AS1 | 0 117035655 | 0 016735632 | 5 80E-08 | -0 38097644 | 0 | 8 35E-08 |
| SFTA1P | 0 005263559 | 0 016721758 | 5 95E-08 | -0 49068864 | 0 | 8 56E-08 |
| BNIP2 | 0 216919533 | 0 016709214 | 6 08E-08 | -0 28699115 | 0 | 8 75E-08 |
| CD38 | 0 067731435 | 0 016707584 | 6 10E-08 | -0 41536056 | 0 | 8 78E-08 |
| BRCA2 | 0 10581371 | 0 016691524 | 6 28E-08 | 0 324162509 | 0 | 9 03E-08 |
| RNF6 | 0 076250214 | 0 016690489 | 6 29E-08 | -0 41176731 | 0 | 9 05E-08 |
| RIF1 | 0 544821153 | 0 01668442 | 6 36E-08 | 0 084022741 | 6 85E-164 | 9 14E-08 |
| MITD1 | 0 205335897 | 0 016676322 | 6 46E-08 | 0 306220356 | 0 | 9 28E-08 |
| STXBP2 | 0 014953075 | 0 016650644 | 6 76E-08 | 0 484877665 | 0 | 9 72E-08 |
| HPN | 0 003493175 | 0 016641375 | 6 88E-08 | 0 495439054 | 0 | 9 88E-08 |
| OS9 | 0 297405342 | 0 016634355 | 6 97E-08 | 0 248127571 | 0 | 1 00E-07 |
| CCDC18 | 0 246958939 | 0 016628986 | 7 03E-08 | 0 256375893 | 0 | 1 01E-07 |
| WRAP53 | 0 030562906 | 0 016614467 | 7 22E-08 | 0 471923178 | 0 | 1 04E-07 |
| PAM | 0 709190764 | 0 016612266 | 7 25E-08 | 0 018016938 | 5 21E-09 | 1 04E-07 |
| NLRP12 | 0 003578839 | 0 016585612 | 7 61E-08 | 0 494855693 | 0 | 1 09E-07 |
| TDRD12 | 0 007100569 | 0 016584142 | 7 63E-08 | -0 48755646 | 0 | 1 09E-07 |
| VGLL3 | 0 154680094 | 0 01658249 | 7 65E-08 | -0 32246763 | 0 | 1 10E-07 |
| SH2D2A | 0 008613961 | 0 01655555 | 8 03E-08 | 0 490551092 | 0 | 1 15E-07 |
| NEB | 0 152462356 | 0 016542081 | 8 22E-08 | 0 308685539 | 0 | 1 18E-07 |
| SYNE4 | 0 006681769 | 0 016540592 | 8 25E-08 | 0 492172952 | 0 | 1 18E-07 |
| PLPPR5 | 0 035445737 | 0 016537139 | 8 30E-08 | -0 45432637 | 0 | 1 19E-07 |
| LINC02223 | 0 017380214 | 0 016501845 | 8 84E-08 | -0 47351353 | 0 | 1 27E-07 |
| PRKX-AS1 | 0 049323257 | 0 01648438 | 9 12E-08 | -0 44255935 | 0 | 1 31E-07 |
| SRSF11 | 0 740991034 | 0 016465718 | 9 43E-08 | 0 039827766 | 3 73E-38 | 1 35E-07 |
| CPED1 | 0 574413204 | 0 016460325 | 9 52E-08 | -0 05700923 | 2 33E-76 | 1 36E-07 |
| CD247 | 0 019559879 | 0 016450752 | 9 69E-08 | -0 47343822 | 0 | 1 38E-07 |
| NWD2 | 0 00977518 | 0 016427518 | 1 01E-07 | -0 48484964 | 0 | 1 44E-07 |
| KCNAB1 | 0 025937066 | 0 016422652 | 1 02E-07 | -0 47005031 | 0 | 1 46E-07 |

| | | | | | | |
|---|---|---|---|---|---|---|
| CDCA7 | 0 067483962 | 0 016422226 | 1 02E-07 | 0 380222498 | 0 | 1 46E-07 |
| PSPN | 0 007376597 | 0 016421397 | 1 02E-07 | 0 491980619 | 0 | 1 46E-07 |
| VAV3-AS1 | 0 002779311 | 0 016418086 | 1 03E-07 | -0 49297985 | 0 | 1 47E-07 |
| OGFRL1 | 0 146266014 | 0 016397618 | 1 06E-07 | -0 34081554 | 0 | 1 52E-07 |
| GPR108 | 0 084550075 | 0 016385832 | 1 09E-07 | -0 40464135 | 0 | 1 55E-07 |
| ERN1 | 0 223163465 | 0 016385683 | 1 09E-07 | -0 27073563 | 0 | 1 55E-07 |
| GTF2F2 | 0 357741143 | 0 016383649 | 1 09E-07 | -0 1815468 | 0 | 1 56E-07 |
| CRAT37 | 0 004483067 | 0 016374936 | 1 11E-07 | -0 49040195 | 0 | 1 58E-07 |
| PLVAP | 0 023690773 | 0 016353692 | 1 15E-07 | 0 476535019 | 0 | 1 64E-07 |
| TP53BP2 | 0 249319449 | 0 016342127 | 1 18E-07 | -0 25136182 | 0 | 1 68E-07 |
| ZNF80 | 0 001066037 | 0 016330012 | 1 20E-07 | 0 497105604 | 0 | 1 71E-07 |
| LINC01356 | 0 004673431 | 0 016326131 | 1 21E-07 | 0 494385056 | 0 | 1 72E-07 |
| GSPT1 | 0 393224953 | 0 016307091 | 1 25E-07 | 0 189262682 | 0 | 1 78E-07 |
| LINC02296 | 0 012335573 | 0 016287707 | 1 29E-07 | -0 48275737 | 0 | 1 84E-07 |
| RAB34 | 0 170499324 | 0 016287661 | 1 29E-07 | -0 32681914 | 0 | 1 84E-07 |
| MTMR14 | 0 16826255 | 0 016287306 | 1 30E-07 | 0 347408191 | 0 | 1 85E-07 |
| WDTC1 | 0 183653462 | 0 01626634 | 1 34E-07 | -0 30789366 | 0 | 1 91E-07 |
| GOLGA7B | 0 020625916 | 0 016253345 | 1 38E-07 | -0 47044231 | 0 | 1 96E-07 |
| LINC02763 | 0 013829929 | 0 01623367 | 1 42E-07 | -0 47879816 | 0 | 2 03E-07 |
| RIMS2 | 0 195160953 | 0 016222891 | 1 45E-07 | 0 33664297 | 0 | 2 07E-07 |
| A1BG | 0 113894653 | 0 016216801 | 1 47E-07 | -0 37638266 | 0 | 2 09E-07 |
| PPP2R3A | 0 492785213 | 0 016192569 | 1 53E-07 | -0 07189683 | 2 00E-120 | 2 18E-07 |
| IL27RA | 0 056804554 | 0 016191305 | 1 53E-07 | -0 43602159 | 0 | 2 18E-07 |
| SLC24A2 | 0 030010851 | 0 016187548 | 1 54E-07 | -0 45916478 | 0 | 2 20E-07 |
| PACSIN1 | 0 002864975 | 0 016174131 | 1 58E-07 | -0 49350266 | 0 | 2 25E-07 |
| LINC01931 | 0 021596771 | 0 016156978 | 1 63E-07 | -0 47304815 | 0 | 2 32E-07 |
| EXOC3L1 | 0 003997639 | 0 016155956 | 1 63E-07 | -0 49121736 | 0 | 2 32E-07 |
| WAS | 0 012659192 | 0 016153671 | 1 64E-07 | -0 48213842 | 0 | 2 33E-07 |
| CCDC88B | 0 015390912 | 0 016152902 | 1 64E-07 | 0 485782903 | 0 | 2 33E-07 |
| DLEU7 | 0 096324075 | 0 016117816 | 1 75E-07 | -0 37625808 | 0 | 2 48E-07 |
| ALCAM | 0 283527822 | 0 016111438 | 1 77E-07 | 0 208655687 | 0 | 2 51E-07 |
| CDT1 | 0 017513468 | 0 016100015 | 1 80E-07 | 0 46867856 | 0 | 2 56E-07 |
| STK4 | 0 305886048 | 0 01609872 | 1 80E-07 | 0 23254044 | 0 | 2 56E-07 |
| NMUR1 | 0 008299861 | 0 016089154 | 1 84E-07 | 0 492004877 | 0 | 2 61E-07 |
| RASSF3 | 0 186413737 | 0 01608852 | 1 84E-07 | 0 280287637 | 0 | 2 61E-07 |
| CHID1 | 0 137566389 | 0 016081149 | 1 86E-07 | -0 34989956 | 0 | 2 64E-07 |
| SVIP | 0 121537759 | 0 016076057 | 1 88E-07 | 0 39408369 | 0 | 2 66E-07 |
| CYSLTR2 | 0 030029887 | 0 016075949 | 1 88E-07 | -0 45585524 | 0 | 2 66E-07 |
| NDUFAF2 | 0 37284651 | 0 016067135 | 1 91E-07 | -0 17601928 | 0 | 2 71E-07 |
| BLNK | 0 022415336 | 0 0160605 | 1 93E-07 | -0 47079135 | 0 | 2 74E-07 |
| PLEKHF2 | 0 10923074 | 0 01603196 | 2 03E-07 | 0 400165011 | 0 | 2 88E-07 |
| NLRC3 | 0 017789496 | 0 016021613 | 2 06E-07 | -0 47550409 | 0 | 2 93E-07 |
| ARL14EP | 0 150396908 | 0 016012179 | 2 10E-07 | -0 344562 | 0 | 2 98E-07 |
| LINC02306 | 0 011555082 | 0 016010427 | 2 11E-07 | 0 488743735 | 0 | 2 98E-07 |

| | | | | | | |
|---|---|---|---|---|---|---|
| CLEC9A | 0 006301041 | 0 01599188 | 2 17E-07 | 0 492883161 | 0 | 3 08E-07 |
| GUCY2D | 0 003597876 | 0 015987155 | 2 19E-07 | 0 495132988 | 0 | 3 11E-07 |
| PPIP5K1 | 0 086548895 | 0 015987015 | 2 19E-07 | -0 40012776 | 0 | 3 11E-07 |
| SLC29A1 | 0 099369896 | 0 015978978 | 2 22E-07 | 0 404339555 | 0 | 3 15E-07 |
| GPSM3 | 0 013392092 | 0 015974931 | 2 24E-07 | -0 48109029 | 0 | 3 17E-07 |
| ARSG | 0 145333232 | 0 015971304 | 2 25E-07 | -0 34737555 | 0 | 3 19E-07 |
| MARK2 | 0 224229503 | 0 015967197 | 2 27E-07 | 0 290753578 | 0 | 3 21E-07 |
| TMEM74B | 0 007452742 | 0 015962043 | 2 29E-07 | 0 494148079 | 0 | 3 24E-07 |
| GEMIN4 | 0 021244598 | 0 015954051 | 2 32E-07 | -0 47137591 | 0 | 3 28E-07 |
| KIF23 | 0 081275818 | 0 015903229 | 2 54E-07 | 0 363470539 | 0 | 3 58E-07 |
| POLA2 | 0 125078525 | 0 015897438 | 2 56E-07 | 0 317523564 | 0 | 3 62E-07 |
| AK9 | 0 11537949 | 0 015867842 | 2 70E-07 | 0 41571953 | 0 | 3 81E-07 |
| TIE1 | 0 013572938 | 0 015858803 | 2 74E-07 | -0 48037789 | 0 | 3 87E-07 |
| LINC02207 | 0 018674687 | 0 015853058 | 2 77E-07 | -0 47702033 | 0 | 3 91E-07 |
| ORC6 | 0 064190668 | 0 015829638 | 2 88E-07 | 0 378899857 | 0 | 4 07E-07 |
| CD302 | 0 298547524 | 0 01582611 | 2 90E-07 | -0 21653201 | 0 | 4 09E-07 |
| MBNL2 | 0 60681312 | 0 015813565 | 2 96E-07 | 0 014408738 | 3 00E-06 | 4 18E-07 |
| BACE2 | 0 32248577 | 0 015798175 | 3 04E-07 | 0 235922387 | 0 | 4 29E-07 |
| CYRIB | 0 180874151 | 0 015792233 | 3 07E-07 | -0 29840249 | 0 | 4 33E-07 |
| AMN | 0 007195751 | 0 01578047 | 3 13E-07 | -0 48860959 | 0 | 4 42E-07 |
| RAB39A | 0 018246369 | 0 015774384 | 3 17E-07 | 0 483723159 | 0 | 4 47E-07 |
| IL18RAP | 0 00441644 | 0 015761859 | 3 24E-07 | -0 49028799 | 0 | 4 56E-07 |
| CCDC77 | 0 108431212 | 0 015749078 | 3 31E-07 | 0 341960795 | 0 | 4 66E-07 |
| PITPNM1 | 0 038719994 | 0 015728408 | 3 43E-07 | 0 461389391 | 0 | 4 83E-07 |
| NOTCH1 | 0 094068264 | 0 015721138 | 3 47E-07 | 0 424276341 | 0 | 4 89E-07 |
| SNX24 | 0 28274733 | 0 015719332 | 3 48E-07 | 0 251398406 | 0 | 4 90E-07 |
| TNFSF4 | 0 012145209 | 0 01570602 | 3 56E-07 | -0 48180757 | 0 | 5 02E-07 |
| GPATCH11 | 0 106365765 | 0 015705327 | 3 56E-07 | 0 390538595 | 0 | 5 02E-07 |
| MICAL3 | 0 328644039 | 0 015704888 | 3 57E-07 | 0 22659351 | 0 | 5 02E-07 |
| LYRM7 | 0 114827435 | 0 015702034 | 3 58E-07 | 0 397734016 | 0 | 5 05E-07 |
| EIF4A3 | 0 114694181 | 0 015696596 | 3 62E-07 | 0 402949301 | 0 | 5 09E-07 |
| BCAR3 | 0 179751004 | 0 015693278 | 3 64E-07 | 0 326163023 | 0 | 5 12E-07 |
| FAM228B | 0 216129524 | 0 015692389 | 3 64E-07 | -0 28303788 | 0 | 5 13E-07 |
| PTEN | 0 700529211 | 0 015667808 | 3 80E-07 | 0 00141711 | 0 645999838 | 5 35E-07 |
| CCNL1 | 0 514857893 | 0 015664073 | 3 82E-07 | -0 0760081 | 2 13E-134 | 5 38E-07 |
| ZNF366 | 0 004359331 | 0 015628269 | 4 07E-07 | -0 49174948 | 0 | 5 72E-07 |
| FBL | 0 120309912 | 0 015605144 | 4 23E-07 | 0 380296386 | 0 | 5 95E-07 |
| NCAPH | 0 047314919 | 0 015597279 | 4 29E-07 | 0 400842584 | 0 | 6 03E-07 |
| MTTP | 0 00493994 | 0 015595856 | 4 30E-07 | 0 493349407 | 0 | 6 04E-07 |
| NT5C | 0 209124136 | 0 015552059 | 4 63E-07 | -0 29604154 | 0 | 6 50E-07 |
| LMNA | 0 513525347 | 0 015549745 | 4 64E-07 | -0 05343486 | 2 69E-67 | 6 52E-07 |
| RADX | 0 071053283 | 0 015539499 | 4 73E-07 | 0 421773205 | 0 | 6 64E-07 |
| LINC01622 | 0 024376083 | 0 01551895 | 4 89E-07 | -0 46417923 | 0 | 6 87E-07 |
| ADORA2A-AS1 | 0 019559879 | 0 015517896 | 4 90E-07 | -0 47203412 | 0 | 6 88E-07 |

| | | | | | | |
|---|---|---|---|---|---|---|
| LINC00840 | 0 006719842 | 0 015506626 | 5 00E-07 | 0 492098004 | 0 | 7 01E-07 |
| SLC39A14 | 0 142182711 | 0 015500585 | 5 05E-07 | -0 34546744 | 0 | 7 08E-07 |
| TOR2A | 0 021453999 | 0 015484848 | 5 18E-07 | 0 480666697 | 0 | 7 27E-07 |
| ALKBH5 | 0 138261217 | 0 015482822 | 5 20E-07 | -0 33903403 | 0 | 7 30E-07 |
| LINC00513 | 0 535150673 | 0 015479244 | 5 23E-07 | 0 096113289 | 4 82E-214 | 7 34E-07 |
| TIFA | 0 019302888 | 0 015476603 | 5 26E-07 | 0 479580011 | 0 | 7 37E-07 |
| ZNFX1 | 0 228103406 | 0 0154459 | 5 54E-07 | -0 27370323 | 0 | 7 76E-07 |
| STK40 | 0 141002456 | 0 015438333 | 5 61E-07 | -0 3494287 | 0 | 7 86E-07 |
| CD200R1 | 0 012554492 | 0 015437597 | 5 61E-07 | -0 48416383 | 0 | 7 87E-07 |
| CADM3-AS1 | 0 001027964 | 0 015429558 | 5 69E-07 | 0 496940145 | 0 | 7 97E-07 |
| CTDSP1 | 0 152747901 | 0 015420333 | 5 78E-07 | -0 33270227 | 0 | 8 09E-07 |
| HEATR9 | 0 003816794 | 0 015414083 | 5 84E-07 | 0 495077038 | 0 | 8 18E-07 |
| HAR1B | 0 005682359 | 0 01538359 | 6 15E-07 | -0 49015385 | 0 | 8 60E-07 |
| POLE4 | 0 080276408 | 0 015357531 | 6 42E-07 | 0 417992608 | 0 | 8 98E-07 |
| ACSM3 | 0 058651082 | 0 015354514 | 6 45E-07 | 0 43984125 | 0 | 9 03E-07 |
| DCSTAMP | 0 004692467 | 0 015343898 | 6 57E-07 | -0 49111794 | 0 | 9 18E-07 |
| BRI3BP | 0 060745084 | 0 015282923 | 7 27E-07 | 0 444456851 | 0 | 1 02E-06 |
| FNDC3B | 0 847223544 | 0 015279631 | 7 31E-07 | -0 00753814 | 0 014551144 | 1 02E-06 |
| NMNAT2 | 0 054015724 | 0 015279422 | 7 32E-07 | -0 42580152 | 0 | 1 02E-06 |
| FKBPL | 0 010089281 | 0 015273676 | 7 39E-07 | 0 488803552 | 0 | 1 03E-06 |
| LNC-LBCS | 0 046953228 | 0 015261897 | 7 53E-07 | 0 47149354 | 0 | 1 05E-06 |
| CDS2 | 0 229464507 | 0 015257005 | 7 59E-07 | -0 27496489 | 0 | 1 06E-06 |
| MLIP | 0 060354838 | 0 015211444 | 8 19E-07 | -0 43835178 | 0 | 1 14E-06 |
| TTC39C-AS1 | 0 005710914 | 0 015200794 | 8 34E-07 | -0 48915185 | 0 | 1 16E-06 |
| KLHL32 | 0 010546154 | 0 015195411 | 8 41E-07 | 0 490131606 | 0 | 1 17E-06 |
| PLXDC1 | 0 225333613 | 0 015194777 | 8 42E-07 | -0 25595654 | 0 | 1 17E-06 |
| MGAT1 | 0 156774095 | 0 01518724 | 8 53E-07 | -0 33500846 | 0 | 1 19E-06 |
| LINC02611 | 0 003569321 | 0 015168651 | 8 79E-07 | -0 49266169 | 0 | 1 23E-06 |
| SYNDIG1L | 0 005158859 | 0 015166939 | 8 82E-07 | -0 48988241 | 0 | 1 23E-06 |
| TSPAN33 | 0 016161885 | 0 015155736 | 8 98E-07 | -0 47739162 | 0 | 1 25E-06 |
| UNC45B | 0 001399174 | 0 015147703 | 9 10E-07 | -0 49507483 | 0 | 1 27E-06 |
| DOCK7 | 0 556890217 | 0 015146515 | 9 12E-07 | -0 0642745 | 1 38E-96 | 1 27E-06 |
| SLC9A3R2 | 0 015819231 | 0 015140506 | 9 21E-07 | 0 478570196 | 0 | 1 28E-06 |
| PCAT19 | 0 008347452 | 0 015132701 | 9 33E-07 | 0 490042586 | 0 | 1 30E-06 |
| ZMYND15 | 0 011754964 | 0 015107325 | 9 73E-07 | -0 48247037 | 0 | 1 35E-06 |
| NCOA3 | 0 477413337 | 0 015104078 | 9 78E-07 | -0 12108966 | 0 | 1 36E-06 |
| GFRA1 | 0 182159106 | 0 015100928 | 9 83E-07 | -0 32072767 | 0 | 1 37E-06 |
| CXCL6 | 0 012630637 | 0 01507741 | 1 02E-06 | 0 486541392 | 0 | 1 42E-06 |
| SLC8A1-AS1 | 0 110258704 | 0 015077141 | 1 02E-06 | 0 337773182 | 0 | 1 42E-06 |
| KMT2E-AS1 | 0 109468695 | 0 01506483 | 1 04E-06 | -0 37425549 | 0 | 1 45E-06 |
| FOXP3 | 0 003683539 | 0 015064014 | 1 05E-06 | 0 495647725 | 0 | 1 45E-06 |
| ACER1 | 0 005996459 | 0 015060481 | 1 05E-06 | -0 48911919 | 0 | 1 46E-06 |
| TMEM255B | 0 023747882 | 0 015059198 | 1 05E-06 | 0 480306915 | 0 | 1 46E-06 |
| SASS6 | 0 101635225 | 0 015054372 | 1 06E-06 | 0 354881799 | 0 | 1 47E-06 |

| | | | | | | |
|---|---|---|---|---|---|---|
| MED29 | 0 087900478 | 0 015035876 | 1 09E-06 | -0 39622081 | 0 | 1 52E-06 |
| C22orf34 | 0 005549104 | 0 015027408 | 1 11E-06 | 0 491455887 | 0 | 1 54E-06 |
| PRKAB1 | 0 058222764 | 0 015026335 | 1 11E-06 | -0 43208656 | 0 | 1 54E-06 |
| GGTA1 | 0 020207116 | 0 015005517 | 1 15E-06 | -0 47074016 | 0 | 1 60E-06 |
| SIPA1L1 | 0 626096971 | 0 015003841 | 1 15E-06 | 0 08703638 | 9 40E-176 | 1 60E-06 |
| EDRF1 | 0 267889437 | 0 014994465 | 1 17E-06 | 0 249772325 | 0 | 1 62E-06 |
| KCNK7 | 0 003150521 | 0 014992412 | 1 18E-06 | 0 496164925 | 0 | 1 63E-06 |
| GIPC2 | 0 053140051 | 0 014981517 | 1 20E-06 | -0 43879073 | 0 | 1 66E-06 |
| ITPKC | 0 185709391 | 0 014905893 | 1 35E-06 | -0 28118385 | 0 | 1 88E-06 |
| APLN | 0 007976243 | 0 014891478 | 1 39E-06 | 0 491221485 | 0 | 1 92E-06 |
| PTGER2 | 0 005739468 | 0 014890277 | 1 39E-06 | -0 4889973 | 0 | 1 92E-06 |
| FAM209B | 0 00289353 | 0 014873165 | 1 43E-06 | -0 49289033 | 0 | 1 98E-06 |
| RBM11 | 0 021482553 | 0 014845495 | 1 49E-06 | -0 46937357 | 0 | 2 07E-06 |
| ATP6V0A1 | 0 403713997 | 0 014840717 | 1 51E-06 | -0 11904156 | 0 | 2 08E-06 |
| HIPK3 | 0 544402353 | 0 014838659 | 1 51E-06 | -0 03980226 | 4 15E-38 | 2 09E-06 |
| SCHLAP1 | 0 052968723 | 0 014825481 | 1 54E-06 | 0 443059553 | 0 | 2 13E-06 |
| NFKBIB | 0 048790238 | 0 01482402 | 1 55E-06 | -0 44007829 | 0 | 2 14E-06 |
| ADGRE2 | 0 023053054 | 0 01480414 | 1 60E-06 | -0 46977888 | 0 | 2 21E-06 |
| TMPO-AS1 | 0 013011365 | 0 01479541 | 1 62E-06 | 0 480930932 | 0 | 2 24E-06 |
| RAB33B-AS1 | 0 133730559 | 0 01479486 | 1 62E-06 | -0 34295212 | 0 | 2 24E-06 |
| IER5 | 0 099160496 | 0 014781883 | 1 66E-06 | 0 404260162 | 0 | 2 29E-06 |
| ACKR2 | 0 00541585 | 0 014776351 | 1 67E-06 | -0 48968036 | 0 | 2 31E-06 |
| U2AF1L4 | 0 045230435 | 0 01475833 | 1 72E-06 | -0 44169152 | 0 | 2 37E-06 |
| SEL1L | 0 29893777 | 0 014757442 | 1 72E-06 | -0 19873627 | 0 | 2 38E-06 |
| GPR146 | 0 013677638 | 0 01474574 | 1 75E-06 | -0 48049142 | 0 | 2 42E-06 |
| TNFSF14 | 0 003816794 | 0 014732885 | 1 79E-06 | -0 49092643 | 0 | 2 47E-06 |
| LINC01331 | 0 011022063 | 0 014732457 | 1 79E-06 | -0 48304291 | 0 | 2 47E-06 |
| ADHFE1 | 0 107469875 | 0 014732457 | 1 79E-06 | -0 37664946 | 0 | 2 47E-06 |
| RIC1 | 0 52013097 | 0 014728648 | 1 80E-06 | 0 108512044 | 1 37E-272 | 2 49E-06 |
| IGFBP5 | 0 451914108 | 0 014708262 | 1 86E-06 | -0 12927497 | 0 | 2 57E-06 |
| TMEM72-AS1 | 0 082284746 | 0 014707309 | 1 87E-06 | -0 40067336 | 0 | 2 57E-06 |
| ANKRD13D | 0 224743485 | 0 01469934 | 1 89E-06 | 0 262581709 | 0 | 2 61E-06 |
| LARP1B | 0 36390893 | 0 014694616 | 1 91E-06 | 0 224761479 | 0 | 2 63E-06 |
| SNX6 | 0 411081076 | 0 014690025 | 1 92E-06 | -0 1461084 | 0 | 2 65E-06 |
| NT5C2 | 0 563933677 | 0 014687163 | 1 93E-06 | 0 103890885 | 6 35E-250 | 2 66E-06 |
| SRRM1 | 0 493470522 | 0 014684372 | 1 94E-06 | 0 126051058 | 0 | 2 67E-06 |
| LSR | 0 0231292 | 0 014673177 | 1 97E-06 | -0 46730967 | 0 | 2 72E-06 |
| BEX3 | 0 272039367 | 0 014664904 | 2 00E-06 | -0 22743973 | 0 | 2 75E-06 |
| GTPBP1 | 0 110372923 | 0 0146628 | 2 01E-06 | 0 394534699 | 0 | 2 76E-06 |
| IQGAP3 | 0 053187642 | 0 014662099 | 2 01E-06 | 0 393082595 | 0 | 2 76E-06 |
| GABRD | 0 009127944 | 0 014654169 | 2 03E-06 | -0 48347303 | 0 | 2 80E-06 |
| CFLAR-AS1 | 0 116778664 | 0 014649428 | 2 05E-06 | -0 37889685 | 0 | 2 82E-06 |
| TMED6 | 0 010841218 | 0 014632381 | 2 11E-06 | -0 4842949 | 0 | 2 90E-06 |
| KDELR2 | 0 280510556 | 0 014622785 | 2 14E-06 | -0 23508676 | 0 | 2 94E-06 |

| | | | | | | |
|---|---|---|---|---|---|---|
| POU2AF1 | 0 004949458 | 0 01462154 | 2 14E-06 | -0 49042152 | 0 | 2 95E-06 |
| CASR | 0 005672841 | 0 014616387 | 2 16E-06 | 0 491293783 | 0 | 2 97E-06 |
| ODC1 | 0 101682816 | 0 014599416 | 2 22E-06 | 0 396273039 | 0 | 3 05E-06 |
| CDKN1A | 0 136100588 | 0 014589289 | 2 26E-06 | -0 35220957 | 0 | 3 10E-06 |
| ZNF669 | 0 026689003 | 0 014583209 | 2 28E-06 | 0 475532197 | 0 | 3 13E-06 |
| XIRP2 | 0 026898403 | 0 014580847 | 2 29E-06 | 0 469279126 | 0 | 3 14E-06 |
| SLC25A25 | 0 12962822 | 0 014580084 | 2 29E-06 | 0 391791438 | 0 | 3 14E-06 |
| APOC4 | 0 001380137 | 0 014577958 | 2 30E-06 | -0 49503921 | 0 | 3 15E-06 |
| GRID2 | 0 119967257 | 0 01457186 | 2 32E-06 | -0 35800557 | 0 | 3 19E-06 |
| ASRGL1 | 0 023766919 | 0 014566698 | 2 34E-06 | 0 471421577 | 0 | 3 21E-06 |
| IL24 | 0 000942301 | 0 014522127 | 2 51E-06 | -0 49550459 | 0 | 3 45E-06 |
| HSD17B7 | 0 071519674 | 0 014516453 | 2 53E-06 | 0 436035005 | 0 | 3 48E-06 |
| KBTBD8 | 0 010108317 | 0 01450633 | 2 57E-06 | 0 48895452 | 0 | 3 53E-06 |
| SAMD9L | 0 190192458 | 0 014504136 | 2 58E-06 | 0 310697973 | 0 | 3 54E-06 |
| ESR2 | 0 201252594 | 0 014503894 | 2 58E-06 | 0 306208337 | 0 | 3 54E-06 |
| BMERB1 | 0 434867031 | 0 014501454 | 2 59E-06 | -0 10928511 | 1 76E-276 | 3 56E-06 |
| RASSF2 | 0 210475719 | 0 014489463 | 2 64E-06 | -0 26345403 | 0 | 3 63E-06 |
| CITED1 | 0 011973882 | 0 014484847 | 2 66E-06 | -0 48372618 | 0 | 3 65E-06 |
| ABT1 | 0 045116217 | 0 01448055 | 2 68E-06 | -0 44540945 | 0 | 3 68E-06 |
| ALKBH6 | 0 033218481 | 0 014479904 | 2 68E-06 | 0 470352133 | 0 | 3 68E-06 |
| EAF1 | 0 070548819 | 0 01446782 | 2 74E-06 | -0 42060014 | 0 | 3 75E-06 |
| CALCB | 0 013506311 | 0 014444868 | 2 84E-06 | 0 482299632 | 0 | 3 89E-06 |
| GJC1 | 0 178323276 | 0 014444269 | 2 84E-06 | 0 274557133 | 0 | 3 89E-06 |
| JADE2 | 0 095838648 | 0 014443647 | 2 84E-06 | 0 407281976 | 0 | 3 89E-06 |
| PLPPR1 | 0 027688413 | 0 01442565 | 2 93E-06 | 0 478327348 | 0 | 4 00E-06 |
| BRICD5 | 0 024100055 | 0 0144239 | 2 93E-06 | 0 472112517 | 0 | 4 01E-06 |
| TXNDC11 | 0 249900059 | 0 014410351 | 3 00E-06 | 0 275968771 | 0 | 4 10E-06 |
| CCDC88C | 0 043203061 | 0 014404578 | 3 02E-06 | 0 442227084 | 0 | 4 14E-06 |
| LINC02649 | 0 071091356 | 0 014378238 | 3 15E-06 | -0 40477884 | 0 | 4 31E-06 |
| LINC02269 | 0 007528888 | 0 014368971 | 3 20E-06 | 0 489858201 | 0 | 4 37E-06 |
| ACSM1 | 0 007014905 | 0 014360593 | 3 24E-06 | -0 48725233 | 0 | 4 43E-06 |
| FAM78A | 0 017741905 | 0 01435104 | 3 29E-06 | -0 47835171 | 0 | 4 50E-06 |
| PLEKHA6 | 0 142373075 | 0 014329509 | 3 40E-06 | -0 34176261 | 0 | 4 65E-06 |
| SPCS3 | 0 239125469 | 0 014308995 | 3 51E-06 | 0 310439424 | 0 | 4 80E-06 |
| PPP1R16B | 0 011545564 | 0 014288864 | 3 63E-06 | 0 48784696 | 0 | 4 95E-06 |
| ZRSR2 | 0 192952733 | 0 014279522 | 3 68E-06 | -0 31173367 | 0 | 5 03E-06 |
| HOXB2 | 0 095305629 | 0 014275905 | 3 70E-06 | 0 399891098 | 0 | 5 05E-06 |
| EFHC2 | 0 050941349 | 0 01427133 | 3 73E-06 | -0 43620319 | 0 | 5 09E-06 |
| TNFSF9 | 0 033256553 | 0 014256464 | 3 82E-06 | -0 46084892 | 0 | 5 21E-06 |
| CLDN19 | 0 002769793 | 0 014240944 | 3 91E-06 | -0 49293772 | 0 | 5 33E-06 |
| NOX4 | 0 112029088 | 0 014233334 | 3 96E-06 | 0 386810783 | 0 | 5 40E-06 |
| CD6 | 0 00746226 | 0 014230027 | 3 98E-06 | 0 491462134 | 0 | 5 42E-06 |
| KANK1 | 0 339980202 | 0 014214792 | 4 07E-06 | -0 16426 | 0 | 5 55E-06 |
| MGME1 | 0 160314862 | 0 014213587 | 4 08E-06 | 0 304756918 | 0 | 5 56E-06 |

| | | | | | | |
|---|---|---|---|---|---|---|
| POLN | 0 062134739 | 0 014212023 | 4 09E-06 | -0 42348201 | 0 | 5 58E-06 |
| PCYOX1L | 0 032342807 | 0 014211263 | 4 09E-06 | 0 476239106 | 0 | 5 58E-06 |
| JAK2 | 0 337172336 | 0 014191902 | 4 22E-06 | 0 194986523 | 0 | 5 75E-06 |
| GP6-AS1 | 0 013230283 | 0 014142538 | 4 56E-06 | 0 488438199 | 0 | 6 20E-06 |
| C2CD2L | 0 025346938 | 0 014138227 | 4 59E-06 | -0 46759378 | 0 | 6 25E-06 |
| DNAH7 | 0 128229046 | 0 014122279 | 4 70E-06 | -0 34373093 | 0 | 6 40E-06 |
| TIAM1 | 0 455883193 | 0 014118121 | 4 73E-06 | -0 09038749 | 1 93E-189 | 6 44E-06 |
| SEM1 | 0 44701224 | 0 01411396 | 4 76E-06 | 0 170631697 | 0 | 6 48E-06 |
| RGS3 | 0 117720965 | 0 014107006 | 4 81E-06 | 0 386819864 | 0 | 6 55E-06 |
| SLC25A19 | 0 046134663 | 0 014096996 | 4 89E-06 | 0 450153503 | 0 | 6 65E-06 |
| ARIH2OS | 0 017246959 | 0 014080627 | 5 01E-06 | -0 47575929 | 0 | 6 82E-06 |
| ZFAS1 | 0 309255487 | 0 014068882 | 5 11E-06 | 0 230107094 | 0 | 6 94E-06 |
| TBX6 | 0 008851916 | 0 014056488 | 5 21E-06 | 0 489573398 | 0 | 7 07E-06 |
| FBXW7 | 0 60287259 | 0 014033865 | 5 39E-06 | 0 111552326 | 4 71E-288 | 7 32E-06 |
| MLLT11 | 0 047057928 | 0 014030847 | 5 42E-06 | 0 459901513 | 0 | 7 36E-06 |
| ZC3H3 | 0 181264396 | 0 014027683 | 5 44E-06 | 0 304292079 | 0 | 7 39E-06 |
| CHPT1 | 0 326473892 | 0 014021503 | 5 49E-06 | -0 19566359 | 0 | 7 46E-06 |
| YPEL5 | 0 16836725 | 0 014018195 | 5 52E-06 | -0 31596481 | 0 | 7 50E-06 |
| GNG12 | 0 359168872 | 0 014011535 | 5 58E-06 | -0 17593759 | 0 | 7 57E-06 |
| LRFN1 | 0 004844758 | 0 014008966 | 5 60E-06 | -0 49031745 | 0 | 7 60E-06 |
| CCDC83 | 0 007052978 | 0 014000168 | 5 68E-06 | 0 49203206 | 0 | 7 70E-06 |
| SKA1 | 0 023091127 | 0 013999145 | 5 69E-06 | 0 448206923 | 0 | 7 72E-06 |
| MEPCE | 0 046191772 | 0 013998408 | 5 69E-06 | 0 458579352 | 0 | 7 72E-06 |
| GPAT2 | 0 017561059 | 0 013987537 | 5 79E-06 | -0 47486573 | 0 | 7 85E-06 |
| RASL11A | 0 009537226 | 0 01398001 | 5 86E-06 | -0 48362579 | 0 | 7 94E-06 |
| SLC52A1 | 0 026774666 | 0 013961121 | 6 03E-06 | 0 47747289 | 0 | 8 17E-06 |
| RRM2 | 0 0927262 | 0 013926138 | 6 36E-06 | 0 332561254 | 0 | 8 62E-06 |
| CITED4 | 0 017846605 | 0 013907496 | 6 54E-06 | -0 47397836 | 0 | 8 86E-06 |
| ATL2 | 0 379385506 | 0 01389337 | 6 69E-06 | 0 184166291 | 0 | 9 05E-06 |
| KIAA0040 | 0 071024728 | 0 013890512 | 6 71E-06 | -0 40914888 | 0 | 9 09E-06 |
| RAD51AP1 | 0 079257962 | 0 013867714 | 6 95E-06 | 0 341307311 | 0 | 9 41E-06 |
| KBTBD7 | 0 040147722 | 0 013862446 | 7 01E-06 | -0 44989995 | 0 | 9 48E-06 |
| TSPAN14 | 0 207667853 | 0 013846567 | 7 18E-06 | -0 28017482 | 0 | 9 71E-06 |
| LYNX1 | 0 022767509 | 0 013837231 | 7 28E-06 | -0 46437684 | 0 | 9 85E-06 |
| IRF8 | 0 021577735 | 0 013832456 | 7 34E-06 | -0 46932715 | 0 | 9 92E-06 |
| ANTXR2 | 0 455692829 | 0 013831112 | 7 35E-06 | 0 112430693 | 1 33E-292 | 9 94E-06 |
| CCDC152 | 0 037377929 | 0 013830552 | 7 36E-06 | -0 45207771 | 0 | 9 95E-06 |
| TRIM66 | 0 182111515 | 0 013819279 | 7 48E-06 | -0 31174542 | 0 | 1 01E-05 |
| TMEM176B | 0 07149112 | 0 013796433 | 7 75E-06 | -0 41222336 | 0 | 1 05E-05 |
| CDC45 | 0 042194133 | 0 013788032 | 7 85E-06 | 0 405143281 | 0 | 1 06E-05 |
| IZUMO1 | 0 002046411 | 0 013778307 | 7 96E-06 | 0 496748662 | 0 | 1 08E-05 |
| RPS20 | 0 353705431 | 0 013767633 | 8 09E-06 | 0 194468871 | 0 | 1 09E-05 |
| BNIPL | 0 020178561 | 0 013745812 | 8 36E-06 | 0 479782664 | 0 | 1 13E-05 |
| UTY | 0 30656184 | 0 013745734 | 8 37E-06 | -0 22296358 | 0 | 1 13E-05 |

| | | | | | | |
|---|---|---|---|---|---|---|
| AKR1C1 | 0 076174069 | 0 013745531 | 8 37E-06 | -0 40051616 | 0 | 1 13E-05 |
| TJP3 | 0 005339704 | 0 013744782 | 8 38E-06 | 0 493757197 | 0 | 1 13E-05 |
| SLC5A2 | 0 007785879 | 0 013727411 | 8 60E-06 | 0 490965352 | 0 | 1 16E-05 |
| PDPN | 0 112124269 | 0 013714314 | 8 77E-06 | -0 38232939 | 0 | 1 18E-05 |
| TCOF1 | 0 108088557 | 0 013701462 | 8 94E-06 | 0 387073813 | 0 | 1 21E-05 |
| EGLN1 | 0 254697226 | 0 013687876 | 9 13E-06 | -0 24278512 | 0 | 1 23E-05 |
| NFE2 | 0 000780492 | 0 013682063 | 9 21E-06 | 0 49663575 | 0 | 1 24E-05 |
| TRIM69 | 0 065418515 | 0 013679977 | 9 24E-06 | 0 446488645 | 0 | 1 25E-05 |
| SLC7A1 | 0 132131503 | 0 013671208 | 9 36E-06 | 0 379877586 | 0 | 1 26E-05 |
| MAST1 | 0 029344577 | 0 013670559 | 9 37E-06 | 0 474947821 | 0 | 1 26E-05 |
| PNCK | 0 022405818 | 0 013662542 | 9 48E-06 | -0 46779271 | 0 | 1 28E-05 |
| C3AR1 | 0 017894196 | 0 013648542 | 9 68E-06 | 0 478892871 | 0 | 1 30E-05 |
| RSPO3 | 0 041984733 | 0 013647179 | 9 70E-06 | -0 44186471 | 0 | 1 31E-05 |
| FBXO5 | 0 054958025 | 0 013643506 | 9 76E-06 | 0 401101699 | 0 | 1 31E-05 |
| ESYT1 | 0 112714397 | 0 013635987 | 9 87E-06 | 0 389199266 | 0 | 1 33E-05 |
| CDC42SE1 | 0 228569797 | 0 013627118 | 1 00E-05 | -0 26498299 | 0 | 1 35E-05 |
| CNR2 | 0 003588357 | 0 013620951 | 1 01E-05 | 0 494846387 | 0 | 1 36E-05 |
| IL26 | 0 003740648 | 0 013620114 | 1 01E-05 | 0 494698105 | 0 | 1 36E-05 |
| POLD3 | 0 189792694 | 0 013580373 | 1 07E-05 | 0 277989792 | 0 | 1 44E-05 |
| MKNK2 | 0 088271687 | 0 013569566 | 1 09E-05 | -0 39272698 | 0 | 1 47E-05 |
| CNPY3 | 0 087557823 | 0 013551251 | 1 12E-05 | 0 425236206 | 0 | 1 51E-05 |
| LINC01882 | 0 000961337 | 0 0135433 | 1 13E-05 | 0 496874068 | 0 | 1 52E-05 |
| PSTPIP1 | 0 009403971 | 0 01353999 | 1 14E-05 | -0 48567905 | 0 | 1 53E-05 |
| BAMBI | 0 089918334 | 0 013537486 | 1 14E-05 | -0 39165162 | 0 | 1 54E-05 |
| MOB4 | 0 237088576 | 0 013530374 | 1 16E-05 | 0 291500886 | 0 | 1 55E-05 |
| GPA33 | 0 00241762 | 0 013519248 | 1 17E-05 | 0 495274174 | 0 | 1 58E-05 |
| UGDH | 0 16947136 | 0 013517746 | 1 18E-05 | -0 32020426 | 0 | 1 58E-05 |
| TRH | 0 000809046 | 0 013511682 | 1 19E-05 | 0 497167553 | 0 | 1 60E-05 |
| AKR1A1 | 0 202737431 | 0 013504807 | 1 20E-05 | 0 310091137 | 0 | 1 61E-05 |
| MME | 0 245331328 | 0 013503151 | 1 20E-05 | 0 233570599 | 0 | 1 62E-05 |
| TERC | 0 004483067 | 0 013492305 | 1 22E-05 | 0 494604502 | 0 | 1 64E-05 |
| MFSD1 | 0 20458396 | 0 013489916 | 1 23E-05 | -0 28506938 | 0 | 1 65E-05 |
| EIF1AX | 0 321334069 | 0 013486758 | 1 23E-05 | 0 24858381 | 0 | 1 65E-05 |
| SVOP | 0 00493994 | 0 013485112 | 1 24E-05 | 0 493632761 | 0 | 1 66E-05 |
| CPB2-AS1 | 0 061858712 | 0 013484449 | 1 24E-05 | 0 449633683 | 0 | 1 66E-05 |
| EGR2 | 0 047419619 | 0 013466989 | 1 27E-05 | -0 43752035 | 0 | 1 70E-05 |
| FAM9B | 0 005282595 | 0 013459943 | 1 28E-05 | -0 49021333 | 0 | 1 72E-05 |
| KLRC2 | 0 014201138 | 0 013455848 | 1 29E-05 | -0 47980879 | 0 | 1 73E-05 |
| UCHL1 | 0 135481906 | 0 013455377 | 1 29E-05 | 0 376448044 | 0 | 1 73E-05 |
| ANKRD49 | 0 090470389 | 0 013443215 | 1 32E-05 | -0 40000397 | 0 | 1 76E-05 |
| CDK1 | 0 071500638 | 0 013434698 | 1 33E-05 | 0 362015502 | 0 | 1 78E-05 |
| TFCP2L1 | 0 015990558 | 0 013404335 | 1 39E-05 | -0 47889424 | 0 | 1 87E-05 |
| NEU4 | 0 001846529 | 0 013391903 | 1 42E-05 | -0 49445098 | 0 | 1 90E-05 |
| FBLN2 | 0 204964688 | 0 013384841 | 1 43E-05 | -0 25269636 | 0 | 1 92E-05 |

| | | | | | | |
|---|---|---|---|---|---|---|
| NR1D2 | 0 250832842 | 0 013370603 | 1 46E-05 | -0 25056323 | 0 | 1 96E-05 |
| LINC01697 | 0 031410025 | 0 013362435 | 1 48E-05 | -0 46413385 | 0 | 1 98E-05 |
| FBXO38 | 0 252479488 | 0 013338596 | 1 53E-05 | -0 25473623 | 0 | 2 05E-05 |
| IRF1 | 0 138527726 | 0 013330171 | 1 55E-05 | -0 35242378 | 0 | 2 08E-05 |
| MLC1 | 0 005634768 | 0 013320963 | 1 58E-05 | -0 48886207 | 0 | 2 11E-05 |
| SLC46A1 | 0 021311226 | 0 013300349 | 1 62E-05 | -0 46954643 | 0 | 2 17E-05 |
| TSPAN5 | 0 435733186 | 0 013273408 | 1 69E-05 | 0 136410428 | 0 | 2 26E-05 |
| ANP32B | 0 263063715 | 0 013268558 | 1 70E-05 | 0 229274944 | 0 | 2 27E-05 |
| CFP | 0 006063087 | 0 01326724 | 1 70E-05 | 0 493289467 | 0 | 2 28E-05 |
| RASGRP3 | 0 042708115 | 0 013253932 | 1 74E-05 | 0 450802966 | 0 | 2 32E-05 |
| FYCO1 | 0 113808989 | 0 013247532 | 1 75E-05 | -0 3699251 | 0 | 2 34E-05 |
| ATXN1L | 0 098237231 | 0 013240698 | 1 77E-05 | -0 37581768 | 0 | 2 37E-05 |
| C15orf39 | 0 025508747 | 0 013237247 | 1 78E-05 | 0 475172374 | 0 | 2 38E-05 |
| SPINK5 | 0 036369001 | 0 013236791 | 1 78E-05 | 0 463522033 | 0 | 2 38E-05 |
| DDX18 | 0 382659763 | 0 013215488 | 1 84E-05 | 0 184285463 | 0 | 2 45E-05 |
| PAX5 | 0 007500333 | 0 013213257 | 1 84E-05 | 0 491826199 | 0 | 2 46E-05 |
| PTDSS1 | 0 140859683 | 0 01318717 | 1 92E-05 | 0 337486717 | 0 | 2 56E-05 |
| CRLF1 | 0 109563877 | 0 013184461 | 1 92E-05 | -0 39525071 | 0 | 2 56E-05 |
| ATG3 | 0 155641431 | 0 013181073 | 1 93E-05 | -0 3393371 | 0 | 2 58E-05 |
| SIGLEC10 | 0 00520645 | 0 013176072 | 1 95E-05 | 0 490913442 | 0 | 2 60E-05 |
| PITPNM2 | 0 192933696 | 0 013160494 | 1 99E-05 | -0 30336385 | 0 | 2 65E-05 |
| XPO6 | 0 353829168 | 0 013142364 | 2 04E-05 | 0 207871319 | 0 | 2 72E-05 |
| LINC01811 | 0 018550951 | 0 013141301 | 2 05E-05 | 0 480765326 | 0 | 2 73E-05 |
| PPP1R14C | 0 029953742 | 0 013113286 | 2 13E-05 | 0 474471361 | 0 | 2 84E-05 |
| RBM38 | 0 024204755 | 0 013101555 | 2 17E-05 | 0 481230849 | 0 | 2 89E-05 |
| GAL | 0 004092821 | 0 013099284 | 2 18E-05 | -0 4919996 | 0 | 2 90E-05 |
| MYOCD | 0 010688927 | 0 013093928 | 2 19E-05 | 0 488774404 | 0 | 2 92E-05 |
| CAPG | 0 059631456 | 0 013084572 | 2 22E-05 | 0 429748041 | 0 | 2 96E-05 |
| PDYN-AS1 | 0 003407512 | 0 013078288 | 2 24E-05 | 0 494181452 | 0 | 2 98E-05 |
| CYP4F22 | 0 009451562 | 0 013073474 | 2 26E-05 | -0 4848239 | 0 | 3 00E-05 |
| EDEM2 | 0 115132017 | 0 013049634 | 2 34E-05 | -0 3757567 | 0 | 3 11E-05 |
| SNX10 | 0 085216348 | 0 013041481 | 2 37E-05 | -0 39510631 | 0 | 3 14E-05 |
| SLC5A5 | 0 002132074 | 0 01302824 | 2 41E-05 | 0 49619114 | 0 | 3 20E-05 |
| SMYD3 | 0 734880356 | 0 013021905 | 2 43E-05 | 0 00883764 | 0 004175496 | 3 23E-05 |
| INTS4 | 0 286250024 | 0 013019953 | 2 44E-05 | 0 214278839 | 0 | 3 24E-05 |
| UBE2D1 | 0 138394472 | 0 013009766 | 2 48E-05 | -0 34704758 | 0 | 3 29E-05 |
| TMEM37 | 0 005710914 | 0 013006842 | 2 49E-05 | 0 493858777 | 0 | 3 30E-05 |
| IRF9 | 0 13319754 | 0 013005389 | 2 49E-05 | -0 35609766 | 0 | 3 31E-05 |
| HES1 | 0 138442063 | 0 012995455 | 2 53E-05 | 0 368028893 | 0 | 3 36E-05 |
| ARF4 | 0 361872037 | 0 012972781 | 2 61E-05 | -0 16797662 | 0 | 3 47E-05 |
| SPATA6L | 0 077811197 | 0 012963314 | 2 65E-05 | -0 40288052 | 0 | 3 51E-05 |
| ZNF98 | 0 023414746 | 0 012962203 | 2 65E-05 | 0 47784429 | 0 | 3 52E-05 |
| SERPINF1 | 0 296929432 | 0 012954862 | 2 68E-05 | -0 21143673 | 0 | 3 55E-05 |
| LAMP5 | 0 036806838 | 0 012942534 | 2 73E-05 | -0 44973836 | 0 | 3 62E-05 |

| | | | | | | |
|---|---|---|---|---|---|---|
| CCR6 | 0 004644876 | 0 012932507 | 2 77E-05 | -0 49137208 | 0 | 3 67E-05 |
| TK1 | 0 039586149 | 0 012917158 | 2 83E-05 | 0 417018633 | 0 | 3 75E-05 |
| GJC2 | 0 008166606 | 0 012913994 | 2 84E-05 | -0 48718471 | 0 | 3 76E-05 |
| TSPAN7 | 0 048856865 | 0 012910812 | 2 85E-05 | -0 44201434 | 0 | 3 78E-05 |
| PLEKHA7 | 0 073052103 | 0 012888592 | 2 94E-05 | -0 40578257 | 0 | 3 90E-05 |
| UHMK1 | 0 228836306 | 0 012888273 | 2 95E-05 | -0 27748749 | 0 | 3 90E-05 |
| CD5 | 0 010136872 | 0 012876701 | 2 99E-05 | -0 48594956 | 0 | 3 97E-05 |
| NOL7 | 0 192933696 | 0 012871917 | 3 01E-05 | -0 30645067 | 0 | 3 99E-05 |
| CNN1 | 0 019683615 | 0 012862335 | 3 06E-05 | -0 47428346 | 0 | 4 05E-05 |
| GRAP2 | 0 005653804 | 0 012855685 | 3 09E-05 | -0 49009962 | 0 | 4 08E-05 |
| ADAMTS19 | 0 104157545 | 0 012841259 | 3 15E-05 | -0 37395966 | 0 | 4 17E-05 |
| MAGOH | 0 13892749 | 0 012823344 | 3 23E-05 | -0 35440213 | 0 | 4 27E-05 |
| LINC00670 | 0 004235594 | 0 012822962 | 3 23E-05 | 0 494711727 | 0 | 4 28E-05 |
| CAGE1 | 0 004797167 | 0 01280593 | 3 31E-05 | -0 49055955 | 0 | 4 38E-05 |
| LRSAM1 | 0 175029982 | 0 012794912 | 3 36E-05 | -0 31914305 | 0 | 4 45E-05 |
| ZBTB7A | 0 215444214 | 0 012769988 | 3 48E-05 | -0 2788411 | 0 | 4 61E-05 |
| HAX1 | 0 116721555 | 0 012769479 | 3 49E-05 | -0 37348697 | 0 | 4 61E-05 |
| STYK1 | 0 010013135 | 0 012766606 | 3 50E-05 | 0 48539593 | 0 | 4 63E-05 |
| FAM177A1 | 0 283518303 | 0 012766428 | 3 50E-05 | -0 22306612 | 0 | 4 63E-05 |
| MAPK1 | 0 493756068 | 0 012733413 | 3 67E-05 | 0 144823925 | 0 | 4 85E-05 |
| ANKRD31 | 0 041604005 | 0 012696243 | 3 87E-05 | 0 45486915 | 0 | 5 11E-05 |
| SHMT1 | 0 078306143 | 0 012691954 | 3 89E-05 | 0 412653443 | 0 | 5 14E-05 |
| MORC1 | 0 002588947 | 0 012691116 | 3 89E-05 | 0 495957005 | 0 | 5 14E-05 |
| FRAT1 | 0 004606804 | 0 012690961 | 3 89E-05 | -0 4915101 | 0 | 5 14E-05 |
| PARD6B | 0 032114371 | 0 012690725 | 3 90E-05 | -0 45956908 | 0 | 5 14E-05 |
| VSIG1 | 0 012326055 | 0 012690716 | 3 90E-05 | 0 483613753 | 0 | 5 14E-05 |
| KLHDC8A | 0 035055491 | 0 012683898 | 3 93E-05 | -0 44538734 | 0 | 5 19E-05 |
| MZT2A | 0 172069825 | 0 012669465 | 4 01E-05 | 0 329794478 | 0 | 5 29E-05 |
| EFHC1 | 0 37347471 | 0 012647975 | 4 14E-05 | -0 14676833 | 0 | 5 45E-05 |
| GRIA3 | 0 084987912 | 0 012646834 | 4 14E-05 | -0 39275568 | 0 | 5 46E-05 |
| ESM1 | 0 03658792 | 0 012646673 | 4 14E-05 | 0 446033571 | 0 | 5 46E-05 |
| MMRN2 | 0 020549771 | 0 012642981 | 4 17E-05 | -0 47464127 | 0 | 5 49E-05 |
| PGM5-AS1 | 0 001694238 | 0 012632072 | 4 23E-05 | 0 496040512 | 0 | 5 57E-05 |
| NTRK2 | 0 179417868 | 0 012630237 | 4 24E-05 | -0 29497306 | 0 | 5 59E-05 |
| BICDL1 | 0 046705755 | 0 012619101 | 4 31E-05 | -0 43859831 | 0 | 5 67E-05 |
| NR1H3 | 0 163189355 | 0 012616683 | 4 32E-05 | -0 31911458 | 0 | 5 69E-05 |
| CAPN8 | 0 002969675 | 0 012610139 | 4 36E-05 | -0 49259899 | 0 | 5 74E-05 |
| GEM | 0 181692715 | 0 012608547 | 4 37E-05 | -0 3179215 | 0 | 5 76E-05 |
| CD46 | 0 503769203 | 0 012605277 | 4 39E-05 | -0 09451806 | 4 91E-207 | 5 78E-05 |
| APOBEC3A | 0 003483657 | 0 012582724 | 4 53E-05 | -0 49233811 | 0 | 5 96E-05 |
| LINC02539 | 0 00220822 | 0 012566742 | 4 63E-05 | 0 495957412 | 0 | 6 10E-05 |
| CASQ2 | 0 003274257 | 0 01256272 | 4 66E-05 | -0 49234995 | 0 | 6 13E-05 |
| CYSLTR1 | 0 013496792 | 0 01253206 | 4 86E-05 | -0 48222129 | 0 | 6 39E-05 |
| SWSAP1 | 0 012697264 | 0 012531123 | 4 87E-05 | -0 48099595 | 0 | 6 40E-05 |

| | | | | | | |
|---|---|---|---|---|---|---|
| LAX1 | 0 002884011 | 0 012528015 | 4 89E-05 | -0 49309484 | 0 | 6 43E-05 |
| PMEL | 0 013791856 | 0 0125219 | 4 93E-05 | 0 480207618 | 0 | 6 48E-05 |
| ITGB2-AS1 | 0 006348632 | 0 012514207 | 4 98E-05 | -0 48934878 | 0 | 6 55E-05 |
| DDX20 | 0 048390474 | 0 012508644 | 5 02E-05 | 0 453174969 | 0 | 6 60E-05 |
| SNX31 | 0 007747806 | 0 012507887 | 5 03E-05 | -0 48742425 | 0 | 6 61E-05 |
| HSPA4 | 0 287249434 | 0 012500844 | 5 08E-05 | 0 260400073 | 0 | 6 67E-05 |
| CCDC50 | 0 353486513 | 0 01248169 | 5 21E-05 | -0 18074599 | 0 | 6 84E-05 |
| NR1I3 | 0 003883421 | 0 012460561 | 5 37E-05 | -0 49204634 | 0 | 7 05E-05 |
| LINC02345 | 0 008071424 | 0 012454655 | 5 41E-05 | -0 48776828 | 0 | 7 10E-05 |
| SGCA | 0 004283185 | 0 012450628 | 5 44E-05 | 0 494495904 | 0 | 7 14E-05 |
| DCX | 0 060716529 | 0 012437362 | 5 54E-05 | -0 43477487 | 0 | 7 27E-05 |
| HUWE1 | 0 571519674 | 0 012422327 | 5 66E-05 | 0 108438206 | 3 20E-272 | 7 42E-05 |
| GPR84 | 0 001560983 | 0 012416662 | 5 70E-05 | 0 49610675 | 0 | 7 48E-05 |
| NRARP | 0 004911386 | 0 012415656 | 5 71E-05 | 0 49322756 | 0 | 7 49E-05 |
| GMEB1 | 0 129342674 | 0 012403136 | 5 81E-05 | 0 377696033 | 0 | 7 62E-05 |
| CYB5D1 | 0 023062573 | 0 012394254 | 5 88E-05 | -0 46990133 | 0 | 7 71E-05 |
| TAF7 | 0 264910243 | 0 012378893 | 6 01E-05 | -0 24079841 | 0 | 7 87E-05 |
| TMEM86A | 0 005948868 | 0 012372913 | 6 06E-05 | -0 48959887 | 0 | 7 93E-05 |
| C20orf197 | 0 000561573 | 0 012365116 | 6 12E-05 | 0 497019822 | 0 | 8 02E-05 |
| FBXO2 | 0 008842398 | 0 012358102 | 6 18E-05 | -0 48460159 | 0 | 8 10E-05 |
| AFF4 | 0 646294569 | 0 012326613 | 6 45E-05 | -0 02219232 | 6 29E-13 | 8 45E-05 |
| LINC00299 | 0 00662466 | 0 012315595 | 6 55E-05 | 0 489978409 | 0 | 8 58E-05 |
| MAP3K14-AS1 | 0 011183872 | 0 012313032 | 6 58E-05 | 0 484993263 | 0 | 8 61E-05 |
| RPS6KA4 | 0 059907483 | 0 01230557 | 6 64E-05 | 0 433175239 | 0 | 8 69E-05 |
| MACO1 | 0 286307133 | 0 012295373 | 6 74E-05 | 0 237489351 | 0 | 8 81E-05 |
| KDM3A | 0 313690963 | 0 012293885 | 6 75E-05 | -0 20136725 | 0 | 8 83E-05 |
| RGS2 | 0 08789096 | 0 012269625 | 6 98E-05 | -0 38331218 | 0 | 9 12E-05 |
| ZNF644 | 0 576783233 | 0 012259601 | 7 07E-05 | -0 06629818 | 1 18E-102 | 9 25E-05 |
| DENND2D | 0 011545564 | 0 012225303 | 7 41E-05 | -0 48377993 | 0 | 9 68E-05 |
| HELLPAR | 0 099541223 | 0 012221225 | 7 45E-05 | 0 380319838 | 0 | 9 73E-05 |
| FMO2 | 0 098265786 | 0 012203384 | 7 63E-05 | -0 3814387 | 0 | 9 97E-05 |
| C16orf87 | 0 121804268 | 0 012199283 | 7 68E-05 | 0 388622278 | 0 | 0 000100237 |
| GYG1 | 0 089851707 | 0 012171415 | 7 97E-05 | -0 40070247 | 0 | 0 000104063 |
| MXD3 | 0 057746854 | 0 012150732 | 8 20E-05 | 0 405184803 | 0 | 0 00010697 |
| SHKBP1 | 0 064038377 | 0 012137087 | 8 35E-05 | 0 434528785 | 0 | 0 000108944 |
| DIO2-AS1 | 0 05198835 | 0 012122792 | 8 51E-05 | 0 43152949 | 0 | 0 000111049 |
| ZNF611 | 0 221840437 | 0 012117518 | 8 57E-05 | 0 284917508 | 0 | 0 000111826 |
| AC144831.1 | 0 003635948 | 0 012097487 | 8 81E-05 | -0 49304055 | 0 | 0 000114867 |
| CCNB2 | 0 047295882 | 0 012078567 | 9 03E-05 | 0 408794413 | 0 | 0 000117798 |
| SIK1 | 0 044402353 | 0 012075479 | 9 07E-05 | -0 43801 | 0 | 0 000118251 |
| HEATR3 | 0 114989244 | 0 012072667 | 9 11E-05 | -0 37819149 | 0 | 0 000118682 |
| LINC01303 | 0 002217738 | 0 012072564 | 9 11E-05 | 0 495854057 | 0 | 0 000118682 |
| PODNL1 | 0 071852811 | 0 012070153 | 9 14E-05 | -0 40195318 | 0 | 0 000119051 |
| CD163 | 0 042203651 | 0 012066637 | 9 18E-05 | -0 45448203 | 0 | 0 000119598 |

| | | | | | | |
|---|---|---|---|---|---|---|
| CHST13 | 0 001265919 | 0 012065178 | 9 20E-05 | -0 49513004 | 0 | 0 000119816 |
| SNN | 0 023528964 | 0 012053663 | 9 34E-05 | 0 481560739 | 0 | 0 000121664 |
| EML2 | 0 131293903 | 0 012052602 | 9 36E-05 | -0 36401104 | 0 | 0 000121821 |
| AURKB | 0 045001999 | 0 012030911 | 9 63E-05 | 0 403195533 | 0 | 0 000125379 |
| EIF4E | 0 280900801 | 0 012020436 | 9 77E-05 | 0 256673272 | 0 | 0 000127131 |
| BCL10 | 0 069130609 | 0 012004443 | 9 98E-05 | -0 42005817 | 0 | 0 000129825 |
| TEPP | 0 004092821 | 0 011997832 | 0 000100665 | 0 49550952 | 0 | 0 000130941 |
| ADCY7 | 0 073109212 | 0 01197569 | 0 000103684 | 0 406172368 | 0 | 0 000134849 |
| C16orf89 | 0 019055415 | 0 011964888 | 0 000105187 | -0 47330263 | 0 | 0 000136785 |
| FAM118A | 0 18176886 | 0 011951053 | 0 000107143 | 0 303111391 | 0 | 0 000139271 |
| PTGDR | 0 004283185 | 0 011933834 | 0 000109625 | -0 49210702 | 0 | 0 000142458 |
| PFKFB2 | 0 05955531 | 0 011920463 | 0 00011159 | -0 42534251 | 0 | 0 000144971 |
| RDH11 | 0 211122956 | 0 011903537 | 0 000114124 | 0 322321871 | 0 | 0 000148203 |
| HK3 | 0 002360511 | 0 011896299 | 0 000115225 | 0 495151814 | 0 | 0 000149612 |
| CEP85 | 0 089965925 | 0 011890262 | 0 00011615 | 0 357842384 | 0 | 0 000150772 |
| ARL11 | 0 002055929 | 0 011886133 | 0 000116787 | -0 49427092 | 0 | 0 000151578 |
| CLDN10 | 0 034065599 | 0 011884202 | 0 000117086 | 0 482934756 | 0 | 0 000151945 |
| ZNF830 | 0 061915821 | 0 011876759 | 0 000118245 | -0 42942082 | 0 | 0 000153407 |
| NPNT | 0 064609469 | 0 01186355 | 0 000120329 | -0 41914721 | 0 | 0 00015609 |
| CDCA2 | 0 057156727 | 0 011860764 | 0 000120773 | 0 385237325 | 0 | 0 000156601 |
| ABRACL | 0 04767661 | 0 011852479 | 0 000122103 | 0 455943112 | 0 | 0 000158303 |
| DCDC1 | 0 087072395 | 0 011852209 | 0 000122147 | -0 39505531 | 0 | 0 000158338 |
| PRKX | 0 248148712 | 0 011834347 | 0 000125061 | -0 2429234 | 0 | 0 000162094 |
| VIPR1-AS1 | 0 002436656 | 0 011833299 | 0 000125234 | 0 495833442 | 0 | 0 000162296 |
| LINC00309 | 0 001703756 | 0 011832153 | 0 000125423 | -0 49412953 | 0 | 0 000162519 |
| MAP4K4 | 0 731529954 | 0 011829625 | 0 000125842 | 0 002819562 | 0 360767753 | 0 000163039 |
| SLC6A20 | 0 029182768 | 0 011813569 | 0 000128533 | -0 45686972 | 0 | 0 00016648 |
| TESC | 0 006377187 | 0 011811956 | 0 000128807 | -0 48848622 | 0 | 0 000166812 |
| ATP5MC2 | 0 362471683 | 0 011798914 | 0 000131037 | 0 193119176 | 0 | 0 000169677 |
| LINC01416 | 0 049475548 | 0 011794331 | 0 000131829 | -0 44405566 | 0 | 0 000170679 |
| EVA1B | 0 110744132 | 0 011793773 | 0 000131926 | -0 37758167 | 0 | 0 000170781 |
| OPCML | 0 069977727 | 0 011793413 | 0 000131989 | 0 435812185 | 0 | 0 000170839 |
| BFSP2 | 0 001922674 | 0 011783379 | 0 000133741 | 0 495837102 | 0 | 0 00017306 |
| CELA1 | 0 001237365 | 0 011781875 | 0 000134006 | -0 49538706 | 0 | 0 000173378 |
| IFIT3 | 0 092573909 | 0 011781335 | 0 000134101 | -0 3969444 | 0 | 0 000173478 |
| AKR1D1 | 0 006110678 | 0 011772704 | 0 00013563 | 0 492428744 | 0 | 0 000175431 |
| PLS1 | 0 016371286 | 0 01177193 | 0 000135768 | 0 480899067 | 0 | 0 000175586 |
| HAR1A | 0 003045821 | 0 011754199 | 0 000138963 | 0 494927624 | 0 | 0 000179669 |
| PRPF40A | 0 471502541 | 0 011742819 | 0 000141051 | 0 121721132 | 0 | 0 000182344 |
| KCNH5 | 0 008899507 | 0 011734386 | 0 000142618 | 0 489265655 | 0 | 0 000184344 |
| SERGEF | 0 21262683 | 0 011732488 | 0 000142973 | -0 29105319 | 0 | 0 000184777 |
| HNRNPA1 | 0 416563553 | 0 011718514 | 0 00014561 | 0 158657002 | 0 | 0 000188161 |
| ARHGEF26 | 0 263606252 | 0 011712331 | 0 000146792 | -0 23487831 | 0 | 0 000189662 |
| KPNA4 | 0 393196398 | 0 011710119 | 0 000147217 | 0 194397288 | 0 | 0 000190185 |

| | | | | | | |
|---|---|---|---|---|---|---|
| BCAS2 | 0 117054692 | 0 011703648 | 0 000148467 | -0 36885402 | 0 | 0 000191773 |
| EMBP1 | 0 006282005 | 0 01169927 | 0 000149318 | 0 493281804 | 0 | 0 00019282 |
| SELENOP | 0 263920352 | 0 011692661 | 0 000150612 | -0 23837855 | 0 | 0 000194464 |
| EYA3 | 0 377596086 | 0 011690714 | 0 000150995 | 0 170841281 | 0 | 0 000194932 |
| AKR1C3 | 0 054225124 | 0 011687865 | 0 000151557 | -0 43252887 | 0 | 0 000195605 |
| XCL1 | 0 007319488 | 0 011687222 | 0 000151685 | 0 491438397 | 0 | 0 000195742 |
| SH3RF1 | 0 345253279 | 0 011680466 | 0 000153027 | 0 210850098 | 0 | 0 000197448 |
| TMIE | 0 001570501 | 0 011661553 | 0 000156845 | 0 496101575 | 0 | 0 00020229 |
| RPP25 | 0 019274333 | 0 011659597 | 0 000157245 | 0 478963715 | 0 | 0 000202751 |
| ZNF90 | 0 074175249 | 0 011658212 | 0 000157528 | 0 385980457 | 0 | 0 000203089 |
| RUBCN | 0 183653462 | 0 011589455 | 0 000172236 | -0 30811133 | 0 | 0 000221868 |
| FPR3 | 0 014905484 | 0 011573601 | 0 000175805 | -0 48163645 | 0 | 0 000226405 |
| CISD3 | 0 051550513 | 0 01152935 | 0 000186141 | -0 44132173 | 0 | 0 000239617 |
| CACYBP | 0 176429156 | 0 011523783 | 0 000187481 | 0 339178585 | 0 | 0 000241309 |
| CNGA1 | 0 011298091 | 0 011515522 | 0 000189486 | 0 488443804 | 0 | 0 000243857 |
| GPN3 | 0 067036607 | 0 011500986 | 0 000193063 | 0 432249467 | 0 | 0 000248393 |
| MAGT1 | 0 260351031 | 0 011500867 | 0 000193093 | 0 297388629 | 0 | 0 000248397 |
| FDPS | 0 255763264 | 0 011497595 | 0 000193907 | 0 28049353 | 0 | 0 000249411 |
| FAS | 0 225533495 | 0 01148198 | 0 000197836 | -0 27591972 | 0 | 0 000254361 |
| GCLM | 0 094943938 | 0 011476154 | 0 000199321 | -0 38955661 | 0 | 0 000256236 |
| GRB2 | 0 372532409 | 0 011465419 | 0 000202086 | 0 177392242 | 0 | 0 000259754 |
| TKTL1 | 0 002008338 | 0 011449487 | 0 000206254 | 0 496206205 | 0 | 0 000265004 |
| TMEM132B | 0 035988274 | 0 011437356 | 0 000209482 | -0 45834995 | 0 | 0 000269115 |
| CNTNAP4 | 0 015486094 | 0 011435507 | 0 000209979 | -0 47958745 | 0 | 0 000269716 |
| BEX5 | 0 008071424 | 0 011421507 | 0 000213771 | -0 48679487 | 0 | 0 000274513 |
| IQCJ | 0 009898917 | 0 011418577 | 0 000214573 | -0 48358854 | 0 | 0 000275505 |
| LY86 | 0 013925111 | 0 01141773 | 0 000214805 | -0 48182875 | 0 | 0 000275766 |
| SUCNR1 | 0 005434886 | 0 011412647 | 0 000216205 | -0 49119267 | 0 | 0 000277525 |
| C1orf56 | 0 121299804 | 0 011396783 | 0 000220627 | -0 37321785 | 0 | 0 000283047 |
| FAM124B | 0 003236184 | 0 011379231 | 0 000225619 | 0 494873216 | 0 | 0 000289412 |
| MUSTN1 | 0 012402201 | 0 011376104 | 0 000226519 | -0 48280615 | 0 | 0 000290488 |
| C9orf78 | 0 090175325 | 0 011344257 | 0 000235883 | -0 40088225 | 0 | 0 000302414 |
| SLC9C1 | 0 120195694 | 0 011338265 | 0 000237685 | 0 365236653 | 0 | 0 000304683 |
| GRAMD4 | 0 111429442 | 0 011337428 | 0 000237938 | -0 37422814 | 0 | 0 000304965 |
| ABCE1 | 0 266633036 | 0 011327207 | 0 000241044 | 0 259974977 | 0 | 0 000308738 |
| TDP1 | 0 137252289 | 0 011325502 | 0 000241566 | 0 328064153 | 0 | 0 000309364 |
| GCC1 | 0 020844834 | 0 011322409 | 0 000242515 | -0 47263508 | 0 | 0 000310538 |
| TNIP3 | 0 007100569 | 0 01132178 | 0 000242709 | 0 491606931 | 0 | 0 000310744 |
| MYD88 | 0 02885915 | 0 011321448 | 0 000242811 | -0 46484915 | 0 | 0 000310832 |
| SIDT1-AS1 | 0 000713864 | 0 01131943 | 0 000243433 | 0 496891914 | 0 | 0 000311587 |
| RANBP2 | 0 490396147 | 0 011316724 | 0 00024427 | 0 150033689 | 0 | 0 000312573 |
| TAFA4 | 0 018408178 | 0 011315083 | 0 000244778 | 0 485749922 | 0 | 0 000313181 |
| NLN | 0 186727837 | 0 011311306 | 0 000245953 | 0 321741937 | 0 | 0 000314641 |
| CNDP1 | 0 00604405 | 0 011299816 | 0 000249558 | -0 48976226 | 0 | 0 000319167 |

| | | | | | | |
|---|---|---|---|---|---|---|
| EFNB1 | 0 119624603 | 0 011294818 | 0 000251142 | -0 346162 | 0 | 0 000321149 |
| SYTL5 | 0 033770535 | 0 011287029 | 0 000253628 | 0 461512497 | 0 | 0 000324285 |
| ATP6V0A4 | 0 004073785 | 0 011282921 | 0 000254949 | -0 49168588 | 0 | 0 00032593 |
| RAB44 | 0 003788239 | 0 011280381 | 0 000255769 | 0 494724682 | 0 | 0 000326933 |
| PAMR1 | 0 278445109 | 0 011266614 | 0 000260256 | -0 2176596 | 0 | 0 000332579 |
| RNF125 | 0 048523729 | 0 011262887 | 0 000261484 | -0 43039634 | 0 | 0 000334102 |
| LINC00892 | 0 001294474 | 0 011256683 | 0 000263539 | -0 49501781 | 0 | 0 000336683 |
| TRHDE | 0 072233538 | 0 011237889 | 0 000269857 | 0 395689406 | 0 | 0 000344662 |
| PAK2 | 0 436304277 | 0 011231546 | 0 000272022 | 0 185393645 | 0 | 0 000347379 |
| HK2 | 0 08478803 | 0 011229786 | 0 000272625 | -0 39471959 | 0 | 0 000348102 |
| KIAA0319L | 0 518122632 | 0 011212423 | 0 000278646 | -0 07200575 | 8 74E-121 | 0 000355694 |
| BBC3 | 0 047219737 | 0 011206027 | 0 000280895 | -0 44017918 | 0 | 0 000358516 |
| PIK3R5 | 0 016732977 | 0 01116091 | 0 000297249 | -0 47730258 | 0 | 0 000379237 |
| HDAC1 | 0 132283794 | 0 01115152 | 0 000300763 | -0 35646461 | 0 | 0 000383617 |
| ATP13A2 | 0 1507586 | 0 011137717 | 0 000306 | -0 3303385 | 0 | 0 000390243 |
| NXF1 | 0 120138585 | 0 011120904 | 0 000312493 | -0 36894855 | 0 | 0 00039847 |
| HRH1 | 0 145694923 | 0 011106093 | 0 00031832 | 0 333302413 | 0 | 0 00040579 |
| KIF2C | 0 063115113 | 0 011093721 | 0 000323265 | 0 385353497 | 0 | 0 000412039 |
| GPIHBP1 | 0 002170147 | 0 011092249 | 0 000323858 | -0 49417705 | 0 | 0 000412739 |
| LINC01619 | 0 198292437 | 0 011054127 | 0 000339578 | 0 293381253 | 0 | 0 000432599 |
| GOT1 | 0 08851916 | 0 011003179 | 0 000361704 | 0 42039746 | 0 | 0 000460723 |
| FRZB | 0 064914051 | 0 011002936 | 0 000361813 | -0 42259686 | 0 | 0 000460799 |
| LINC01748 | 0 077116369 | 0 010973495 | 0 00037521 | -0 39402208 | 0 | 0 000477797 |
| CD300A | 0 00746226 | 0 010965347 | 0 000378998 | 0 490692575 | 0 | 0 000482557 |
| PRSS57 | 0 003179075 | 0 010960894 | 0 000381084 | -0 4923748 | 0 | 0 000485081 |
| FLT3 | 0 016000076 | 0 010957824 | 0 000382528 | -0 47912095 | 0 | 0 000486854 |
| LCP1 | 0 026555748 | 0 010949507 | 0 000386466 | 0 469661846 | 0 | 0 000491734 |
| DNAJB4 | 0 128457482 | 0 010945067 | 0 000388584 | -0 35891872 | 0 | 0 000494362 |
| SMURF1 | 0 505434886 | 0 010922439 | 0 000399547 | 0 133592633 | 0 | 0 000508172 |
| UNC80 | 0 09787554 | 0 010922222 | 0 000399653 | -0 36282293 | 0 | 0 000508239 |
| CLDN23 | 0 002170147 | 0 010909682 | 0 000405854 | 0 495727785 | 0 | 0 000516055 |
| DLGAP5 | 0 049313739 | 0 010907358 | 0 000407013 | 0 404911907 | 0 | 0 000517389 |
| LINC00964 | 0 011859664 | 0 010907099 | 0 000407142 | -0 48260626 | 0 | 0 000517484 |
| GRIA2 | 0 014058366 | 0 010898901 | 0 000411257 | 0 486503979 | 0 | 0 000522643 |
| ARRB2 | 0 042003769 | 0 010895672 | 0 000412887 | 0 459004847 | 0 | 0 000524575 |
| CYP27A1 | 0 273133959 | 0 010866808 | 0 00042774 | -0 21592343 | 0 | 0 000543371 |
| SMPDL3B | 0 005025604 | 0 010859957 | 0 000431337 | 0 493244603 | 0 | 0 000547868 |
| CLDN16 | 0 029106623 | 0 010839212 | 0 000442404 | -0 4597524 | 0 | 0 000561697 |
| INHBA-AS1 | 0 008071424 | 0 010836558 | 0 000443839 | -0 48724014 | 0 | 0 000563444 |
| AMY2B | 0 094192001 | 0 010817383 | 0 000454334 | -0 39072528 | 0 | 0 000576613 |
| TRAC | 0 017970341 | 0 01078835 | 0 000470668 | -0 47798053 | 0 | 0 000597183 |
| TRAF3 | 0 273438541 | 0 010779669 | 0 000475657 | 0 228045684 | 0 | 0 000603431 |
| KIF18B | 0 076174069 | 0 010773845 | 0 000479032 | 0 340843517 | 0 | 0 00060755 |
| MAPRE1 | 0 135101178 | 0 010761061 | 0 000486519 | 0 385723776 | 0 | 0 000616797 |

| | | | | | | |
|---|---|---|---|---|---|---|
| FOS | 0 348546573 | 0 010757899 | 0 000488387 | -0 13718375 | 0 | 0 000619082 |
| CMTM8 | 0 169775942 | 0 010751089 | 0 000492434 | -0 31674472 | 0 | 0 000624045 |
| HSPA1L | 0 005016086 | 0 010748804 | 0 000493799 | -0 48989433 | 0 | 0 00062569 |
| HILPDA | 0 099427005 | 0 010748292 | 0 000494105 | -0 3885387 | 0 | 0 000625995 |
| CASC11 | 0 00693876 | 0 010732091 | 0 000503889 | 0 490893415 | 0 | 0 000638219 |
| KCNMB1 | 0 012430755 | 0 010716472 | 0 000513493 | -0 48267177 | 0 | 0 000650209 |
| HAUS3 | 0 080114599 | 0 010711324 | 0 000516696 | 0 41732712 | 0 | 0 000654089 |
| DNASE1L1 | 0 046458282 | 0 010698802 | 0 000524563 | -0 4422813 | 0 | 0 000663871 |
| SIPA1L3 | 0 43892178 | 0 010694817 | 0 00052709 | 0 117018154 | 0 | 0 000666979 |
| SLIRP | 0 297357751 | 0 01068609 | 0 000532664 | 0 236833675 | 0 | 0 000673942 |
| RAB3A | 0 007005387 | 0 010664886 | 0 000546437 | -0 48761269 | 0 | 0 000691183 |
| GNG7 | 0 158991833 | 0 010662624 | 0 000547925 | -0 29220327 | 0 | 0 000692973 |
| GPR171 | 0 013163656 | 0 010643892 | 0 000560399 | -0 4812253 | 0 | 0 00070856 |
| LINC02456 | 0 074889113 | 0 010634535 | 0 000566729 | -0 41106281 | 0 | 0 000716467 |
| MED12L | 0 266071463 | 0 010627927 | 0 000571239 | 0 255605069 | 0 | 0 000721975 |
| SELENOW | 0 255154099 | 0 010598643 | 0 000591631 | -0 24062764 | 0 | 0 000747549 |
| RXFP1 | 0 018265405 | 0 010567388 | 0 000614142 | 0 475644915 | 0 | 0 000775784 |
| ATF7IP | 0 519854943 | 0 010565703 | 0 000615378 | 0 099626371 | 6 73E-230 | 0 000777242 |
| ABI1 | 0 420285165 | 0 010561489 | 0 000618479 | -0 11506027 | 1 94E-306 | 0 000781053 |
| REXO2 | 0 191296568 | 0 01056089 | 0 000618921 | -0 29548989 | 0 | 0 000781507 |
| ZNF252P-AS1 | 0 005606213 | 0 010524765 | 0 000646126 | -0 48978515 | 0 | 0 00081575 |
| SH3D19 | 0 61816832 | 0 010511561 | 0 000656344 | -0 03305194 | 8 56E-27 | 0 00082843 |
| C11orf21 | 0 002017856 | 0 010481327 | 0 000680311 | -0 49419663 | 0 | 0 000858451 |
| SAMD12-AS1 | 0 008404561 | 0 010478371 | 0 000682697 | -0 48673801 | 0 | 0 000861232 |
| SLC1A1 | 0 099912433 | 0 010466266 | 0 00069255 | -0 37431963 | 0 | 0 000873545 |
| GNS | 0 250956578 | 0 010439049 | 0 00071519 | -0 26111951 | 0 | 0 000901981 |
| PNO1 | 0 097380594 | 0 010438095 | 0 000715996 | 0 401250673 | 0 | 0 000902877 |
| ESRRB | 0 03027736 | 0 010437454 | 0 000716538 | 0 47322337 | 0 | 0 00090344 |
| ESPN | 0 008014315 | 0 010435853 | 0 000717893 | -0 48549808 | 0 | 0 000905028 |
| TMCO6 | 0 029468314 | 0 010423147 | 0 000728734 | 0 467344521 | 0 | 0 000918572 |
| SSH2 | 0 607136738 | 0 010416492 | 0 000734472 | 0 078000506 | 1 89E-141 | 0 000925558 |
| IMPA2 | 0 085549485 | 0 010411093 | 0 000739158 | 0 394886961 | 0 | 0 000931339 |
| CXCL1 | 0 012916183 | 0 010404172 | 0 000745206 | -0 48242043 | 0 | 0 000938835 |
| TANGO2 | 0 10097847 | 0 010401109 | 0 000747897 | 0 398734066 | 0 | 0 000942099 |
| ARRDC3-AS1 | 0 110468105 | 0 01038227 | 0 00076465 | 0 389675371 | 0 | 0 000962817 |
| A2M-AS1 | 0 019474215 | 0 010381947 | 0 00076494 | -0 47232367 | 0 | 0 000963054 |
| IQCK | 0 224848185 | 0 010376042 | 0 000770264 | -0 27894638 | 0 | 0 000969628 |
| PNOC | 0 002255811 | 0 010375338 | 0 000770901 | 0 495335514 | 0 | 0 0009703 |
| NCK1-DT | 0 118234947 | 0 01037034 | 0 000775438 | 0 386167729 | 0 | 0 00097562 |
| TMEM252-DT | 0 004035712 | 0 010357364 | 0 000787332 | -0 49147917 | 0 | 0 000990321 |
| USP38 | 0 121042813 | 0 010356674 | 0 000787969 | 0 386197214 | 0 | 0 000990992 |
| MAP1S | 0 047219737 | 0 01035232 | 0 000792001 | -0 44503133 | 0 | 0 000995796 |
| FAM151B | 0 089375797 | 0 010349476 | 0 000794645 | -0 39781097 | 0 | 0 000998856 |
| IGSF6 | 0 04305077 | 0 010341523 | 0 000802081 | 0 451493954 | 0 | 0 001008069 |

| | | | | | | |
|---|---|---|---|---|---|---|
| PLN | 0 031000742 | 0 010337712 | 0 000805668 | -0 45970842 | 0 | 0 001012442 |
| FAM83D | 0 031362434 | 0 010333898 | 0 000809272 | 0 429896875 | 0 | 0 001016836 |
| SLC12A3 | 0 00189412 | 0 010321165 | 0 000821414 | 0 495402459 | 0 | 0 001031818 |
| VIPR1 | 0 00483524 | 0 010320138 | 0 0008224 | 0 493357919 | 0 | 0 001032919 |
| LRRFIP1 | 0 531619425 | 0 010314216 | 0 000828111 | 0 117564964 | 0 | 0 001039816 |
| BMX | 0 015276694 | 0 010299171 | 0 000842785 | -0 48146296 | 0 | 0 0010581 |
| P2RX1 | 0 003578839 | 0 010284952 | 0 000856874 | -0 49234411 | 0 | 0 001075646 |
| STON2 | 0 160790771 | 0 010250405 | 0 000892022 | 0 343605408 | 0 | 0 001119321 |
| DIPK2B | 0 008899507 | 0 010248654 | 0 000893838 | -0 48681714 | 0 | 0 001121451 |
| DDAH2 | 0 110134968 | 0 010233929 | 0 00090925 | -0 38014619 | 0 | 0 001140334 |
| RAP1GAP2 | 0 139365327 | 0 010225707 | 0 000917963 | 0 37526386 | 0 | 0 001150955 |
| HPSE | 0 008404561 | 0 010219751 | 0 000924322 | 0 487669625 | 0 | 0 001158775 |
| LINC00603 | 0 002646057 | 0 010183304 | 0 000964137 | -0 49375595 | 0 | 0 001208048 |
| CA5B | 0 262644914 | 0 010183025 | 0 000964448 | 0 269888334 | 0 | 0 001208277 |
| ETV5 | 0 229721498 | 0 010171024 | 0 000977905 | 0 308140389 | 0 | 0 001224974 |
| PLA2G2C | 0 001960747 | 0 010169723 | 0 000979375 | 0 496250651 | 0 | 0 001226652 |
| ANXA2R | 0 025946584 | 0 010165684 | 0 000983949 | 0 468826823 | 0 | 0 001232218 |
| UBXN11 | 0 068921208 | 0 010160457 | 0 000989898 | 0 423940759 | 0 | 0 001239504 |
| MS4A4E | 0 014686566 | 0 010154097 | 0 000997182 | -0 47985677 | 0 | 0 00124846 |
| ERCC1 | 0 232767318 | 0 010151165 | 0 001000557 | -0 2556327 | 0 | 0 001252353 |
| IRAK4 | 0 155308294 | 0 010143233 | 0 001009739 | 0 341659502 | 0 | 0 001263511 |
| GZMK | 0 006795987 | 0 010140869 | 0 001012492 | 0 491187052 | 0 | 0 001266788 |
| DMTN | 0 011488454 | 0 010081469 | 0 001083955 | -0 48209143 | 0 | 0 00135602 |
| FAM162A | 0 141573547 | 0 010076379 | 0 00109029 | -0 33652036 | 0 | 0 001363764 |
| SOD2 | 0 365726904 | 0 010057062 | 0 001114648 | -0 18721744 | 0 | 0 001393526 |
| OSCAR | 0 013487274 | 0 010057042 | 0 001114673 | 0 481980184 | 0 | 0 001393526 |
| NUCB2 | 0 257752565 | 0 010050205 | 0 001123415 | 0 261601192 | 0 | 0 001404269 |
| NAPSA | 0 005196931 | 0 010044466 | 0 001130801 | -0 48983827 | 0 | 0 001413314 |
| GLOD4 | 0 113085607 | 0 010041962 | 0 001134037 | 0 408033784 | 0 | 0 001416984 |
| CLEC6A | 0 001227846 | 0 010040891 | 0 001135424 | -0 49509745 | 0 | 0 00141853 |
| NLRP6 | 0 001075555 | 0 010039532 | 0 001137187 | -0 49560632 | 0 | 0 001420544 |
| HERC5 | 0 080733281 | 0 010038612 | 0 001138381 | 0 402347141 | 0 | 0 001421848 |
| LRRIQ3 | 0 011031581 | 0 010017919 | 0 001165556 | -0 48341402 | 0 | 0 001455597 |
| SIL1 | 0 495716815 | 0 009998866 | 0 001191104 | -0 07101791 | 1 54E-117 | 0 001487306 |
| SELP | 0 005396813 | 0 009978864 | 0 001218482 | -0 49009097 | 0 | 0 001520889 |
| SARAF | 0 301736118 | 0 009978014 | 0 001219659 | -0 19395193 | 0 | 0 001522158 |
| OMD | 0 234813729 | 0 009974589 | 0 001224408 | 0 272137306 | 0 | 0 001527882 |
| PDE4D | 0 785231578 | 0 009965481 | 0 001237123 | -0 00379969 | 0 218101076 | 0 001543546 |
| HTRA1 | 0 2210885 | 0 009960481 | 0 001244156 | 0 29211592 | 0 | 0 001552115 |
| LINC01695 | 0 025727666 | 0 009955484 | 0 00125122 | -0 46921269 | 0 | 0 001560721 |
| IDO1 | 0 018122632 | 0 009941951 | 0 001270538 | 0 478895316 | 0 | 0 00158461 |
| FCRL6 | 0 004226076 | 0 009922015 | 0 0012995 | 0 494546391 | 0 | 0 001620303 |
| TMEM245 | 0 4535322 | 0 00990695 | 0 001321787 | 0 12533872 | 0 | 0 001647658 |
| LINC01033 | 0 035093564 | 0 009905708 | 0 00132364 | 0 467452339 | 0 | 0 001649749 |

| | | | | | | |
|---|---|---|---|---|---|---|
| EDNRB-AS1 | 0 001846529 | 0 009905031 | 0 001324652 | -0 49470672 | 0 | 0 001650792 |
| ENY2 | 0 249595477 | 0 009901417 | 0 001330061 | 0 290577614 | 0 | 0 001657315 |
| DOCK1 | 0 753307571 | 0 009893685 | 0 001341703 | -0 00823822 | 0 007578546 | 0 001671381 |
| WBP2 | 0 160333898 | 0 009885793 | 0 001353684 | -0 32469639 | 0 | 0 001686083 |
| ENC1 | 0 111638842 | 0 0098846 | 0 001355503 | 0 362674979 | 0 | 0 001688126 |
| LRG1 | 0 001570501 | 0 009879682 | 0 001363028 | -0 49468037 | 0 | 0 001697275 |
| SPNS2 | 0 00835697 | 0 009873314 | 0 001372828 | -0 48629541 | 0 | 0 001709027 |
| S1PR1 | 0 011202909 | 0 009863059 | 0 001388748 | -0 4841002 | 0 | 0 001728163 |
| RGSL1 | 0 002798348 | 0 009855172 | 0 001401106 | 0 495273684 | 0 | 0 001743082 |
| CTSG | 0 0005235 | 0 009849148 | 0 001410615 | -0 49622294 | 0 | 0 001754681 |
| CDK17 | 0 501047001 | 0 009844586 | 0 001417855 | 0 090437231 | 1 20E-189 | 0 001763455 |
| RHOH | 0 008851916 | 0 009833143 | 0 001436166 | 0 489707646 | 0 | 0 001785759 |
| TRDN | 0 022120272 | 0 009824037 | 0 001450893 | 0 478772502 | 0 | 0 001803358 |
| CKAP4 | 0 218261598 | 0 009821051 | 0 001455751 | 0 297586862 | 0 | 0 00180916 |
| CCDC183 | 0 014819821 | 0 00981807 | 0 001460618 | -0 48016388 | 0 | 0 001814731 |
| TMCO4 | 0 151148845 | 0 009807458 | 0 001478064 | 0 345305976 | 0 | 0 001835923 |
| SAG | 0 002341475 | 0 009800361 | 0 001489837 | -0 49403606 | 0 | 0 001850303 |
| GTF2B | 0 191410786 | 0 009763373 | 0 001552611 | -0 2990286 | 0 | 0 001927505 |
| IL2RB | 0 004444994 | 0 009738552 | 0 00159609 | -0 49181173 | 0 | 0 001981222 |
| PDE6G | 0 003931012 | 0 009738309 | 0 001596522 | 0 493162664 | 0 | 0 001981499 |
| HLA-DRB5 | 0 02775504 | 0 009737023 | 0 001598804 | -0 47006215 | 0 | 0 001984071 |
| REM2 | 0 004149931 | 0 009736566 | 0 001599617 | 0 493434723 | 0 | 0 001984819 |
| SPAG5 | 0 036140565 | 0 009735508 | 0 001601499 | 0 421917677 | 0 | 0 001986893 |
| NOCT | 0 149949554 | 0 009706529 | 0 001653843 | 0 335504906 | 0 | 0 002051295 |
| P2RX4 | 0 10844073 | 0 009676019 | 0 001710648 | -0 38392378 | 0 | 0 002120638 |
| MIAT | 0 026641412 | 0 009671024 | 0 001720117 | 0 461721145 | 0 | 0 002132097 |
| NSD2 | 0 345957625 | 0 009670226 | 0 001721635 | 0 150651684 | 0 | 0 002133698 |
| MRC1 | 0 040728332 | 0 00966317 | 0 001735104 | 0 450951167 | 0 | 0 002149827 |
| IGHG1 | 0 007500333 | 0 009658814 | 0 001743468 | -0 48995391 | 0 | 0 002159624 |
| LINC01970 | 0 006005977 | 0 009658421 | 0 001744225 | -0 48911351 | 0 | 0 002160278 |
| CD28 | 0 00341703 | 0 009654153 | 0 001752457 | 0 493991721 | 0 | 0 002170018 |
| NDFIP1 | 0 438464906 | 0 009654107 | 0 001752548 | -0 11405529 | 4 12E-301 | 0 002170018 |
| ARHGAP20 | 0 182216215 | 0 009629432 | 0 001800857 | -0 27826113 | 0 | 0 002228667 |
| CNN3 | 0 515428985 | 0 009623285 | 0 001813081 | -0 0826409 | 1 36E-158 | 0 002243501 |
| CERT1 | 0 471426396 | 0 009610777 | 0 001838191 | -0 10091455 | 7 40E-236 | 0 002273976 |
| NUPR1 | 0 184957454 | 0 009597401 | 0 001865395 | -0 31601551 | 0 | 0 002307328 |
| POT1-AS1 | 0 081171118 | 0 009579301 | 0 001902796 | 0 429583473 | 0 | 0 002352974 |
| BCL2L15 | 0 008594925 | 0 009568352 | 0 001925756 | 0 488885592 | 0 | 0 002380743 |
| DYNLT1 | 0 197216881 | 0 009560687 | 0 001941978 | 0 325214962 | 0 | 0 002400484 |
| KIF16B | 0 377139213 | 0 009550525 | 0 001963681 | -0 14708384 | 0 | 0 002426675 |
| MDGA2 | 0 126430108 | 0 009547865 | 0 001969398 | -0 34560929 | 0 | 0 002433422 |
| RASIP1 | 0 016571168 | 0 00954362 | 0 001978554 | 0 482329347 | 0 | 0 002444416 |
| BISPR | 0 012173764 | 0 009543039 | 0 00197981 | -0 48374545 | 0 | 0 002445648 |
| TBC1D10C | 0 00593935 | 0 009541149 | 0 0019839 | -0 49011105 | 0 | 0 00245038 |

| | | | | | | |
|---|---|---|---|---|---|---|
| EPPK1 | 0 001608574 | 0 009526263 | 0 002016397 | 0 496225665 | 0 | 0 002489867 |
| ZNF672 | 0 033780054 | 0 009515527 | 0 002040135 | 0 474446953 | 0 | 0 002518521 |
| HIF1A-AS3 | 0 236565076 | 0 009511815 | 0 002048403 | -0 29208723 | 0 | 0 002528397 |
| IL27 | 0 001979783 | 0 009510647 | 0 002051012 | 0 496144095 | 0 | 0 002531286 |
| SDCBP | 0 384563401 | 0 009508856 | 0 002055016 | -0 14319041 | 0 | 0 002535897 |
| NCF2 | 0 015885858 | 0 009498881 | 0 00207745 | 0 481038179 | 0 | 0 002563246 |
| OR7C1 | 0 008080943 | 0 00949399 | 0 002088532 | -0 48807405 | 0 | 0 002576247 |
| GLP2R | 0 002503284 | 0 009489867 | 0 002097917 | -0 49384631 | 0 | 0 002587486 |
| ZNF207 | 0 610030268 | 0 00948825 | 0 002101607 | 0 087846534 | 5 15E-179 | 0 002591361 |
| NUDT1 | 0 049113857 | 0 009487585 | 0 002103128 | 0 447852499 | 0 | 0 002592897 |
| LINC02540 | 0 003816794 | 0 009480433 | 0 002119535 | -0 49225491 | 0 | 0 002612103 |
| KRT7 | 0 002903048 | 0 009473297 | 0 002136023 | 0 49454572 | 0 | 0 00263208 |
| RUFY4 | 0 002017856 | 0 009472434 | 0 002138024 | -0 4946463 | 0 | 0 002634201 |
| RPGRIP1 | 0 005958386 | 0 009457818 | 0 002172192 | 0 491785562 | 0 | 0 00267595 |
| UTP6 | 0 229454988 | 0 009442499 | 0 002208538 | 0 30076415 | 0 | 0 002720016 |
| LSM7 | 0 229207516 | 0 009426881 | 0 002246169 | 0 301767156 | 0 | 0 002765281 |
| PGLYRP2 | 0 003321848 | 0 009410959 | 0 002285135 | 0 494037175 | 0 | 0 002812153 |
| CCDC69 | 0 015695494 | 0 009409408 | 0 002288962 | -0 47822993 | 0 | 0 002816496 |
| SESTD1 | 0 503816794 | 0 00938487 | 0 002350323 | 0 144351658 | 0 | 0 002891246 |
| ARHGAP9 | 0 009080353 | 0 009383394 | 0 00235406 | 0 4890399 | 0 | 0 002895467 |
| SGO2 | 0 090613162 | 0 009376786 | 0 002370864 | 0 351913025 | 0 | 0 002915376 |
| ALOX15B | 0 004188003 | 0 009376549 | 0 002371469 | -0 49188863 | 0 | 0 00291574 |
| BNIP3L | 0 388827549 | 0 009373374 | 0 002379583 | -0 12821995 | 0 | 0 002925337 |
| PXK | 0 299299461 | 0 009369034 | 0 002390719 | 0 221860167 | 0 | 0 002938644 |
| TROAP | 0 038853249 | 0 009343542 | 0 002457081 | 0 410050537 | 0 | 0 003019038 |
| TGM1 | 0 00756696 | 0 009339235 | 0 002468458 | 0 491441188 | 0 | 0 003032621 |
| TTF2 | 0 134739487 | 0 00932961 | 0 002494059 | 0 330890228 | 0 | 0 003063676 |
| CAPZB | 0 493318231 | 0 009328871 | 0 002496035 | -0 10224205 | 4 43E-242 | 0 003065704 |
| PPFIA2 | 0 329843331 | 0 009322958 | 0 002511892 | -0 2071293 | 0 | 0 00308478 |
| OTUD7B | 0 381222516 | 0 009309313 | 0 002548841 | 0 195277066 | 0 | 0 003129343 |
| BORA | 0 057289981 | 0 009307112 | 0 002554848 | 0 414671893 | 0 | 0 00313631 |
| SAP30 | 0 064809351 | 0 00929232 | 0 002595548 | 0 433285864 | 0 | 0 003185446 |
| SMIM2-AS1 | 0 006462851 | 0 009291416 | 0 002598056 | -0 48859234 | 0 | 0 00318811 |
| IGFLR1 | 0 010450972 | 0 009290994 | 0 002599226 | 0 487230605 | 0 | 0 00318913 |
| ARRDC1 | 0 059412537 | 0 009289929 | 0 002602185 | -0 42788948 | 0 | 0 003192347 |
| MICALL2 | 0 164074546 | 0 00928497 | 0 002615993 | -0 31743144 | 0 | 0 00320887 |
| CD3E | 0 004245112 | 0 009267539 | 0 002665068 | 0 493640317 | 0 | 0 003268643 |
| CD320 | 0 069520854 | 0 009238497 | 0 002748706 | 0 431051973 | 0 | 0 003369474 |
| TM2D2 | 0 068235899 | 0 009210464 | 0 002831703 | -0 4208168 | 0 | 0 003469864 |
| MATN1-AS1 | 0 020083379 | 0 009207522 | 0 002840545 | 0 474675338 | 0 | 0 003480248 |
| HEXA-AS1 | 0 004340294 | 0 00920336 | 0 002853099 | -0 49122805 | 0 | 0 003494723 |
| KLHL6-AS1 | 0 00089471 | 0 009201279 | 0 002859391 | 0 496255861 | 0 | 0 003501976 |
| CSK | 0 087672041 | 0 009200901 | 0 002860536 | 0 419291505 | 0 | 0 003502924 |
| LINC02211 | 0 002864975 | 0 009165742 | 0 002968862 | 0 495272772 | 0 | 0 003632753 |

| | | | | | | |
|---|---|---|---|---|---|---|
| SMCR8 | 0 040433268 | 0 009157418 | 0 002995052 | -0 4503211 | 0 | 0 003664325 |
| FZD2 | 0 159039424 | 0 009142663 | 0 003041989 | -0 32993113 | 0 | 0 003721268 |
| SCHIP1 | 0 075041404 | 0 009127303 | 0 003091565 | -0 4058197 | 0 | 0 003780446 |
| CTC1 | 0 065780206 | 0 009116765 | 0 003126002 | -0 42148554 | 0 | 0 003822063 |
| CCDC22 | 0 032085816 | 0 009104539 | 0 003166393 | -0 46252436 | 0 | 0 003870446 |
| VAMP8 | 0 028849632 | 0 00909526 | 0 003197368 | -0 4655379 | 0 | 0 003907802 |
| EPS15-AS1 | 0 00946108 | 0 009088766 | 0 003219207 | -0 48478929 | 0 | 0 003933985 |
| HEMGN | 0 000856637 | 0 009083934 | 0 003235547 | 0 496620864 | 0 | 0 003952931 |
| IL19 | 0 003369439 | 0 009081934 | 0 003242333 | 0 494809215 | 0 | 0 003960709 |
| FGF22 | 0 004435476 | 0 009071816 | 0 003276854 | -0 49096689 | 0 | 0 004002362 |
| TFR2 | 0 034398736 | 0 009053662 | 0 003339639 | 0 443475325 | 0 | 0 004077993 |
| AKT3 | 0 80146009 | 0 009002974 | 0 003520781 | 0 009768636 | 0 001543532 | 0 004296408 |
| TNFRSF8 | 0 025080429 | 0 008994209 | 0 003552995 | -0 46454085 | 0 | 0 004335159 |
| RILPL2 | 0 072461975 | 0 008971442 | 0 00363793 | -0 41835355 | 0 | 0 004438218 |
| PSMA8 | 0 002293884 | 0 008971136 | 0 003639087 | 0 495648425 | 0 | 0 004439056 |
| LAMP3 | 0 006005977 | 0 008939331 | 0 003760865 | 0 491955246 | 0 | 0 004585237 |
| BTD | 0 301612381 | 0 008935313 | 0 003776508 | 0 219560986 | 0 | 0 004603716 |
| ANO5 | 0 058908073 | 0 008924647 | 0 003818329 | 0 451932436 | 0 | 0 004652297 |
| FMNL2 | 0 772258286 | 0 008919642 | 0 003838095 | -0 01844687 | 2 24E-09 | 0 004675776 |
| LINC01727 | 0 002094002 | 0 008891717 | 0 003950107 | -0 49434142 | 0 | 0 004811616 |
| DIPK1A | 0 156545659 | 0 008890126 | 0 003956577 | 0 348874804 | 0 | 0 004818875 |
| ASB15 | 0 001970265 | 0 008881656 | 0 003991184 | 0 495668368 | 0 | 0 004860398 |
| CCSER2 | 0 527450458 | 0 008870892 | 0 004035563 | -0 07029328 | 3 46E-115 | 0 004912543 |
| NEU1 | 0 082246673 | 0 008862877 | 0 004068898 | -0 40064259 | 0 | 0 004952484 |
| IGSF10 | 0 067255525 | 0 008856675 | 0 004094861 | 0 403896765 | 0 | 0 004983443 |
| HSH2D | 0 00583465 | 0 008855353 | 0 004100417 | 0 492470725 | 0 | 0 004989562 |
| AP4B1-AS1 | 0 029953742 | 0 008840183 | 0 00416464 | 0 459005812 | 0 | 0 005066406 |
| LINC02156 | 0 000780492 | 0 00883948 | 0 004167638 | 0 496454846 | 0 | 0 005069402 |
| WNT2 | 0 010032171 | 0 008832993 | 0 004195401 | 0 488054864 | 0 | 0 005102514 |
| RAP1A | 0 502760275 | 0 008825206 | 0 004228944 | -0 06808628 | 3 58E-108 | 0 005142649 |
| NEK10 | 0 118320611 | 0 008808753 | 0 00430062 | -0 36159833 | 0 | 0 005227792 |
| CLIP1 | 0 506948278 | 0 008806958 | 0 004308503 | 0 118426993 | 0 | 0 005236701 |
| TRG-AS1 | 0 005453922 | 0 008792651 | 0 004371829 | 0 492026885 | 0 | 0 005312304 |
| PRC1 | 0 100968952 | 0 008790943 | 0 004379444 | 0 333514583 | 0 | 0 005320873 |
| RPP21 | 0 081837391 | 0 008778954 | 0 004433245 | -0 41084427 | 0 | 0 005384854 |
| ATF3 | 0 138270735 | 0 008754344 | 0 004545556 | -0 32899837 | 0 | 0 005520563 |
| KISS1R | 0 002750757 | 0 008752353 | 0 004554754 | -0 49338224 | 0 | 0 005531023 |
| NFAM1 | 0 007614551 | 0 008748056 | 0 004574664 | -0 48790899 | 0 | 0 005553774 |
| CLDN5 | 0 018341551 | 0 008746716 | 0 004580889 | -0 47813949 | 0 | 0 005560616 |
| PTPN6 | 0 022491481 | 0 008745212 | 0 004587886 | 0 470767832 | 0 | 0 005568395 |
| TRIM59 | 0 062220403 | 0 008733564 | 0 004642392 | 0 384315565 | 0 | 0 005633103 |
| CHAC1 | 0 002084483 | 0 008727857 | 0 004669313 | 0 495198818 | 0 | 0 005664314 |
| CYP2S1 | 0 002922084 | 0 008724249 | 0 004686403 | 0 494931919 | 0 | 0 005684316 |
| PDLIM2 | 0 091108108 | 0 008717623 | 0 004717938 | -0 39922079 | 0 | 0 005721832 |

| | | | | | | |
|---|---|---|---|---|---|---|
| TMEM98 | 0 197864118 | 0 008707141 | 0 004768218 | -0 27739779 | 0 | 0 005782069 |
| SCARF1 | 0 020083379 | 0 008704552 | 0 004780714 | -0 47238995 | 0 | 0 005796478 |
| LINC01375 | 0 001085074 | 0 008685444 | 0 004873839 | 0 496535942 | 0 | 0 005907116 |
| FDFT1 | 0 197235918 | 0 008650319 | 0 005049322 | 0 328656404 | 0 | 0 006117448 |
| LINC01692 | 0 003007748 | 0 008642916 | 0 005087029 | -0 49317811 | 0 | 0 006162342 |
| CCL19 | 0 001465801 | 0 008639518 | 0 005104421 | 0 496041054 | 0 | 0 006182617 |
| RNPC3 | 0 33563039 | 0 008630979 | 0 005148362 | -0 19525588 | 0 | 0 006234242 |
| TTC22 | 0 004797167 | 0 008606543 | 0 00527601 | 0 492327799 | 0 | 0 006387176 |
| CCDC59 | 0 138032781 | 0 008602582 | 0 005296969 | -0 35528554 | 0 | 0 006410906 |
| CATSPERG | 0 051912204 | 0 00860175 | 0 005301381 | -0 43569183 | 0 | 0 006415425 |
| IRAG2 | 0 031191106 | 0 00858423 | 0 005395055 | 0 474178319 | 0 | 0 006527112 |
| DGLUCY | 0 272953113 | 0 008576087 | 0 0054391 | -0 25457302 | 0 | 0 006579556 |
| ZNF792 | 0 00473054 | 0 008572295 | 0 005459722 | -0 49123647 | 0 | 0 006603657 |
| ELF1 | 0 47787021 | 0 008544334 | 0 005613975 | -0 07973829 | 9 40E-148 | 0 00678936 |
| ZPLD1 | 0 008813843 | 0 008511883 | 0 005797914 | 0 486573761 | 0 | 0 007007327 |
| LINC01505 | 0 167710495 | 0 008511168 | 0 005802024 | 0 3093832 | 0 | 0 0070105 |
| MRO | 0 006805505 | 0 008505498 | 0 005834735 | -0 48876947 | 0 | 0 007049124 |
| IGSF21 | 0 008080943 | 0 008482671 | 0 005968119 | -0 48703739 | 0 | 0 007208426 |
| IL20RB | 0 049028193 | 0 008472055 | 0 006031087 | 0 444846209 | 0 | 0 007282619 |
| PYGL | 0 182044888 | 0 008450912 | 0 006158271 | -0 32094674 | 0 | 0 007434296 |
| RNF135 | 0 10633721 | 0 008447897 | 0 0061766 | -0 38787032 | 0 | 0 007455469 |
| LGR4 | 0 262111896 | 0 008434213 | 0 006260421 | -0 24296579 | 0 | 0 007554715 |
| CCR9 | 0 003150521 | 0 008419224 | 0 006353408 | -0 49307555 | 0 | 0 007665948 |
| RAB33A | 0 002446175 | 0 008415875 | 0 006374354 | -0 49381488 | 0 | 0 007689258 |
| PTRHD1 | 0 035141155 | 0 008411695 | 0 006400588 | -0 45881998 | 0 | 0 007718756 |
| FAM110D | 0 003388475 | 0 008411588 | 0 006401258 | 0 494545877 | 0 | 0 007718756 |
| SRCIN1 | 0 04163256 | 0 008407376 | 0 006427787 | -0 4430474 | 0 | 0 007749756 |
| CLNK | 0 010955436 | 0 008385186 | 0 006569207 | -0 48458114 | 0 | 0 00791824 |
| ABHD15 | 0 022063163 | 0 008381391 | 0 006593674 | -0 47046164 | 0 | 0 007946718 |
| TIGD3 | 0 000818564 | 0 008380197 | 0 006601384 | 0 496505794 | 0 | 0 007953982 |
| MIR325HG | 0 005139822 | 0 008372109 | 0 006653847 | 0 493658346 | 0 | 0 008016173 |
| OTOF | 0 001722792 | 0 008370547 | 0 006664027 | 0 496180354 | 0 | 0 008027413 |
| NSA2 | 0 198463764 | 0 008368379 | 0 006678172 | 0 321241423 | 0 | 0 008043426 |
| TMEM105 | 0 001189774 | 0 008360323 | 0 006730979 | -0 49525723 | 0 | 0 008105996 |
| ANK1 | 0 086120576 | 0 008356482 | 0 006756283 | 0 412493574 | 0 | 0 008132324 |
| RASSF5 | 0 077468542 | 0 008354846 | 0 006767089 | -0 39133068 | 0 | 0 008144294 |
| UCN | 0 031524243 | 0 00833545 | 0 006896388 | -0 4640283 | 0 | 0 008297794 |
| ECSCR | 0 009423007 | 0 008327602 | 0 006949331 | 0 488414272 | 0 | 0 00836043 |
| HPGDS | 0 007233824 | 0 008327154 | 0 006952365 | 0 490401825 | 0 | 0 008363015 |
| STRC | 0 004711504 | 0 008293507 | 0 007183601 | -0 49128725 | 0 | 0 008637871 |
| LINC02084 | 0 001798938 | 0 008284471 | 0 007246865 | 0 495469083 | 0 | 0 008712834 |
| DIRAS3 | 0 024880547 | 0 00828139 | 0 007268551 | -0 46821049 | 0 | 0 008737796 |
| C5AR2 | 0 004816204 | 0 008280209 | 0 007276879 | -0 49067715 | 0 | 0 008745582 |
| BIRC7 | 0 003683539 | 0 008261344 | 0 00741107 | -0 49172483 | 0 | 0 008905724 |

| | | | | | | |
|---|---|---|---|---|---|---|
| VIM-AS1 | 0 032409434 | 0 008252546 | 0 007474402 | -0 4603484 | 0 | 0 008980686 |
| MATK | 0 004197521 | 0 008246787 | 0 007516129 | 0 493579801 | 0 | 0 009029674 |
| HDC | 0 001256401 | 0 008217071 | 0 007734741 | -0 49539476 | 0 | 0 009288767 |
| LINC00609 | 0 00714816 | 0 008216865 | 0 007736281 | -0 48806288 | 0 | 0 009289436 |
| SERPINI2 | 0 008909025 | 0 008209995 | 0 007787638 | -0 48718374 | 0 | 0 009348727 |
| PPIC | 0 111762578 | 0 008206431 | 0 007814398 | -0 37977775 | 0 | 0 00937966 |
| PCED1B | 0 075602977 | 0 008184081 | 0 007984107 | 0 409375661 | 0 | 0 009580928 |
| ERI2 | 0 153252365 | 0 008171044 | 0 008084623 | 0 304872541 | 0 | 0 009700316 |
| TMEM19 | 0 086777331 | 0 00816484 | 0 008132847 | 0 399283493 | 0 | 0 009756939 |
| FOXO4 | 0 009651444 | 0 008158138 | 0 008185238 | -0 48364932 | 0 | 0 009818545 |
| VPS29 | 0 327892102 | 0 00815389 | 0 008218603 | 0 206813121 | 0 | 0 009857317 |
| CCL5 | 0 016285622 | 0 008150673 | 0 008243949 | 0 48083358 | 0 | 0 009886461 |
| S1PR3 | 0 100312197 | 0 008110504 | 0 008566371 | -0 38457548 | 0 | 0 010271819 |
| CPNE8 | 0 368334888 | 0 008110259 | 0 00856837 | 0 201102536 | 0 | 0 010272913 |
| CLN8 | 0 09877025 | 0 008104428 | 0 008616109 | -0 38802582 | 0 | 0 010326613 |
| SEPHS2 | 0 051588586 | 0 008104388 | 0 008616438 | -0 43866627 | 0 | 0 010326613 |
| PNPLA6 | 0 086843959 | 0 00809491 | 0 008694544 | -0 39870989 | 0 | 0 0104189 |
| PNPLA8 | 0 335925454 | 0 008081567 | 0 008805575 | 0 246717944 | 0 | 0 010547939 |
| CA12 | 0 069159163 | 0 008078012 | 0 008835371 | 0 411020306 | 0 | 0 010582289 |
| CENPM | 0 025070911 | 0 008068721 | 0 008913668 | 0 444113473 | 0 | 0 010674713 |
| OCM | 0 002988711 | 0 008059425 | 0 008992626 | -0 49292469 | 0 | 0 010767906 |
| KRT81 | 0 001275437 | 0 008043554 | 0 009128867 | 0 496237728 | 0 | 0 010925505 |
| HTR1F | 0 00625345 | 0 008042314 | 0 009139592 | 0 491904822 | 0 | 0 010936956 |
| FRAT2 | 0 011402791 | 0 007987121 | 0 009628379 | -0 48373284 | 0 | 0 011516033 |
| LINC00402 | 0 003597876 | 0 007974015 | 0 00974781 | -0 49221331 | 0 | 0 011657404 |
| GRPEL1 | 0 057899145 | 0 007972188 | 0 009764567 | -0 43521988 | 0 | 0 011675966 |
| PLK1 | 0 033313662 | 0 007968878 | 0 00979498 | 0 438626429 | 0 | 0 01171085 |
| NMB | 0 024509337 | 0 007939949 | 0 01006445 | -0 46919346 | 0 | 0 012028094 |
| LINC01440 | 0 007100569 | 0 007939847 | 0 010065415 | -0 487966 | 0 | 0 012028094 |
| LINC01170 | 0 017703832 | 0 007937105 | 0 010091302 | 0 477843076 | 0 | 0 012057505 |
| AKR1B1 | 0 096305039 | 0 007935484 | 0 010106628 | 0 414928137 | 0 | 0 012074291 |
| MMP19 | 0 062391731 | 0 007929928 | 0 010159325 | -0 41943276 | 0 | 0 012135713 |
| MIER1 | 0 412251813 | 0 007928819 | 0 010169874 | 0 172905305 | 0 | 0 012146778 |
| RTKN2 | 0 087919514 | 0 007927075 | 0 010186484 | 0 349447232 | 0 | 0 012165079 |
| MAML3 | 0 489577583 | 0 007920905 | 0 010245429 | -0 0892224 | 1 27E-184 | 0 012233928 |
| RPS6KL1 | 0 009346862 | 0 007919647 | 0 010257489 | 0 489078048 | 0 | 0 01224678 |
| VCPIP1 | 0 148502789 | 0 007919168 | 0 010262079 | 0 34808908 | 0 | 0 012250714 |
| MCMDC2 | 0 041613523 | 0 007917355 | 0 010279489 | 0 461572214 | 0 | 0 012269947 |
| TMEM170B | 0 047457692 | 0 007908099 | 0 01036875 | -0 44066614 | 0 | 0 012374928 |
| PBK | 0 048695056 | 0 007904399 | 0 01040463 | 0 389203403 | 0 | 0 012416183 |
| PRANCR | 0 170299442 | 0 007901887 | 0 010429048 | -0 29506849 | 0 | 0 01244375 |
| BEX1 | 0 018931678 | 0 00789156 | 0 010529977 | 0 478595646 | 0 | 0 012561005 |
| KLHL6 | 0 008947098 | 0 007883528 | 0 010609073 | 0 488593641 | 0 | 0 01265376 |
| FRMD6-AS2 | 0 099198568 | 0 007870526 | 0 010738229 | -0 38893715 | 0 | 0 012806192 |

| | | | | | | |
|---|---|---|---|---|---|---|
| BIK | 0 002826902 | 0 007868054 | 0 010762946 | 0 494314476 | 0 | 0 01283405 |
| ATP2B4 | 0 52985856 | 0 007865869 | 0 010784827 | -0 0450271 | 2 73E-48 | 0 012858519 |
| PVRIG | 0 003778721 | 0 007846697 | 0 010978584 | -0 49231733 | 0 | 0 013087879 |
| PRKAA2 | 0 141697284 | 0 007844456 | 0 011001426 | -0 35095213 | 0 | 0 013113455 |
| HBA1 | 0 00956578 | 0 00783668 | 0 011081036 | -0 48601374 | 0 | 0 013206683 |
| LINC02237 | 0 006339114 | 0 007800862 | 0 011454369 | -0 48974109 | 0 | 0 013643029 |
| TP63 | 0 166063848 | 0 007793774 | 0 011529559 | 0 308195243 | 0 | 0 013729124 |
| MMRN1 | 0 005853686 | 0 007786823 | 0 011603717 | -0 49084168 | 0 | 0 013815689 |
| PPARGC1A | 0 091174735 | 0 007775309 | 0 011727489 | -0 39743803 | 0 | 0 013961296 |
| LINC01579 | 0 033370772 | 0 007757562 | 0 011920559 | -0 46075168 | 0 | 0 014189354 |
| LINC02109 | 0 003378957 | 0 007756882 | 0 01192801 | 0 494452745 | 0 | 0 014196435 |
| FRMD6 | 0 453208582 | 0 007750239 | 0 012001045 | -0 11086279 | 1 65E-284 | 0 014277965 |
| GDPD5 | 0 083664884 | 0 007746492 | 0 012042417 | -0 39854797 | 0 | 0 01432358 |
| HIVEP3 | 0 392882298 | 0 007726499 | 0 012265292 | 0 174250318 | 0 | 0 014585002 |
| F2R | 0 142068493 | 0 007721586 | 0 012320615 | 0 351592416 | 0 | 0 014648945 |
| PRDX6 | 0 277474253 | 0 00770698 | 0 012486392 | -0 22701983 | 0 | 0 014844183 |
| CXCL14 | 0 068312044 | 0 007695162 | 0 01262198 | -0 43016808 | 0 | 0 015001598 |
| SLC39A12 | 0 002008338 | 0 007693917 | 0 012636339 | 0 495646479 | 0 | 0 015016776 |
| FCN3 | 0 00231292 | 0 007690312 | 0 012677991 | 0 495217138 | 0 | 0 01506438 |
| LINC00612 | 0 010089281 | 0 007656546 | 0 013074132 | -0 48493556 | 0 | 0 015531182 |
| SHB | 0 109763759 | 0 007652468 | 0 013122703 | 0 382264684 | 0 | 0 015584963 |
| MYO5A | 0 391197579 | 0 007650249 | 0 013149202 | 0 185223423 | 0 | 0 01561447 |
| RAMP2 | 0 07332813 | 0 007644238 | 0 013221217 | -0 42055595 | 0 | 0 015698015 |
| WNT9A | 0 048656983 | 0 007643082 | 0 013235116 | -0 44120885 | 0 | 0 01571057 |
| TMEM87A | 0 48149664 | 0 007608148 | 0 013661 | -0 09664121 | 2 17E-216 | 0 016210001 |
| LINC01150 | 0 001656165 | 0 007593709 | 0 013840536 | -0 49458267 | 0 | 0 016420975 |
| TJP1 | 0 566579734 | 0 007589059 | 0 013898799 | 0 10624738 | 2 34E-261 | 0 01648803 |
| PLEC | 0 290913937 | 0 007566442 | 0 014185268 | -0 22740623 | 0 | 0 016825754 |
| MEFV | 0 001237365 | 0 00756621 | 0 014188246 | 0 496129013 | 0 | 0 016827174 |
| DCK | 0 046896118 | 0 007565705 | 0 0141947 | 0 446442634 | 0 | 0 016832716 |
| LINC01644 | 0 002017856 | 0 007563586 | 0 014221822 | -0 49440043 | 0 | 0 016862762 |
| TNFRSF13B | 0 001817974 | 0 00755937 | 0 014275914 | -0 49522058 | 0 | 0 016924776 |
| ARHGAP26-AS | 0 033427881 | 0 007559161 | 0 014278597 | -0 45646599 | 0 | 0 016925833 |
| ZAP70 | 0 004530658 | 0 007548716 | 0 01441342 | -0 49156447 | 0 | 0 017081367 |
| LINC00989 | 0 003902458 | 0 007543009 | 0 014487569 | -0 49211865 | 0 | 0 017167088 |
| TLR5 | 0 022367745 | 0 00753479 | 0 014594928 | -0 47223091 | 0 | 0 017292135 |
| IPMK | 0 168824123 | 0 007499354 | 0 015065899 | -0 32880838 | 0 | 0 017843432 |
| FKBP4 | 0 095029602 | 0 007475973 | 0 015383931 | 0 405082267 | 0 | 0 01821553 |
| RIMS3 | 0 006577069 | 0 007463195 | 0 015560228 | 0 491464309 | 0 | 0 018421968 |
| DCC | 0 078563134 | 0 007444853 | 0 01581639 | -0 40841593 | 0 | 0 018718208 |
| LINC00426 | 0 005568141 | 0 007440271 | 0 015880967 | -0 49037404 | 0 | 0 018792279 |
| MS4A6E | 0 000932783 | 0 00743899 | 0 015899053 | -0 49581442 | 0 | 0 018811325 |
| ATP1A1-AS1 | 0 129970874 | 0 00741423 | 0 016252356 | -0 35794839 | 0 | 0 01922453 |
| CTNND1 | 0 640726428 | 0 007360539 | 0 017042243 | -0 06097357 | 4 28E-87 | 0 020151301 |

| | | | | | | |
|---|---|---|---|---|---|---|
| DYSF | 0 038862767 | 0 007360271 | 0 017046263 | -0 44631538 | 0 | 0 020153532 |
| ZCCHC12 | 0 000828082 | 0 007359221 | 0 017062042 | 0 496840393 | 0 | 0 020169665 |
| ARG2 | 0 067560107 | 0 007356601 | 0 017101469 | 0 419143163 | 0 | 0 020213744 |
| SLC43A2 | 0 055748034 | 0 007337217 | 0 017395707 | -0 42618274 | 0 | 0 020551248 |
| BFSP2-AS1 | 0 000951819 | 0 00732745 | 0 01754564 | 0 496404873 | 0 | 0 020725787 |
| TMPRSS3 | 0 000694828 | 0 007324466 | 0 01759167 | 0 496734123 | 0 | 0 020777563 |
| TSIX | 0 133663932 | 0 007318465 | 0 017684557 | 0 350616152 | 0 | 0 020882052 |
| SRFBP1 | 0 296853287 | 0 007311596 | 0 017791417 | 0 240392827 | 0 | 0 021005608 |
| CASP3 | 0 108079039 | 0 00731086 | 0 017802894 | 0 384683911 | 0 | 0 021016534 |
| TNFAIP8 | 0 195789153 | 0 007299994 | 0 017973187 | 0 273299653 | 0 | 0 021214915 |
| MAT2A | 0 340075384 | 0 007297066 | 0 018019316 | 0 202857291 | 0 | 0 021266708 |
| UBE2D2 | 0 471064705 | 0 007284942 | 0 018211437 | 0 142826257 | 0 | 0 021490769 |
| TUBD1 | 0 165292875 | 0 007280462 | 0 018282873 | 0 301292492 | 0 | 0 021572375 |
| MORF4L1 | 0 5113552 | 0 007278256 | 0 018318141 | -0 09568526 | 3 76E-212 | 0 02161129 |
| PKN3 | 0 048971084 | 0 007268037 | 0 018482297 | 0 427955984 | 0 | 0 021796793 |
| PCK1 | 0 002703166 | 0 007265424 | 0 018524476 | 0 494621491 | 0 | 0 021843811 |
| IGHA1 | 0 005910796 | 0 007234812 | 0 019024944 | -0 49094087 | 0 | 0 022419968 |
| MCM5 | 0 060754602 | 0 007234371 | 0 019032244 | 0 385626402 | 0 | 0 022425774 |
| LINC02363 | 0 004245112 | 0 007224178 | 0 01920154 | 0 494095193 | 0 | 0 022619615 |
| HCG27 | 0 029525423 | 0 007219221 | 0 019284345 | 0 463014765 | 0 | 0 022714329 |
| ERMP1 | 0 203260932 | 0 00720722 | 0 019486132 | 0 315325711 | 0 | 0 022949146 |
| GUK1 | 0 24584531 | 0 007201254 | 0 019587112 | -0 26686594 | 0 | 0 02305945 |
| GIMAP1 | 0 006282005 | 0 007200256 | 0 019604059 | -0 49010666 | 0 | 0 023076527 |
| ANO9 | 0 006577069 | 0 007198677 | 0 019630889 | -0 48905631 | 0 | 0 023105231 |
| SLC16A12 | 0 185300109 | 0 007176218 | 0 020015935 | -0 28950486 | 0 | 0 023549625 |
| LENG1 | 0 038786621 | 0 00716782 | 0 020161589 | -0 45266178 | 0 | 0 023715088 |
| FNDC3A | 0 699529801 | 0 007165223 | 0 020206822 | -0 0286182 | 1 73E-20 | 0 023765336 |
| ASPM | 0 113123679 | 0 007148211 | 0 020505319 | 0 315700017 | 0 | 0 024113397 |
| NBEAL1 | 0 633892368 | 0 007125431 | 0 020911026 | -0 04871606 | 3 13E-56 | 0 024581316 |
| PHKG2 | 0 082180046 | 0 007119088 | 0 021025246 | 0 407222198 | 0 | 0 02471251 |
| LINC00996 | 0 000942301 | 0 007115975 | 0 021081486 | -0 49580019 | 0 | 0 02477553 |
| KLRC3 | 0 008385525 | 0 00710103 | 0 021353366 | -0 4870393 | 0 | 0 025088811 |
| IDS | 0 305077002 | 0 00709382 | 0 021485606 | -0 1754649 | 0 | 0 025241046 |
| IGKC | 0 031200624 | 0 007071088 | 0 021907257 | -0 46917581 | 0 | 0 025723606 |
| DELEC1 | 0 054977061 | 0 007070238 | 0 02192316 | -0 43023791 | 0 | 0 025739081 |
| CNTN1 | 0 209790409 | 0 007057783 | 0 022157364 | -0 25338194 | 0 | 0 026007588 |
| GPR18 | 0 020045307 | 0 007049816 | 0 022308311 | 0 477730169 | 0 | 0 026181513 |
| MARS2 | 0 007795397 | 0 007041172 | 0 022473103 | 0 489727699 | 0 | 0 026368366 |
| PRR14 | 0 05309246 | 0 007030971 | 0 022668936 | -0 43365536 | 0 | 0 026594841 |
| DPYD-AS1 | 0 227418096 | 0 007018429 | 0 02291173 | 0 223112557 | 0 | 0 026873011 |
| TNFRSF10A | 0 03569321 | 0 007007243 | 0 023130186 | 0 464383266 | 0 | 0 027122504 |
| ADGRE3 | 0 004711504 | 0 006992436 | 0 023422137 | 0 493242687 | 0 | 0 027458032 |
| LRRC8A | 0 099179532 | 0 006992241 | 0 023426019 | -0 39121985 | 0 | 0 027459176 |
| C4BPB | 0 001808456 | 0 006989298 | 0 023484423 | -0 49477236 | 0 | 0 027524221 |

| | | | | | | |
|---|---|---|---|---|---|---|
| LINC01933 | 0 040300013 | 0 006963667 | 0 023998549 | -0 45192489 | 0 | 0 028112841 |
| NUDT4 | 0 318830786 | 0 006962024 | 0 024031828 | -0 22031855 | 0 | 0 028148336 |
| ACSL6 | 0 006586587 | 0 006960767 | 0 024057314 | 0 493331235 | 0 | 0 028174695 |
| TNFSF8 | 0 003664503 | 0 00695389 | 0 024197197 | 0 494346689 | 0 | 0 028335008 |
| LINC02516 | 0 084940321 | 0 006953542 | 0 024204294 | -0 39697285 | 0 | 0 028339806 |
| TFEC | 0 020283261 | 0 006944015 | 0 02439931 | 0 475426168 | 0 | 0 028561065 |
| LINC01366 | 0 002617502 | 0 00694052 | 0 024471179 | -0 49377726 | 0 | 0 028638098 |
| BIN2 | 0 008499743 | 0 006921338 | 0 024868963 | -0 48795224 | 0 | 0 029085606 |
| TBXAS1 | 0 113561516 | 0 006918549 | 0 024927274 | 0 34580319 | 0 | 0 029150197 |
| PLBD1 | 0 109906531 | 0 006909981 | 0 025107098 | -0 38471842 | 0 | 0 02935322 |
| RIPK2 | 0 17068017 | 0 006899719 | 0 025323974 | 0 311004559 | 0 | 0 029603112 |
| GPR160 | 0 019997716 | 0 006898869 | 0 025342018 | 0 473216001 | 0 | 0 029620541 |
| RASSF8 | 0 499076736 | 0 006895903 | 0 025405045 | -0 09683031 | 3 12E-217 | 0 029690536 |
| CDK9 | 0 078039634 | 0 006880652 | 0 025731244 | -0 4102623 | 0 | 0 030064325 |
| ADA2 | 0 034627173 | 0 006878968 | 0 025767478 | -0 45570193 | 0 | 0 030102939 |
| ACKR3 | 0 070244237 | 0 006815392 | 0 027168332 | -0 4115539 | 0 | 0 031712047 |
| FAM221A | 0 064714169 | 0 006778373 | 0 028013927 | -0 42617361 | 0 | 0 03268695 |
| LINC00449 | 0 001760865 | 0 006769976 | 0 028208859 | 0 495754572 | 0 | 0 032904451 |
| TRMT10C | 0 075069959 | 0 006767685 | 0 028262228 | 0 42820122 | 0 | 0 032960392 |
| KIAA0513 | 0 078353734 | 0 006759927 | 0 028443677 | -0 40239365 | 0 | 0 03316791 |
| CEP152 | 0 165407093 | 0 006758931 | 0 028467039 | 0 285590686 | 0 | 0 033191058 |
| KCTD5 | 0 055681407 | 0 006754273 | 0 028576526 | 0 446375908 | 0 | 0 033310494 |
| ZNF700 | 0 202528031 | 0 006744126 | 0 028816308 | 0 261714272 | 0 | 0 033585855 |
| NEURL2 | 0 017789496 | 0 006731357 | 0 029120481 | -0 47697798 | 0 | 0 033923638 |
| XKR8 | 0 019540843 | 0 006708379 | 0 029674855 | -0 47566669 | 0 | 0 034565189 |
| CCR4 | 0 000666273 | 0 006694372 | 0 030017196 | 0 49657022 | 0 | 0 03495533 |
| OSBPL9 | 0 546696237 | 0 006689921 | 0 030126705 | -0 05095335 | 2 38E-61 | 0 035078532 |
| IRF7 | 0 035950201 | 0 006669611 | 0 030630713 | -0 45563279 | 0 | 0 035660987 |
| NUAK2 | 0 007119606 | 0 006667504 | 0 030683421 | 0 488977905 | 0 | 0 035717951 |
| SRSF3 | 0 40616969 | 0 006660923 | 0 030848517 | -0 14308325 | 0 | 0 035905713 |
| ATP1A3 | 0 003150521 | 0 006633479 | 0 031545288 | 0 494215727 | 0 | 0 036694113 |
| CARD9 | 0 003778721 | 0 006605715 | 0 032263875 | -0 49258449 | 0 | 0 037506903 |
| CSRNP1 | 0 06186823 | 0 006591331 | 0 032641646 | -0 41718006 | 0 | 0 037936731 |
| CCL7 | 0 000361691 | 0 006585564 | 0 032794162 | 0 49665405 | 0 | 0 0381093 |
| ADARB2 | 0 059269765 | 0 006582706 | 0 03286997 | -0 41681921 | 0 | 0 038192698 |
| NAGA | 0 038310712 | 0 006578801 | 0 03297381 | 0 463299032 | 0 | 0 038303934 |
| CORO1A | 0 012307019 | 0 006570521 | 0 033194887 | -0 48435876 | 0 | 0 03855127 |
| CDKN2C | 0 073623194 | 0 006567312 | 0 033280902 | 0 394179423 | 0 | 0 038641666 |
| GPRC5B | 0 084645257 | 0 006539273 | 0 03404067 | -0 40696566 | 0 | 0 039509252 |
| MMP26 | 0 003141002 | 0 006527055 | 0 034376347 | 0 494759071 | 0 | 0 039893955 |
| SLC38A8 | 0 002855457 | 0 006522933 | 0 034490241 | -0 49331095 | 0 | 0 040016302 |
| ATP13A4 | 0 020940016 | 0 00652205 | 0 034514688 | -0 47055111 | 0 | 0 04003975 |
| VNN1 | 0 002950639 | 0 006521844 | 0 03452038 | 0 493746483 | 0 | 0 040041438 |
| ANKRD29 | 0 104871409 | 0 006521398 | 0 034532723 | 0 366502445 | 0 | 0 040050838 |

| | | | | | | |
|---|---|---|---|---|---|---|
| ELF2 | 0 560668938 | 0 006509616 | 0 034860375 | 0 086087852 | 5 66E-172 | 0 040420926 |
| AGPS | 0 41179494 | 0 006501865 | 0 035077359 | 0 164371871 | 0 | 0 040667531 |
| LINC02018 | 0 081180636 | 0 00649841 | 0 035174453 | -0 4089033 | 0 | 0 040775096 |
| SGTB | 0 095134302 | 0 006490547 | 0 035396278 | 0 39867501 | 0 | 0 041027209 |
| LINC02147 | 0 090061107 | 0 006485792 | 0 035530998 | -0 3912759 | 0 | 0 041178309 |
| LCNL1 | 0 020578325 | 0 006446885 | 0 036649886 | -0 46637249 | 0 | 0 042437289 |
| RBBP8 | 0 172736099 | 0 006433094 | 0 037053613 | 0 299991614 | 0 | 0 042885076 |
| CRYBB1 | 0 001837011 | 0 006402364 | 0 037966887 | 0 495600262 | 0 | 0 043920564 |
| RAP2B | 0 083455483 | 0 006398244 | 0 038090765 | 0 41998278 | 0 | 0 044058473 |
| GPBAR1 | 0 00189412 | 0 006394796 | 0 038194707 | -0 49436618 | 0 | 0 044173294 |
| MS4A14 | 0 001875083 | 0 006371738 | 0 038896036 | -0 49465099 | 0 | 0 044973393 |
| CDKN2AIP | 0 039072167 | 0 006370612 | 0 03893057 | -0 45398843 | 0 | 0 045007816 |
| ATP6V1B2 | 0 15990558 | 0 006340273 | 0 039870698 | -0 32548117 | 0 | 0 046083427 |
| MYBPC3 | 0 001284956 | 0 006332361 | 0 040119018 | 0 495955328 | 0 | 0 046364769 |
| FCGRT | 0 270573566 | 0 006314815 | 0 040674373 | -0 22211304 | 0 | 0 046989343 |
| ANKRD20A11P | 0 037159011 | 0 00631386 | 0 040704788 | -0 45861431 | 0 | 0 047018732 |
| MYBPC1 | 0 024042946 | 0 006311923 | 0 040766549 | -0 47117314 | 0 | 0 047084318 |
| TMC8 | 0 009032762 | 0 006295814 | 0 041283124 | 0 489074305 | 0 | 0 047669296 |
| JRKL | 0 055253089 | 0 006293719 | 0 041350716 | 0 451544321 | 0 | 0 047735679 |
| THAP11 | 0 030324951 | 0 006283699 | 0 041675292 | -0 46121284 | 0 | 0 048104496 |
| IL23R | 0 001532428 | 0 006278248 | 0 041852768 | -0 4952953 | 0 | 0 048303452 |
| KCNK3 | 0 001142183 | 0 006274184 | 0 041985489 | 0 49611116 | 0 | 0 048450711 |
| GZMM | 0 004178485 | 0 006240082 | 0 043113344 | -0 49211066 | 0 | 0 049727951 |
| CMPK2 | 0 01698045 | 0 006235638 | 0 043262207 | 0 476103256 | 0 | 0 049893563 |
| FPR1 | 0 009242162 | 0 006225926 | 0 043589 | 0 486250837 | 0 | 0 050264314 |
| CDV3 | 0 213968895 | 0 006212517 | 0 044043624 | -0 28423681 | 0 | 0 050782363 |
| VPS13A | 0 417486817 | 0 006209186 | 0 044157194 | 0 130867911 | 0 | 0 050907098 |
| MAFF | 0 07595515 | 0 00619144 | 0 044766323 | -0 40488989 | 0 | 0 05159675 |
| HMGB2 | 0 149121471 | 0 006186785 | 0 04492728 | 0 293483753 | 0 | 0 051769638 |
| LONRF1 | 0 337476918 | 0 006184917 | 0 044992023 | -0 15279343 | 0 | 0 051837919 |
| YDJC | 0 014467648 | 0 006184626 | 0 045002107 | 0 481959886 | 0 | 0 051843217 |
| EFEMP2 | 0 161181017 | 0 006171295 | 0 045466407 | -0 32672658 | 0 | 0 05235257 |
| HYAL3 | 0 00473054 | 0 006167908 | 0 045584997 | 0 493905627 | 0 | 0 052482727 |
| ARRDC4 | 0 054777179 | 0 006156768 | 0 045976926 | -0 43400247 | 0 | 0 052921066 |
| RLN1 | 0 00120881 | 0 006154547 | 0 046055413 | 0 496155633 | 0 | 0 053004951 |
| TOPORS | 0 105442501 | 0 006152278 | 0 046135722 | 0 400094708 | 0 | 0 053090913 |
| TMOD2 | 0 140041119 | 0 006148348 | 0 046275058 | -0 34398218 | 0 | 0 053242287 |
| PLCH2 | 0 004330776 | 0 00614825 | 0 046278534 | -0 49170491 | 0 | 0 053242287 |
| KLHL13 | 0 323323371 | 0 006141177 | 0 046530235 | -0 19314393 | 0 | 0 053518832 |
| PTCRA | 0 000723382 | 0 006124638 | 0 047123309 | -0 49601697 | 0 | 0 054194386 |
| LINC02446 | 0 001408692 | 0 006122691 | 0 047193517 | 0 495721601 | 0 | 0 054261922 |
| GPD1L | 0 071824256 | 0 00611122 | 0 047609106 | -0 41354727 | 0 | 0 054726439 |
| HK1 | 0 21120862 | 0 006109158 | 0 047684129 | -0 28808711 | 0 | 0 054799345 |
| ZDHHC19 | 0 020492661 | 0 006106732 | 0 04777253 | -0 46978804 | 0 | 0 054894262 |

| | | | | | | |
|---|---|---|---|---|---|---|
| SPOCK2 | 0 019655061 | 0 0061064 | 0 04778463 | -0 47304675 | 0 | 0 05490149 |
| SLC4A1 | 0 002227256 | 0 006103127 | 0 047904119 | -0 49444029 | 0 | 0 05503082 |
| B3GNT8 | 0 003074375 | 0 006099223 | 0 048047007 | -0 49323223 | 0 | 0 055182817 |
| CLIC3 | 0 00352173 | 0 006072371 | 0 04903942 | -0 49259036 | 0 | 0 056295252 |
| GNMT | 0 004340294 | 0 006072022 | 0 049052432 | 0 492200584 | 0 | 0 056301741 |
| CD40LG | 0 000970855 | 0 006067806 | 0 049209832 | -0 49576108 | 0 | 0 056470299 |
| IL1RL1 | 0 002303402 | 0 006066083 | 0 049274298 | -0 49392189 | 0 | 0 056530547 |
| PPME1 | 0 370628772 | 0 006062125 | 0 049422623 | 0 186143704 | 0 | 0 056693832 |
| LINC02580 | 0 061839676 | 0 006061154 | 0 049459062 | -0 41879642 | 0 | 0 056728746 |
| PRPF38B | 0 282899621 | 0 006043478 | 0 050126436 | 0 237834301 | 0 | 0 057473285 |
| ZNF845 | 0 037311302 | 0 006042436 | 0 050166017 | 0 45515128 | 0 | 0 057511691 |
| THBS2 | 0 27044983 | 0 00602977 | 0 050649189 | -0 25161636 | 0 | 0 058051526 |
| TMUB2 | 0 125935162 | 0 006022091 | 0 050944001 | 0 384694471 | 0 | 0 058375265 |
| TRBV28 | 5 71E-05 | 0 005984784 | 0 052396914 | 0 496943584 | 0 | 0 060018283 |
| PZP | 0 010089281 | 0 005966269 | 0 053130722 | -0 48520493 | 0 | 0 060851451 |
| KLF3 | 0 236022539 | 0 005947933 | 0 053865878 | 0 277281835 | 0 | 0 061671013 |
| VWA5A | 0 027183949 | 0 005943139 | 0 054059494 | -0 4639848 | 0 | 0 061885187 |
| OR51E1 | 0 000571091 | 0 00593825 | 0 054257494 | -0 49618823 | 0 | 0 062096805 |
| PLD4 | 0 005444404 | 0 005936638 | 0 054322954 | -0 49057168 | 0 | 0 062156667 |
| CLVS1 | 0 021615808 | 0 005911905 | 0 055335102 | -0 4741907 | 0 | 0 063291784 |
| N4BP2L1 | 0 151234509 | 0 00588292 | 0 056541256 | -0 32574803 | 0 | 0 06465572 |
| CCR3 | 0 001541947 | 0 005877946 | 0 056750396 | 0 495478437 | 0 | 0 064879173 |
| TRGC2 | 0 002607984 | 0 005849827 | 0 057944985 | 0 494205164 | 0 | 0 066236859 |
| LINC02315 | 0 084940321 | 0 00584704 | 0 058064516 | -0 41678826 | 0 | 0 066365466 |
| PPP1R1C | 0 03369439 | 0 005845666 | 0 058123542 | 0 464281633 | 0 | 0 066424897 |
| SPSB1 | 0 177799775 | 0 005829778 | 0 058809465 | -0 31332002 | 0 | 0 067176289 |
| LINC00115 | 0 039433858 | 0 005826325 | 0 058959402 | 0 452884992 | 0 | 0 067339418 |
| SERTAD1 | 0 059688565 | 0 005818586 | 0 059296633 | -0 42460477 | 0 | 0 067716397 |
| LINC02357 | 0 000713864 | 0 005814112 | 0 059492353 | 0 496671585 | 0 | 0 067931698 |
| F11R | 0 027117321 | 0 005808325 | 0 059746256 | -0 46501203 | 0 | 0 068196898 |
| LINC00461 | 0 081247264 | 0 005793935 | 0 060381559 | 0 400612981 | 0 | 0 068897095 |
| PADI4 | 0 002141593 | 0 005791177 | 0 060503959 | -0 49425843 | 0 | 0 069028421 |
| RAPGEF4 | 0 212303211 | 0 005766028 | 0 061629559 | -0 27184171 | 0 | 0 07027867 |
| TNIP1 | 0 152129219 | 0 005760818 | 0 0618649 | -0 3449675 | 0 | 0 070530016 |
| LINC01819 | 0 03611201 | 0 005752108 | 0 062260014 | -0 44888021 | 0 | 0 070954792 |
| MAMDC2-AS1 | 0 203051531 | 0 005750278 | 0 062343279 | 0 28873571 | 0 | 0 071041118 |
| LRRFIP2 | 0 539157831 | 0 005742349 | 0 062705194 | 0 107181668 | 5 81E-266 | 0 071436297 |
| ATG16L1 | 0 134806114 | 0 005734085 | 0 063084206 | 0 367751213 | 0 | 0 071850759 |
| DYRK4 | 0 064895014 | 0 005699356 | 0 064697781 | 0 421358637 | 0 | 0 073653057 |
| LINC02356 | 0 012240391 | 0 005692 | 0 065043913 | -0 4826947 | 0 | 0 074038181 |
| HIRIP3 | 0 02927795 | 0 005657066 | 0 066708532 | 0 465403919 | 0 | 0 075923838 |
| PLCXD2 | 0 021777617 | 0 005648394 | 0 067127139 | 0 465213341 | 0 | 0 076381874 |
| ARMC3 | 0 024157164 | 0 005645062 | 0 067288526 | -0 46876672 | 0 | 0 076556293 |
| TMEM70 | 0 064133559 | 0 005642264 | 0 067424329 | 0 441821033 | 0 | 0 076701566 |

| | | | | | | |
|---|---|---|---|---|---|---|
| LINC01136 | 0 000218918 | 0 005621261 | 0 068450864 | -0 49670619 | 0 | 0 077831869 |
| IFNG | 0 000885192 | 0 005617153 | 0 068653122 | -0 49597603 | 0 | 0 078052454 |
| SLC46A3 | 0 052797396 | 0 005614938 | 0 068762423 | -0 43499971 | 0 | 0 078167316 |
| WDR54 | 0 047391064 | 0 005612544 | 0 068880665 | 0 453912256 | 0 | 0 078292312 |
| IL6R-AS1 | 0 001408692 | 0 005565733 | 0 071226844 | -0 49480937 | 0 | 0 080929864 |
| ADRA2B | 0 001351583 | 0 005555203 | 0 07176352 | -0 49521547 | 0 | 0 081520046 |
| KDR | 0 05461537 | 0 005534093 | 0 07284943 | -0 43256897 | 0 | 0 082743644 |
| PPP1CC | 0 269155356 | 0 005526231 | 0 073257292 | 0 241909753 | 0 | 0 083186903 |
| LINC02232 | 0 021568217 | 0 005514028 | 0 073894005 | -0 47279685 | 0 | 0 083899837 |
| LINC01762 | 0 014258248 | 0 005495025 | 0 074894572 | -0 48025093 | 0 | 0 084995041 |
| GSKIP | 0 059307837 | 0 005492175 | 0 075045593 | 0 442895111 | 0 | 0 085135756 |
| CA5A | 0 003017266 | 0 005487838 | 0 075275827 | -0 49339124 | 0 | 0 085386697 |
| TRPM2 | 0 013163656 | 0 005486559 | 0 075343859 | -0 48220469 | 0 | 0 085453609 |
| FLACC1 | 0 051550513 | 0 005475782 | 0 075919008 | 0 448858087 | 0 | 0 086095599 |
| FCMR | 0 002712684 | 0 005457463 | 0 076904831 | 0 494863881 | 0 | 0 087203102 |
| NME8 | 0 002322438 | 0 005454456 | 0 077067682 | -0 49400163 | 0 | 0 087377276 |
| STPG2 | 0 078972416 | 0 005445625 | 0 077547444 | -0 40735731 | 0 | 0 087900126 |
| SPI1 | 0 023348118 | 0 005435479 | 0 078101652 | -0 46950084 | 0 | 0 088507088 |
| LINC01645 | 0 00210352 | 0 005425197 | 0 078666627 | -0 49430956 | 0 | 0 089125959 |
| ARRB1 | 0 12179475 | 0 005418968 | 0 079010483 | -0 36141562 | 0 | 0 089504803 |
| BROX | 0 148902553 | 0 005379759 | 0 081203121 | -0 34347952 | 0 | 0 091953635 |
| R3HDM4 | 0 062125221 | 0 005379617 | 0 081211115 | -0 42555468 | 0 | 0 091953635 |
| MRPL10 | 0 061649312 | 0 005373672 | 0 08154788 | -0 43309495 | 0 | 0 092323884 |
| FILNC1 | 0 001856047 | 0 005364682 | 0 082059255 | -0 49441634 | 0 | 0 092858336 |
| KIR3DL2 | 0 0001047 | 0 005340757 | 0 083432937 | -0 49677194 | 0 | 0 09439019 |
| EPHA1 | 0 004844758 | 0 005323635 | 0 084427333 | -0 49128925 | 0 | 0 095480887 |
| CLEC3B | 0 016447431 | 0 005303465 | 0 085611087 | -0 47811082 | 0 | 0 096796455 |
| TMEM191C | 0 005615732 | 0 005278273 | 0 087108341 | 0 490628599 | 0 | 0 098453992 |
| KRT19 | 0 007823952 | 0 005276593 | 0 087208974 | -0 48602089 | 0 | 0 098555944 |
| LTB | 0 004530658 | 0 005275594 | 0 087268799 | -0 49188067 | 0 | 0 09861176 |
| RASGRF1 | 0 019455179 | 0 005262652 | 0 088047376 | -0 47190698 | 0 | 0 099452492 |
| NLRC4 | 0 019912052 | 0 005262527 | 0 088054921 | 0 469435269 | 0 | 0 099452492 |
| EXPH5 | 0 035816946 | 0 005249781 | 0 088827161 | 0 455724593 | 0 | 0 1003127 |
| NCAM2 | 0 250671032 | 0 005232915 | 0 089857415 | -0 25576441 | 0 | 0 1014398 |
| SUSD2 | 0 005663323 | 0 005230415 | 0 090010975 | -0 48957997 | 0 | 0 101601017 |
| PLK3 | 0 092307399 | 0 005221705 | 0 090547516 | -0 38851303 | 0 | 0 102170033 |
| NUP62 | 0 112276561 | 0 005219216 | 0 090701292 | 0 361118659 | 0 | 0 102319113 |
| LEAP2 | 0 02402391 | 0 00516291 | 0 094236954 | 0 469940016 | 0 | 0 106269596 |
| SLC9A3R1 | 0 007757324 | 0 005151781 | 0 094948692 | 0 487446724 | 0 | 0 107059437 |
| LINC00886 | 0 109068931 | 0 005151433 | 0 094971023 | -0 38277301 | 0 | 0 107071841 |
| RASGRF2 | 0 210675601 | 0 005145017 | 0 095383383 | -0 29065285 | 0 | 0 107511089 |
| USP32 | 0 659344006 | 0 005140527 | 0 095672781 | 0 036480626 | 2 79E-32 | 0 107824422 |
| TCAF2 | 0 01771335 | 0 005108141 | 0 097781238 | 0 480710252 | 0 | 0 110135006 |
| LINC01094 | 0 005653804 | 0 005097675 | 0 098470522 | 0 490721429 | 0 | 0 110898157 |

| | | | | | | |
|---|---|---|---|---|---|---|
| CCNI2 | 0 000837601 | 0 005096306 | 0 098560951 | 0 496541695 | 0 | 0 110986771 |
| FBXL2 | 0 358664408 | 0 005085109 | 0 099303225 | -0 18473244 | 0 | 0 111809302 |
| KIF18A | 0 067360225 | 0 005081909 | 0 09951618 | 0 365854256 | 0 | 0 112035728 |
| IL1R2 | 0 005739468 | 0 005077812 | 0 099789313 | 0 491095631 | 0 | 0 11232984 |
| PMEPA1 | 0 228617388 | 0 005076331 | 0 099888186 | 0 225476942 | 0 | 0 112427746 |
| DUSP6 | 0 230892235 | 0 005074321 | 0 10002255 | -0 2581029 | 0 | 0 112565571 |
| FAM177B | 0 004425958 | 0 005072846 | 0 100121223 | 0 492090932 | 0 | 0 1126632 |
| MAP4K3-DT | 0 303535055 | 0 005066858 | 0 100522573 | -0 20081803 | 0 | 0 113101359 |
| CD80 | 0 00493994 | 0 005058155 | 0 101108279 | 0 49247912 | 0 | 0 113719737 |
| C17orf107 | 0 003540766 | 0 005053713 | 0 101408223 | -0 49284972 | 0 | 0 114016384 |
| VPS37B | 0 276370143 | 0 005052613 | 0 101482616 | 0 262069042 | 0 | 0 114086453 |
| FGF20 | 0 001284956 | 0 005049214 | 0 10171274 | 0 49586824 | 0 | 0 114317958 |
| STX7 | 0 31160648 | 0 005037831 | 0 102486539 | -0 1738875 | 0 | 0 115160261 |
| PRND | 0 002512802 | 0 005032435 | 0 102854969 | 0 49441379 | 0 | 0 115546773 |
| CX3CL1 | 0 015657421 | 0 00503053 | 0 10298531 | 0 480603762 | 0 | 0 115679446 |
| DSG2 | 0 060145438 | 0 005013385 | 0 104164072 | 0 429237619 | 0 | 0 116961797 |
| IFIT2 | 0 042755706 | 0 005004141 | 0 104804057 | -0 45133346 | 0 | 0 11766643 |
| TMEM150B | 0 003179075 | 0 004998672 | 0 10518417 | -0 4933073 | 0 | 0 118065139 |
| DGCR6L | 0 045515981 | 0 004997155 | 0 105289823 | -0 44973058 | 0 | 0 118169694 |
| GFI1 | 0 002541357 | 0 004989611 | 0 105816355 | 0 494339358 | 0 | 0 118746533 |
| JMJD6 | 0 095695875 | 0 00498226 | 0 106331468 | 0 400036793 | 0 | 0 119296257 |
| RMI1 | 0 098760732 | 0 00497827 | 0 106611849 | 0 342707562 | 0 | 0 119582429 |
| RBP1 | 0 066532143 | 0 004972806 | 0 106996777 | -0 42005827 | 0 | 0 119997786 |
| RPA2 | 0 125792389 | 0 004972474 | 0 107020247 | 0 316976615 | 0 | 0 119997786 |
| CIDEB | 0 010755554 | 0 004971439 | 0 107093259 | -0 48431618 | 0 | 0 120065405 |
| VWC2 | 0 063362586 | 0 004964375 | 0 107592948 | -0 42006712 | 0 | 0 120611311 |
| USP31 | 0 191315604 | 0 004937976 | 0 109476603 | 0 295679536 | 0 | 0 122592001 |
| NUGGC | 0 001932192 | 0 004929368 | 0 110096416 | -0 49471693 | 0 | 0 123271461 |
| FGR | 0 007290933 | 0 004922535 | 0 110590348 | -0 48812739 | 0 | 0 123795166 |
| NECAB1 | 0 091241362 | 0 004919347 | 0 110821443 | 0 392604365 | 0 | 0 124039161 |
| ARRDC5 | 0 00299823 | 0 004918655 | 0 110871662 | -0 49331854 | 0 | 0 124080674 |
| IL4 | 0 001837011 | 0 004916395 | 0 111035758 | -0 49488422 | 0 | 0 124249605 |
| PKD2L1 | 0 002132074 | 0 004905938 | 0 111797489 | -0 49428495 | 0 | 0 125087174 |
| C11orf45 | 0 001027964 | 0 00489541 | 0 112568588 | 0 495249152 | 0 | 0 125905215 |
| FSTL3 | 0 136452761 | 0 004891937 | 0 112823861 | -0 33766847 | 0 | 0 126175797 |
| PSME2 | 0 136928671 | 0 004885149 | 0 113324145 | 0 362738747 | 0 | 0 12672029 |
| ZNF330 | 0 137499762 | 0 004872048 | 0 114294649 | -0 35878543 | 0 | 0 127775279 |
| GRIK3 | 0 11364718 | 0 004861901 | 0 115050807 | -0 33398003 | 0 | 0 128574991 |
| CLEC4D | 0 000694828 | 0 004854153 | 0 115630819 | -0 49595078 | 0 | 0 129207903 |
| CD3G | 0 003226666 | 0 004845112 | 0 116310502 | 0 493827709 | 0 | 0 129952027 |
| THEMIS | 0 01004169 | 0 004835503 | 0 117036373 | 0 486420811 | 0 | 0 130747574 |
| C1orf162 | 0 017751423 | 0 004828587 | 0 117560988 | -0 47787786 | 0 | 0 131302604 |
| FOXO1 | 0 536854429 | 0 004824285 | 0 117888318 | -0 02591515 | 4 42E-17 | 0 13163708 |
| OTULINL | 0 050113266 | 0 004809009 | 0 11905613 | 0 45995899 | 0 | 0 132909679 |

| | | | | | | |
|---|---|---|---|---|---|---|
| EIF4G2 | 0 410300584 | 0 004801978 | 0 119596724 | -0 15016164 | 0 | 0 13348164 |
| KHK | 0 011117245 | 0 004801632 | 0 119623355 | 0 485234259 | 0 | 0 133495595 |
| SIGLEC1 | 0 010917363 | 0 00479931 | 0 119802344 | -0 48512886 | 0 | 0 133679555 |
| ANKRD62 | 0 009870362 | 0 004797739 | 0 11992353 | 0 479279922 | 0 | 0 13379898 |
| CLEC10A | 0 002588947 | 0 004784341 | 0 120961164 | 0 494904072 | 0 | 0 134892966 |
| FGD2 | 0 006443814 | 0 004779208 | 0 121360555 | -0 48965931 | 0 | 0 135322388 |
| TRIM60 | 0 002065447 | 0 004762514 | 0 122666604 | 0 49513336 | 0 | 0 136762554 |
| NLRP4 | 0 0006282 | 0 004751692 | 0 123519158 | -0 4961272 | 0 | 0 13768059 |
| PIK3IP1-DT | 0 018455769 | 0 004729505 | 0 125281392 | -0 47401272 | 0 | 0 139595467 |
| ITPKB-AS1 | 0 017694314 | 0 004717683 | 0 12622834 | -0 47670063 | 0 | 0 140634027 |
| B3GALT1 | 0 213911785 | 0 004711682 | 0 126711133 | -0 24170237 | 0 | 0 141155276 |
| CSF1R | 0 033494508 | 0 004692527 | 0 12826195 | -0 46160313 | 0 | 0 142832362 |
| DNAJC5B | 0 001389656 | 0 004678711 | 0 129389602 | -0 49539278 | 0 | 0 144071136 |
| CASC2 | 0 114798881 | 0 004669347 | 0 130158187 | 0 396380013 | 0 | 0 144909857 |
| MCOLN1 | 0 09051798 | 0 004646268 | 0 132067779 | -0 40494752 | 0 | 0 147018554 |
| LRP8 | 0 067522035 | 0 00464094 | 0 132511715 | 0 386039362 | 0 | 0 147495371 |
| CCN5 | 0 011022063 | 0 00463551 | 0 132965332 | -0 48405033 | 0 | 0 147982851 |
| GRAP | 0 002017856 | 0 004633732 | 0 133114064 | -0 49483765 | 0 | 0 148122939 |
| SLAMF6 | 0 003045821 | 0 004574744 | 0 138123753 | 0 494802724 | 0 | 0 153579182 |
| AQP4-AS1 | 0 035807428 | 0 00456831 | 0 138678893 | 0 458819105 | 0 | 0 15414203 |
| LINC02615 | 0 142572957 | 0 004564234 | 0 139031413 | -0 33411451 | 0 | 0 154461187 |
| GSTO2 | 0 046315509 | 0 004563802 | 0 139068826 | 0 439733875 | 0 | 0 15448459 |
| LINC00607 | 0 112248006 | 0 0045446 | 0 140739307 | -0 34824221 | 0 | 0 156266766 |
| NFYC-AS1 | 0 012716301 | 0 004537188 | 0 14138823 | -0 48256742 | 0 | 0 156950403 |
| P2RX5 | 0 003978603 | 0 004529976 | 0 142021853 | 0 493852515 | 0 | 0 157635249 |
| TRBC2 | 0 008128534 | 0 004518807 | 0 143007413 | -0 48779469 | 0 | 0 158710517 |
| COX4I2 | 0 003854867 | 0 004517859 | 0 143091248 | -0 49314832 | 0 | 0 15878491 |
| LAIR1 | 0 015200548 | 0 004507934 | 0 143971836 | -0 48056572 | 0 | 0 159705816 |
| SCAMP2 | 0 12742 | 0 004505881 | 0 144154498 | -0 36334832 | 0 | 0 159889672 |
| MUS81 | 0 142230302 | 0 004503991 | 0 144322879 | 0 332750735 | 0 | 0 160057647 |
| VPS26A | 0 278359445 | 0 004502604 | 0 144446504 | -0 24343815 | 0 | 0 160175952 |
| IGHG3 | 0 011060136 | 0 004501854 | 0 144513375 | -0 48543034 | 0 | 0 160231303 |
| COPE | 0 197968818 | 0 004501549 | 0 144540511 | -0 30100238 | 0 | 0 160242589 |
| LINC02422 | 0 035845501 | 0 004489291 | 0 145637176 | 0 453423234 | 0 | 0 161420516 |
| JAKMIP1 | 0 000437837 | 0 004474681 | 0 146952506 | -0 49642358 | 0 | 0 162802014 |
| APOBEC3B | 0 005168377 | 0 004453763 | 0 14885149 | 0 4869933 | 0 | 0 164867157 |
| ARSD | 0 06124003 | 0 00445154 | 0 149054353 | -0 42321362 | 0 | 0 165072499 |
| LINC00540 | 0 00383583 | 0 004443961 | 0 149747779 | -0 4920416 | 0 | 0 165801583 |
| LINC02361 | 0 003264739 | 0 004436543 | 0 150428755 | -0 49274245 | 0 | 0 166477542 |
| TCL1A | 0 000133255 | 0 004435527 | 0 150522256 | -0 49677153 | 0 | 0 166542012 |
| DBNDD2 | 0 027593231 | 0 004415875 | 0 152338707 | -0 46481818 | 0 | 0 168512323 |
| LTA | 0 000961337 | 0 004409004 | 0 152977742 | -0 49587071 | 0 | 0 169199398 |
| C10orf95 | 0 003759685 | 0 004389518 | 0 15480105 | 0 493495574 | 0 | 0 171135929 |
| LINC01580 | 0 007528888 | 0 004384847 | 0 155240514 | -0 48897107 | 0 | 0 171601691 |

| | | | | | | |
|---|---|---|---|---|---|---|
| VAV1 | 0 010955436 | 0 004378879 | 0 155803488 | -0 48561167 | 0 | 0 172192667 |
| EFCAB5 | 0 044164398 | 0 004378793 | 0 155811587 | 0 443004852 | 0 | 0 172192667 |
| SULT1B1 | 0 012782928 | 0 004375749 | 0 156099387 | 0 479259164 | 0 | 0 172490552 |
| LCN10 | 0 000799528 | 0 004365036 | 0 157115201 | 0 496351331 | 0 | 0 173572439 |
| KLRF1 | 0 00946108 | 0 004354017 | 0 158165242 | 0 483770155 | 0 | 0 174691624 |
| HLA-DMB | 0 023795473 | 0 004350551 | 0 158496682 | -0 47275944 | 0 | 0 174996337 |
| IL2 | 6 66E-05 | 0 004345811 | 0 158950766 | -0 49687338 | 0 | 0 175477189 |
| APLNR | 0 016837677 | 0 004344604 | 0 159066526 | -0 48234293 | 0 | 0 175584473 |
| TBCC | 0 033227999 | 0 004343776 | 0 159145956 | -0 46027346 | 0 | 0 175651633 |
| MBD2 | 0 347699454 | 0 004339784 | 0 159529462 | 0 210823537 | 0 | 0 176033795 |
| NAP1L1 | 0 65722145 | 0 004338078 | 0 15969358 | 0 042379536 | 5 64E-43 | 0 176194318 |
| RSU1 | 0 337238964 | 0 00433775 | 0 159725201 | -0 16504494 | 0 | 0 176208633 |
| CSF1 | 0 066008643 | 0 004333797 | 0 160106042 | -0 42019282 | 0 | 0 176608158 |
| GEMIN7 | 0 026898403 | 0 004327207 | 0 160742412 | -0 46844744 | 0 | 0 177268736 |
| CACNA1E | 0 05755649 | 0 004318202 | 0 161615103 | -0 4253906 | 0 | 0 178210353 |
| SEMA3G | 0 008204679 | 0 004307369 | 0 162669628 | -0 48656603 | 0 | 0 179352232 |
| DGKZ | 0 125069007 | 0 004302774 | 0 163118497 | 0 391252909 | 0 | 0 179826155 |
| LINC01678 | 0 001989302 | 0 004296094 | 0 163772758 | -0 49490914 | 0 | 0 180526369 |
| BTLA | 0 003607394 | 0 004283571 | 0 165004586 | 0 4939183 | 0 | 0 181862997 |
| CXCR6 | 0 003131484 | 0 004277141 | 0 165639752 | -0 49375533 | 0 | 0 182520482 |
| IL17C | 0 000276027 | 0 004271639 | 0 166184748 | -0 49669808 | 0 | 0 18309967 |
| CKB | 0 065618397 | 0 004261804 | 0 167162235 | 0 431645179 | 0 | 0 184112251 |
| LILRA1 | 0 001037483 | 0 00426138 | 0 167204425 | -0 49555465 | 0 | 0 184137259 |
| ISG20 | 0 047352992 | 0 004257047 | 0 167636601 | -0 44654241 | 0 | 0 184570183 |
| TSSK6 | 0 003654985 | 0 004246689 | 0 168672908 | 0 494241128 | 0 | 0 185667907 |
| LPAR5 | 0 005606213 | 0 004242552 | 0 16908819 | -0 49105478 | 0 | 0 186103354 |
| CD5L | 0 001085074 | 0 004234346 | 0 169914086 | -0 49556604 | 0 | 0 186947039 |
| SERINC1 | 0 309074642 | 0 004231775 | 0 170173507 | -0 18662889 | 0 | 0 187210668 |
| DNTTIP2 | 0 078696389 | 0 004228293 | 0 170525279 | -0 41111406 | 0 | 0 187575822 |
| CPNE9 | 0 008909025 | 0 004218184 | 0 171549691 | -0 48816019 | 0 | 0 188658743 |
| LAP3 | 0 157640251 | 0 004214882 | 0 171885374 | -0 33860708 | 0 | 0 18900591 |
| CXorf65 | 0 000152291 | 0 004201355 | 0 173265385 | -0 49672212 | 0 | 0 190501208 |
| JAML | 0 006910205 | 0 00418234 | 0 17521922 | -0 48933253 | 0 | 0 192604586 |
| UCK2 | 0 177057357 | 0 004180487 | 0 175410426 | 0 323480584 | 0 | 0 192769918 |
| DNASE2B | 0 000704346 | 0 004170375 | 0 176456992 | -0 4962168 | 0 | 0 193897509 |
| ARID5A | 0 074774895 | 0 004168769 | 0 176623687 | -0 40486052 | 0 | 0 194058114 |
| GIMAP4 | 0 009280235 | 0 004167967 | 0 176706929 | 0 486956997 | 0 | 0 194127003 |
| DKK3 | 0 213464431 | 0 004167057 | 0 176801461 | -0 26723794 | 0 | 0 194208277 |
| SLC25A39 | 0 106422874 | 0 004163388 | 0 177182819 | 0 394918702 | 0 | 0 194604561 |
| KCNMA1 | 0 284670004 | 0 004126898 | 0 181009749 | 0 213155719 | 0 | 0 198692313 |
| ITGA1 | 0 299899107 | 0 004123114 | 0 18141014 | -0 22801945 | 0 | 0 199062451 |
| GALNT7 | 0 322038415 | 0 004115015 | 0 182269227 | 0 219587866 | 0 | 0 19998191 |
| LINC00221 | 0 000218918 | 0 004090093 | 0 184931615 | -0 49670955 | 0 | 0 202761778 |
| ZBTB44-DT | 0 011869182 | 0 004084859 | 0 185494376 | -0 48204521 | 0 | 0 203355204 |

| | | | | | | |
|---|---|---|---|---|---|---|
| LY9 | 0 00131351 | 0 004082196 | 0 185781209 | 0 495586972 | 0 | 0 203622411 |
| CSF2RB | 0 002541357 | 0 004076092 | 0 186439878 | 0 494297022 | 0 | 0 204320636 |
| NAGPA | 0 033932345 | 0 00407043 | 0 187052354 | -0 4594256 | 0 | 0 204968083 |
| ZC3H12D | 0 009442044 | 0 004053289 | 0 188915744 | -0 48670911 | 0 | 0 20696195 |
| LIPT2 | 0 00331233 | 0 00403552 | 0 190861909 | 0 492757085 | 0 | 0 20892448 |
| ARRDC3 | 0 317450648 | 0 004034812 | 0 190939738 | -0 20962977 | 0 | 0 208985467 |
| NAAA | 0 079857608 | 0 004031873 | 0 191263166 | -0 40905411 | 0 | 0 209315219 |
| LINC01258 | 0 006025014 | 0 004024405 | 0 192086729 | 0 492406209 | 0 | 0 210167835 |
| ADGRG5 | 0 002408102 | 0 004023026 | 0 192239027 | 0 494542365 | 0 | 0 210310118 |
| LRRIQ1 | 0 104681045 | 0 004011729 | 0 193490512 | 0 402324662 | 0 | 0 211654745 |
| HS3ST1 | 0 019017342 | 0 004011354 | 0 193532176 | -0 474851 | 0 | 0 211675818 |
| C4orf50 | 0 001989302 | 0 00401083 | 0 193590405 | 0 495415217 | 0 | 0 211715001 |
| SH2D1B | 0 001646647 | 0 004006534 | 0 194068064 | 0 495510559 | 0 | 0 212212822 |
| TMEM167B | 0 066322743 | 0 003984313 | 0 196552602 | -0 42111994 | 0 | 0 214879927 |
| BRDT | 0 003616912 | 0 003983866 | 0 196602792 | 0 493064452 | 0 | 0 214909935 |
| AOC3 | 0 006596105 | 0 003969287 | 0 198245828 | -0 48911003 | 0 | 0 216680901 |
| FDX1 | 0 158763397 | 0 003958003 | 0 199524333 | -0 33605271 | 0 | 0 218027861 |
| SLC46A2 | 0 005463441 | 0 003946673 | 0 200814163 | -0 49061424 | 0 | 0 219354246 |
| NPFFR1 | 0 001941711 | 0 003946525 | 0 200830998 | -0 49460897 | 0 | 0 219354246 |
| GNGT2 | 0 000809046 | 0 00394589 | 0 200903456 | 0 496416689 | 0 | 0 219408027 |
| CAPN3 | 0 10187318 | 0 003941473 | 0 201408114 | -0 37209457 | 0 | 0 21993375 |
| CNKSR1 | 0 00751937 | 0 003931641 | 0 202534719 | -0 4884556 | 0 | 0 22111288 |
| CD8B | 0 005130304 | 0 003930558 | 0 202659085 | 0 492525797 | 0 | 0 221223097 |
| ACTG2 | 0 004425958 | 0 003919831 | 0 203894026 | -0 49143316 | 0 | 0 222545455 |
| LSM4 | 0 093925492 | 0 003915037 | 0 204447765 | 0 400605023 | 0 | 0 223124077 |
| CALHM6 | 0 017380214 | 0 00390933 | 0 205108284 | 0 474965377 | 0 | 0 223819086 |
| SNX20 | 0 001637129 | 0 003905755 | 0 205522916 | 0 495064529 | 0 | 0 224219759 |
| LAMC2 | 0 010374826 | 0 003905471 | 0 205555831 | -0 48535745 | 0 | 0 224229782 |
| MORN3 | 0 022757991 | 0 003902646 | 0 205883934 | 0 474482969 | 0 | 0 22456177 |
| MDK | 0 204526851 | 0 003895983 | 0 206659179 | -0 30390052 | 0 | 0 225381332 |
| LINC02099 | 0 003217148 | 0 003872825 | 0 209370216 | -0 4931496 | 0 | 0 228311625 |
| SET | 0 26451048 | 0 003865564 | 0 21022562 | 0 273373073 | 0 | 0 229191522 |
| TOMM40L | 0 012030991 | 0 003842945 | 0 212906183 | -0 48248064 | 0 | 0 231953358 |
| SLC37A2 | 0 010308199 | 0 003838761 | 0 213404731 | 0 485526963 | 0 | 0 232469707 |
| RHPN1 | 0 025385011 | 0 003814206 | 0 216347556 | 0 476536514 | 0 | 0 235593962 |
| EIF3K | 0 246559175 | 0 003798412 | 0 218255776 | -0 27245338 | 0 | 0 237589804 |
| SMPDL3A | 0 061792085 | 0 003780382 | 0 220448921 | -0 42648194 | 0 | 0 239866702 |
| RCC2 | 0 219280044 | 0 003773343 | 0 221309402 | 0 266794352 | 0 | 0 240775254 |
| RACGAP1 | 0 099931469 | 0 003745305 | 0 224760837 | 0 32267183 | 0 | 0 244417711 |
| PSMC4 | 0 135320097 | 0 003739087 | 0 225531432 | 0 368971104 | 0 | 0 245227481 |
| TBX21 | 0 001085074 | 0 003722705 | 0 227570774 | 0 496208828 | 0 | 0 247387992 |
| SLC10A6 | 0 033094744 | 0 003713701 | 0 228697171 | -0 45245341 | 0 | 0 248555292 |
| AMD1 | 0 266509299 | 0 003712051 | 0 228903987 | -0 21707498 | 0 | 0 248751458 |
| P2RY14 | 0 178427976 | 0 003708207 | 0 229386444 | -0 29585946 | 0 | 0 249247085 |

| | | | | | | |
|---|---|---|---|---|---|---|
| SRSF2 | 0 427709353 | 0 003690461 | 0 231622883 | 0 142735874 | 0 | 0 251590373 |
| SOX18 | 0 01287811 | 0 003668326 | 0 234434038 | -0 48190737 | 0 | 0 254585343 |
| SMC3 | 0 302659382 | 0 003667713 | 0 234512229 | 0 203564778 | 0 | 0 254640994 |
| USP30-AS1 | 0 00173231 | 0 003667378 | 0 234554977 | -0 49492179 | 0 | 0 254658149 |
| CLEC5A | 0 008928062 | 0 003648968 | 0 2369123 | -0 4867084 | 0 | 0 257187965 |
| KCNE1 | 0 010679408 | 0 003634595 | 0 23876429 | 0 48743955 | 0 | 0 259109157 |
| CRX | 0 003083893 | 0 003626375 | 0 239828119 | -0 49359317 | 0 | 0 260203872 |
| RNASET2 | 0 149502199 | 0 003624496 | 0 240071765 | -0 34202404 | 0 | 0 260438317 |
| KITLG | 0 134320687 | 0 003608948 | 0 242094426 | 0 346294252 | 0 | 0 262572286 |
| TNFSF18 | 0 001199292 | 0 003606564 | 0 242405575 | -0 49543532 | 0 | 0 262849419 |
| TSPYL1 | 0 083388856 | 0 003592558 | 0 244239508 | -0 40159675 | 0 | 0 264746886 |
| FCRL3 | 0 001237365 | 0 003586047 | 0 24509539 | 0 496171381 | 0 | 0 265644161 |
| GLS | 0 645780587 | 0 003584433 | 0 245307861 | -0 06595159 | 1 33E-101 | 0 265832413 |
| CHRNA6 | 0 001960747 | 0 0035843 | 0 245325341 | -0 49441771 | 0 | 0 265832413 |
| RPL31 | 0 408235137 | 0 003579453 | 0 245964341 | 0 146332635 | 0 | 0 266463719 |
| TMEM45B | 0 002931602 | 0 003577962 | 0 246161077 | -0 49315659 | 0 | 0 266615722 |
| MACC1 | 0 011992918 | 0 003560331 | 0 24849618 | 0 485669844 | 0 | 0 269052349 |
| LINC02312 | 0 0002094 | 0 003554151 | 0 249318466 | -0 49663944 | 0 | 0 269911732 |
| LPXN | 0 125325998 | 0 003534441 | 0 251953378 | 0 361005869 | 0 | 0 272639348 |
| ADAM17 | 0 512211837 | 0 003532602 | 0 252200176 | -0 09180131 | 2 25E-195 | 0 272875162 |
| EXD1 | 0 00373113 | 0 003525671 | 0 253131963 | 0 493314149 | 0 | 0 273851981 |
| SORBS1 | 0 452923036 | 0 003524362 | 0 253308145 | -0 07072433 | 1 39E-116 | 0 274011215 |
| MS4A2 | 0 001237365 | 0 003512618 | 0 254893295 | -0 49557712 | 0 | 0 275662811 |
| ERMN | 0 000656755 | 0 003492286 | 0 257653693 | 0 496435213 | 0 | 0 278520636 |
| MS4A7 | 0 025385011 | 0 00345492 | 0 262780902 | 0 467719731 | 0 | 0 283803374 |
| CCSAP | 0 063743313 | 0 003453846 | 0 262929216 | 0 426724802 | 0 | 0 283931104 |
| TRBC1 | 0 003693057 | 0 003452844 | 0 263067741 | 0 493028883 | 0 | 0 284015783 |
| PARP10 | 0 082218119 | 0 003452198 | 0 263157049 | -0 4117156 | 0 | 0 284079748 |
| TTLL10 | 0 001503874 | 0 003445174 | 0 264129645 | -0 49516424 | 0 | 0 285064544 |
| CYLD | 0 227913042 | 0 003441994 | 0 264570809 | -0 27293435 | 0 | 0 285508068 |
| PRR13 | 0 116883364 | 0 003440447 | 0 264785513 | -0 38022169 | 0 | 0 285707136 |
| XIRP1 | 0 001846529 | 0 003435897 | 0 265417988 | -0 49508864 | 0 | 0 286356889 |
| UNC13A | 0 027431421 | 0 003428401 | 0 266462167 | -0 46612161 | 0 | 0 287417815 |
| FAM168A | 0 50352173 | 0 00342593 | 0 266806903 | 0 100417531 | 1 50E-233 | 0 287756818 |
| VSTM1 | 0 000837601 | 0 003424085 | 0 267064529 | 0 496164656 | 0 | 0 287992231 |
| FCGR2C | 0 005368259 | 0 003421363 | 0 267444957 | -0 49001518 | 0 | 0 288346246 |
| LINC02520 | 0 000742419 | 0 003414798 | 0 268364158 | 0 496293944 | 0 | 0 289304277 |
| EGFL8 | 0 109468695 | 0 003413063 | 0 268607334 | -0 36348997 | 0 | 0 289533398 |
| SYNGR3 | 0 001560983 | 0 003406266 | 0 269561845 | -0 49491168 | 0 | 0 290529133 |
| GCSAM | 0 004359331 | 0 003405632 | 0 269650996 | -0 49226064 | 0 | 0 290592076 |
| GPR141 | 0 00751937 | 0 003404406 | 0 269823414 | -0 48816145 | 0 | 0 290744728 |
| USP2 | 0 011126763 | 0 003375363 | 0 273930657 | -0 48436986 | 0 | 0 295069498 |
| C1orf141 | 0 002027374 | 0 00336622 | 0 275232435 | -0 49469728 | 0 | 0 296437943 |
| TLR3 | 0 013887038 | 0 003364066 | 0 275539679 | -0 48187391 | 0 | 0 296735039 |

| | | | | | | |
|---|---|---|---|---|---|---|
| HP | 0 001075555 | 0 003359444 | 0 276199846 | -0 49572892 | 0 | 0 29737821 |
| TPX2 | 0 100007615 | 0 003329279 | 0 28053497 | 0 317541642 | 0 | 0 30190815 |
| NEURL1 | 0 106099256 | 0 003302787 | 0 284380166 | 0 35761393 | 0 | 0 305906952 |
| SMG1 | 0 515781158 | 0 003300489 | 0 284715358 | 0 101810111 | 4 79E-240 | 0 306232658 |
| IFIT1 | 0 079457844 | 0 003292799 | 0 285839041 | -0 414602 | 0 | 0 307371295 |
| ELAPOR2 | 0 075041404 | 0 003290334 | 0 286199984 | -0 41732575 | 0 | 0 307724411 |
| CLEC4G | 0 0002094 | 0 003281886 | 0 287438964 | -0 49667878 | 0 | 0 30898626 |
| MGAM2 | 0 003350403 | 0 003257138 | 0 291089224 | -0 49279966 | 0 | 0 312803408 |
| GPR183 | 0 025166092 | 0 003255615 | 0 291314859 | -0 47138257 | 0 | 0 31301028 |
| KLF16 | 0 022291599 | 0 003236781 | 0 294115126 | -0 47052914 | 0 | 0 315947248 |
| CDKN3 | 0 042717633 | 0 003222488 | 0 296252272 | 0 408750823 | 0 | 0 318170698 |
| ENO1-AS1 | 0 002360511 | 0 003220925 | 0 296486573 | -0 49428031 | 0 | 0 318349973 |
| HSPA2 | 0 047172146 | 0 003202375 | 0 299277223 | -0 43879629 | 0 | 0 321309902 |
| CCL8 | 0 002712684 | 0 003201851 | 0 299356255 | -0 49383558 | 0 | 0 321358242 |
| MRPS24 | 0 101692334 | 0 003188484 | 0 301378275 | 0 400704196 | 0 | 0 323455388 |
| LINC01215 | 0 000409282 | 0 003175591 | 0 303337293 | 0 49674235 | 0 | 0 325520943 |
| FABP4 | 0 019978679 | 0 003148168 | 0 307531909 | 0 472040352 | 0 | 0 32983504 |
| SOCS2 | 0 060564238 | 0 00313598 | 0 309408478 | -0 42124512 | 0 | 0 331772394 |
| NTM | 0 317060402 | 0 003127871 | 0 310661209 | -0 21218793 | 0 | 0 333077879 |
| HCK | 0 032228589 | 0 003117984 | 0 312193079 | -0 45817554 | 0 | 0 334644351 |
| RAB8A | 0 085730331 | 0 003115494 | 0 312579654 | 0 424358203 | 0 | 0 334993996 |
| BID | 0 069349527 | 0 003115426 | 0 312590165 | 0 41015834 | 0 | 0 334993996 |
| RIPK3 | 0 001779901 | 0 00311163 | 0 313180178 | -0 49490341 | 0 | 0 33558824 |
| GPRC5A | 0 04173726 | 0 003103926 | 0 314379876 | -0 4504631 | 0 | 0 336835582 |
| NXT2 | 0 015524167 | 0 003064542 | 0 320559502 | 0 485473804 | 0 | 0 343339827 |
| LINC02889 | 0 00846167 | 0 003061975 | 0 320965034 | -0 48658303 | 0 | 0 343735219 |
| CD300LB | 0 000675791 | 0 003060073 | 0 321265714 | -0 49607232 | 0 | 0 344018244 |
| PHC2 | 0 439749862 | 0 003056678 | 0 321802956 | 0 137846467 | 0 | 0 34455449 |
| ABTB1 | 0 081761246 | 0 003051848 | 0 322568169 | -0 40379424 | 0 | 0 345295559 |
| PRSS21 | 0 002550875 | 0 003041242 | 0 324252577 | -0 49468476 | 0 | 0 347020029 |
| SSB | 0 25436409 | 0 00303696 | 0 324934373 | -0 26068158 | 0 | 0 347670949 |
| SEC62 | 0 47503379 | 0 003030164 | 0 326018135 | -0 11337002 | 1 66E-297 | 0 348791053 |
| LAGE3 | 0 047657574 | 0 003008155 | 0 329544354 | 0 456656923 | 0 | 0 352404 |
| MYO1G | 0 006320078 | 0 002994686 | 0 331714465 | -0 48968803 | 0 | 0 354644382 |
| RCL1 | 0 206363861 | 0 002993189 | 0 331956191 | 0 285668644 | 0 | 0 35486267 |
| CLDN7 | 0 007138642 | 0 00299244 | 0 332077343 | -0 48991718 | 0 | 0 354952029 |
| FAM3B | 0 002531838 | 0 002988443 | 0 332723515 | 0 495536497 | 0 | 0 35560249 |
| XRRA1 | 0 40433268 | 0 002977011 | 0 334576182 | 0 145262586 | 0 | 0 357501687 |
| AC023590.1 | 0 031095924 | 0 002970601 | 0 335617999 | -0 46204587 | 0 | 0 358533811 |
| SHCBP1 | 0 065152005 | 0 0029581 | 0 337655699 | 0 378313642 | 0 | 0 360629109 |
| SAMD12 | 0 134330205 | 0 002939343 | 0 340727996 | -0 32875834 | 0 | 0 363745997 |
| CD7 | 0 004368849 | 0 002933934 | 0 341617275 | 0 492047393 | 0 | 0 364654157 |
| TRIB1 | 0 055148389 | 0 002922454 | 0 343509481 | -0 42871315 | 0 | 0 366591145 |
| CEBPA | 0 00804287 | 0 002894956 | 0 348069343 | -0 48827735 | 0 | 0 37133556 |

| | | | | | | |
|---|---|---|---|---|---|---|
| DDX28 | 0 009927471 | 0 002894934 | 0 348073057 | -0 48563294 | 0 | 0 37133556 |
| CSTA | 0 004978013 | 0 002891897 | 0 348579052 | 0 492562333 | 0 | 0 371833394 |
| GIMAP6 | 0 003911976 | 0 002888418 | 0 349159153 | 0 493217897 | 0 | 0 372410157 |
| PIK3AP1 | 0 016856713 | 0 002886936 | 0 349406494 | -0 48035465 | 0 | 0 372631912 |
| ZNF341-AS1 | 0 004435476 | 0 002884484 | 0 349815996 | -0 49201038 | 0 | 0 373026536 |
| LINC00639 | 0 010498563 | 0 00286325 | 0 353374445 | -0 4847224 | 0 | 0 376651083 |
| MEIKIN | 0 005225486 | 0 002862061 | 0 35357443 | -0 4912689 | 0 | 0 376821739 |
| UBE2B | 0 459319259 | 0 002861738 | 0 353628733 | -0 11921019 | 0 | 0 376837114 |
| TRIM44 | 0 501456283 | 0 002857351 | 0 354367056 | 0 1001552 | 2 46E-232 | 0 377581314 |
| LINC00881 | 0 002179665 | 0 002843088 | 0 356774446 | 0 494728974 | 0 | 0 380060705 |
| PECAM1 | 0 087129504 | 0 002838503 | 0 357550497 | 0 383840773 | 0 | 0 380844477 |
| ABI3 | 0 005511032 | 0 002828008 | 0 35933084 | 0 491114808 | 0 | 0 382654546 |
| HDX | 0 153062001 | 0 002826854 | 0 359527068 | 0 349217047 | 0 | 0 382820371 |
| HAUS6 | 0 175296492 | 0 002824055 | 0 360003018 | 0 288094312 | 0 | 0 38328397 |
| SAMD10 | 0 006091641 | 0 002819328 | 0 360807527 | -0 48973606 | 0 | 0 384053968 |
| SRSF7 | 0 407397537 | 0 00281373 | 0 361761942 | 0 187381778 | 0 | 0 38489646 |
| PTCSC1 | 0 00210352 | 0 002805278 | 0 363205915 | -0 4943204 | 0 | 0 386345778 |
| CEBPE | 0 000133255 | 0 002798475 | 0 364370716 | 0 496905727 | 0 | 0 387497554 |
| CXCR2 | 0 000571091 | 0 002774297 | 0 368529404 | -0 49625819 | 0 | 0 391611702 |
| LMOD1 | 0 049446993 | 0 002773539 | 0 368660349 | -0 42796772 | 0 | 0 391706803 |
| TCEAL9 | 0 223220575 | 0 002769089 | 0 369429119 | -0 27482444 | 0 | 0 392479502 |
| TOR3A | 0 0698064 | 0 002747894 | 0 37310447 | 0 429970807 | 0 | 0 39629507 |
| NAPSB | 0 004378367 | 0 002715932 | 0 378689447 | -0 4914801 | 0 | 0 402091604 |
| FCAR | 0 001770383 | 0 002697695 | 0 381899276 | -0 49472933 | 0 | 0 405454236 |
| IL23A | 0 00431174 | 0 002695717 | 0 382248332 | 0 491635807 | 0 | 0 405733656 |
| ME2 | 0 229778607 | 0 002692077 | 0 382891285 | 0 298729605 | 0 | 0 406370468 |
| S100A12 | 0 000466391 | 0 002687923 | 0 383625884 | -0 49645882 | 0 | 0 407104391 |
| ZNF66 | 0 131417639 | 0 00268729 | 0 383737896 | 0 294204146 | 0 | 0 407177534 |
| CSMD2 | 0 075050922 | 0 00268285 | 0 384524107 | -0 38873236 | 0 | 0 407965961 |
| NPM1 | 0 429394072 | 0 00265632 | 0 3892428 | 0 143903189 | 0 | 0 412879612 |
| LGALS9C | 1 90E-05 | 0 002652699 | 0 38988959 | 0 496921279 | 0 | 0 413380077 |
| SLC41A2 | 0 25488759 | 0 002651198 | 0 390157789 | -0 2599144 | 0 | 0 413571633 |
| SNHG25 | 0 407492719 | 0 002614975 | 0 396666268 | -0 16399027 | 0 | 0 420423535 |
| CCDC159 | 0 03658792 | 0 002596152 | 0 400073887 | -0 4509533 | 0 | 0 423797582 |
| STEAP3 | 0 040328568 | 0 002589175 | 0 401341624 | 0 451365858 | 0 | 0 425045203 |
| NCOA4 | 0 13156993 | 0 002586931 | 0 401749837 | -0 35875732 | 0 | 0 425429848 |
| NDST3 | 0 135358169 | 0 002581373 | 0 402761846 | 0 33068273 | 0 | 0 426453719 |
| IL2RA | 0 010555672 | 0 002578049 | 0 403367895 | -0 48569329 | 0 | 0 427047569 |
| UPK3A | 0 00079001 | 0 002573575 | 0 404184432 | -0 49614421 | 0 | 0 427864107 |
| SEMA3D | 0 095619729 | 0 002549221 | 0 408646693 | -0 37380565 | 0 | 0 432539343 |
| ACY3 | 0 000761455 | 0 00254834 | 0 408808662 | 0 496457675 | 0 | 0 43266232 |
| ZNF536 | 0 216710133 | 0 002543902 | 0 409625263 | -0 27126204 | 0 | 0 433467488 |
| GJA4 | 0 005320668 | 0 002543707 | 0 409661179 | -0 49167173 | 0 | 0 433467488 |
| TNFAIP8L2 | 0 001532428 | 0 002541323 | 0 410100232 | -0 49515284 | 0 | 0 433786343 |

| | | | | | | |
|---|---|---|---|---|---|---|
| LYPD3 | 0 004721022 | 0 002537931 | 0 410725371 | -0 49093403 | 0 | 0 434350353 |
| CD19 | 0 000875673 | 0 002530329 | 0 41212831 | 0 496243589 | 0 | 0 435785222 |
| PIP4K2A | 0 376615713 | 0 002527184 | 0 412709686 | -0 16356076 | 0 | 0 436351145 |
| SIGLEC11 | 0 00141821 | 0 002503408 | 0 41711988 | 0 495663481 | 0 | 0 440865996 |
| CD79A | 0 000828082 | 0 002498393 | 0 418053671 | 0 495876659 | 0 | 0 44175413 |
| TDRD9 | 0 018598542 | 0 00248008 | 0 421474063 | -0 4690632 | 0 | 0 445268854 |
| CLEC12A | 0 006339114 | 0 002470283 | 0 423310411 | -0 49086405 | 0 | 0 447158885 |
| NXF3 | 0 000942301 | 0 002467707 | 0 423794141 | -0 4958932 | 0 | 0 447569807 |
| CATSPERB | 0 019379033 | 0 002459747 | 0 425290709 | -0 47431029 | 0 | 0 449100145 |
| NTNG2 | 0 032418953 | 0 002458195 | 0 425582871 | -0 46472996 | 0 | 0 449308249 |
| DCLRE1A | 0 026536712 | 0 002442817 | 0 428484036 | 0 465450696 | 0 | 0 452226744 |
| NCKAP1L | 0 01319221 | 0 002426967 | 0 431486153 | -0 48356608 | 0 | 0 455337146 |
| MTF1 | 0 097989758 | 0 00241732 | 0 433319234 | 0 403476976 | 0 | 0 45716945 |
| MIR3945HG | 0 001560983 | 0 002415031 | 0 433754951 | 0 495440594 | 0 | 0 457578063 |
| COL5A3 | 0 175106128 | 0 002410179 | 0 434679156 | -0 32393354 | 0 | 0 458501844 |
| INHBB | 0 01319221 | 0 00240814 | 0 435067966 | -0 47979679 | 0 | 0 458758343 |
| SURF2 | 0 027831185 | 0 002398776 | 0 436855798 | -0 46627123 | 0 | 0 460540751 |
| HBA2 | 0 017304068 | 0 002393468 | 0 437871131 | -0 47797226 | 0 | 0 461559641 |
| FDXACB1 | 0 007157678 | 0 002378526 | 0 440736551 | -0 48863933 | 0 | 0 464424666 |
| LCP2 | 0 021948944 | 0 002375931 | 0 441235306 | -0 47498066 | 0 | 0 464898388 |
| FAM111A | 0 175458301 | 0 002367986 | 0 442764356 | 0 295352652 | 0 | 0 466405436 |
| VPS18 | 0 037244675 | 0 002361217 | 0 444069307 | -0 45597643 | 0 | 0 467623685 |
| CEACAM4 | 0 000590128 | 0 002354544 | 0 44535803 | -0 49624224 | 0 | 0 468876268 |
| DNASE1L3 | 0 001646647 | 0 002349796 | 0 446276311 | -0 49504495 | 0 | 0 469790703 |
| IRS2 | 0 213426358 | 0 002336923 | 0 448771149 | -0 25445631 | 0 | 0 472311767 |
| PDZD7 | 0 013687156 | 0 002332894 | 0 449553737 | -0 48051181 | 0 | 0 473030043 |
| FFAR4 | 0 001656165 | 0 002320621 | 0 451942032 | 0 495353696 | 0 | 0 475437181 |
| TRIM22 | 0 310940207 | 0 002314876 | 0 453062633 | -0 21031832 | 0 | 0 476509947 |
| TMEM176A | 0 038463003 | 0 002311446 | 0 453732331 | -0 44970067 | 0 | 0 477161198 |
| PAPLN | 0 150358836 | 0 002303257 | 0 455333347 | -0 34138975 | 0 | 0 478738332 |
| SOX9 | 0 123774533 | 0 002301962 | 0 45558688 | -0 37301847 | 0 | 0 47895161 |
| AK1 | 0 082960538 | 0 002297889 | 0 456384829 | -0 39660614 | 0 | 0 479737111 |
| TMEM273 | 0 004178485 | 0 002278954 | 0 460103916 | -0 49222131 | 0 | 0 483485156 |
| ACADSB | 0 149778226 | 0 002277939 | 0 460303751 | -0 34867503 | 0 | 0 483641366 |
| VSIG8 | 0 006691287 | 0 002277057 | 0 460477466 | -0 48938249 | 0 | 0 483770101 |
| TAF5 | 0 045525499 | 0 002269716 | 0 4619246 | 0 431278644 | 0 | 0 48523649 |
| CYTIP | 0 010746036 | 0 002266975 | 0 462465553 | 0 485850014 | 0 | 0 485750748 |
| ICOS | 0 002960157 | 0 002260123 | 0 46381963 | 0 494175455 | 0 | 0 48711886 |
| TPSB2 | 0 002217738 | 0 002255998 | 0 464635645 | -0 49461861 | 0 | 0 487867427 |
| FLT3LG | 0 028706859 | 0 002250628 | 0 465699378 | -0 46562838 | 0 | 0 488875708 |
| CXorf58 | 0 003597876 | 0 002235956 | 0 468612357 | -0 49263411 | 0 | 0 491824387 |
| CYTH4 | 0 009127944 | 0 002235468 | 0 468709335 | -0 48733878 | 0 | 0 491871539 |
| SLC22A18 | 0 036188156 | 0 002219302 | 0 471930977 | -0 45303258 | 0 | 0 495087448 |
| DENND4C | 0 513058956 | 0 002215117 | 0 47276704 | -0 0976259 | 8 46E-221 | 0 495900663 |

| | | | | | | |
|---|---|---|---|---|---|---|
| MBNL1-AS1 | 0 011593155 | 0 002214897 | 0 472811109 | -0 48438496 | 0 | 0 495900663 |
| IATPR | 0 002636538 | 0 002213045 | 0 473181181 | 0 493616912 | 0 | 0 496233732 |
| SPN | 0 00310293 | 0 002210571 | 0 473676088 | 0 493967435 | 0 | 0 496697629 |
| MS4A1 | 0 000732901 | 0 002201372 | 0 475518387 | -0 49621829 | 0 | 0 498574144 |
| TREML1 | 0 000961337 | 0 002198885 | 0 476017248 | -0 49559097 | 0 | 0 499041823 |
| KLF11 | 0 047562392 | 0 002192448 | 0 477309573 | 0 43937764 | 0 | 0 500285655 |
| KHDRBS2 | 0 079486399 | 0 002180489 | 0 479715587 | -0 40608539 | 0 | 0 502695974 |
| NFKBIE | 0 016209476 | 0 002160453 | 0 4837612 | -0 4767296 | 0 | 0 506766804 |
| PARM1 | 0 049037711 | 0 002160143 | 0 483823996 | 0 450662399 | 0 | 0 506776409 |
| GFPT2 | 0 20989511 | 0 002151394 | 0 485596538 | -0 30440094 | 0 | 0 508576669 |
| SLCO1A2 | 0 025898993 | 0 002147289 | 0 486429317 | -0 47346183 | 0 | 0 509392403 |
| NYAP2 | 0 005910796 | 0 002136241 | 0 488674778 | -0 48872301 | 0 | 0 511687164 |
| TTC16 | 0 001694238 | 0 002131787 | 0 489581664 | 0 495721486 | 0 | 0 51257996 |
| MAD2L1 | 0 027802631 | 0 002125321 | 0 490899716 | 0 442840471 | 0 | 0 513902993 |
| LPO | 0 001951229 | 0 002094456 | 0 497217451 | -0 49470497 | 0 | 0 520286226 |
| TRDC | 0 001085074 | 0 002085458 | 0 499067342 | -0 49571634 | 0 | 0 522164125 |
| SNHG9 | 0 124659725 | 0 002082726 | 0 499629869 | -0 36688938 | 0 | 0 522694814 |
| LINC01806 | 0 000809046 | 0 002069308 | 0 502397013 | 0 496298588 | 0 | 0 525531522 |
| NRBF2 | 0 091974263 | 0 002046711 | 0 507075108 | -0 40228081 | 0 | 0 530307633 |
| CX3CR1 | 0 006301041 | 0 002044454 | 0 507543737 | -0 4888268 | 0 | 0 530738997 |
| TINAGL1 | 0 013972702 | 0 002036744 | 0 509145964 | -0 48194936 | 0 | 0 532355539 |
| INKA2 | 0 174544555 | 0 002034747 | 0 509561401 | -0 31335936 | 0 | 0 53267204 |
| SIGLEC6 | 0 0004188 | 0 002025666 | 0 511452683 | -0 49646899 | 0 | 0 53453084 |
| KDM2A | 0 607860121 | 0 002017205 | 0 513218115 | 0 059722622 | 1 25E-83 | 0 536316619 |
| ZSCAN16 | 0 01235461 | 0 002016199 | 0 51342831 | -0 4822108 | 0 | 0 536476949 |
| XCR1 | 0 000732901 | 0 002015774 | 0 513517025 | -0 49611647 | 0 | 0 536510324 |
| CPA3 | 0 002170147 | 0 002011742 | 0 514359827 | -0 49480876 | 0 | 0 537331458 |
| IL1RL2 | 0 023671737 | 0 001994434 | 0 517985191 | 0 458509485 | 0 | 0 540939336 |
| HEG1 | 0 15722145 | 0 001991139 | 0 518677002 | 0 341970065 | 0 | 0 541601953 |
| CHRAC1 | 0 050541585 | 0 001988457 | 0 519240395 | 0 451920329 | 0 | 0 542070452 |
| GP6 | 0 022377263 | 0 001982677 | 0 520455582 | 0 466567176 | 0 | 0 543219047 |
| MDFIC | 0 542898479 | 0 001972605 | 0 522576526 | 0 095456477 | 3 83E-211 | 0 545312298 |
| HAVCR1 | 0 00173231 | 0 00196517 | 0 524145161 | 0 495373569 | 0 | 0 546856714 |
| MAGIX | 0 007072015 | 0 001964199 | 0 524350236 | -0 48887087 | 0 | 0 546981978 |
| ADAMTSL1 | 0 395252327 | 0 001950178 | 0 527315253 | 0 142697444 | 0 | 0 549953553 |
| GSDMA | 0 001580019 | 0 001924712 | 0 532722648 | -0 49511467 | 0 | 0 555225435 |
| PAK1 | 0 30645714 | 0 001900947 | 0 537794084 | 0 176174083 | 0 | 0 560263929 |
| IDH1 | 0 114361044 | 0 001899538 | 0 538095442 | 0 388831539 | 0 | 0 560516086 |
| KCNJ5 | 0 003198112 | 0 001899158 | 0 538176835 | 0 493341653 | 0 | 0 560539082 |
| NPL | 0 114370562 | 0 001888676 | 0 540422006 | -0 37172871 | 0 | 0 562691489 |
| ADAM19 | 0 265671699 | 0 001880108 | 0 542260798 | 0 220006515 | 0 | 0 564543852 |
| DOCK5 | 0 497239725 | 0 001879649 | 0 542359498 | -0 10462273 | 1 91E-253 | 0 564584408 |
| RRAS2 | 0 273781196 | 0 001855468 | 0 547566237 | 0 218840679 | 0 | 0 569939812 |
| RCSD1 | 0 071919438 | 0 001848193 | 0 549137438 | 0 407267625 | 0 | 0 571451249 |

| | | | | | | |
|---|---|---|---|---|---|---|
| GDAP2 | 0 210751747 | 0 00184483 | 0 549864578 | 0 302553938 | 0 | 0 572144931 |
| GNAI3 | 0 299423198 | 0 001836421 | 0 551684686 | 0 22672021 | 0 | 0 57397559 |
| MZB1 | 0 00231292 | 0 001820836 | 0 555066084 | 0 494638262 | 0 | 0 577430041 |
| ASCL2 | 0 005787059 | 0 001804784 | 0 558559245 | -0 48992638 | 0 | 0 58093604 |
| F2RL1 | 0 00877577 | 0 00179944 | 0 55972436 | 0 485995699 | 0 | 0 582083768 |
| ZNF600 | 0 069263863 | 0 001781647 | 0 563612767 | 0 418745521 | 0 | 0 585998531 |
| S100P | 0 001570501 | 0 001754001 | 0 569680084 | -0 49510179 | 0 | 0 592046277 |
| CNDP2 | 0 089404352 | 0 001752076 | 0 57010362 | -0 40292873 | 0 | 0 59242129 |
| SASH3 | 0 004987531 | 0 001750253 | 0 57050496 | -0 49184982 | 0 | 0 592773158 |
| LILRA5 | 0 00079001 | 0 001726212 | 0 575809977 | 0 496190323 | 0 | 0 598153708 |
| DENND1C | 0 003474139 | 0 001721033 | 0 576955839 | -0 4929887 | 0 | 0 599212296 |
| TMEM232 | 0 162094763 | 0 001711375 | 0 579095414 | -0 30439039 | 0 | 0 601368314 |
| UVRAG | 0 545877672 | 0 00170312 | 0 58092728 | 0 093897033 | 2 44E-204 | 0 603204351 |
| CD300C | 0 000513982 | 0 001672359 | 0 587777073 | -0 49631846 | 0 | 0 610182724 |
| CNFN | 0 00531115 | 0 001668199 | 0 588706248 | 0 49304322 | 0 | 0 611013074 |
| MICALL1 | 0 085720812 | 0 001661063 | 0 59030185 | 0 387982925 | 0 | 0 612601854 |
| DHRS1 | 0 060878339 | 0 001659857 | 0 590571661 | -0 43117881 | 0 | 0 61281456 |
| GIN1 | 0 087500714 | 0 001649205 | 0 592957488 | -0 40892941 | 0 | 0 61522269 |
| CREG1 | 0 079400735 | 0 001646892 | 0 593476091 | -0 41258092 | 0 | 0 615693167 |
| IFNAR2 | 0 276760389 | 0 001637754 | 0 59552721 | -0 23470327 | 0 | 0 617617662 |
| DHRS13 | 0 01729455 | 0 001633641 | 0 596451349 | 0 483858683 | 0 | 0 618508203 |
| PAFAH2 | 0 054910434 | 0 001631358 | 0 596964653 | -0 43826725 | 0 | 0 618972566 |
| PWWP2B | 0 014115475 | 0 001625566 | 0 598267751 | 0 484692789 | 0 | 0 62025565 |
| IL10 | 0 001865565 | 0 001625031 | 0 598388191 | -0 49495037 | 0 | 0 620312463 |
| PKIB | 0 09162209 | 0 001602524 | 0 603464805 | -0 39996908 | 0 | 0 625369273 |
| SATB2 | 0 296758105 | 0 001595123 | 0 605138211 | 0 178015994 | 0 | 0 62703466 |
| C5orf64 | 0 003645466 | 0 001572589 | 0 610246374 | -0 49215797 | 0 | 0 63225833 |
| FCRL5 | 0 001199292 | 0 00156746 | 0 611411718 | -0 49568417 | 0 | 0 633396265 |
| FCGR3B | 0 000371209 | 0 001566335 | 0 611667549 | 0 496579318 | 0 | 0 633591838 |
| SIRPG | 0 00383583 | 0 001563641 | 0 612279998 | 0 491597607 | 0 | 0 634156727 |
| ZNF165 | 0 003712094 | 0 001559367 | 0 613252574 | 0 494043318 | 0 | 0 635024855 |
| CLMN | 0 188241229 | 0 001549159 | 0 615577877 | -0 29145551 | 0 | 0 637362873 |
| DOK2 | 0 003892939 | 0 001543398 | 0 616892005 | 0 492407568 | 0 | 0 638583574 |
| ARHGEF15 | 0 009860844 | 0 001542437 | 0 617111275 | -0 48724257 | 0 | 0 638740586 |
| CATSPERD | 0 009546744 | 0 001540638 | 0 617521812 | 0 485202605 | 0 | 0 639095512 |
| RAB39B | 0 005558623 | 0 00152218 | 0 621742295 | -0 49011542 | 0 | 0 643322532 |
| SPAG1 | 0 147084579 | 0 001511372 | 0 624219094 | 0 329314801 | 0 | 0 645673209 |
| PIF1 | 0 036702138 | 0 001502746 | 0 626199048 | 0 42644997 | 0 | 0 64750859 |
| MTHFD2L | 0 210294874 | 0 001494722 | 0 628043261 | 0 277234855 | 0 | 0 649273473 |
| LILRA4 | 0 000142773 | 0 001491711 | 0 62873589 | -0 49680224 | 0 | 0 649847332 |
| C7 | 0 080485808 | 0 001485792 | 0 630098508 | -0 39518966 | 0 | 0 651184481 |
| CCL17 | 0 001541947 | 0 001470559 | 0 633610873 | -0 49538458 | 0 | 0 65431349 |
| CCL4L2 | 0 012944737 | 0 001461204 | 0 63577197 | -0 48171736 | 0 | 0 656473461 |
| DPEP2 | 0 003531248 | 0 001451027 | 0 638126525 | -0 49304744 | 0 | 0 658832696 |

| | | | | | | |
|---|---|---|---|---|---|---|
| TNFRSF1B | 0 040928214 | 0 001450352 | 0 638282795 | 0 450467865 | 0 | 0 658922046 |
| ZMYM6 | 0 255077954 | 0 001447572 | 0 638926567 | -0 2580613 | 0 | 0 659514588 |
| SLC26A2 | 0 130732329 | 0 001440062 | 0 640667375 | -0 35055477 | 0 | 0 661239263 |
| LINC00880 | 0 003978603 | 0 001436699 | 0 64144749 | -0 49225 | 0 | 0 661972128 |
| PPP1CA | 0 100236051 | 0 001425598 | 0 644025636 | 0 392717767 | 0 | 0 664487637 |
| MIR4422HG | 0 00604405 | 0 001419528 | 0 645437246 | 0 489897991 | 0 | 0 665871394 |
| CEBPD | 0 282376121 | 0 001405468 | 0 648711531 | -0 2169393 | 0 | 0 669176288 |
| LRP2 | 0 004892349 | 0 001373908 | 0 656085991 | -0 49177073 | 0 | 0 676577591 |
| SERBP1 | 0 346880889 | 0 001373843 | 0 656101274 | 0 208418046 | 0 | 0 676577591 |
| FAM30A | 0 0005235 | 0 001369155 | 0 657199566 | -0 49639752 | 0 | 0 677636214 |
| TMEM143 | 0 053901506 | 0 001364932 | 0 65818961 | 0 437986217 | 0 | 0 678583004 |
| CD209 | 0 002331956 | 0 001352183 | 0 661182031 | 0 49379554 | 0 | 0 681296499 |
| TFF3 | 0 010431935 | 0 001350607 | 0 661552354 | -0 48473906 | 0 | 0 681603767 |
| FSIP1 | 0 067731435 | 0 001349514 | 0 661809226 | -0 42338002 | 0 | 0 68179409 |
| LINC00544 | 0 000980373 | 0 001346144 | 0 662601487 | -0 49587291 | 0 | 0 682461477 |
| CD1C | 0 000951819 | 0 001329891 | 0 6664274 | -0 49568465 | 0 | 0 6861777 |
| CCR2 | 0 00120881 | 0 001329572 | 0 666502653 | -0 49569879 | 0 | 0 68618042 |
| TTC24 | 0 000466391 | 0 001311723 | 0 670714716 | -0 4963657 | 0 | 0 690140904 |
| CASP5 | 0 001513392 | 0 001301748 | 0 673072901 | -0 49535151 | 0 | 0 692416603 |
| EMP2 | 0 220679218 | 0 001294388 | 0 674815158 | 0 273943449 | 0 | 0 694133367 |
| MS4A4A | 0 020140488 | 0 001293962 | 0 674915985 | 0 474961164 | 0 | 0 694161521 |
| TIPARP | 0 204745769 | 0 001278103 | 0 678676306 | -0 28254335 | 0 | 0 697953106 |
| PDZD8 | 0 348727418 | 0 001277296 | 0 678867865 | 0 193412211 | 0 | 0 698074138 |
| FANCB | 0 112638252 | 0 001271642 | 0 680210464 | 0 30536181 | 0 | 0 699242319 |
| ATP8B4 | 0 03721612 | 0 001271577 | 0 680225887 | -0 45450372 | 0 | 0 699242319 |
| JARID2-AS1 | 0 001180255 | 0 001263318 | 0 682188918 | 0 495948166 | 0 | 0 701031456 |
| TIMD4 | 0 001047001 | 0 001255716 | 0 683997857 | 0 495888102 | 0 | 0 702813934 |
| BTK | 0 004997049 | 0 001254251 | 0 684346616 | -0 49191051 | 0 | 0 703095838 |
| SPATA1 | 0 160552816 | 0 001234521 | 0 689050385 | 0 334107248 | 0 | 0 707697657 |
| ADAM8 | 0 020730616 | 0 001227279 | 0 690780091 | 0 473489775 | 0 | 0 709319989 |
| QPCT | 0 039871695 | 0 001220098 | 0 692496733 | -0 44626415 | 0 | 0 710928205 |
| ARHGEF37 | 0 063933677 | 0 001219133 | 0 692727541 | -0 42162892 | 0 | 0 711010673 |
| EMB | 0 149368944 | 0 001215818 | 0 693520551 | 0 338093666 | 0 | 0 711747308 |
| FCRL2 | 0 000723382 | 0 001212456 | 0 694325336 | -0 49614706 | 0 | 0 712495866 |
| SPTA1 | 0 001741829 | 0 00121082 | 0 694717044 | -0 49511974 | 0 | 0 71282042 |
| IRF5 | 0 00641526 | 0 001178654 | 0 702434635 | -0 4892752 | 0 | 0 720504428 |
| FPR2 | 0 00120881 | 0 001168171 | 0 704956754 | 0 495713112 | 0 | 0 722951435 |
| LPCAT2 | 0 272305877 | 0 001167695 | 0 705071271 | -0 21728006 | 0 | 0 722973471 |
| LINC02541 | 0 009527707 | 0 001152467 | 0 708740758 | 0 486957372 | 0 | 0 726657282 |
| UXT-AS1 | 0 00877577 | 0 001127042 | 0 714882378 | -0 48753597 | 0 | 0 732636208 |
| TULP2 | 0 011916773 | 0 001124028 | 0 715611524 | -0 48250097 | 0 | 0 733303937 |
| IL13RA2 | 0 026708039 | 0 001113767 | 0 718096274 | 0 468943844 | 0 | 0 735665415 |
| LINC01060 | 0 094801165 | 0 001113547 | 0 718149572 | -0 38501843 | 0 | 0 735665415 |
| IGLC2 | 0 011916773 | 0 00110361 | 0 720558842 | 0 481759499 | 0 | 0 738053442 |

| | | | | | | |
|---|---|---|---|---|---|---|
| RETN | 0 000333137 | 0 001098809 | 0 721724022 | -0 49658063 | 0 | 0 739166794 |
| ASCL4 | 0 000133255 | 0 001097052 | 0 722150412 | -0 4967771 | 0 | 0 739500375 |
| MT3 | 0 006548514 | 0 001096822 | 0 722206239 | -0 48738064 | 0 | 0 739500375 |
| PPP6R3 | 0 71556795 | 0 001093934 | 0 72290759 | 0 016514865 | 8 64E-08 | 0 739977982 |
| ANO3 | 0 040909177 | 0 001092566 | 0 723239755 | -0 44923915 | 0 | 0 740237808 |
| SLAMF1 | 0 001427728 | 0 00108914 | 0 72407219 | 0 495509676 | 0 | 0 741009551 |
| SLC4A8 | 0 073775485 | 0 001081864 | 0 725840949 | 0 38759677 | 0 | 0 742739249 |
| STAP1 | 0 003026784 | 0 001072811 | 0 728043555 | 0 493280452 | 0 | 0 744912472 |
| EIF4A2 | 0 509242162 | 0 001067534 | 0 72932873 | -0 09667155 | 1 59E-216 | 0 746146638 |
| STIM1 | 0 401591441 | 0 00106106 | 0 730906281 | -0 15207082 | 0 | 0 747598697 |
| NALCN | 0 157192896 | 0 001060706 | 0 730992634 | -0 31928051 | 0 | 0 747606103 |
| ACKR1 | 0 011069654 | 0 001048901 | 0 733872154 | -0 48478886 | 0 | 0 750388644 |
| ZNF430 | 0 20258514 | 0 001042978 | 0 735318528 | 0 253493946 | 0 | 0 751786226 |
| FLOT1 | 0 233252746 | 0 001028584 | 0 738836964 | 0 282445249 | 0 | 0 755301743 |
| TEX264 | 0 077801679 | 0 001024999 | 0 739714179 | -0 4150551 | 0 | 0 756116711 |
| CD3D | 0 004035712 | 0 001021672 | 0 740528573 | -0 49320074 | 0 | 0 756785445 |
| RAC1 | 0 440949154 | 0 001011755 | 0 742957858 | -0 13378227 | 0 | 0 75918596 |
| GPR55 | 0 002132074 | 0 001009038 | 0 743623727 | 0 493663113 | 0 | 0 759784217 |
| C12orf42 | 0 01881746 | 0 001004519 | 0 744731773 | -0 47775766 | 0 | 0 760834082 |
| NOXA1 | 0 021739544 | 0 000992074 | 0 747786149 | -0 46948173 | 0 | 0 763789354 |
| PNMA5 | 0 000904228 | 0 000959076 | 0 75590386 | 0 496083893 | 0 | 0 771913926 |
| TRGV10 | 0 000152291 | 0 000950555 | 0 758004386 | -0 49682671 | 0 | 0 773808086 |
| SLC4A7 | 0 558156136 | 0 0009467 | 0 758955487 | -0 07091988 | 3 21E-117 | 0 774611659 |
| A2MP1 | 0 000989892 | 0 00094078 | 0 760416309 | -0 49560776 | 0 | 0 775935009 |
| C1orf167 | 0 005577659 | 0 000935486 | 0 761723613 | -0 49058072 | 0 | 0 777185073 |
| HBB | 0 021682435 | 0 000933437 | 0 762229926 | -0 47407529 | 0 | 0 777617705 |
| FCER2 | 0 000818564 | 0 000928161 | 0 763533691 | -0 49603194 | 0 | 0 778863707 |
| CD2 | 0 006205859 | 0 000923542 | 0 76467561 | 0 489628178 | 0 | 0 779944362 |
| CCL20 | 0 00252232 | 0 000919921 | 0 765571255 | 0 494594361 | 0 | 0 780773619 |
| DCDC2 | 0 020740134 | 0 000919336 | 0 765715784 | -0 46802507 | 0 | 0 780836749 |
| CT69 | 0 006462851 | 0 000914417 | 0 766933021 | -0 48953729 | 0 | 0 78199364 |
| FYB1 | 0 054596334 | 0 000895243 | 0 771682994 | -0 44188069 | 0 | 0 78666713 |
| KCTD9 | 0 347756563 | 0 000892155 | 0 772448874 | -0 19574055 | 0 | 0 787362944 |
| HELZ2 | 0 06455236 | 0 000890782 | 0 772789287 | 0 423816618 | 0 | 0 787624974 |
| MAP3K8 | 0 139622318 | 0 000877128 | 0 776178579 | -0 31862253 | 0 | 0 790823458 |
| PLK2 | 0 172612362 | 0 00087173 | 0 77751954 | 0 332530637 | 0 | 0 792104317 |
| ZNF385A | 0 029791932 | 0 000867148 | 0 778658391 | -0 46637326 | 0 | 0 793093532 |
| VASP | 0 08657745 | 0 000861172 | 0 780144421 | -0 40900591 | 0 | 0 794521476 |
| TMEM238 | 0 006586587 | 0 000854213 | 0 781875917 | -0 48853868 | 0 | 0 796113287 |
| ITGAM | 0 011421827 | 0 000845526 | 0 784039012 | 0 484494212 | 0 | 0 798143775 |
| P2RY6 | 0 005901277 | 0 000802972 | 0 794658224 | -0 49082073 | 0 | 0 808605601 |
| ARC | 0 004759095 | 0 000797454 | 0 796038132 | -0 49104697 | 0 | 0 809835326 |
| SH2D1A | 0 001389656 | 0 000784347 | 0 799318177 | 0 495576141 | 0 | 0 812831326 |
| MIR23AHG | 0 114913099 | 0 000784311 | 0 799327144 | -0 35598374 | 0 | 0 812831326 |

| | | | | | | |
|---|---|---|---|---|---|---|
| UHRF1 | 0 076488169 | 0 000774089 | 0 801887986 | 0 331012872 | 0 | 0 815172275 |
| SLC2A6 | 0 017580096 | 0 000753771 | 0 806983741 | 0 483641577 | 0 | 0 82017599 |
| TLDC2 | 0 005701395 | 0 000735857 | 0 811483709 | -0 48970711 | 0 | 0 824572156 |
| CCNB3 | 0 010412899 | 0 000735157 | 0 811659492 | 0 481804117 | 0 | 0 824662101 |
| CAB39 | 0 406588491 | 0 000709763 | 0 818049254 | 0 172244872 | 0 | 0 830787029 |
| TLR10 | 0 00152291 | 0 000708378 | 0 818398161 | -0 49519434 | 0 | 0 830972668 |
| HIGD1B | 0 021587253 | 0 00070732 | 0 818664617 | 0 468464047 | 0 | 0 831153914 |
| CYP4F3 | 0 002788829 | 0 000696167 | 0 821475362 | -0 49367779 | 0 | 0 833828375 |
| CKS1B | 0 075964668 | 0 000634621 | 0 837025971 | 0 393160035 | 0 | 0 849065632 |
| PRPS2 | 0 095714911 | 0 000629149 | 0 838411947 | 0 386558525 | 0 | 0 850288999 |
| XKR7 | 0 028050104 | 0 000628487 | 0 838579613 | -0 45926724 | 0 | 0 85036778 |
| SIRPB1 | 0 001351583 | 0 00062659 | 0 839060101 | -0 49558435 | 0 | 0 850763729 |
| PRMT1 | 0 163731892 | 0 00062099 | 0 840478956 | -0 3347578 | 0 | 0 852110946 |
| ITGAD | 0 006900687 | 0 000610114 | 0 843236347 | 0 486194099 | 0 | 0 854539784 |
| YWHAQ | 0 468085511 | 0 000604022 | 0 844781722 | 0 138426418 | 0 | 0 856014076 |
| UBE2C | 0 039338676 | 0 000601687 | 0 845374165 | 0 408828934 | 0 | 0 856522554 |
| SVOPL | 0 003740648 | 0 000600023 | 0 845796411 | 0 493184899 | 0 | 0 8568585 |
| GRK5 | 0 366878605 | 0 000597335 | 0 846478535 | -0 16625113 | 0 | 0 857457622 |
| UTS2 | 0 003121966 | 0 000580722 | 0 850697351 | -0 49358731 | 0 | 0 861334991 |
| GRK6 | 0 053806324 | 0 000574781 | 0 852207178 | 0 432595275 | 0 | 0 862705713 |
| LDLRAD3 | 0 615779254 | 0 000552899 | 0 857772336 | -0 04816392 | 5 29E-55 | 0 86815343 |
| CABP4 | 0 00441644 | 0 000547232 | 0 859214915 | -0 49181638 | 0 | 0 869520342 |
| GLYATL1 | 0 013420647 | 0 000529477 | 0 863737186 | 0 474030867 | 0 | 0 873909679 |
| EBI3 | 0 002331956 | 0 000527542 | 0 864230259 | -0 49438515 | 0 | 0 87431495 |
| FCRL1 | 0 000485428 | 0 00051998 | 0 866158179 | 0 496535216 | 0 | 0 87607779 |
| CDHR1 | 0 00462584 | 0 000516591 | 0 867022294 | 0 492660564 | 0 | 0 87685795 |
| SIGLEC8 | 0 000704346 | 0 000514516 | 0 867551559 | 0 496226442 | 0 | 0 877299329 |
| MED11 | 0 041613523 | 0 000487757 | 0 874381142 | -0 4523035 | 0 | 0 883921886 |
| PCED1B-AS1 | 0 006082123 | 0 000475408 | 0 877536156 | 0 489638581 | 0 | 0 886826721 |
| LDHC | 0 007233824 | 0 000466865 | 0 879719872 | 0 490505773 | 0 | 0 888748427 |
| VNN2 | 0 002684129 | 0 000428946 | 0 889423488 | -0 49414912 | 0 | 0 897783803 |
| DIAPH2-AS1 | 0 156412404 | 0 000427487 | 0 889797187 | 0 335051964 | 0 | 0 898065088 |
| LACC1 | 0 015238621 | 0 00042039 | 0 891615226 | 0 482693126 | 0 | 0 899707836 |
| ATP13A3 | 0 464801736 | 0 000396963 | 0 897620951 | 0 152655071 | 0 | 0 905478008 |
| PCP2 | 0 000742419 | 0 000391476 | 0 899028406 | 0 496049872 | 0 | 0 906704208 |
| BATF | 0 002674611 | 0 00038989 | 0 89943544 | 0 494411536 | 0 | 0 906921139 |
| SPINT2 | 0 015552721 | 0 000380759 | 0 901778468 | -0 47981707 | 0 | 0 909089665 |
| SIRPB2 | 0 003616912 | 0 000351785 | 0 909218931 | -0 4929787 | 0 | 0 916297206 |
| SMIM35 | 0 006167787 | 0 000345174 | 0 910917957 | -0 48960982 | 0 | 0 917911569 |
| PIK3CG | 0 007119606 | 0 000337355 | 0 912927699 | 0 489496456 | 0 | 0 919838656 |
| AXL | 0 367516324 | 0 000335831 | 0 913319577 | 0 166830931 | 0 | 0 920135395 |
| IL10RA | 0 01498163 | 0 000292178 | 0 924550366 | -0 48145516 | 0 | 0 930755402 |
| ADAMTS8 | 0 018465287 | 0 000288473 | 0 925504441 | -0 47135155 | 0 | 0 931616635 |
| LINC00924 | 0 182673088 | 0 000286536 | 0 926003238 | -0 29412631 | 0 | 0 932019448 |

| | | | | | | |
|---|---|---|---|---|---|---|
| C8G | 0 002132074 | 0 000280614 | 0 927528354 | -0 4943579 | 0 | 0 933455053 |
| PIM2 | 0 013696674 | 0 000263782 | 0 931864423 | 0 476919717 | 0 | 0 937619123 |
| SH3GLB1 | 0 352363366 | 0 000250371 | 0 935320825 | -0 18527336 | 0 | 0 940896508 |
| SC5D | 0 082389446 | 0 000249869 | 0 935450074 | -0 39917463 | 0 | 0 940926365 |
| CLEC4C | 0 001189774 | 0 000246982 | 0 9361943 | 0 495612977 | 0 | 0 941574727 |
| ERAP2 | 0 133702005 | 0 00022286 | 0 942414579 | -0 35993056 | 0 | 0 947729889 |
| CD300E | 0 001646647 | 0 000208136 | 0 94621329 | 0 495268231 | 0 | 0 951246339 |
| SCN1A | 0 007966724 | 0 00019552 | 0 94946887 | 0 492582943 | 0 | 0 954316191 |
| FCGR2A | 0 029934705 | 0 000191874 | 0 950410096 | 0 46128164 | 0 | 0 95505906 |
| GMFG | 0 017446841 | 0 000183889 | 0 952471158 | 0 477720752 | 0 | 0 95692669 |
| DUSP28 | 0 017599132 | 0 000180643 | 0 953309218 | 0 479909165 | 0 | 0 957666855 |
| ADAP1 | 0 00725286 | 0 000174091 | 0 955000843 | -0 48753099 | 0 | 0 959060357 |
| NCR3 | 0 000485428 | 0 000172578 | 0 955391657 | -0 49639081 | 0 | 0 959350883 |
| TATDN2 | 0 114513335 | 0 000166764 | 0 956892856 | 0 354980254 | 0 | 0 960756215 |
| SCN1B | 0 016780568 | 0 000165021 | 0 957343058 | -0 47648443 | 0 | 0 961106119 |
| MIF4GD | 0 03994784 | 0 000163372 | 0 957768885 | 0 459948009 | 0 | 0 961431481 |
| TRGV2 | 2 86E-05 | 0 000162424 | 0 958013665 | -0 49694063 | 0 | 0 961575054 |
| NKX3-1 | 0 006424778 | 0 00015524 | 0 959869201 | -0 48997751 | 0 | 0 963232872 |
| IL21R-AS1 | 0 000856637 | 0 000150805 | 0 96101481 | -0 49601616 | 0 | 0 964280097 |
| ZNF683 | 0 000475909 | 0 000149826 | 0 96126753 | 0 496135537 | 0 | 0 964431273 |
| DYNC1H1 | 0 514705602 | 0 000146288 | 0 962181496 | -0 0889212 | 2 18E-183 | 0 965245768 |
| TALDO1 | 0 132607413 | 0 00013897 | 0 96407198 | -0 36226169 | 0 | 0 966731772 |
| PRSS36 | 0 003816794 | 0 000126984 | 0 967169057 | -0 49344454 | 0 | 0 969631614 |
| TPSAB1 | 0 000885192 | 0 00011664 | 0 969841965 | 0 495980666 | 0 | 0 972208186 |
| LINC00324 | 0 012326055 | 0 000106726 | 0 972404276 | -0 48298709 | 0 | 0 974673357 |
| FAM111B | 0 041223278 | 0 000101709 | 0 973700968 | 0 402671062 | 0 | 0 975869567 |
| FUCA1 | 0 060944966 | 9 95E-05 | 0 974275933 | -0 43559608 | 0 | 0 976238741 |
| BTG2 | 0 154822866 | 8 09E-05 | 0 979082608 | -0 31352737 | 0 | 0 980743125 |
| NXT1 | 0 045135253 | 7 47E-05 | 0 980678476 | 0 456879188 | 0 | 0 982029416 |
| LINC02133 | 0 006538996 | 6 66E-05 | 0 982773728 | 0 489035303 | 0 | 0 984023281 |
| UBASH3A | 0 001170737 | 4 23E-05 | 0 98906326 | 0 495735719 | 0 | 0 989691635 |
| LINC01146 | 0 001846529 | 3 68E-05 | 0 990486644 | 0 494252217 | 0 | 0 990906075 |
| HNRNPA0 | 0 166558794 | 1 05E-05 | 0 997277486 | 0 361846048 | 0 | 0 997488595 |
| CHD1 | 0 492118939 | 9 26E-06 | 0 997605465 | 0 124739363 | 0 | 0 997711043 |
| APOBEC3H | 0 002798348 | -6 96E-07 | 0 999820122 | -0 49385389 | 0 | 0 999820122 |
| DOC2B | 0 033532581 | -2 82E-05 | 0 992715526 | -0 45970805 | 0 | 0 993030773 |
| FCN1 | 0 003892939 | -3 76E-05 | 0 990266582 | 0 492549975 | 0 | 0 99079081 |
| LINC01010 | 0 01077459 | -4 58E-05 | 0 988165983 | -0 48455058 | 0 | 0 988898501 |
| APOBEC3D | 0 023205345 | -4 89E-05 | 0 987365923 | -0 47284192 | 0 | 0 988202496 |
| CACNA1F | 0 001294474 | -5 52E-05 | 0 985718343 | -0 49555252 | 0 | 0 986658017 |
| SLC25A30-AS1 | 0 003236184 | -5 78E-05 | 0 985050374 | -0 49322081 | 0 | 0 986093859 |
| HES6 | 0 018465287 | -6 44E-05 | 0 983355773 | 0 481698 | 0 | 0 984501754 |
| MOB1A | 0 231510917 | -7 76E-05 | 0 979936485 | 0 287337929 | 0 | 0 981390397 |
| SEC61G | 0 243342027 | -8 00E-05 | 0 979322767 | -0 27338328 | 0 | 0 980879719 |

| | | | | | | |
|---|---|---|---|---|---|---|
| LINC02653 | 0 005139822 | -8 65E-05 | 0 977628839 | 0 492587732 | 0 | 0 979390706 |
| INPP1 | 0 249557404 | -9 66E-05 | 0 975031264 | -0 26351662 | 0 | 0 976892011 |
| PPM1N | 0 014248729 | -0 00010098 | 0 97389002 | 0 479266756 | 0 | 0 975955535 |
| MPP1 | 0 047609983 | -0 00013304 | 0 96560352 | 0 449052071 | 0 | 0 968164802 |
| MLKL | 0 04094725 | -0 00013905 | 0 964051021 | -0 45257475 | 0 | 0 966731772 |
| JADE3 | 0 200881384 | -0 00014038 | 0 963706769 | 0 304909614 | 0 | 0 966570683 |
| ASGR2 | 0 0005235 | -0 0001448 | 0 962565769 | -0 49636458 | 0 | 0 965528768 |
| CD34 | 0 105728046 | -0 00016168 | 0 958206386 | -0 37318128 | 0 | 0 96166635 |
| RIT1 | 0 124935752 | -0 0001755 | 0 954636372 | -0 36579899 | 0 | 0 958796229 |
| CD1E | 0 000571091 | -0 00017779 | 0 954045188 | -0 49642435 | 0 | 0 958304318 |
| CD83 | 0 052921132 | -0 00019075 | 0 950699966 | -0 43908624 | 0 | 0 95524877 |
| GTSE1 | 0 05340656 | -0 00019423 | 0 949802434 | 0 375139232 | 0 | 0 954549931 |
| AIF1 | 0 029125659 | -0 00019733 | 0 949000733 | -0 46983076 | 0 | 0 953947126 |
| LINC01141 | 0 001380137 | -0 00021072 | 0 945547349 | 0 494798464 | 0 | 0 950677992 |
| MPDU1 | 0 068369153 | -0 00022039 | 0 943050676 | -0 42428181 | 0 | 0 948268662 |
| PTAFR | 0 011840627 | -0 00025243 | 0 934790578 | 0 484064035 | 0 | 0 940463214 |
| GIMAP7 | 0 009099389 | -0 0002705 | 0 930133863 | -0 48736889 | 0 | 0 935977533 |
| LRIF1 | 0 122955969 | -0 00029273 | 0 924407516 | 0 365533236 | 0 | 0 930710742 |
| STARD4 | 0 105927928 | -0 00030003 | 0 922529013 | 0 377866093 | 0 | 0 928918399 |
| SIT1 | 0 000552055 | -0 00030032 | 0 922454802 | 0 496273842 | 0 | 0 928918399 |
| PIGA | 0 045725381 | -0 00030103 | 0 922271536 | 0 44666588 | 0 | 0 928857083 |
| LRRC26 | 0 000171327 | -0 00030267 | 0 921849263 | -0 49673878 | 0 | 0 928530754 |
| ABLIM2 | 0 034979346 | -0 00031449 | 0 918809046 | -0 45274307 | 0 | 0 925567156 |
| TMIGD3 | 0 004168967 | -0 00036816 | 0 905013467 | -0 49239495 | 0 | 0 912156278 |
| NBEAL2 | 0 032552207 | -0 00037897 | 0 902236584 | -0 46096448 | 0 | 0 909454477 |
| DCDC2C | 0 091507872 | -0 00038621 | 0 900378973 | -0 37895231 | 0 | 0 907775663 |
| CREB3L2 | 0 540471341 | -0 00039109 | 0 899126797 | -0 09525102 | 3 06E-210 | 0 906706673 |
| LINC00847 | 0 02575622 | -0 00039378 | 0 898436483 | -0 46831963 | 0 | 0 906203946 |
| LCN6 | 0 000618682 | -0 00039981 | 0 896889968 | -0 49620431 | 0 | 0 904837216 |
| DRAM2 | 0 161409453 | -0 00041094 | 0 894036281 | 0 337231386 | 0 | 0 902054544 |
| CYTOR | 0 33394567 | -0 00042382 | 0 890735987 | 0 163859408 | 0 | 0 898916603 |
| MFSD6 | 0 284479641 | -0 00043792 | 0 887125198 | 0 240799329 | 0 | 0 895559569 |
| STUM | 0 070624964 | -0 00045048 | 0 883910972 | -0 41810462 | 0 | 0 892410116 |
| IL6 | 0 004797167 | -0 0004514 | 0 883675302 | -0 49117501 | 0 | 0 892267508 |
| ZCWPW1 | 0 082256192 | -0 00045469 | 0 882833222 | -0 40949577 | 0 | 0 891512497 |
| ANKRD28 | 0 585273458 | -0 00045889 | 0 881759622 | 0 073255875 | 5 88E-125 | 0 890523504 |
| ADGRD1 | 0 274057223 | -0 00046411 | 0 880424691 | -0 21406548 | 0 | 0 889270343 |
| KCNK17 | 0 004759095 | -0 00046586 | 0 879976062 | -0 49063893 | 0 | 0 888912217 |
| CSF3R | 0 011440863 | -0 00046948 | 0 879050119 | -0 48422096 | 0 | 0 888166751 |
| NLRP3 | 0 009737108 | -0 00047252 | 0 878274204 | -0 48676803 | 0 | 0 887477676 |
| TMOD1 | 0 08699625 | -0 00047952 | 0 876484324 | -0 39649772 | 0 | 0 885858487 |
| CCDC153 | 0 003369439 | -0 00048314 | 0 875560429 | 0 494402759 | 0 | 0 885019366 |
| GML | 0 002646057 | -0 00048822 | 0 874263761 | 0 493771778 | 0 | 0 883897779 |
| RNF139 | 0 075365023 | -0 0004929 | 0 873067551 | -0 41833868 | 0 | 0 882782833 |

| | | | | | | |
|---|---|---|---|---|---|---|
| WNT6 | 0 025879957 | -0 00052074 | 0 865964614 | -0 46238879 | 0 | 0 875975766 |
| PTPRC | 0 066874798 | -0 00054132 | 0 860719875 | 0 420068258 | 0 | 0 870950082 |
| DSCAM | 0 042089433 | -0 00055439 | 0 857393283 | -0 44894053 | 0 | 0 867862739 |
| SLC15A4 | 0 150168472 | -0 00058047 | 0 85076199 | 0 35246076 | 0 | 0 861334991 |
| GLOD5 | 0 003064857 | -0 00058469 | 0 849688623 | -0 49321092 | 0 | 0 86043265 |
| SFI1 | 0 184329253 | -0 00059029 | 0 848267314 | 0 286556548 | 0 | 0 859085427 |
| LAIR2 | 0 000342655 | -0 00059083 | 0 848129448 | 0 496713923 | 0 | 0 859037866 |
| GASK1B | 0 249214749 | -0 0006133 | 0 842427961 | -0 26257491 | 0 | 0 853812123 |
| IL21R | 0 005025604 | -0 00061336 | 0 84241314 | 0 490754862 | 0 | 0 853812123 |
| WDR76 | 0 083655365 | -0 00061577 | 0 841801442 | 0 333989438 | 0 | 0 853360183 |
| DBP | 0 034294036 | -0 00063205 | 0 837677077 | -0 46028657 | 0 | 0 849634902 |
| PFKFB3 | 0 238687632 | -0 00064413 | 0 83461877 | -0 24068546 | 0 | 0 846714694 |
| FUT7 | 0 000285546 | -0 00064673 | 0 833960183 | -0 49665549 | 0 | 0 846137398 |
| JCHAIN | 0 003131484 | -0 00065152 | 0 832749628 | 0 493476055 | 0 | 0 844999891 |
| PIK3R6 | 0 009603853 | -0 00065294 | 0 832389906 | -0 48424931 | 0 | 0 844725581 |
| FOXM1 | 0 070253755 | -0 00067582 | 0 826607952 | 0 352851782 | 0 | 0 838948034 |
| NCF1 | 0 004778131 | -0 00069755 | 0 82112751 | 0 491155625 | 0 | 0 833564827 |
| TMPPE | 0 003150521 | -0 00070945 | 0 818127418 | -0 49332896 | 0 | 0 830787029 |
| PDGFRL | 0 158830024 | -0 00071483 | 0 816774304 | -0 31737561 | 0 | 0 82959127 |
| VMA21 | 0 096895167 | -0 00071714 | 0 816192526 | 0 418063408 | 0 | 0 829089474 |
| DPEP3 | 0 000237955 | -0 00073369 | 0 81202908 | 0 496715606 | 0 | 0 824948915 |
| ICAM3 | 0 015095848 | -0 00075083 | 0 807721777 | 0 482233098 | 0 | 0 82083781 |
| MOSMO | 0 416163789 | -0 00076967 | 0 80299617 | -0 1470946 | 0 | 0 816211015 |
| LRRC8C | 0 124602616 | -0 00077771 | 0 800981063 | 0 348723699 | 0 | 0 814337929 |
| ZDHHC18 | 0 088871333 | -0 00078145 | 0 800044231 | 0 386872809 | 0 | 0 813472991 |
| FFAR3 | 0 000304582 | -0 00078607 | 0 798885817 | 0 496554459 | 0 | 0 812557418 |
| PDCD1 | 0 001151701 | -0 00079465 | 0 796739619 | -0 49603768 | 0 | 0 810461722 |
| PPP1R3B | 0 060811711 | -0 00080196 | 0 794910873 | -0 42875072 | 0 | 0 808775597 |
| FFAR2 | 0 000276027 | -0 00082037 | 0 790311988 | 0 49658687 | 0 | 0 804269684 |
| GALR1 | 0 006719842 | -0 00083334 | 0 787074878 | -0 4900371 | 0 | 0 801061669 |
| LYAR | 0 067988426 | -0 00083786 | 0 785948564 | 0 422035029 | 0 | 0 800001501 |
| PNRC1 | 0 228950524 | -0 00085359 | 0 78203164 | -0 25123933 | 0 | 0 796186059 |
| SLCO1C1 | 0 118225429 | -0 0008572 | 0 781131867 | -0 37809207 | 0 | 0 795441395 |
| NRG2 | 0 125354553 | -0 00086856 | 0 778306559 | 0 334049644 | 0 | 0 79282063 |
| PFDN1 | 0 268013173 | -0 00088678 | 0 773781427 | -0 2459761 | 0 | 0 788466086 |
| CAPZA1 | 0 433905694 | -0 00088937 | 0 773139111 | 0 146282851 | 0 | 0 787896538 |
| BTN3A2 | 0 081304373 | -0 00091054 | 0 76789334 | 0 41314194 | 0 | 0 782888344 |
| GZMA | 0 006558032 | -0 00094487 | 0 759407586 | 0 487492649 | 0 | 0 774989383 |
| CLSPN | 0 067455407 | -0 00094902 | 0 758384062 | 0 351012515 | 0 | 0 774112053 |
| LINC02416 | 0 000295064 | -0 00095571 | 0 756734475 | 0 496693714 | 0 | 0 772595159 |
| YPEL3 | 0 138670499 | -0 00095774 | 0 756233687 | -0 34917627 | 0 | 0 772167298 |
| ENPP4 | 0 03200967 | -0 00097303 | 0 752468048 | -0 45582939 | 0 | 0 768488388 |
| KLF9 | 0 327406674 | -0 00099407 | 0 747296695 | -0 17741082 | 0 | 0 763371935 |
| ZNF431 | 0 300222726 | -0 00102195 | 0 740461623 | 0 18679918 | 0 | 0 756785445 |

| | | | | | | |
|---|---|---|---|---|---|---|
| TMIGD2 | 0 000742419 | -0 00105757 | 0 73175627 | 0 496169807 | 0 | 0 748306109 |
| F13A1 | 0 047076964 | -0 00106536 | 0 729859296 | 0 436151415 | 0 | 0 746608611 |
| KIR2DL4 | 0 000237955 | -0 00109507 | 0 722632845 | 0 496604815 | 0 | 0 739776881 |
| ADAMDEC1 | 0 000856637 | -0 00109637 | 0 722314956 | 0 495623009 | 0 | 0 739531564 |
| VTN | 0 003645466 | -0 00111549 | 0 717678173 | -0 49283351 | 0 | 0 735341943 |
| TASL | 0 001589538 | -0 00113165 | 0 713768571 | -0 49490973 | 0 | 0 731574077 |
| KIR3DL1 | 3 81E-05 | -0 00114217 | 0 711227 | 0 496852126 | 0 | 0 729048178 |
| PDIK1L | 0 041194723 | -0 00115058 | 0 70919565 | 0 443622686 | 0 | 0 727044792 |
| TXLNB | 0 042194133 | -0 0011681 | 0 704973278 | -0 45306952 | 0 | 0 722951435 |
| LILRB1 | 0 005768023 | -0 00119033 | 0 699630331 | -0 49075525 | 0 | 0 717705886 |
| PLD1 | 0 36249072 | -0 00120116 | 0 697031821 | -0 20634452 | 0 | 0 715117871 |
| MAPK6 | 0 357722107 | -0 00121941 | 0 692661801 | 0 198969773 | 0 | 0 711010673 |
| OAS1 | 0 05009423 | -0 00122454 | 0 691433763 | -0 44867179 | 0 | 0 709914066 |
| FRYL | 0 587748187 | -0 00123313 | 0 689382256 | 0 076826237 | 2 85E-137 | 0 707961565 |
| MLEC | 0 257467019 | -0 00124359 | 0 686887251 | 0 287959332 | 0 | 0 705552665 |
| CCL22 | 0 000456873 | -0 0012488 | 0 685645182 | -0 49644211 | 0 | 0 704353405 |
| CCL23 | 0 000380728 | -0 00126536 | 0 681703865 | 0 496390094 | 0 | 0 700609193 |
| OSBP2 | 0 149045326 | -0 00126983 | 0 68064199 | 0 333603387 | 0 | 0 699593953 |
| KANSL1-AS1 | 0 023243418 | -0 0012723 | 0 680054818 | -0 47418383 | 0 | 0 699218587 |
| CXCL2 | 0 016856713 | -0 0013105 | 0 671004709 | 0 476491089 | 0 | 0 690364126 |
| EVI2B | 0 086920104 | -0 00131346 | 0 670303872 | 0 392940761 | 0 | 0 689793269 |
| NMRK1 | 0 095857684 | -0 00131623 | 0 669650121 | 0 406138904 | 0 | 0 689195561 |
| TRGV7 | 5 71E-05 | -0 00132115 | 0 66848936 | 0 496912954 | 0 | 0 688075858 |
| EOMES | 0 002170147 | -0 00132341 | 0 667956622 | 0 494431829 | 0 | 0 687602405 |
| SLC45A3 | 0 008632998 | -0 00133925 | 0 664222531 | -0 48617731 | 0 | 0 683982011 |
| B3GALT2 | 0 024004873 | -0 00134055 | 0 663916308 | -0 47244631 | 0 | 0 683741184 |
| CST6 | 0 002065447 | -0 00134803 | 0 662157272 | -0 49422198 | 0 | 0 682078289 |
| LRIG1 | 0 291465992 | -0 00135281 | 0 661035504 | -0 25046433 | 0 | 0 681219794 |
| SLC2A13 | 0 371019017 | -0 00135502 | 0 66051487 | -0 18820618 | 0 | 0 680757501 |
| ADO | 0 03238088 | -0 00135847 | 0 659705532 | -0 46040454 | 0 | 0 679997522 |
| TNFRSF19 | 0 282090575 | -0 00136213 | 0 658847491 | -0 21076507 | 0 | 0 67918717 |
| TINF2 | 0 047086482 | -0 00139297 | 0 651628611 | -0 44224985 | 0 | 0 672112025 |
| MRTFA | 0 521863281 | -0 00142631 | 0 643859917 | 0 102247883 | 4 16E-242 | 0 664389191 |
| ARHGAP30 | 0 014439093 | -0 00147117 | 0 633469759 | -0 48205002 | 0 | 0 654239259 |
| BPGM | 0 052064495 | -0 00147218 | 0 633236078 | -0 442612 | 0 | 0 654069399 |
| GLA | 0 061401839 | -0 00147518 | 0 632543499 | -0 4294822 | 0 | 0 653425456 |
| LST1 | 0 013677638 | -0 00147739 | 0 632034744 | -0 48185805 | 0 | 0 652971283 |
| SLC9C2 | 0 020873389 | -0 00147953 | 0 631541877 | 0 474541614 | 0 | 0 652533428 |
| SIGLEC7 | 0 001465801 | -0 00147974 | 0 631493261 | 0 495144104 | 0 | 0 652533428 |
| SHOC2 | 0 427823571 | -0 00149367 | 0 628284446 | 0 162627071 | 0 | 0 649451763 |
| DNAAF1 | 0 006995869 | -0 00149637 | 0 62766511 | -0 48754124 | 0 | 0 648953533 |
| NR2F2 | 0 407730673 | -0 00150315 | 0 626106309 | -0 14879792 | 0 | 0 647483543 |
| S1PR4 | 0 000875673 | -0 00150588 | 0 625479357 | 0 495922314 | 0 | 0 646905978 |
| CLPB | 0 105642383 | -0 00151218 | 0 624034689 | 0 381611346 | 0 | 0 645553127 |

| | | | | | | |
|---|---|---|---|---|---|---|
| TMEM74 | 0 015428985 | -0 0015188 | 0 622517408 | -0 47829585 | 0 | 0 644054029 |
| SBSPON | 0 07995279 | -0 00153415 | 0 619004142 | -0 39310276 | 0 | 0 640559476 |
| BZW2 | 0 247491957 | -0 00154692 | 0 616089254 | 0 279601143 | 0 | 0 637822464 |
| SLC27A2 | 0 003255221 | -0 00156201 | 0 612651733 | 0 493951799 | 0 | 0 634472205 |
| HNF1A | 0 001437247 | -0 00160652 | 0 602561146 | 0 494896717 | 0 | 0 624501297 |
| MTA2 | 0 047029373 | -0 00160723 | 0 602401704 | 0 445324858 | 0 | 0 62440453 |
| FAM210A | 0 138346881 | -0 00163862 | 0 59533367 | 0 357950428 | 0 | 0 617484709 |
| KTN1 | 0 671908016 | -0 00163933 | 0 595173698 | 0 065425173 | 5 16E-100 | 0 617386548 |
| CD244 | 0 001037483 | -0 00166929 | 0 588463259 | 0 495568521 | 0 | 0 610827965 |
| JOSD1 | 0 151510537 | -0 0017001 | 0 581598808 | 0 343701804 | 0 | 0 603835282 |
| HERC6 | 0 106965411 | -0 00172402 | 0 576294421 | 0 371851923 | 0 | 0 59859115 |
| ZNF296 | 0 004483067 | -0 00173184 | 0 574565516 | 0 491071265 | 0 | 0 596926575 |
| ST6GALNAC6 | 0 088071805 | -0 00176016 | 0 56832475 | -0 39284131 | 0 | 0 590702693 |
| LCK | 0 004502104 | -0 00177433 | 0 565215616 | 0 491982044 | 0 | 0 587535757 |
| LINC00698 | 0 001389656 | -0 00177678 | 0 564679764 | 0 494969341 | 0 | 0 587043319 |
| TMEM236 | 0 00593935 | -0 00178297 | 0 563323929 | 0 491814897 | 0 | 0 585762668 |
| ARPP19 | 0 254002399 | -0 00180652 | 0 558181197 | 0 277790056 | 0 | 0 580606749 |
| C15orf48 | 0 006710323 | -0 0018552 | 0 547624708 | -0 48978181 | 0 | 0 569939812 |
| AMDHD1 | 0 019426624 | -0 00189629 | 0 538791604 | 0 479357509 | 0 | 0 56105572 |
| RNF144B | 0 054748625 | -0 00189657 | 0 538730971 | 0 440303081 | 0 | 0 561054405 |
| PRX | 0 031448097 | -0 00190757 | 0 536378501 | -0 4519141 | 0 | 0 558850809 |
| TRAV5 | 9 52E-06 | -0 00191862 | 0 534019666 | -0 49695594 | 0 | 0 556454498 |
| AHR | 0 351887457 | -0 00191936 | 0 533862596 | -0 14791396 | 0 | 0 556352176 |
| CENPA | 0 068854581 | -0 00192962 | 0 531679249 | 0 38257968 | 0 | 0 554199085 |
| CXCR3 | 0 001113628 | -0 00193096 | 0 531393202 | 0 495571039 | 0 | 0 553962025 |
| B4GALT1 | 0 2792256 | -0 00193295 | 0 530970383 | -0 22018208 | 0 | 0 553582317 |
| NEU3 | 0 097485294 | -0 00193935 | 0 529610184 | 0 37404693 | 0 | 0 552225118 |
| FOSB | 0 172260189 | -0 00194449 | 0 528520335 | -0 28520419 | 0 | 0 551149544 |
| SLCO2B1 | 0 056214426 | -0 00195357 | 0 526597268 | -0 42974841 | 0 | 0 549265362 |
| HP1BP3 | 0 635491424 | -0 00196504 | 0 524172287 | 0 049023364 | 6 39E-57 | 0 546856714 |
| PRMT2 | 0 410462394 | -0 00197359 | 0 522368342 | -0 15581377 | 0 | 0 545155255 |
| SYNE2 | 0 572747521 | -0 00198616 | 0 51972408 | 0 133352175 | 0 | 0 54251547 |
| MOGAT1 | 0 004149931 | -0 00199056 | 0 518798945 | -0 49257633 | 0 | 0 541669432 |
| INIP | 0 128619291 | -0 00200391 | 0 515998274 | 0 361046987 | 0 | 0 538923927 |
| TDRKH | 0 027003103 | -0 00201114 | 0 514485306 | -0 46745625 | 0 | 0 537403133 |
| FAM222A-AS1 | 0 012925701 | -0 00203073 | 0 510398047 | 0 480192758 | 0 | 0 533487617 |
| CATSPER1 | 0 001151701 | -0 00203539 | 0 509428055 | 0 49538072 | 0 | 0 532591561 |
| TOE1 | 0 03269498 | -0 00204763 | 0 506884392 | 0 453282285 | 0 | 0 530166852 |
| RAB13 | 0 124269479 | -0 00210281 | 0 495503089 | 0 371294688 | 0 | 0 518549744 |
| NSMCE3 | 0 056785517 | -0 00211393 | 0 493226727 | -0 43277173 | 0 | 0 516224673 |
| DDX58 | 0 116464564 | -0 00211418 | 0 493174461 | 0 322676073 | 0 | 0 516224673 |
| KIF6 | 0 041175687 | -0 00216678 | 0 482482258 | -0 44684715 | 0 | 0 505483075 |
| RAD9B | 0 022044126 | -0 0021773 | 0 480357747 | 0 46036461 | 0 | 0 503313084 |
| ARPC1B | 0 343587596 | -0 0021814 | 0 479531806 | 0 202824083 | 0 | 0 502559117 |

| | | | | | | |
|---|---|---|---|---|---|---|
| GRPEL2 | 0 034817536 | -0 00219832 | 0 47613087 | 0 456194949 | 0 | 0 499105571 |
| RAB32 | 0 093478137 | -0 00222557 | 0 470681271 | 0 401446024 | 0 | 0 493831244 |
| APOBR | 0 001380137 | -0 00222963 | 0 469870531 | -0 49563419 | 0 | 0 493035368 |
| PTGES3 | 0 464811254 | -0 0022393 | 0 467948296 | 0 132247348 | 0 | 0 491181983 |
| XRCC4 | 0 226047477 | -0 00225181 | 0 465464739 | 0 264608851 | 0 | 0 488683678 |
| AGTR1 | 0 068997354 | -0 00225901 | 0 464039226 | -0 43252798 | 0 | 0 487295331 |
| RLN2 | 0 000809046 | -0 00228286 | 0 459334792 | 0 496106809 | 0 | 0 482730625 |
| LIPN | 0 000704346 | -0 0022956 | 0 456832777 | -0 49597899 | 0 | 0 48015457 |
| PLA1A | 0 006291523 | -0 00230495 | 0 45500198 | -0 48963157 | 0 | 0 478443163 |
| SDF4 | 0 210285355 | -0 00231672 | 0 452703515 | -0 29559707 | 0 | 0 476185243 |
| CCL3-AS1 | 0 00110411 | -0 00233152 | 0 449819939 | -0 49581585 | 0 | 0 473257451 |
| MTCH1 | 0 380632388 | -0 00233345 | 0 44944483 | -0 17766973 | 0 | 0 472968112 |
| LAPTM4A | 0 433182311 | -0 00234407 | 0 447385529 | -0 14543221 | 0 | 0 470905908 |
| CP | 0 102130171 | -0 00235494 | 0 44528157 | -0 37641651 | 0 | 0 468848004 |
| BOLA2B | 0 023643182 | -0 00236161 | 0 443994211 | 0 472613565 | 0 | 0 467596711 |
| SLC16A3 | 0 18193067 | -0 00236264 | 0 443795853 | 0 300104438 | 0 | 0 467439904 |
| ZNF674-AS1 | 0 030915079 | -0 0023713 | 0 442126202 | 0 462369385 | 0 | 0 465785129 |
| DMC1 | 0 020787725 | -0 00238264 | 0 439947019 | 0 456889407 | 0 | 0 4636444 |
| LINC00671 | 0 001303992 | -0 0023902 | 0 438497829 | -0 49578368 | 0 | 0 462168691 |
| IKBKE | 0 052473777 | -0 0024067 | 0 4353426 | -0 43218496 | 0 | 0 458996716 |
| WDR49 | 0 024499819 | -0 00240862 | 0 434976611 | -0 4719629 | 0 | 0 458713199 |
| AGAP1 | 0 75036645 | -0 00240951 | 0 434807139 | 0 023164357 | 5 95E-14 | 0 458585655 |
| SAMSN1 | 0 028249986 | -0 00241965 | 0 432876505 | 0 464713479 | 0 | 0 456753346 |
| SLC22A18AS | 0 001446765 | -0 00244278 | 0 428490822 | -0 49545098 | 0 | 0 452226744 |
| THBD | 0 054710552 | -0 00244434 | 0 428196013 | -0 43421414 | 0 | 0 452016568 |
| FASTKD1 | 0 202927795 | -0 00245829 | 0 425565716 | 0 289297702 | 0 | 0 449308249 |
| TMPRSS13 | 0 001722792 | -0 00246832 | 0 42367864 | 0 495300906 | 0 | 0 447497837 |
| SNHG8 | 0 209047991 | -0 00248479 | 0 420592789 | -0 30913419 | 0 | 0 444387506 |
| MCEMP1 | 0 0003141 | -0 00249977 | 0 417796611 | -0 49671943 | 0 | 0 441531869 |
| LRRC69 | 0 116474082 | -0 00250624 | 0 416593369 | -0 37667466 | 0 | 0 440358763 |
| CCL11 | 0 006605623 | -0 00251532 | 0 414907612 | -0 48966758 | 0 | 0 438625902 |
| CXCL12 | 0 191144277 | -0 00254104 | 0 410152648 | -0 29892787 | 0 | 0 433793232 |
| SFTA2 | 0 000723382 | -0 00254143 | 0 410079695 | 0 495907448 | 0 | 0 433786343 |
| SPNS3 | 0 003445585 | -0 00254316 | 0 409762422 | -0 49313269 | 0 | 0 433526073 |
| FMNL1-DT | 0 017037559 | -0 00259189 | 0 400848168 | 0 483556382 | 0 | 0 424570185 |
| LIMS1 | 0 589118806 | -0 00260515 | 0 398442482 | 0 050500251 | 2 70E-60 | 0 422116755 |
| OAS3 | 0 057651672 | -0 00261016 | 0 3975364 | 0 413845935 | 0 | 0 421204057 |
| LAMA5 | 0 081618473 | -0 00261394 | 0 396852336 | -0 39977006 | 0 | 0 420526416 |
| LINC01504 | 0 043345834 | -0 00261444 | 0 39676376 | -0 44546132 | 0 | 0 420479705 |
| CD300LF | 0 002046411 | -0 0026512 | 0 39015727 | -0 49493093 | 0 | 0 413571633 |
| ENHO | 0 004159449 | -0 00265413 | 0 389634034 | 0 49491579 | 0 | 0 413155478 |
| ADGRE1 | 0 002569911 | -0 00265502 | 0 389475743 | 0 493863764 | 0 | 0 413033977 |
| SPTSSB | 0 002132074 | -0 00265591 | 0 389315498 | -0 49447624 | 0 | 0 412910376 |
| RPN2 | 0 340113457 | -0 00267926 | 0 385160449 | 0 228033552 | 0 | 0 408595223 |

| | | | | | | |
|---|---|---|---|---|---|---|
| MREG | 0 067788544 | -0 00269693 | 0 382034735 | 0 427605658 | 0 | 0 405552487 |
| NOTCH4 | 0 02002627 | -0 00272322 | 0 377410702 | -0 47475702 | 0 | 0 400778867 |
| SERAC1 | 0 076069369 | -0 0027325 | 0 37578835 | 0 408077058 | 0 | 0 399100911 |
| TICAM1 | 0 040338086 | -0 0027616 | 0 370725576 | -0 45549892 | 0 | 0 393812578 |
| STAMBPL1 | 0 184709981 | -0 00277767 | 0 367948102 | -0 31189888 | 0 | 0 391037963 |
| LRRC25 | 0 003198112 | -0 00277789 | 0 367909196 | -0 49423573 | 0 | 0 391037963 |
| DUSP23 | 0 038862767 | -0 00277947 | 0 367637137 | 0 459842672 | 0 | 0 390795381 |
| RRM2B | 0 149092917 | -0 0027896 | 0 36589322 | 0 340916145 | 0 | 0 388985367 |
| ITGAX | 0 011364718 | -0 00278982 | 0 365856579 | -0 4854027 | 0 | 0 388985367 |
| GZMH | 0 001227846 | -0 00279394 | 0 365149019 | 0 495218256 | 0 | 0 388281561 |
| SORBS2 | 0 197388209 | -0 00280043 | 0 364035116 | -0 2894896 | 0 | 0 387184226 |
| AURKA | 0 037282747 | -0 00280573 | 0 363128774 | 0 414129877 | 0 | 0 386307206 |
| TLR8 | 0 00120881 | -0 00281514 | 0 361521205 | 0 494998481 | 0 | 0 384683637 |
| PDK1 | 0 157878205 | -0 00281624 | 0 361334031 | 0 324595144 | 0 | 0 384527769 |
| ILDR1 | 0 000847119 | -0 00281797 | 0 361038894 | 0 496049782 | 0 | 0 38425696 |
| NCF4 | 0 004740058 | -0 00282043 | 0 360619157 | -0 49225404 | 0 | 0 383896703 |
| LINC01871 | 0 001380137 | -0 00283396 | 0 358320512 | 0 495471784 | 0 | 0 381621643 |
| HSPA1B | 0 135548533 | -0 00285388 | 0 354951967 | -0 35738078 | 0 | 0 378161904 |
| GHRLOS | 0 022672327 | -0 00286686 | 0 35276756 | 0 463208358 | 0 | 0 376046638 |
| LINC01876 | 0 214026004 | -0 00286745 | 0 352668163 | 0 296742671 | 0 | 0 375983093 |
| GINS3 | 0 037558775 | -0 00287681 | 0 351098687 | 0 458452242 | 0 | 0 374352092 |
| CD69 | 0 007681179 | -0 00290743 | 0 345996268 | -0 48943301 | 0 | 0 369203335 |
| POMGNT1 | 0 133406941 | -0 0029298 | 0 342298371 | -0 36207576 | 0 | 0 365339915 |
| BLK | 0 000942301 | -0 00294653 | 0 339548398 | -0 49616464 | 0 | 0 362527665 |
| TRGV9 | 0 000114218 | -0 0029553 | 0 338113443 | 0 496747289 | 0 | 0 361036388 |
| DMGDH | 0 209914146 | -0 00295619 | 0 337967447 | -0 28170113 | 0 | 0 360921277 |
| FAM83G | 0 007481297 | -0 00295812 | 0 337652341 | -0 48863814 | 0 | 0 360629109 |
| TNXB | 0 269003065 | -0 00297607 | 0 334729122 | -0 2081338 | 0 | 0 35762467 |
| ANKRD12 | 0 628923874 | -0 00298224 | 0 333727634 | -0 04868797 | 3 61E-56 | 0 356635321 |
| MBOAT1 | 0 252289125 | -0 00299544 | 0 331592413 | -0 23487707 | 0 | 0 354554006 |
| RNPEPL1 | 0 05966001 | -0 0030088 | 0 329441396 | -0 43157264 | 0 | 0 35233377 |
| CTSA | 0 179636786 | -0 003013 | 0 328766155 | -0 31211457 | 0 | 0 351651405 |
| CCL14 | 0 01066989 | -0 00302342 | 0 327095952 | -0 48650092 | 0 | 0 349904544 |
| TLR7 | 0 003750167 | -0 00303937 | 0 324551334 | -0 49309422 | 0 | 0 347300431 |
| LILRA2 | 0 001798938 | -0 00304973 | 0 322904138 | 0 495007921 | 0 | 0 34561605 |
| MBNL1 | 0 842654813 | -0 00305387 | 0 322247704 | -0 00945624 | 0 002175912 | 0 344991595 |
| ZFYVE16 | 0 538053721 | -0 00308639 | 0 317121665 | -0 0660853 | 5 24E-102 | 0 339696183 |
| EEF1D | 0 382669281 | -0 00309466 | 0 315826926 | 0 17518096 | 0 | 0 33834763 |
| LARP6 | 0 11900592 | -0 00311909 | 0 312022092 | -0 37921982 | 0 | 0 334499009 |
| APOA1 | 0 006177305 | -0 00313647 | 0 309333383 | 0 49269742 | 0 | 0 331729513 |
| CRISPLD2 | 0 221440673 | -0 0031489 | 0 307419999 | -0 26163291 | 0 | 0 32975244 |
| PI4K2B | 0 188793284 | -0 00316174 | 0 30545092 | 0 301009072 | 0 | 0 327677511 |
| GPATCH3 | 0 033085226 | -0 00316423 | 0 305070577 | 0 458099405 | 0 | 0 327306648 |
| NOP53 | 0 167301213 | -0 00317337 | 0 303675689 | -0 3358599 | 0 | 0 325847084 |

| | | | | | | |
|---|---|---|---|---|---|---|
| LARGE1 | 0 562334621 | -0 00319499 | 0 300393663 | -0 05485625 | 7 88E-71 | 0 322435269 |
| HIF1A-AS1 | 0 013630047 | -0 00322207 | 0 296314547 | -0 48046266 | 0 | 0 318201417 |
| TUBA4A | 0 017618168 | -0 00323504 | 0 294375385 | 0 481143782 | 0 | 0 316190883 |
| CASP8 | 0 129076165 | -0 00324435 | 0 292987682 | 0 349773982 | 0 | 0 314771896 |
| VENTX | 0 004178485 | -0 00325743 | 0 2910468 | -0 49195123 | 0 | 0 312793388 |
| LINC02842 | 0 001960747 | -0 00325953 | 0 290734753 | -0 49468192 | 0 | 0 312493564 |
| HEXIM1 | 0 109792313 | -0 00328206 | 0 287413182 | -0 38258821 | 0 | 0 30898626 |
| PSMD1 | 0 435523786 | -0 00329958 | 0 28484864 | 0 14437316 | 0 | 0 306341146 |
| ETF1 | 0 247453884 | -0 00330479 | 0 284088382 | -0 26212836 | 0 | 0 30562787 |
| TNFRSF18 | 0 004721022 | -0 00330771 | 0 283662563 | -0 49210768 | 0 | 0 305204511 |
| KLB | 0 009299271 | -0 00332308 | 0 281431142 | 0 48313532 | 0 | 0 302838111 |
| MYOT | 0 044830671 | -0 00333275 | 0 280033215 | 0 444728081 | 0 | 0 301402492 |
| CDC42SE2 | 0 348384763 | -0 00334144 | 0 278781556 | 0 174950942 | 0 | 0 300089498 |
| RCAN2 | 0 164893111 | -0 0033522 | 0 277236758 | -0 34132824 | 0 | 0 298460624 |
| JADE1 | 0 214168777 | -0 00335974 | 0 276157847 | 0 323123586 | 0 | 0 29736687 |
| SLBP | 0 127458072 | -0 00339436 | 0 271239546 | 0 352653778 | 0 | 0 292204025 |
| BSPRY | 0 001741829 | -0 0034005 | 0 270373492 | -0 49477046 | 0 | 0 291304241 |
| TNNT2 | 0 010279644 | -0 00342393 | 0 267086129 | -0 48349806 | 0 | 0 287992231 |
| BCL7B | 0 083150901 | -0 00342854 | 0 266442603 | 0 414287882 | 0 | 0 287417815 |
| MYADM | 0 187822429 | -0 00344845 | 0 263675778 | -0 28990077 | 0 | 0 284607207 |
| KCNS3 | 0 076202623 | -0 00345345 | 0 262983937 | -0 41385526 | 0 | 0 283957747 |
| BVES-AS1 | 0 039624222 | -0 00345715 | 0 262472568 | 0 455443682 | 0 | 0 283502774 |
| ABCA9 | 0 515409948 | -0 00346128 | 0 261903599 | -0 08595409 | 1 92E-171 | 0 282920554 |
| LILRB2 | 0 004111858 | -0 00346667 | 0 261161648 | 0 491488591 | 0 | 0 282151317 |
| RUNX1 | 0 463231235 | -0 00346774 | 0 26101372 | 0 088856642 | 4 01E-183 | 0 282023743 |
| OTUD1 | 0 027850222 | -0 00348389 | 0 258799908 | -0 46575924 | 0 | 0 279663708 |
| PLPP1 | 0 504444994 | -0 00348776 | 0 258270975 | -0 07743415 | 1 99E-139 | 0 279124052 |
| TAF9 | 0 101939807 | -0 0034904 | 0 257911596 | -0 39257052 | 0 | 0 278767538 |
| CENPE | 0 086834441 | -0 00350406 | 0 256052747 | 0 33495832 | 0 | 0 276821697 |
| APMAP | 0 130694257 | -0 00350554 | 0 255851382 | -0 36716617 | 0 | 0 276635648 |
| EFHD2 | 0 073927776 | -0 00350854 | 0 255445816 | 0 404779206 | 0 | 0 276228741 |
| LINC01708 | 0 006453332 | -0 00351833 | 0 254121983 | -0 48843696 | 0 | 0 274860105 |
| CTSW | 0 002370029 | -0 00354191 | 0 2509526 | 0 492999059 | 0 | 0 271587502 |
| TYR | 0 00804287 | -0 00354697 | 0 250276169 | 0 489124042 | 0 | 0 270886473 |
| SOWAHD | 0 000447355 | -0 0035509 | 0 24975124 | -0 49660776 | 0 | 0 270349281 |
| CCPG1 | 0 311482744 | -0 0035706 | 0 247134039 | -0 18717176 | 0 | 0 26760819 |
| CLEC12B | 0 000409282 | -0 00357111 | 0 247066904 | 0 496599131 | 0 | 0 267566152 |
| CST7 | 0 004892349 | -0 00357814 | 0 246138259 | -0 49285859 | 0 | 0 266615722 |
| SLAMF7 | 0 002512802 | -0 00358002 | 0 245889551 | 0 493023404 | 0 | 0 266413237 |
| HSPD1 | 0 403418934 | -0 00359625 | 0 243755588 | 0 157769915 | 0 | 0 264252645 |
| NES | 0 156041195 | -0 00359979 | 0 243291572 | -0 33397563 | 0 | 0 263779871 |
| PRKAG2-AS1 | 0 008014315 | -0 00360803 | 0 242213946 | -0 48889297 | 0 | 0 262671768 |
| RFX4 | 0 052559441 | -0 00361651 | 0 241109817 | -0 43239769 | 0 | 0 261534409 |
| CARD18 | 0 004854277 | -0 00362981 | 0 239382845 | -0 49106949 | 0 | 0 25975059 |

| | | | | | | |
|---|---|---|---|---|---|---|
| PPIF | 0 041537378 | -0 00364228 | 0 237772397 | 0 443357551 | 0 | 0 258062381 |
| TIPARP-AS1 | 0 002465211 | -0 00364472 | 0 237458138 | -0 49450026 | 0 | 0 257750908 |
| MILR1 | 0 021035198 | -0 00367191 | 0 233977209 | -0 47576289 | 0 | 0 254118449 |
| FASLG | 0 000609164 | -0 00369402 | 0 23117324 | 0 495844065 | 0 | 0 251130833 |
| PDP1 | 0 146151796 | -0 00370367 | 0 229957144 | -0 34407801 | 0 | 0 249838469 |
| GCA | 0 097275894 | -0 00371995 | 0 227915003 | 0 386769643 | 0 | 0 247733698 |
| CRYBG1 | 0 100616779 | -0 0037329 | 0 226300231 | 0 373608452 | 0 | 0 246035111 |
| KCNC4 | 0 026917439 | -0 00376095 | 0 222830054 | -0 4630714 | 0 | 0 242345955 |
| TRAT1 | 0 001827492 | -0 00376548 | 0 222274127 | 0 494998767 | 0 | 0 241769164 |
| TRGC1 | 0 000428319 | -0 00377299 | 0 221352425 | 0 496117764 | 0 | 0 240794338 |
| CRYBG2 | 0 003369439 | -0 00378595 | 0 219770318 | -0 49336876 | 0 | 0 239155862 |
| XCL2 | 0 000875673 | -0 00378925 | 0 219367816 | 0 495821407 | 0 | 0 238745349 |
| LINC02273 | 0 000409282 | -0 0037918 | 0 219058914 | -0 49650406 | 0 | 0 23843662 |
| SIGLEC9 | 0 002826902 | -0 00380774 | 0 217127045 | 0 493442511 | 0 | 0 236388315 |
| LINC01182 | 0 020854353 | -0 00381126 | 0 216702679 | -0 47436895 | 0 | 0 235953487 |
| IBSP | 0 000609164 | -0 00381507 | 0 216243223 | -0 49645504 | 0 | 0 235507487 |
| SELL | 0 010403381 | -0 00383507 | 0 213845786 | -0 48439414 | 0 | 0 232923315 |
| LRRC38 | 0 00641526 | -0 00384673 | 0 212455431 | -0 48964158 | 0 | 0 231488968 |
| TMEM144 | 0 141335592 | -0 00385023 | 0 212040125 | -0 36210395 | 0 | 0 231063097 |
| CD84 | 0 010660372 | -0 00385562 | 0 211400916 | 0 484526581 | 0 | 0 23039311 |
| LDHD | 0 003892939 | -0 00386385 | 0 210427307 | -0 49183487 | 0 | 0 229358483 |
| MS4A6A | 0 03658792 | -0 00386397 | 0 210413341 | 0 454023327 | 0 | 0 229358483 |
| IL6ST | 0 581218709 | -0 00386909 | 0 209810345 | -0 02505903 | 4 52E-16 | 0 228765173 |
| LINC00482 | 0 004958977 | -0 00390741 | 0 205331113 | -0 49119019 | 0 | 0 224036372 |
| PLEK | 0 009679998 | -0 00393435 | 0 202224273 | -0 48741277 | 0 | 0 220799466 |
| PHEX | 0 061363766 | -0 00394671 | 0 200809611 | 0 410103182 | 0 | 0 219354246 |
| SIRT7 | 0 111924388 | -0 00395159 | 0 200254087 | 0 374175755 | 0 | 0 218799991 |
| ANLN | 0 110220632 | -0 00396582 | 0 198638104 | 0 288811695 | 0 | 0 217084548 |
| ARHGAP15-AS | 0 002303402 | -0 00398768 | 0 19617449 | -0 49463104 | 0 | 0 214491372 |
| NCR1 | 0 001303992 | -0 00402945 | 0 191529622 | 0 494778771 | 0 | 0 209582553 |
| RAPSN | 0 006310559 | -0 00403559 | 0 1908543 | -0 49038033 | 0 | 0 20892448 |
| IGFBP4 | 0 332270469 | -0 00404557 | 0 189759819 | -0 16787792 | 0 | 0 207766225 |
| KRTAP5-AS1 | 0 009908435 | -0 00404677 | 0 189627919 | -0 48482398 | 0 | 0 207645867 |
| GPR42 | 0 000133255 | -0 00404703 | 0 189599978 | 0 496672108 | 0 | 0 207639332 |
| ACP5 | 0 009108907 | -0 00404963 | 0 189315636 | -0 48723698 | 0 | 0 207351966 |
| MAP3K11 | 0 07974339 | -0 00404989 | 0 189287239 | -0 40806717 | 0 | 0 207344894 |
| IAH1 | 0 213683349 | -0 00405871 | 0 188324634 | -0 30397461 | 0 | 0 206338294 |
| MSL3 | 0 124393215 | -0 00408376 | 0 185612898 | 0 367679383 | 0 | 0 203461534 |
| GNB4 | 0 290114409 | -0 00409319 | 0 184598873 | 0 226891214 | 0 | 0 20242044 |
| CSF2 | 8 57E-05 | -0 00409845 | 0 184035799 | -0 49690932 | 0 | 0 201826425 |
| BRI3 | 0 330481049 | -0 00409995 | 0 183875051 | -0 21109992 | 0 | 0 201673542 |
| SOX13 | 0 129000019 | -0 00410281 | 0 183569565 | -0 35483143 | 0 | 0 201361856 |
| RASAL3 | 0 005739468 | -0 00410586 | 0 183243469 | 0 488941872 | 0 | 0 201027488 |
| CD33 | 0 004216558 | -0 00412437 | 0 181277302 | -0 4925888 | 0 | 0 198939787 |

| | | | | | | |
|---|---|---|---|---|---|---|
| CCDC60 | 0 019597952 | -0 00412648 | 0 181053982 | -0 4729678 | 0 | 0 198717786 |
| HCCS | 0 025575375 | -0 00413315 | 0 180350045 | -0 47064681 | 0 | 0 197991162 |
| HIPK1 | 0 224210466 | -0 00415562 | 0 177992017 | 0 280696806 | 0 | 0 195425184 |
| RELL2 | 0 010765072 | -0 00416117 | 0 177413325 | -0 48528794 | 0 | 0 194812447 |
| PGM2 | 0 082560774 | -0 00416177 | 0 17735174 | 0 394398409 | 0 | 0 194767454 |
| CD300LG | 0 002579429 | -0 00418149 | 0 175306444 | 0 492204895 | 0 | 0 192678053 |
| IDH1-AS1 | 0 00231292 | -0 00419509 | 0 173907121 | -0 49472649 | 0 | 0 191184539 |
| IL32 | 0 036949611 | -0 00422515 | 0 170843479 | 0 456031758 | 0 | 0 187903966 |
| TWSG1 | 0 198768346 | -0 0042386 | 0 169485453 | 0 301668892 | 0 | 0 186497151 |
| PECR | 0 09225029 | -0 00424221 | 0 169123071 | 0 40358808 | 0 | 0 186120068 |
| DEGS1 | 0 166530239 | -0 00425538 | 0 167803274 | 0 34832972 | 0 | 0 184732169 |
| STAB1 | 0 048428547 | -0 00425837 | 0 167504665 | -0 44443369 | 0 | 0 184446409 |
| PIGR | 0 000304582 | -0 00426571 | 0 166773424 | -0 49674695 | 0 | 0 183705427 |
| ANKH | 0 555833698 | -0 00426931 | 0 166415728 | -0 03684652 | 6 73E-33 | 0 183332785 |
| AQP9 | 0 003074375 | -0 00428247 | 0 16511367 | -0 4939042 | 0 | 0 181962004 |
| VHL | 0 177580857 | -0 00433223 | 0 160257314 | 0 315732437 | 0 | 0 17675439 |
| FGF18 | 0 017722868 | -0 00434046 | 0 159464712 | 0 479194654 | 0 | 0 175982894 |
| ESRRA | 0 094287183 | -0 00435119 | 0 15843551 | -0 40254621 | 0 | 0 174949237 |
| ABHD11 | 0 035512364 | -0 00435309 | 0 15825356 | 0 459620798 | 0 | 0 174768744 |
| CCR5 | 0 001389656 | -0 00435579 | 0 157996006 | -0 49591122 | 0 | 0 174525104 |
| HCAR2 | 0 001008928 | -0 00437432 | 0 156234875 | 0 49523456 | 0 | 0 172620083 |
| IL18 | 0 096933239 | -0 00439385 | 0 154394483 | -0 37609671 | 0 | 0 170706431 |
| UTF1 | 0 000228437 | -0 00439839 | 0 153968947 | 0 496509834 | 0 | 0 170255856 |
| INSYN2B | 0 050579658 | -0 00440103 | 0 153722215 | 0 400807136 | 0 | 0 170002918 |
| FAM3C | 0 413432069 | -0 00442652 | 0 151352433 | 0 1626008 | 0 | 0 167440938 |
| ARHGAP5 | 0 547295882 | -0 00443622 | 0 150458461 | -0 06253933 | 1 55E-91 | 0 16649092 |
| ARL6IP6 | 0 225257467 | -0 00443723 | 0 150365819 | 0 238556338 | 0 | 0 166427382 |
| CLEC7A | 0 018274923 | -0 00444157 | 0 149967003 | -0 47798006 | 0 | 0 166005409 |
| NXPE3 | 0 117768556 | -0 00444197 | 0 149930812 | 0 37295564 | 0 | 0 165984791 |
| IGHGP | 0 013887038 | -0 00445038 | 0 14916051 | -0 48277784 | 0 | 0 165170708 |
| MPEG1 | 0 008547334 | -0 00447027 | 0 147351644 | 0 485341966 | 0 | 0 163225066 |
| EEA1 | 0 569825436 | -0 00447995 | 0 146477239 | 0 098147505 | 3 74E-223 | 0 162294513 |
| LINC00528 | 0 001113628 | -0 00448194 | 0 146298187 | 0 494961477 | 0 | 0 162115135 |
| KIF14 | 0 071757629 | -0 00448449 | 0 146068877 | 0 354768999 | 0 | 0 161880015 |
| IDI1 | 0 102539453 | -0 00449501 | 0 145124702 | -0 37936639 | 0 | 0 16087137 |
| EIF5AL1 | 0 004083303 | -0 00450832 | 0 143937577 | -0 4932174 | 0 | 0 159686558 |
| CLEC17A | 0 001875083 | -0 00451358 | 0 143470251 | -0 49490398 | 0 | 0 159186788 |
| LINC02712 | 0 002227256 | -0 00453797 | 0 141319493 | -0 49485846 | 0 | 0 156892529 |
| POTEG | 0 001837011 | -0 00455647 | 0 139705107 | -0 49515439 | 0 | 0 155136693 |
| BCL2L1-AS1 | 0 055643334 | -0 00455666 | 0 139688096 | -0 42282836 | 0 | 0 155136034 |
| CLEC4E | 0 00178942 | -0 00455692 | 0 139665484 | -0 49520691 | 0 | 0 155129152 |
| PKD2L2 | 0 07743047 | -0 00456739 | 0 13875872 | 0 388886699 | 0 | 0 154176356 |
| ITK | 0 00893758 | -0 00456771 | 0 138731013 | 0 486625892 | 0 | 0 154163696 |
| PLLP | 0 01808456 | -0 00456786 | 0 138717572 | -0 47586631 | 0 | 0 154163696 |

| | | | | | | |
|---|---|---|---|---|---|---|
| OSTF1 | 0 127962536 | -0 00457084 | 0 138460233 | -0 36755248 | 0 | 0 153917092 |
| MMP2-AS1 | 0 001475319 | -0 00457173 | 0 138383556 | -0 49573967 | 0 | 0 153849954 |
| CD93 | 0 02160629 | -0 00459476 | 0 136408235 | 0 472056417 | 0 | 0 151689553 |
| ZNF208 | 0 038091793 | -0 00459937 | 0 136015333 | 0 46098324 | 0 | 0 151270436 |
| SHISA6 | 0 194152025 | -0 00461215 | 0 134930406 | -0 27003112 | 0 | 0 150081489 |
| C16orf54 | 0 000704346 | -0 00461469 | 0 134715715 | 0 495991572 | 0 | 0 14986033 |
| FAM72A | 0 008880471 | -0 00461575 | 0 13462618 | 0 483458267 | 0 | 0 149778362 |
| KIFC2 | 0 05067484 | -0 00463363 | 0 133122553 | 0 450550543 | 0 | 0 148122939 |
| PHLDA2 | 0 024404637 | -0 00470373 | 0 127353162 | 0 467695859 | 0 | 0 141837052 |
| ERMAP | 0 067464925 | -0 0047071 | 0 127081046 | 0 424934393 | 0 | 0 14155067 |
| METRNL | 0 098627477 | -0 00474336 | 0 12417862 | -0 39890047 | 0 | 0 138383014 |
| PGRMC1 | 0 163912737 | -0 00474442 | 0 124094659 | -0 33201268 | 0 | 0 138305758 |
| NCMAP | 0 003321848 | -0 00475177 | 0 123512985 | -0 49348449 | 0 | 0 13768059 |
| METTL7B | 0 00310293 | -0 00478825 | 0 120658119 | 0 490969565 | 0 | 0 134570899 |
| PCAT1 | 0 129970874 | -0 00479114 | 0 120434099 | 0 329292349 | 0 | 0 134336902 |
| MEPE | 0 000704346 | -0 00479115 | 0 120433 | -0 49593327 | 0 | 0 134336902 |
| FLVCR2 | 0 095258038 | -0 00480329 | 0 119495555 | -0 36835882 | 0 | 0 133384478 |
| C10orf67 | 0 004987531 | -0 00481243 | 0 118794022 | -0 49166198 | 0 | 0 13263274 |
| SPHKAP | 0 018332033 | -0 00482812 | 0 117596437 | -0 47637906 | 0 | 0 131326676 |
| SLC30A2 | 0 001951229 | -0 00483031 | 0 117429953 | -0 49451514 | 0 | 0 131171756 |
| PTP4A2 | 0 318031258 | -0 00486387 | 0 114904031 | 0 186962977 | 0 | 0 128426149 |
| LINC00589 | 0 003169557 | -0 00487015 | 0 114435874 | -0 49344498 | 0 | 0 127918028 |
| LOXL3 | 0 200386438 | -0 00488068 | 0 113654742 | -0 30479952 | 0 | 0 12707493 |
| TWF2 | 0 054329824 | -0 00489602 | 0 112523875 | -0 44107524 | 0 | 0 125870102 |
| MX1 | 0 125116598 | -0 00489958 | 0 112262971 | -0 38102932 | 0 | 0 125593119 |
| GARRE1 | 0 175125164 | -0 00492273 | 0 110576023 | -0 29589359 | 0 | 0 123793795 |
| CD86 | 0 013868002 | -0 00493889 | 0 10941105 | -0 4837662 | 0 | 0 122533115 |
| CXADR | 0 024595001 | -0 00495122 | 0 10852807 | -0 47053766 | 0 | 0 121558642 |
| EXOC6 | 0 282699739 | -0 00495316 | 0 10839015 | -0 23402994 | 0 | 0 121418554 |
| VPS35 | 0 232215263 | -0 00495507 | 0 108254268 | 0 299397248 | 0 | 0 121280715 |
| CD27 | 0 011764482 | -0 00495669 | 0 10813884 | 0 479952651 | 0 | 0 121165762 |
| SCN1A-AS1 | 0 071348347 | -0 00496178 | 0 107776683 | -0 41866843 | 0 | 0 120774298 |
| LY96 | 0 086958177 | -0 00496318 | 0 107677431 | 0 397748553 | 0 | 0 120677387 |
| RBM8A | 0 178104357 | -0 00496324 | 0 107673212 | 0 332976079 | 0 | 0 120677387 |
| CMTM7 | 0 192353087 | -0 00497249 | 0 107019389 | 0 288269802 | 0 | 0 119997786 |
| C6orf226 | 0 03931964 | -0 00498088 | 0 106428478 | -0 45049218 | 0 | 0 119390921 |
| HEPH | 0 085968285 | -0 00498455 | 0 106170841 | -0 38182915 | 0 | 0 119130188 |
| PI4K2A | 0 113732843 | -0 00500097 | 0 105024149 | -0 37891415 | 0 | 0 117899526 |
| CSF3 | 0 000437837 | -0 00501743 | 0 103885343 | -0 49651256 | 0 | 0 116662685 |
| CEP55 | 0 044031144 | -0 00502708 | 0 103221664 | 0 389434179 | 0 | 0 115931154 |
| CD180 | 0 001646647 | -0 00503737 | 0 102518276 | -0 4954254 | 0 | 0 115182226 |
| DRAM1 | 0 177485675 | -0 00504157 | 0 102232053 | -0 31993779 | 0 | 0 114887966 |
| IFFO2 | 0 060326284 | -0 00505107 | 0 101587166 | -0 43509205 | 0 | 0 114190403 |
| ZBP1 | 0 003445585 | -0 00505679 | 0 101200465 | 0 491570863 | 0 | 0 113796335 |

| | | | | | | |
|---|---|---|---|---|---|---|
| FOLR2 | 0 012706783 | -0 00505685 | 0 101196489 | 0 478545656 | 0 | 0 113796335 |
| GPR176 | 0 161990063 | -0 00505821 | 0 101104566 | 0 292048036 | 0 | 0 113719737 |
| PTTG2 | 0 000323618 | -0 00506035 | 0 100960486 | 0 496574904 | 0 | 0 113580547 |
| DEFB124 | 0 010193981 | -0 00510961 | 0 097684978 | -0 48681015 | 0 | 0 1100397 |
| PARD6A | 0 003017266 | -0 00511651 | 0 097233198 | -0 49382942 | 0 | 0 109543838 |
| MNDA | 0 009794217 | -0 00513225 | 0 096207905 | -0 48679498 | 0 | 0 108401658 |
| PXT1 | 0 007909615 | -0 00513508 | 0 096024715 | -0 48809686 | 0 | 0 108208151 |
| FUT11 | 0 105632864 | -0 00515009 | 0 095057143 | -0 3791628 | 0 | 0 10715615 |
| DEPDC1 | 0 041899069 | -0 00516585 | 0 094049924 | 0 393634936 | 0 | 0 106071343 |
| ARPC5L | 0 089547125 | -0 0051717 | 0 093678034 | 0 40348641 | 0 | 0 105664529 |
| SULT2B1 | 0 005672841 | -0 00521993 | 0 090657192 | -0 49039606 | 0 | 0 102281574 |
| PRSS12 | 0 097437703 | -0 00522434 | 0 090384708 | 0 355965744 | 0 | 0 101998506 |
| RSRP1 | 0 541366051 | -0 00522618 | 0 090271759 | -0 09209473 | 1 29E-196 | 0 10188321 |
| KLHL2 | 0 203022977 | -0 00524288 | 0 08924767 | 0 315220989 | 0 | 0 100763499 |
| CD48 | 0 003978603 | -0 00524518 | 0 0891073 | -0 49338252 | 0 | 0 100617037 |
| RHEB | 0 422169766 | -0 00527003 | 0 087603079 | 0 155878552 | 0 | 0 098965822 |
| DHRS9 | 0 003169557 | -0 00527221 | 0 087471881 | 0 49269621 | 0 | 0 098829421 |
| PRDX2 | 0 117987474 | -0 00527923 | 0 087051035 | -0 37499989 | 0 | 0 09840099 |
| REEP3 | 0 429812872 | -0 00529462 | 0 086134673 | -0 15370567 | 0 | 0 097376798 |
| CD52 | 0 004149931 | -0 00531084 | 0 085176756 | 0 491076031 | 0 | 0 096316901 |
| KCNG2 | 0 019969161 | -0 00532473 | 0 084363409 | -0 4743892 | 0 | 0 095420014 |
| CHIT1 | 0 004197521 | -0 00533619 | 0 083697169 | 0 490412945 | 0 | 0 094677789 |
| IL4I1 | 0 010155908 | -0 00535212 | 0 082778151 | 0 482354007 | 0 | 0 093660624 |
| CPD | 0 271439721 | -0 00536587 | 0 081991847 | -0 2330161 | 0 | 0 092793169 |
| TGOLN2 | 0 274790124 | -0 00537302 | 0 081584864 | -0 24469773 | 0 | 0 092343629 |
| LILRA6 | 0 001741829 | -0 00537338 | 0 081564501 | 0 493912968 | 0 | 0 09233164 |
| DEDD2 | 0 032428471 | -0 00538025 | 0 081175383 | -0 46218769 | 0 | 0 091935207 |
| SLC10A3 | 0 028116731 | -0 00540998 | 0 079508556 | -0 46534948 | 0 | 0 090058234 |
| YWHAB | 0 300632008 | -0 00543286 | 0 078245203 | -0 22335566 | 0 | 0 088659132 |
| NFATC2 | 0 231767908 | -0 005439 | 0 077908785 | -0 226218 | 0 | 0 088299115 |
| LINC00239 | 0 000247473 | -0 00545255 | 0 077170824 | -0 49675584 | 0 | 0 08748372 |
| CCR5AS | 0 001589538 | -0 00549242 | 0 075032622 | 0 49474605 | 0 | 0 085131261 |
| ELOVL5 | 0 418971655 | -0 00549455 | 0 074919854 | -0 15144968 | 0 | 0 085013523 |
| VDR | 0 027450458 | -0 00549679 | 0 07480112 | 0 457274803 | 0 | 0 084899182 |
| MOAP1 | 0 03348499 | -0 00550249 | 0 074500034 | -0 45644923 | 0 | 0 084567607 |
| SHTN1 | 0 319658868 | -0 00550268 | 0 074490133 | 0 180617475 | 0 | 0 084566525 |
| SP140 | 0 043555234 | -0 00552681 | 0 073227371 | 0 433560852 | 0 | 0 08316292 |
| AGMAT | 0 019597952 | -0 0055564 | 0 071702349 | 0 469210618 | 0 | 0 081460351 |
| UAP1L1 | 0 066674916 | -0 00556701 | 0 071162027 | -0 42575174 | 0 | 0 08086594 |
| ATAD3C | 0 012135691 | -0 00557348 | 0 070834097 | -0 48249404 | 0 | 0 080502972 |
| CAMK2N1 | 0 058022882 | -0 00563156 | 0 067945913 | -0 43927598 | 0 | 0 077267013 |
| C17orf49 | 0 058641564 | -0 00563449 | 0 067802879 | -0 43273731 | 0 | 0 077113636 |
| PLS3-AS1 | 0 030724715 | -0 00563944 | 0 067561566 | -0 46485663 | 0 | 0 076848435 |
| IFRD1 | 0 294854467 | -0 00565646 | 0 066737477 | -0 21959922 | 0 | 0 075947634 |

| | | | | | | |
|---|---|---|---|---|---|---|
| C9orf72 | 0 117435419 | -0 00570317 | 0 06451906 | -0 38184421 | 0 | 0 073458448 |
| MTFP1 | 0 021406408 | -0 00571565 | 0 063936652 | 0 468480141 | 0 | 0 072804116 |
| ACAT1 | 0 143838876 | -0 00571706 | 0 063870982 | -0 35322163 | 0 | 0 072738103 |
| CLECL1 | 0 001570501 | -0 00574167 | 0 062736496 | 0 494114579 | 0 | 0 071463343 |
| SLC28A3 | 0 003369439 | -0 00574724 | 0 062481909 | -0 49325874 | 0 | 0 071190504 |
| KIR2DL3 | 8 57E-05 | -0 00575609 | 0 062079173 | 0 496649979 | 0 | 0 070757229 |
| C22orf46 | 8 57E-05 | -0 00575609 | 0 062079173 | -0 49695324 | 0 | 0 070757229 |
| XACT | 0 023785955 | -0 0057626 | 0 061784574 | -0 46983229 | 0 | 0 070446939 |
| HMMR | 0 046134663 | -0 00577216 | 0 061353464 | 0 38715935 | 0 | 0 06997227 |
| PSAT1 | 0 070653519 | -0 00577605 | 0 061179091 | 0 437758643 | 0 | 0 069781824 |
| KAT2B | 0 231815499 | -0 00578076 | 0 060968261 | -0 2671286 | 0 | 0 069549743 |
| PJA2 | 0 509556262 | -0 00580257 | 0 059999581 | -0 07217513 | 2 40E-121 | 0 068469513 |
| SLC24A4 | 0 043954998 | -0 00580532 | 0 059878465 | -0 43435956 | 0 | 0 068339552 |
| IL5RA | 0 013544383 | -0 00580898 | 0 059717392 | -0 48237811 | 0 | 0 068172186 |
| ADGRG3 | 0 001227846 | -0 00581152 | 0 059605913 | -0 49556686 | 0 | 0 068053145 |
| PGBD4 | 0 022329672 | -0 0058298 | 0 058808314 | 0 473854082 | 0 | 0 067176289 |
| PTPRB | 0 077021187 | -0 0058397 | 0 058380305 | -0 41835639 | 0 | 0 066702199 |
| ACAT2 | 0 063943195 | -0 00584322 | 0 058228925 | 0 442492531 | 0 | 0 066537284 |
| SYT14 | 0 12862881 | -0 00588159 | 0 056597035 | -0 34128145 | 0 | 0 064711673 |
| SLC18A2 | 0 027850222 | -0 00588716 | 0 056363389 | 0 456475097 | 0 | 0 064460127 |
| THEM6 | 0 017037559 | -0 00591432 | 0 055235697 | -0 4781228 | 0 | 0 063185732 |
| NLRP2 | 0 003045821 | -0 005934 | 0 054430217 | -0 49379373 | 0 | 0 062271858 |
| TANC1 | 0 502398584 | -0 00593753 | 0 054286611 | 0 112554192 | 3 03E-293 | 0 062122605 |
| EIF3L | 0 235080238 | -0 00594006 | 0 054184282 | 0 265104978 | 0 | 0 062020526 |
| KLRC1 | 0 001922674 | -0 00594803 | 0 053862155 | 0 493804467 | 0 | 0 061671013 |
| ARL4D | 0 036016828 | -0 00595202 | 0 053701446 | -0 45510187 | 0 | 0 061497657 |
| ST18 | 0 008661552 | -0 00599102 | 0 05215164 | -0 48729525 | 0 | 0 059744575 |
| LINC00852 | 0 008785289 | -0 0060146 | 0 051233128 | 0 481983402 | 0 | 0 058699449 |
| RNF43 | 0 030724715 | -0 00602412 | 0 050865893 | -0 45779159 | 0 | 0 058292831 |
| SLCO4A1 | 0 010022653 | -0 00603696 | 0 05037433 | 0 479469101 | 0 | 0 0577435 |
| TOP2A | 0 122042223 | -0 00604521 | 0 050060687 | 0 28319099 | 0 | 0 057404865 |
| GLUD1 | 0 336287145 | -0 00605558 | 0 049668649 | -0 17612912 | 0 | 0 056962225 |
| SIRPD | 0 000770973 | -0 00606636 | 0 04926408 | 0 495824128 | 0 | 0 056525687 |
| NR4A2 | 0 204536369 | -0 0060719 | 0 049056988 | -0 27832131 | 0 | 0 056301741 |
| ARHGAP10 | 0 722040319 | -0 00609041 | 0 048371018 | -0 01409301 | 4 92E-06 | 0 055534701 |
| IL13RA1 | 0 191163313 | -0 00609507 | 0 048199431 | -0 30189477 | 0 | 0 055344426 |
| IL2RG | 0 004825722 | -0 00609601 | 0 048164868 | 0 490108676 | 0 | 0 05531146 |
| GLRA2 | 0 048228665 | -0 006103 | 0 047908842 | 0 404472619 | 0 | 0 05503082 |
| CCDC186 | 0 272591422 | -0 00611052 | 0 047634447 | -0 2504017 | 0 | 0 054748909 |
| LMOD3 | 0 012107137 | -0 00611766 | 0 047375483 | 0 480370973 | 0 | 0 054464515 |
| FCER1A | 0 002341475 | -0 00612301 | 0 047181978 | -0 49457462 | 0 | 0 054255256 |
| PIK3IP1 | 0 04484019 | -0 00614478 | 0 046402055 | -0 44381699 | 0 | 0 053377896 |
| MMADHC-DT | 0 078306143 | -0 00616303 | 0 045756349 | -0 39982884 | 0 | 0 052673589 |
| PSMB8 | 0 069558927 | -0 00617521 | 0 04532953 | 0 418773922 | 0 | 0 052201323 |

| | | | | | | |
|---|---|---|---|---|---|---|
| IL1RN | 0 002427138 | -0 00617667 | 0 045278887 | 0 493525876 | 0 | 0 052149358 |
| CNPPD1 | 0 05088424 | -0 00618001 | 0 045162547 | -0 43780534 | 0 | 0 052021705 |
| CRIP2 | 0 10281548 | -0 00618828 | 0 044875513 | -0 37542173 | 0 | 0 051716292 |
| TMX1 | 0 154061411 | -0 00620227 | 0 044393833 | -0 34911799 | 0 | 0 051173666 |
| HCFC1R1 | 0 104852373 | -0 00625223 | 0 042708576 | -0 38433602 | 0 | 0 049267096 |
| VAMP1 | 0 045554054 | -0 00625586 | 0 042588514 | 0 431865361 | 0 | 0 049134593 |
| UHRF2 | 0 397346329 | -0 00626307 | 0 042350346 | 0 130770228 | 0 | 0 048865784 |
| RNF113A | 0 017837087 | -0 00629461 | 0 041321888 | -0 47678153 | 0 | 0 047708227 |
| RGS9 | 0 063353068 | -0 00630176 | 0 041091673 | 0 451638443 | 0 | 0 047454028 |
| LILRB4 | 0 006005977 | -0 00631854 | 0 040556093 | -0 49144152 | 0 | 0 046858427 |
| INPP5D | 0 064961642 | -0 00633165 | 0 040141542 | -0 42434173 | 0 | 0 046385128 |
| KLF4 | 0 097475776 | -0 00634512 | 0 039719165 | -0 38070019 | 0 | 0 045913897 |
| PTPN9 | 0 5352173 | -0 0063895 | 0 038354767 | -0 07859424 | 1 37E-143 | 0 044352979 |
| RPSA | 0 348460909 | -0 00641154 | 0 037692218 | 0 186449921 | 0 | 0 043608161 |
| P2RY13 | 0 00420704 | -0 0064224 | 0 03736933 | -0 4929043 | 0 | 0 043239888 |
| DPYSL3 | 0 424844378 | -0 00642339 | 0 037340097 | 0 124694524 | 0 | 0 043211354 |
| PRKAG2 | 0 208296054 | -0 00644173 | 0 036800269 | 0 268615245 | 0 | 0 042597078 |
| PKHD1L1 | 0 003198112 | -0 00644541 | 0 0366929 | -0 49392291 | 0 | 0 042477999 |
| WNT10A | 0 002569911 | -0 00644677 | 0 036653244 | 0 49276079 | 0 | 0 042437289 |
| CDK5R1 | 0 009032762 | -0 00645356 | 0 036455731 | -0 48681913 | 0 | 0 042218953 |
| TXLNG | 0 242561535 | -0 00646047 | 0 036255861 | 0 272547396 | 0 | 0 041992632 |
| SDS | 0 008680589 | -0 00646477 | 0 036132011 | -0 48853173 | 0 | 0 041854315 |
| ATP6V0D2 | 0 004930422 | -0 00647005 | 0 035980033 | -0 49098826 | 0 | 0 041683377 |
| NOD2 | 0 008385525 | -0 00647071 | 0 035961158 | 0 483817978 | 0 | 0 041666619 |
| ARHGAP11A | 0 07153871 | -0 00647469 | 0 035847397 | 0 343990379 | 0 | 0 041539902 |
| LINC02853 | 0 041880033 | -0 00651318 | 0 034761132 | -0 44650173 | 0 | 0 040310799 |
| LYST | 0 476651882 | -0 00652389 | 0 034463836 | -0 08094661 | 3 22E-152 | 0 039990576 |
| SLC17A5 | 0 178865813 | -0 00654408 | 0 033909497 | -0 295984 | 0 | 0 039361841 |
| NME3 | 0 227741714 | -0 00654811 | 0 03379964 | -0 27779188 | 0 | 0 03923914 |
| FCGR1B | 0 002017856 | -0 00656759 | 0 033273433 | 0 49380327 | 0 | 0 038637742 |
| ARF1 | 0 298376197 | -0 00657738 | 0 033011685 | -0 20846779 | 0 | 0 038343218 |
| MELK | 0 10639432 | -0 00658147 | 0 032902709 | 0 303758438 | 0 | 0 038226039 |
| MIR223HG | 0 001332546 | -0 00659268 | 0 032606076 | -0 49533467 | 0 | 0 037900051 |
| SNRPB | 0 158573033 | -0 00661156 | 0 032111355 | 0 341165835 | 0 | 0 037334191 |
| LINC01608 | 0 005948868 | -0 00662077 | 0 031872501 | -0 49025591 | 0 | 0 037061048 |
| SORCS3 | 0 052902096 | -0 00662405 | 0 031787887 | -0 43259247 | 0 | 0 036967208 |
| APH1B | 0 133454532 | -0 00662613 | 0 031734238 | -0 35254326 | 0 | 0 03690936 |
| KMO | 0 010631817 | -0 00664118 | 0 03134839 | 0 487232493 | 0 | 0 036469566 |
| HLA-DQA1 | 0 032723535 | -0 00664367 | 0 031284964 | -0 46625277 | 0 | 0 03640026 |
| CXCL11 | 0 004130894 | -0 00664733 | 0 031191926 | 0 491719511 | 0 | 0 036296479 |
| OTOGL | 0 055700444 | -0 00664944 | 0 031138359 | 0 44251057 | 0 | 0 036238607 |
| TNFSF13 | 0 017694314 | -0 00669984 | 0 029883209 | -0 4759075 | 0 | 0 034803589 |
| MZT1 | 0 066199006 | -0 00673179 | 0 029110056 | 0 416089081 | 0 | 0 033915674 |
| BCL6B | 0 004502104 | -0 00673598 | 0 02901002 | 0 490548468 | 0 | 0 03380329 |

| | | | | | | |
|---|---|---|---|---|---|---|
| HSD17B3 | 0 02554682 | -0 00674305 | 0 028841868 | -0 464199 | 0 | 0 0336115 |
| FOXD1 | 0 082037273 | -0 006757 | 0 028512497 | -0 41048753 | 0 | 0 033239957 |
| RABGGTB | 0 133920923 | -0 00676989 | 0 028210779 | -0 36485777 | 0 | 0 032904451 |
| LINC02646 | 0 0469913 | -0 00677312 | 0 028135841 | -0 44184468 | 0 | 0 032825148 |
| RRAD | 0 014334393 | -0 00680559 | 0 027390065 | -0 48196199 | 0 | 0 031962968 |
| ZNF367 | 0 071605338 | -0 00680716 | 0 027354466 | 0 345708099 | 0 | 0 031925367 |
| SLC30A1 | 0 046800937 | -0 00684281 | 0 02655634 | -0 44335969 | 0 | 0 031001533 |
| ZBTB32 | 0 030924597 | -0 0068445 | 0 026518983 | -0 46030665 | 0 | 0 030961748 |
| ACSL1 | 0 220108127 | -0 00684742 | 0 026454541 | -0 26822545 | 0 | 0 030890327 |
| HMGCS1 | 0 218090271 | -0 0068501 | 0 026395582 | 0 287515496 | 0 | 0 03082529 |
| TREM1 | 0 004425958 | -0 00686578 | 0 026052842 | -0 49174019 | 0 | 0 030428792 |
| SPTAN1 | 0 386495593 | -0 00686597 | 0 026048811 | -0 14873829 | 0 | 0 030427845 |
| HNRNPH2 | 0 08268451 | -0 00688398 | 0 025659845 | -0 40315328 | 0 | 0 029984609 |
| SORL1 | 0 097970722 | -0 00691784 | 0 024942097 | -0 38266414 | 0 | 0 029163921 |
| DIRC3-AS1 | 0 107260475 | -0 00692923 | 0 02470472 | -0 38415662 | 0 | 0 028897092 |
| LAMB2 | 0 221326455 | -0 00692974 | 0 024694021 | -0 26485637 | 0 | 0 028888153 |
| PRF1 | 0 002084483 | -0 00693004 | 0 024687843 | -0 49544593 | 0 | 0 028884501 |
| FAM72B | 0 009432526 | -0 0069357 | 0 02457057 | 0 476528175 | 0 | 0 028750853 |
| LINC02725 | 0 006091641 | -0 00694378 | 0 024404074 | 0 489573544 | 0 | 0 028563104 |
| SFRP4 | 0 213207439 | -0 00695326 | 0 024210126 | -0 27938071 | 0 | 0 028343123 |
| ARPC3 | 0 34409206 | -0 00697929 | 0 023683998 | 0 20760415 | 0 | 0 027747803 |
| ZBTB7B | 0 026508157 | -0 00698647 | 0 023540661 | -0 46659354 | 0 | 0 027583292 |
| ZC3H12A | 0 012468828 | -0 00698699 | 0 023530282 | -0 48465101 | 0 | 0 027574549 |
| PPP1R18 | 0 088395424 | -0 0070008 | 0 023256814 | 0 404083904 | 0 | 0 027267605 |
| SNHG12 | 0 111029678 | -0 00701177 | 0 023041539 | 0 360394939 | 0 | 0 02702191 |
| PYCARD | 0 033627763 | -0 00702851 | 0 022716387 | 0 456071964 | 0 | 0 026647202 |
| BUB1 | 0 072795111 | -0 00704367 | 0 022425289 | 0 337475535 | 0 | 0 026315532 |
| RAB4B | 0 137661571 | -0 00706597 | 0 022003248 | -0 3451114 | 0 | 0 025829899 |
| PHPT1 | 0 19885401 | -0 00707991 | 0 021742786 | -0 31501927 | 0 | 0 025533655 |
| TM4SF18 | 0 016038149 | -0 00708615 | 0 021627082 | -0 47865408 | 0 | 0 025400935 |
| CRADD | 0 415430888 | -0 00709095 | 0 021538441 | -0 1586476 | 0 | 0 025299971 |
| PSMA4 | 0 203013459 | -0 00711102 | 0 021171281 | 0 302410391 | 0 | 0 024877966 |
| SLC16A13 | 0 02034037 | -0 00713265 | 0 020781634 | -0 47441977 | 0 | 0 024432252 |
| TNF | 0 002027374 | -0 00713342 | 0 020767993 | -0 49495262 | 0 | 0 024419253 |
| OBI1-AS1 | 0 015095848 | -0 00716857 | 0 020148508 | 0 472507048 | 0 | 0 023702652 |
| CLDND1 | 0 199253774 | -0 00717816 | 0 019982414 | -0 30586857 | 0 | 0 023513113 |
| BDH1 | 0 058527346 | -0 00719364 | 0 019716665 | -0 43175226 | 0 | 0 023203298 |
| DNAJB9 | 0 070843883 | -0 00720299 | 0 019557681 | -0 41631296 | 0 | 0 023027671 |
| PIWIL2 | 0 07743047 | -0 00720508 | 0 019522276 | -0 42115948 | 0 | 0 022988848 |
| AIM2 | 0 049209038 | -0 0072333 | 0 019050042 | 0 425345027 | 0 | 0 022443947 |
| SLC11A1 | 0 019103006 | -0 00724663 | 0 018830416 | -0 4771656 | 0 | 0 022193493 |
| MAP7D1 | 0 120709676 | -0 00724681 | 0 018827351 | 0 372801437 | 0 | 0 022192649 |
| HAGHL | 0 050789058 | -0 00724992 | 0 018776535 | -0 43379266 | 0 | 0 022135511 |
| STAT4 | 0 037672993 | -0 00726349 | 0 018555707 | 0 448622945 | 0 | 0 021877908 |

| | | | | | | |
|---|---|---|---|---|---|---|
| NANS | 0 125030934 | -0 00727279 | 0 018405741 | 0 350411621 | 0 | 0 021709217 |
| MDS2 | 0 004740058 | -0 00727469 | 0 018375351 | -0 49299927 | 0 | 0 021676079 |
| DUSP1 | 0 228560279 | -0 00732154 | 0 017636875 | -0 25281583 | 0 | 0 020828351 |
| SLC15A1 | 0 014039329 | -0 00733792 | 0 017385031 | 0 478501132 | 0 | 0 020541204 |
| MYL5 | 0 023976319 | -0 00734167 | 0 017327808 | -0 47069468 | 0 | 0 020476152 |
| RSL1D1 | 0 174944319 | -0 00735085 | 0 017188249 | 0 316959987 | 0 | 0 020313776 |
| SFRP5 | 0 017903714 | -0 007385 | 0 016678258 | -0 47677594 | 0 | 0 019723382 |
| S100A1 | 0 007024424 | -0 0073985 | 0 016480323 | -0 48723283 | 0 | 0 019491746 |
| VIT | 0 319125849 | -0 00742901 | 0 016040666 | -0 19248583 | 0 | 0 018976501 |
| GAP43 | 0 104928518 | -0 00744542 | 0 015808499 | -0 39031009 | 0 | 0 018711212 |
| DNAH6 | 0 123289105 | -0 00745343 | 0 015696214 | -0 36530228 | 0 | 0 018580636 |
| LINC00971 | 0 008585407 | -0 00748423 | 0 015270904 | -0 48822494 | 0 | 0 018083965 |
| SUPT4H1 | 0 091222326 | -0 00750652 | 0 014969659 | -0 4106404 | 0 | 0 017731672 |
| RNASE2 | 0 000856637 | -0 00753333 | 0 014614112 | 0 495713448 | 0 | 0 017312694 |
| FKBP1A | 0 256277246 | -0 00755245 | 0 01436516 | -0 26159167 | 0 | 0 017026309 |
| IL7R | 0 008099979 | -0 0076163 | 0 013560533 | 0 484633186 | 0 | 0 016092809 |
| PATL2 | 0 050969903 | -0 00762156 | 0 013496103 | 0 423102921 | 0 | 0 016018359 |
| PTP4A1 | 0 004901867 | -0 00764395 | 0 013224735 | -0 49295279 | 0 | 0 01570022 |
| SLAMF8 | 0 004054749 | -0 00765252 | 0 013122054 | 0 489712288 | 0 | 0 015584963 |
| NFKB1 | 0 345234243 | -0 00767289 | 0 012881097 | -0 18507837 | 0 | 0 015303793 |
| GK | 0 123812606 | -0 00770377 | 0 012523094 | 0 40456637 | 0 | 0 014885942 |
| PPL | 0 092821382 | -0 00773247 | 0 012198373 | -0 39417615 | 0 | 0 014507252 |
| PLEK2 | 0 003055339 | -0 00774943 | 0 012009985 | 0 491141569 | 0 | 0 014286803 |
| CAB39L | 0 186128191 | -0 00775354 | 0 011964699 | -0 30475642 | 0 | 0 014236515 |
| HCAR3 | 0 001075555 | -0 00775523 | 0 011946191 | 0 494935212 | 0 | 0 014216283 |
| ZDHHC14 | 0 231748872 | -0 00779744 | 0 011490634 | 0 285006036 | 0 | 0 013684498 |
| CRACR2A | 0 10660372 | -0 00781178 | 0 01133945 | 0 348347848 | 0 | 0 013507854 |
| MYCL | 0 001684719 | -0 00782086 | 0 011244574 | -0 49476326 | 0 | 0 013396523 |
| LINC01891 | 0 002541357 | -0 00783065 | 0 011143133 | -0 49424205 | 0 | 0 013277343 |
| SREBF1 | 0 207877253 | -0 00783626 | 0 011085303 | -0 28459942 | 0 | 0 013210102 |
| LINC00987 | 0 017589614 | -0 00789898 | 0 010457364 | -0 4761572 | 0 | 0 012475961 |
| MMP28 | 0 081904019 | -0 00794776 | 0 009991068 | -0 38613127 | 0 | 0 011942271 |
| MRAS | 0 187632065 | -0 00795234 | 0 009948285 | -0 26479031 | 0 | 0 011892637 |
| SLC3A2 | 0 246330738 | -0 00799095 | 0 009593751 | 0 248831895 | 0 | 0 01147607 |
| GPR68 | 0 018436733 | -0 00799823 | 0 009528128 | 0 467558663 | 0 | 0 011399014 |
| LRRC4 | 0 059174583 | -0 00801953 | 0 009338588 | 0 417015666 | 0 | 0 011173671 |
| DNAJC8 | 0 196740972 | -0 00804536 | 0 009113268 | 0 310059761 | 0 | 0 010908218 |
| HLA-DOB | 0 00404523 | -0 00804901 | 0 009081846 | 0 489544403 | 0 | 0 010871985 |
| SYAP1 | 0 24694942 | -0 00805572 | 0 009024251 | 0 268743572 | 0 | 0 010804405 |
| DPP6 | 0 071967029 | -0 00808633 | 0 008765799 | -0 41650409 | 0 | 0 010501623 |
| CALU | 0 584540557 | -0 00808963 | 0 008738316 | 0 074813919 | 2 94E-130 | 0 010470024 |
| ARHGAP4 | 0 055938398 | -0 00810591 | 0 008603938 | 0 411905329 | 0 | 0 010314247 |
| MICB | 0 01456283 | -0 00820347 | 0 00783671 | 0 470134468 | 0 | 0 009405246 |
| DNMBP | 0 315461347 | -0 00821512 | 0 007749324 | 0 206523091 | 0 | 0 009303915 |

| | | | | | | |
|---|---|---|---|---|---|---|
| RPS2 | 0 522120272 | -0 00823169 | 0 007626511 | -0 12649368 | 0 | 0 009159956 |
| FBP1 | 0 014201138 | -0 00824454 | 0 007532472 | -0 48259848 | 0 | 0 009048159 |
| LAMTOR4 | 0 204412633 | -0 00828115 | 0 007270224 | -0 31091495 | 0 | 0 008738694 |
| ZEB2-AS1 | 0 008880471 | -0 00831499 | 0 007035155 | 0 484984453 | 0 | 0 00846045 |
| ADRA2A | 0 007119606 | -0 00832107 | 0 006993645 | -0 48858072 | 0 | 0 008411601 |
| HJURP | 0 051027013 | -0 00834675 | 0 006820809 | 0 371395223 | 0 | 0 008207901 |
| MINDY2 | 0 300984181 | -0 00835714 | 0 006751936 | -0 20645695 | 0 | 0 008128126 |
| DERL3 | 0 004530658 | -0 00835837 | 0 006743805 | 0 487389311 | 0 | 0 008119373 |
| TMEM156 | 0 009917953 | -0 00836019 | 0 00673186 | -0 48784902 | 0 | 0 008106024 |
| SSR2 | 0 277340999 | -0 0083809 | 0 006596869 | -0 2521611 | 0 | 0 007949555 |
| PARP14 | 0 314994955 | -0 00838754 | 0 006554092 | -0 22507302 | 0 | 0 00790103 |
| GFRA2 | 0 049799166 | -0 0084133 | 0 006390535 | -0 43759112 | 0 | 0 007707793 |
| SERPINB1 | 0 111924388 | -0 0084185 | 0 006357906 | -0 38291129 | 0 | 0 007670395 |
| VSIG4 | 0 013887038 | -0 00844167 | 0 006214641 | 0 477809351 | 0 | 0 007500429 |
| SCNN1B | 0 003217148 | -0 00846317 | 0 006084264 | -0 49318242 | 0 | 0 007345892 |
| BAX | 0 166206621 | -0 00847306 | 0 006025109 | 0 343604144 | 0 | 0 007276329 |
| MARCHF1 | 0 081428109 | -0 00849414 | 0 005900774 | 0 433145722 | 0 | 0 007127997 |
| MYT1L | 0 016837677 | -0 00851156 | 0 005799757 | 0 472512666 | 0 | 0 007008658 |
| SYT1 | 0 336744018 | -0 00851734 | 0 005766627 | 0 123323554 | 0 | 0 006970405 |
| LAMB3 | 0 006158268 | -0 00851836 | 0 005760751 | 0 486604296 | 0 | 0 006964193 |
| HECW2-AS1 | 0 001951229 | -0 00851891 | 0 005757624 | 0 492864604 | 0 | 0 006961304 |
| ASAP2 | 0 37731054 | -0 00854102 | 0 005632497 | 0 157359406 | 0 | 0 006810889 |
| MIR548A1HG | 0 003931012 | -0 00859456 | 0 005339655 | -0 49232626 | 0 | 0 006460914 |
| PMP2 | 0 01224991 | -0 00860401 | 0 00528938 | -0 48337026 | 0 | 0 006402542 |
| BIRC5 | 0 053711142 | -0 00862543 | 0 005177112 | 0 36838883 | 0 | 0 006268252 |
| SERPINH1 | 0 332375169 | -0 00863854 | 0 005109453 | -0 21400994 | 0 | 0 006187919 |
| EML2-AS1 | 0 00404523 | -0 00866244 | 0 004988128 | 0 489456161 | 0 | 0 006044084 |
| RNASE6 | 0 007424188 | -0 00866608 | 0 004969895 | 0 486021352 | 0 | 0 006022763 |
| VPS13C | 0 685985418 | -0 0086952 | 0 004826104 | -0 04255606 | 2 55E-43 | 0 00585001 |
| CDCA8 | 0 043707525 | -0 00869892 | 0 004808018 | 0 386374567 | 0 | 0 005828835 |
| C19orf33 | 0 002246293 | -0 00872842 | 0 004666646 | -0 49457043 | 0 | 0 005661805 |
| LILRB3 | 0 011326645 | -0 00873585 | 0 004631632 | 0 478123685 | 0 | 0 005620768 |
| GRIN2A | 0 018303478 | -0 00875217 | 0 004555603 | -0 47816511 | 0 | 0 005531344 |
| KCNMA1-AS1 | 0 039776513 | -0 00878355 | 0 004412541 | 0 42515538 | 0 | 0 005360395 |
| CORO1B | 0 071424492 | -0 00879616 | 0 004356207 | -0 4190183 | 0 | 0 005294002 |
| KCNK5 | 0 011774 | -0 00881751 | 0 004262317 | -0 48163509 | 0 | 0 005181898 |
| DDX24 | 0 363413984 | -0 00882504 | 0 004229673 | -0 16576859 | 0 | 0 005142873 |
| GIMAP5 | 0 00867107 | -0 00885435 | 0 004104651 | -0 48607834 | 0 | 0 004994072 |
| HCST | 0 012897146 | -0 00887486 | 0 004019135 | 0 480008149 | 0 | 0 004893175 |
| ISG15 | 0 118986884 | -0 00887869 | 0 004003356 | 0 37202938 | 0 | 0 004874592 |
| ANK2 | 0 644695513 | -0 00892538 | 0 003815462 | 0 014106876 | 4 82E-06 | 0 004649403 |
| RBP7 | 0 021929908 | -0 00892892 | 0 003801544 | -0 47049748 | 0 | 0 00463304 |
| CXCL16 | 0 037663475 | -0 00892912 | 0 003800745 | -0 4567087 | 0 | 0 004632663 |
| TYMP | 0 124497916 | -0 00894103 | 0 003754267 | -0 38360783 | 0 | 0 004577784 |

| | | | | | | |
|---|---|---|---|---|---|---|
| AFDN | 0 487159963 | -0 0089576 | 0 003690472 | 0 117299588 | 0 | 0 004500576 |
| HAMP | 0 006110678 | -0 00896604 | 0 003658358 | -0 4904534 | 0 | 0 004461988 |
| RNF169 | 0 388608631 | -0 0090052 | 0 003512647 | 0 130936028 | 0 | 0 004287035 |
| ZZZ3 | 0 257067256 | -0 00902132 | 0 003454202 | -0 21006143 | 0 | 0 004216251 |
| PSMB10 | 0 050560621 | -0 00902881 | 0 003427376 | -0 44212361 | 0 | 0 004184046 |
| AREG | 0 008813843 | -0 00905303 | 0 003341861 | 0 48459325 | 0 | 0 004080179 |
| VPS28 | 0 201490548 | -0 00905846 | 0 003322958 | -0 3106153 | 0 | 0 004058148 |
| RPS6KA5 | 0 257552683 | -0 00908427 | 0 003234401 | 0 23226814 | 0 | 0 003952041 |
| LINC01949 | 0 03159087 | -0 00911529 | 0 003130852 | 0 446819014 | 0 | 0 003827497 |
| HLA-DQB2 | 0 001408692 | -0 00913808 | 0 003056712 | -0 49574521 | 0 | 0 003738311 |
| GHRL | 0 041927624 | -0 00913969 | 0 003051535 | 0 426411857 | 0 | 0 003732463 |
| GPR157 | 0 058184691 | -0 00916759 | 0 002963067 | -0 43606973 | 0 | 0 003626131 |
| FNIP1 | 0 668072186 | -0 00917112 | 0 002952065 | 0 056390425 | 9 53E-75 | 0 003613134 |
| ADCY8 | 0 042479679 | -0 00918435 | 0 002911082 | -0 45050774 | 0 | 0 003563435 |
| DDIT4 | 0 279082827 | -0 00919309 | 0 002884299 | -0 24235274 | 0 | 0 003531108 |
| ITPRIP | 0 113428261 | -0 00919997 | 0 002863356 | 0 358481144 | 0 | 0 003505923 |
| B3GALT4 | 0 02233919 | -0 00920711 | 0 002841798 | -0 47186875 | 0 | 0 003481332 |
| RAB20 | 0 042603415 | -0 00921785 | 0 002809606 | -0 44880071 | 0 | 0 003443234 |
| ATP6AP1 | 0 153338029 | -0 00922546 | 0 002787016 | -0 33562004 | 0 | 0 003415992 |
| FERMT3 | 0 033894272 | -0 00923915 | 0 002746806 | -0 45933042 | 0 | 0 003367582 |
| RORC | 0 00141821 | -0 00926264 | 0 00267901 | -0 49564664 | 0 | 0 003284889 |
| CCRL2 | 0 001123146 | -0 00926421 | 0 002674528 | 0 494506363 | 0 | 0 00327982 |
| CCNB1 | 0 048971084 | -0 00929387 | 0 002591249 | 0 389612992 | 0 | 0 003180583 |
| TNS4 | 0 005101749 | -0 0093196 | 0 002520949 | 0 490701718 | 0 | 0 0030955 |
| CD53 | 0 020140488 | -0 00934427 | 0 00245517 | 0 472018116 | 0 | 0 003017082 |
| PSMB8-AS1 | 0 008575889 | -0 0093569 | 0 002422096 | 0 48514192 | 0 | 0 002976826 |
| INMT | 0 061411357 | -0 0093787 | 0 002365978 | -0 42710293 | 0 | 0 002909747 |
| ADAP2 | 0 037644439 | -0 00940002 | 0 002312259 | -0 45576371 | 0 | 0 002844791 |
| SLC7A5 | 0 048580838 | -0 00941757 | 0 002268884 | 0 447529169 | 0 | 0 002792517 |
| CDKL2 | 0 040852068 | -0 00941989 | 0 002263213 | 0 4672459 | 0 | 0 002785901 |
| ANXA4 | 0 407073918 | -0 00943322 | 0 00223083 | -0 12715544 | 0 | 0 002746755 |
| SNRPN | 0 129770992 | -0 00943873 | 0 002217572 | -0 34394331 | 0 | 0 002730786 |
| EIF5B | 0 444918239 | -0 00944408 | 0 002204768 | 0 111852456 | 1 33E-289 | 0 002715727 |
| NUDT3 | 0 332127696 | -0 00948378 | 0 002111838 | -0 16101435 | 0 | 0 002602957 |
| DLGAP1-AS2 | 0 041337496 | -0 00948407 | 0 002111181 | 0 458024218 | 0 | 0 002602486 |
| PHF11 | 0 154242257 | -0 00948867 | 0 002100656 | -0 337879 | 0 | 0 002590526 |
| C11orf96 | 0 086606004 | -0 00949694 | 0 002081845 | -0 39732935 | 0 | 0 002568334 |
| CTBP2 | 0 510384344 | -0 00952339 | 0 002022726 | 0 111773008 | 3 42E-289 | 0 002497356 |
| RAB10 | 0 552673659 | -0 00953681 | 0 001993326 | -0 09286677 | 6 62E-200 | 0 002461701 |
| OSM | 0 001599056 | -0 00955286 | 0 001958671 | 0 494398283 | 0 | 0 0024208 |
| CASC19 | 0 035969237 | -0 009572 | 0 001918083 | 0 43778143 | 0 | 0 002371567 |
| STAT1 | 0 36764006 | -0 00958057 | 0 001900164 | 0 185281827 | 0 | 0 002350026 |
| KLKB1 | 0 027469494 | -0 00961215 | 0 001835412 | -0 46672558 | 0 | 0 002270836 |
| TSPAN13 | 0 040880623 | -0 00963093 | 0 001797884 | 0 446912933 | 0 | 0 002225278 |

| | | | | | | |
|---|---|---|---|---|---|---|
| LINC02233 | 0 043802707 | -0 00963243 | 0 001794926 | -0 45270588 | 0 | 0 002221908 |
| HCAR1 | 0 008395043 | -0 00963745 | 0 001785033 | -0 48782312 | 0 | 0 002209952 |
| IFFO1 | 0 151853191 | -0 0096608 | 0 001739642 | 0 304443034 | 0 | 0 002155168 |
| IL6R | 0 096495403 | -0 00966811 | 0 001725659 | -0 38008945 | 0 | 0 002138405 |
| KIF4A | 0 09051798 | -0 00967987 | 0 001703381 | 0 332856139 | 0 | 0 002111907 |
| CXCL9 | 0 007985761 | -0 00968695 | 0 001690101 | 0 48148562 | 0 | 0 002095716 |
| G3BP1 | 0 344939179 | -0 00970464 | 0 001657304 | 0 194301269 | 0 | 0 002055318 |
| HSPA9 | 0 347813672 | -0 00971488 | 0 001638594 | 0 206325446 | 0 | 0 002032648 |
| COX5A | 0 132426567 | -0 00977765 | 0 001528092 | 0 364898473 | 0 | 0 001897316 |
| NINJ1 | 0 045696827 | -0 00979387 | 0 00150068 | -0 45138314 | 0 | 0 001863525 |
| FREM2 | 0 059069883 | -0 00981627 | 0 001463561 | -0 41916297 | 0 | 0 001818148 |
| ATG16L2 | 0 11537949 | -0 00981881 | 0 001459417 | 0 346503024 | 0 | 0 001813477 |
| LINC02320 | 0 005815614 | -0 00982465 | 0 001449898 | 0 486359855 | 0 | 0 001802359 |
| ANAPC15 | 0 02985856 | -0 00983274 | 0 001436818 | 0 456945164 | 0 | 0 001786335 |
| TMEM102 | 0 00178942 | -0 00983985 | 0 001425411 | -0 49527456 | 0 | 0 001772619 |
| AKAP1 | 0 10660372 | -0 00986116 | 0 00139171 | 0 374633859 | 0 | 0 001731621 |
| SLITRK4 | 0 02034037 | -0 00986423 | 0 001386928 | -0 47123901 | 0 | 0 001726126 |
| RNF19B | 0 090908226 | -0 00987058 | 0 001377053 | 0 393683194 | 0 | 0 001714061 |
| EZR | 0 281167311 | -0 00987725 | 0 001366762 | 0 227163134 | 0 | 0 001701699 |
| TAPBP | 0 266271345 | -0 00989932 | 0 001333211 | -0 24518514 | 0 | 0 001661021 |
| SYP | 0 034865127 | -0 00991875 | 0 001304294 | -0 45637917 | 0 | 0 001626066 |
| ZDHHC12 | 0 013515829 | -0 00993086 | 0 001286573 | -0 48241498 | 0 | 0 001604397 |
| HLA-DRB6 | 0 009308789 | -0 00998758 | 0 001206488 | -0 48627362 | 0 | 0 001506117 |
| DDIT3 | 0 139203518 | -0 00999651 | 0 0011943 | -0 36266874 | 0 | 0 0014911 |
| POLE3 | 0 069473263 | -0 01004237 | 0 001133507 | 0 407228174 | 0 | 0 001416508 |
| OTUD7A | 0 120709676 | -0 01005906 | 0 00111211 | -0 35283249 | 0 | 0 001390689 |
| RP2 | 0 110068341 | -0 01007598 | 0 001090787 | 0 381883623 | 0 | 0 001364206 |
| TBC1D1 | 0 241647789 | -0 01014794 | 0 001004285 | 0 240738909 | 0 | 0 001256853 |
| FLII | 0 109630504 | -0 01015369 | 0 000997654 | -0 38613489 | 0 | 0 001248885 |
| SWAP70 | 0 243989263 | -0 01018679 | 0 000960265 | -0 26853728 | 0 | 0 001203356 |
| MIR29B2CHG | 0 216167596 | -0 01019344 | 0 000952907 | -0 27629752 | 0 | 0 001194293 |
| F2RL2 | 0 031981116 | -0 0101968 | 0 000949213 | -0 45362974 | 0 | 0 001189822 |
| ST6GAL1 | 0 261873941 | -0 01023133 | 0 000911994 | -0 20931792 | 0 | 0 001143623 |
| ACSM5 | 0 006025014 | -0 01023887 | 0 00090405 | -0 49082113 | 0 | 0 001133963 |
| IRF2 | 0 223429975 | -0 01024536 | 0 000897264 | -0 29153542 | 0 | 0 0011256 |
| LRRTM4 | 0 076773715 | -0 01025447 | 0 000887814 | -0 41487668 | 0 | 0 001114189 |
| GLIPR1 | 0 115817327 | -0 01027669 | 0 000865158 | 0 356609282 | 0 | 0 001085901 |
| EFNB2 | 0 130475338 | -0 01031684 | 0 00082558 | 0 348193703 | 0 | 0 001036775 |
| FOCAD | 0 42477775 | -0 01032715 | 0 000815688 | 0 166088493 | 0 | 0 001024761 |
| RBM3 | 0 359463936 | -0 01035213 | 0 000792178 | -0 1858205 | 0 | 0 000995887 |
| MED15 | 0 324751099 | -0 01035396 | 0 000790478 | 0 207685086 | 0 | 0 000994015 |
| MANF | 0 137366507 | -0 01035863 | 0 000786163 | -0 3598208 | 0 | 0 000988983 |
| B3GNT2 | 0 072052693 | -0 01037068 | 0 000775131 | -0 41818601 | 0 | 0 000975365 |
| SERPINB6 | 0 266271345 | -0 01037327 | 0 000772772 | -0 24690637 | 0 | 0 000972526 |

| | | | | | | |
|---|---|---|---|---|---|---|
| NCL | 0 323152044 | -0 0103849 | 0 000762287 | 0 214128332 | 0 | 0 000959969 |
| ITPRID2 | 0 346033771 | -0 01038549 | 0 000761765 | -0 19735013 | 0 | 0 00095944 |
| HOOK2 | 0 189373894 | -0 01042303 | 0 000728838 | 0 26978236 | 0 | 0 00091858 |
| ATP5IF1 | 0 176705184 | -0 01048028 | 0 000681151 | -0 32489848 | 0 | 0 000859397 |
| FCGR1A | 0 006757914 | -0 01050282 | 0 000663189 | -0 49123616 | 0 | 0 000836958 |
| NEURL1B | 0 1280482 | -0 0105162 | 0 000652736 | 0 304737996 | 0 | 0 000823985 |
| HLA-DQA2 | 0 002788829 | -0 01058673 | 0 000600122 | -0 49495175 | 0 | 0 000758176 |
| FCGR2B | 0 00751937 | -0 01060367 | 0 000588082 | -0 48982204 | 0 | 0 000743164 |
| TENT5C | 0 022643772 | -0 01062912 | 0 000570423 | -0 47285137 | 0 | 0 00072104 |
| ARHGAP11B | 0 094572728 | -0 01066169 | 0 000548543 | 0 303554532 | 0 | 0 000693661 |
| LYL1 | 0 010650854 | -0 0106708 | 0 000542562 | 0 480888102 | 0 | 0 000686373 |
| ABCB1 | 0 039310122 | -0 01070232 | 0 00052234 | -0 45534611 | 0 | 0 000661146 |
| PXN | 0 3489273 | -0 01071607 | 0 000513745 | -0 18343466 | 0 | 0 00065044 |
| ZNF699 | 0 088880851 | -0 01071885 | 0 000512021 | 0 378730558 | 0 | 0 000648432 |
| MTRNR2L12 | 0 005644286 | -0 01074253 | 0 000497566 | 0 488187119 | 0 | 0 000630295 |
| SNHG29 | 0 317298357 | -0 01075255 | 0 000491565 | -0 2136201 | 0 | 0 000623027 |
| PAQR3 | 0 189231121 | -0 01076245 | 0 0004857 | 0 29465296 | 0 | 0 000615842 |
| BRINP2 | 0 016675868 | -0 01077308 | 0 000479477 | 0 463116081 | 0 | 0 000608033 |
| LIMA1 | 0 524061982 | -0 01077666 | 0 000477396 | -0 07482245 | 2 75E-130 | 0 000605556 |
| DOP1B | 0 15680265 | -0 01079169 | 0 000468761 | 0 310147466 | 0 | 0 000594842 |
| NRROS | 0 005301631 | -0 0108278 | 0 000448606 | -0 48969466 | 0 | 0 000569419 |
| CHP1 | 0 156250595 | -0 01084957 | 0 000436848 | -0 33605758 | 0 | 0 000554719 |
| BCL2A1 | 0 003769203 | -0 01085771 | 0 000432525 | -0 49391874 | 0 | 0 000549303 |
| PLAC9 | 0 221907064 | -0 01089574 | 0 000412851 | -0 25800133 | 0 | 0 000524575 |
| NMI | 0 074974777 | -0 01090925 | 0 00040607 | 0 404054448 | 0 | 0 00051626 |
| TAGLN | 0 127762654 | -0 01094444 | 0 000388883 | -0 35308777 | 0 | 0 000494676 |
| BCL3 | 0 046448764 | -0 01095291 | 0 000384849 | 0 434384994 | 0 | 0 000489742 |
| BDKRB2 | 0 072423902 | -0 01096448 | 0 000379404 | -0 41376026 | 0 | 0 000483007 |
| CLEC2B | 0 197426282 | -0 01107022 | 0 000332858 | 0 257334896 | 0 | 0 000424095 |
| GCNT4 | 0 022481963 | -0 01108842 | 0 000325407 | -0 47156688 | 0 | 0 000414657 |
| NOS3 | 0 006738878 | -0 01110707 | 0 000317932 | -0 48720265 | 0 | 0 00040535 |
| LDLRAP1 | 0 069749291 | -0 0111538 | 0 000299907 | 0 40163877 | 0 | 0 000382576 |
| IL12RB1 | 0 004168967 | -0 01116738 | 0 00029485 | 0 489744786 | 0 | 0 000376226 |
| CYFIP2 | 0 1510727 | -0 01117582 | 0 000291748 | -0 32984265 | 0 | 0 000372318 |
| NDRG1 | 0 278978127 | -0 01122444 | 0 000274465 | -0 22824216 | 0 | 0 000350405 |
| SCO2 | 0 035074527 | -0 01124142 | 0 000268661 | -0 46233674 | 0 | 0 00034318 |
| RANGAP1 | 0 098798805 | -0 01127459 | 0 000257648 | 0 357174551 | 0 | 0 000329291 |
| COQ10B | 0 149683044 | -0 01130621 | 0 000247545 | -0 33974807 | 0 | 0 000316636 |
| RSL24D1 | 0 161694999 | -0 01131881 | 0 000243626 | -0 34418019 | 0 | 0 000311791 |
| HNRNPF | 0 276903162 | -0 01132943 | 0 000240366 | 0 263388926 | 0 | 0 000307911 |
| FOXN2 | 0 282718776 | -0 01133184 | 0 000239632 | 0 239637531 | 0 | 0 000307012 |
| SAT1 | 0 381945899 | -0 01133294 | 0 000239298 | -0 16650086 | 0 | 0 000306626 |
| ANKRD22 | 0 003759685 | -0 01133581 | 0 000238426 | -0 49373742 | 0 | 0 00030555 |
| CXCL8 | 0 014201138 | -0 01135006 | 0 000234151 | -0 48193514 | 0 | 0 000300234 |

| | | | | | | |
|---|---|---|---|---|---|---|
| PCGF5 | 0 354324113 | -0 0113791 | 0 000225657 | -0 16793767 | 0 | 0 000289421 |
| TRMT1 | 0 118282538 | -0 01139779 | 0 000220345 | 0 342892915 | 0 | 0 000282724 |
| KLRG1 | 0 048428547 | -0 01139957 | 0 000219843 | 0 434918814 | 0 | 0 000282118 |
| PBDC1 | 0 059174583 | -0 01140527 | 0 000218252 | -0 43506994 | 0 | 0 000280114 |
| KRT10 | 0 194980107 | -0 01143304 | 0 000210642 | -0 31145577 | 0 | 0 000270532 |
| SH2B3 | 0 151729455 | -0 01145571 | 0 000204616 | 0 318362979 | 0 | 0 000262935 |
| CD55 | 0 165388057 | -0 0114653 | 0 000202118 | -0 33514263 | 0 | 0 000259759 |
| NR2F1 | 0 190087758 | -0 01148573 | 0 000196887 | -0 31243526 | 0 | 0 000253175 |
| SEC11A | 0 364746531 | -0 01149286 | 0 000195091 | 0 183277166 | 0 | 0 000250899 |
| CHCHD10 | 0 065132969 | -0 01150361 | 0 000192413 | -0 42016709 | 0 | 0 00024759 |
| LYSMD2 | 0 03069616 | -0 0115372 | 0 000184267 | 0 459519561 | 0 | 0 000237238 |
| SFR1 | 0 028763968 | -0 01155885 | 0 000179188 | 0 458154264 | 0 | 0 00023073 |
| NSG1 | 0 019141079 | -0 01158241 | 0 000173814 | -0 47476898 | 0 | 0 000223871 |
| TAGAP | 0 035997792 | -0 01160361 | 0 000169106 | -0 44805378 | 0 | 0 000217866 |
| POLD4 | 0 105461537 | -0 0116118 | 0 000167319 | -0 39358416 | 0 | 0 000215594 |
| GRIK2 | 0 262445032 | -0 01162326 | 0 00016485 | -0 21443769 | 0 | 0 000212442 |
| FLOT2 | 0 093335364 | -0 01162743 | 0 00016396 | -0 39898725 | 0 | 0 000211323 |
| ST8SIA6-AS1 | 0 078820125 | -0 01164929 | 0 000159368 | -0 40568746 | 0 | 0 000205433 |
| RAB9A | 0 163903219 | -0 01166057 | 0 000157045 | -0 33668973 | 0 | 0 000202521 |
| TXNRD1 | 0 273666978 | -0 01167133 | 0 000154861 | 0 247559613 | 0 | 0 000199759 |
| ZMAT1 | 0 152643201 | -0 01167812 | 0 000153496 | -0 32495985 | 0 | 0 000198026 |
| FAM72D | 0 010726999 | -0 01168837 | 0 000151457 | 0 471520345 | 0 | 0 000195502 |
| NAT8L | 0 005491995 | -0 01170057 | 0 000149066 | -0 48884644 | 0 | 0 00019252 |
| COL6A2 | 0 69612229 | -0 01176444 | 0 000137108 | -0 04825554 | 3 32E-55 | 0 000177295 |
| PHTF1 | 0 196074699 | -0 01179295 | 0 000132069 | 0 277460674 | 0 | 0 00017092 |
| PAOX | 0 033047153 | -0 01181796 | 0 000127793 | 0 457281038 | 0 | 0 000165543 |
| CALM3 | 0 150587272 | -0 01186288 | 0 000120437 | 0 358980247 | 0 | 0 000156186 |
| CPM | 0 174487446 | -0 01186293 | 0 000120428 | -0 32797685 | 0 | 0 000156186 |
| NETO2 | 0 080876054 | -0 01187937 | 0 000117838 | 0 395436685 | 0 | 0 0001529 |
| GNB1 | 0 625716244 | -0 01189072 | 0 000116079 | 0 063341776 | 7 46E-94 | 0 000150701 |
| ICAM1 | 0 036730692 | -0 01190891 | 0 000113314 | -0 45887801 | 0 | 0 000147172 |
| PTPN7 | 0 022757991 | -0 01191432 | 0 000112503 | 0 46481948 | 0 | 0 000146138 |
| CCL3L1 | 0 00252232 | -0 01192637 | 0 000110718 | 0 492371448 | 0 | 0 000143858 |
| MFNG | 0 006234414 | -0 01194637 | 0 000107813 | -0 48990541 | 0 | 0 000140123 |
| DAPP1 | 0 006586587 | -0 01195479 | 0 000106611 | -0 49083463 | 0 | 0 000138599 |
| DHRS7 | 0 128847728 | -0 01196227 | 0 000105555 | -0 36314053 | 0 | 0 000137244 |
| TUBB4A | 0 004578249 | -0 01200156 | 0 000100165 | -0 4911441 | 0 | 0 000130308 |
| ADAMTS1 | 0 141935238 | -0 01201286 | 9 87E-05 | -0 33187149 | 0 | 0 000128391 |
| TRAK2 | 0 292503474 | -0 01202031 | 9 77E-05 | 0 23631504 | 0 | 0 000127136 |
| ARHGAP15 | 0 086025395 | -0 01203458 | 9 58E-05 | 0 392952479 | 0 | 0 000124781 |
| RIOK3 | 0 324846281 | -0 01207704 | 9 05E-05 | 0 202043526 | 0 | 0 00011802 |
| H2AZ1 | 0 091783899 | -0 01207854 | 9 03E-05 | 0 419431468 | 0 | 0 000117798 |
| ERO1A | 0 21804268 | -0 01215363 | 8 17E-05 | -0 29603005 | 0 | 0 000106567 |
| ATP5PF | 0 196817118 | -0 01216612 | 8 03E-05 | -0 31259176 | 0 | 0 000104797 |

| | | | | | | |
|---|---|---|---|---|---|---|
| BLVRB | 0 14008871 | -0 01217693 | 7 91E-05 | -0 36394178 | 0 | 0 000103304 |
| SNRPG | 0 202023567 | -0 01223448 | 7 32E-05 | 0 30442535 | 0 | 9 56E-05 |
| OASL | 0 00872818 | -0 01224321 | 7 23E-05 | 0 481792675 | 0 | 9 45E-05 |
| CYP4B1 | 0 022301117 | -0 01225288 | 7 14E-05 | -0 46718281 | 0 | 9 33E-05 |
| TMEM107 | 0 061259066 | -0 01225301 | 7 14E-05 | 0 439247862 | 0 | 9 33E-05 |
| EEF2 | 0 375216539 | -0 01226038 | 7 06E-05 | -0 19346762 | 0 | 9 24E-05 |
| WIPI1 | 0 280558147 | -0 01226623 | 7 01E-05 | -0 21710617 | 0 | 9 16E-05 |
| TMEM160 | 0 074898631 | -0 0122974 | 6 72E-05 | 0 423860995 | 0 | 8 79E-05 |
| SLC16A1 | 0 150320763 | -0 01230117 | 6 68E-05 | 0 314640295 | 0 | 8 74E-05 |
| MT1F | 0 017694314 | -0 012308 | 6 62E-05 | -0 47045233 | 0 | 8 67E-05 |
| PTPN14 | 0 471683387 | -0 01237371 | 6 05E-05 | 0 135748275 | 0 | 7 93E-05 |
| DCBLD1 | 0 125668653 | -0 01238266 | 5 98E-05 | 0 33991523 | 0 | 7 83E-05 |
| HMOX1 | 0 028154804 | -0 01239883 | 5 85E-05 | -0 47006224 | 0 | 7 66E-05 |
| SLC2A5 | 0 020759171 | -0 01240387 | 5 80E-05 | -0 47636115 | 0 | 7 61E-05 |
| TMEM9B | 0 093373437 | -0 01246736 | 5 32E-05 | -0 39830939 | 0 | 6 98E-05 |
| TOR4A | 0 010831699 | -0 01248677 | 5 18E-05 | 0 480556951 | 0 | 6 80E-05 |
| FAM53B | 0 19953932 | -0 01250233 | 5 07E-05 | -0 29870922 | 0 | 6 65E-05 |
| LNCAROD | 0 02875445 | -0 01250667 | 5 04E-05 | 0 453475873 | 0 | 6 62E-05 |
| VCL | 0 590565571 | -0 01251574 | 4 97E-05 | -0 06323862 | 1 49E-93 | 6 54E-05 |
| CCDC78 | 0 007033942 | -0 0125265 | 4 90E-05 | -0 48803477 | 0 | 6 44E-05 |
| TSTD1 | 0 013696674 | -0 01254106 | 4 80E-05 | -0 48206662 | 0 | 6 32E-05 |
| LINC01592 | 0 041794369 | -0 01256361 | 4 65E-05 | 0 423885655 | 0 | 6 12E-05 |
| ERCC6L2 | 0 420399383 | -0 01257913 | 4 55E-05 | -0 14921192 | 0 | 5 99E-05 |
| SLC38A2 | 0 464183054 | -0 01259798 | 4 44E-05 | 0 115017542 | 3 28E-306 | 5 84E-05 |
| ZNF714 | 0 15307152 | -0 01264151 | 4 17E-05 | 0 285518625 | 0 | 5 50E-05 |
| HPS5 | 0 17903714 | -0 01264976 | 4 13E-05 | -0 30014174 | 0 | 5 44E-05 |
| SYPL1 | 0 217861834 | -0 0126657 | 4 03E-05 | 0 280816481 | 0 | 5 32E-05 |
| MAST2 | 0 434638594 | -0 0126876 | 3 91E-05 | 0 132877767 | 0 | 5 16E-05 |
| LINC02343 | 0 029449278 | -0 01270021 | 3 84E-05 | -0 45812012 | 0 | 5 08E-05 |
| GADD45G | 0 029515905 | -0 0127906 | 3 38E-05 | -0 46279216 | 0 | 4 47E-05 |
| ERP44 | 0 375016657 | -0 01280804 | 3 30E-05 | 0 171377666 | 0 | 4 37E-05 |
| LINC01088 | 0 424159068 | -0 01281096 | 3 29E-05 | -0 12715619 | 0 | 4 35E-05 |
| CFAP61 | 0 019198188 | -0 01282921 | 3 20E-05 | -0 47358596 | 0 | 4 24E-05 |
| LINC01709 | 0 005920314 | -0 01288189 | 2 97E-05 | -0 48956502 | 0 | 3 94E-05 |
| IL15RA | 0 023500409 | -0 01289205 | 2 93E-05 | 0 456874805 | 0 | 3 88E-05 |
| GNA13 | 0 29825246 | -0 01289998 | 2 90E-05 | 0 215531419 | 0 | 3 84E-05 |
| DTNA | 0 274238069 | -0 01293211 | 2 77E-05 | 0 25842825 | 0 | 3 67E-05 |
| GNGT1 | 0 202623213 | -0 01294834 | 2 70E-05 | -0 32614576 | 0 | 3 59E-05 |
| AC104653.1 | 0 003816794 | -0 01295983 | 2 66E-05 | 0 489531743 | 0 | 3 53E-05 |
| CASP1 | 0 013392092 | -0 01296133 | 2 65E-05 | -0 48369609 | 0 | 3 52E-05 |
| STX4 | 0 103405608 | -0 01298993 | 2 55E-05 | -0 39488445 | 0 | 3 38E-05 |
| MAL | 0 007376597 | -0 01299965 | 2 51E-05 | -0 48937363 | 0 | 3 34E-05 |
| PARK7 | 0 25993223 | -0 01300165 | 2 50E-05 | 0 252506543 | 0 | 3 33E-05 |
| RGS16 | 0 085130685 | -0 01304912 | 2 34E-05 | -0 39803071 | 0 | 3 11E-05 |

| | | | | | | |
|---|---|---|---|---|---|---|
| NKG7 | 0 005282595 | -0 013081 | 2 23E-05 | 0 487526216 | 0 | 2 97E-05 |
| ASF1B | 0 045487426 | -0 01308882 | 2 21E-05 | 0 38305774 | 0 | 2 94E-05 |
| TRAF6 | 0 113828025 | -0 01311269 | 2 13E-05 | -0 38656349 | 0 | 2 84E-05 |
| GK-AS1 | 0 027041176 | -0 01316248 | 1 99E-05 | 0 474599221 | 0 | 2 65E-05 |
| MYOC | 0 016057185 | -0 01316617 | 1 97E-05 | -0 48280576 | 0 | 2 63E-05 |
| YWHAZ | 0 568883136 | -0 01316901 | 1 97E-05 | 0 099986945 | 1 47E-231 | 2 62E-05 |
| SNCG | 0 016656831 | -0 01317544 | 1 95E-05 | -0 47473178 | 0 | 2 60E-05 |
| ADORA3 | 0 002693648 | -0 01319728 | 1 89E-05 | -0 4950376 | 0 | 2 52E-05 |
| WDSUB1 | 0 218470998 | -0 01320269 | 1 87E-05 | 0 278876526 | 0 | 2 50E-05 |
| FARP2 | 0 378947669 | -0 01320605 | 1 86E-05 | 0 181732366 | 0 | 2 49E-05 |
| TLR2 | 0 056747444 | -0 01325049 | 1 75E-05 | -0 43377827 | 0 | 2 33E-05 |
| ATP5F1D | 0 158563515 | -0 01329932 | 1 63E-05 | -0 33945538 | 0 | 2 17E-05 |
| TMEM123 | 0 219660772 | -0 01333072 | 1 55E-05 | -0 28629213 | 0 | 2 08E-05 |
| KYNU | 0 024423674 | -0 01333304 | 1 55E-05 | -0 4735853 | 0 | 2 07E-05 |
| C4orf19 | 0 107574575 | -0 0133738 | 1 46E-05 | -0 38680068 | 0 | 1 95E-05 |
| CASP4LP | 0 00877577 | -0 01345812 | 1 29E-05 | -0 48762117 | 0 | 1 72E-05 |
| RGS1 | 0 027793113 | -0 01346013 | 1 28E-05 | -0 47164424 | 0 | 1 72E-05 |
| SIK3 | 0 6740972 | -0 01346537 | 1 27E-05 | -0 02907738 | 4 23E-21 | 1 71E-05 |
| FEM1C | 0 169214369 | -0 01347588 | 1 25E-05 | -0 33037076 | 0 | 1 68E-05 |
| AZGP1 | 0 004759095 | -0 01349514 | 1 22E-05 | -0 49193757 | 0 | 1 63E-05 |
| POMP | 0 224324684 | -0 01350435 | 1 20E-05 | -0 28962888 | 0 | 1 61E-05 |
| E2F7 | 0 112552588 | -0 01350791 | 1 19E-05 | 0 292619679 | 0 | 1 60E-05 |
| DGAT2 | 0 009089871 | -0 01352475 | 1 17E-05 | 0 482343671 | 0 | 1 57E-05 |
| SMIM26 | 0 099369896 | -0 01352893 | 1 16E-05 | -0 39541198 | 0 | 1 56E-05 |
| GIMAP8 | 0 00772877 | -0 01353546 | 1 15E-05 | -0 48665167 | 0 | 1 54E-05 |
| TGIF1 | 0 137947117 | -0 01356444 | 1 10E-05 | 0 351397396 | 0 | 1 48E-05 |
| SERPINA1 | 0 006129714 | -0 013672 | 9 35E-06 | 0 487420339 | 0 | 1 26E-05 |
| SGMS1 | 0 468675639 | -0 01367664 | 9 28E-06 | -0 08575089 | 1 22E-170 | 1 25E-05 |
| CLDN1 | 0 196636272 | -0 01371696 | 8 74E-06 | -0 2928309 | 0 | 1 18E-05 |
| SMAP2 | 0 187070492 | -0 01376645 | 8 11E-06 | 0 306758047 | 0 | 1 09E-05 |
| MACF1 | 0 750109459 | -0 01378943 | 7 83E-06 | 0 021365675 | 4 33E-12 | 1 06E-05 |
| CENPF | 0 12148065 | -0 0137923 | 7 80E-06 | 0 283473567 | 0 | 1 05E-05 |
| HHEX | 0 010393863 | -0 01380165 | 7 69E-06 | 0 480601157 | 0 | 1 04E-05 |
| PYHIN1 | 0 013477756 | -0 0138313 | 7 35E-06 | -0 48291434 | 0 | 9 94E-06 |
| ITGB2 | 0 026936476 | -0 01383702 | 7 28E-06 | 0 457800439 | 0 | 9 85E-06 |
| IFI16 | 0 536378519 | -0 01386727 | 6 96E-06 | -0 09124309 | 5 09E-193 | 9 42E-06 |
| TNFRSF14-AS1 | 0 007214787 | -0 0138857 | 6 76E-06 | -0 48922635 | 0 | 9 16E-06 |
| WASF2 | 0 493594259 | -0 01391473 | 6 47E-06 | -0 11908833 | 0 | 8 77E-06 |
| ZNF490 | 0 197673755 | -0 01392799 | 6 34E-06 | 0 249139147 | 0 | 8 59E-06 |
| NCAPD2 | 0 164540938 | -0 01393307 | 6 29E-06 | 0 239766313 | 0 | 8 53E-06 |
| SEMA3B | 0 130960766 | -0 01393505 | 6 27E-06 | -0 33807569 | 0 | 8 50E-06 |
| BST2 | 0 122936932 | -0 01394537 | 6 17E-06 | -0 37811225 | 0 | 8 37E-06 |
| VAMP2 | 0 210437646 | -0 01394755 | 6 15E-06 | -0 28400486 | 0 | 8 34E-06 |
| MRPL54 | 0 053054387 | -0 01397678 | 5 88E-06 | -0 44422658 | 0 | 7 98E-06 |

| | | | | | | |
|---|---|---|---|---|---|---|
| IRF6 | 0 002684129 | -0 01401472 | 5 55E-06 | -0 49444378 | 0 | 7 54E-06 |
| EIF3G | 0 154746721 | -0 01404819 | 5 27E-06 | -0 35055263 | 0 | 7 16E-06 |
| POMC | 0 007481297 | -0 01411379 | 4 76E-06 | -0 48891888 | 0 | 6 48E-06 |
| SPAG9 | 0 731168263 | -0 01411906 | 4 73E-06 | 0 045527929 | 2 48E-49 | 6 43E-06 |
| CHMP1B | 0 058955664 | -0 01413182 | 4 63E-06 | -0 43450991 | 0 | 6 31E-06 |
| STX1A | 0 06197293 | -0 01413507 | 4 61E-06 | 0 398194524 | 0 | 6 27E-06 |
| MIR34AHG | 0 328310902 | -0 01415238 | 4 49E-06 | 0 182466154 | 0 | 6 11E-06 |
| MATN2 | 0 434552931 | -0 01415377 | 4 48E-06 | -0 13241199 | 0 | 6 10E-06 |
| CPA6 | 0 094401401 | -0 01416266 | 4 42E-06 | 0 307559181 | 0 | 6 02E-06 |
| CCR1 | 0 005120786 | -0 01417327 | 4 34E-06 | 0 488346207 | 0 | 5 92E-06 |
| GRM1 | 0 019302888 | -0 01417671 | 4 32E-06 | -0 47520934 | 0 | 5 89E-06 |
| STX11 | 0 010213017 | -0 01424669 | 3 87E-06 | 0 484143667 | 0 | 5 29E-06 |
| PLEKHA1 | 0 319516095 | -0 01424916 | 3 86E-06 | -0 21261595 | 0 | 5 27E-06 |
| CHST7 | 0 046372618 | -0 01425884 | 3 80E-06 | -0 45463564 | 0 | 5 19E-06 |
| ARHGAP21 | 0 608050484 | -0 01430761 | 3 52E-06 | -0 06019068 | 6 44E-85 | 4 81E-06 |
| PAQR8 | 0 038282157 | -0 01430805 | 3 52E-06 | -0 45291381 | 0 | 4 81E-06 |
| CCL18 | 0 002189184 | -0 01434024 | 3 35E-06 | 0 491620791 | 0 | 4 57E-06 |
| PLEKHF1 | 0 023662218 | -0 01438151 | 3 14E-06 | -0 47370035 | 0 | 4 29E-06 |
| SLC26A8 | 0 013525347 | -0 0144167 | 2 97E-06 | 0 473214808 | 0 | 4 06E-06 |
| RNF149 | 0 451866517 | -0 01443153 | 2 90E-06 | 0 126293691 | 0 | 3 97E-06 |
| CXorf38 | 0 096923721 | -0 01445492 | 2 79E-06 | 0 388057909 | 0 | 3 83E-06 |
| SF3B2 | 0 225676267 | -0 01446134 | 2 76E-06 | 0 282399017 | 0 | 3 79E-06 |
| SNRPB2 | 0 185271554 | -0 01447168 | 2 72E-06 | -0 31361435 | 0 | 3 73E-06 |
| CIITA | 0 039786031 | -0 01447251 | 2 72E-06 | 0 438084372 | 0 | 3 72E-06 |
| ADAM15 | 0 075507795 | -0 01452528 | 2 50E-06 | 0 420921554 | 0 | 3 43E-06 |
| NCKAP5 | 0 369010679 | -0 01457069 | 2 32E-06 | -0 15707618 | 0 | 3 19E-06 |
| OAT | 0 20852449 | -0 0145804 | 2 29E-06 | -0 2867107 | 0 | 3 14E-06 |
| SDF2L1 | 0 071405456 | -0 01458335 | 2 28E-06 | -0 42066454 | 0 | 3 13E-06 |
| CCL13 | 0 004530658 | -0 01460714 | 2 19E-06 | -0 4937602 | 0 | 3 01E-06 |
| CCDC71L | 0 061411357 | -0 01461635 | 2 16E-06 | 0 420496876 | 0 | 2 97E-06 |
| AVPI1 | 0 035065009 | -0 01464798 | 2 05E-06 | -0 4541958 | 0 | 2 83E-06 |
| HACD1 | 0 053901506 | -0 01466573 | 2 00E-06 | -0 42629265 | 0 | 2 75E-06 |
| ANXA3 | 0 038919876 | -0 01469384 | 1 91E-06 | -0 4526213 | 0 | 2 63E-06 |
| EFNA1 | 0 036492738 | -0 01475516 | 1 73E-06 | -0 45206741 | 0 | 2 39E-06 |
| AHNAK2 | 0 076564314 | -0 01478762 | 1 64E-06 | -0 42197697 | 0 | 2 27E-06 |
| DTX1 | 0 117111801 | -0 0148089 | 1 58E-06 | -0 35327034 | 0 | 2 19E-06 |
| PNP | 0 028526013 | -0 01481714 | 1 56E-06 | 0 457474727 | 0 | 2 16E-06 |
| ABHD6 | 0 077316251 | -0 01483254 | 1 53E-06 | -0 41289673 | 0 | 2 11E-06 |
| TCEAL4 | 0 200405475 | -0 01483498 | 1 52E-06 | -0 28598725 | 0 | 2 10E-06 |
| C12orf40 | 0 125592507 | -0 0148353 | 1 52E-06 | -0 38834373 | 0 | 2 10E-06 |
| H1-10 | 0 137223735 | -0 01484023 | 1 51E-06 | 0 38740614 | 0 | 2 08E-06 |
| WDR47 | 0 134482496 | -0 01486468 | 1 45E-06 | 0 351858815 | 0 | 2 00E-06 |
| DPF3 | 0 147979288 | -0 01490479 | 1 36E-06 | 0 260313608 | 0 | 1 88E-06 |
| NUP210L | 0 076516723 | -0 01490832 | 1 35E-06 | 0 387681772 | 0 | 1 87E-06 |

| | | | | | | |
|---|---|---|---|---|---|---|
| COTL1 | 0 169423769 | -0 01496427 | 1 23E-06 | -0 30761422 | 0 | 1 71E-06 |
| EPOR | 0 025185129 | -0 01498779 | 1 18E-06 | 0 460785062 | 0 | 1 64E-06 |
| SERPINE2 | 0 282357084 | -0 01498829 | 1 18E-06 | -0 25376004 | 0 | 1 64E-06 |
| RNF167 | 0 079781462 | -0 01499837 | 1 16E-06 | -0 41018106 | 0 | 1 61E-06 |
| NDUFB1 | 0 204098532 | -0 01500195 | 1 16E-06 | -0 30118355 | 0 | 1 61E-06 |
| GPX3 | 0 085844549 | -0 01502679 | 1 11E-06 | -0 40166701 | 0 | 1 54E-06 |
| UQCRH | 0 231472845 | -0 01506036 | 1 05E-06 | 0 271333729 | 0 | 1 46E-06 |
| EIF5 | 0 28774438 | -0 01511065 | 9 68E-07 | -0 23453829 | 0 | 1 35E-06 |
| C12orf56 | 0 065999124 | -0 01512022 | 9 53E-07 | 0 392984248 | 0 | 1 33E-06 |
| P2RX7 | 0 109145076 | -0 01513885 | 9 24E-07 | -0 38930093 | 0 | 1 29E-06 |
| GM2A | 0 052388114 | -0 01516083 | 8 91E-07 | -0 4397839 | 0 | 1 24E-06 |
| KIAA0930 | 0 095172374 | -0 01516363 | 8 87E-07 | -0 40172341 | 0 | 1 24E-06 |
| GDF10 | 0 022205936 | -0 01517062 | 8 76E-07 | -0 46612418 | 0 | 1 22E-06 |
| WDR74 | 0 083246083 | -0 01518834 | 8 51E-07 | 0 396417557 | 0 | 1 19E-06 |
| C1S | 0 473006415 | -0 01518956 | 8 49E-07 | -0 12714615 | 0 | 1 18E-06 |
| CLEC14A | 0 026689003 | -0 01520879 | 8 23E-07 | -0 46931901 | 0 | 1 15E-06 |
| LACTB | 0 12148065 | -0 01521129 | 8 19E-07 | -0 37509643 | 0 | 1 14E-06 |
| TOB1 | 0 126544326 | -0 01521289 | 8 17E-07 | -0 35231814 | 0 | 1 14E-06 |
| PA2G4 | 0 175163237 | -0 01524136 | 7 79E-07 | 0 308157569 | 0 | 1 09E-06 |
| KRT15 | 0 004806686 | -0 01525376 | 7 64E-07 | 0 48866295 | 0 | 1 07E-06 |
| HIGD2A | 0 089537606 | -0 0152754 | 7 36E-07 | -0 41170358 | 0 | 1 03E-06 |
| C10orf55 | 0 00320763 | -0 01532347 | 6 80E-07 | 0 487989358 | 0 | 9 50E-07 |
| RAPGEF6 | 0 543954998 | -0 01532963 | 6 73E-07 | 0 082565539 | 2 63E-158 | 9 40E-07 |
| CDH1 | 0 017237441 | -0 01534404 | 6 57E-07 | -0 47922813 | 0 | 9 18E-07 |
| ZNF791 | 0 244550837 | -0 01534726 | 6 53E-07 | 0 232993526 | 0 | 9 14E-07 |
| CFB | 0 035997792 | -0 01535665 | 6 43E-07 | -0 45599239 | 0 | 9 00E-07 |
| GAPT | 0 002960157 | -0 01538738 | 6 11E-07 | -0 49344608 | 0 | 8 55E-07 |
| TMEM38B | 0 137680608 | -0 01539342 | 6 05E-07 | 0 337094119 | 0 | 8 46E-07 |
| ST14 | 0 00614875 | -0 01540461 | 5 93E-07 | -0 49142539 | 0 | 8 31E-07 |
| LCTL | 0 013287392 | -0 0154188 | 5 79E-07 | 0 47008809 | 0 | 8 11E-07 |
| CETP | 0 004997049 | -0 01542149 | 5 77E-07 | 0 486981219 | 0 | 8 08E-07 |
| PIK3CD | 0 04263197 | -0 0154685 | 5 33E-07 | 0 436230985 | 0 | 7 47E-07 |
| JUN | 0 405617635 | -0 01548065 | 5 22E-07 | -0 12530498 | 0 | 7 32E-07 |
| FAM78B | 0 127619882 | -0 01548091 | 5 22E-07 | -0 3463987 | 0 | 7 32E-07 |
| ACTN2 | 0 060459538 | -0 01554316 | 4 70E-07 | -0 4288027 | 0 | 6 60E-07 |
| RCN1 | 0 236479412 | -0 01555616 | 4 59E-07 | -0 28730263 | 0 | 6 46E-07 |
| SMKR1 | 0 003978603 | -0 01556826 | 4 50E-07 | 0 490586876 | 0 | 6 33E-07 |
| ALOX5AP | 0 013077992 | -0 01557137 | 4 48E-07 | 0 478101093 | 0 | 6 29E-07 |
| GRB14 | 0 032123889 | -0 01558091 | 4 41E-07 | -0 46570593 | 0 | 6 19E-07 |
| RASGEF1A | 0 005634768 | -0 01560611 | 4 22E-07 | -0 49006402 | 0 | 5 94E-07 |
| IGLC3 | 0 004540176 | -0 01561319 | 4 17E-07 | 0 488618941 | 0 | 5 87E-07 |
| GNAI2 | 0 226380613 | -0 01563611 | 4 01E-07 | -0 27485267 | 0 | 5 64E-07 |
| MAP1LC3B | 0 150063772 | -0 01571296 | 3 52E-07 | -0 34390481 | 0 | 4 96E-07 |
| RPL23 | 0 365212922 | -0 015729 | 3 42E-07 | -0 20891066 | 0 | 4 83E-07 |

| | | | | | | |
|---|---|---|---|---|---|---|
| MIR31HG | 0 18445299 | -0 0157795 | 3 14E-07 | 0 229992225 | 0 | 4 43E-07 |
| TNNC1 | 0 024747292 | -0 01585595 | 2 75E-07 | -0 47141606 | 0 | 3 89E-07 |
| CAVIN2 | 0 038805658 | -0 01591454 | 2 49E-07 | -0 45405618 | 0 | 3 52E-07 |
| FOXJ1 | 0 003455103 | -0 01592949 | 2 42E-07 | -0 49365032 | 0 | 3 43E-07 |
| ABCA1 | 0 502094002 | -0 01596556 | 2 28E-07 | -0 14145075 | 0 | 3 22E-07 |
| SYTL2 | 0 186470846 | -0 01597192 | 2 25E-07 | -0 31232063 | 0 | 3 19E-07 |
| KIF13A | 0 612562106 | -0 01598465 | 2 20E-07 | -0 05140372 | 2 08E-62 | 3 12E-07 |
| SPARCL1 | 0 415307152 | -0 0159862 | 2 20E-07 | -0 11774471 | 0 | 3 11E-07 |
| TACSTD2 | 0 002094002 | -0 01599946 | 2 15E-07 | -0 49485068 | 0 | 3 04E-07 |
| GNA15 | 0 006091641 | -0 01600342 | 2 13E-07 | 0 483835726 | 0 | 3 02E-07 |
| SPTLC2 | 0 349964783 | -0 01602248 | 2 06E-07 | -0 18292692 | 0 | 2 92E-07 |
| ABHD5 | 0 116331309 | -0 01607129 | 1 89E-07 | -0 38007071 | 0 | 2 69E-07 |
| RPL17 | 0 446231749 | -0 01608425 | 1 85E-07 | -0 17461994 | 0 | 2 63E-07 |
| CDKN2D | 0 032980526 | -0 01609336 | 1 82E-07 | 0 442998307 | 0 | 2 59E-07 |
| PET100 | 0 17120367 | -0 01612022 | 1 74E-07 | -0 33720038 | 0 | 2 47E-07 |
| AP1B1 | 0 121166549 | -0 01619317 | 1 53E-07 | 0 350789633 | 0 | 2 18E-07 |
| LILRB5 | 0 006634178 | -0 01623924 | 1 41E-07 | 0 483858622 | 0 | 2 01E-07 |
| ZFAND5 | 0 322990234 | -0 01627025 | 1 33E-07 | -0 17405716 | 0 | 1 90E-07 |
| LINC00861 | 0 016066703 | -0 01635579 | 1 15E-07 | 0 474407857 | 0 | 1 64E-07 |
| SECTM1 | 0 009660962 | -0 01638295 | 1 09E-07 | 0 47712198 | 0 | 1 56E-07 |
| SH3PXD2A | 0 49143363 | -0 01640799 | 1 05E-07 | 0 081086779 | 9 69E-153 | 1 49E-07 |
| XBP1 | 0 141754393 | -0 01642121 | 1 02E-07 | -0 34397779 | 0 | 1 46E-07 |
| IKZF2 | 0 184138889 | -0 01645323 | 9 64E-08 | 0 309695665 | 0 | 1 38E-07 |
| HCRT | 0 003426548 | -0 01645358 | 9 64E-08 | 0 491189624 | 0 | 1 38E-07 |
| NIPA1 | 0 090194361 | -0 01646883 | 9 38E-08 | -0 40520185 | 0 | 1 34E-07 |
| SLC1A5 | 0 099940987 | -0 0165189 | 8 57E-08 | -0 39709064 | 0 | 1 23E-07 |
| HKDC1 | 0 003950049 | -0 01652136 | 8 54E-08 | -0 49362183 | 0 | 1 22E-07 |
| KIF20B | 0 156421922 | -0 01652502 | 8 48E-08 | 0 283547099 | 0 | 1 21E-07 |
| ANKRD55 | 0 023795473 | -0 01652578 | 8 47E-08 | -0 47029021 | 0 | 1 21E-07 |
| ADIPOR2 | 0 395404618 | -0 0165491 | 8 12E-08 | -0 1733659 | 0 | 1 16E-07 |
| C1QTNF1 | 0 051464849 | -0 01656493 | 7 89E-08 | -0 44820587 | 0 | 1 13E-07 |
| PALM2AKAP2 | 0 251813215 | -0 01656925 | 7 83E-08 | -0 23451005 | 0 | 1 12E-07 |
| ANG | 0 014658011 | -0 01656973 | 7 83E-08 | -0 47920322 | 0 | 1 12E-07 |
| RNF220 | 0 164245874 | -0 0165867 | 7 59E-08 | 0 269962707 | 0 | 1 09E-07 |
| FAM214B | 0 026079839 | -0 01661187 | 7 25E-08 | 0 463365503 | 0 | 1 04E-07 |
| PROS1 | 0 249919095 | -0 01661902 | 7 16E-08 | -0 23183315 | 0 | 1 03E-07 |
| CDC20 | 0 02160629 | -0 01663071 | 7 01E-08 | 0 43797241 | 0 | 1 01E-07 |
| RGS10 | 0 088262169 | -0 0167 | 6 19E-08 | -0 41674411 | 0 | 8 89E-08 |
| PLEKHO1 | 0 120490758 | -0 01670173 | 6 17E-08 | 0 37764928 | 0 | 8 87E-08 |
| ASB2 | 0 002465211 | -0 01670276 | 6 16E-08 | 0 492388102 | 0 | 8 85E-08 |
| RPS6KA1 | 0 0467819 | -0 01672494 | 5 91E-08 | 0 436685826 | 0 | 8 51E-08 |
| PRCP | 0 147389161 | -0 01673887 | 5 77E-08 | 0 339513059 | 0 | 8 30E-08 |
| FCER1G | 0 022596181 | -0 01676919 | 5 46E-08 | 0 463076747 | 0 | 7 86E-08 |
| GLYATL2 | 0 147988807 | -0 01682998 | 4 88E-08 | 0 268896276 | 0 | 7 04E-08 |

| | | | | | | |
|---|---|---|---|---|---|---|
| S100A9 | 0 009061316 | -0 0168572 | 4 65E-08 | 0 481570152 | 0 | 6 71E-08 |
| PACSIN2 | 0 388323085 | -0 01693811 | 4 01E-08 | -0 15177963 | 0 | 5 79E-08 |
| TLR6 | 0 014105956 | -0 01694003 | 3 99E-08 | -0 48220972 | 0 | 5 77E-08 |
| SLC25A5 | 0 104433573 | -0 01695539 | 3 88E-08 | 0 39756147 | 0 | 5 61E-08 |
| HYAL2 | 0 079353144 | -0 01697517 | 3 74E-08 | -0 40863652 | 0 | 5 41E-08 |
| IGHM | 0 003931012 | -0 01700013 | 3 58E-08 | 0 490121788 | 0 | 5 17E-08 |
| COX7A2 | 0 226723268 | -0 01701445 | 3 48E-08 | -0 28595515 | 0 | 5 04E-08 |
| RCAN1 | 0 170584988 | -0 01702514 | 3 41E-08 | -0 32029286 | 0 | 4 94E-08 |
| OLA1 | 0 438626716 | -0 017039 | 3 33E-08 | 0 105419184 | 2 62E-257 | 4 82E-08 |
| RHOG | 0 05019893 | -0 01709124 | 3 02E-08 | -0 44833444 | 0 | 4 38E-08 |
| SUMO2 | 0 351906493 | -0 01709662 | 2 99E-08 | 0 196348967 | 0 | 4 33E-08 |
| KLF7 | 0 451000362 | -0 01710719 | 2 93E-08 | -0 15263712 | 0 | 4 25E-08 |
| PHLDA3 | 0 15801146 | -0 01710756 | 2 93E-08 | -0 31202876 | 0 | 4 25E-08 |
| HSPG2 | 0 381289144 | -0 01712354 | 2 85E-08 | -0 1810588 | 0 | 4 13E-08 |
| AK5 | 0 058774819 | -0 01716678 | 2 63E-08 | -0 42954188 | 0 | 3 81E-08 |
| SOCS3 | 0 13335935 | -0 01724178 | 2 28E-08 | -0 34893039 | 0 | 3 32E-08 |
| CTTNBP2NL | 0 244113 | -0 01724343 | 2 28E-08 | 0 249578899 | 0 | 3 31E-08 |
| SERPINB8 | 0 022605699 | -0 01727398 | 2 15E-08 | -0 475157 | 0 | 3 13E-08 |
| PRR7-AS1 | 0 016180922 | -0 01735517 | 1 85E-08 | 0 471081273 | 0 | 2 69E-08 |
| IGSF8 | 0 060316765 | -0 01736841 | 1 80E-08 | -0 42535457 | 0 | 2 63E-08 |
| IL1B | 0 008975652 | -0 01737569 | 1 78E-08 | 0 479739786 | 0 | 2 59E-08 |
| NDRG2 | 0 158268451 | -0 0173887 | 1 73E-08 | -0 31701339 | 0 | 2 53E-08 |
| HLA-DQB1 | 0 04578249 | -0 01742116 | 1 63E-08 | -0 45288366 | 0 | 2 38E-08 |
| SCPEP1 | 0 20558337 | -0 01747681 | 1 47E-08 | -0 30927063 | 0 | 2 14E-08 |
| PDCL3 | 0 037178047 | -0 01747727 | 1 47E-08 | -0 45874788 | 0 | 2 14E-08 |
| EHF | 0 003331366 | -0 01748708 | 1 44E-08 | -0 49374259 | 0 | 2 10E-08 |
| APCDD1L-DT | 0 117092764 | -0 01754671 | 1 29E-08 | 0 321318253 | 0 | 1 88E-08 |
| BAALC-AS1 | 0 188279302 | -0 01761692 | 1 13E-08 | 0 280908084 | 0 | 1 65E-08 |
| CYCS | 0 146979879 | -0 01763323 | 1 09E-08 | -0 34841676 | 0 | 1 60E-08 |
| CST2 | 0 004026194 | -0 01765209 | 1 05E-08 | -0 49316375 | 0 | 1 54E-08 |
| C1QC | 0 038872285 | -0 01766143 | 1 03E-08 | 0 440958825 | 0 | 1 52E-08 |
| PER3 | 0 307780168 | -0 01768181 | 9 95E-09 | -0 22584726 | 0 | 1 46E-08 |
| UXT | 0 114332489 | -0 01768636 | 9 86E-09 | 0 380807314 | 0 | 1 45E-08 |
| HPGD | 0 042279797 | -0 01769833 | 9 64E-09 | -0 44697047 | 0 | 1 42E-08 |
| HSPB8 | 0 027307685 | -0 01776776 | 8 44E-09 | 0 459587515 | 0 | 1 24E-08 |
| WWC1 | 0 081437627 | -0 01778844 | 8 11E-09 | -0 40378656 | 0 | 1 19E-08 |
| TFAP2A | 0 065637433 | -0 01779794 | 7 96E-09 | -0 41486246 | 0 | 1 17E-08 |
| ALDOA | 0 035645619 | -0 01780886 | 7 80E-09 | -0 462574 | 0 | 1 15E-08 |
| PHLDB1 | 0 422055548 | -0 01781196 | 7 75E-09 | -0 12162214 | 0 | 1 14E-08 |
| CELF2 | 0 595153338 | -0 01782653 | 7 53E-09 | -0 00832029 | 0 00699902 | 1 11E-08 |
| PRDM8 | 0 084321639 | -0 01786145 | 7 04E-09 | 0 35087474 | 0 | 1 04E-08 |
| FN1 | 0 755411091 | -0 01786722 | 6 97E-09 | -0 07878502 | 2 80E-144 | 1 03E-08 |
| OAS2 | 0 030058442 | -0 01787119 | 6 91E-09 | 0 44918185 | 0 | 1 02E-08 |
| ZFP36 | 0 209657155 | -0 0178993 | 6 55E-09 | -0 25994306 | 0 | 9 66E-09 |

| | | | | | | |
|---|---|---|---|---|---|---|
| HNRNPK | 0 469684567 | -0 01790861 | 6 43E-09 | 0 125426616 | 0 | 9 49E-09 |
| APOL3 | 0 048190592 | -0 01791153 | 6 39E-09 | -0 44937057 | 0 | 9 43E-09 |
| ARF6 | 0 081561364 | -0 01791252 | 6 38E-09 | 0 415925519 | 0 | 9 42E-09 |
| SNU13 | 0 196512535 | -0 01793139 | 6 15E-09 | -0 31610722 | 0 | 9 08E-09 |
| CYSTM1 | 0 35003141 | -0 01794736 | 5 97E-09 | -0 18491818 | 0 | 8 81E-09 |
| RPL10A | 0 387790067 | -0 01796849 | 5 73E-09 | -0 20211021 | 0 | 8 46E-09 |
| DDX21 | 0 258314138 | -0 0179726 | 5 68E-09 | -0 26183447 | 0 | 8 39E-09 |
| RDH5 | 0 068464335 | -0 01801117 | 5 27E-09 | 0 386654803 | 0 | 7 79E-09 |
| RAB7A | 0 561249548 | -0 01803651 | 5 02E-09 | -0 0559333 | 1 44E-73 | 7 41E-09 |
| MROCKI | 0 004083303 | -0 01809561 | 4 47E-09 | -0 4925352 | 0 | 6 62E-09 |
| ODF3B | 0 0238621 | -0 01811241 | 4 33E-09 | -0 47417012 | 0 | 6 40E-09 |
| C3 | 0 071814738 | -0 01813289 | 4 16E-09 | -0 41723641 | 0 | 6 16E-09 |
| CST4 | 0 002170147 | -0 01814607 | 4 05E-09 | -0 4955688 | 0 | 6 00E-09 |
| HLA-DPB1 | 0 113409225 | -0 01817221 | 3 85E-09 | -0 39744738 | 0 | 5 70E-09 |
| SPTBN5 | 0 016095258 | -0 01818922 | 3 72E-09 | 0 473913215 | 0 | 5 52E-09 |
| ITGA4 | 0 154318402 | -0 01827582 | 3 14E-09 | 0 243110116 | 0 | 4 66E-09 |
| TRHDE-AS1 | 0 031333879 | -0 01833394 | 2 80E-09 | 0 437435688 | 0 | 4 16E-09 |
| ADH1C | 0 032314253 | -0 01835502 | 2 68E-09 | -0 45701202 | 0 | 3 99E-09 |
| MICOS10 | 0 186461328 | -0 01836467 | 2 63E-09 | 0 316328412 | 0 | 3 92E-09 |
| ICAM2 | 0 029001923 | -0 01838172 | 2 54E-09 | -0 46799014 | 0 | 3 79E-09 |
| CALM2 | 0 501180255 | -0 01840906 | 2 41E-09 | 0 098289844 | 8 47E-224 | 3 59E-09 |
| ME3 | 0 16463612 | -0 01842151 | 2 35E-09 | -0 32536466 | 0 | 3 50E-09 |
| KLK10 | 0 00872818 | -0 01843028 | 2 31E-09 | -0 48615328 | 0 | 3 44E-09 |
| NAV2 | 0 360111172 | -0 0184464 | 2 24E-09 | 0 14261755 | 0 | 3 34E-09 |
| CSTF3-DT | 0 023985837 | -0 01849899 | 2 02E-09 | 0 439369936 | 0 | 3 01E-09 |
| ERP29 | 0 187822429 | -0 01853811 | 1 86E-09 | 0 306099774 | 0 | 2 78E-09 |
| SCIN | 0 027564676 | -0 01856091 | 1 78E-09 | -0 4624622 | 0 | 2 66E-09 |
| ECPAS | 0 564780796 | -0 01856454 | 1 77E-09 | 0 070606992 | 3 34E-116 | 2 64E-09 |
| SAXO1 | 0 104804782 | -0 01858477 | 1 70E-09 | 0 339109438 | 0 | 2 54E-09 |
| CBLN4 | 0 022986427 | -0 0185919 | 1 67E-09 | -0 46736766 | 0 | 2 50E-09 |
| FBXO32 | 0 248872095 | -0 01863033 | 1 55E-09 | -0 22592187 | 0 | 2 32E-09 |
| SMDT1 | 0 07784927 | -0 01868031 | 1 40E-09 | -0 42470982 | 0 | 2 10E-09 |
| HERPUD1 | 0 222687556 | -0 01869601 | 1 36E-09 | -0 26999846 | 0 | 2 03E-09 |
| ABCA7 | 0 026117911 | -0 01871313 | 1 31E-09 | 0 456296702 | 0 | 1 96E-09 |
| DYNLL2 | 0 105899374 | -0 01872782 | 1 27E-09 | -0 39249447 | 0 | 1 91E-09 |
| ASAH1 | 0 190639813 | -0 01874235 | 1 24E-09 | -0 29974926 | 0 | 1 85E-09 |
| BCAP31 | 0 167758086 | -0 01876033 | 1 19E-09 | -0 33521534 | 0 | 1 79E-09 |
| IER2 | 0 220022463 | -0 01876751 | 1 17E-09 | 0 25945988 | 0 | 1 76E-09 |
| DAPL1 | 0 016114294 | -0 01881708 | 1 06E-09 | -0 47598265 | 0 | 1 59E-09 |
| GPC1 | 0 112371742 | -0 01881776 | 1 06E-09 | -0 37822721 | 0 | 1 59E-09 |
| SERPING1 | 0 240657897 | -0 0188699 | 9 55E-10 | -0 25726306 | 0 | 1 43E-09 |
| OSMR | 0 353743504 | -0 01888833 | 9 19E-10 | -0 18642177 | 0 | 1 38E-09 |
| C9orf16 | 0 026708039 | -0 01890924 | 8 81E-10 | 0 455228905 | 0 | 1 32E-09 |
| TNFRSF11B | 0 011602673 | -0 01892016 | 8 62E-10 | -0 48013944 | 0 | 1 30E-09 |

| | | | | | | |
|---|---|---|---|---|---|---|
| LINC01877 | 0 021349299 | -0 0189465 | 8 17E-10 | -0 46891347 | 0 | 1 23E-09 |
| PCDH1 | 0 013630047 | -0 01896146 | 7 92E-10 | -0 48112016 | 0 | 1 19E-09 |
| CXCL3 | 0 013553902 | -0 01905346 | 6 56E-10 | 0 466952283 | 0 | 9 88E-10 |
| MKI67 | 0 094287183 | -0 01905636 | 6 52E-10 | 0 319942566 | 0 | 9 83E-10 |
| PRR11 | 0 074032476 | -0 01916762 | 5 19E-10 | 0 33470691 | 0 | 7 83E-10 |
| TREM2 | 0 00693876 | -0 01916836 | 5 18E-10 | -0 49163929 | 0 | 7 82E-10 |
| CKAP2L | 0 0580324 | -0 01919376 | 4 91E-10 | 0 346668725 | 0 | 7 42E-10 |
| CASP4 | 0 116654928 | -0 01919779 | 4 87E-10 | -0 37206831 | 0 | 7 36E-10 |
| DSG3 | 0 005025604 | -0 0192038 | 4 81E-10 | 0 489655896 | 0 | 7 27E-10 |
| SBDS | 0 149292799 | -0 01923952 | 4 47E-10 | -0 34778882 | 0 | 6 76E-10 |
| GALNT2 | 0 506434296 | -0 01933407 | 3 67E-10 | -0 09630286 | 6 95E-215 | 5 56E-10 |
| FJX1 | 0 041337496 | -0 01933581 | 3 66E-10 | -0 45716796 | 0 | 5 54E-10 |
| S100A8 | 0 004444994 | -0 01934567 | 3 59E-10 | -0 49387079 | 0 | 5 43E-10 |
| IL16 | 0 096647694 | -0 01935544 | 3 51E-10 | -0 37706853 | 0 | 5 32E-10 |
| PI16 | 0 16053378 | -0 0193997 | 3 20E-10 | -0 32490677 | 0 | 4 86E-10 |
| GIMAP2 | 0 016342731 | -0 01944516 | 2 91E-10 | -0 47972484 | 0 | 4 42E-10 |
| SOAT1 | 0 236688812 | -0 01947438 | 2 74E-10 | 0 264235368 | 0 | 4 16E-10 |
| PDE2A | 0 010203499 | -0 01949144 | 2 64E-10 | 0 479403555 | 0 | 4 01E-10 |
| MT1A | 0 00672936 | -0 01951409 | 2 52E-10 | -0 48920618 | 0 | 3 83E-10 |
| ATP1B3 | 0 28905789 | -0 0195197 | 2 49E-10 | 0 223711959 | 0 | 3 79E-10 |
| KLF13 | 0 138603872 | -0 0195516 | 2 33E-10 | -0 33228852 | 0 | 3 54E-10 |
| MCOLN2 | 0 032361843 | -0 01959339 | 2 13E-10 | 0 445831128 | 0 | 3 25E-10 |
| SEC11C | 0 088642897 | -0 01963154 | 1 97E-10 | -0 40351741 | 0 | 3 00E-10 |
| TLN1 | 0 2097333 | -0 01964416 | 1 92E-10 | -0 30342544 | 0 | 2 92E-10 |
| GAB2 | 0 499762045 | -0 01966167 | 1 85E-10 | -0 11124694 | 1 76E-286 | 2 81E-10 |
| OLR1 | 0 008109497 | -0 01966347 | 1 84E-10 | -0 48942491 | 0 | 2 80E-10 |
| VSIR | 0 075802859 | -0 01969287 | 1 73E-10 | -0 40171173 | 0 | 2 64E-10 |
| RPL21 | 0 488083227 | -0 01977523 | 1 45E-10 | 0 084379868 | 2 83E-165 | 2 22E-10 |
| MCL1 | 0 449020578 | -0 01978423 | 1 42E-10 | -0 10890452 | 1 46E-274 | 2 18E-10 |
| STXBP5L | 0 073489939 | -0 01979802 | 1 38E-10 | -0 41104679 | 0 | 2 11E-10 |
| MAN2A1 | 0 489691801 | -0 01981114 | 1 35E-10 | -0 10052762 | 4 65E-234 | 2 06E-10 |
| TRABD2A | 0 036882983 | -0 01982475 | 1 31E-10 | -0 45935212 | 0 | 2 00E-10 |
| KMT2A | 0 499400354 | -0 01982672 | 1 30E-10 | 0 087129431 | 3 98E-176 | 1 99E-10 |
| ATP6V0E2 | 0 048266738 | -0 01983057 | 1 29E-10 | -0 44217438 | 0 | 1 97E-10 |
| ALDH1A3 | 0 349936228 | -0 01984886 | 1 24E-10 | -0 21418281 | 0 | 1 90E-10 |
| NDUFB2 | 0 22208791 | -0 01989208 | 1 13E-10 | 0 288550428 | 0 | 1 73E-10 |
| IFIH1 | 0 090898707 | -0 01989278 | 1 13E-10 | 0 380124738 | 0 | 1 73E-10 |
| FCGBP | 0 030458206 | -0 0198992 | 1 11E-10 | 0 456217936 | 0 | 1 71E-10 |
| GNLY | 0 010717481 | -0 01994143 | 1 02E-10 | 0 475170532 | 0 | 1 56E-10 |
| CPXM2 | 0 134587196 | -0 01994242 | 1 02E-10 | -0 34894582 | 0 | 1 56E-10 |
| RAB38 | 0 031895452 | -0 01996584 | 9 66E-11 | -0 46630784 | 0 | 1 48E-10 |
| QPCTL | 0 040471341 | -0 02000615 | 8 86E-11 | 0 449106619 | 0 | 1 36E-10 |
| ATF4 | 0 240962479 | -0 02006025 | 7 89E-11 | -0 27661685 | 0 | 1 21E-10 |
| DCXR | 0 094896347 | -0 02006849 | 7 75E-11 | 0 412318369 | 0 | 1 19E-10 |

| | | | | | | |
|---|---|---|---|---|---|---|
| TUBA1C | 0 167643867 | -0 02010349 | 7 18E-11 | 0 254851748 | 0 | 1 11E-10 |
| BEST1 | 0 026879366 | -0 02017041 | 6 21E-11 | -0 46462535 | 0 | 9 58E-11 |
| CTNNA1 | 0 654394548 | -0 02018012 | 6 09E-11 | -0 0331658 | 5 74E-27 | 9 38E-11 |
| ZNF708 | 0 279530182 | -0 02019917 | 5 84E-11 | 0 239584399 | 0 | 9 01E-11 |
| TKT | 0 142201748 | -0 02022359 | 5 54E-11 | -0 35773903 | 0 | 8 55E-11 |
| CH25H | 0 023214864 | -0 02028338 | 4 86E-11 | -0 4752983 | 0 | 7 52E-11 |
| ADM | 0 07574575 | -0 02035178 | 4 19E-11 | -0 41511462 | 0 | 6 48E-11 |
| COX17 | 0 127439036 | -0 02035588 | 4 15E-11 | -0 37149013 | 0 | 6 43E-11 |
| PLEKHO2 | 0 053701624 | -0 02037736 | 3 96E-11 | -0 43375425 | 0 | 6 14E-11 |
| KPNA2 | 0 114370562 | -0 02037739 | 3 96E-11 | 0 313139383 | 0 | 6 14E-11 |
| TAP2 | 0 079933753 | -0 02040853 | 3 70E-11 | 0 386376773 | 0 | 5 74E-11 |
| RARRES2 | 0 14124041 | -0 02052402 | 2 87E-11 | -0 35779739 | 0 | 4 46E-11 |
| PKIA | 0 165978184 | -0 0205716 | 2 58E-11 | 0 260571257 | 0 | 4 02E-11 |
| RPL7 | 0 394357617 | -0 02058795 | 2 49E-11 | 0 153746848 | 0 | 3 88E-11 |
| CTSF | 0 107279511 | -0 02059073 | 2 48E-11 | -0 37789945 | 0 | 3 85E-11 |
| FTCDNL1 | 0 05871771 | -0 02059077 | 2 48E-11 | -0 43848558 | 0 | 3 85E-11 |
| EPS15 | 0 580238335 | -0 02066113 | 2 12E-11 | -0 07695319 | 1 01E-137 | 3 31E-11 |
| ADPRH | 0 012402201 | -0 02067244 | 2 07E-11 | -0 4820553 | 0 | 3 22E-11 |
| UBB | 0 24595001 | -0 02068582 | 2 01E-11 | -0 26619216 | 0 | 3 13E-11 |
| CCL2 | 0 11558889 | -0 02070671 | 1 91E-11 | -0 3796134 | 0 | 2 99E-11 |
| RGCC | 0 154004302 | -0 02070864 | 1 91E-11 | -0 325542 | 0 | 2 98E-11 |
| DENND3 | 0 049446993 | -0 02071452 | 1 88E-11 | 0 434035014 | 0 | 2 94E-11 |
| APP | 0 728655461 | -0 02073587 | 1 79E-11 | -0 04643272 | 3 04E-51 | 2 81E-11 |
| B3GAT1 | 0 011783518 | -0 02074467 | 1 76E-11 | -0 48219527 | 0 | 2 75E-11 |
| C12orf57 | 0 210951629 | -0 02078838 | 1 60E-11 | 0 268210756 | 0 | 2 50E-11 |
| SLC9A7 | 0 125173707 | -0 02079961 | 1 56E-11 | -0 37610064 | 0 | 2 44E-11 |
| LY6K | 0 023643182 | -0 02080913 | 1 52E-11 | -0 4760699 | 0 | 2 39E-11 |
| CYBB | 0 028145286 | -0 02081244 | 1 51E-11 | 0 454072451 | 0 | 2 37E-11 |
| DTHD1 | 0 00531115 | -0 02081581 | 1 50E-11 | -0 49199045 | 0 | 2 35E-11 |
| STX12 | 0 35896899 | -0 02082126 | 1 48E-11 | -0 18706765 | 0 | 2 32E-11 |
| APRT | 0 113228379 | -0 02089064 | 1 27E-11 | -0 38975953 | 0 | 1 99E-11 |
| LINC00702 | 0 019350479 | -0 02091069 | 1 21E-11 | 0 468474424 | 0 | 1 91E-11 |
| PPP1R1A | 0 009004207 | -0 02109079 | 8 09E-12 | -0 4874095 | 0 | 1 27E-11 |
| OAZ1 | 0 28889608 | -0 0211191 | 7 59E-12 | -0 24886516 | 0 | 1 20E-11 |
| CMTM6 | 0 197178809 | -0 02113379 | 7 34E-12 | -0 31835586 | 0 | 1 16E-11 |
| L1TD1 | 0 056709372 | -0 02115522 | 6 99E-12 | -0 42897507 | 0 | 1 10E-11 |
| BVES | 0 055129352 | -0 02121794 | 6 06E-12 | 0 424877812 | 0 | 9 56E-12 |
| PLAA | 0 168852677 | -0 02124757 | 5 67E-12 | 0 313068934 | 0 | 8 95E-12 |
| PFDN2 | 0 187489292 | -0 02124808 | 5 66E-12 | -0 32303459 | 0 | 8 94E-12 |
| REEP5 | 0 30466772 | -0 02126604 | 5 44E-12 | -0 21625279 | 0 | 8 58E-12 |
| KANK4 | 0 184966972 | -0 02127833 | 5 29E-12 | -0 28715238 | 0 | 8 34E-12 |
| ELOVL7 | 0 047105519 | -0 02132232 | 4 78E-12 | 0 452203097 | 0 | 7 55E-12 |
| POU3F1 | 0 015724049 | -0 02134105 | 4 58E-12 | -0 4801842 | 0 | 7 24E-12 |
| KIF1B | 0 607926748 | -0 0214182 | 3 84E-12 | -0 05122835 | 5 38E-62 | 6 07E-12 |

| | | | | | | |
|---|---|---|---|---|---|---|
| GADD45B | 0 115703109 | -0 02145892 | 3 49E-12 | -0 37179833 | 0 | 5 53E-12 |
| NUP50-DT | 0 169804496 | -0 02147132 | 3 40E-12 | 0 273839377 | 0 | 5 38E-12 |
| PDIA4 | 0 223953475 | -0 02160427 | 2 50E-12 | -0 28741664 | 0 | 3 97E-12 |
| LIPC | 0 03190497 | -0 02160526 | 2 49E-12 | 0 434829772 | 0 | 3 96E-12 |
| ABL2 | 0 481401458 | -0 02160949 | 2 47E-12 | -0 12855026 | 0 | 3 92E-12 |
| AQP1 | 0 112581143 | -0 021641 | 2 29E-12 | -0 35775176 | 0 | 3 65E-12 |
| TMEM52B | 0 005139822 | -0 02164716 | 2 26E-12 | 0 48289858 | 0 | 3 60E-12 |
| ARPC2 | 0 431021682 | -0 02164858 | 2 25E-12 | -0 15547505 | 0 | 3 59E-12 |
| PMAIP1 | 0 066893834 | -0 02169864 | 2 01E-12 | 0 372595212 | 0 | 3 20E-12 |
| GRAMD2B | 0 297157869 | -0 02174308 | 1 81E-12 | -0 19261925 | 0 | 2 89E-12 |
| ACER3 | 0 334136034 | -0 02176222 | 1 73E-12 | 0 192487874 | 0 | 2 76E-12 |
| RASD1 | 0 02150159 | -0 02177453 | 1 68E-12 | -0 47176606 | 0 | 2 69E-12 |
| RNF181 | 0 084464411 | -0 02184167 | 1 44E-12 | -0 41667365 | 0 | 2 30E-12 |
| HDAC9 | 0 38151758 | -0 02189867 | 1 26E-12 | 0 190375549 | 0 | 2 01E-12 |
| RAB27B | 0 050636767 | -0 02196462 | 1 08E-12 | 0 408780339 | 0 | 1 73E-12 |
| RELL1 | 0 193038396 | -0 02196611 | 1 07E-12 | 0 306546112 | 0 | 1 72E-12 |
| TYROBP | 0 039833622 | -0 02197581 | 1 05E-12 | 0 442484901 | 0 | 1 68E-12 |
| CKS2 | 0 073518494 | -0 02203884 | 9 05E-13 | 0 344278759 | 0 | 1 45E-12 |
| NFE2L2 | 0 524423674 | -0 02209905 | 7 85E-13 | -0 09312809 | 5 02E-201 | 1 26E-12 |
| LRRC23 | 0 142420666 | -0 02212757 | 7 33E-13 | 0 290992087 | 0 | 1 18E-12 |
| NCKAP5-AS2 | 0 044116807 | -0 02212974 | 7 30E-13 | -0 4471581 | 0 | 1 18E-12 |
| BUB3 | 0 199453656 | -0 02215794 | 6 82E-13 | 0 238882601 | 0 | 1 10E-12 |
| TUBB2A | 0 071357865 | -0 0221679 | 6 67E-13 | -0 41200812 | 0 | 1 07E-12 |
| GLUL | 0 341398412 | -0 02219037 | 6 32E-13 | -0 16313355 | 0 | 1 02E-12 |
| TUT7 | 0 401401077 | -0 02220793 | 6 06E-13 | 0 136190537 | 0 | 9 78E-13 |
| GNPTAB | 0 308227523 | -0 02230072 | 4 86E-13 | -0 21417328 | 0 | 7 84E-13 |
| MIR22HG | 0 107174811 | -0 02235266 | 4 29E-13 | -0 38123358 | 0 | 6 93E-13 |
| RPL5 | 0 439331062 | -0 02247894 | 3 17E-13 | -0 16130511 | 0 | 5 14E-13 |
| IL1RAP | 0 232767318 | -0 02253079 | 2 80E-13 | 0 255733074 | 0 | 4 54E-13 |
| TBCA | 0 53238088 | -0 0225339 | 2 78E-13 | 0 094824514 | 2 26E-208 | 4 51E-13 |
| ARFGEF3 | 0 116455046 | -0 02255565 | 2 63E-13 | -0 3710092 | 0 | 4 28E-13 |
| CITED2 | 0 071091356 | -0 02255614 | 2 63E-13 | -0 41903721 | 0 | 4 27E-13 |
| CDK2AP2 | 0 04946603 | -0 02256243 | 2 59E-13 | -0 43928275 | 0 | 4 21E-13 |
| PANX1 | 0 051712322 | -0 02258561 | 2 45E-13 | -0 44560014 | 0 | 3 98E-13 |
| CCL4 | 0 023890655 | -0 02261109 | 2 30E-13 | -0 47591184 | 0 | 3 75E-13 |
| PTBP3 | 0 435894995 | -0 02263602 | 2 17E-13 | 0 129343619 | 0 | 3 53E-13 |
| EBP | 0 060307247 | -0 02267813 | 1 96E-13 | 0 410908715 | 0 | 3 19E-13 |
| TMEM154 | 0 089918334 | -0 02269741 | 1 87E-13 | 0 376648989 | 0 | 3 05E-13 |
| PLTP | 0 126334926 | -0 02272475 | 1 75E-13 | -0 35669299 | 0 | 2 85E-13 |
| CD151 | 0 172012716 | -0 02273861 | 1 69E-13 | -0 33571812 | 0 | 2 76E-13 |
| DHH | 0 008804325 | -0 02274643 | 1 66E-13 | -0 48558236 | 0 | 2 71E-13 |
| IFITM1 | 0 098951096 | -0 02276095 | 1 60E-13 | -0 3904493 | 0 | 2 61E-13 |
| EDF1 | 0 254145171 | -0 02276251 | 1 60E-13 | -0 27384578 | 0 | 2 60E-13 |
| SMIM3 | 0 096124193 | -0 02276707 | 1 58E-13 | -0 40309031 | 0 | 2 58E-13 |

| | | | | | | |
|---|---|---|---|---|---|---|
| NAMPT | 0 287715825 | -0 02280399 | 1 44E-13 | -0 25495113 | 0 | 2 36E-13 |
| LINC00944 | 0 08410272 | -0 02286483 | 1 24E-13 | 0 352965835 | 0 | 2 03E-13 |
| SMTN | 0 185576136 | -0 02286967 | 1 23E-13 | -0 31394953 | 0 | 2 01E-13 |
| LDHB | 0 231786945 | -0 02288922 | 1 17E-13 | 0 287849201 | 0 | 1 92E-13 |
| LRRC63 | 0 030106033 | -0 02289263 | 1 16E-13 | 0 456643527 | 0 | 1 90E-13 |
| ZNF503 | 0 064999714 | -0 02289641 | 1 15E-13 | -0 41380079 | 0 | 1 88E-13 |
| BMP3 | 0 011212427 | -0 02297833 | 9 41E-14 | -0 48394385 | 0 | 1 54E-13 |
| LY6E | 0 178856294 | -0 02298774 | 9 20E-14 | -0 33791187 | 0 | 1 51E-13 |
| EML5 | 0 036121528 | -0 02303292 | 8 23E-14 | -0 4599086 | 0 | 1 35E-13 |
| STARD13-AS | 0 329224648 | -0 02303597 | 8 17E-14 | -0 23226913 | 0 | 1 34E-13 |
| DYNLL1 | 0 31712703 | -0 02310561 | 6 88E-14 | -0 20698913 | 0 | 1 13E-13 |
| PPT1 | 0 142839466 | -0 02313202 | 6 44E-14 | -0 36268722 | 0 | 1 06E-13 |
| APOC2 | 0 007024424 | -0 02317226 | 5 83E-14 | -0 49022115 | 0 | 9 59E-14 |
| CLEC11A | 0 185062154 | -0 0232305 | 5 05E-14 | -0 32603546 | 0 | 8 31E-14 |
| SPAAR | 0 03753022 | -0 02326029 | 4 69E-14 | -0 45413729 | 0 | 7 72E-14 |
| ADTRP | 0 010688927 | -0 0232618 | 4 67E-14 | -0 48858679 | 0 | 7 69E-14 |
| NTN4 | 0 221907064 | -0 02329497 | 4 30E-14 | 0 265627066 | 0 | 7 09E-14 |
| RELB | 0 110639432 | -0 02343995 | 2 99E-14 | 0 362224815 | 0 | 4 95E-14 |
| NFKB2 | 0 054348861 | -0 02344956 | 2 92E-14 | 0 427883422 | 0 | 4 83E-14 |
| LINC02257 | 0 012326055 | -0 02345621 | 2 87E-14 | -0 48372349 | 0 | 4 75E-14 |
| MARCKSL1 | 0 119805448 | -0 0234933 | 2 62E-14 | -0 36914031 | 0 | 4 33E-14 |
| GALNT6 | 0 015448021 | -0 02349953 | 2 58E-14 | 0 464087734 | 0 | 4 27E-14 |
| ITPK1-AS1 | 0 015210066 | -0 02350485 | 2 54E-14 | -0 47901596 | 0 | 4 21E-14 |
| MAN2B1 | 0 155251185 | -0 02353673 | 2 35E-14 | 0 299780952 | 0 | 3 89E-14 |
| MT-ND6 | 0 167034703 | -0 02359906 | 2 01E-14 | -0 34042871 | 0 | 3 33E-14 |
| P4HB | 0 463402562 | -0 02362755 | 1 87E-14 | 0 122999125 | 0 | 3 10E-14 |
| RGS17 | 0 190144867 | -0 02362795 | 1 87E-14 | 0 260678676 | 0 | 3 10E-14 |
| CDCP1 | 0 01918867 | -0 02363842 | 1 82E-14 | -0 47833272 | 0 | 3 02E-14 |
| CHML | 0 075517314 | -0 02365032 | 1 76E-14 | -0 41348626 | 0 | 2 93E-14 |
| GYPC | 0 1513868 | -0 02365612 | 1 74E-14 | -0 34358097 | 0 | 2 89E-14 |
| CAPS | 0 082855838 | -0 02367375 | 1 66E-14 | -0 39097562 | 0 | 2 76E-14 |
| CIB1 | 0 111162932 | -0 02371851 | 1 48E-14 | 0 37457607 | 0 | 2 47E-14 |
| CSNK1A1 | 0 678294721 | -0 02372507 | 1 46E-14 | -0 05101171 | 1 74E-61 | 2 43E-14 |
| NLRC5 | 0 140878719 | -0 0238513 | 1 06E-14 | 0 285708833 | 0 | 1 77E-14 |
| EVI2A | 0 022539072 | -0 02386323 | 1 03E-14 | 0 455312023 | 0 | 1 71E-14 |
| PGK1 | 0 293550475 | -0 02389304 | 9 52E-15 | -0 25026784 | 0 | 1 59E-14 |
| CLTC | 0 634349241 | -0 02389903 | 9 37E-15 | 0 025398936 | 1 81E-16 | 1 57E-14 |
| ROMO1 | 0 127372409 | -0 02390862 | 9 15E-15 | -0 3840086 | 0 | 1 53E-14 |
| GABARAPL2 | 0 177666521 | -0 02393158 | 8 63E-15 | -0 32133481 | 0 | 1 44E-14 |
| MPZ | 0 020521216 | -0 02396426 | 7 93E-15 | -0 47525978 | 0 | 1 33E-14 |
| YBX3 | 0 535807428 | -0 02398488 | 7 53E-15 | -0 10675676 | 7 31E-264 | 1 26E-14 |
| SELPLG | 0 008690107 | -0 0240326 | 6 66E-15 | -0 48888089 | 0 | 1 12E-14 |
| GCNT2 | 0 037653957 | -0 02404945 | 6 38E-15 | 0 429764193 | 0 | 1 07E-14 |
| ACVR1C | 0 029810969 | -0 02405461 | 6 29E-15 | 0 438611201 | 0 | 1 06E-14 |

| | | | | | | |
|---|---|---|---|---|---|---|
| MAP2K3 | 0 081142563 | -0 02406838 | 6 07E-15 | -0 40865837 | 0 | 1 02E-14 |
| ACTR3C | 0 030220251 | -0 02408423 | 5 83E-15 | 0 446534278 | 0 | 9 79E-15 |
| H3-3A | 0 195084807 | -0 02409562 | 5 66E-15 | -0 30018888 | 0 | 9 51E-15 |
| RPL4 | 0 352639394 | -0 02409672 | 5 65E-15 | 0 175635629 | 0 | 9 48E-15 |
| MOSPD2 | 0 218785098 | -0 02413706 | 5 09E-15 | -0 28673547 | 0 | 8 56E-15 |
| NDP | 0 013411129 | -0 02414452 | 4 99E-15 | -0 4820062 | 0 | 8 39E-15 |
| SHISA3 | 0 051112676 | -0 02415821 | 4 82E-15 | -0 45054667 | 0 | 8 11E-15 |
| LRMDA | 0 595619729 | -0 0241864 | 4 48E-15 | -0 07809361 | 8 74E-142 | 7 54E-15 |
| SPCS1 | 0 184481544 | -0 02419143 | 4 42E-15 | -0 31853482 | 0 | 7 45E-15 |
| THEMIS2 | 0 022881727 | -0 02419716 | 4 36E-15 | 0 459231894 | 0 | 7 34E-15 |
| C3orf70 | 0 168833641 | -0 02421019 | 4 21E-15 | 0 353677717 | 0 | 7 10E-15 |
| GNG11 | 0 134149359 | -0 02426754 | 3 63E-15 | 0 349552366 | 0 | 6 12E-15 |
| GNG2 | 0 365755459 | -0 02427194 | 3 59E-15 | -0 15225775 | 0 | 6 05E-15 |
| ZDHHC2 | 0 147779406 | -0 02434544 | 2 97E-15 | -0 34736354 | 0 | 5 01E-15 |
| STXBP5-AS1 | 0 052683178 | -0 02435156 | 2 92E-15 | 0 412812332 | 0 | 4 93E-15 |
| SYF2 | 0 141583065 | -0 02439732 | 2 59E-15 | -0 36987489 | 0 | 4 39E-15 |
| XKR4 | 0 06891169 | -0 0244406 | 2 32E-15 | -0 42498401 | 0 | 3 92E-15 |
| OSTC | 0 244484209 | -0 02447907 | 2 10E-15 | -0 28142919 | 0 | 3 55E-15 |
| YBX1 | 0 321743352 | -0 02448183 | 2 08E-15 | 0 210056942 | 0 | 3 52E-15 |
| TRPV2 | 0 025480193 | -0 02449322 | 2 02E-15 | -0 46795025 | 0 | 3 42E-15 |
| CCL3 | 0 024928138 | -0 02452899 | 1 84E-15 | -0 47401206 | 0 | 3 12E-15 |
| PPTC7 | 0 180541014 | -0 02455405 | 1 72E-15 | 0 298723689 | 0 | 2 92E-15 |
| MAMLD1 | 0 112657288 | -0 02457814 | 1 62E-15 | -0 38167654 | 0 | 2 74E-15 |
| EIF3A | 0 466191392 | -0 02459904 | 1 53E-15 | 0 10780152 | 4 86E-269 | 2 60E-15 |
| CD68 | 0 03027736 | -0 02473565 | 1 07E-15 | 0 453049898 | 0 | 1 82E-15 |
| NAV3 | 0 506710323 | -0 02474629 | 1 04E-15 | -0 13744006 | 0 | 1 77E-15 |
| DNASE2 | 0 071262683 | -0 02482164 | 8 51E-16 | 0 404706636 | 0 | 1 45E-15 |
| GBE1 | 0 439569016 | -0 02482423 | 8 45E-16 | 0 106975696 | 6 07E-265 | 1 44E-15 |
| ECHDC2 | 0 259313548 | -0 02487947 | 7 30E-16 | -0 24458923 | 0 | 1 25E-15 |
| TAMALIN | 0 060183511 | -0 02492098 | 6 54E-16 | -0 42094619 | 0 | 1 12E-15 |
| POLR2I | 0 092640536 | -0 02495605 | 5 95E-16 | -0 4072276 | 0 | 1 02E-15 |
| TACC1 | 0 628771582 | -0 02497334 | 5 69E-16 | -0 0309899 | 9 46E-24 | 9 73E-16 |
| RCAN3AS | 0 01881746 | -0 02503367 | 4 84E-16 | 0 460511846 | 0 | 8 29E-16 |
| KIAA0319 | 0 014924521 | -0 02506361 | 4 47E-16 | 0 483558942 | 0 | 7 66E-16 |
| TAOK3 | 0 46430679 | -0 02511831 | 3 86E-16 | 0 096639105 | 2 22E-216 | 6 63E-16 |
| HLA-E | 0 23081609 | -0 02513937 | 3 65E-16 | -0 28832389 | 0 | 6 27E-16 |
| SQOR | 0 046829491 | -0 02516928 | 3 37E-16 | -0 44753191 | 0 | 5 79E-16 |
| PPP1R15A | 0 14534275 | -0 02518104 | 3 26E-16 | -0 33517258 | 0 | 5 61E-16 |
| TMEM51 | 0 080904609 | -0 02526364 | 2 61E-16 | 0 377013069 | 0 | 4 50E-16 |
| C1QA | 0 069597 | -0 02526752 | 2 59E-16 | 0 402286876 | 0 | 4 45E-16 |
| EPHA5 | 0 141278483 | -0 02530223 | 2 35E-16 | 0 294514524 | 0 | 4 06E-16 |
| RAC2 | 0 018684206 | -0 02534163 | 2 12E-16 | 0 469779306 | 0 | 3 65E-16 |
| LINC01828 | 0 020140488 | -0 02536868 | 1 97E-16 | 0 45759452 | 0 | 3 40E-16 |
| ANGPTL4 | 0 112286079 | -0 02544829 | 1 59E-16 | 0 338185003 | 0 | 2 74E-16 |

| | | | | | | |
|---|---|---|---|---|---|---|
| CLIP4 | 0 422283985 | -0 02545395 | 1 56E-16 | -0 14134995 | 0 | 2 70E-16 |
| SGPP2 | 0 017684796 | -0 0254829 | 1 44E-16 | 0 462926906 | 0 | 2 50E-16 |
| NENF | 0 167700977 | -0 02548327 | 1 44E-16 | -0 34261941 | 0 | 2 50E-16 |
| PARP8 | 0 363461575 | -0 02549857 | 1 38E-16 | 0 168340132 | 0 | 2 40E-16 |
| ACSL4 | 0 305419657 | -0 02551466 | 1 32E-16 | -0 21337962 | 0 | 2 29E-16 |
| LINC02742 | 0 022186899 | -0 02553878 | 1 24E-16 | 0 452525933 | 0 | 2 15E-16 |
| ARHGEF16 | 0 010289163 | -0 02554754 | 1 21E-16 | -0 48443113 | 0 | 2 10E-16 |
| PTCHD1-AS | 0 086691668 | -0 02559261 | 1 07E-16 | 0 32718977 | 0 | 1 86E-16 |
| PEA15 | 0 094696465 | -0 02560131 | 1 05E-16 | -0 3961797 | 0 | 1 81E-16 |
| PTPRJ | 0 355228341 | -0 02560288 | 1 04E-16 | -0 19325802 | 0 | 1 81E-16 |
| ANGPTL2 | 0 141859093 | -0 02563441 | 9 55E-17 | -0 358511 | 0 | 1 66E-16 |
| UCN2 | 0 006129714 | -0 02565592 | 9 01E-17 | 0 484269455 | 0 | 1 57E-16 |
| NKAPL | 0 009232644 | -0 02577685 | 6 47E-17 | -0 48484306 | 0 | 1 13E-16 |
| LINC01684 | 0 097704213 | -0 0258323 | 5 55E-17 | 0 344048843 | 0 | 9 67E-17 |
| GPAT3 | 0 033256553 | -0 02591844 | 4 38E-17 | -0 46387149 | 0 | 7 64E-17 |
| APOBEC3C | 0 045373208 | -0 0259198 | 4 36E-17 | -0 44937342 | 0 | 7 61E-17 |
| BSG | 0 257447983 | -0 02592509 | 4 30E-17 | -0 25009161 | 0 | 7 50E-17 |
| CDK18 | 0 049332775 | -0 02593954 | 4 13E-17 | 0 435899294 | 0 | 7 21E-17 |
| CTSS | 0 068302526 | -0 0259804 | 3 69E-17 | 0 411847372 | 0 | 6 44E-17 |
| UCP2 | 0 016152367 | -0 02599953 | 3 50E-17 | 0 469895328 | 0 | 6 11E-17 |
| PPFIA1 | 0 444185338 | -0 02609922 | 2 65E-17 | 0 102608257 | 8 23E-244 | 4 64E-17 |
| ETS1 | 0 222125983 | -0 02623263 | 1 83E-17 | 0 261568977 | 0 | 3 21E-17 |
| SLC2A1 | 0 108183739 | -0 02624073 | 1 79E-17 | -0 38622737 | 0 | 3 14E-17 |
| INSC | 0 028535531 | -0 02630131 | 1 51E-17 | -0 46816822 | 0 | 2 65E-17 |
| S100A13 | 0 206506634 | -0 02631824 | 1 44E-17 | 0 273519429 | 0 | 2 52E-17 |
| VEPH1 | 0 102282462 | -0 02640419 | 1 13E-17 | 0 323002306 | 0 | 1 99E-17 |
| TESPA1 | 0 015685976 | -0 02643523 | 1 04E-17 | -0 48416288 | 0 | 1 82E-17 |
| JUND | 0 351649502 | -0 02644698 | 1 00E-17 | -0 16835143 | 0 | 1 76E-17 |
| FOSL2 | 0 194389979 | -0 02645366 | 9 83E-18 | -0 28871418 | 0 | 1 73E-17 |
| MOCOS | 0 060107365 | -0 02647819 | 9 18E-18 | 0 423588965 | 0 | 1 62E-17 |
| HLA-G | 0 003407512 | -0 02653977 | 7 71E-18 | -0 49306523 | 0 | 1 36E-17 |
| CTNNAL1 | 0 225343131 | -0 02654281 | 7 64E-18 | 0 250986555 | 0 | 1 35E-17 |
| ECE1 | 0 285393387 | -0 0265681 | 7 12E-18 | -0 22475551 | 0 | 1 25E-17 |
| GPR65 | 0 007595515 | -0 02666382 | 5 43E-18 | 0 477791562 | 0 | 9 58E-18 |
| DUSP18 | 0 06344825 | -0 02678137 | 3 88E-18 | -0 43342511 | 0 | 6 87E-18 |
| HSPA12A | 0 222478156 | -0 02683801 | 3 30E-18 | 0 234017044 | 0 | 5 85E-18 |
| NEGR1 | 0 439045516 | -0 02688015 | 2 93E-18 | -0 11346703 | 5 13E-298 | 5 19E-18 |
| STXBP3 | 0 321467324 | -0 02691527 | 2 65E-18 | -0 21403997 | 0 | 4 70E-18 |
| NIPAL3 | 0 134111287 | -0 02694886 | 2 40E-18 | 0 34211633 | 0 | 4 27E-18 |
| PARP9 | 0 159791361 | -0 02696893 | 2 27E-18 | 0 317500553 | 0 | 4 03E-18 |
| TNIK | 0 403285679 | -0 02697452 | 2 23E-18 | -0 11341318 | 9 84E-298 | 3 97E-18 |
| PTTG1 | 0 062163294 | -0 02701271 | 2 00E-18 | 0 376407194 | 0 | 3 56E-18 |
| HLA-DPA1 | 0 10029316 | -0 02703888 | 1 86E-18 | -0 41320665 | 0 | 3 30E-18 |
| GDNF | 0 038615294 | -0 0270521 | 1 79E-18 | 0 449079967 | 0 | 3 18E-18 |

| | | | | | | |
|---|---|---|---|---|---|---|
| HLA-DRB1 | 0 128457482 | -0 0270559 | 1 77E-18 | -0 38401228 | 0 | 3 15E-18 |
| KIF21A | 0 205231197 | -0 02706755 | 1 71E-18 | 0 252695918 | 0 | 3 04E-18 |
| KLF2 | 0 116217091 | -0 02707756 | 1 66E-18 | -0 35765187 | 0 | 2 96E-18 |
| CYBA | 0 13177933 | -0 0271914 | 1 20E-18 | -0 36458583 | 0 | 2 14E-18 |
| ANKFN1 | 0 070691592 | -0 02722961 | 1 07E-18 | 0 379535982 | 0 | 1 91E-18 |
| PRKCA-AS1 | 0 214539986 | -0 02724925 | 1 01E-18 | -0 31729796 | 0 | 1 81E-18 |
| A2M | 0 416144753 | -0 02725597 | 9 91E-19 | -0 12166881 | 0 | 1 78E-18 |
| WDR1 | 0 194304316 | -0 02726963 | 9 53E-19 | -0 30011287 | 0 | 1 71E-18 |
| CDCA3 | 0 0461537 | -0 02727188 | 9 47E-19 | 0 373970164 | 0 | 1 70E-18 |
| YWHAG | 0 290304772 | -0 02727721 | 9 32E-19 | 0 221627711 | 0 | 1 67E-18 |
| FCGR3A | 0 021158935 | -0 02728066 | 9 23E-19 | 0 460288362 | 0 | 1 65E-18 |
| OPTN | 0 178599303 | -0 02733708 | 7 83E-19 | -0 33167708 | 0 | 1 41E-18 |
| APOBEC3G | 0 043145952 | -0 02733927 | 7 78E-19 | -0 45613538 | 0 | 1 40E-18 |
| CXCL10 | 0 013106547 | -0 02735241 | 7 49E-19 | 0 469303216 | 0 | 1 34E-18 |
| ECT2 | 0 130370638 | -0 02742401 | 6 08E-19 | 0 262808655 | 0 | 1 09E-18 |
| GADD45A | 0 129057128 | -0 02744886 | 5 65E-19 | 0 343618991 | 0 | 1 02E-18 |
| UBE2L6 | 0 066427443 | -0 02745139 | 5 61E-19 | -0 43696922 | 0 | 1 01E-18 |
| GBP5 | 0 011117245 | -0 02745396 | 5 57E-19 | 0 469883843 | 0 | 1 00E-18 |
| DHCR7 | 0 021625326 | -0 02751832 | 4 61E-19 | 0 468546531 | 0 | 8 31E-19 |
| LIMD1 | 0 348479945 | -0 0275244 | 4 53E-19 | -0 18241157 | 0 | 8 17E-19 |
| TNFRSF9 | 0 006795987 | -0 02752821 | 4 48E-19 | 0 482884686 | 0 | 8 08E-19 |
| ARPC4 | 0 176467229 | -0 02754076 | 4 32E-19 | -0 3416106 | 0 | 7 80E-19 |
| RPS12 | 0 484799452 | -0 02755836 | 4 10E-19 | -0 14975123 | 0 | 7 41E-19 |
| C1QB | 0 068740363 | -0 02762746 | 3 35E-19 | 0 394641916 | 0 | 6 06E-19 |
| RAB6A | 0 535017418 | -0 02766649 | 2 99E-19 | 0 067565936 | 1 50E-106 | 5 41E-19 |
| NADSYN1 | 0 158772915 | -0 02769552 | 2 74E-19 | 0 287112143 | 0 | 4 97E-19 |
| TMED10 | 0 385258228 | -0 02785495 | 1 71E-19 | -0 14613091 | 0 | 3 11E-19 |
| FOLR3 | 0 003321848 | -0 0278556 | 1 71E-19 | -0 49373193 | 0 | 3 10E-19 |
| PALMD | 0 105832746 | -0 02788982 | 1 54E-19 | -0 38291408 | 0 | 2 81E-19 |
| BZW1 | 0 263111306 | -0 02793201 | 1 36E-19 | -0 25563407 | 0 | 2 48E-19 |
| BHLHE40 | 0 14634216 | -0 02795054 | 1 29E-19 | 0 32530101 | 0 | 2 35E-19 |
| FKBP2 | 0 050855685 | -0 02797533 | 1 20E-19 | -0 44478211 | 0 | 2 18E-19 |
| RPS29 | 0 439540462 | -0 02801985 | 1 05E-19 | -0 18697162 | 0 | 1 91E-19 |
| CTSB | 0 31271059 | -0 028035 | 1 00E-19 | 0 196971774 | 0 | 1 83E-19 |
| SNRPD2 | 0 157811578 | -0 0280671 | 9 10E-20 | 0 31965173 | 0 | 1 66E-19 |
| DOCK10 | 0 398621766 | -0 02809083 | 8 48E-20 | 0 144789069 | 0 | 1 55E-19 |
| DENND2B | 0 289038853 | -0 02815604 | 6 98E-20 | -0 16110842 | 0 | 1 28E-19 |
| CHST9 | 0 040728332 | -0 02821375 | 5 87E-20 | -0 45485908 | 0 | 1 08E-19 |
| HSPA1A | 0 192695742 | -0 028218 | 5 79E-20 | -0 30145609 | 0 | 1 06E-19 |
| LAMP2 | 0 354933277 | -0 02822538 | 5 67E-20 | -0 18062505 | 0 | 1 04E-19 |
| SELENOK | 0 120300394 | -0 02829515 | 4 60E-20 | -0 3738181 | 0 | 8 44E-20 |
| CD79B | 0 012963774 | -0 02847502 | 2 67E-20 | -0 47921087 | 0 | 4 92E-20 |
| PLAT | 0 13077992 | -0 02848078 | 2 62E-20 | -0 35684835 | 0 | 4 83E-20 |
| F5 | 0 014953075 | -0 02850937 | 2 41E-20 | -0 47523534 | 0 | 4 43E-20 |

| | | | | | | |
|---|---|---|---|---|---|---|
| SERPINE1 | 0 214282995 | -0 02858025 | 1 94E-20 | -0 3061039 | 0 | 3 58E-20 |
| DNAH5 | 0 03590261 | -0 02859571 | 1 85E-20 | -0 46319448 | 0 | 3 42E-20 |
| PDIA6 | 0 329300794 | -0 02860476 | 1 80E-20 | 0 198997094 | 0 | 3 33E-20 |
| AIP | 0 075793341 | -0 02863537 | 1 64E-20 | -0 41938738 | 0 | 3 03E-20 |
| CYP51A1 | 0 203156232 | -0 02864141 | 1 61E-20 | -0 30886752 | 0 | 2 98E-20 |
| IL1A | 0 009632408 | -0 02866212 | 1 51E-20 | 0 465494594 | 0 | 2 80E-20 |
| LRRC8B | 0 104443091 | -0 02867831 | 1 44E-20 | 0 378656391 | 0 | 2 66E-20 |
| GPNMB | 0 189802212 | -0 02868763 | 1 40E-20 | -0 31422862 | 0 | 2 59E-20 |
| RNF38 | 0 456273439 | -0 02873425 | 1 21E-20 | -0 14559185 | 0 | 2 25E-20 |
| CASC9 | 0 064990196 | -0 02875445 | 1 14E-20 | -0 44722192 | 0 | 2 12E-20 |
| PLOD1 | 0 252003579 | -0 02878184 | 1 05E-20 | -0 29007574 | 0 | 1 95E-20 |
| DEPDC7 | 0 035578991 | -0 02878944 | 1 03E-20 | 0 454173919 | 0 | 1 90E-20 |
| GATM | 0 049894348 | -0 02880391 | 9 81E-21 | -0 44083235 | 0 | 1 82E-20 |
| NOP2 | 0 102339571 | -0 02882496 | 9 19E-21 | 0 324065882 | 0 | 1 71E-20 |
| LAMP1 | 0 267803773 | -0 02883875 | 8 81E-21 | -0 23518253 | 0 | 1 64E-20 |
| FST | 0 060716529 | -0 02884462 | 8 66E-21 | -0 42503978 | 0 | 1 61E-20 |
| TRAPPC1 | 0 083208011 | -0 0288771 | 7 83E-21 | -0 41018493 | 0 | 1 46E-20 |
| SRI | 0 187889056 | -0 02892281 | 6 81E-21 | -0 33057877 | 0 | 1 27E-20 |
| ZC3HAV1 | 0 287468352 | -0 02893401 | 6 58E-21 | 0 202833076 | 0 | 1 23E-20 |
| LINC01436 | 0 036606956 | -0 02896908 | 5 90E-21 | 0 429062288 | 0 | 1 10E-20 |
| C1R | 0 280272601 | -0 02898328 | 5 65E-21 | -0 23459624 | 0 | 1 05E-20 |
| SIGIRR | 0 077763606 | -0 02900374 | 5 31E-21 | -0 40806397 | 0 | 9 91E-21 |
| RELT | 0 030600978 | -0 02903069 | 4 88E-21 | 0 427093122 | 0 | 9 13E-21 |
| ACSBG1 | 0 03400849 | -0 02909913 | 3 95E-21 | -0 45829096 | 0 | 7 40E-21 |
| IGSF11 | 0 092754754 | -0 02910989 | 3 82E-21 | -0 38386244 | 0 | 7 16E-21 |
| WFDC1 | 0 123603206 | -0 02915272 | 3 35E-21 | -0 33553416 | 0 | 6 27E-21 |
| GPX1 | 0 101454379 | -0 02934892 | 1 82E-21 | -0 39153313 | 0 | 3 42E-21 |
| TTLL7 | 0 253098171 | -0 02935449 | 1 79E-21 | 0 216152898 | 0 | 3 36E-21 |
| AKAP6 | 0 170242333 | -0 02939759 | 1 56E-21 | -0 32332478 | 0 | 2 94E-21 |
| PLCG2 | 0 306171594 | -0 02945135 | 1 32E-21 | -0 20831783 | 0 | 2 49E-21 |
| LINC01829 | 0 063191258 | -0 02948033 | 1 21E-21 | 0 381610814 | 0 | 2 28E-21 |
| SCRG1 | 0 085673222 | -0 02948669 | 1 18E-21 | -0 42006304 | 0 | 2 23E-21 |
| PDIA3 | 0 397669947 | -0 02950102 | 1 13E-21 | -0 15807104 | 0 | 2 13E-21 |
| CCDC107 | 0 098732177 | -0 02951451 | 1 08E-21 | -0 40028611 | 0 | 2 05E-21 |
| CUTA | 0 159600998 | -0 02958216 | 8 76E-22 | -0 34708591 | 0 | 1 66E-21 |
| ATP6V0E1 | 0 344805924 | -0 02960288 | 8 21E-22 | -0 21037981 | 0 | 1 55E-21 |
| SEMA3A | 0 410081666 | -0 02962843 | 7 58E-22 | 0 117170601 | 0 | 1 43E-21 |
| CADM4 | 0 044088253 | -0 02963522 | 7 42E-22 | -0 44675515 | 0 | 1 40E-21 |
| SYCE2 | 0 0702252 | -0 02973143 | 5 47E-22 | 0 3650736 | 0 | 1 04E-21 |
| TOMM7 | 0 289134035 | -0 02974226 | 5 29E-22 | -0 25008553 | 0 | 1 00E-21 |
| RPS26 | 0 316432202 | -0 02976599 | 4 91E-22 | -0 24433418 | 0 | 9 33E-22 |
| FHL2 | 0 312263235 | -0 02978559 | 4 61E-22 | 0 166833275 | 0 | 8 77E-22 |
| RPL27A | 0 436761151 | -0 02984717 | 3 80E-22 | -0 18432472 | 0 | 7 23E-22 |
| SAP18 | 0 159781843 | -0 02986696 | 3 56E-22 | -0 34376295 | 0 | 6 79E-22 |

| | | | | | | |
|---|---|---|---|---|---|---|
| LINC01117 | 0 175781919 | -0 02988805 | 3 33E-22 | 0 245842111 | 0 | 6 36E-22 |
| AFAP1L1 | 0 095476956 | -0 02991394 | 3 07E-22 | -0 38128768 | 0 | 5 86E-22 |
| C4orf3 | 0 170137633 | -0 02991442 | 3 07E-22 | -0 33299722 | 0 | 5 85E-22 |
| DLGAP1-AS1 | 0 061706421 | -0 02997859 | 2 50E-22 | -0 43220268 | 0 | 4 78E-22 |
| BCAS1 | 0 016466467 | -0 03007243 | 1 85E-22 | -0 47808477 | 0 | 3 55E-22 |
| CHCHD2 | 0 291189964 | -0 03007892 | 1 81E-22 | -0 26000251 | 0 | 3 48E-22 |
| PTPRQ | 0 065751651 | -0 03009491 | 1 72E-22 | 0 368761445 | 0 | 3 30E-22 |
| BAG3 | 0 136243361 | -0 03010621 | 1 66E-22 | -0 36945232 | 0 | 3 19E-22 |
| ATP1B2 | 0 041851478 | -0 03013166 | 1 53E-22 | -0 43749506 | 0 | 2 94E-22 |
| FABP5 | 0 084730921 | -0 03015701 | 1 41E-22 | 0 376086023 | 0 | 2 71E-22 |
| LYZ | 0 038796139 | -0 03015804 | 1 41E-22 | -0 46366513 | 0 | 2 71E-22 |
| MSMO1 | 0 104747673 | -0 03019326 | 1 26E-22 | -0 39538698 | 0 | 2 42E-22 |
| ASPA | 0 122537169 | -0 0302444 | 1 07E-22 | -0 35660146 | 0 | 2 06E-22 |
| SH3KBP1 | 0 449125278 | -0 0302738 | 9 72E-23 | -0 12935541 | 0 | 1 87E-22 |
| PLS3 | 0 347737526 | -0 03031948 | 8 39E-23 | -0 20842749 | 0 | 1 62E-22 |
| SP100 | 0 46651501 | -0 03038428 | 6 81E-23 | -0 15836478 | 0 | 1 31E-22 |
| RPL6 | 0 480011803 | -0 03043223 | 5 83E-23 | -0 16415427 | 0 | 1 13E-22 |
| NDUFA3 | 0 12563058 | -0 03046505 | 5 24E-23 | -0 37458848 | 0 | 1 01E-22 |
| ACTR3 | 0 487531172 | -0 03048282 | 4 95E-23 | 0 095939562 | 2 83E-213 | 9 58E-23 |
| BTNL8 | 0 004958977 | -0 03052905 | 4 26E-23 | 0 483607104 | 0 | 8 26E-23 |
| RPE65 | 0 005073195 | -0 0306126 | 3 25E-23 | -0 49335837 | 0 | 6 30E-23 |
| GIPR | 0 030781824 | -0 0306529 | 2 85E-23 | -0 45949259 | 0 | 5 53E-23 |
| NDE1 | 0 184129371 | -0 03070422 | 2 41E-23 | 0 265115343 | 0 | 4 69E-23 |
| NOP10 | 0 102330053 | -0 03074627 | 2 10E-23 | -0 40458035 | 0 | 4 09E-23 |
| PRELID1 | 0 123193924 | -0 03074717 | 2 10E-23 | -0 3864485 | 0 | 4 08E-23 |
| TMEM244 | 0 01235461 | -0 03074785 | 2 09E-23 | 0 467045152 | 0 | 4 07E-23 |
| TPMT | 0 053463669 | -0 03075426 | 2 05E-23 | 0 418129825 | 0 | 3 99E-23 |
| ATP8A1 | 0 107983857 | -0 03093645 | 1 13E-23 | -0 38382353 | 0 | 2 20E-23 |
| ITIH5 | 0 136557461 | -0 03098612 | 9 58E-24 | -0 34790417 | 0 | 1 87E-23 |
| MAP3K9 | 0 053016314 | -0 03101009 | 8 85E-24 | 0 398474651 | 0 | 1 73E-23 |
| FOXS1 | 0 072614266 | -0 03106769 | 7 32E-24 | -0 42224697 | 0 | 1 44E-23 |
| GJB2 | 0 043869334 | -0 03113706 | 5 82E-24 | -0 44986831 | 0 | 1 14E-23 |
| SKP1 | 0 37962346 | -0 03116845 | 5 25E-24 | -0 1792489 | 0 | 1 03E-23 |
| UQCR11 | 0 193028878 | -0 03121494 | 4 50E-24 | -0 32126785 | 0 | 8 85E-24 |
| CAPN12 | 0 035322 | -0 03129907 | 3 40E-24 | -0 4621346 | 0 | 6 71E-24 |
| CHST1 | 0 02002627 | -0 03131226 | 3 26E-24 | 0 468903791 | 0 | 6 42E-24 |
| MLPH | 0 047590946 | -0 03136339 | 2 75E-24 | -0 43103842 | 0 | 5 42E-24 |
| PAQR5 | 0 039043612 | -0 03141187 | 2 34E-24 | 0 420373318 | 0 | 4 62E-24 |
| CD81 | 0 369867316 | -0 03142268 | 2 25E-24 | -0 16483073 | 0 | 4 45E-24 |
| CARD16 | 0 017799014 | -0 03143506 | 2 16E-24 | -0 48118218 | 0 | 4 28E-24 |
| ACSL5 | 0 033504026 | -0 03147001 | 1 93E-24 | 0 432722193 | 0 | 3 81E-24 |
| CCL26 | 0 007814433 | -0 0314969 | 1 76E-24 | 0 47738256 | 0 | 3 48E-24 |
| SGSM1 | 0 064561878 | -0 03156768 | 1 39E-24 | -0 41752472 | 0 | 2 75E-24 |
| MAP4 | 0 731596581 | -0 03163394 | 1 11E-24 | -0 03802749 | 6 22E-35 | 2 20E-24 |

| | | | | | | |
|---|---|---|---|---|---|---|
| KIRREL3 | 0 115941063 | -0 03167973 | 9 53E-25 | 0 367892895 | 0 | 1 89E-24 |
| CALM1 | 0 327815956 | -0 03168615 | 9 33E-25 | -0 21921305 | 0 | 1 85E-24 |
| CADM3 | 0 018750833 | -0 03170921 | 8 63E-25 | -0 47172332 | 0 | 1 72E-24 |
| PRRG4 | 0 029896632 | -0 03171302 | 8 52E-25 | 0 440784945 | 0 | 1 69E-24 |
| BCL2L1 | 0 304134701 | -0 031729 | 8 08E-25 | -0 22785676 | 0 | 1 61E-24 |
| KIAA1549L | 0 125573471 | -0 03175563 | 7 38E-25 | -0 37317762 | 0 | 1 47E-24 |
| TGFBR2 | 0 4884354 | -0 03179226 | 6 52E-25 | -0 10399234 | 2 07E-250 | 1 30E-24 |
| IGFBP6 | 0 226704232 | -0 03181114 | 6 12E-25 | -0 27852538 | 0 | 1 22E-24 |
| ZSWIM6 | 0 567645771 | -0 03181156 | 6 11E-25 | 0 072806653 | 1 89E-123 | 1 22E-24 |
| FHAD1 | 0 110401477 | -0 03182534 | 5 84E-25 | 0 304560488 | 0 | 1 16E-24 |
| TMEM109 | 0 08468333 | -0 03186826 | 5 05E-25 | -0 4170526 | 0 | 1 01E-24 |
| IFITM2 | 0 150806191 | -0 031929 | 4 11E-25 | -0 34235838 | 0 | 8 21E-25 |
| ALDOC | 0 012725819 | -0 03201985 | 3 02E-25 | -0 48085051 | 0 | 6 04E-25 |
| CD70 | 0 104281282 | -0 03204934 | 2 73E-25 | 0 292785163 | 0 | 5 47E-25 |
| SIPA1 | 0 095277074 | -0 0320583 | 2 65E-25 | -0 40496207 | 0 | 5 31E-25 |
| CEBPB | 0 158887133 | -0 03206204 | 2 62E-25 | -0 33453558 | 0 | 5 24E-25 |
| TSPAN15 | 0 029296987 | -0 03211054 | 2 22E-25 | -0 46104518 | 0 | 4 45E-25 |
| ORMDL2 | 0 041584969 | -0 03211149 | 2 21E-25 | 0 439591812 | 0 | 4 44E-25 |
| PLAAT4 | 0 037206602 | -0 03212274 | 2 13E-25 | -0 45902199 | 0 | 4 27E-25 |
| COL20A1 | 0 022472445 | -0 03213808 | 2 02E-25 | -0 47479452 | 0 | 4 06E-25 |
| PLA2G2A | 0 010574708 | -0 03219026 | 1 69E-25 | -0 48932365 | 0 | 3 40E-25 |
| TAP1 | 0 076783233 | -0 03224604 | 1 40E-25 | 0 374543771 | 0 | 2 81E-25 |
| NDUFA2 | 0 125230816 | -0 03230491 | 1 14E-25 | 0 363047087 | 0 | 2 30E-25 |
| SMIM5 | 0 026251166 | -0 03232667 | 1 06E-25 | -0 45949519 | 0 | 2 14E-25 |
| EMP3 | 0 23990596 | -0 0323454 | 9 93E-26 | 0 240405764 | 0 | 2 01E-25 |
| ABCA6 | 0 550427367 | -0 03239259 | 8 45E-26 | -0 08541153 | 2 64E-169 | 1 71E-25 |
| ICOSLG | 0 044488017 | -0 03248449 | 6 16E-26 | -0 4524689 | 0 | 1 25E-25 |
| BDNF | 0 061002075 | -0 03253607 | 5 15E-26 | 0 417709553 | 0 | 1 04E-25 |
| COMMD6 | 0 173821172 | -0 03254407 | 5 01E-26 | -0 34366147 | 0 | 1 02E-25 |
| TIMP2 | 0 573661267 | -0 03255209 | 4 87E-26 | -0 06699482 | 8 71E-105 | 9 89E-26 |
| SERPINA3 | 0 083017647 | -0 03264694 | 3 51E-26 | -0 38599186 | 0 | 7 13E-26 |
| SOX8 | 0 100635815 | -0 03266737 | 3 27E-26 | -0 36769235 | 0 | 6 65E-26 |
| IKZF3 | 0 025765738 | -0 0327014 | 2 91E-26 | 0 435577742 | 0 | 5 91E-26 |
| PSMA7 | 0 254402163 | -0 03270713 | 2 85E-26 | 0 242894517 | 0 | 5 80E-26 |
| IFI30 | 0 030296396 | -0 03278778 | 2 15E-26 | 0 445320463 | 0 | 4 39E-26 |
| MX2 | 0 131417639 | -0 03279903 | 2 07E-26 | -0 36334455 | 0 | 4 22E-26 |
| CEROX1 | 0 197112181 | -0 03281056 | 1 99E-26 | -0 25480581 | 0 | 4 06E-26 |
| ZSWIM4 | 0 1276294 | -0 03282437 | 1 90E-26 | 0 308597837 | 0 | 3 87E-26 |
| RPL36A | 0 398869239 | -0 03283354 | 1 84E-26 | -0 20298494 | 0 | 3 75E-26 |
| KCP | 0 06034532 | -0 0328469 | 1 75E-26 | -0 43119806 | 0 | 3 58E-26 |
| GPX4 | 0 199263292 | -0 03285001 | 1 73E-26 | -0 31375869 | 0 | 3 55E-26 |
| TMEM47 | 0 122489578 | -0 03293445 | 1 29E-26 | 0 366680825 | 0 | 2 65E-26 |
| SLC31A2 | 0 026060802 | -0 03305613 | 8 44E-27 | -0 47430998 | 0 | 1 73E-26 |
| RPL15 | 0 431963983 | -0 0330591 | 8 35E-27 | -0 17940313 | 0 | 1 72E-26 |

| | | | | | | |
|---|---|---|---|---|---|---|
| B3GNT7 | 0 007595515 | -0 03307888 | 7 79E-27 | -0 48948048 | 0 | 1 60E-26 |
| NID1 | 0 469646494 | -0 03309831 | 7 28E-27 | -0 11483725 | 2 98E-305 | 1 50E-26 |
| COBLL1 | 0 295225676 | -0 03313615 | 6 37E-27 | 0 176012951 | 0 | 1 31E-26 |
| PKHD1 | 0 040509413 | -0 03317083 | 5 64E-27 | -0 43886387 | 0 | 1 16E-26 |
| ITGB8 | 0 444585102 | -0 03321703 | 4 79E-27 | -0 14457367 | 0 | 9 88E-27 |
| PCA3 | 0 080067008 | -0 03332864 | 3 23E-27 | -0 41414502 | 0 | 6 68E-27 |
| SREBF2 | 0 424387505 | -0 03340088 | 2 50E-27 | -0 15779074 | 0 | 5 18E-27 |
| CCDC73 | 0 030896042 | -0 03345078 | 2 10E-27 | 0 43067849 | 0 | 4 34E-27 |
| HSPA8 | 0 273524205 | -0 0334701 | 1 96E-27 | -0 25637748 | 0 | 4 05E-27 |
| CCL21 | 0 005891759 | -0 03348373 | 1 87E-27 | 0 482117366 | 0 | 3 87E-27 |
| NDUFB11 | 0 134377796 | -0 03349471 | 1 79E-27 | 0 356524126 | 0 | 3 72E-27 |
| KLHL4 | 0 085073576 | -0 03355275 | 1 46E-27 | -0 41879889 | 0 | 3 03E-27 |
| SYNM | 0 08920447 | -0 0335547 | 1 45E-27 | -0 39609508 | 0 | 3 01E-27 |
| SRM | 0 20568807 | -0 03356349 | 1 41E-27 | -0 3119615 | 0 | 2 92E-27 |
| RPS11 | 0 424301841 | -0 03359623 | 1 25E-27 | 0 107626317 | 3 62E-268 | 2 60E-27 |
| BICD1 | 0 599446041 | -0 03364414 | 1 05E-27 | 0 053937158 | 1 56E-68 | 2 19E-27 |
| ST3GAL6-AS1 | 0 023957282 | -0 03365911 | 9 99E-28 | -0 46977096 | 0 | 2 08E-27 |
| ITPK1 | 0 141316556 | -0 03368924 | 8 97E-28 | -0 33446454 | 0 | 1 87E-27 |
| AGA | 0 030220251 | -0 03388455 | 4 46E-28 | -0 46858329 | 0 | 9 33E-28 |
| C12orf75 | 0 15049209 | -0 03392919 | 3 80E-28 | 0 239431068 | 0 | 7 96E-28 |
| ST8SIA6 | 0 114180198 | -0 03403871 | 2 56E-28 | -0 35945804 | 0 | 5 37E-28 |
| BTG1 | 0 287258952 | -0 03404639 | 2 49E-28 | 0 211046955 | 0 | 5 23E-28 |
| APLP2 | 0 300860444 | -0 03404763 | 2 48E-28 | -0 21901624 | 0 | 5 21E-28 |
| ADGRL4 | 0 144685995 | -0 03411293 | 1 96E-28 | 0 279726818 | 0 | 4 12E-28 |
| CRHBP | 0 011240982 | -0 03411795 | 1 92E-28 | 0 472948397 | 0 | 4 05E-28 |
| PFDN5 | 0 357979098 | -0 03415219 | 1 70E-28 | -0 22830023 | 0 | 3 58E-28 |
| CSTB | 0 19874931 | -0 03415714 | 1 67E-28 | -0 30891586 | 0 | 3 51E-28 |
| MYBL1 | 0 134396832 | -0 03416317 | 1 63E-28 | 0 248920659 | 0 | 3 44E-28 |
| MXI1 | 0 232462736 | -0 03427819 | 1 07E-28 | -0 25697438 | 0 | 2 27E-28 |
| RPS13 | 0 337886201 | -0 03430068 | 9 90E-29 | -0 23408467 | 0 | 2 09E-28 |
| SLC15A3 | 0 035598028 | -0 03431019 | 9 57E-29 | 0 437500678 | 0 | 2 02E-28 |
| TPPP3 | 0 033037635 | -0 0344155 | 6 52E-29 | -0 46254219 | 0 | 1 38E-28 |
| UBXN1 | 0 140555101 | -0 03442957 | 6 19E-29 | -0 37404689 | 0 | 1 31E-28 |
| TMED5 | 0 195817708 | -0 03443668 | 6 03E-29 | -0 30962712 | 0 | 1 28E-28 |
| SHISA9 | 0 026793703 | -0 03445629 | 5 62E-29 | -0 46866531 | 0 | 1 19E-28 |
| FAM167B | 0 01025109 | -0 03446156 | 5 51E-29 | -0 48640881 | 0 | 1 17E-28 |
| TFRC | 0 179103767 | -0 03450515 | 4 70E-29 | 0 287838123 | 0 | 9 97E-29 |
| DAZAP2 | 0 203146713 | -0 03452264 | 4 41E-29 | 0 275615184 | 0 | 9 36E-29 |
| NDUFS5 | 0 19139175 | -0 03454458 | 4 07E-29 | -0 32569008 | 0 | 8 64E-29 |
| ORAI1 | 0 039157831 | -0 03459306 | 3 40E-29 | -0 45622379 | 0 | 7 24E-29 |
| UQCRB | 0 2435895 | -0 03460553 | 3 25E-29 | -0 29516367 | 0 | 6 92E-29 |
| RPS5 | 0 306761722 | -0 0346083 | 3 22E-29 | 0 176946469 | 0 | 6 86E-29 |
| FGD4 | 0 378928633 | -0 03463043 | 2 97E-29 | -0 1467441 | 0 | 6 33E-29 |
| HSPE1 | 0 23496602 | -0 03467007 | 2 57E-29 | 0 261524932 | 0 | 5 47E-29 |

| | | | | | |
|---|---|---|---|---|---|
| SCCPDH | 0 242637681 | -0 03475845 | 1 85E-29 | -0 26359 | 0 | 3 96E-29 |
| DNAJA1 | 0 22781786 | -0 03479476 | 1 62E-29 | -0 28898018 | 0 | 3 47E-29 |
| NDUFA13 | 0 256420019 | -0 03481014 | 1 53E-29 | -0 27613014 | 0 | 3 28E-29 |
| SORT1 | 0 157411814 | -0 03487132 | 1 22E-29 | -0 32212946 | 0 | 2 62E-29 |
| CCDC112 | 0 109145076 | -0 03498685 | 7 97E-30 | 0 360002397 | 0 | 1 71E-29 |
| SQLE | 0 158040014 | -0 03505234 | 6 25E-30 | 0 319508022 | 0 | 1 34E-29 |
| RAP1GDS1 | 0 408158992 | -0 03518023 | 3 88E-30 | 0 156962484 | 0 | 8 38E-30 |
| RPS17 | 0 345862443 | -0 03521956 | 3 35E-30 | 0 153355308 | 0 | 7 24E-30 |
| DPP7 | 0 176314938 | -0 03525008 | 2 99E-30 | -0 32056632 | 0 | 6 47E-30 |
| CPEB2 | 0 226171213 | -0 03531475 | 2 35E-30 | -0 27492819 | 0 | 5 09E-30 |
| DSTN | 0 369791171 | -0 03531504 | 2 35E-30 | -0 17924303 | 0 | 5 08E-30 |
| MYO7A | 0 026470084 | -0 03534473 | 2 10E-30 | -0 47410641 | 0 | 4 55E-30 |
| PERP | 0 100426415 | -0 03534537 | 2 09E-30 | -0 3959777 | 0 | 4 54E-30 |
| TMED9 | 0 201595249 | -0 03536105 | 1 97E-30 | -0 31525635 | 0 | 4 28E-30 |
| RPS27L | 0 360929737 | -0 03537071 | 1 90E-30 | -0 19701696 | 0 | 4 13E-30 |
| TRAF1 | 0 075336468 | -0 03551131 | 1 12E-30 | 0 410172171 | 0 | 2 45E-30 |
| PCDH20 | 0 023843064 | -0 03552004 | 1 09E-30 | -0 4673285 | 0 | 2 37E-30 |
| HMGCR | 0 160467153 | -0 03567789 | 5 99E-31 | -0 33985581 | 0 | 1 31E-30 |
| LINC01705 | 0 018836497 | -0 03569004 | 5 72E-31 | -0 4789349 | 0 | 1 25E-30 |
| SLC17A3 | 0 005986941 | -0 03570552 | 5 40E-31 | -0 48816554 | 0 | 1 18E-30 |
| EIF5A | 0 11417068 | -0 03572612 | 4 99E-31 | -0 3887561 | 0 | 1 09E-30 |
| SLC38A5 | 0 013334983 | -0 03573519 | 4 82E-31 | 0 469822283 | 0 | 1 05E-30 |
| TCERG1L | 0 055310198 | -0 03574096 | 4 72E-31 | -0 42585982 | 0 | 1 03E-30 |
| TMCC3 | 0 127581809 | -0 03575898 | 4 41E-31 | 0 298133902 | 0 | 9 64E-31 |
| HPCAL1 | 0 134653823 | -0 03576046 | 4 38E-31 | 0 306643188 | 0 | 9 59E-31 |
| RPL34 | 0 471207477 | -0 03581275 | 3 60E-31 | -0 16938569 | 0 | 7 87E-31 |
| MOXD1 | 0 092488245 | -0 03583158 | 3 35E-31 | 0 377207604 | 0 | 7 33E-31 |
| SV2B | 0 039567113 | -0 03586583 | 2 94E-31 | -0 44673347 | 0 | 6 44E-31 |
| KAZN-AS1 | 0 155032267 | -0 03591699 | 2 42E-31 | 0 256430495 | 0 | 5 31E-31 |
| CYB5R3 | 0 228036778 | -0 0359272 | 2 33E-31 | -0 28161028 | 0 | 5 11E-31 |
| CHI3L2 | 0 007852506 | -0 03609672 | 1 22E-31 | -0 48897384 | 0 | 2 68E-31 |
| ADH1B | 0 079438808 | -0 03610212 | 1 19E-31 | -0 40006244 | 0 | 2 63E-31 |
| HSP90AB1 | 0 423749786 | -0 03612484 | 1 09E-31 | -0 16182577 | 0 | 2 41E-31 |
| PCBP1 | 0 186204336 | -0 03613635 | 1 05E-31 | -0 32043898 | 0 | 2 31E-31 |
| DGKG | 0 101273534 | -0 03617365 | 9 08E-32 | -0 40472064 | 0 | 2 00E-31 |
| GAL3ST1 | 0 012078582 | -0 03626733 | 6 34E-32 | -0 48102201 | 0 | 1 40E-31 |
| C4orf45 | 0 067560107 | -0 03630941 | 5 39E-32 | -0 43009889 | 0 | 1 19E-31 |
| RBM47 | 0 098322895 | -0 03632643 | 5 05E-32 | 0 352509502 | 0 | 1 12E-31 |
| NFKBIA | 0 192276941 | -0 03634487 | 4 70E-32 | -0 31446096 | 0 | 1 04E-31 |
| RHOB | 0 179408349 | -0 03645567 | 3 07E-32 | -0 30139154 | 0 | 6 81E-32 |
| CFD | 0 065209115 | -0 03646106 | 3 00E-32 | -0 43107381 | 0 | 6 67E-32 |
| TPM3 | 0 361224801 | -0 03648528 | 2 74E-32 | 0 164683465 | 0 | 6 08E-32 |
| PFKFB4 | 0 101149797 | -0 03653677 | 2 24E-32 | -0 3981064 | 0 | 4 99E-32 |
| ACP6 | 0 136490834 | -0 03658367 | 1 87E-32 | 0 321003307 | 0 | 4 16E-32 |

| | | | | | | |
|---|---|---|---|---|---|---|
| EIF4A1 | 0 413755687 | -0 03658966 | 1 83E-32 | -0 1639497 | 0 | 4 07E-32 |
| TUBB2B | 0 06344825 | -0 03659089 | 1 82E-32 | -0 41479417 | 0 | 4 05E-32 |
| PIM3 | 0 074660677 | -0 03670669 | 1 16E-32 | -0 41792812 | 0 | 2 59E-32 |
| CD44 | 0 425310769 | -0 03673827 | 1 03E-32 | 0 082386424 | 1 26E-157 | 2 29E-32 |
| SLC2A3 | 0 321505397 | -0 0367389 | 1 02E-32 | -0 22236136 | 0 | 2 29E-32 |
| SELENOS | 0 249861986 | -0 03678629 | 8 51E-33 | -0 27070333 | 0 | 1 90E-32 |
| MZT2B | 0 210904038 | -0 03694465 | 4 58E-33 | 0 264231324 | 0 | 1 03E-32 |
| PTPN22 | 0 031333879 | -0 03698785 | 3 87E-33 | 0 426177699 | 0 | 8 68E-33 |
| MYLK | 0 113142716 | -0 03699282 | 3 80E-33 | -0 37659599 | 0 | 8 52E-33 |
| ELOB | 0 223867811 | -0 03703875 | 3 17E-33 | -0 30059451 | 0 | 7 12E-33 |
| NPC2 | 0 189630885 | -0 03707516 | 2 75E-33 | -0 32314251 | 0 | 6 18E-33 |
| CLSTN2 | 0 09939845 | -0 03708322 | 2 66E-33 | -0 38333797 | 0 | 5 99E-33 |
| IFITM10 | 0 023548 | -0 03715744 | 1 99E-33 | -0 46761857 | 0 | 4 47E-33 |
| MT1G | 0 005101749 | -0 03722735 | 1 51E-33 | -0 49276624 | 0 | 3 40E-33 |
| P2RY12 | 0 042565342 | -0 03732899 | 1 01E-33 | 0 436483716 | 0 | 2 28E-33 |
| GCNT1 | 0 211741638 | -0 03740806 | 7 39E-34 | 0 26197218 | 0 | 1 67E-33 |
| NDUFA4 | 0 237469304 | -0 03745017 | 6 25E-34 | -0 2936958 | 0 | 1 42E-33 |
| DUSP5 | 0 089128324 | -0 03749569 | 5 22E-34 | -0 39829492 | 0 | 1 18E-33 |
| SCML1 | 0 266775809 | -0 03756561 | 3 95E-34 | 0 198311459 | 0 | 8 97E-34 |
| GABARAP | 0 196284099 | -0 03763521 | 3 00E-34 | -0 313913 | 0 | 6 81E-34 |
| MYH11 | 0 06828349 | -0 03773908 | 1 98E-34 | 0 365059095 | 0 | 4 51E-34 |
| C6orf141 | 0 015286212 | -0 0377783 | 1 69E-34 | 0 453789693 | 0 | 3 86E-34 |
| CTSL | 0 225105176 | -0 03778589 | 1 64E-34 | -0 30564904 | 0 | 3 74E-34 |
| ATP6V0C | 0 205183606 | -0 03778687 | 1 64E-34 | -0 30720779 | 0 | 3 73E-34 |
| PPFIBP1 | 0 673183454 | -0 0378957 | 1 06E-34 | -0 08320811 | 9 33E-161 | 2 42E-34 |
| M6PR | 0 209285945 | -0 0379059 | 1 01E-34 | 0 244639605 | 0 | 2 32E-34 |
| RPL8 | 0 443204965 | -0 03794706 | 8 60E-35 | -0 18421397 | 0 | 1 97E-34 |
| APOC1 | 0 045715863 | -0 03807934 | 5 05E-35 | 0 432327598 | 0 | 1 16E-34 |
| TMA7 | 0 296348823 | -0 0383094 | 1 99E-35 | -0 25541799 | 0 | 4 60E-35 |
| PDCD1LG2 | 0 072500048 | -0 03835317 | 1 67E-35 | 0 35555268 | 0 | 3 85E-35 |
| MYEF2 | 0 237012431 | -0 03839227 | 1 42E-35 | 0 238345245 | 0 | 3 29E-35 |
| PTPN12 | 0 64855038 | -0 03841936 | 1 27E-35 | 0 02728655 | 9 07E-19 | 2 95E-35 |
| RPL32 | 0 477518037 | -0 03848092 | 9 92E-36 | -0 17057365 | 0 | 2 30E-35 |
| SLC44A1 | 0 475462108 | -0 03866032 | 4 77E-36 | -0 14983818 | 0 | 1 11E-35 |
| RTN3 | 0 313576745 | -0 03875472 | 3 24E-36 | 0 168648068 | 0 | 7 53E-36 |
| APOE | 0 239867888 | -0 03884161 | 2 27E-36 | -0 29826 | 0 | 5 28E-36 |
| RPS3A | 0 463031353 | -0 03888288 | 1 91E-36 | -0 17855337 | 0 | 4 46E-36 |
| COX5B | 0 187089528 | -0 03889355 | 1 83E-36 | -0 33383274 | 0 | 4 27E-36 |
| VAT1L | 0 064000305 | -0 03899577 | 1 20E-36 | -0 42880352 | 0 | 2 81E-36 |
| TPM4 | 0 48722659 | -0 03914975 | 6 35E-37 | -0 12238451 | 0 | 1 49E-36 |
| ACSL3 | 0 446774286 | -0 03924649 | 4 25E-37 | 0 112833098 | 1 07E-294 | 1 00E-36 |
| TNFRSF1A | 0 265129162 | -0 03928695 | 3 59E-37 | 0 210798767 | 0 | 8 47E-37 |
| COX7B | 0 17341189 | -0 03940658 | 2 18E-37 | -0 34227674 | 0 | 5 15E-37 |
| UNC93B1 | 0 044725971 | -0 03945121 | 1 81E-37 | 0 429401883 | 0 | 4 28E-37 |

| | | | | | | |
|---|---|---|---|---|---|---|
| TUBB6 | 0 318716567 | -0 03947712 | 1 63E-37 | 0 192300368 | 0 | 3 85E-37 |
| RP1 | 0 012630637 | -0 03961837 | 9 00E-38 | 0 461877366 | 0 | 2 13E-37 |
| TM4SF1 | 0 119224839 | -0 03969965 | 6 40E-38 | -0 37907843 | 0 | 1 52E-37 |
| ELL2 | 0 368277779 | -0 03984188 | 3 52E-38 | -0 21357812 | 0 | 8 37E-38 |
| LIF | 0 027764558 | -0 0398919 | 2 85E-38 | -0 47211687 | 0 | 6 79E-38 |
| DEPP1 | 0 041984733 | -0 03991353 | 2 60E-38 | -0 45706562 | 0 | 6 20E-38 |
| USP54 | 0 413289296 | -0 0399321 | 2 40E-38 | -0 14355269 | 0 | 5 74E-38 |
| RPL22L1 | 0 127039272 | -0 03994256 | 2 30E-38 | -0 38466808 | 0 | 5 49E-38 |
| ZNF267 | 0 123450915 | -0 04001148 | 1 72E-38 | 0 320606893 | 0 | 4 11E-38 |
| GAS2L3 | 0 20621157 | -0 04002192 | 1 64E-38 | 0 259003909 | 0 | 3 93E-38 |
| TBCB | 0 154451657 | -0 04008983 | 1 23E-38 | -0 35003137 | 0 | 2 95E-38 |
| CCNDBP1 | 0 076897451 | -0 04016588 | 8 92E-39 | -0 42093957 | 0 | 2 14E-38 |
| PRDX4 | 0 197283509 | -0 04017785 | 8 47E-39 | 0 26210481 | 0 | 2 03E-38 |
| RGS7 | 0 127619882 | -0 04033682 | 4 30E-39 | 0 371891502 | 0 | 1 04E-38 |
| SYNGR2 | 0 032780644 | -0 04033769 | 4 29E-39 | 0 438373687 | 0 | 1 03E-38 |
| LUM | 0 446222231 | -0 04037185 | 3 71E-39 | -0 16682338 | 0 | 8 94E-39 |
| ANXA5 | 0 444271002 | -0 04049442 | 2 19E-39 | -0 12693942 | 0 | 5 31E-39 |
| GNG5 | 0 20085283 | -0 04049921 | 2 15E-39 | -0 31888667 | 0 | 5 20E-39 |
| ATP6V1F | 0 120547867 | -0 04051467 | 2 01E-39 | -0 38643239 | 0 | 4 87E-39 |
| TPT1 | 0 565018751 | -0 04052186 | 1 95E-39 | -0 11559858 | 0 | 4 72E-39 |
| KRTCAP2 | 0 21846148 | -0 04052771 | 1 90E-39 | -0 29699491 | 0 | 4 61E-39 |
| ATOX1 | 0 172897908 | -0 04059806 | 1 41E-39 | -0 34536914 | 0 | 3 41E-39 |
| CALB2 | 0 009442044 | -0 04060481 | 1 37E-39 | -0 48735752 | 0 | 3 32E-39 |
| COX4I1 | 0 248538958 | -0 04065396 | 1 11E-39 | -0 28565563 | 0 | 2 69E-39 |
| RACK1 | 0 433353639 | -0 04065574 | 1 10E-39 | -0 1769795 | 0 | 2 67E-39 |
| SCN9A | 0 130970284 | -0 04073664 | 7 74E-40 | -0 35838812 | 0 | 1 88E-39 |
| CTSC | 0 148959662 | -0 04074155 | 7 58E-40 | -0 36102563 | 0 | 1 85E-39 |
| PSMB9 | 0 073613676 | -0 04077287 | 6 62E-40 | 0 381768344 | 0 | 1 61E-39 |
| JUNB | 0 18860292 | -0 04078953 | 6 16E-40 | -0 31553401 | 0 | 1 50E-39 |
| RPL9 | 0 460394814 | -0 04081505 | 5 52E-40 | -0 18040522 | 0 | 1 35E-39 |
| CLGN | 0 042717633 | -0 04085908 | 4 56E-40 | 0 403967886 | 0 | 1 12E-39 |
| SH3TC2 | 0 055652853 | -0 04090238 | 3 78E-40 | -0 43384282 | 0 | 9 26E-40 |
| IFI27 | 0 181283433 | -0 04091858 | 3 53E-40 | -0 33520792 | 0 | 8 64E-40 |
| EFL1 | 0 203936723 | -0 04110656 | 1 56E-40 | 0 252664863 | 0 | 3 83E-40 |
| CNR1 | 0 025832366 | -0 04116729 | 1 20E-40 | -0 46884633 | 0 | 2 94E-40 |
| FADS1 | 0 119358093 | -0 04119063 | 1 08E-40 | 0 308969851 | 0 | 2 66E-40 |
| WARS1 | 0 09939845 | -0 04123397 | 8 96E-41 | -0 37517637 | 0 | 2 21E-40 |
| IFI6 | 0 156336259 | -0 04124377 | 8 58E-41 | -0 37684261 | 0 | 2 11E-40 |
| TMEM233 | 0 04578249 | -0 04132964 | 5 90E-41 | -0 44464298 | 0 | 1 46E-40 |
| CPPED1 | 0 089566161 | -0 04142737 | 3 84E-41 | -0 38571048 | 0 | 9 52E-41 |
| PADI2 | 0 019493252 | -0 04144455 | 3 57E-41 | -0 46975161 | 0 | 8 84E-41 |
| C4orf48 | 0 105804192 | -0 04148055 | 3 04E-41 | -0 39091566 | 0 | 7 55E-41 |
| INSIG1 | 0 111619805 | -0 04148405 | 3 00E-41 | 0 349535007 | 0 | 7 44E-41 |
| CCN3 | 0 053549333 | -0 04148483 | 2 99E-41 | -0 44173114 | 0 | 7 41E-41 |

| | | | | | | |
|---|---|---|---|---|---|---|
| MYC | 0 076135996 | -0 04163982 | 1 51E-41 | -0 42758092 | 0 | 3 76E-41 |
| SOX5 | 0 666111439 | -0 04172595 | 1 03E-41 | -0 06674981 | 4 92E-104 | 2 58E-41 |
| TRIM14 | 0 106898784 | -0 04188897 | 5 03E-42 | 0 326839488 | 0 | 1 26E-41 |
| MT1M | 0 038681921 | -0 0420428 | 2 54E-42 | -0 45301361 | 0 | 6 37E-42 |
| PRDX5 | 0 140621728 | -0 04205831 | 2 37E-42 | -0 37533943 | 0 | 5 95E-42 |
| TNFAIP6 | 0 098227713 | -0 04217871 | 1 39E-42 | 0 354781195 | 0 | 3 49E-42 |
| RAN | 0 194742152 | -0 04229423 | 8 27E-43 | 0 277892606 | 0 | 2 08E-42 |
| B3GNT5 | 0 103386572 | -0 04231584 | 7 50E-43 | 0 327490899 | 0 | 1 89E-42 |
| SNAI1 | 0 029211323 | -0 04234884 | 6 47E-43 | -0 45793592 | 0 | 1 64E-42 |
| TRIM58 | 0 012611601 | -0 04240058 | 5 13E-43 | 0 465872375 | 0 | 1 30E-42 |
| EPCAM-DT | 0 037711066 | -0 04242611 | 4 58E-43 | 0 430466351 | 0 | 1 16E-42 |
| TXK | 0 038386857 | -0 0424436 | 4 23E-43 | 0 421273239 | 0 | 1 07E-42 |
| DDX39A | 0 139879309 | -0 04248688 | 3 48E-43 | 0 266103318 | 0 | 8 84E-43 |
| ITGA7 | 0 11033485 | -0 04250301 | 3 24E-43 | 0 367066245 | 0 | 8 22E-43 |
| CCDC85B | 0 122499096 | -0 04251586 | 3 06E-43 | -0 39117217 | 0 | 7 76E-43 |
| GZMB | 0 006063087 | -0 04272832 | 1 17E-43 | -0 49228567 | 0 | 2 99E-43 |
| RPL14 | 0 399782985 | -0 04275637 | 1 03E-43 | -0 20675195 | 0 | 2 64E-43 |
| CTSK | 0 199605947 | -0 04280784 | 8 19E-44 | -0 2953226 | 0 | 2 09E-43 |
| TDRD7 | 0 128571701 | -0 04285773 | 6 53E-44 | 0 310536982 | 0 | 1 67E-43 |
| IGF1 | 0 194856371 | -0 0428704 | 6 17E-44 | -0 28443913 | 0 | 1 58E-43 |
| RPS10 | 0 364660867 | -0 042919 | 4 95E-44 | -0 22278127 | 0 | 1 27E-43 |
| RPLP0 | 0 497734671 | -0 0429559 | 4 18E-44 | 0 039884821 | 2 93E-38 | 1 07E-43 |
| ITGA5 | 0 190363785 | -0 04298656 | 3 64E-44 | 0 268030215 | 0 | 9 36E-44 |
| MCF2L2 | 0 191306086 | -0 04301522 | 3 19E-44 | 0 273246227 | 0 | 8 22E-44 |
| TMEM59 | 0 32248577 | -0 04309586 | 2 21E-44 | -0 21996726 | 0 | 5 70E-44 |
| CCR7 | 0 012830519 | -0 04313344 | 1 86E-44 | -0 48230152 | 0 | 4 81E-44 |
| METRN | 0 11816832 | -0 04329501 | 8 90E-45 | -0 36129717 | 0 | 2 31E-44 |
| RUBCNL | 0 037282747 | -0 04336149 | 6 56E-45 | -0 45381575 | 0 | 1 70E-44 |
| KCND3 | 0 045563572 | -0 04339592 | 5 61E-45 | -0 43855273 | 0 | 1 45E-44 |
| CDH6 | 0 042927034 | -0 04341606 | 5 11E-45 | -0 45612848 | 0 | 1 33E-44 |
| RNF157 | 0 259741867 | -0 04379296 | 8 98E-46 | 0 22796258 | 0 | 2 35E-45 |
| DAD1 | 0 258761493 | -0 04390178 | 5 42E-46 | -0 27076351 | 0 | 1 42E-45 |
| RPS4X | 0 445517885 | -0 04391003 | 5 22E-46 | -0 1791537 | 0 | 1 37E-45 |
| CDC42 | 0 405998363 | -0 04391517 | 5 10E-46 | -0 18638078 | 0 | 1 34E-45 |
| FADS3 | 0 114932135 | -0 04411451 | 2 01E-46 | -0 38955257 | 0 | 5 31E-46 |
| NFE2L3 | 0 082560774 | -0 0441932 | 1 39E-46 | 0 36492102 | 0 | 3 68E-46 |
| SQSTM1 | 0 436475605 | -0 04424235 | 1 11E-46 | -0 13930041 | 0 | 2 93E-46 |
| DNER | 0 072005102 | -0 04427344 | 9 58E-47 | -0 41794009 | 0 | 2 53E-46 |
| IL12RB2 | 0 054681997 | -0 04437919 | 5 84E-47 | 0 410513903 | 0 | 1 55E-46 |
| UGCG | 0 335344844 | -0 04441081 | 5 03E-47 | 0 149528114 | 0 | 1 33E-46 |
| UBE2S | 0 123784051 | -0 04445672 | 4 06E-47 | 0 297271316 | 0 | 1 08E-46 |
| HSBP1 | 0 183206107 | -0 04454581 | 2 67E-47 | -0 32568498 | 0 | 7 08E-47 |
| LINC01435 | 0 019912052 | -0 04460541 | 2 01E-47 | -0 47751459 | 0 | 5 35E-47 |
| RPL24 | 0 405075099 | -0 04462597 | 1 83E-47 | -0 19841052 | 0 | 4 86E-47 |

| | | | | | | |
|---|---|---|---|---|---|---|
| COMTD1 | 0 026555748 | -0 04471634 | 1 19E-47 | -0 47194868 | 0 | 3 18E-47 |
| FIGN | 0 296015686 | -0 04472865 | 1 12E-47 | 0 177867867 | 0 | 3 00E-47 |
| SLC20A1 | 0 304286992 | -0 04479341 | 8 28E-48 | 0 178293592 | 0 | 2 21E-47 |
| SERINC5 | 0 51529573 | -0 04481098 | 7 62E-48 | -0 09246233 | 3 53E-198 | 2 04E-47 |
| RPL27 | 0 297548114 | -0 04483155 | 6 91E-48 | -0 26048667 | 0 | 1 85E-47 |
| ERAP1 | 0 172250671 | -0 04490056 | 4 98E-48 | 0 28376849 | 0 | 1 33E-47 |
| TUBB | 0 298614152 | -0 04493242 | 4 28E-48 | 0 18838036 | 0 | 1 15E-47 |
| SPECC1 | 0 225428794 | -0 04495562 | 3 83E-48 | -0 26820405 | 0 | 1 03E-47 |
| SP140L | 0 160714626 | -0 04502561 | 2 75E-48 | 0 285790883 | 0 | 7 37E-48 |
| RASEF | 0 072423902 | -0 04510033 | 1 92E-48 | -0 42566128 | 0 | 5 18E-48 |
| FAM102A | 0 054624888 | -0 04511784 | 1 77E-48 | -0 44081022 | 0 | 4 77E-48 |
| HTR7 | 0 032352325 | -0 04529422 | 7 62E-49 | 0 424172074 | 0 | 2 05E-48 |
| NDUFA1 | 0 1738878 | -0 04534305 | 6 03E-49 | -0 33770542 | 0 | 1 63E-48 |
| TRIM9 | 0 146351678 | -0 04540196 | 4 54E-49 | -0 34876815 | 0 | 1 23E-48 |
| H3-3B | 0 356989206 | -0 04549613 | 2 89E-49 | -0 17122066 | 0 | 7 85E-49 |
| UGT8 | 0 039728922 | -0 04555244 | 2 20E-49 | 0 430211454 | 0 | 5 99E-49 |
| LUCAT1 | 0 040490377 | -0 04562796 | 1 53E-49 | 0 431379735 | 0 | 4 17E-49 |
| OSBPL3 | 0 352829758 | -0 04567948 | 1 19E-49 | 0 139720651 | 0 | 3 26E-49 |
| RPL7A | 0 445213303 | -0 04574567 | 8 67E-50 | -0 18651049 | 0 | 2 37E-49 |
| MAPK4 | 0 04857132 | -0 04574933 | 8 51E-50 | -0 44402263 | 0 | 2 33E-49 |
| SLC25A3 | 0 318849822 | -0 04580224 | 6 59E-50 | 0 190680231 | 0 | 1 80E-49 |
| RPLP2 | 0 427690316 | -0 04596683 | 2 97E-50 | -0 19498419 | 0 | 8 16E-50 |
| AMPD3 | 0 0585559 | -0 0459862 | 2 70E-50 | -0 43903555 | 0 | 7 42E-50 |
| SSR4 | 0 22040319 | -0 04605616 | 1 92E-50 | -0 29852437 | 0 | 5 29E-50 |
| ATP5ME | 0 227170623 | -0 04619117 | 9 94E-51 | -0 29982174 | 0 | 2 75E-50 |
| LRAT | 0 157107232 | -0 04622315 | 8 50E-51 | 0 296623806 | 0 | 2 35E-50 |
| RPL18A | 0 383697245 | -0 04623724 | 7 94E-51 | -0 22234213 | 0 | 2 20E-50 |
| HBEGF | 0 030467724 | -0 04623855 | 7 88E-51 | -0 46138637 | 0 | 2 19E-50 |
| SUN2 | 0 127001199 | -0 04626655 | 6 88E-51 | 0 311024098 | 0 | 1 91E-50 |
| BTF3 | 0 257543165 | -0 04626874 | 6 80E-51 | -0 2842071 | 0 | 1 89E-50 |
| MGST3 | 0 310959243 | -0 04627936 | 6 46E-51 | -0 22490686 | 0 | 1 79E-50 |
| RPL10 | 0 630951248 | -0 04638379 | 3 87E-51 | -0 10123579 | 2 35E-237 | 1 08E-50 |
| TPD52 | 0 180103177 | -0 04641816 | 3 27E-51 | -0 33617809 | 0 | 9 12E-51 |
| HTATIP2 | 0 054101388 | -0 04660008 | 1 34E-51 | 0 418091074 | 0 | 3 74E-51 |
| RPS25 | 0 350497801 | -0 04660159 | 1 33E-51 | -0 22473001 | 0 | 3 71E-51 |
| CC2D1A | 0 227408578 | -0 04666984 | 9 48E-52 | -0 29999824 | 0 | 2 66E-51 |
| RPL22 | 0 415973425 | -0 04681767 | 4 56E-52 | 0 102164319 | 1 03E-241 | 1 28E-51 |
| PPIA | 0 391825779 | -0 04684487 | 3 99E-52 | 0 160296702 | 0 | 1 12E-51 |
| TMBIM1 | 0 118520493 | -0 04685098 | 3 87E-52 | -0 37227956 | 0 | 1 09E-51 |
| SAMD5 | 0 268298719 | -0 04700971 | 1 76E-52 | 0 211593932 | 0 | 4 98E-52 |
| HEXB | 0 258285584 | -0 0470137 | 1 73E-52 | -0 26942777 | 0 | 4 88E-52 |
| KIF5C | 0 108840494 | -0 04713424 | 9 47E-53 | -0 37860308 | 0 | 2 69E-52 |
| RTN4 | 0 681597533 | -0 0473854 | 2 70E-53 | 0 016714656 | 6 02E-08 | 7 70E-53 |
| PPDPF | 0 220555482 | -0 0474056 | 2 44E-53 | -0 29619777 | 0 | 6 96E-53 |

| | | | | | | |
|---|---|---|---|---|---|---|
| TUBB4B | 0 144885877 | -0 0474082 | 2 41E-53 | 0 321195269 | 0 | 6 87E-53 |
| RRAGA | 0 083036683 | -0 04762708 | 8 02E-54 | -0 40787242 | 0 | 2 30E-53 |
| DUSP4 | 0 069568445 | -0 04766727 | 6 55E-54 | -0 40883333 | 0 | 1 88E-53 |
| MACORIS | 0 01408692 | -0 04768357 | 6 03E-54 | -0 47924715 | 0 | 1 73E-53 |
| EEF1A1 | 0 74732063 | -0 04771782 | 5 07E-54 | -0 05649478 | 5 11E-75 | 1 46E-53 |
| TMEM62 | 0 124459843 | -0 04778595 | 3 60E-54 | 0 349751304 | 0 | 1 04E-53 |
| MALL | 0 031638461 | -0 04782926 | 2 89E-54 | -0 45601226 | 0 | 8 34E-54 |
| PPA1 | 0 191068131 | -0 04790684 | 1 95E-54 | -0 32981744 | 0 | 5 64E-54 |
| FNIP2 | 0 390160096 | -0 0479874 | 1 30E-54 | -0 16223494 | 0 | 3 76E-54 |
| UQCR10 | 0 142373075 | -0 04804389 | 9 74E-55 | -0 37856356 | 0 | 2 82E-54 |
| OPN3 | 0 067607698 | -0 04805001 | 9 44E-55 | -0 41943976 | 0 | 2 74E-54 |
| RPL36AL | 0 197045554 | -0 04807706 | 8 23E-55 | -0 33243645 | 0 | 2 39E-54 |
| SAMHD1 | 0 300089471 | -0 04816815 | 5 18E-55 | -0 18929203 | 0 | 1 50E-54 |
| COX6B1 | 0 203118159 | -0 04819584 | 4 50E-55 | -0 32690225 | 0 | 1 31E-54 |
| CANX | 0 442643392 | -0 04835195 | 2 03E-55 | -0 14892725 | 0 | 5 91E-55 |
| TPP1 | 0 132769222 | -0 04835802 | 1 96E-55 | -0 37042364 | 0 | 5 73E-55 |
| TMEM158 | 0 050113266 | -0 04839542 | 1 62E-55 | -0 45647502 | 0 | 4 74E-55 |
| SRP14 | 0 286526051 | -0 04843974 | 1 29E-55 | -0 25711351 | 0 | 3 78E-55 |
| ARPC5 | 0 268603301 | -0 04854017 | 7 73E-56 | -0 26737916 | 0 | 2 26E-55 |
| CNP | 0 100407379 | -0 04870892 | 3 25E-56 | -0 38415313 | 0 | 9 54E-56 |
| CST1 | 0 023386191 | -0 04872039 | 3 06E-56 | -0 4786619 | 0 | 8 99E-56 |
| ARAP2 | 0 202537549 | -0 04890103 | 1 20E-56 | -0 30777053 | 0 | 3 55E-56 |
| CAVIN1 | 0 30135539 | -0 04894191 | 9 75E-57 | -0 21117939 | 0 | 2 88E-56 |
| RHOA | 0 435676077 | -0 04894829 | 9 43E-57 | -0 15807745 | 0 | 2 79E-56 |
| PSME1 | 0 125183225 | -0 04911827 | 3 91E-57 | 0 342544225 | 0 | 1 16E-56 |
| UQCRQ | 0 179217986 | -0 04912834 | 3 71E-57 | 0 302520926 | 0 | 1 10E-56 |
| RPL35A | 0 416011498 | -0 0492513 | 1 96E-57 | -0 20702788 | 0 | 5 81E-57 |
| ATP6AP2 | 0 25005235 | -0 04934253 | 1 22E-57 | -0 27111567 | 0 | 3 62E-57 |
| NACA | 0 43250652 | -0 04937984 | 1 00E-57 | -0 1995991 | 0 | 2 98E-57 |
| TRPM8 | 0 015019703 | -0 04941274 | 8 43E-58 | 0 453223454 | 0 | 2 51E-57 |
| IL15 | 0 129742438 | -0 04943559 | 7 48E-58 | 0 280425902 | 0 | 2 23E-57 |
| LHFPL2 | 0 505330186 | -0 04950824 | 5 12E-58 | -0 14137404 | 0 | 1 53E-57 |
| SOD1 | 0 214911195 | -0 04952239 | 4 75E-58 | -0 3062465 | 0 | 1 42E-57 |
| FXYD5 | 0 077259142 | -0 04954034 | 4 32E-58 | 0 387556077 | 0 | 1 30E-57 |
| RPL39 | 0 459452514 | -0 04954093 | 4 31E-58 | -0 18305381 | 0 | 1 29E-57 |
| EID1 | 0 291951419 | -0 04956241 | 3 85E-58 | -0 24117824 | 0 | 1 16E-57 |
| USP53 | 0 363642421 | -0 04958955 | 3 34E-58 | -0 18131484 | 0 | 1 00E-57 |
| MRPL41 | 0 093287773 | -0 04959082 | 3 32E-58 | -0 40881556 | 0 | 9 98E-58 |
| VAMP5 | 0 070358455 | -0 04970227 | 1 85E-58 | -0 43392639 | 0 | 5 58E-58 |
| CD4 | 0 044906817 | -0 04972158 | 1 67E-58 | -0 44529722 | 0 | 5 04E-58 |
| FAM180A | 0 049237593 | -0 04980451 | 1 08E-58 | 0 388389019 | 0 | 3 26E-58 |
| LDLR | 0 167015667 | -0 04986925 | 7 68E-59 | 0 285771309 | 0 | 2 33E-58 |
| TNFRSF14 | 0 124773943 | -0 05004405 | 3 05E-59 | -0 37198262 | 0 | 9 30E-59 |
| TMEFF2 | 0 137395062 | -0 05012039 | 2 04E-59 | 0 222998179 | 0 | 6 22E-59 |

| | | | | | | |
|---|---|---|---|---|---|---|
| AQP7 | 0 017684796 | -0 05018921 | 1 41E-59 | -0 47261929 | 0 | 4 33E-59 |
| UBA52 | 0 333802897 | -0 05031074 | 7 41E-60 | -0 25370703 | 0 | 2 28E-59 |
| COX7C | 0 241514534 | -0 05047892 | 3 03E-60 | -0 29980196 | 0 | 9 32E-60 |
| HLA-DRA | 0 184824199 | -0 05054223 | 2 16E-60 | -0 35387146 | 0 | 6 66E-60 |
| RPL12 | 0 405379681 | -0 05056282 | 1 93E-60 | -0 20947701 | 0 | 5 97E-60 |
| RPL13A | 0 524737774 | -0 05065387 | 1 19E-60 | 0 035234724 | 3 17E-30 | 3 68E-60 |
| TMEM50A | 0 231282481 | -0 05072436 | 8 14E-61 | 0 244081451 | 0 | 2 53E-60 |
| MYL12A | 0 237916659 | -0 05074337 | 7 35E-61 | -0 29963404 | 0 | 2 28E-60 |
| TFPI | 0 522453408 | -0 05084029 | 4 37E-61 | -0 15071066 | 0 | 1 36E-60 |
| ITGAV | 0 496021397 | -0 05097463 | 2 12E-61 | -0 12999077 | 0 | 6 61E-61 |
| EHBP1 | 0 752203461 | -0 05104944 | 1 42E-61 | -0 04238319 | 5 55E-43 | 4 42E-61 |
| CLIC1 | 0 170613542 | -0 05132514 | 3 19E-62 | 0 298413308 | 0 | 1 00E-61 |
| TRIM54 | 0 013287392 | -0 05138232 | 2 34E-62 | -0 48320193 | 0 | 7 35E-62 |
| RAB8B | 0 321867088 | -0 05141709 | 1 93E-62 | -0 2114848 | 0 | 6 09E-62 |
| RGL3 | 0 030315433 | -0 0514206 | 1 90E-62 | 0 440253128 | 0 | 5 98E-62 |
| SLC7A14 | 0 036844911 | -0 05146288 | 1 51E-62 | -0 45201966 | 0 | 4 76E-62 |
| COL8A1 | 0 387504521 | -0 0515255 | 1 07E-62 | -0 1687916 | 0 | 3 39E-62 |
| RPS8 | 0 470569759 | -0 05159086 | 7 51E-63 | -0 17494444 | 0 | 2 37E-62 |
| APOL1 | 0 045582608 | -0 05163418 | 5 93E-63 | -0 46043241 | 0 | 1 88E-62 |
| STMN1 | 0 203213341 | -0 05168668 | 4 45E-63 | 0 235114262 | 0 | 1 41E-62 |
| ECM1 | 0 115293827 | -0 05169491 | 4 25E-63 | -0 37541521 | 0 | 1 35E-62 |
| RPL29 | 0 359016581 | -0 0518071 | 2 30E-63 | -0 23079255 | 0 | 7 33E-63 |
| YWHAH | 0 109773277 | -0 0518539 | 1 78E-63 | 0 352363994 | 0 | 5 68E-63 |
| POLR2L | 0 179884259 | -0 05187599 | 1 58E-63 | -0 3415066 | 0 | 5 04E-63 |
| CHN1 | 0 463926063 | -0 05195881 | 1 00E-63 | -0 15820152 | 0 | 3 20E-63 |
| FXYD1 | 0 113314043 | -0 05198118 | 8 86E-64 | -0 37995373 | 0 | 2 83E-63 |
| BMP8B | 0 053968133 | -0 05203955 | 6 43E-64 | -0 43710723 | 0 | 2 06E-63 |
| PDGFB | 0 035198264 | -0 05208466 | 5 01E-64 | -0 45123101 | 0 | 1 61E-63 |
| SYT11 | 0 09808494 | -0 05210817 | 4 41E-64 | -0 39102835 | 0 | 1 41E-63 |
| IFITM3 | 0 288515353 | -0 05211995 | 4 13E-64 | -0 24708375 | 0 | 1 33E-63 |
| GBP1 | 0 107469875 | -0 05215653 | 3 37E-64 | -0 39653176 | 0 | 1 09E-63 |
| TIMP1 | 0 450514934 | -0 05230758 | 1 47E-64 | -0 16131022 | 0 | 4 72E-64 |
| MIR155HG | 0 076088405 | -0 05232054 | 1 36E-64 | 0 33042623 | 0 | 4 40E-64 |
| RPS9 | 0 451247835 | -0 05233431 | 1 26E-64 | -0 1916356 | 0 | 4 08E-64 |
| SLFN5 | 0 136052997 | -0 05235091 | 1 15E-64 | -0 37449983 | 0 | 3 72E-64 |
| CSPG4 | 0 147950734 | -0 05237363 | 1 02E-64 | -0 34408723 | 0 | 3 28E-64 |
| EEF1B2 | 0 265376635 | -0 05245581 | 6 45E-65 | 0 198211788 | 0 | 2 09E-64 |
| ABHD2 | 0 338114637 | -0 05262783 | 2 48E-65 | -0 20322976 | 0 | 8 07E-65 |
| SNCA | 0 167720013 | -0 05272162 | 1 47E-65 | 0 301131514 | 0 | 4 80E-65 |
| FLNA | 0 40175325 | -0 05283201 | 7 95E-66 | -0 16789801 | 0 | 2 60E-65 |
| DNAJB1 | 0 200967048 | -0 0528512 | 7 14E-66 | 0 245696649 | 0 | 2 34E-65 |
| RPL3 | 0 475728617 | -0 05291175 | 5 09E-66 | -0 17562355 | 0 | 1 67E-65 |
| COL15A1 | 0 217490625 | -0 05297472 | 3 58E-66 | -0 27669581 | 0 | 1 17E-65 |
| CD14 | 0 069587482 | -0 05303159 | 2 60E-66 | -0 43081233 | 0 | 8 55E-66 |

| | | | | | | |
|---|---|---|---|---|---|---|
| CMTM5 | 0 014515239 | -0 05314769 | 1 36E-66 | -0 47876874 | 0 | 4 47E-66 |
| RPL37 | 0 556985399 | -0 05328141 | 6 39E-67 | -0 15067802 | 0 | 2 11E-66 |
| RPL18 | 0 41258495 | -0 05331032 | 5 43E-67 | -0 21137637 | 0 | 1 80E-66 |
| SOX2-OT | 0 126591917 | -0 0533207 | 5 12E-67 | -0 36170873 | 0 | 1 69E-66 |
| TGFBI | 0 43692296 | -0 05336547 | 3 98E-67 | -0 18473637 | 0 | 1 32E-66 |
| RPL30 | 0 531866898 | -0 05338755 | 3 51E-67 | 0 022484825 | 3 12E-13 | 1 16E-66 |
| ALK | 0 104576345 | -0 05349473 | 1 92E-67 | -0 37680978 | 0 | 6 37E-67 |
| RPS16 | 0 418952618 | -0 05359466 | 1 09E-67 | 0 074302606 | 1 66E-128 | 3 62E-67 |
| RHBDF2 | 0 085882622 | -0 05361932 | 9 47E-68 | 0 346057494 | 0 | 3 15E-67 |
| CADPS2 | 0 438474425 | -0 0536318 | 8 83E-68 | 0 074694892 | 7 53E-130 | 2 93E-67 |
| ARL6IP5 | 0 203070568 | -0 05364394 | 8 24E-68 | -0 31299306 | 0 | 2 74E-67 |
| CDKN2B | 0 048913975 | -0 05370843 | 5 71E-68 | -0 45358023 | 0 | 1 90E-67 |
| ST8SIA4 | 0 150711009 | -0 05374317 | 4 69E-68 | 0 271958253 | 0 | 1 56E-67 |
| ARL4C | 0 110791723 | -0 05391015 | 1 81E-68 | 0 314164991 | 0 | 6 08E-68 |
| ATP1A1 | 0 306095448 | -0 05395383 | 1 41E-68 | -0 22939675 | 0 | 4 74E-68 |
| TBC1D2 | 0 081085454 | -0 05404774 | 8 27E-69 | 0 33696025 | 0 | 2 78E-68 |
| CSRP1 | 0 140688355 | -0 0540742 | 7 11E-69 | -0 33585212 | 0 | 2 39E-68 |
| PFN1 | 0 227684605 | -0 05408214 | 6 80E-69 | 0 262743693 | 0 | 2 29E-68 |
| MAMDC2 | 0 315090137 | -0 054104 | 6 00E-69 | 0 175436727 | 0 | 2 02E-68 |
| GPSM2 | 0 370476481 | -0 05419313 | 3 60E-69 | 0 137346147 | 0 | 1 22E-68 |
| RPS19 | 0 548523729 | -0 05439116 | 1 16E-69 | 0 004842902 | 0 116477164 | 3 91E-69 |
| RPS27A | 0 510232053 | -0 05440923 | 1 04E-69 | -0 18164138 | 0 | 3 53E-69 |
| GPR37 | 0 029515905 | -0 0544796 | 6 95E-70 | -0 45272686 | 0 | 2 36E-69 |
| CLDN11 | 0 085482858 | -0 05458896 | 3 70E-70 | -0 41004625 | 0 | 1 26E-69 |
| MET | 0 265804953 | -0 05472443 | 1 69E-70 | 0 179576146 | 0 | 5 77E-70 |
| DDX5 | 0 848946336 | -0 05474874 | 1 47E-70 | -0 0697241 | 2 34E-113 | 5 01E-70 |
| COX6C | 0 311663589 | -0 05484619 | 8 35E-71 | -0 24939578 | 0 | 2 86E-70 |
| LPL | 0 07432754 | -0 05488581 | 6 64E-71 | -0 41371479 | 0 | 2 28E-70 |
| EHD1 | 0 084711884 | -0 05493389 | 5 02E-71 | -0 41287777 | 0 | 1 72E-70 |
| BCHE | 0 132655004 | -0 05495732 | 4 38E-71 | -0 37305737 | 0 | 1 50E-70 |
| RPS6 | 0 443024119 | -0 05502868 | 2 90E-71 | -0 1954244 | 0 | 9 94E-71 |
| POLR2F | 0 17703832 | -0 05523024 | 8 94E-72 | -0 3289623 | 0 | 3 08E-71 |
| PLAAT3 | 0 11327597 | -0 05538202 | 3 68E-72 | -0 36865284 | 0 | 1 28E-71 |
| TMEM169 | 0 022672327 | -0 05540721 | 3 18E-72 | 0 451083395 | 0 | 1 10E-71 |
| OST4 | 0 204993242 | -0 05552227 | 1 62E-72 | -0 3235783 | 0 | 5 64E-72 |
| ATP6V0B | 0 185204927 | -0 0555676 | 1 24E-72 | -0 33953328 | 0 | 4 32E-72 |
| MYL9 | 0 22502903 | -0 05559044 | 1 08E-72 | -0 27523923 | 0 | 3 78E-72 |
| RAMP3 | 0 029449278 | -0 05568687 | 6 15E-73 | -0 45589908 | 0 | 2 15E-72 |
| RPL11 | 0 518427214 | -0 05576163 | 3 96E-73 | -0 16466152 | 0 | 1 39E-72 |
| IGFBP7 | 0 645199977 | -0 05584305 | 2 45E-73 | -0 06878313 | 2 31E-110 | 8 61E-73 |
| ABCA8 | 0 598408559 | -0 05588458 | 1 92E-73 | -0 06471246 | 6 96E-98 | 6 75E-73 |
| GFRA3 | 0 024471265 | -0 0559141 | 1 61E-73 | -0 46525361 | 0 | 5 68E-73 |
| KCNH8 | 0 071462565 | -0 05592229 | 1 53E-73 | -0 42171846 | 0 | 5 41E-73 |
| UNC13B | 0 234689993 | -0 05610353 | 5 25E-74 | -0 29906078 | 0 | 1 86E-73 |

| | | | | | | |
|---|---|---|---|---|---|---|
| RABAC1 | 0 174363709 | -0 0563065 | 1 57E-74 | -0 33865034 | 0 | 5 60E-74 |
| PLAU | 0 072481011 | -0 05634148 | 1 28E-74 | 0 360830125 | 0 | 4 55E-74 |
| NPC1 | 0 313119872 | -0 0563432 | 1 26E-74 | 0 184564087 | 0 | 4 51E-74 |
| SCN7A | 0 339371038 | -0 05649221 | 5 19E-75 | -0 18540321 | 0 | 1 85E-74 |
| RPS15A | 0 387057166 | -0 05683001 | 6 86E-76 | -0 22913174 | 0 | 2 47E-75 |
| NRXN1 | 0 183339362 | -0 05687798 | 5 14E-76 | -0 30233061 | 0 | 1 85E-75 |
| IFI27L2 | 0 095905275 | -0 05688893 | 4 82E-76 | -0 41206751 | 0 | 1 74E-75 |
| RPS23 | 0 476042718 | -0 05700706 | 2 37E-76 | -0 18557794 | 0 | 8 54E-76 |
| APOL6 | 0 067464925 | -0 05702244 | 2 16E-76 | 0 395695608 | 0 | 7 80E-76 |
| ITGB1 | 0 585016466 | -0 05702767 | 2 09E-76 | -0 09245773 | 3 69E-198 | 7 56E-76 |
| MCUB | 0 168985932 | -0 05702869 | 2 08E-76 | 0 278817414 | 0 | 7 51E-76 |
| TCIRG1 | 0 096048048 | -0 05717236 | 8 72E-77 | 0 337633809 | 0 | 3 17E-76 |
| PLP2 | 0 095020083 | -0 05724893 | 5 49E-77 | 0 355423807 | 0 | 2 00E-76 |
| TMBIM6 | 0 442405437 | -0 05729508 | 4 15E-77 | 0 110558287 | 5 98E-283 | 1 51E-76 |
| MLXIP | 0 320943824 | -0 05736546 | 2 71E-77 | 0 150504273 | 0 | 9 89E-77 |
| SLC7A14-AS1 | 0 081694618 | -0 05755194 | 8 73E-78 | -0 39962278 | 0 | 3 20E-77 |
| TAGLN2 | 0 223477566 | -0 05774448 | 2 70E-78 | 0 252333215 | 0 | 9 93E-78 |
| TLR1 | 0 046420209 | -0 05780459 | 1 87E-78 | -0 45219514 | 0 | 6 88E-78 |
| RNASEK | 0 21404504 | -0 05781541 | 1 75E-78 | -0 30329346 | 0 | 6 45E-78 |
| PLA2G4C | 0 116892882 | -0 05784957 | 1 42E-78 | -0 38333542 | 0 | 5 24E-78 |
| LDHA | 0 238126059 | -0 05808482 | 3 36E-79 | -0 29723387 | 0 | 1 25E-78 |
| RPL13 | 0 611467514 | -0 05814267 | 2 36E-79 | -0 12927463 | 0 | 8 77E-79 |
| MYL12B | 0 26424397 | -0 05825683 | 1 17E-79 | -0 26441805 | 0 | 4 36E-79 |
| ZNF365 | 0 056233462 | -0 05829005 | 9 51E-80 | 0 392832673 | 0 | 3 56E-79 |
| PPIB | 0 321315033 | -0 05833329 | 7 29E-80 | -0 23618387 | 0 | 2 73E-79 |
| RPS24 | 0 568749881 | -0 0585724 | 1 66E-80 | -0 14235843 | 0 | 6 25E-80 |
| RCAN3 | 0 085235385 | -0 05857732 | 1 61E-80 | 0 366723558 | 0 | 6 07E-80 |
| PTMS | 0 330090804 | -0 05870585 | 7 27E-81 | 0 149361698 | 0 | 2 75E-80 |
| ZC3H12C | 0 250099941 | -0 05872505 | 6 46E-81 | 0 244706293 | 0 | 2 44E-80 |
| CERKL | 0 021910872 | -0 05874431 | 5 73E-81 | 0 449337563 | 0 | 2 17E-80 |
| TNFSF10 | 0 082398964 | -0 05875217 | 5 45E-81 | 0 360475374 | 0 | 2 07E-80 |
| PTMA | 0 514972112 | -0 0587597 | 5 21E-81 | 0 059285917 | 1 95E-82 | 1 97E-80 |
| VWA1 | 0 069768327 | -0 05879707 | 4 13E-81 | -0 41495784 | 0 | 1 56E-80 |
| DRAP1 | 0 158677733 | -0 05882689 | 3 43E-81 | 0 293222148 | 0 | 1 30E-80 |
| CPNE7 | 0 019407588 | -0 05899965 | 1 17E-81 | 0 436218743 | 0 | 4 45E-81 |
| VKORC1 | 0 143619958 | -0 0590779 | 7 17E-82 | -0 37242527 | 0 | 2 74E-81 |
| DBI | 0 161314272 | -0 05909048 | 6 63E-82 | -0 3557364 | 0 | 2 54E-81 |
| GBP4 | 0 056747444 | -0 059118 | 5 58E-82 | -0 43732826 | 0 | 2 14E-81 |
| CTSZ | 0 172421998 | -0 05921868 | 2 97E-82 | -0 34566594 | 0 | 1 14E-81 |
| GAS7 | 0 511431345 | -0 05934859 | 1 32E-82 | -0 07617073 | 5 75E-135 | 5 08E-82 |
| TRMT112 | 0 150349318 | -0 05939131 | 1 01E-82 | -0 37059271 | 0 | 3 88E-82 |
| ITM2B | 0 563857532 | -0 05968611 | 1 58E-83 | -0 10201417 | 5 26E-241 | 6 13E-83 |
| XIST | 0 375073766 | -0 05977821 | 8 81E-84 | -0 21824387 | 0 | 3 44E-83 |
| ARHGDIA | 0 23759304 | -0 06022382 | 5 22E-85 | -0 2930255 | 0 | 2 05E-84 |

| | | | | | | |
|---|---|---|---|---|---|---|
| MMP17 | 0 11979593 | -0 06044435 | 1 28E-85 | -0 36347668 | 0 | 5 04E-85 |
| GRN | 0 214606613 | -0 0604681 | 1 10E-85 | -0 31100684 | 0 | 4 33E-85 |
| SAMD9 | 0 113761398 | -0 06048119 | 1 01E-85 | 0 311778347 | 0 | 3 99E-85 |
| RXRG | 0 028868668 | -0 06058045 | 5 35E-86 | -0 45958196 | 0 | 2 13E-85 |
| DAG1 | 0 176657593 | -0 06061064 | 4 41E-86 | -0 32073045 | 0 | 1 75E-85 |
| PKN1 | 0 171555843 | -0 06067376 | 2 94E-86 | 0 269541085 | 0 | 1 17E-85 |
| OCIAD2 | 0 059888447 | -0 06077557 | 1 53E-86 | 0 365660984 | 0 | 6 11E-86 |
| FMNL1 | 0 047905047 | -0 06130694 | 4 96E-88 | 0 380525527 | 0 | 2 01E-87 |
| BIRC3 | 0 036330928 | -0 06149261 | 1 49E-88 | 0 419303336 | 0 | 6 05E-88 |
| SPTBN1 | 0 792598656 | -0 06162728 | 6 19E-89 | -0 0340518 | 2 44E-28 | 2 52E-88 |
| MCTP1 | 0 431002646 | -0 06165188 | 5 27E-89 | -0 19295576 | 0 | 2 15E-88 |
| RASAL2 | 0 626135044 | -0 06165236 | 5 26E-89 | -0 09207805 | 1 52E-196 | 2 15E-88 |
| SCARB2 | 0 399944795 | -0 06175075 | 2 77E-89 | -0 17262081 | 0 | 1 13E-88 |
| ENTPD2 | 0 081218709 | -0 06176125 | 2 58E-89 | -0 40054345 | 0 | 1 06E-88 |
| SEC61B | 0 268546192 | -0 0618968 | 1 06E-89 | -0 2877143 | 0 | 4 38E-89 |
| SEMA4B | 0 088252651 | -0 06207734 | 3 26E-90 | 0 368637696 | 0 | 1 35E-89 |
| RETREG1 | 0 193266833 | -0 06208805 | 3 04E-90 | -0 31193251 | 0 | 1 26E-89 |
| MID1IP1 | 0 051826541 | -0 06214207 | 2 13E-90 | 0 409860386 | 0 | 8 81E-90 |
| CRIP1 | 0 095781539 | -0 06221893 | 1 29E-90 | -0 39709603 | 0 | 5 33E-90 |
| FGFBP2 | 0 038386857 | -0 06225291 | 1 03E-90 | -0 4428096 | 0 | 4 27E-90 |
| PLEKHN1 | 0 018474805 | -0 0622654 | 9 48E-91 | 0 442987074 | 0 | 3 93E-90 |
| RPS7 | 0 446574404 | -0 06239153 | 4 13E-91 | -0 20389349 | 0 | 1 72E-90 |
| ADGRG6 | 0 128904837 | -0 06240113 | 3 87E-91 | -0 36663539 | 0 | 1 61E-90 |
| SH3BGRL3 | 0 150901373 | -0 06243741 | 3 05E-91 | 0 300855437 | 0 | 1 27E-90 |
| MIR3142HG | 0 021910872 | -0 06248644 | 2 21E-91 | -0 48262165 | 0 | 9 21E-91 |
| CD58 | 0 158382669 | -0 06250646 | 1 93E-91 | -0 35525116 | 0 | 8 07E-91 |
| EEF1G | 0 335173517 | -0 06260018 | 1 04E-91 | 0 136421175 | 0 | 4 36E-91 |
| SRRM4 | 0 028526013 | -0 06265052 | 7 45E-92 | -0 47358775 | 0 | 3 13E-91 |
| LMO4 | 0 32700691 | -0 06271123 | 4 98E-92 | -0 2123355 | 0 | 2 09E-91 |
| SDC4 | 0 091964745 | -0 06275733 | 3 67E-92 | -0 41527491 | 0 | 1 54E-91 |
| MT-ATP8 | 0 241295616 | -0 06281511 | 2 50E-92 | -0 27529717 | 0 | 1 05E-91 |
| RPL23A | 0 388018503 | -0 06290294 | 1 39E-92 | 0 108636369 | 3 25E-273 | 5 90E-92 |
| MEGF9 | 0 221688146 | -0 06296347 | 9 32E-93 | -0 30273798 | 0 | 3 95E-92 |
| WDR83OS | 0 1744113 | -0 06296578 | 9 18E-93 | 0 257374015 | 0 | 3 90E-92 |
| C10orf90 | 0 034027527 | -0 06305994 | 4 90E-93 | -0 46014362 | 0 | 2 09E-92 |
| UBL5 | 0 212388875 | -0 06345598 | 3 47E-94 | -0 32730002 | 0 | 1 49E-93 |
| TPST2 | 0 085911176 | -0 06349703 | 2 63E-94 | 0 33293497 | 0 | 1 13E-93 |
| HSPA5 | 0 364717976 | -0 06351399 | 2 35E-94 | -0 19218873 | 0 | 1 01E-93 |
| RPS14 | 0 452152063 | -0 06352809 | 2 14E-94 | -0 19662003 | 0 | 9 18E-94 |
| PTGDS | 0 163808037 | -0 0636129 | 1 21E-94 | -0 31726642 | 0 | 5 21E-94 |
| CD74 | 0 221926101 | -0 06365638 | 9 02E-95 | -0 33134297 | 0 | 3 89E-94 |
| RPL36 | 0 438141288 | -0 06367901 | 7 75E-95 | -0 20959382 | 0 | 3 34E-94 |
| RNASE1 | 0 096362148 | -0 06369453 | 6 98E-95 | -0 41790403 | 0 | 3 01E-94 |
| RPL37A | 0 601435343 | -0 0637237 | 5 73E-95 | -0 01541031 | 5 88E-07 | 2 48E-94 |

| | | | | | | |
|---|---|---|---|---|---|---|
| FA2H | 0 01918867 | -0 06378462 | 3 80E-95 | -0 473605 | 0 | 1 64E-94 |
| RDX | 0 517132741 | -0 06389408 | 1 82E-95 | 0 077894099 | 4 54E-141 | 7 87E-95 |
| S100Z | 0 020987607 | -0 06408056 | 5 14E-96 | 0 431902498 | 0 | 2 24E-95 |
| UBALD2 | 0 110934496 | -0 06413986 | 3 44E-96 | 0 31423805 | 0 | 1 50E-95 |
| COX7A1 | 0 053711142 | -0 06415573 | 3 09E-96 | -0 44497254 | 0 | 1 35E-95 |
| S100A4 | 0 179570159 | -0 06445729 | 3 98E-97 | -0 3400866 | 0 | 1 75E-96 |
| PEBP1 | 0 269707411 | -0 064485 | 3 29E-97 | -0 26996315 | 0 | 1 45E-96 |
| ECRG4 | 0 082627401 | -0 06449917 | 2 99E-97 | -0 39583838 | 0 | 1 32E-96 |
| ITPR3 | 0 24542651 | -0 0649029 | 1 89E-98 | -0 26746659 | 0 | 8 39E-98 |
| NRP2 | 0 489901201 | -0 06492703 | 1 60E-98 | -0 1514401 | 0 | 7 12E-98 |
| FOLH1 | 0 049856275 | -0 06495181 | 1 35E-98 | -0 44039628 | 0 | 6 01E-98 |
| ZEB2 | 0 551693286 | -0 06496947 | 1 19E-98 | 0 039919812 | 2 53E-38 | 5 33E-98 |
| SHC3 | 0 106470465 | -0 06497414 | 1 16E-98 | 0 279453069 | 0 | 5 16E-98 |
| ADI1 | 0 161314272 | -0 06502672 | 8 06E-99 | -0 34885721 | 0 | 3 60E-98 |
| TSPO | 0 123412842 | -0 06505469 | 6 65E-99 | -0 39190191 | 0 | 2 97E-98 |
| TECR | 0 193723706 | -0 06505546 | 6 61E-99 | 0 226364699 | 0 | 2 96E-98 |
| CLDN14 | 0 035236337 | -0 06509168 | 5 16E-99 | 0 428522618 | 0 | 2 31E-98 |
| CTXND1 | 0 036054901 | -0 06522743 | 2 02E-99 | -0 44812292 | 0 | 9 11E-99 |
| RPS21 | 0 433772439 | -0 06531825 | 1 08E-99 | 0 045036259 | 2 61E-48 | 4 88E-99 |
| STARD13 | 0 721231273 | -0 06549666 | 3 15E-100 | -0 05055054 | 2 06E-60 | 1 43E-99 |
| C19orf53 | 0 15196741 | -0 06553604 | 2 40E-100 | 0 295585411 | 0 | 1 09E-99 |
| SELENOM | 0 143144048 | -0 06555468 | 2 11E-100 | -0 35536901 | 0 | 9 58E-100 |
| ARHGEF4 | 0 117130837 | -0 06557237 | 1 86E-100 | 0 312968027 | 0 | 8 49E-100 |
| MRPL51 | 0 153614056 | -0 06585906 | 2 54E-101 | 0 268514645 | 0 | 1 16E-100 |
| EHD4 | 0 118986884 | -0 06588097 | 2 18E-101 | 0 336130169 | 0 | 1 00E-100 |
| RPS28 | 0 511478936 | -0 06594549 | 1 39E-101 | -0 1855438 | 0 | 6 40E-101 |
| IQGAP1 | 0 556804554 | -0 06598679 | 1 04E-101 | 0 063129055 | 3 09E-93 | 4 80E-101 |
| COX8A | 0 148921589 | -0 06618198 | 2 67E-102 | 0 301259505 | 0 | 1 23E-101 |
| EREG | 0 019103006 | -0 06624194 | 1 75E-102 | 0 436376007 | 0 | 8 11E-102 |
| RASGEF1C | 0 041166169 | -0 06643089 | 4 65E-103 | -0 45732594 | 0 | 2 16E-102 |
| TUBA1A | 0 274504578 | -0 06653763 | 2 20E-103 | -0 2519878 | 0 | 1 02E-102 |
| PMP22 | 0 354228931 | -0 06655722 | 1 91E-103 | -0 17670335 | 0 | 8 94E-103 |
| CLDN10-AS1 | 0 017227923 | -0 06661731 | 1 25E-103 | -0 47136748 | 0 | 5 87E-103 |
| TEC | 0 078772534 | -0 06688946 | 1 84E-104 | 0 348912641 | 0 | 8 67E-104 |
| CALR | 0 392758562 | -0 06707814 | 4 82E-105 | 0 119679526 | 0 | 2 28E-104 |
| RPS18 | 0 540280977 | -0 06712646 | 3 42E-105 | -0 16985828 | 0 | 1 62E-104 |
| LY75 | 0 088090842 | -0 06714079 | 3 09E-105 | -0 422245 | 0 | 1 47E-104 |
| RPS15 | 0 443937865 | -0 06734069 | 7 46E-106 | 0 068252924 | 1 08E-108 | 3 56E-105 |
| RPS27 | 0 549294702 | -0 06743556 | 3 80E-106 | -0 17694408 | 0 | 1 82E-105 |
| QKI | 0 790932973 | -0 06757585 | 1 39E-106 | -0 04014266 | 9 84E-39 | 6 70E-106 |
| RPL26 | 0 45090518 | -0 06763375 | 9 21E-107 | -0 19210207 | 0 | 4 44E-106 |
| CDKN2B-AS1 | 0 14834098 | -0 06771123 | 5 29E-107 | 0 26428029 | 0 | 2 55E-106 |
| HSP90B1 | 0 566208524 | -0 06774392 | 4 19E-107 | 0 051815587 | 2 20E-63 | 2 02E-106 |
| ARHGEF28 | 0 358635853 | -0 06778771 | 3 06E-107 | -0 21109833 | 0 | 1 48E-106 |

| | | | | | | |
|---|---|---|---|---|---|---|
| PDLIM4 | 0 134615751 | -0 06786844 | 1 71E-107 | -0 35375154 | 0 | 8 32E-107 |
| COL28A1 | 0 142306448 | -0 06788922 | 1 48E-107 | -0 31722687 | 0 | 7 18E-107 |
| MT-ND5 | 0 374674002 | -0 06796518 | 8 56E-108 | -0 20308254 | 0 | 4 17E-107 |
| NGFR | 0 199006301 | -0 06798876 | 7 22E-108 | -0 29756011 | 0 | 3 52E-107 |
| SERF2 | 0 409653348 | -0 06802994 | 5 37E-108 | -0 19705589 | 0 | 2 62E-107 |
| LGALS3 | 0 301203099 | -0 06807198 | 3 97E-108 | -0 23253244 | 0 | 1 94E-107 |
| RPL38 | 0 417848508 | -0 0681272 | 2 67E-108 | 0 075263087 | 8 29E-132 | 1 31E-107 |
| PARP15 | 0 027383831 | -0 06817905 | 1 84E-108 | 0 413530198 | 0 | 9 00E-108 |
| DOCK2 | 0 102491862 | -0 06854881 | 1 27E-109 | 0 264910079 | 0 | 6 29E-109 |
| SDC3 | 0 133226095 | -0 06856242 | 1 15E-109 | -0 35934142 | 0 | 5 71E-109 |
| PCSK2 | 0 032352325 | -0 06865297 | 5 95E-110 | -0 45649632 | 0 | 2 96E-109 |
| RHOC | 0 181054996 | -0 06867562 | 5 04E-110 | -0 34052326 | 0 | 2 52E-109 |
| ANGPTL7 | 0 162979955 | -0 06892851 | 8 01E-111 | -0 31645614 | 0 | 4 02E-110 |
| ANXA1 | 0 440121071 | -0 06895824 | 6 45E-111 | -0 14641903 | 0 | 3 24E-110 |
| CADPS | 0 154251775 | -0 06921253 | 1 01E-111 | 0 253999391 | 0 | 5 07E-111 |
| SMS | 0 268955474 | -0 06935165 | 3 63E-112 | 0 156032549 | 0 | 1 84E-111 |
| MT-ND4L | 0 280424892 | -0 06981554 | 1 19E-113 | -0 25488177 | 0 | 6 11E-113 |
| ENO2 | 0 144743104 | -0 06989264 | 6 75E-114 | 0 269532728 | 0 | 3 47E-113 |
| COX6A1 | 0 212826712 | -0 06993134 | 5 07E-114 | 0 248996394 | 0 | 2 61E-113 |
| PRDX1 | 0 217957016 | -0 06995114 | 4 38E-114 | -0 31223613 | 0 | 2 26E-113 |
| SPCS2 | 0 313424454 | -0 07003837 | 2 30E-114 | 0 160247451 | 0 | 1 19E-113 |
| DNAJC6 | 0 090784489 | -0 07007637 | 1 73E-114 | 0 352079394 | 0 | 8 98E-114 |
| LGI4 | 0 06759818 | -0 07008261 | 1 65E-114 | -0 41422137 | 0 | 8 58E-114 |
| CAP1 | 0 261835868 | -0 07013928 | 1 09E-114 | -0 27390848 | 0 | 5 64E-114 |
| ALDH3B1 | 0 030838933 | -0 07028331 | 3 73E-115 | 0 437637531 | 0 | 1 94E-114 |
| MYOF | 0 445784394 | -0 07033362 | 2 56E-115 | -0 13616105 | 0 | 1 34E-114 |
| CD109 | 0 311121052 | -0 07044976 | 1 08E-115 | -0 25195678 | 0 | 5 65E-115 |
| NRG3 | 0 050037121 | -0 07046017 | 9 99E-116 | -0 44502705 | 0 | 5 23E-115 |
| PRNP | 0 242494908 | -0 07056199 | 4 68E-116 | -0 27424262 | 0 | 2 46E-115 |
| KLF6 | 0 438055624 | -0 07069949 | 1 67E-116 | -0 14736226 | 0 | 8 84E-116 |
| AHNAK | 0 525356456 | -0 07086949 | 4 69E-117 | -0 13240639 | 0 | 2 49E-116 |
| CIT | 0 182577906 | -0 07101308 | 1 60E-117 | 0 203912242 | 0 | 8 51E-117 |
| RPL19 | 0 419961547 | -0 07102521 | 1 46E-117 | -0 21178756 | 0 | 7 78E-117 |
| PDGFA | 0 084492966 | -0 07106624 | 1 07E-117 | -0 40512346 | 0 | 5 72E-117 |
| MT1X | 0 105670937 | -0 07112366 | 6 95E-118 | -0 38686006 | 0 | 3 72E-117 |
| LINC00511 | 0 397793684 | -0 07164375 | 1 37E-119 | -0 1966196 | 0 | 7 41E-119 |
| ITGA6 | 0 289800308 | -0 07213013 | 3 38E-121 | -0 25560784 | 0 | 1 86E-120 |
| PAG1 | 0 231111153 | -0 07214124 | 3 11E-121 | 0 20994395 | 0 | 1 71E-120 |
| RPL28 | 0 500276027 | -0 07226177 | 1 24E-121 | 0 038287014 | 2 18E-35 | 6 83E-121 |
| SNX22 | 0 038710476 | -0 07247656 | 2 39E-122 | -0 4438276 | 0 | 1 32E-121 |
| RPL41 | 0 69322876 | -0 07255559 | 1 30E-122 | -0 05206713 | 5 52E-64 | 7 25E-122 |
| HSP90AA1 | 0 556633226 | -0 0726571 | 5 98E-123 | -0 13133771 | 0 | 3 34E-122 |
| EDNRB | 0 061525575 | -0 07267647 | 5 15E-123 | 0 392587454 | 0 | 2 88E-122 |
| ENO1 | 0 331689859 | -0 07277859 | 2 35E-123 | 0 122418608 | 0 | 1 31E-122 |

| | | | | | | |
|---|---|---|---|---|---|---|
| ABCB5 | 0 06407645 | -0 07280013 | 1 99E-123 | -0 43226952 | 0 | 1 11E-122 |
| HINT1 | 0 256086882 | -0 07300388 | 4 13E-124 | 0 204279325 | 0 | 2 32E-123 |
| ART3 | 0 048637947 | -0 07317672 | 1 09E-124 | -0 43760468 | 0 | 6 12E-124 |
| ATP5F1E | 0 318259694 | -0 07323289 | 7 02E-125 | 0 155856699 | 0 | 3 97E-124 |
| WDR64 | 0 036673583 | -0 07366407 | 2 46E-126 | -0 44967512 | 0 | 1 41E-125 |
| CCND1 | 0 202356704 | -0 07376358 | 1 13E-126 | 0 181297043 | 0 | 6 50E-126 |
| NDUFC2 | 0 176695665 | -0 07395267 | 2 58E-127 | -0 35851985 | 0 | 1 49E-126 |
| MTAP | 0 340322857 | -0 07418605 | 4 15E-128 | -0 21558719 | 0 | 2 41E-127 |
| IRAK2 | 0 075983705 | -0 0742452 | 2 61E-128 | 0 360323289 | 0 | 1 52E-127 |
| CAST | 0 489273001 | -0 07447094 | 4 41E-129 | 0 061430696 | 2 22E-88 | 2 59E-128 |
| AKAP12 | 0 481943995 | -0 0747602 | 4 49E-130 | -0 13145903 | 0 | 2 66E-129 |
| NABP1 | 0 146599151 | -0 07481745 | 2 86E-130 | 0 282196609 | 0 | 1 69E-129 |
| MANCR | 0 032637871 | -0 07494672 | 1 03E-130 | 0 396870033 | 0 | 6 09E-130 |
| KCNK10 | 0 0345891 | -0 07494853 | 1 01E-130 | 0 433582171 | 0 | 6 01E-130 |
| STX3 | 0 119653157 | -0 07527174 | 7 74E-132 | 0 318528316 | 0 | 4 63E-131 |
| S100A16 | 0 109421104 | -0 07568676 | 2 81E-133 | 0 328364447 | 0 | 1 70E-132 |
| AJAP1 | 0 043688489 | -0 07574332 | 1 79E-133 | -0 44843471 | 0 | 1 09E-132 |
| RPGR | 0 214920713 | -0 07586799 | 6 57E-134 | 0 235768442 | 0 | 4 00E-133 |
| TMEM229B | 0 03248558 | -0 07595488 | 3 27E-134 | -0 46057345 | 0 | 1 99E-133 |
| PLAUR | 0 099883878 | -0 07610046 | 1 01E-134 | 0 313832214 | 0 | 6 20E-134 |
| TRIO | 0 831984923 | -0 07619041 | 4 91E-135 | -0 06477699 | 4 47E-98 | 3 02E-134 |
| RPL35 | 0 429013345 | -0 07640682 | 8 55E-136 | 0 080811653 | 1 02E-151 | 5 27E-135 |
| GALNT13 | 0 152233919 | -0 07644998 | 6 03E-136 | 0 261456273 | 0 | 3 73E-135 |
| LINC02006 | 0 058165655 | -0 0766186 | 1 54E-136 | 0 348382523 | 0 | 9 53E-136 |
| ARL6IP1 | 0 179674859 | -0 0767727 | 4 40E-137 | 0 223837574 | 0 | 2 74E-136 |
| COL7A1 | 0 123241515 | -0 07693347 | 1 19E-137 | 0 30311406 | 0 | 7 43E-137 |
| TMEM178B | 0 146494451 | -0 07705837 | 4 30E-138 | -0 33662632 | 0 | 2 70E-137 |
| SHC4 | 0 115036835 | -0 0771238 | 2 52E-138 | -0 37752086 | 0 | 1 58E-137 |
| KIF13B | 0 310521406 | -0 07729812 | 6 07E-139 | -0 2280946 | 0 | 3 82E-138 |
| LINC-PINT | 0 558565419 | -0 07742254 | 2 19E-139 | -0 06324425 | 1 43E-93 | 1 39E-138 |
| TMEM71 | 0 026393939 | -0 07754168 | 8 24E-140 | 0 407730109 | 0 | 5 24E-139 |
| GPR155 | 0 217214597 | -0 07791117 | 3 94E-141 | -0 29249373 | 0 | 2 54E-140 |
| ZBTB38 | 0 326635701 | -0 07798032 | 2 23E-141 | 0 165008001 | 0 | 1 44E-140 |
| SH3TC2-DT | 0 045772972 | -0 07803042 | 1 47E-141 | -0 44368012 | 0 | 9 53E-141 |
| CDH13 | 0 226590014 | -0 07841601 | 6 04E-143 | 0 162171539 | 0 | 3 94E-142 |
| ST3GAL6 | 0 253231425 | -0 07854728 | 2 03E-143 | -0 2858423 | 0 | 1 33E-142 |
| RPS3 | 0 422931222 | -0 07857769 | 1 58E-143 | -0 23029304 | 0 | 1 04E-142 |
| TMEM258 | 0 193342978 | -0 07878546 | 2 79E-144 | -0 34605961 | 0 | 1 85E-143 |
| MFSD2A | 0 034665245 | -0 07890113 | 1 06E-144 | -0 45771789 | 0 | 7 07E-144 |
| CD274 | 0 035274409 | -0 07890232 | 1 05E-144 | 0 385027292 | 0 | 7 01E-144 |
| RENBP | 0 032742571 | -0 07928216 | 4 37E-146 | -0 46338721 | 0 | 2 94E-145 |
| MT-ND2 | 0 509975062 | -0 0795236 | 5 75E-147 | -0 15896348 | 0 | 3 88E-146 |
| NT5DC3 | 0 214539986 | -0 07957181 | 3 83E-147 | 0 211263382 | 0 | 2 59E-146 |
| UPP1 | 0 092440654 | -0 079817 | 4 84E-148 | -0 41450774 | 0 | 3 29E-147 |

| | | | | | | |
|---|---|---|---|---|---|---|
| SOD3 | 0 121575831 | -0 07986859 | 3 13E-148 | -0 37562296 | 0 | 2 13E-147 |
| GSTP1 | 0 233490701 | -0 08009901 | 4 44E-149 | -0 31882787 | 0 | 3 05E-148 |
| PRELP | 0 143220194 | -0 08031391 | 7 16E-150 | -0 33641971 | 0 | 4 96E-149 |
| ATP5MG | 0 247834612 | -0 08032698 | 6 40E-150 | 0 184043924 | 0 | 4 44E-149 |
| MT-ND3 | 0 659505816 | -0 08042989 | 2 67E-150 | -0 12030054 | 0 | 1 85E-149 |
| MT-ATP6 | 0 631284384 | -0 08067236 | 3 37E-151 | -0 11702094 | 0 | 2 37E-150 |
| SRGN | 0 111772096 | -0 08103557 | 1 50E-152 | -0 41610873 | 0 | 1 07E-151 |
| PKM | 0 332536978 | -0 08110674 | 8 16E-153 | 0 109836051 | 2 85E-279 | 5 81E-152 |
| DYNLT3 | 0 107565057 | -0 0811124 | 7 77E-153 | 0 322202138 | 0 | 5 54E-152 |
| FHOD3 | 0 263254079 | -0 0813691 | 8 54E-154 | 0 178000073 | 0 | 6 11E-153 |
| DCBLD2 | 0 482239059 | -0 08184721 | 1 37E-155 | 0 050145918 | 1 78E-59 | 9 91E-155 |
| ABCB4 | 0 08058099 | -0 08192563 | 6 94E-156 | -0 43574815 | 0 | 5 03E-155 |
| NTNG1 | 0 218794617 | -0 0819755 | 4 50E-156 | 0 1472396 | 0 | 3 27E-155 |
| S100A11 | 0 286516533 | -0 08207092 | 1 96E-156 | -0 28505164 | 0 | 1 43E-155 |
| NME2 | 0 270307057 | -0 08222549 | 5 12E-157 | 0 179879993 | 0 | 3 74E-156 |
| NCEH1 | 0 116759628 | -0 08256714 | 2 59E-158 | 0 280642166 | 0 | 1 91E-157 |
| NIM1K | 0 079105671 | -0 08283723 | 2 44E-159 | 0 377551384 | 0 | 1 80E-158 |
| KHDRBS3 | 0 327787402 | -0 08338473 | 1 96E-161 | -0 21666558 | 0 | 1 48E-160 |
| AQP3 | 0 03327559 | -0 08342157 | 1 42E-161 | -0 45124065 | 0 | 1 07E-160 |
| CRYM | 0 090489425 | -0 08385772 | 2 97E-163 | -0 39046453 | 0 | 2 26E-162 |
| SKAP1-AS1 | 0 027155394 | -0 08387316 | 2 59E-163 | 0 40016117 | 0 | 1 97E-162 |
| ITGB4 | 0 14917858 | -0 08397087 | 1 09E-163 | -0 31942144 | 0 | 8 32E-163 |
| FOXD3 | 0 037263711 | -0 08400029 | 8 37E-164 | -0 45947356 | 0 | 6 41E-163 |
| MT-CYB | 0 567655289 | -0 08414358 | 2 34E-164 | -0 1428145 | 0 | 1 80E-163 |
| SRPX | 0 284384459 | -0 08420615 | 1 34E-164 | 0 156920238 | 0 | 1 03E-163 |
| GBP2 | 0 105994556 | -0 08452 | 8 08E-166 | -0 39460014 | 0 | 6 29E-165 |
| ST6GALNAC2 | 0 065856351 | -0 08472965 | 1 23E-166 | -0 43055563 | 0 | 9 66E-166 |
| PRIMA1 | 0 045449354 | -0 08478115 | 7 77E-167 | -0 44058968 | 0 | 6 09E-166 |
| CACNA2D4 | 0 040985323 | -0 08479921 | 6 61E-167 | 0 387065967 | 0 | 5 19E-166 |
| PRKCA | 0 726494832 | -0 08501243 | 9 70E-168 | -0 07441085 | 7 08E-129 | 7 67E-167 |
| CTNNA2 | 0 095905275 | -0 08555748 | 7 04E-170 | -0 42459829 | 0 | 5 61E-169 |
| HMGA1 | 0 050636767 | -0 0860207 | 1 04E-171 | 0 36256856 | 0 | 8 37E-171 |
| FOSL1 | 0 068293008 | -0 08610749 | 4 73E-172 | 0 31077733 | 0 | 3 81E-171 |
| MT-ND1 | 0 527688413 | -0 086183 | 2 37E-172 | -0 16432147 | 0 | 1 92E-171 |
| ADGRE5 | 0 145514077 | -0 08618345 | 2 36E-172 | 0 224458879 | 0 | 1 91E-171 |
| MYL6 | 0 442177 | -0 08632134 | 6 70E-173 | -0 1992066 | 0 | 5 44E-172 |
| GLIS3 | 0 355713769 | -0 08632763 | 6 33E-173 | 0 131882039 | 0 | 5 14E-172 |
| NRCAM | 0 259732349 | -0 0865861 | 5 92E-174 | 0 159777618 | 0 | 4 87E-173 |
| LGALS3BP | 0 175439264 | -0 08677567 | 1 04E-174 | -0 33944752 | 0 | 8 61E-174 |
| TNFRSF12A | 0 139355809 | -0 0868685 | 4 42E-175 | 0 251537366 | 0 | 3 68E-174 |
| LINC00466 | 0 048790238 | -0 08686969 | 4 37E-175 | 0 381881785 | 0 | 3 64E-174 |
| CST3 | 0 387190421 | -0 08688969 | 3 64E-175 | -0 2076178 | 0 | 3 03E-174 |
| CORO1C | 0 37636824 | -0 08696713 | 1 78E-175 | 0 119376437 | 0 | 1 49E-174 |
| RASSF4 | 0 114161162 | -0 08736046 | 4 70E-177 | 0 308565083 | 0 | 3 97E-176 |

| | | | | | | |
|---|---|---|---|---|---|---|
| FRMPD4 | 0 10702252 | -0 0874023 | 3 19E-177 | 0 306872721 | 0 | 2 70E-176 |
| FTL | 0 545297063 | -0 08771209 | 1 80E-178 | -0 1746197 | 0 | 1 53E-177 |
| CTSD | 0 213083703 | -0 08778695 | 8 98E-179 | -0 3114858 | 0 | 7 66E-178 |
| HSPB1 | 0 313995545 | -0 08830932 | 6 85E-181 | -0 25292997 | 0 | 5 91E-180 |
| MYH9 | 0 308122823 | -0 08837682 | 3 64E-181 | 0 116238158 | 0 | 3 15E-180 |
| CCBE1 | 0 109268813 | -0 08846091 | 1 65E-181 | 0 235976537 | 0 | 1 44E-180 |
| VMP1 | 0 741543089 | -0 088513 | 1 01E-181 | -0 05780251 | 1 89E-78 | 8 85E-181 |
| LINC02391 | 0 073299575 | -0 08853496 | 8 26E-182 | 0 327420113 | 0 | 7 21E-181 |
| AFAP1L2 | 0 206040243 | -0 08857126 | 5 87E-182 | -0 3153397 | 0 | 5 13E-181 |
| LAMC1 | 0 653871048 | -0 0887069 | 1 64E-182 | -0 09341231 | 3 02E-202 | 1 44E-181 |
| BAALC | 0 148217243 | -0 08887479 | 3 38E-183 | 0 272974391 | 0 | 2 97E-182 |
| SPHK1 | 0 104452609 | -0 08893908 | 1 84E-183 | 0 317657153 | 0 | 1 63E-182 |
| CD82 | 0 087538787 | -0 0889999 | 1 04E-183 | 0 319108833 | 0 | 9 17E-183 |
| MT-CO3 | 0 685309627 | -0 08904862 | 6 56E-184 | -0 11631869 | 0 | 5 79E-183 |
| MIF | 0 272305877 | -0 08916416 | 2 20E-184 | 0 167025317 | 0 | 1 95E-183 |
| CLIC4 | 0 513030401 | -0 08919948 | 1 58E-184 | 0 065581951 | 1 74E-100 | 1 40E-183 |
| FXYD3 | 0 030762788 | -0 08941992 | 1 96E-185 | -0 45672942 | 0 | 1 75E-184 |
| TFPI2 | 0 061439912 | -0 08947324 | 1 18E-185 | 0 314143035 | 0 | 1 06E-184 |
| GPR37L1 | 0 028973368 | -0 08963858 | 2 45E-186 | -0 45563168 | 0 | 2 21E-185 |
| PTPRZ1 | 0 161247644 | -0 08969936 | 1 38E-186 | -0 33744236 | 0 | 1 25E-185 |
| ITIH6 | 0 036483219 | -0 09089283 | 1 50E-191 | -0 4513839 | 0 | 1 40E-190 |
| RND3 | 0 314157355 | -0 09093489 | 9 99E-192 | -0 24630455 | 0 | 9 34E-191 |
| RPLP1 | 0 597380594 | -0 09120412 | 7 41E-193 | -0 04149195 | 2 90E-41 | 7 01E-192 |
| ENDOD1 | 0 108573985 | -0 09141575 | 9 55E-194 | -0 37684184 | 0 | 9 10E-193 |
| HLA-C | 0 229835716 | -0 09163497 | 1 14E-194 | 0 182978847 | 0 | 1 09E-193 |
| TUBA1B | 0 340475148 | -0 09168832 | 6 77E-195 | 0 104276793 | 8 88E-252 | 6 52E-194 |
| CD59 | 0 324084826 | -0 09207465 | 1 57E-196 | -0 23801166 | 0 | 1 52E-195 |
| TPI1 | 0 229645352 | -0 09211825 | 1 02E-196 | 0 168984036 | 0 | 9 92E-196 |
| CHL1 | 0 11064895 | -0 09273584 | 2 40E-199 | -0 36668731 | 0 | 2 36E-198 |
| SH3RF2 | 0 063353068 | -0 09279608 | 1 33E-199 | -0 44183406 | 0 | 1 30E-198 |
| FAU | 0 345081952 | -0 09340705 | 3 18E-202 | -0 27449493 | 0 | 3 16E-201 |
| BTC | 0 044088253 | -0 09349161 | 1 37E-202 | -0 46361635 | 0 | 1 37E-201 |
| CFL1 | 0 267851364 | -0 09353337 | 9 08E-203 | -0 29932473 | 0 | 9 08E-202 |
| EIF1 | 0 452685081 | -0 09365527 | 2 71E-203 | -0 17537637 | 0 | 2 72E-202 |
| LINC02240 | 0 057889627 | -0 09391551 | 2 03E-204 | -0 45674547 | 0 | 2 05E-203 |
| PPP1R14B | 0 132340903 | -0 09421563 | 1 02E-205 | 0 264119668 | 0 | 1 04E-204 |
| CD9 | 0 167777122 | -0 09440581 | 1 51E-206 | -0 35435692 | 0 | 1 56E-205 |
| ADIRF | 0 105908892 | -0 09544573 | 4 27E-211 | -0 39243043 | 0 | 4 53E-210 |
| PGM2L1 | 0 156098304 | -0 09589273 | 4 56E-213 | 0 257822021 | 0 | 4 92E-212 |
| ABCA13 | 0 073566085 | -0 09713198 | 1 40E-218 | 0 361175492 | 0 | 1 56E-217 |
| ESYT2 | 0 645904323 | -0 09730437 | 2 36E-219 | -0 08822271 | 1 54E-180 | 2 65E-218 |
| ANKRD30B | 0 04252727 | -0 09769658 | 4 06E-221 | -0 45551976 | 0 | 4 63E-220 |
| ADGRG1 | 0 046382136 | -0 09801491 | 1 49E-222 | -0 44703147 | 0 | 1 71E-221 |
| ENTHD1 | 0 033418362 | -0 09817847 | 2 71E-223 | -0 46265691 | 0 | 3 13E-222 |

| | | | | | | |
|---|---|---|---|---|---|---|
| ERBB3 | 0 06113533 | -0 09846782 | 1 32E-224 | -0 42457643 | 0 | 1 53E-223 |
| MT-CO2 | 0 654603948 | -0 09872215 | 9 21E-226 | -0 13121886 | 0 | 1 08E-224 |
| UBC | 0 452294835 | -0 09877069 | 5 53E-226 | -0 17589571 | 0 | 6 50E-225 |
| SLC17A9 | 0 0461537 | -0 09927905 | 2 64E-228 | 0 353417376 | 0 | 3 12E-227 |
| TXN | 0 260046449 | -0 09950062 | 2 54E-229 | -0 29391687 | 0 | 3 05E-228 |
| PLEKHB1 | 0 057527936 | -0 09957733 | 1 13E-229 | -0 43200952 | 0 | 1 36E-228 |
| PLP1 | 0 109183149 | -0 09974767 | 1 86E-230 | -0 37599916 | 0 | 2 26E-229 |
| GPM6B | 0 291932383 | -0 1015573 | 7 36E-239 | -0 28359523 | 0 | 9 37E-238 |
| LAPTM5 | 0 077811197 | -0 10225432 | 3 88E-242 | 0 342781632 | 0 | 5 06E-241 |
| MAP1B | 0 498201062 | -0 10233795 | 1 56E-242 | -0 15781139 | 0 | 2 05E-241 |
| PPP2R2B | 0 140574137 | -0 10236181 | 1 21E-242 | -0 35427467 | 0 | 1 58E-241 |
| SPOCD1 | 0 035683692 | -0 10256298 | 1 35E-243 | 0 388746165 | 0 | 1 78E-242 |
| APOD | 0 495659706 | -0 10262317 | 7 00E-244 | -0 1802203 | 0 | 9 26E-243 |
| SCD | 0 148950144 | -0 10272627 | 2 27E-244 | -0 36413278 | 0 | 3 02E-243 |
| ARNTL2 | 0 075879005 | -0 10273279 | 2 12E-244 | 0 287084111 | 0 | 2 82E-243 |
| AP1S2 | 0 247196893 | -0 10284487 | 6 22E-245 | 0 206818972 | 0 | 8 31E-244 |
| MT-ND4 | 0 587710114 | -0 10339784 | 1 45E-247 | -0 14996626 | 0 | 1 96E-246 |
| MT-CO1 | 0 662332718 | -0 10377737 | 2 22E-249 | -0 13580185 | 0 | 3 04E-248 |
| CAVIN3 | 0 130665702 | -0 10400383 | 1 82E-250 | -0 38290746 | 0 | 2 50E-249 |
| MIR222HG | 0 207353753 | -0 10474648 | 4 81E-254 | 0 173332516 | 0 | 6 72E-253 |
| MIA | 0 109097485 | -0 10483434 | 1 81E-254 | -0 37966623 | 0 | 2 54E-253 |
| MCAM | 0 1040814 | -0 10486721 | 1 25E-254 | -0 39062063 | 0 | 1 77E-253 |
| SOX10 | 0 062277512 | -0 10494573 | 5 21E-255 | -0 42350625 | 0 | 7 38E-254 |
| ARHGAP22 | 0 152538501 | -0 10523872 | 1 97E-256 | 0 215638766 | 0 | 2 81E-255 |
| ACTN4 | 0 468114066 | -0 10526407 | 1 49E-256 | 0 045838928 | 5 52E-50 | 2 12E-255 |
| FGL2 | 0 102720298 | -0 10570272 | 1 09E-258 | -0 38473479 | 0 | 1 57E-257 |
| TMC7 | 0 070977137 | -0 10688737 | 1 66E-264 | 0 311263659 | 0 | 2 48E-263 |
| GPR39 | 0 086910586 | -0 10694188 | 8 92E-265 | 0 264824579 | 0 | 1 34E-263 |
| PEX5L | 0 165131065 | -0 10716244 | 7 24E-266 | 0 236083744 | 0 | 1 10E-264 |
| PTPRE | 0 20248044 | -0 10747913 | 1 95E-267 | 0 225733485 | 0 | 2 99E-266 |
| CRYAB | 0 124079115 | -0 10799491 | 5 28E-270 | -0 36580375 | 0 | 8 24E-269 |
| KCNN4 | 0 04888542 | -0 1089797 | 6 10E-275 | -0 45135395 | 0 | 9 81E-274 |
| FTH1 | 0 537768175 | -0 10947675 | 1 89E-277 | -0 17539015 | 0 | 3 07E-276 |
| S100B | 0 096838058 | -0 10948993 | 1 62E-277 | -0 38993259 | 0 | 2 64E-276 |
| GLDC | 0 05551008 | -0 10954896 | 8 15E-278 | 0 35613923 | 0 | 1 33E-276 |
| DKK1 | 0 045991891 | -0 11036278 | 5 96E-282 | 0 344999286 | 0 | 9 89E-281 |
| L1CAM | 0 044088253 | -0 11073787 | 7 21E-284 | -0 47110957 | 0 | 1 21E-282 |
| ACTG1 | 0 551141231 | -0 11195177 | 4 06E-290 | -0 17235724 | 0 | 7 14E-289 |
| ABLIM3 | 0 194171061 | -0 11274038 | 3 25E-294 | -0 32735202 | 0 | 5 82E-293 |
| S100A10 | 0 262873351 | -0 11283252 | 1 07E-294 | -0 32053735 | 0 | 1 94E-293 |
| MALAT1 | 0 997601416 | -0 1138534 | 4 78E-300 | -0 09115236 | 1 22E-192 | 9 01E-299 |
| ACTB | 0 585777922 | -0 11499495 | 4 32E-306 | -0 14580363 | 0 | 8 37E-305 |
| RFX8 | 0 152329101 | -0 11543332 | 0 | 0 174201976 | 0 | 3 92E-307 |
| GSAP | 0 187413147 | -0 11613243 | 0 | 0 210495544 | 0 | 0 |

| | | | | | | |
|---|---|---|---|---|---|---|
| VEGFC | 0 112923797 | -0 11672621 | 0 | 0 268823104 | 0 | 0 |
| MT1E | 0 118349165 | -0 11830399 | 0 | -0 4060486 | 0 | 0 |
| HLA-B | 0 348698864 | -0 11847363 | 0 | 0 065354488 | 8 42E-100 | 0 |
| NEAT1 | 0 91331785 | -0 11873207 | 0 | -0 10348911 | 5 32E-248 | 0 |
| MTCL1 | 0 142106566 | -0 1196814 | 0 | 0 239694198 | 0 | 0 |
| PHLDA1 | 0 296577259 | -0 12006452 | 0 | -0 26630973 | 0 | 0 |
| ATP10B | 0 086272867 | -0 12080569 | 0 | -0 39957866 | 0 | 0 |
| ITGA2 | 0 138527726 | -0 12219294 | 0 | 0 281733122 | 0 | 0 |
| SOX2 | 0 063038967 | -0 12257343 | 0 | -0 447745 | 0 | 0 |
| COBL | 0 146846624 | -0 12275814 | 0 | 0 285564755 | 0 | 0 |
| CDKN2A | 0 147036988 | -0 12293106 | 0 | 0 21734725 | 0 | 0 |
| DST | 0 807408959 | -0 12300079 | 0 | -0 10324912 | 7 43E-247 | 0 |
| SKAP1 | 0 10549961 | -0 12388604 | 0 | 0 214518498 | 0 | 0 |
| INPP5F | 0 29420723 | -0 12506579 | 0 | -0 27561874 | 0 | 0 |
| CFAP299 | 0 126982163 | -0 12583112 | 0 | 0 235700989 | 0 | 0 |
| CAV2 | 0 211427538 | -0 12633361 | 0 | 0 185262548 | 0 | 0 |
| GSN | 0 68687061 | -0 12658278 | 0 | -0 07409321 | 8 59E-128 | 0 |
| PSAP | 0 478441301 | -0 12788871 | 0 | -0 17604504 | 0 | 0 |
| ANXA2 | 0 603529345 | -0 12918164 | 0 | -0 01232655 | 6 45E-05 | 0 |
| GAPDH | 0 370485999 | -0 13041086 | 0 | 0 044187376 | 1 43E-46 | 0 |
| LGALS1 | 0 413393996 | -0 13062909 | 0 | 0 015616913 | 4 14E-07 | 0 |
| SLC35F1 | 0 158810988 | -0 13104375 | 0 | -0 35904267 | 0 | 0 |
| HLA-A | 0 386210047 | -0 13154644 | 0 | 0 0662894 | 1 26E-102 | 0 |
| CD63 | 0 568197826 | -0 13219847 | 0 | -0 18881869 | 0 | 0 |
| TMSB4X | 0 594049228 | -0 14599241 | 0 | -0 0605514 | 6 45E-86 | 0 |
| CAV1 | 0 212683939 | -0 1465469 | 0 | 0 169894806 | 0 | 0 |
| FRMD5 | 0 230720908 | -0 15031433 | 0 | 0 145335206 | 0 | 0 |
| TMSB10 | 0 556071653 | -0 15340872 | 0 | -0 2105418 | 0 | 0 |
| B2M | 0 616436009 | -0 15360942 | 0 | -0 0776074 | 4 80E-140 | 0 |
| VIM | 0 705516742 | -0 15714965 | 0 | -0 149944 | 0 | 0 |
| ITGA3 | 0 153119111 | -0 15867581 | 0 | 0 201279321 | 0 | 0 |
| FGF5 | 0 095296111 | -0 16119024 | 0 | 0 233909908 | 0 | 0 |
| CDH19 | 0 419162019 | -0 16363143 | 0 | -0 17549827 | 0 | 0 |
| SPP1 | 0 186527955 | -0 16709448 | 0 | -0 3873135 | 0 | 0 |
| MT2A | 0 276760389 | -0 19420653 | 0 | 0 100789869 | 2 81E-235 | 0 |
| S100A6 | 0 489101673 | -0 19508655 | 0 | -0 24991819 | 0 | 0 |
| IFNL1 | 0 | | | | | |
